# Neural Inflammation in Thoracic Dorsal Root Ganglia Mediates Cardiopulmonary Spinal Afferent Sensitization in Chronic Heart Failure

**DOI:** 10.1101/2025.10.22.683960

**Authors:** Juan Hong, Samuel Gillman, Peter Pellegrino, Gang Zhao, Rongguo Ren, Steven J. Lisco, Irving H. Zucker, Dong Wang, Han-Jun Wang

## Abstract

The cardiac sympathetic afferent reflex (CSAR) and pulmonary spinal afferent reflex (PSAR) amplify sympathetic outflow, and their sensitization contributes to chronic heart failure (CHF). Using a myocardial infarction (MI) rat model, molecular profiling, imaging, and functional assays revealed that thoracic dorsal root ganglia (DRGs) undergo marked macrophage and glial activation and suppression of voltage-gated potassium (Kv) channels after MI. In vitro studies confirmed that pro-inflammatory cytokines and activated macrophages directly reduce Kv channel expression and activity in DRG neurons. Cardiac afferents mediated cytokine transport from the heart to DRGs, driving macrophage infiltration in a cytokine receptor-dependent manner. Anti-inflammatory strategies including systemic minocycline, liposomal clodronate–induced macrophage depletion, or local epidural dexamethasone prodrug delivery reduced neuroinflammation, restored Kv channel levels, attenuated the exaggerated CSAR and PSAR, and improved cardiac remodeling. These findings highlight a cytokine uptake–driven inflammatory pathway in cardiopulmonary spinal afferent sensitization and support targeted DRG anti-inflammatory therapy as a potential cardioprotective approach.

**Abstract:** The cardiac sympathetic afferent reflex (CSAR) and pulmonary spinal afferent reflex (PSAR) amplify sympathetic activity and may contribute to chronic heart failure (CHF). We hypothesized that neural inflammation in thoracic dorsal root ganglia (DRGs) drives cardiopulmonary afferent sensitization through suppression of voltage-gated potassium (Kv) channels after myocardial infarction (MI). MI was induced in rats by coronary ligation. Molecular profiling, immunofluorescence, tissue clearing, and functional assays were used to assess neuroinflammation and reflex responses. Post-MI, thoracic DRGs showed macrophage infiltration, glial activation, cytokine upregulation, and reduced Kv channel expression. Bulk RNA-seq identified enrichment of macrophage activation–related genes, and in vitro studies confirmed that pro-inflammatory cytokines and activated macrophages suppressed Kv channels and increased DRG neuron excitability. Epicardial injection of biotinylated TNF-α demonstrated cardiac afferent–mediated cytokine transport to DRGs, inducing macrophage infiltration via a cytokine receptor-dependent mechanism. Anti-inflammatory interventions including oral minocycline, systemic macrophage depletion, and local epidural delivery of thermo-responsive hydrogel-forming dexamethasone prodrug (ProGel-Dex) significantly reduced DRG neuroinflammation, restored Kv channel levels, and attenuated exaggerated CSAR and PSAR responses. ProGel-Dex also improved cardiac chamber dilation in the post-MI rats. These findings identify a cytokine uptake–glial activation– macrophage activation pathway as a driver of cardiopulmonary afferent sensitization after MI. Targeting DRG inflammation, particularly with sustained local dexamethasone delivery using ProGel-Dex, offers a precision medicine to dampen pathological sympathetic activation and improve cardiac outcomes in CHF.

**Highlights:**

1. Both cardiac (CSAR) and pulmonary (PSAR) spinal afferent reflexes are sensitized after myocardial infarction, contributing to sympathetic overactivation.
2. Thoracic dorsal root ganglia (T1–T4) exhibit macrophage activation, glial responses, pro-inflammatory cytokine upregulation, and suppression of Kv channels following MI.
3. Cardiac afferents mediate receptor-dependent uptake and transport of cytokines (e.g., TNF-α) from the heart to DRGs, driving macrophage infiltration and inflammation.
4. Activated macrophages and pro-inflammatory cytokines reduce Kv channel expression and Kv current density (Ito) in DRG neurons, enhancing excitability.
5. Anti-inflammatory strategies including minocycline, liposomal clodronate–induced macrophage depletion, and local epidural dexamethasone prodrug attenuate neuroinflammation, restore Kv channel expression, and suppress exaggerated CSAR/PSAR.
6. Targeting DRG inflammation, particularly via sustained epidural dexamethasone delivery, represents a promising cardioprotective precise medicine.

## Introduction

Chronic heart failure (CHF) is a progressive disease that affects approximately six million Americans with over 660,000 new diagnoses each year.^1^ Exaggerated sympatho-excitation, a hallmark of CHF is a critical factor in the development and progression of the CHF state.^2–6^ Sympathetic dysfunction takes the form of persistent and adverse activation of sympathetic nerve outflow to the heart, kidneys and other organs, causing an array of chronic effects that aggravate the disease progression. Previous studies from our laboratory have shown that the enhanced cardiac sympathetic afferent reflex (CSAR), a sympathoexcitatory reflex originating in the heart, is a critical contributor to the elevated sympathetic tone in CHF.^7–13^ More recently we published new evidence showing that chronic and selective ablation of cardiac spinal afferents with the potent neurotoxin resiniferatoxin (RTX) markedly attenuates the deleterious cardiac remodeling and cardiac dysfunction in CHF.^9^ However, the mechanisms underlying the enhanced CSAR in the CHF state are not fully understood.

Inflammation plays an important role in the development and progression of CHF, contributing to cardiac remodeling and peripheral vascular disturbances.^14,15^ Increased levels of inflammatory cytokines, especially that of TNFα, IL-6 and IL-1β have been found in CHF, with particularly high concentrations in patients with the most severe heart failure.^16,17^ It is also well recognized that inflammation, such as microglia activation in the central nervous system (CNS), is involved in the pathogenesis of sympatho-excitation in CHF ^18,19^. Microglia, resident macrophages in the brain, are thought to modulate the pathologic and regenerative states of the brain by producing a variety of molecules including neurotrophic and neurotoxic factors.^20–22^ Studies have demonstrated that microglial activation in the hypothalamic paraventricular nucleus (PVN) following MI mediated the increase of sympathetic nerve activity in animals with reduced left ventricular function.^18,19^ On the other hand, whether inflammation occurred in the peripheral nervous system, such as in sensory ganglia, in CHF remains unclear. It is well established that macrophages can infiltrate into the injured nerve tissue such as dorsal root ganglia (DRG) and be activated in the development of neuropathic pain following peripheral nerve injury.^23–26^ Whether similar macrophage activation occurs in thoracic T1-T4 DRGs (the most predominant sensory ganglia innervating both heart and lung) post-MI has not been examined. In this study, our first goal was to determine whether macrophages are activated in the thoracic T1-T4 DRGs in a time-dependent manner in the post-MI state. Our second goal was to define how activated macrophages influence thoracic visceral sensory afferents. We employed an *in vitro* co-culture system to probe macrophage-neuron interactions. We also tested a cardiac afferent–mediated cytokine uptake hypothesis by injecting biotinylated cytokines into the epicardium and tracking their transport to thoracic DRGs.

Because the lungs and heart anatomically share common sympathetic afferent (i.e. T1-T4 DRGs) input, macrophage infiltration and activation in these DRGs could also sensitize pulmonary afferents post-MI. Previous work from our laboratory has shown that activation of pulmonary spinal afferents by application of bradykinin and capsaicin to the surface of the lung resulted in a reflex increase in mean arterial pressure (MAP), heart rate, and renal sympathetic nerve activity (RSNA) in the anesthetized, vagotomized rat, demonstrating an excitatory pulmonary spinal afferent reflex (PSAR) in normal animals.^27,28^ We therefore hypothesized that the PSAR is sensitized in CHF and that this sensitization is induced by macrophage activation in T1-T4 DRGs common to both heart and lung.

## Results

### 1. Myocardial infarction causes time-dependent macrophage activation in thoracic T1- T4 DRGs

We examined whether macrophages infiltrate and activate within the DRGs at different time points after MI. T1–T4 DRG sections were prepared and stained with ionized calcium-binding adaptor molecule-1 (Iba-1), a macrophage-specific marker in the DRG.^29^ Compared with sham-operated rats, the number of Iba-1–immunoreactive cells was significantly increased at 1 week post-MI and remained elevated through 8 weeks, the latest time point examined, indicating sustained macrophage infiltration into thoracic DRGs after MI (**Fig. 1A and 1B**).

**Figure 1.**
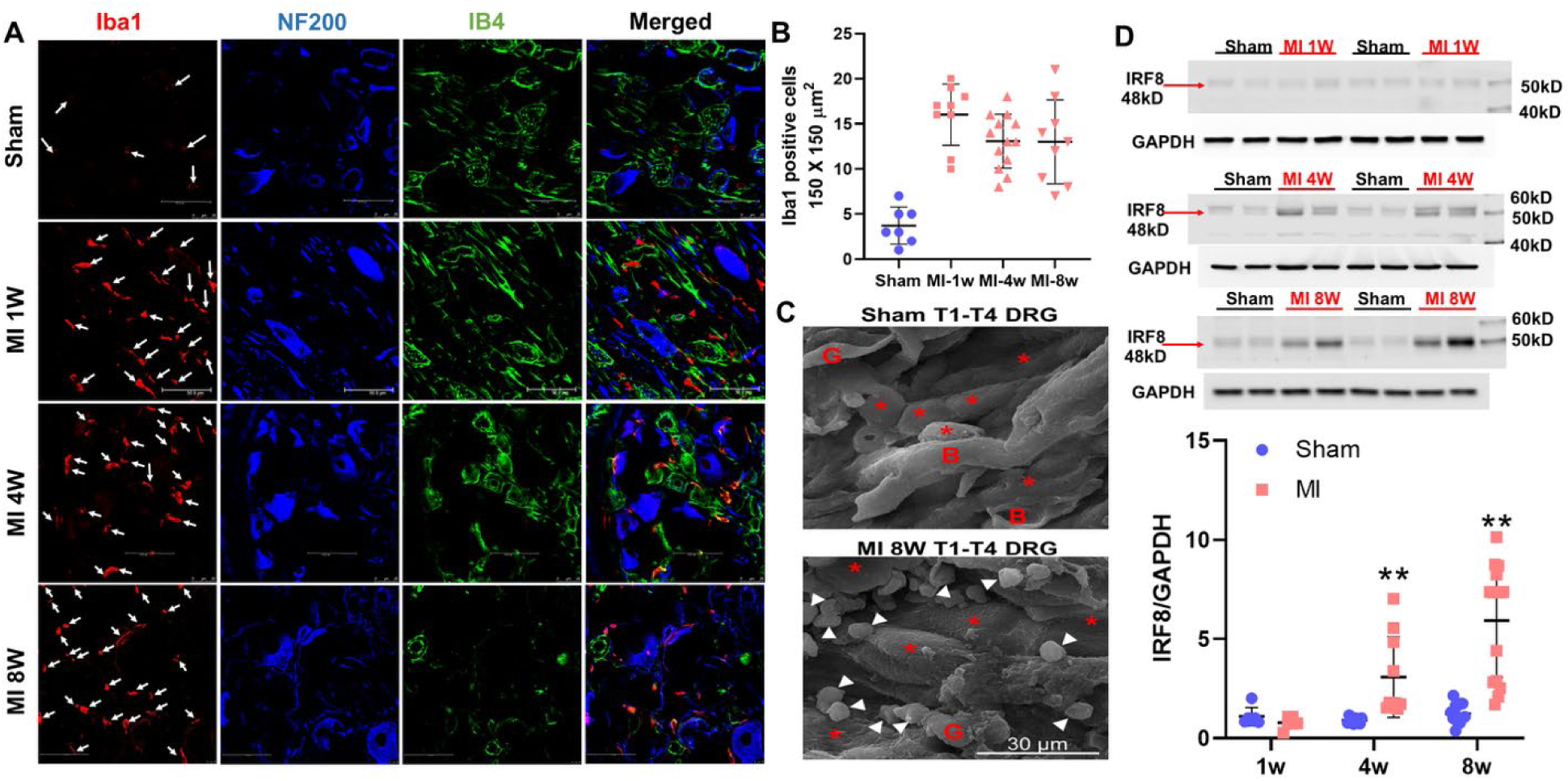
Macrophage activation in the post-MI rats. (**A**) Triple immunofluorescence of Iba1 (red) with NF200 (blue) and IB4 (green) in the T1-T4 DRGs of sham and myocardial infarction (MI) rats 1 week, 4 weeks and 8 weeks post MI. The white arrows point to the positive Iba1 cells. Scale bar=50 μm. (**B**) quantitative data comparing the Iba1-positive cells in in the T1-T4 DRGs of sham and MI rats 1 week, 4 weeks and 8 weeks post MI. **C**) Scanning electron microscope (SEM) results show that the number of macrophages is higher in 8w-MI rats than in the sham rats. Red asterisks and white arrows indicate DRG neurons and macrophages, respectively. “B” and “G” indicate blood vessels and glial cells, respectively. (**D**) Western blot analysis of IRF8 protein in the T1-T4 DRG of sham rats and MI rats at each time point. A histogram of the relative band density ratio of IRF8 (normalized to GAPDH) in MI rats to the sham rats at each time point (n=6).

Additional staining with NF200 (A-fiber neuronal marker) and IB4 (C-fiber neuronal marker) revealed that macrophages accumulated around both A- and C-fiber DRG neurons, suggesting potential influence on both subpopulations. To further confirm macrophage infiltration and activation, we employed a tissue clearing technique for high-resolution 3D imaging of thoracic DRGs in CX3CR1CreER-tdTomato reporter mice. Using this modified protocol, previously applied for imaging sensory afferents in muscle, DRG, nodose ganglia, stellate ganglia, and vagus/sciatic nerves,^30^ we observed abundant CX3CR1-positive macrophages in thoracic DRGs of male mice at 8 weeks post-MI. In contrast, sham-operated mice displayed only scattered CX3CR1-positive macrophages (**Supplemental Video**).

Scanning electron microscopy (SEM) provided higher-resolution confirmation of macrophages surrounding DRG neurons. Male rats at 8 weeks post-MI exhibited a marked increase in macrophages (white arrows) in T1–T4 DRGs compared to sham rats (**Figure 1C**).

To quantify macrophage activation, we assessed Interferon Regulatory Factor 8 (IRF8), a transcription factor critical for driving macrophages toward a reactive phenotype.^31^ IRF8 protein expression was significantly increased in male rats at 4 and 8 weeks post-MI (*P*<0.01, n=6; **Figure 1D**), and similar upregulation was observed in female rats at 4 weeks post-MI (**Supplemental Figure S1**).

### 2. Myocardial infarction upregulates pro-inflammatory gene expression in thoracic T1–T4 DRGs

To determine whether MI-induced macrophage activation is associated with increased inflammatory gene expression, we performed RNA-seq analysis on T1–T4 DRGs from 8-week post-MI rats (**Figure 2**). Differential Gene Expression (DEG) analysis identified 41 DEGs (29 upregulated and 12 downregulated; FDR≤0.15, log2FC≥0.5), and notably the expression of *Irf7*, which promotes conversion of anti-inflammatory M2 macrophages into pro-inflammatory M1 macrophages during chronic inflammation, was significantly elevated in post-MI rats (Log2FC=0.67; *P*<0.01, MI n=5; sham n = 4) (**Figure 2A**). Gene Ontology (GO) analysis highlighted gene groups associated with terms related to contractile proteins and immune responses (**Figure 2B**). KEGG pathway analysis again revealed enrichment in pathways centered on immune responses, contractile components, and phagosome (**Figure 2C**). At the gene level, there was increased expression of contractile genes (*Myl9, Actg2, Acta2, Myh11, Cnn1*) and immune/inflammatory response genes (*Cst3, Slfn4, Irf7, C4b, Pla2g2a, Ifit1*), and genes associated with negative regulation of neuronal growth and inhibitory neurotransmission (*Cbln4, Rtn4lp1*). In contrast, sham DRG exhibited relatively higher levels of ECM/adhesion markers (*Thbs4, Pcdhga3, Fndc7*) and positive regulations of neuronal growth (*Vgf, Elk1*). (**Figure 2D**).

**Figure 2.**
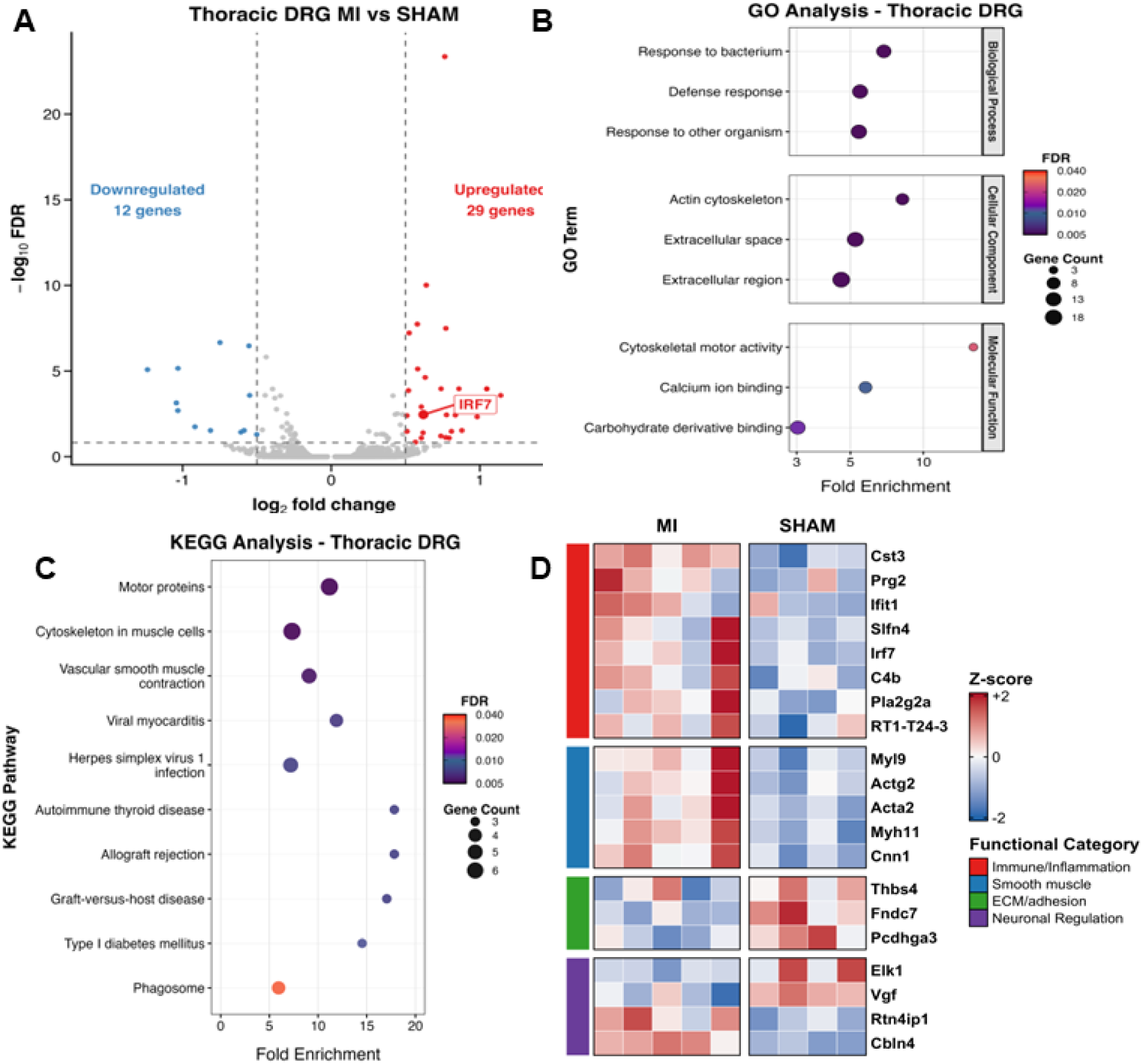
T1-T4 DRG transcriptional changes 8 weeks post-MI. (A) Volcano plot of differentially expressed genes (DEGs) in T1-T4 DRGs from rats 8 weeks post-MI versus sham (MI n =5; sham n =4). Thresholds: FDR≤0.15 and |log2FC|≥0.5. Upregulated (red) and downregulated (blue) genes are shown with *Irf7* highlighted. (B) Gene Ontology enrichment for DEGs. Enrichment includes clusters for immune/defense responses, extracellular/ECM, and cytoskeleton/motor activity. (C) KEGG pathway enrichment. Immune/inflammatory pathways (phagosome, viral myocarditis) and cytoskeleton/vascular-smooth muscle contraction are prominent. (D) Heatmap of representative DEGs groups by functional category. Values are gene- wise Z-scores across samples; columns are individual DRGs (MI, left block; sham, right block). Immune/Inflammation genes (*Cst3, Ifit1, Slfn4, Irf7, C4b, Pla2g2a, RT1-T24-3, Prg2*) and smooth-muscle/contractile genes (*Myl9, Actg2, Acta2, Myh11, Cnn1*) are increased in MI. Adhesion genes (*Thbs4, Fndc7, Pcdhga3*) are upregulated in the sham group.

### 3. Macrophage activation decreases Kv channel protein levels in 50B11 DRG neurons in vitro

Because voltage-gated K^+^ (Kv) channel isoforms (Kv1.4, Kv4.2, Kv4.3, Kv3.4) are known regulators of DRG neuronal excitability and nociceptive signaling,^32^ we asked whether macrophage activation could directly suppress these channels. We used an indirect co-culture design in which 50B11 DRG neurons were incubated for 72h with BV2 microglia or RAW264.7 macrophages, briefly primed with lipopolysaccharide (LPS, 25 ng/ml, 4 hours; LPS washed before co-culture; **Figure 3A**).

**Figure 3.**
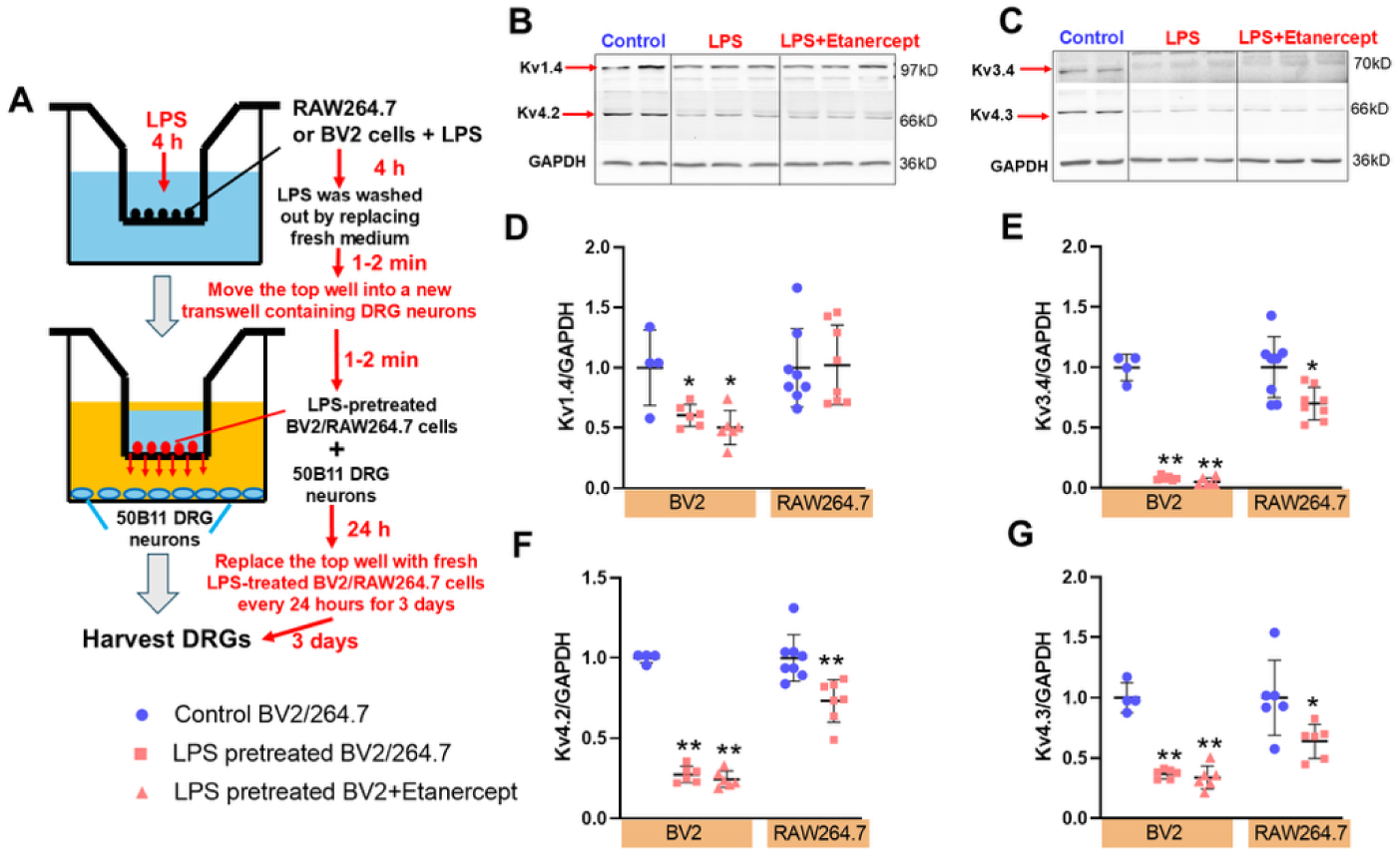
The protein expression of Kv channels in 50B11 DRG neurons after co-culture with BV2 or RAW264.7 cells. **(A)** A schematic diagram describing how 50B11 DRG neurons and BV2/ RAW264.7 cells were co-cultured. Original tracing (**B&C**) and mean data (**D-G**) showing protein expression of Kv channel isoforms in 50B11 DRG neurons after co-culture with BV2 or RAW264.7 cells with or without the treatment of Etanercept (10 ng/ml). \*\**P<*0.01 and \**P<*0.05 *vs.* control BV2 or RAW264.7 treated 50B11 neurons.

LPS-pretreated BV2 and RAW264.7 cells significantly reduced Kv channel protein expression in 50B11 neurons (BV2: Kv1.4 *P*<0.05, Kv4.2/4.3/3.4 *P*<0.01; RAW264.7: Kv1.4 *P*>0.05, Kv4.2/4.3/3.4 *P*<0.05, n=6; **Figure 3B–G**). Notably, the TNFα inhibitor Etanercept (10 ng/ml) failed to block these effects (*P*>0.05, n=6). These findings indicate that activated macrophages sensitize DRG neurons by downregulating Kv channel expression.

### 4. Pro-inflammatory cytokines reduce Kv channel expression in 50B11 cells *in vitro*

Considering that TNFα inhibitor Etanercept did not block any effects of activated microglia on protein expression, and that activated microglia/macrophages can also release many other pro-inflammatory cytokines, such as IL-1β, IL-6, and IFN-γ, and the anti-inflammatory cytokines IL- 10 and IL-4, we used multiple cytokines (including pro-inflammatory cytokines: TNFα, IL-6, IL- 1β, and IFNγ; anti-inflammatory cytokines: IL-4 and IL-10) to create an inflammatory environment and investigated their effects on protein levels of multiple Kv channels (Kv1.4, Kv3.4, Kv4.2, and Kv4.3). Compared to the control group, the protein expression of Kv1.4 significantly increased in 50B11 cells after three days of exposure to TNFα (10 ng/ml) (*P*<0.05, n=8, **Figure 4A and B**) while it decreased after treatment of IFNγ (50 ng/ml) (*P*<0.05, n=8). The protein expression of Kv3.4 significantly decreased in 50B11 with treatment of TNFα, IL-1β, IL-6, IFNγ and IL-4 (TNFα and IL-6: *P*<0.05; IL-1β, IFNγ and IL-4: *P*<0.01; n=8) Protein expression of Kv4.2 decreased following treatment with IL-6 and IFNγ, while it increased post-treatment with IL-4 (IL-6, IFNγ, and IL-4: *P*<0.05, n=8; **Figure 4 A, D, E**). Protein expression of Kv4.3 was significantly reduced by three-day exposure to IL-6 (10 ng/ml) (*P*<0.01; n=8; **Figure 4 E**). These data suggest that, in general, chronic exposure to pro-inflammatory cytokines causes a reduction in Kv channels; anti-inflammatory cytokines appear to have no clear effect on Kv channel isoforms.

**Figure 4.**
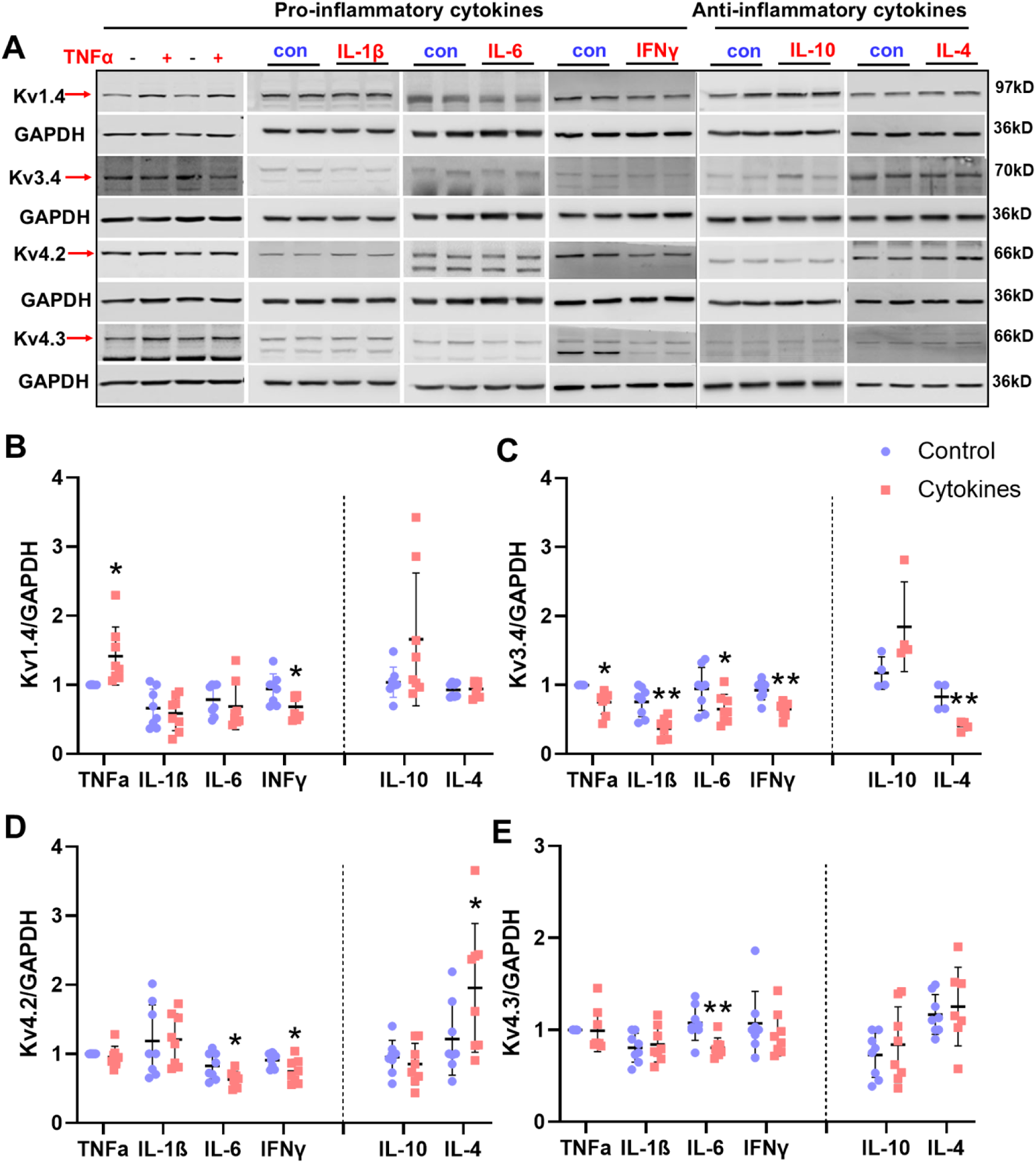
Original tracing (**A**) and mean data (**B-E**) showing protein expression of Kv channel isoforms in 50B11 DRG neurons after multiple pro-inflammatory (red) and anti-inflammatory (blue) cytokines treatment for 3 days. \*\**P<*0.01 and \**P<*0.05 *vs.* vehicle treated 50B11 neurons.

### 5. Protein expression of Kv channel isoforms is decreased in T1-T4 DRGs of post-MI rats

We next assessed Kv channel expression in T1–T4 DRGs in vivo at 1-, 4-, and 8-weeks post-MI (**Figure 5A**). Compared to sham rats, Kv1.4, Kv3.4, Kv4.2, and Kv4.3 protein levels were significantly reduced at 4 weeks (Kv1.4/Kv3.4 P<0.01; Kv4.2/4.3 *P*<0.05, n=6) and 8 weeks post-MI (Kv3.4/Kv4.2 *P*<0.05; Kv1.4/Kv4.3 *P*<0.01, n=6), but not at 1 week. Immunofluorescence confirmed a significant reduction in Kv-positive DRG neurons at 4 weeks post-MI (*P*<0.01, **Table S1**; **Supplemental Figure S2A–D**). These findings demonstrate progressive Kv channel downregulation in thoracic DRGs post-MI.

**Figure 5.**
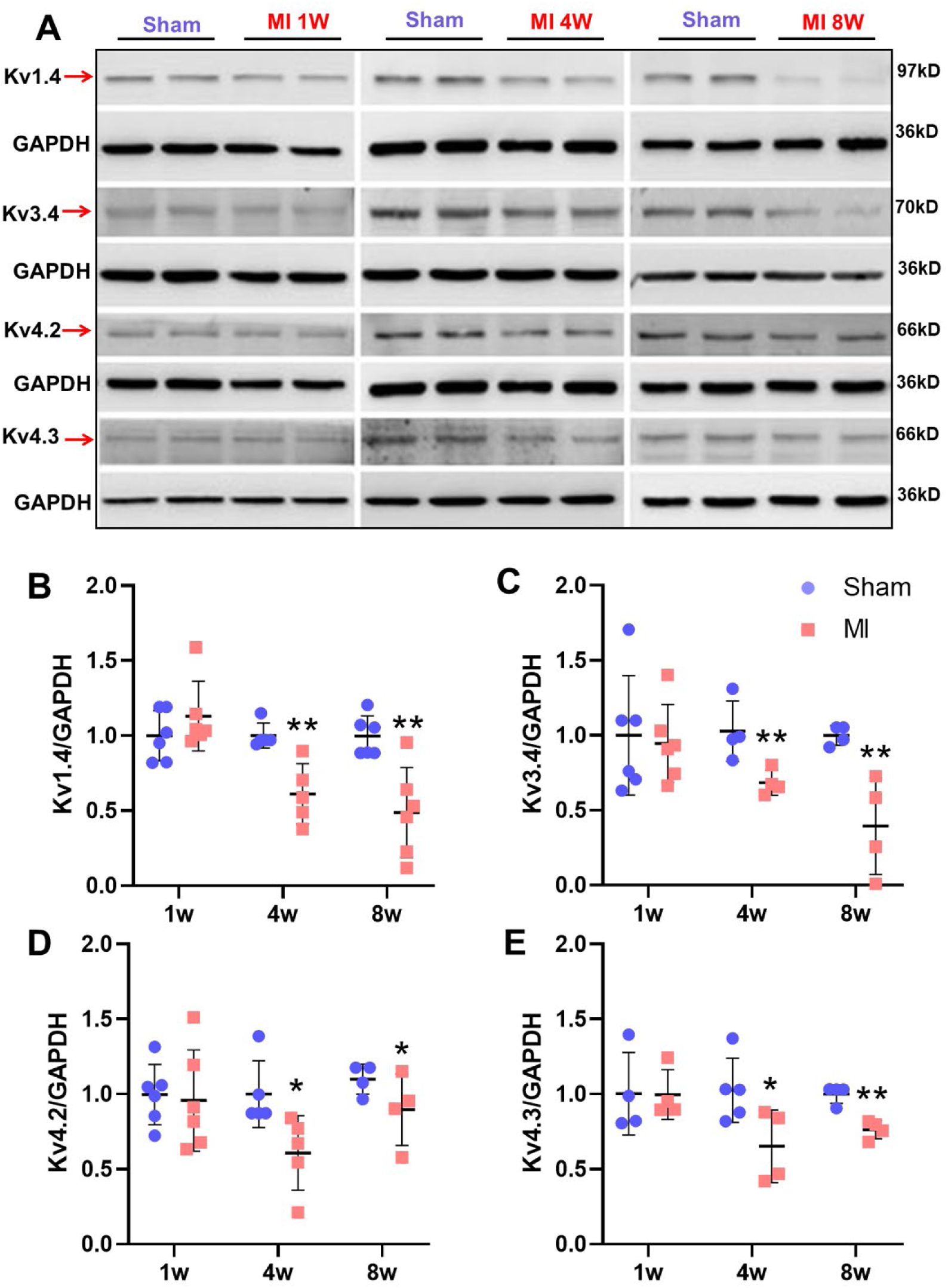
Original tracing (**A**) and mean data (**B-E**) showing protein expression of Kv channel isoforms in sham and 1w, 4w and 8w post-MI rats. \*\**P*<0.01 and \**P*<0.05 *vs.* sham rats.

### 6. Chronic oral minocycline attenuates macrophage activation, restores Kv channel function, and normalizes exaggerated CSAR and PSAR in post-MI rats

To determine the functional contribution of macrophage activation to DRG neuronal dysfunction, we co-cultured acutely dissociated T1–T4 DRG neurons with LPS-pretreated BV2 or RAW264.7 cells for 24 hours (non-LPS-treated cells as control). Two weeks prior to dissociation, DiI was injected into the subepicardium of the left ventricle to retrogradely label cardiac-specific afferent DRG neurons.^33^ Whole-cell patch-clamp recordings were performed in IB4-positive cardiac C-fiber neurons.^34^ Kv current density (Ito) was significantly decreased after co-culture with LPS-pretreated BV2 or RAW264.7 cells, but not controls (**Figure 6E, F**), confirming that activated macrophages suppress Kv currents in cardiac DRG neurons.

**Figure 6.**
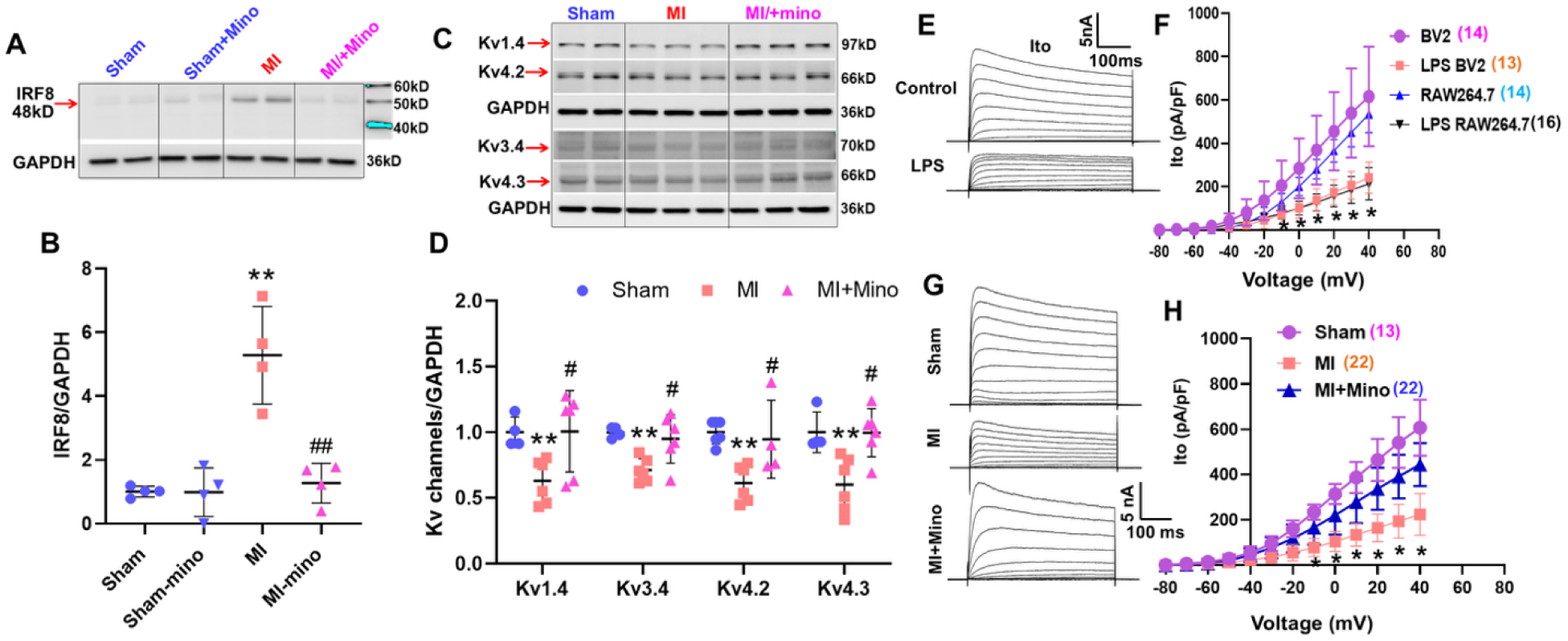
The effects of minocycline on protein expressions of IRF8 (**A, B**) and Kv channels (**C, D**) as well as Kv currents (**G, H**) in cardiac DRG neurons of sham and 4w-MI rats with or without the treatment of minocycline (20 mg/kg). **E and F,** the effects of LPS-pretreated microglia (BV2) or macrophages (RAW264.7) on Kv currents in cardiac DRG neurons of normal rats. \*\**P<*0.05 *vs.* sham rats, ^#^*P<*0.05 *vs.* MI rats.

To test whether macrophage inhibition could rescue these effects *in vivo*, we administered minocycline (20 mg/kg/day in drinking water), a known macrophage deactivator.^35–38^ Minocycline reduced IRF8 expression in T1–T4 DRGs by ∼76% in 4-week post-MI rats, restoring levels to those of sham controls (**Figure 6A, B**). IMinocycline had no effect in sham rats.

Minocycline also restored Kv channel protein expression (**Figure 6C, D**) and Kv current density (**Figure 6G, H**) in DRGs in 4-week post-MI rats. Functionally, minocycline prevented the exaggerated CSAR and PSAR responses induced by bradykinin (BK, 10 μg/ml) in vagotomized 4-week post-MI rats, bringing them to sham levels (**Figure 7A-D**). There was no significant difference in infarct size (34.50±2.67% vs. 36.13±2.82%, Veh vs. Minocycline, n=6-8/each group, *P*>0.05) between MI rats treated with vehicle or minocycline.

**Figure 7.**
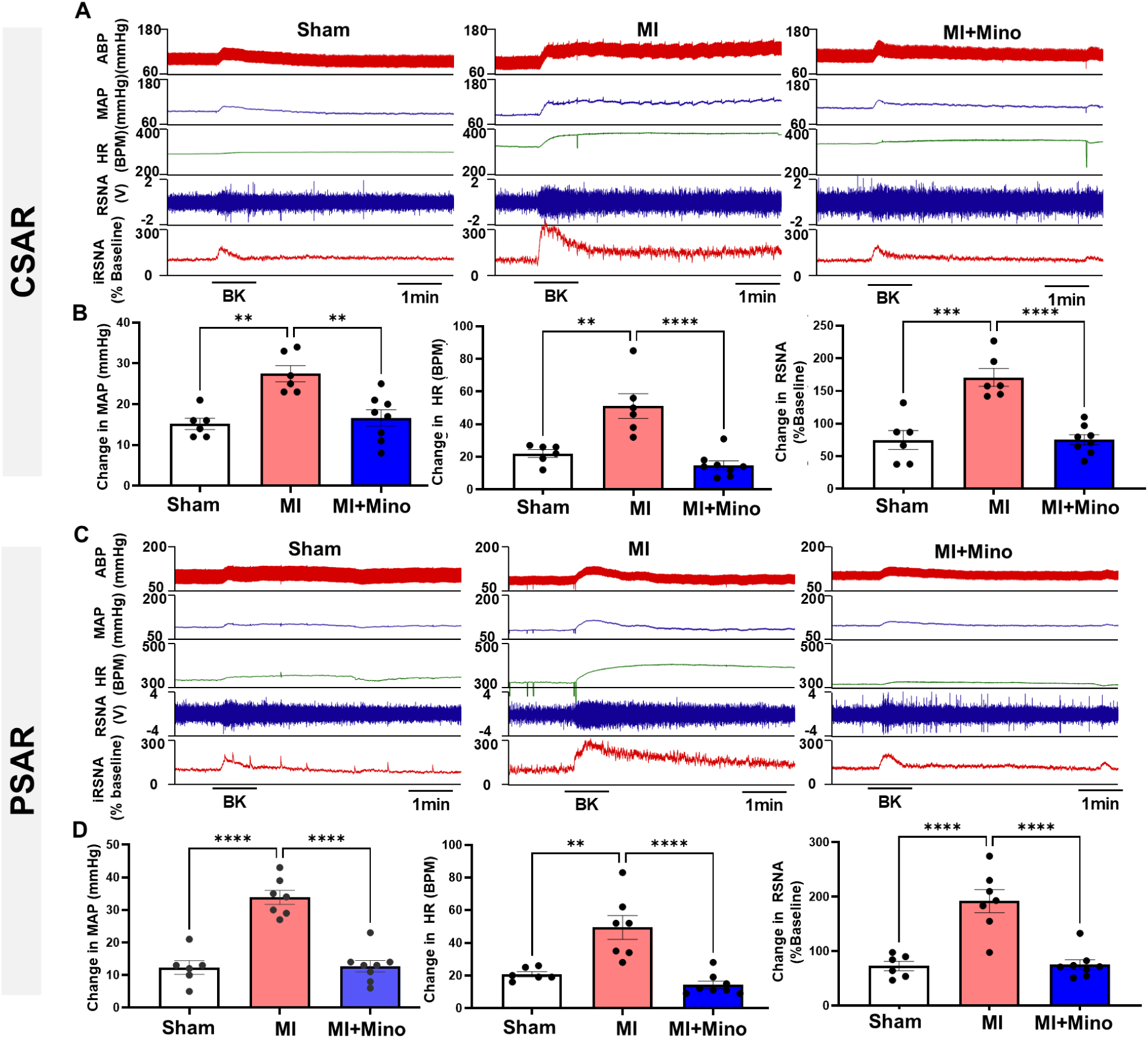
Representative tracing (**A** and **C**) and summary data (**B** and **D**) show cardiac and pulmonary functional changes including mean arterial pressure (MAP), heart rate and renal sympathetic nerve activity (RSNA) in response to epicardial or pulmonary topic application of bradykinin (BK) in sham, veh-treated MI and minocycline-treated MI rats. ** *P<*0.01 and *** *P<*0.001.

### 7. Liposomal clodronate–mediated macrophage depletion attenuates exaggerated CSAR and PSAR in post-MI rats

To independently confirm the role of macrophages, we depleted peripheral macrophages using tail vein injection of clodronate liposomes. CD68-positive macrophages in the heart, lung, and spleen were completely depleted in post-MI rats following treatment (**Supplemental Figure S3**). In thoracic DRGs, Iba-1–positive cells were substantially reduced in 4-week post-MI rats treated with clodronate (**Figure 8A, B**).

**Figure 8.**
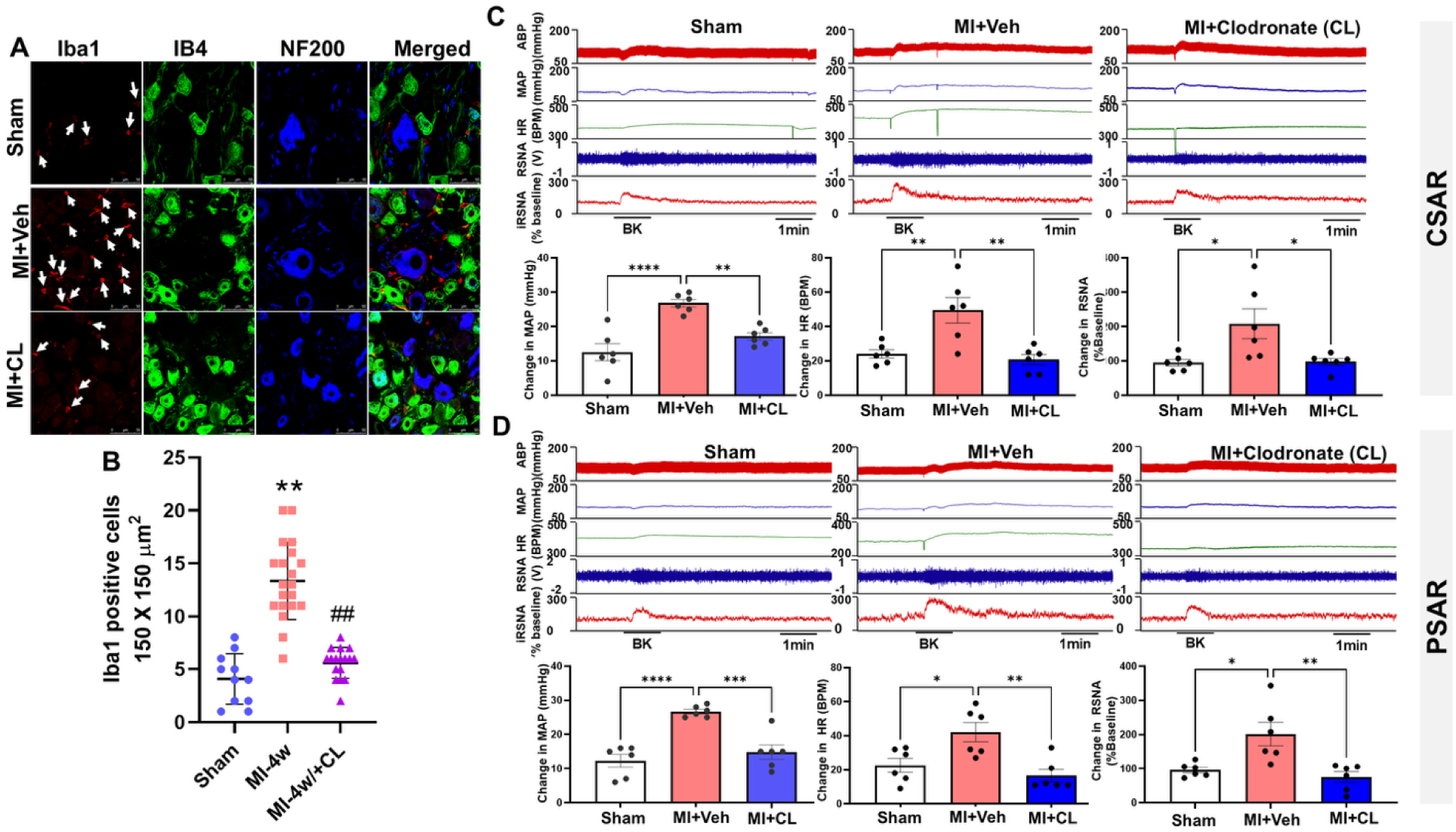
The effects of clodronate liposomes on macrophage infiltration in thoracic DRGs (**A** and **B**) as well as cardiac (**C**) and pulmonary (**D**) spinal afferent reflexes in in sham, veh-treated MI and clodronate liposomes (CL)-treated MI rats. * *P<*0.05 and ** *P<*0.01.

Functionally, clodronate liposome treatment prevented the exaggerated CSAR and PSAR responses to BK (10 μg/ml) observed in vagotomized post-MI rats, consistent with the protective effects of minocycline (**Figure 8C, D**). There was no significant difference in infarct size (36.83±3.28% vs. 37.33±2.70%, Veh vs. Liposomal Clodronate, n=6/each group, *P*>0.05) between MI rats treated with vehicle or liposomal clodronate.

### 8. Local epidural delivery of ProGel-Dex restores Kv channel function, reduces neuronal hyperexcitability, and delays cardiac remodeling in post-MI rats

We next tested local modulation of macrophage activation by epidural delivery of ProGel- Dex (a thermoresponsive dexamethasone prodrug that forms hydrogel upon injection *in vivo* ^39,40^) to T1–T4 DRGs. ProGel-Dex normalized IRF8 protein expression (**Figure 9A, B**) and restored downregulated Kv channel expression (**Figure 9C, D**) in thoracic DRGs of post-MI rats.

**Figure 9.**
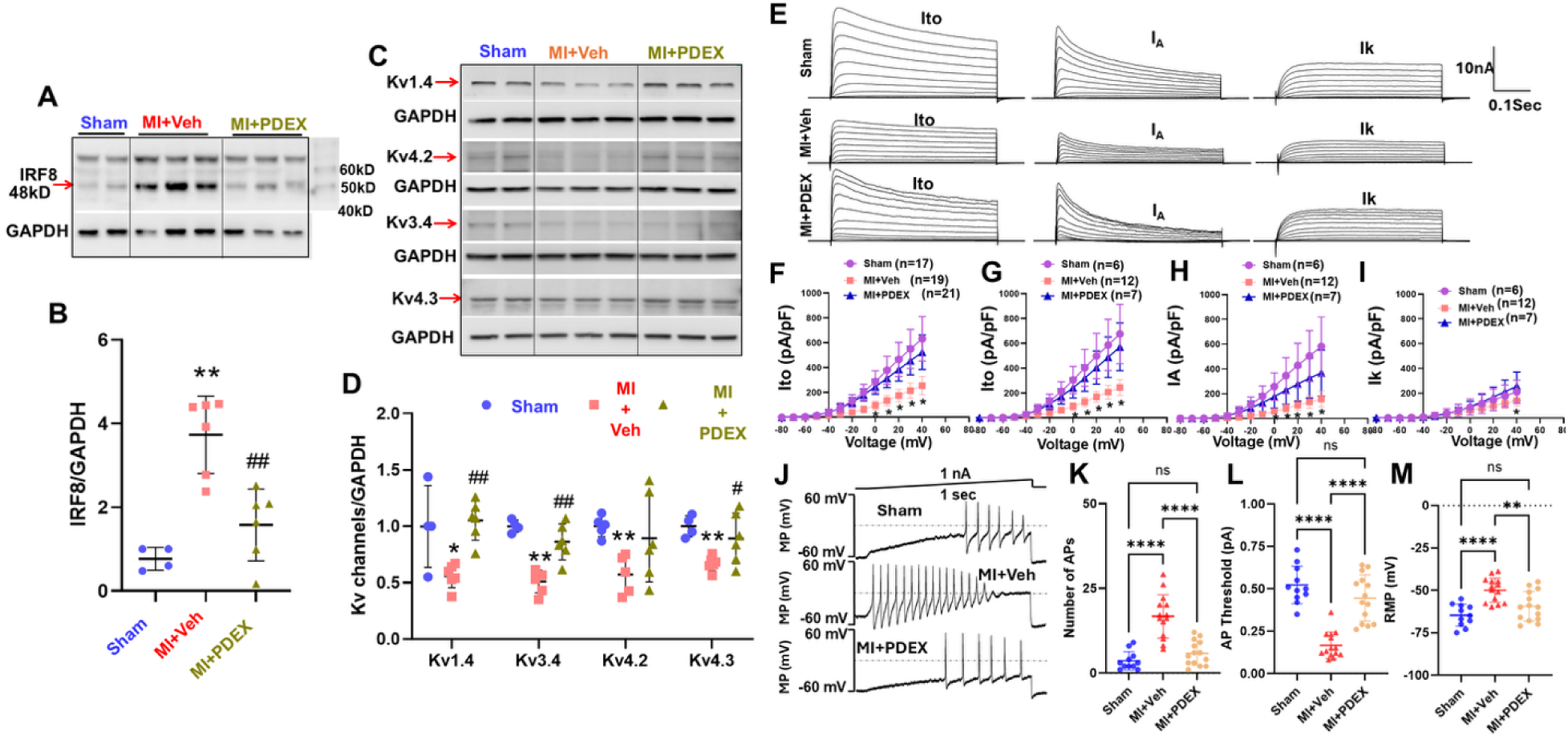
The effects of epidural delivery of ProGel-Dex on protein expressions of IRF8 (**A, B**) and Kv channels (**C, D**) as well as Kv currents (**E, F**) and cardiac DRG neuronal excitability in sham and MI rats with or without the treatment of ProGel Dex. \*\**P<*0.05 *vs.* sham rats, ^#^*P<*0.05 *vs.* MI rats.

Patch-clamp recordings in cardiac DRG neurons showed that MI reduced Ito, IA, and IK current densities (**Figure 9E–I**), increased action potential number (**Figure 9J, K**), lowered threshold potential (**Figure 9L**), and depolarized resting membrane potential (**Figure 9M**). ProGel-Dex treatment largely reversed these electrophysiological abnormalities, reducing neuronal hyperexcitability.

Functionally, ProGel-Dex prevented the exaggerated CSAR and PSAR responses to BK (10 μg/ml) in 4-week post-MI rats (**Figure 10A–D**), similar to systemic macrophage inhibition. Echocardiography revealed that ProGel-Dex delayed adverse cardiac remodeling, particularly left ventricular chamber dilation, although basal systolic function was not significantly improved (**Figure 10E–J**). There was no significant difference in infarct size (36.50±1.95% vs. 35.57±1.74%, Veh vs. ProGel-Dex, n=7-10/each group, *P*>0.05) between MI rats treated with vehicle or ProGel-Dex. These results demonstrate that local epidural therapy targeting thoracic DRGs is sufficient to mitigate neuronal dysfunction, afferent hyperexcitability, and reflex exaggeration after MI.

**Figure 10.**
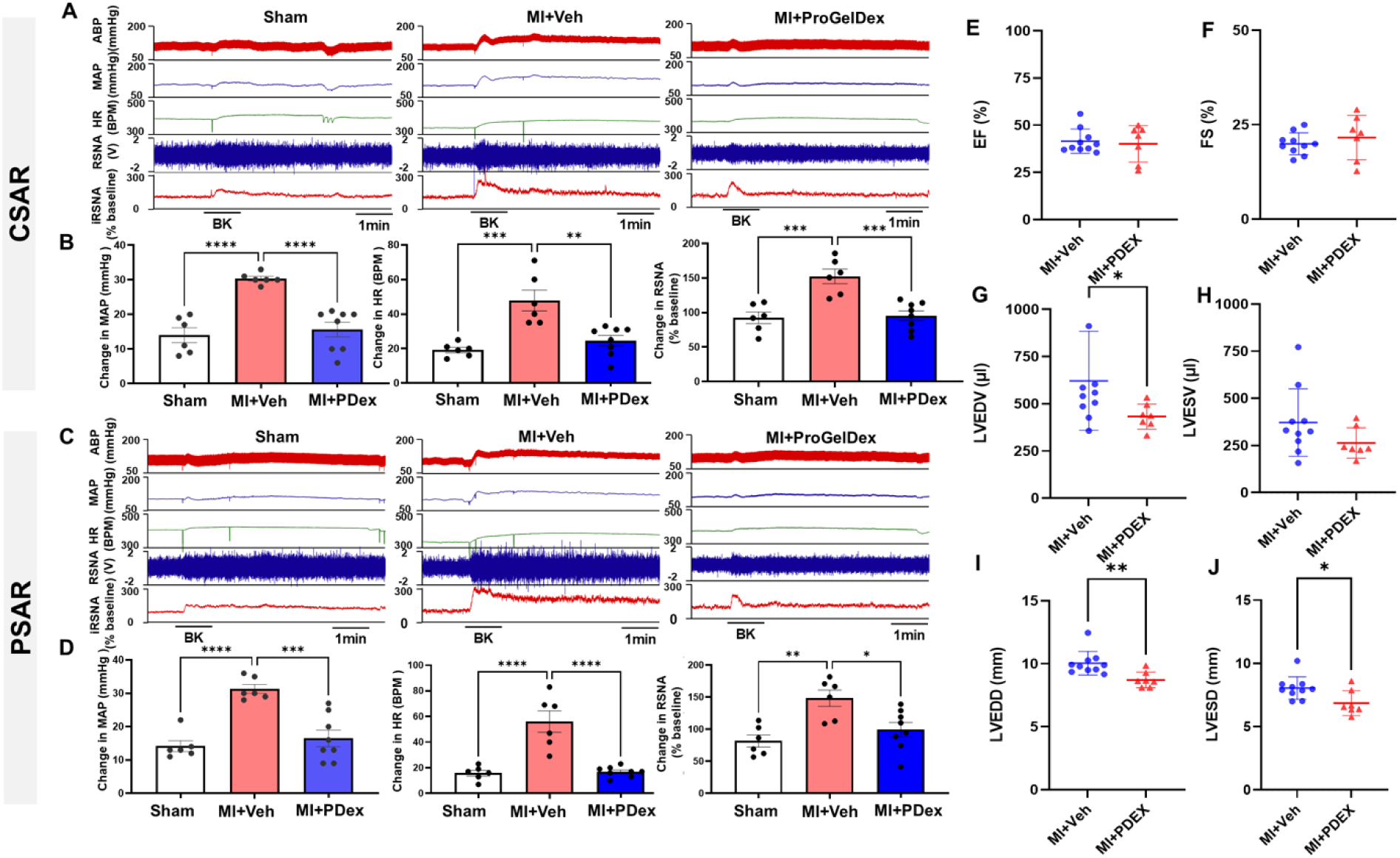
The effects of epidural delivery of ProGel-Dex on cardiac (**A** and **B**) and pulmonary (**C** and **D**) spinal afferent reflexes as well as cardiac function (**E-J**) in MI rats with or without the treatment of ProGel Dex. \**P<*0.05, *\*\*P<*0.01, and *\*\*\*P<*0.001.

### 9. Cytokine uptake by cardiac sensory afferents triggers macrophage recruitment to thoracic DRGs

To investigate why macrophages are activated in DRGs after MI, we hypothesized that cytokines produced by the injured heart are taken up by cardiac sensory nerve terminals and retrogradely transported to DRG somata, where they are released to recruit macrophages. To test this, we injected biotin-labeled TNFα (1 µL, 1.25 ng/µL, three injection sites in the left ventricular subepicardium) in rats. Saline-injected rats and L4/L5 DRGs served as controls. Biotin-TNFα was detected in T1–T4 DRG neuronal somata at day 1 post-injection, but not in controls (**Figure 11A, B**). Uptake occurred predominantly in NF200-positive A-fiber neurons and, to a lesser extent, CGRP-positive peptidergic C-fiber neurons, but not IB4-positive non-peptidergic C-fiber neurons. Similar results were obtained with biotinylated IL-6. Biotin-TNFα was also detected in stellate ganglia neurons, indicating sympathetic uptake (**Supplemental Figure S4**).

**Figure 11.**
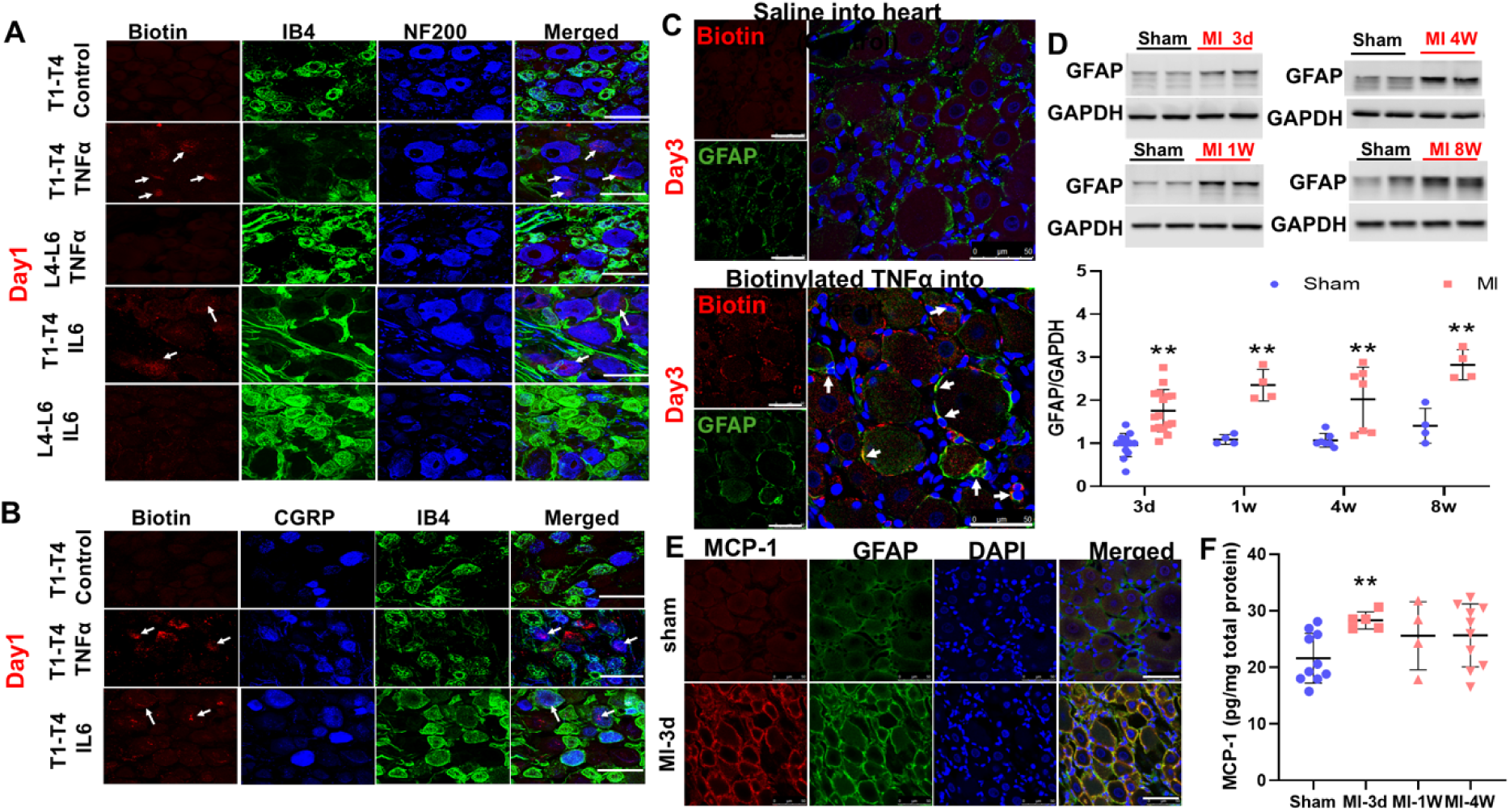
Immunofluorescence staining data (**A** and **B**) demonstrate that exogenous biotinylated TNFα in the sub epicardium of the left ventricle in normal rats can be taken up by NF200-positive and CGRP-positive spinal <u>afferents</u> (but not IB4-positive) and transported back to the DRG soma. **C**, biotin-labeled TNFα were taken up by satellite glia cells (SGCs) in T1-T4 DRGs 3 days after sub epicardial injection of cytokines. **D,** western blotting data show a time-dependent SGC activation in T1-T4 DRGs post MI. **E,** representative immunofluorescence images show the increased MCP-1 protein expression in SGCs rather than DRG neurons 3 days post MI. **F,** MCP- 1 protein expression measured by ELISA assay in T1-T4 DRGs in sham and myocardial infarction (MI) rats. IB4, a non-peptidergic C-fiber neuron marker; NF200, an A-fiber neuron marker; CGRP, a peptidergic C-fiber neuron marker. GFAP, a satellite glia cell marker.

At day 3, biotin-TNFα was released into adjacent GFAP-positive satellite glial cells (SGCs, **Figure 11C**). GFAP expression in DRGs was elevated from day 3 through 8 weeks post-MI (P<0.01 vs. sham, **Figure 11D**), preceding the peak of IRF8 expression. MCP-1 levels, assessed by ELISA, were transiently increased at day 3 post-MI (P<0.05, **Figure 11E, F**) and localized primarily in GFAP-positive cells, implicating SGCs as an early cytokine source. Biotin-TNFα injection also increased Iba-1–positive macrophages in DRGs by day 7 (**Supplemental Figure S5**).

To determine whether cytokine uptake depends on TNFα receptors, we injected biotin- TNFα into wild-type (WT) and p55/p75 double knockout (KO) mice. In WT mice, biotin-TNFα was detected in A-fiber DRG neurons at day 1, and macrophage numbers increased by day 7 (**Figure 12A, B**). In contrast, neither cytokine uptake nor macrophage recruitment was observed in p55/p75 KO mice. No changes were seen in saline-injected controls or in L4–L6 DRGs.

**Figure 12.**
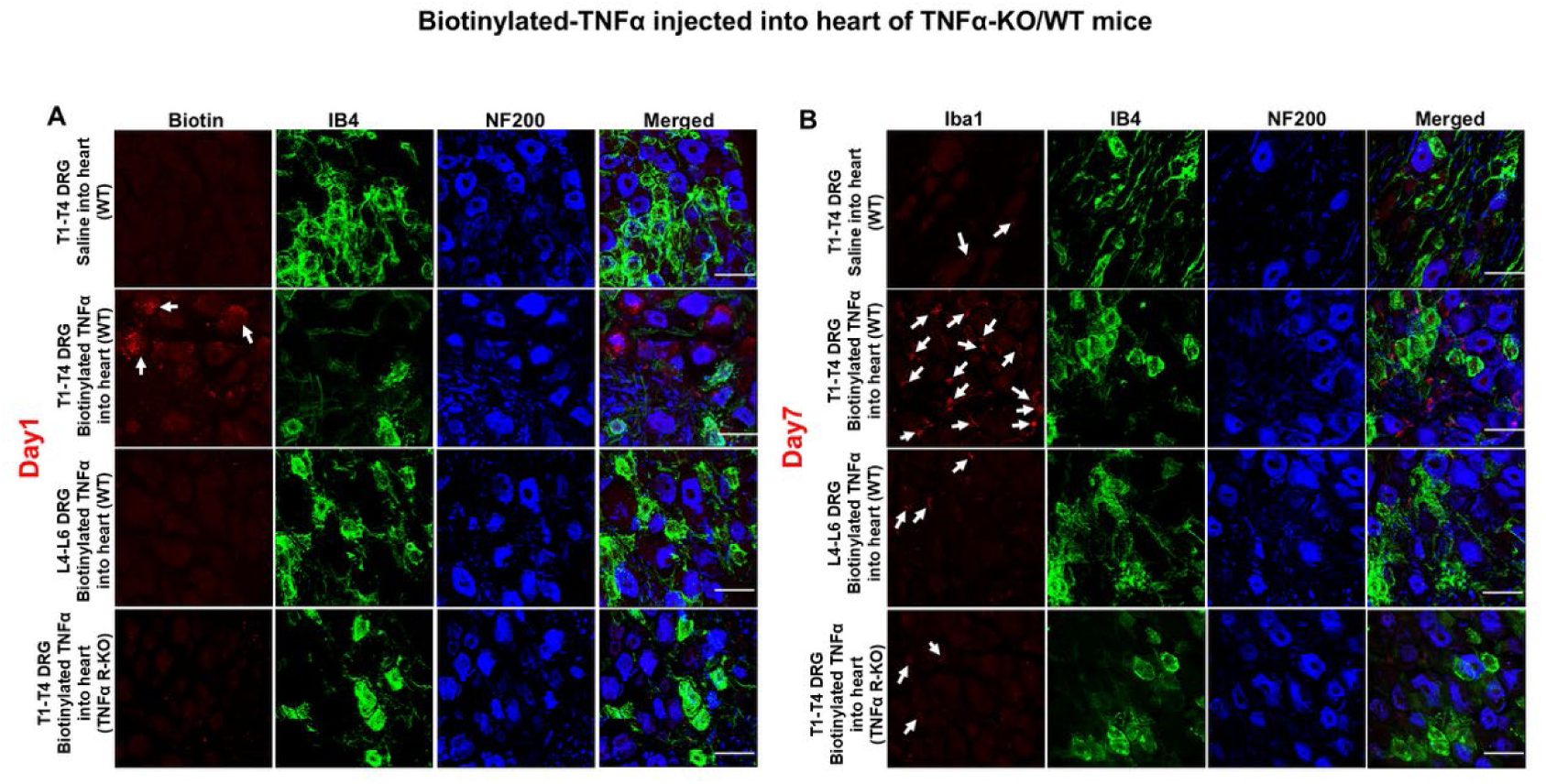
Immunofluorescence staining data demonstrate that injection of exogenous biotinylated TNFα into sub epicardium of the left ventricle caused retrograde TNFα transport (**A**) to T1-T4 DRGs at day 1 after injection and subsequently increased Iba1-positive cells (**B**) in T1- T4 DRGs at day 7 after injection in wide type mice but not in p55/p75 KO mice. IB4, a C-fiber neuron marker; NF200, an A-fiber neuron marker. White bar =50 μm.

## Discussion

This study provides very solid evidence that macrophages are activated in a time-dependent manner in cardiopulmonary sensory ganglia after MI, shaping the inflammatory milieu within the DRG microenvironment. Our *in vitro* co-culture experiments demonstrated that activated macrophages sensitize cardiopulmonary DRG neurons, most likely through downregulation of Kv channels. *In vivo*, we found that MI produced a time-dependent downregulation of Kv channels in T1–T4 DRG neurons. Importantly, pretreatment with a microglia/macrophage inhibitor (minocycline), systemic macrophage depletion (clodronate liposomes), or local epidural delivery of ProGel-Dex completely prevented macrophage activation in T1–T4 DRGs at 4 weeks post-MI. Functionally, these interventions attenuated exaggerated CSAR and PSAR responses. These results suggest that MI-induced macrophage activation in cardiopulmonary sensory ganglia sensitizes cardiac afferents while simultaneously exaggerating pulmonary afferent sensitivity. We propose a new mechanism of pathological heart–lung communication mediated by macrophage activation in thoracic DRGs.

Our laboratory has long focused on the role of cardiac spinal afferents in mediating autonomic dysfunction in CHF.^7–13^ These afferents are silent in the normal state and were originally considered to be essential pathways only for transmission of cardiac nociception to the central nervous system during myocardial ischemia. However, studies from our laboratory and others^7–11,41,42^ have demonstrated that stimulation of these afferents increases sympathetic outflow, blood pressure (BP) and heart rate, and decreases baroreflex sensitivity.^7–11,39,40^ We have also demonstrated that the discharge of cardiac sympathetic afferents is increased and cardiac reflex responses of BP, HR, and sympathetic nerve activity (SNA) are exaggerated in CHF animals.^10,11,13^ These data suggest that these afferents become sensitized and are tonically active in CHF. Our studies confirmed that an enhanced cardiac spinal reflex contributes to autonomic dysfunction, including increased global sympathetic outflow and impaired baroreflex function in the CHF state.^13,43^ Recent work from our group using resiniferatoxin (RTX)-mediated ablation of TRPV1- positive afferents confirmed their critical role in promoting adverse cardiac remodeling, dysfunction, and renal impairment in CHF.^12,44^

The present study advances this field by proposing that MI-induced macrophage activation in T1–T4 DRGs underlies chronic sensitization of cardiac spinal afferents. Using Iba1 staining, IRF8 expression, tissue clearing, electron microscopy, and RNAseq analysis, we established a rigorous chain of evidence that macrophages infiltrate and become proinflammatory in thoracic DRGs after MI. IRF8 is a critical transcription factor that transforms microglia or macrophages into a reactive phenotype.^31^ Interestingly, macrophage numbers increased as early as 1 week post-MI, whereas IRF8 expression rose only at 4 and 8 weeks. IRF8 is involved in the commitment of proinflammatory M1 polarization.^45^ We speculate that the early increased number of macrophages in T1-T4 DRGs detected by Iba1 staining may be anti-inflammatory M2 macrophages or non-activated monocytes, which are eventually converted into pro-inflammatory M1 macrophages in T1-T4 DRGs at 4 and 8-weeks post MI because IRF8 protein expression was significantly elevated at both later time points. Future studies should characterize macrophage phenotypes across different stages of post-MI remodeling.

Comparing to the well-documented function of macrophages in DRGs of neuropathic pain, the impact of macrophage activation on visceral DRG neuron excitability is less well defined. For many years, the manipulation of neuron–macrophage/microglia communication has proven to be a viable approach for halting the development of neuropathic pain, and both macrophage and microglia targets are being considered for novel therapeutic approaches.^46–48^ Macrophages were also shown to invade DRGs of the segments that supply the inflamed tissue.^49,50^ However, whether macrophage infiltration occurs in the cardiac sensory nervous system post-MI remains unknown. Most importantly, there are very few studies examining how macrophage activation affects DRG neuronal excitability in both neuropathic pain and cardiopulmonary diseases. Our co-culture and patch-clamp studies show that activated macrophages reduce Kv channel expression and suppress Ito currents in cardiac DRG neurons, thereby increasing excitability. Individual cytokines (TNF- α, IL-1β, IL-6, IFN-γ) each contributed to Kv downregulation, albeit with distinct effects, underscoring the complexity of macrophage-derived signaling. Collectively, these findings provide strong evidence that macrophage activation sensitizes DRG neurons via Kv channel inhibition, a mechanism that contributes to exaggerated reflexes after MI.

We next investigated how macrophages are recruited to DRGs post-MI. Traditional models of neuropathic pain emphasize ATP-driven microglial purinergic signaling, which activates cytokine production and leads to the sensitization of secondary dorsal root neurons in the spinal cord, ultimately resulting in hyperalgesia.^27,51–53^ However, this does not explain macrophage infiltration into DRGs, where synaptic transmission among DRG neurons is absent. A recent study by Simeoli et al. ^54^ suggested a hypothesis of macrophage activation via nociceptive stimuli-triggered extracellular vesicles from DRG neuronal soma. Complementing this, extensive work demonstrates that sensory nerve terminals internalize nerve growth factor (NGF) from peripheral innervated organs and retrogradely transport it back to the DRG soma,^54–57^ and likewise take up some retrograde neuronal tracers such as 1,1-dioctadecyl-3,3,3,3-tetramethylindocarbocyanine percholate (DiI) and fast blue.^58–61^ Inspired by these observations, we proposed that organ-derived cytokines are directly taken up by peripheral nerve terminals and conveyed retrogradely to DRG soma. Cytokines within DRG soma can be communicated with surrounding satellite glial cells to attract macrophage infiltration around DRG ganglia via releasing the chemokine MCP-1. Our data strongly support this model: biotinylated TNFα and IL-6 injected into the heart were retrogradely transported to T1–T4 DRG neurons and stellate ganglia, released into GFAP-positive satellite glial cells, and induced MCP-1 production, ultimately leading to macrophage recruitment. Importantly, TNFα uptake and macrophage infiltration were abolished in TNFR knockout mice, confirming receptor dependence. Myers et al. showed that exogenous cytokines, including biotin-labeled TNFα, when injected directly into the sciatic nerve trunk, were retrogradely transported to the DRG soma within six hours.^62^ However, those studies did not address whether end-organ sensory nerve terminals can directly take up cytokines, transport them back to the DRG, and initiate macrophage infiltration. Building on that foundation, our results demonstrate that sensory nerve terminals within peripherally innervated target organs can indeed mediate cytokine uptake and retrograde transport to DRG somata *in vivo*. Moreover, while Myers et al. ^62^ focused on neuropathic pain mechanisms by targeting the sciatic nerve, our work revealed a broader phenomenon: autonomic nerve endings, such as cardiac sympathetic efferents, can also take up cytokines from the heart and transport them back to the stellate ganglia soma.Thus, our study expands the concept beyond sensory nerves to show that both sensory and autonomic nerve terminals can internalize cytokines from a damaged organ in a non-neuropathic pain context. Beyond documenting this nerve-mediated cytokine uptake after MI, we further elucidated downstream mechanisms, including a role for the satellite glia cells in mediating cytokine- triggered macrophage infiltration. We observed that GFAP expression increased as early as three days post-MI, coinciding with elevated MCP-1 levels in GFAP-positive cells. In TNFR knockout mice, biotinylated TNFα failed to enter T1–T4 DRGs post-MI, and no macrophage infiltration was detected at day seven after TNFα injection. Collectively, our findings suggest a new paradigm: nerve terminals can directly internalize cytokines and transport them back to the neuronal soma, where they are transferred to adjacent satellite glial cells, stimulate chemokine release (e.g., MCP- 1), and ultimately drive macrophage infiltration around DRG neurons following MI.

To investigate whether cardiac ganglionic inflammation contributes to the sensitization of CSAR after MI, we employed three anti-inflammatory strategies: systemic administration of minocycline, systemic delivery of clodronate-containing liposomes, and local application of ProGel-Dex to attenuate neural inflammation in thoracic DRGs. Minocycline, which crosses the blood–brain barrier (BBB) and inhibits microglial activation in the brain,^35,36,38,63^ may exert its effects in part via the central nervous system, a possibility we cannot exclude. Similar issues exist with the liposomal clodronate-induced systemic macrophage depletion approach. Nevertheless, we found that minocycline completely prevented the post-MI upregulation of IRF8 protein in T1– T4 DRGs and significantly attenuated the exaggerated CSAR in MI rats at four weeks.

To selectively deplete peripheral macrophages, we next used intravenous injection of clodronate-containing liposomes.^64–71^ Prior work has shown that liposomal clodronate reduce macrophages and TNFα expression in DRGs of paclitaxel-treated rats.^67^ In contrast, Pent et al. reported that intraperitoneal injections of liposomal clodronate liposomes failed to deplete macrophages in DRGs of CX3CR1GFP/+ mice.^72^ Consistent with these mixed findings, our data demonstrated that clodronate-containing liposomes largely, but not completely, depleted macrophages in the DRGs of MI rats, despite robust depletion in peripheral organs, such as the heart, lungs, and spleen. This suggests that while tail-vein injections effectively eliminate circulating macrophages, they may not fully target locally proliferating macrophages within the DRGs after MI. In other words, both infiltrating macrophages and local macrophage proliferation contribute to the heightened macrophage activity in thoracic DRGs post-MI. Importantly, even with incomplete macrophage depletion in DRGs, clodronate treatment still significantly reduced the CSAR in MI rats, reinforcing the role of macrophage-driven inflammation in this process.

To overcome the limitations of systemic minocycline and liposomal clodronate administration, we delivered ProGel-Dex epidurally to T1–T4 DRGs to locally suppress thoracic ganglionic inflammation. ProGel-Dex is a *N*-(2-hydroxypropyl)-methacrylamide (HPMA) copolymer-based thermoresponsive macromolecular prodrug of dexamethasone (Dex), which forms hydrogel upon *in vivo* injection. HPMA copolymer prodrugs have been widely use as cancer chemotherapeutic agents with improved efficacy and safety.^73–80^ As a new member of the HPMA copolymer-based prodrugs, ProGel-Dex is a thermoresponsive macromolecular prodrug of Dex. Aqueous solution of ProGel-Dex remains a free-flowing liquid at room temperature but transitions into a hydrogel at physiological temperature (>30 °C). This phase-transition property enables local deposition of the prodrug with sustained retention (>1 month), producing outstanding therapeutic efficacy and minimal adverse effect observed.^39,81^ Once administered *in vivo*, ProGel-Dex will gradually solubilizes when exposed to interstitial fluid. It may then be activated by the pathology-associated tissue acidosis or be activated intracellularly to release Dex when internalized by phagocytic cells. These unique prodrug design features make ProGel-Dex ideally suited for selective and chronic suppression of macrophage activation within thoracic DRGs post-MI.

Our experimental findings support ProGel-Dex’ therapeutic potential. When ProGel-Dex was epidurally administered, it effectively suppressed macrophage activation in thoracic DRGs post-MI. Patch-clamp recordings further showed that ProGel-Dex largely restored neuronal excitability and normalized Kv channel function. Notably, a single epidural injection of ProGel- Dex immediately after MI significantly limited adverse cardiac remodeling, including attenuation of chamber dilation and reductions in systolic and diastolic diameters and volumes, without altering resting ejection fraction. These effects parallel our previous findings with RTX-mediated cardiac afferent ablation,^12^ supporting the idea that silencing cardiac spinal afferents benefits cardiac remodeling post-MI. Clinically, White et al. .^3^ reported that left ventricular end-systolic volume, rather than ejection fraction, is the primary determinant of survival in post-MI patients Thus, the ability of ProGel-Dex to reduce cardiac volumes suggests a promising therapeutic avenue for improving long-term outcomes after MI.

Finally, our study demonstrates, for the first time, that activation of pulmonary spinal afferents by topical bradykinin produces exaggerated pressor and sympatho-excitatory responses in vagotomized rats at 4 weeks post-MI compared with sham controls. While most research on pulmonary sensory input has focused on vagal afferents in regulating respiratory function,^82^ little attention has been given to the role of pulmonary spinal afferents in cardiovascular and respiratory regulation. Our previous work suggested that pulmonary spinal afferents are sympatho- excitatory,^28^ and the present study extends this by showing that myocardial injury chronically sensitizes these afferents. This finding has important clinical implications. The heightened sensitivity of pulmonary spinal afferents observed at 4 weeks post-MI suggests that patients recovering from MI may be particularly vulnerable to pulmonary irritants such as smoke or air pollution during a critical time window (several months in humans). Importantly, treatment with minocycline, clodronate, or ProGel-Dex prevented the exaggerated PSAR in MI rats, indicating that macrophage activation in thoracic DRGs enhances both CSAR and PSAR. Together, these data support a novel paradigm in which myocardial injury initiates a neuro-inflammatory cascade involving macrophage activation in autonomic ganglia shared by the heart and lungs, leading to cross-organ afferent sensitization. Given that DRGs contain heterogeneous neurons that innervate multiple organs, cross-sensitization of afferent pathways provides a plausible mechanism by which pathology in one organ can influence sensory function in another. To our knowledge, this represents a new form of organ-to-organ interaction mediated by the peripheral nervous system, which may extend beyond the heart–lung axis.

In summary, this study identifies a novel cytokine uptake–glial activation–macrophage activation pathway as a key driver of cardiopulmonary afferent sensitization after MI (**Figure 13**). A paradigm-shifting concept emerging from our work is the existence of a previously unrecognized neural crosstalk mechanism mediating heart–lung interactions post-MI. From a clinical perspective, targeting DRG inflammation, particularly through localized dexamethasone delivery, represents a precise therapeutic strategy to disrupt this pathological neural crosstalk, attenuate sympathetic activation, and improve cardiac outcomes.

**Figure 13.**
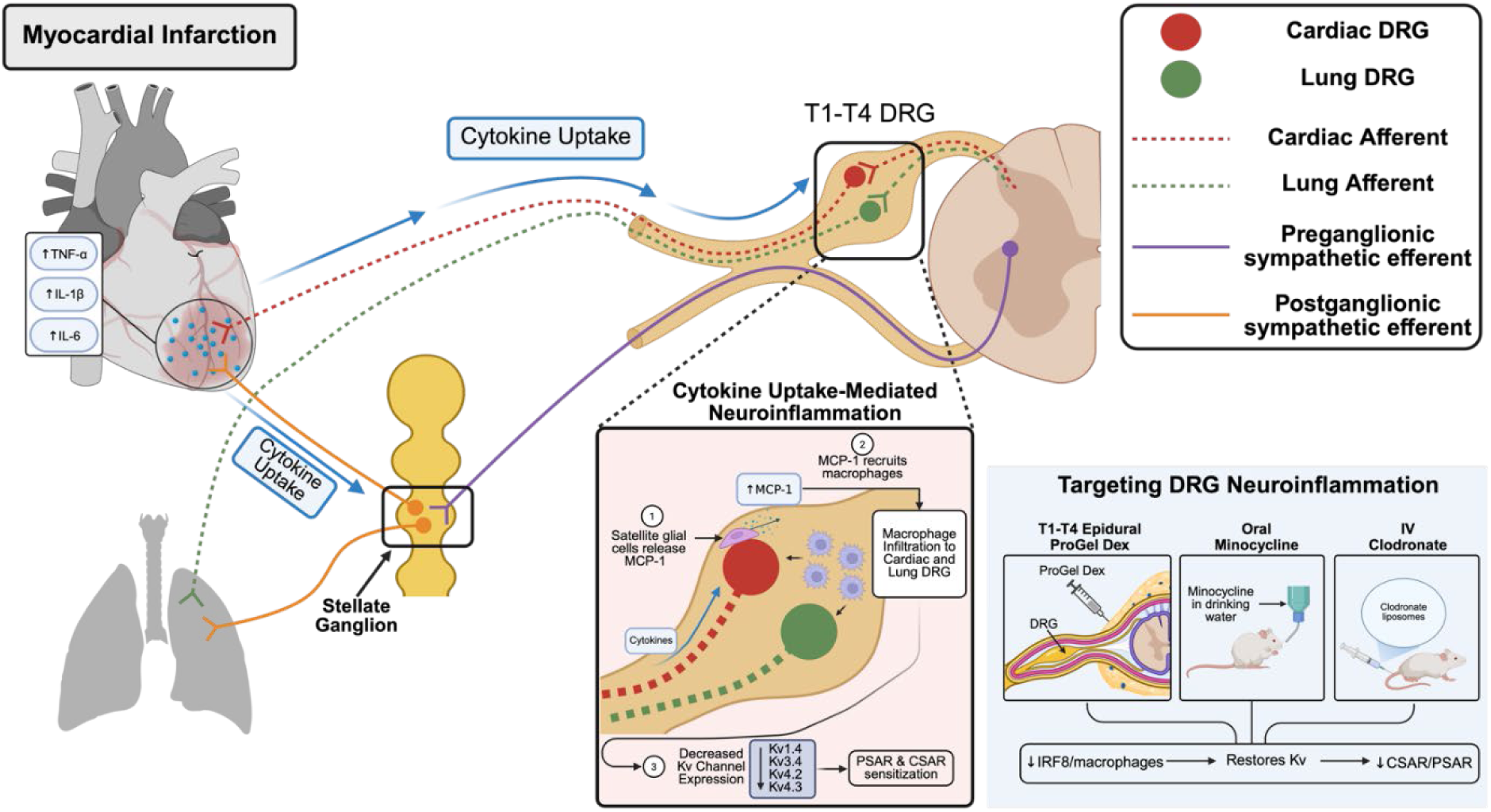
A schematic figure summarizing the molecular mechanisms underlying heart-lung neural crosstalk.

## Acknowledgments

We sincerely thank Dr. Bryan Hackfort and the UNMC Echocardiography Core Facility for their support with all echocardiography studies.

## Methods

All animal experimentation was approved by the Institutional Animal Care and Use Committee of the University of Nebraska Medical Center and performed in accordance with the National Institutes of Health’s Guide for Use and Care of Laboratory Animals and in accordance with the ARRIVE guidelines.^83,84^ Adult male and female Sprague-Dawley rats weighing 180-200g were purchased from Charles Rivers.

For cytokine uptake experiments, ∼10-week-old male TNFα receptor knockout (p55/p75 KO; B6.129S, Jackson Laboratory) and their wild-type controls (B6129SF2/J, Jackson Laboratory) were used. For tissue-clearing experiments, male heterozygous Cx3cr1CreER::TdTomato (mT/mG) mice were generated by crossing homozygous Cx3cr1CreER males (Stock No. 020940, Jackson Laboratory) with homozygous TdTomato (mT/mG) females (Stock No. 007676, Jackson Laboratory).

### Myocardial Infarct (MI)

MI was produced by left coronary artery ligation as described in our previous studies.^85,86^ Sham-operated rats were prepared in the same manner but did not undergo coronary artery ligation. Briefly, rats (180-200 grams) were chosen at random to undergo either MI surgery or sham surgery. Rat were ventilated at a rate of 60 breaths/min under 2%–3% isoflurane during the surgical procedure. A left thoracotomy was performed through the fifth intercostal space, the pericardium was opened, the heart was exteriorized, and the left anterior descending coronary artery was ligated. Buprenorphine SR (1.0 mg/kg) was subcutaneously injected at the beginning of surgery for pain management. All sham rats survived, and∼70% of rats survived from coronary artery ligation surgery. At the end of terminal experiments, infarct size (IS) was measured by taking the ratio of the infarct area to whole left ventricle (LV) minus septum in rats 4 and 8 weeks post sham/MI. In rats 3 days or 1 week post sham/MI, we did not measure IS because the scar has not completely formed although each infarcted heart exhibited a pale ischemic region.

In some experiments, four weeks after myocardial infarction (MI) or sham surgery, hemodynamics were measured by echocardiography (VEVO 2100, Visual Sonics, Toronto, ON, Canada). One, four and eight weeks after sham or MI surgery, the T1-T4 DRGs were collected to analyze the protein expression. RNA seq was performed eight weeks after MI surgery. In some experiments, the sham and MI rats were fed daily with minocycline in drinking water 3 days prior to the surgery (see below). In other experiments, epidural delivery of ProGel-Dex was performed immediately after MI. Four weeks after sham or MI surgery, the cardiac spinal afferent reflex (CSAR) and pulmonary spinal afferent reflex (PSAR) were determined.

### Treatment with drugs

Minocycline (macrophage activation inhibitor)^35,38,87^ and bradykinin (BK)^28^ were purchased from Sigma-Aldrich (Sigma, St. Louis, MO, USA). Minocycline were dissolved in drinking water to a final concentration of 20 mg/kg. BK was dissolved in sterile saline at a stock concentration of 100 µg/ml and was further diluted in sterile saline to a working concentration of10 µg/ml.^28^

Liposome-encapsulated clodronate was used to specifically deplete peripheral phagocytic macrophages.^67,72^ Clodronate liposomes (15 ml/kg/per day, ClodronateLiposomes.com) or control (PBS) were intravenously administrated in a volume of 2 mL for 3 continuous days prior to the MI surgery.

### General Surgical Preparation for Acute Experiments

For acute terminal experiments that were performed 4 weeks post-MI, rats were anesthetized with urethane (800 mg/kg ip) and α-chloralose (40 mg/kg ip). Anesthetic plane was monitored by establishing that rats were unresponsives to pedal withdrawal and corneal reflexes. The cervical vagal nerves were cut bilaterally to prevent any reflex responses observed from vagal afferent activation, so as to observe only responses related to cardiac spinal afferent activation. The trachea was cannulated, and rats were ventilated artificially with room air supplemented with oxygen (∼40% O2). A Millar catheter (SPR 524; size, 3.5-Fr; Millar Instruments, Houston, TX) was advanced through the right common carotid artery and progressed into the aorta and left in place to record arterial pressure (AP). Mean arterial pressure (MAP) and heart rate ^27^ were derived from the arterial pressure pulse using Chart 7.1 software and an analog to digital converter (PowerLab model 16S; AD Instruments, Colorado Springs, CO). The right jugular vein was cannulated for intravenous injections and administration of saline at a rate of 3 mL/h. Body temperature was maintained between 37 and 38 °C by a heating pad.

### Recording renal sympathetic nerve activity (RSNA)

In acute surgical preparations described above, RSNA was recorded as previously described.^7,88,89^ In brief, the left kidney, renal artery, and nerves were exposed through a left retroperitoneal flank incision. Sympathetic nerves running on or beside the renal artery were identified. The renal sympathetic nerves were placed on a pair of platinum–iridium recording electrodes and cut distally to avoid recording afferent activity. Nerve activity was amplified (×10000) and filtered (bandwidth: 100 to 3000 Hz) using a Grass P55C preamplifier. The nerve signal was displayed on a computer where it was rectified, integrated, sampled (1 kHz), and converted to a digital signal by the PowerLab data acquisition system. At the end of the experiment, the rat was euthanized with an overdose of pentobarbital sodium. Respective noise levels were subtracted from the nerve recording data before percent changes from baseline were calculated. Integrated RSNA (iRSNA) was normalized as 100% of mean baseline during the control period.^88^

### Activation of cardiac or pulmonary spinal afferents

Epicardial application of bradykinin has been demonstrated to effectively stimulate cardiac sympathetic afferents in vagotomized rats.^85,99,100^ The chest was opened through the fourth intercostal space. The pericardium was removed to expose the left ventricle. A square of filter paper (3 mm × 3 mm) saturated with BK (10 μg/ml) was applied to the anterior surface of the left ventricle and ventral surface of the left lung to stimulate regional cardiac sympathetic afferents and pulmonary spinal afferents. Hemodynamic and neural parameters were continuously recorded. After the peak response, the epicardium was rinsed three times with 10 ml of warm normal saline (38°C). The time interval between BK treatment was at least 20 min to allow the AP, HR and RSNA to return to, and stabilize at their control levels.

### Epidural delivery of ProGel Dex

To selectively inhibit macrophage activation in T1-T4 DRGs, rats were anesthetized using 2%-3% isoflurane:oxygen mixture. Rats were placed in the prone position and a small midline incision was made in the region of the T13-L1 thoracic vertebrae. Following dissection of the superficial muscles, two small holes (approximately 2 mm*2 mm) were made in the left and right sides of T13 vertebral. A polyethylene catheter (PE-10) was inserted into the epidural space via the left hole and gently advanced about 4 cm to the left T1 level in which the first injection (20% w/v ProGel-Dex concentration, 10 µl/per ganglion) was made at a slow speed to minimize the diffusion of ProGel Dex. The catheter was then pulled back to the left T2, T3 and T4 respectively to perform serial injections (10 ul/each) at each segment. The same injections were repeated on the right side. Silicon gel was used to seal the hole at the T13 vertebra. The skin overlying the muscle was closed with 3-0 polypropylene suture. Simple interrupted sutures were used to close the skin. Skin suture was removed 10-14 days after surgery. The rats were allowed to recover from the anesthesia.

Buprenorphine SR (1.0 mg/kg) was subcutaneously injected at the beginning of surgery for pain management.

### *In vitro* electrophysiology recording: Patch clamp

Rats were given an overdose of pentobarbital sodium (150 mg/kg, i.p.) and T1-T4 DRGs were dissociated by a collagenase digestion procedure described previously.^90–92^ Dissociated DRGs were suspended in Dulbecco’s modified Eagle’s medium and stored in an incubator at 37° C until used, usually within 6 h of isolation. Aliquots of DRGs were transferred to a cell chamber mounted on the stage of an inverted microscope and superfused with an external solution.

Voltage gated potassium currents (Kv currents) were recorded using the whole-cell patch-clamp technique using Axopatch 200B patch-clamp amplifier (Axon Instruments, Burlingame, CA, USA) only in DiI-labeled small and medium-sized (<35 µm) DRG neurons., we used Griffonia simplicifolia isolectin B4 (IB4, Alexa Fluor® 488 conjugate, Invitrogen, Carlsbad, CA, USA) to further confirm cardiac C fiber-positive neurons from these DRG subpopulations. Only DiI-labeled IB4-positive DRG neurons were recorded. Briefly, DRG neurons were incubated with IB4 for 20 min before recording. The IB4-stained neurons were easily recognized under epifluorescence illumination.^92^ In voltage-clamp experiments, borosilicate glass capillaries were pulled (Sutter Instruments, Novato, CA) to an internal tip diameter of 1 to 2 μm and filled with a pipette solution containing (in mmol/l) 135 KCl, 3 MgCl2, 10 HEPES, 3 Na2-ATP, 10 EGTA, and 0.5 Na-GTP, pH 7.2, 320 mosm liter^-1^. The extracellular solution consisted of (in mM): 138 choline-Cl, 0.5 CdCl2 (to block Ca^2+^ channels), 4 KCl, 1.2 MgCl2, 5 HEPES, 1 CaCl2 and 18 glucose (pH 7.4; 330 mosm liter^-1^). Filled pipettes with a resistance of 2–4 MΩ were coupled to a patch-clamp amplifier (Axopatch 200B, Molecular Devices, Sunnyvale, CA). After correction of the liquid junction potential and creation of a GΩ seal, the membrane within the pipette was ruptured, and at least 5 min were allowed for the contents of the pipette and cytoplasm to equilibrate. A computer program (pClamp, Molecular Devices) controlled command potentials and acquired current signals that were filtered at 2 kHz. Currents were sampled at 4 kHz by a 12- bit resolution analog-to-digital converter and stored on the hard disk of a computer. All electrophysiological experiments were done at room temperature (22–24°C). The holding potential was initially set at −80 mV. During recording, the *I*to was evoked by 500-ms depolarizing pulses to test potentials between −80 and +40 mV. For each test pulse, *I*to amplitude was measured as the difference between the peak outward current and the steady-state current level at the end of the depolarizing pulse. 4-aminopyridine (4 AP) and Tetraethylammonium (TEA) were applied to the bath solution to test for the presence of the transient (IA) and delayed rectifier (IK) K^+^ currents, respectively.^93–96^ The TEA or 4 AP-sensitive current were defined as the difference between the outward current recorded in a drug-free bath solution and the current elicited after cell superfusion of 1 mM TEA or 4 AP, respectively. All electrophysiological data were normalized as current densities by dividing measured current amplitude by whole cell capacitance.

In the current-clamp experiments, APs were elicited by a ramp current injection of 0–1 nA and the current threshold-induced APs or threshold potential was measured at the beginning of the first AP. The number of APs was also measured. The patch pipette solution was composed of (in mM) 105 K-aspartate, 20 KCl, 1 CaCl2, 5 MgATP, 10 HEPES, 10 EGTA and 25 glucose (pH 7.2; 320 mosM). The bath solution was composed of (in mM) 140 NaCl, 5.4 KCl, 0.5 MgCl2, 2.5 CaCl2, 5.5 HEPES, 11 glucose and 10 sucrose (pH 7.4; 330 mosM).

The P-clamp 10.2 program (Axon Instruments, San Jose, CA, USA) was used for data acquisition and analysis. All experiments were done at room temperature (22–24 °C).

### Cell culture and treatment

50B11 cells (DRG neurons), which were kindly supplied by Professor Ahmet Hoke of Johns Hopkins University, was cultured in NeuroBasal medium (Life technologies, Waltham, MA,USA) supplemented with 10% fetal bovine serum (FBS) (Life technologies), 0.27% L-Glutamine (Life technologies), 1% penicillin and streptomycin (Life technologies), 2% B-27 (Life technologies) and 1.1% Glucose (20%, Sigma). The BV2 cell line was purchased from Banca Biologica e Cell Factory (ICLC ATL03001) and was cultured in RPMI 1640 (Life technologies) supplemented with 10% FBS and 2mM L-Glutamine. The RAW264.7 cell line was purchased from ATCC (TIB-71) and was cultured in DMEM (high glucose, no HEPES) supplemented with 10% FBS. All cell lines were maintained at 37°C in a CO2 incubator.

To identify the chronic effect of multiple cytokines on 50B11 cells, the 50B11 cells were seeded into 25 cm^2^ flasks. All the cytokines such as TNFα, IL-1β, IL-6, IFNγ, IL-10 and IL-4 are purchased from R&D Systems, Inc. (Minneapolis, MN, USA). After attaching onto the bottom, 50B11 cells were differentiated with Fosklin (75 µM) for 24 hours, then the NeuroBasal medium of 50B11 cells containing Fosklin (100 µM) with or without cytokines were applied every 24 hours twice, and experiments were carried out for 72 hours. Then the adherent cell monolayer at the bottom of flasks was washed with PBS and total cell protein was extracted.

### Co-culture of 50B11 DRG with BV2 or RAW264.7 cell line

Microglia and macrophages, which had been stimulated by LPS (25 ng/ml) were co-cultured with 50B11 cells in a Transwell chamber consisting of a 10-µm thick porous membrane with 0.4-µm pores in the culture inserts (Corning Incorporated, corning, NY, USA). This methodology was used to construct an indirect co-culture system. Briefly, the 50B11 cells were seeded in the wells of a 12-well (5x10^4^ cells/well) plate (lower layer) at 37°C prior to co-culture with macrophages. Before co-culture, the 50B11 cells were differentiated with Forskolin (75 µM) for 24 hours. The BV2/RAW264.7 (microglia/macrophage) cells were individually grown in the upper layer of the Transwell, and pretreated with LPS for 4 hours. Then, macrophage cells in fresh medium containing the inserts were added to the upper chamber of the Transwell system, with the 50B11 cells located at the bottom. NeuroBasal medium of 50B11 cells containing Forskolin (100 µM) and the BV2/RAW264.7 treated with or without LPS were changed every 24 h, and experiments were carried out for co-culture at 72 h. The adherent cell monolayer at the bottom of transwells was washed with PBS, then total cell protein was extracted.

### Western Blot Analysis

Rats (n=4-6/each group) were anesthetized with pentobarbital sodium (40 mg/kg, i.p.). The T1-T4 dorsal root ganglions (DRGs) were rapidly removed and lysed with 20 mM Tris-HCl buffer, PH 8.0, containing 1% NP-40, 150 mM NaCl, 1 mM EDTA, 10% glycerol, 0.1% β- mercaptoethanol, 0.5 mM dithiothreitol, and a mixture of proteinase and phosphatase inhibitors (Sigma). Protein concentration was measured by the BCA protein assay method using bovine serum albumin as standard (Thermo Scientific, Waltham, MA, USA). The proteins were loaded onto a 10% SDS-PAGE gel along with protein standards (Bio-Rad Laboratories, Berkeley, California, USA) in a separate lane for electrophoresis and then transferred to polyvinylidene fluoride membrane. The membrane was probed with mouse antibody against IRF8 (1:500, Santa Cruz Biotechnology, Dallas, USA) and rabbit antibodies against Kv1.4, Kv4.2, Kv4.3, and Kv3.4 (1:200, Alomone labs, Jerusalem, Israel) and secondary antibody of goat anti-mouse (1:5000, Invitrogen, Carlsbad, CA, USA), goat anti-rabbit IgG (1:5000, Invitrogen). The protein signals were detected by enhanced chemiluminescence reagent (Thermo Scientific) and analyzed using UVP BioImaging Systems. GAPDH (1:1000, Santa Cruz Biotechnology) was used to verify equal protein loading, and the densitometric results of IRF8 and Kv channel isoforms (Kv1.4, Kv4.2, Kv4.3, Kv3.4) were reported as the ratio to GAPDH and normalized to age-matched sham rats.

### Immunofluorescence staining

Rats were anesthetized with pentobarbital sodium (40 mg/kg, intraperitoneally) and then perfused with 4% paraformaldehyde (PFA). The T1-T4 DRGs were immediately dissected and post-fixed with 4% PFA overnight at 4 °C, and then were transferred to 30% sucrose until they sank to the bottom. 14-μm-thick sections were cut using a Leica cryostat and thawed onto gelatin-coated slides. Since the slices were 14 µm thick, and the DRG neuron soma ranged from less than 10 µm to over 100 µm, the same neuron could potentially be counted twice. Therefore, we counted every 7^th^ slice.

For triple-immunostaining of potassium channels, sections were stained with the isolectin B4 (a C-fiber neuronal marker, Invitrogen, I21411) and NF200 (an A-fiber neuronal marker, Sigma-Aldrich, N5389).^89^ After washing with PBS, sections were incubated in blocking serum (10% donkey serum in PBS) for 1 h and further incubated overnight with rabbit antibodies of Kv1.4, Kv4.2, Kv4.3, and Kv3.4 (1:100, Alomone labs) and mouse anti-NF200 antibody (1:200, Sigma-Aldrich, N5389) overnight at 4°C. After being washed in PBS, the sections were treated with PBS and sections were incubated with fluorescence-conjugated secondary antibody (Alexa 568- conjugated goat anti-rabbit IgG and pacific blue-conjugated goat anti-mouse IgG, 1:200, Invitrogen) and Alexa FluorR 488 conjugated isolectin-B4 (1:200, Invitrogen) for 60 min at room temperature. After three washes with PBS, the sections were mounted with an Aqua-Mount Mounting Medium and then were examined with a laser confocal microscope (Leica TSC STED). No staining was seen when a negative control was performed with PBS instead of the primary antibody (data not shown).

Counts were obtained from the total number of IB4- (C fiber) or NF200-positive (A fiber) neurons. The percentage of neurons positive for Kv4.2, Kv4.3, Kv3.4 and Kv1.4 for A or C fiber neurons was calculated. A neuron was considered to be “positive” when the measured intensity of immunostaining was more than five times greater than the background. Ten sections from each DRG were analyzed for a total of 5 rats in each group.

For triple-immunostaining of macrophages in DRGs, sections were stained with the isolectin B4 (a C-fiber neuronal marker, Invitrogen, I21411) and NF200 (an A-fiber neuronal marker, Sigma-Aldrich, N5389). After washing with PBS, sections were incubated in blocking serum (10% donkey serum in PBS) for 1 h and further incubated overnight with rabbit monoclonal antibody of Ionized calcium-binding adapter molecule 1 (Iba1, 1:100, Abcam, ab178846) and mouse anti- NF200 antibody (1:200, Sigma-Aldrich, N5389) overnight at 4°C. After being washed in PBS, the sections were treated with PBS, and sections were incubated with fluorescence-conjugated secondary antibody (Alexa 568-conjugated goat anti-rabbit IgG and Pacific blue-conjugated goat anti-mouse IgG, 1:200, Invitrogen) and Alexa Fluor 488 conjugated isolectin-B4 (1:200, Invitrogen) for 60 min at room temperature. After three washes with PBS, the sections were mounted with an Aqua-Mount Mounting Medium and then were examined with a laser confocal microscope (Leica TSC STED). No staining was seen when a negative control was performed with PBS instead of the primary antibody.

### Tissue Clearing

We recently developed a tissue-clearing protocol that combines passive clarity technique (PACT) and Clear Unobstructed Brain Imaging Cocktails and Computational Analysis (CUBIC) into a single method (CPC).^30^ This approach enables rapid tissue clearance while preserving endogenous fluorescence by limiting exposure to harsh detergents. The CPC protocol was validated by assessing TRPV1+ cell lineages in nervous system structures of varying sizes.^30^ To assess macrophage infiltration in thoracic DRGs after MI, we applied CPC for high-resolution three-dimensional (3D) mapping in CX3CR1CreER-tdTomato reporter mice (male, 4 weeks post-MI or sham; n = 2/each). Mice were anesthetized with urethane, transcardially perfused with PBS followed by 4% PFA, and T1–T4 DRGs were dissected, post-fixed, and processed with CPC to optimize imaging depth and fluorescence retention. Cleared tissues were mounted in CUBIC-R+ with spacers and imaged at the UNMC Multiphoton Intravital and Tissue Imaging Core using an Olympus FVMPE-RS multiphoton microscope with dual-line laser excitation for GFP and TdTomato. Volumetric image stacks were reconstructed and analyzed using Olympus FLUOVIEW software.

### Scanning electron microscope (SEM)

Rats were anesthetized and then perfused in 2% Glutaraldehyde, 2% PFA in 0.1M Sodium Cacodylate Buffer. T1-T4 DRGs were resected and then submersion fixed after being washed 3 times in Sodium Cacodylate buffer 15 minutes each wash and post fixed in 1% Osmium Tetroxide for 1 hour. Then all samples were washed 3 times in Sodium Cacodylate buffer, 15 minutes each wash, and then dehydrated through a graded Ethanol series (50%, 70%, 90%, 95%, 100% x 3 changes), 15 minutes each step. Subsequently, samples were then soaked in 100% Hexamethyldisilazane (HMDS) for three changes, 15 minutes for each change, and were left overnight in a fume hood in the third change of HMDS to allow the HMDS to evaporate, leaving the samples dry. Finally, samples were mounted on SEM stubs with double-sided adhesive carbon tabs and silver paste, then sputter-coated with approximately 50nm of gold/palladium alloy. Samples were observed on a FEI Quanta 200 SEM operated at 25 kV using the EM Core Facility at the University of Nebraska Medical Center (UNMC).

### Quantification of MCP-1 production by enzyme linked immunosorbent assay (ELISA)

Tissue protein for western blot was quick-thawed for analysis by ELISA, and the level of MCP-1 protein was detected using the Mouse CCL2/JE/MCP-1 Quantikine ELISA Kit (MJE00, R&D Systems, Minneapolis, MN) according to the manufacturer’s instructions. The lowest level of detection for MCP-1 was 15.6 pg/ml.

### Biotinylated TNFα injection and staining

Biotin-labeled TNFα or IL-6 (1.25 ng/µL, 1 µL for rats, 300 nL for mice) was injected into the subepicardium of the left ventricle (base, middle, and apex, at three injection sites). Saline injection served as the control. Buprenorphine SR (1.0 mg/kg) was subcutaneously injected at the beginning of surgery for pain management. Animals were anesthetized with pentobarbital sodium (40 mg/kg, intraperitoneally) and perfused with 4% PFA 1 day, 3 days, and 7 days after biotinylated TNFα or IL6 injection, respectively. Bilateral T1-T4 DRGs and stellate ganglia were post-fixed, dehydrated, and sectioned for biotin staining, respectively. Bilateral L4/L6 DRGs, which primarily innervate the hindlimb, were collected to serve as another type of control for T1- T4 DRGs, respectively. Then DRGs and stellate ganglia were removed and continuously fixed in 4% PFA for 24 hours and then were transferred into 30% sucrose. The experimenter was blind to all samples. For triple immunostaining of biotin, the DRG sections were stained with the isolectin IB4 (a non-peptidergic C-fiber neuronal marker), CGRP (a peptidergic C-fiber neuronal marker, Abcam, ab47027), or GFAP (a satellite glial cell marker, Abcam, ab7260) and NF200 (an A-fiber neuronal marker, Sigma-Aldrich, N5389). After being pre-incubated in 10% donkey serum for 60 min, the ganglia were incubated with mouse anti-NF200 antibody (1:200, Sigma-Aldrich) or rabbit anti-CGRP antibody (1:200, Abcam, ab47027) overnight at 4 °C. After PBS rinses, the sections were treated with fluorescence-conjugated secondary antibodies including Alexa Fluor 594 IgG Fraction Monoclonal Mouse Anti-Biotin (1:200, Jackson ImmunoResearch, West Grove, PA, USA), Pacific blue-conjugated goat anti-mouse IgG (1:200, Invitrogen, Carlsbad, CA, USA) for NF200 staining or Pacific blue-conjugated goat anti-rabbit IgG (1:200, Invitrogen, Carlsbad, CA, USA) for CGRP staining and Alexa Fluor 488 conjugated isolectin-B4 (1:200, Invitrogen) for 60 min at room temperature. After three washes with PBS, the sections of DRGs were mounted in standard mounting medium.

For double staining of biotin with satellite glial cells, the ganglia were incubated overnight at 4 °C with rabbit anti-GFAP (1:200, Abcam, ab7260) after pre-incubation in 10% donkey serum for 60 min. After PBS rinses, the sections were treated with fluorescence-conjugated secondary antibodies, including Alexa Fluor 594 IgG Fraction Monoclonal Mouse Anti-Biotin (1:200, Jackson ImmunoResearch, West Grove, PA, USA) and Alexa Fluor 488-conjugated goat anti- rabbit IgG (1:200, Invitrogen, Carlsbad, CA, USA) for 60 min at room temperature.

For double staining of MCP-1 with satellite glial cells, the ganglia were incubated with rabbit anti-MCP1 (1:200, Abcam, ab25124) and mouse anti-GFAP (1:200, Abcam, ab4648) overnight at 4 °C after being pre-incubated in 10% donkey serum for 60 min. After PBS rinses, the sections were treated with fluorescence-conjugated secondary antibodies (Alexa 568- conjugated goat anti-rabbit IgG and Alexa FluorR 488-conjugated goat anti-mouse IgG) (1:200, Invitrogen, Carlsbad, CA, USA) for 60 min at room temperature. After three washes with PBS, the sections of DRGs were mounted in standard mounting medium.

For double staining of biotin with stellate sympathetic neurons, the ganglia were incubated with rabbit anti-tyrosine hydroxylase (TH, 1:200, Abcam, ab137869) overnight at 4 °C after being pre-incubated in 10% donkey serum for 60 min. After PBS rinses, the sections were treated with fluorescence-conjugated secondary antibodies, including Alexa Fluor 594 IgG Fraction Monoclonal Mouse Anti-Biotin (1:200, Jackson ImmunoResearch, West Grove, PA, USA) and Alexa Fluor 488-conjugated goat anti-rabbit IgG (1:200, Invitrogen, Carlsbad, CA, USA) for 60 min at room temperature.

All slides were observed under a confocal laser scanning microscopy (Leica TSC STED).

No staining was seen when a negative control was performed.

### Thoracic DRG RNA Sequencing and Analysis

Thoracic DRGs were collected 8 weeks after MI or sham surgery (MI, n = 5; sham, n = 4). mRNA libraries were prepared on an Illumina NextSeq, generating 75bp reads. Demultiplexing used bcl2fastq v2.16.0.10. Adapters and low-quality bases were trimmed with cutadapt.^97^ Reads were aligned to the *Rattus norvegicus* rn6 reference with STAR v2.3.0.^98^ Gene-level counts were summarized against Partek v6.6 annotations (Partek Inc., St. Louis, MO, USA). Library sizes were normalized with the trimmed mean of M-values (TMM) method.^99^ Differential expression for MI versus SHAM was modeled in edgeR using a generalized linear model with likelihood-ratio tests and Benjamini-Hochberg FDR control.^100–102^ Differentially expressed genes were defined by FDR ≤ 0.15 and |log2FC| ≥ 0.5. Gene Ontology and KEGG enrichment were performed in ShinyGO v0.76.1 (*Rattus norvegicus*) using hypergeometric tests with FDR ≤ 0.05.^103^

### Data Acquisition and Statistical Analysis

MAP, HR, and RSNA were recorded using PowerLab software. Baseline values were calculated from at least 30 seconds of data prior to muscle contraction. Peak responses were defined as the greatest change from baseline. Data are presented as mean ± SEM. Comparisons between disease and treatment groups were performed using two-way ANOVA with Tukey’s multiple comparisons test (GraphPad Prism, version 10.5.0, San Diego, CA, USA). Western blot data were analyzed using Student’s *t*-test when data were normally distributed or the Mann– Whitney U test when they were not. Repeated-measures one-way ANOVA was used to evaluate treatment effects across multiple group measurements. A *P* value < 0.05 was considered statistically significant.

