## Supplemental Figures for "Neural Inflammation in Thoracic Dorsal Root Ganglia Mediates Cardiopulmonary Spinal Afferent Sensitization in Chronic Heart Failure"

### Both authors contribute equally to this work

**Running title: Neural Inflammation and Cardiopulmonary Afferent Sensitization**

*Correspondence to: Hanjun Wang, Department of Anesthesiology, University of Nebraska Medical Center, Omaha, NE, 68198, USA

**Disclosure:** Drs. Hanjun Wang, Dong Wang, Irving H. Zucker, Steven J. Lisco, and Juan Hong are holding a US patent related to the utility of ProGel-Dex in treating chronic heart failure (US Patent issued No. 12,138,274 B2).

**Figure S1**

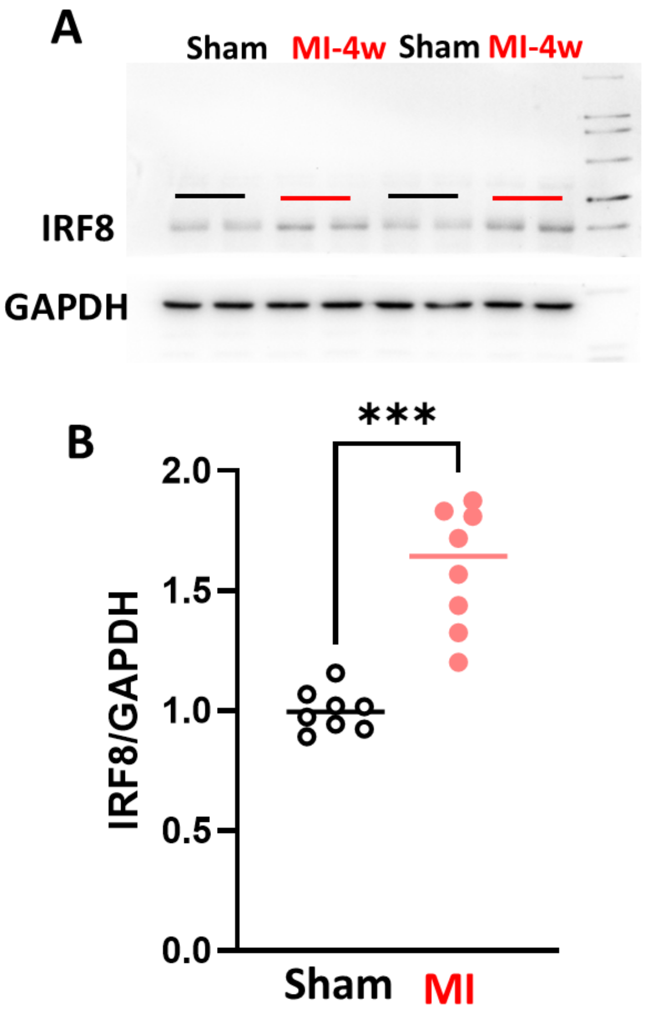


**Figure S1.** Western blot analysis of IRF8 protein in the T1-T4 DRG of female sham and MI rats at 4 weeks post sham and MI. Values are means ± SD. N=8 each group. *** $P$<0.001 vs. Sham.

**Figure S2**

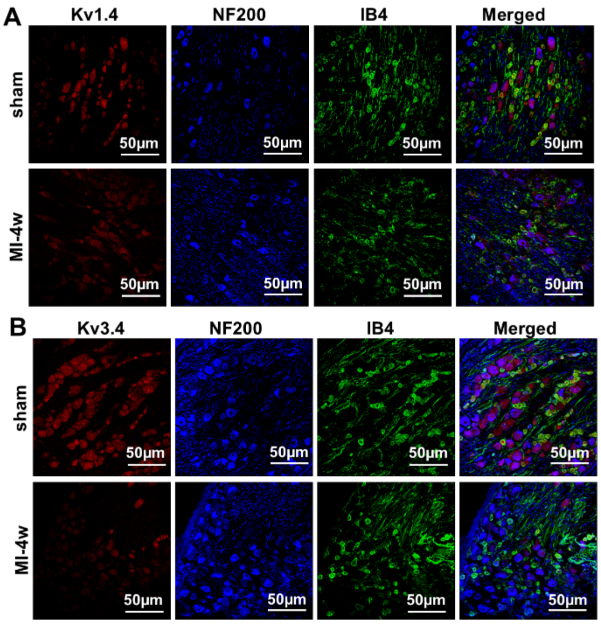


**Figure S2.** Immunofluorescence straining of Kv channels including Kv1.4, Kv3.4, Kv4.2 and Kv4.3 in T1-T4 DRG between sham and MI rats (**A-D**). Kv channels (red color); Isolectin B4 (IB4, green color), a C-fiber neuron marker; NF200 (blue color), an A-fiber neuronal marker. Scale bar = 50 μm.

### Figure S3

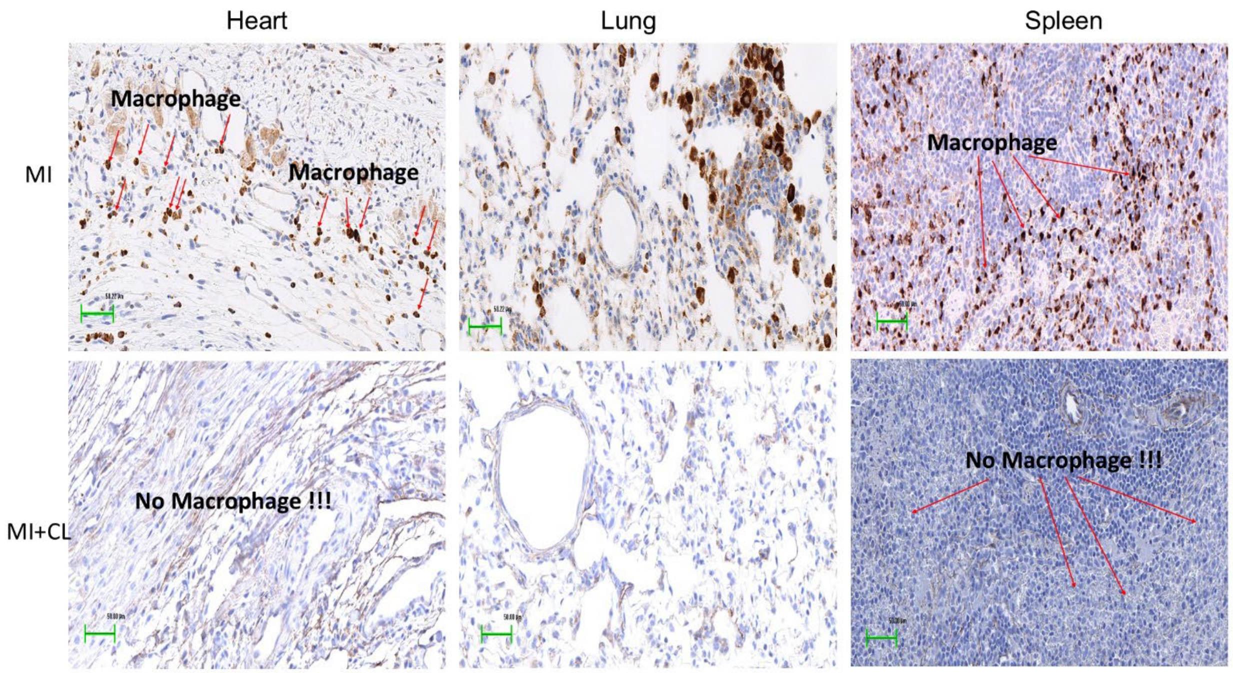


**Figure S3.** Immunohistological straining of CD68-positive cells in the heart, lungs and spleen in MI rats treated with and without clodronate liposomes (CL).

**Figure S4**

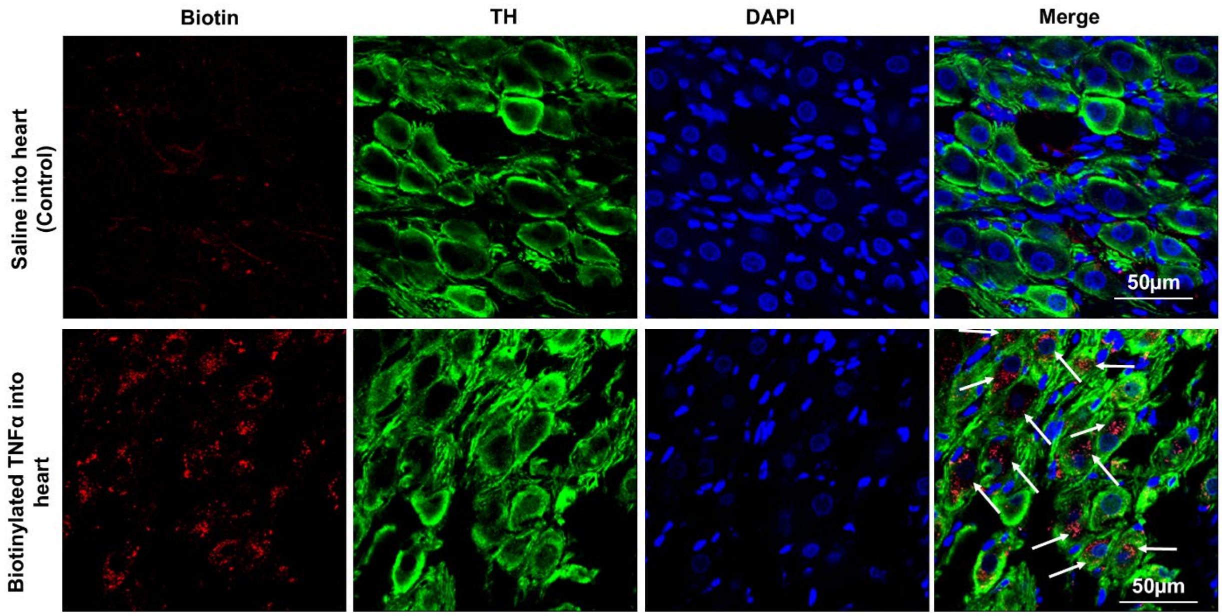


**Figure S4.** Immunofluorescence staining demonstrating that exogenous biotinylated TNFα in the sub epicardium of the left ventricle in normal rats can be taken up by Tyrosine hydroxylase (TH)-positive cardiac sympathetic efferents and transported back to the stellate ganglia soma.

**Figure S5**

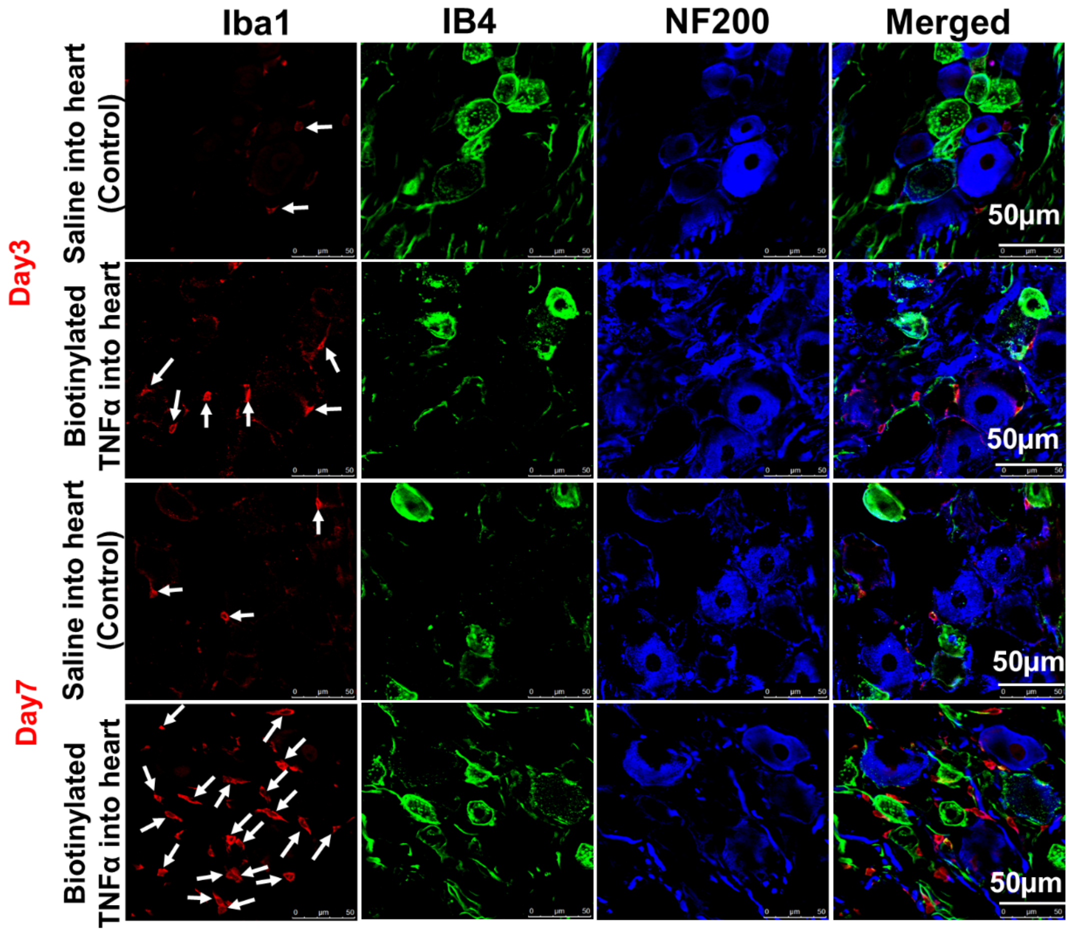


**Figure S5.** Immunofluorescence staining data demonstrate that injection of exogenous biotinylated TNFα into sub epicardium of the left ventricle caused gradually increased Iba1-positive cells in T1-T4 DRGs from day 3 to day 7 after injection in normal rats. IB4, a C-fiber neuron marker; NF200, an A-fiber neuron marker. White bar =50 μm.

**Table S1.** The percent of Kv1.4, Kv3.4, Kv4.2 and Kv4.3-positive DRG neurons to IB4- and NF200-positive neurons in T1-T4 DRGs in sham and MI rats

**Table 1.** The percent of potassium channels (Kv1.4, Kv3.4, Kv4.2 and Kv4.3)-positive DRG neurons to IB4-and NF200-neurons in intact ganglia in sham and CHF rats

| | sham | | CHF | |
|---|---|---|---|---|
| | Number | % | Number | % |
| Total IB4-positive | 2534±251 | | 2450±461 | |
| Kv1.4 with IB4 | 1806±346 | 70.2±8.1 | 562±137 | 23.8±5.5** |
| Total NF200-positive | 2520±331 | | 2379±442 | |
| Kv1.4 with NF200 | 2184±222 | 88.2±5.1 | 897±207 | 36.8±4.6** |
| Total IB4-positive | 2622±420 | | 2600±348 | |
| Kv3.4 with IB4 | 2152±314 | 82.1±4.3 | 910±217 | 35.0±5.2** |
| Total NF200-positive | 2428±479 | | 2405±384 | |
| Kv3.4 with NF200 | 1858±375 | 76.5±5.5 | 1332±425 | 55.4±8.3* |
| Total IB4-positive | 2691±372 | | 2604±474 | |
| Kv4.2 with IB4 | 2076±240 | 77.1±5.1 | 984±207 | 37.7±4.9** |
| Total NF200-positive | 2576±192 | | 2430±528 | |
| Kv4.2 with NF200 | 2240±141 | 86.9±3.3 | 1176±304 | 48.3±6.0** |
| Total IB4-positive | 2556±163 | | 2488±231 | |
| Kv4.3 with IB4 | 2214±144 | 86.6±5.7 | 680±168 | 27.3±5.8** |
| Total NF200-positive | 2362±453 | | 2325±295 | |
| Kv4.3 with NF200 | 1990±364 | 84.2±6.2 | 934±152 | 40.2±7.7** |

Values are Mean±S.E.M. n=5 per group. *P<0.05 and **P<0.01 vs. sham.

**Supplemental Video**

**Sham T1-T4 DRG**

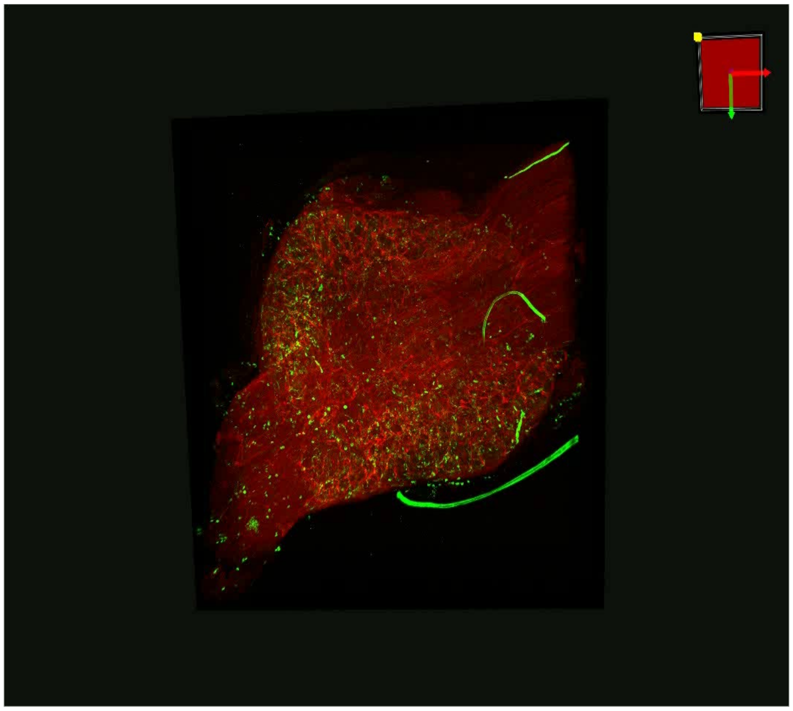

**MI T1-T4 DRGs**

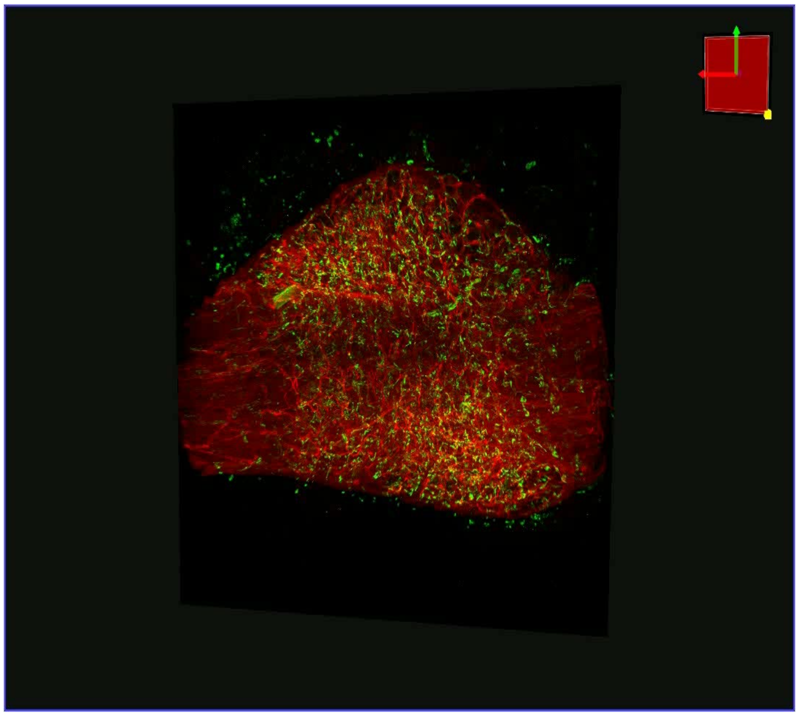

Table S2. Thoracic DRG MI vs Sham mRNA-Seq. All expressed genes.

| Gene | FC | log2FC | PValue | FDR |
|---|---|---|---|---|
| Cst3 | 1.69874052 | 0.7644655 | 7.43E-28 | 4.39E-24 |
| Necab1 | 1.45358639 | 0.539616817 | 3.03E-11 | 5.97E-08 |
| Myh11 | 1.57150815 | 0.652149758 | 2.45E-14 | 9.66E-11 |
| Gpx2 | 1.51593488 | 0.600207785 | 6.19E-12 | 1.83E-08 |
| Thbs4 | 0.67676144 | -0.563280727 | 2.31E-10 | 3.40E-07 |
| Myl9 | 1.45277314 | 0.538809437 | 2.21E-07 | 1.38E-04 |
| Igfbp3 | 1.52763667 | 0.61130146 | 7.04E-09 | 7.56E-06 |
| Vgf | 0.67322972 | -0.570829218 | 4.67E-07 | 2.62E-04 |
| Slc6a20 | 1.42959924 | 0.515610772 | 9.77E-06 | 0.0033923 |
| Acta2 | 1.7542968 | 0.810892852 | 1.36E-11 | 3.21E-08 |
| Slfn4 | 1.5848652 | 0.664360141 | 2.60E-08 | 2.36E-05 |
| Myom2 | 1.39867978 | 0.484065698 | 4.73E-05 | 0.0136129 |
| Anpep | 1.44187988 | 0.527950986 | 1.28E-05 | 0.0039761 |
| Elk1 | 0.57985511 | -0.786235644 | 1.29E-10 | 2.18E-07 |
| C4b | 1.55707195 | 0.638835612 | 2.90E-06 | 0.0012243 |
| Rtn4ip1 | 1.42078908 | 0.506692397 | 2.47E-04 | 0.0495239 |
| Pcdha7 | 1.45465914 | 0.540681138 | 1.48E-04 | 0.0332059 |
| Cbln4 | 1.72729259 | 0.788512481 | 1.48E-07 | 1.09E-04 |
| Acta1 | 0.6864839 | -0.542702212 | 2.65E-04 | 0.0504073 |
| Irf7 | 1.5951439 | 0.673686577 | 1.03E-05 | 0.0034881 |
| Fam111a | 0.14177968 | -2.818277272 | 2.12E-51 | 2.50E-47 |
| Cfp | 0.64197603 | -0.639408661 | 1.22E-04 | 0.0293905 |
| Pla2g2a | 1.95536552 | 0.967438315 | 1.29E-07 | 1.09E-04 |
| Pcdhga3 | 0.62875694 | -0.669425673 | 1.69E-04 | 0.036983 |
| Lgals5 | 1.61534417 | 0.691841585 | 1.87E-04 | 0.0401627 |
| Aox3 | 1.607574 | 0.684885152 | 4.79E-04 | 0.0816015 |
| Smoc1 | 1.57514624 | 0.655485775 | 9.16E-04 | 0.1424064 |
| RGD1309362 | 1.84562647 | 0.884110604 | 1.11E-05 | 0.0036253 |
| Prg2 | 1.9226729 | 0.943113342 | 1.17E-05 | 0.0037483 |
| Cnn1 | 2.42399331 | 1.277385718 | 1.46E-07 | 1.09E-04 |
| Actg2 | 2.42589906 | 1.278519524 | 1.71E-07 | 1.12E-04 |
| RT1-CE11 | 1.83861905 | 0.878622597 | 3.42E-04 | 0.0621292 |
| Epx | 1.9679175 | 0.976669743 | 4.24E-04 | 0.0758517 |
| RT1-T24-3 | 2.09250891 | 1.065233767 | 1.49E-04 | 0.0332059 |
| Myh4 | 0.4584093 | -1.125291797 | 1.25E-04 | 0.0295485 |
| Ifit1 | 3.56932506 | 1.835651295 | 4.91E-07 | 2.64E-04 |
| Fndc7 | 0.32380424 | -1.626806232 | 5.16E-06 | 0.0020328 |
| Col11a1 | 0.71853918 | -0.476861273 | 1.06E-04 | 0.0271388 |

| | | | | |
|---|---|---|---|---|
| Gbp2 | 1.36978714 | 0.45395172 | 8.11E-06 | 0.0030712 |
| Hspa1b | 0.73080028 | -0.452450916 | 4.44E-07 | 2.62E-04 |
| Ass1 | 1.35817497 | 0.441669352 | 9.75E-06 | 0.0033923 |
| Hba2 | 0.74022178 | -0.433970513 | 1.15E-09 | 1.51E-06 |
| C4a | 1.34310512 | 0.425572222 | 5.38E-07 | 2.76E-04 |
| Sfrp2 | 1.33639946 | 0.418351304 | 4.84E-04 | 0.0816015 |
| Trpa1 | 0.75747432 | -0.400731117 | 1.67E-07 | 1.12E-04 |
| Aldh1a2 | 1.31289276 | 0.392749077 | 3.12E-04 | 0.0576039 |
| Lypd1 | 1.30874472 | 0.388183722 | 5.91E-04 | 0.0968581 |
| Ctgf | 0.76647442 | -0.383690456 | 8.04E-07 | 3.80E-04 |
| Herpud1 | 0.7735873 | -0.37036399 | 4.18E-06 | 0.0017028 |
| Mylk | 1.28806181 | 0.365201826 | 2.36E-04 | 0.0480532 |
| Mx2 | 1.26772699 | 0.342244093 | 2.35E-04 | 0.0480532 |
| Vcl | 0.78899961 | -0.341903506 | 5.63E-07 | 2.77E-04 |
| Gpm6a | 0.79231854 | -0.335847533 | 6.55E-05 | 0.0179966 |
| Ppp1r10 | 0.79482784 | -0.331285688 | 4.35E-04 | 0.0766883 |
| Tagln | 1.23908272 | 0.309272507 | 2.54E-04 | 0.0499833 |
| Dpysl4 | 0.81053845 | -0.303047471 | 3.81E-05 | 0.0112357 |
| Col1a1 | 0.81345256 | -0.297869879 | 8.32E-06 | 0.0030712 |
| Mfap4 | 0.81710776 | -0.291401739 | 1.22E-04 | 0.0293905 |
| Ddit4 | 0.82514829 | -0.277274678 | 9.76E-05 | 0.0256189 |
| Itm2a | 0.82703744 | -0.273975448 | 2.63E-04 | 0.0504073 |
| Basp1 | 0.82783533 | -0.27258428 | 6.47E-04 | 0.1046 |
| Col1a2 | 0.82880391 | -0.27089728 | 6.87E-05 | 0.0184507 |
| Igfbp6 | 1.19174774 | 0.253078885 | 1.45E-04 | 0.0332059 |
| Phgdh | 0.83911262 | -0.253063643 | 5.11E-04 | 0.0850186 |
| Scd | 0.84423457 | -0.24428418 | 2.10E-04 | 0.0443635 |
| Serpinh1 | 0.84672026 | -0.240042693 | 2.73E-04 | 0.0512234 |
| Clk1 | 0.85156426 | -0.231812697 | 8.45E-04 | 0.1348625 |
| Col15a1 | 0.85401401 | -0.22766836 | 9.03E-04 | 0.1422239 |
| Coch | 2.57927595 | 1.366966129 | 4.72E-04 | 0.0816015 |
| RT1-N2 | 3.97586685 | 1.991269442 | 2.42E-06 | 0.0010565 |
| Car4 | 0.32402478 | -1.625823955 | 6.52E-05 | 0.0179966 |
| Ttr | 4.15713445 | 2.055589408 | 1.55E-05 | 0.0047067 |
| Alb | 0.15618921 | -2.678633334 | 8.46E-09 | 8.33E-06 |
| Ear11 | 3.52355004 | 1.817029702 | 1.20E-04 | 0.0293905 |
| Rnase1l2 | 0.19776215 | -2.338161783 | 1.60E-06 | 7.27E-04 |
| Myh6 | 0.04492928 | -4.476200147 | 5.91E-09 | 6.98E-06 |
| Per1 | 0.8167531 | -0.292028072 | 0.00109793 | 0.1683831 |
| Zdhhc21 | 0.81899391 | -0.288075376 | 0.00116447 | 0.1762983 |

| | | | | |
|---|---|---|---|---|
| Lss | 0.8396468 | -0.252145507 | 0.00132294 | 0.1977551 |
| Lect1 | 0.85170327 | -0.231577214 | 0.00142595 | 0.2082519 |
| Osmr | 1.35236463 | 0.435484188 | 0.00142844 | 0.2082519 |
| C1qtnf1 | 0.77031053 | -0.376487945 | 0.00146154 | 0.2093046 |
| Cartpt | 0.41957286 | -1.25300672 | 0.0014757 | 0.2093046 |
| Pcp4l1 | 0.78827452 | -0.343229948 | 0.00148883 | 0.2093046 |
| Anxa6 | 1.16321798 | 0.21812147 | 0.00151285 | 0.2101795 |
| Plau | 1.53556437 | 0.618768987 | 0.00153541 | 0.210833 |
| Kcnh6 | 0.77868715 | -0.360884279 | 0.00163404 | 0.2217973 |
| Nfkbia | 0.82272074 | -0.281525282 | 0.00166265 | 0.2231165 |
| Kcnj8 | 2.11256476 | 1.07899557 | 0.00170151 | 0.2257653 |
| Art2b | 0.40026627 | -1.320968032 | 0.00174452 | 0.2289 |
| Sphk1 | 0.76974257 | -0.377552065 | 0.00176591 | 0.2289798 |
| Pcdhgb8 | 1.46162653 | 0.547574721 | 0.00178391 | 0.2289798 |
| Sv2a | 0.81386687 | -0.297135267 | 0.00181874 | 0.2309405 |
| Oas2 | 2.50135267 | 1.322708477 | 0.0018601 | 0.2336806 |
| Chad | 0.5523622 | -0.856313511 | 0.00195322 | 0.2427954 |
| Mmp13 | 0.51479704 | -0.95792433 | 0.00208531 | 0.2565144 |
| Zcchc9 | 0.81409335 | -0.296733856 | 0.00214968 | 0.2617064 |
| Usp18 | 1.71982673 | 0.782263226 | 0.00219027 | 0.2621344 |
| Fos | 0.76794025 | -0.380934034 | 0.00219759 | 0.2621344 |
| Cacna2d2 | 0.79807704 | -0.325400068 | 0.00228933 | 0.2703464 |
| Chrne | 0.74538768 | -0.423937122 | 0.00234556 | 0.2730769 |
| Lyz2 | 1.14951564 | 0.201026099 | 0.00239606 | 0.2730769 |
| Cyr61 | 1.24312094 | 0.313966661 | 0.00239655 | 0.2730769 |
| Steap4 | 1.22186781 | 0.289088207 | 0.00240494 | 0.2730769 |
| Npy1r | 0.8309899 | -0.267097157 | 0.00257043 | 0.2871258 |
| Fabp4 | 1.74633794 | 0.804332771 | 0.0025773 | 0.2871258 |
| C2 | 1.24514114 | 0.316309287 | 0.00265404 | 0.2929119 |
| Fasn | 0.87063559 | -0.199859103 | 0.00268835 | 0.2939508 |
| Il2rg | 0.67379529 | -0.569617746 | 0.00275959 | 0.2970162 |
| Nqo1 | 1.19987203 | 0.262880552 | 0.00276668 | 0.2970162 |
| Sostdc1 | 0.83405496 | -0.261785648 | 0.00284243 | 0.3011822 |
| RT1-A1 | 1.27047038 | 0.345362741 | 0.0028565 | 0.3011822 |
| Cd163 | 1.28303602 | 0.359561675 | 0.00290886 | 0.3039883 |
| Col3a1 | 0.86682717 | -0.206183727 | 0.00301281 | 0.3120901 |
| Xdh | 1.23186604 | 0.300845377 | 0.00308556 | 0.3161321 |
| Mri1 | 1.31658647 | 0.396802276 | 0.00311828 | 0.3161321 |
| Mbp | 0.86588188 | -0.207757869 | 0.00317067 | 0.3161321 |
| C3 | 1.39935969 | 0.484766842 | 0.00317654 | 0.3161321 |

| | | | | |
|---|---|---|---|---|
| Krt12 | 0.17782466 | -2.491472727 | 0.00318568 | 0.3161321 |
| Scn11a | 0.87185769 | -0.197835425 | 0.00326749 | 0.3215482 |
| Tpbg | 0.57244311 | -0.804795778 | 0.00330162 | 0.3222218 |
| Ccr5 | 1.50383792 | 0.58864909 | 0.00345202 | 0.3341386 |
| Tmem87b | 1.2225127 | 0.289849457 | 0.00349783 | 0.3358205 |
| Crabp1 | 0.60955947 | -0.714161103 | 0.00354042 | 0.3371682 |
| Rn45s | 1.54097248 | 0.623841096 | 0.0036257 | 0.3425274 |
| Fkbp1a | 0.87419178 | -0.193978281 | 0.003659 | 0.3429294 |
| Mrgprx1 | 1.59823717 | 0.676481517 | 0.0038173 | 0.3549486 |
| Pygb | 1.14880891 | 0.200138844 | 0.00389341 | 0.3564187 |
| Top2a | 0.64819287 | -0.625504938 | 0.00389347 | 0.3564187 |
| Mpp2 | 0.70637326 | -0.501497359 | 0.00396739 | 0.360392 |
| Larp6 | 1.28475694 | 0.361495443 | 0.00400572 | 0.3610954 |
| Fkbp5 | 0.84695603 | -0.23964102 | 0.0040382 | 0.361266 |
| Txnip | 1.14101711 | 0.190320425 | 0.00407634 | 0.3619364 |
| Pi15 | 0.53797084 | -0.894400125 | 0.00424938 | 0.3742142 |
| Map3k11 | 0.81741972 | -0.290851046 | 0.004278 | 0.3742142 |
| P2rx4 | 1.2332484 | 0.302463414 | 0.00437146 | 0.379578 |
| Lrrc4b | 0.82331877 | -0.280476979 | 0.00443783 | 0.3825277 |
| Cd36 | 1.271248 | 0.346245505 | 0.00449859 | 0.3849551 |
| Per2 | 0.82832849 | -0.271725093 | 0.00454259 | 0.3859244 |
| C1qb | 1.222556 | 0.289900552 | 0.00479541 | 0.4044931 |
| Slc4a1 | 1.30603181 | 0.385190032 | 0.00485914 | 0.4045826 |
| Pmp2 | 1.14135213 | 0.190743958 | 0.00489223 | 0.4045826 |
| Wee1 | 0.83877609 | -0.253642357 | 0.00489926 | 0.4045826 |
| Adamts1 | 1.19918551 | 0.262054853 | 0.00494296 | 0.4053574 |
| Tenm3 | 0.85409935 | -0.227524206 | 0.00498537 | 0.4060151 |
| Slc16a11 | 1.17303671 | 0.230248167 | 0.00506673 | 0.4070793 |
| Zer1 | 0.84801568 | -0.237837162 | 0.0050777 | 0.4070793 |
| Slc3a1 | 0.42923878 | -1.220147655 | 0.00512818 | 0.4070793 |
| Capn5 | 0.87827462 | -0.187255984 | 0.00513632 | 0.4070793 |
| Rspo1 | 1.91168498 | 0.934844803 | 0.00529174 | 0.4161276 |
| Cxcl13 | 1.48996888 | 0.575282199 | 0.00532096 | 0.4161276 |
| Tpm2 | 1.22881388 | 0.297266415 | 0.00538301 | 0.4182101 |
| Net1 | 1.18169595 | 0.240858881 | 0.00561093 | 0.4330683 |
| Prim1 | 0.81735688 | -0.290961958 | 0.00565149 | 0.4333665 |
| Inmt | 1.4056888 | 0.491277235 | 0.00571283 | 0.434137 |
| Slc13a4 | 1.27045206 | 0.345341934 | 0.00575652 | 0.434137 |
| Abhd4 | 0.8788821 | -0.186258454 | 0.00581926 | 0.434137 |
| Spp1 | 1.14206816 | 0.191648755 | 0.005837 | 0.434137 |

| | | | | |
|---|---|---|---|---|
| Tnfrsf25 | 0.61362785 | -0.704564137 | 0.00584535 | 0.434137 |
| Dnm1 | 0.88106224 | -0.182684156 | 0.00599589 | 0.4366669 |
| Celsr2 | 0.84319696 | -0.246058434 | 0.00600091 | 0.4366669 |
| Ifi27l2b | 1.14453118 | 0.194756772 | 0.00602148 | 0.4366669 |
| Dmbt1 | 1.33103623 | 0.412549841 | 0.00602733 | 0.4366669 |
| Mcmdc2 | 1.57414991 | 0.654572942 | 0.00612369 | 0.4387971 |
| Nptxr | 0.86802808 | -0.204186382 | 0.00615223 | 0.4387971 |
| Fbxl4 | 1.22284263 | 0.290238753 | 0.0061682 | 0.4387971 |
| Oaf | 0.84813075 | -0.237641407 | 0.0062518 | 0.4420807 |
| Ptgds | 1.14079421 | 0.190038564 | 0.00631676 | 0.4440157 |
| Myl1 | 0.62125515 | -0.686742178 | 0.00643865 | 0.4499054 |
| Cybb | 1.34621144 | 0.428905021 | 0.00663354 | 0.460797 |
| Mmd2 | 0.8707522 | -0.199665889 | 0.00677964 | 0.4674423 |
| Gstm2 | 1.20936142 | 0.274245466 | 0.00680837 | 0.4674423 |
| Ramp3 | 0.78110903 | -0.356404149 | 0.00696944 | 0.4742751 |
| Mrgprc | 0.6493126 | -0.623014892 | 0.00698822 | 0.4742751 |
| Serping1 | 1.13538471 | 0.183181218 | 0.00703116 | 0.474463 |
| Plcb3 | 0.88065077 | -0.183358075 | 0.00732742 | 0.4916449 |
| C1r | 1.14278101 | 0.192548974 | 0.00737142 | 0.491803 |
| Ncam2 | 0.8121737 | -0.300139787 | 0.00753502 | 0.4946294 |
| Sfrp4 | 1.20879454 | 0.273569053 | 0.00755983 | 0.4946294 |
| Ddah1 | 0.86200358 | -0.214234232 | 0.00757769 | 0.4946294 |
| Scara5 | 1.15921015 | 0.213142128 | 0.00758133 | 0.4946294 |
| Pi16 | 1.14041887 | 0.18956382 | 0.00776276 | 0.5036835 |
| Ankrd9 | 0.75015368 | -0.414741915 | 0.00783128 | 0.5053529 |
| Nov | 1.21639515 | 0.282611965 | 0.00811747 | 0.5206653 |
| Otof | 0.75865187 | -0.39849009 | 0.00818535 | 0.5206653 |
| Frzb | 1.39990888 | 0.485332922 | 0.00820084 | 0.5206653 |
| Tril | 0.84455296 | -0.2437402 | 0.00841115 | 0.5294645 |
| Acsl5 | 1.17851496 | 0.236970076 | 0.00842911 | 0.5294645 |
| Cfd | 1.41477146 | 0.500569017 | 0.00849066 | 0.5305089 |
| RT1-DOb | 2.49132793 | 1.316914934 | 0.00861647 | 0.5317813 |
| Rab7b | 0.75514174 | -0.405180641 | 0.00862217 | 0.5317813 |
| Eif2ak2 | 1.18755458 | 0.247993818 | 0.00864612 | 0.5317813 |
| Cd247 | 0.48002326 | -1.058823788 | 0.00878015 | 0.5372268 |
| Mras | 0.85260375 | -0.230052694 | 0.00882769 | 0.5373517 |
| Ubash3b | 0.87780381 | -0.188029556 | 0.00893506 | 0.5410979 |
| Sdc3 | 0.88648293 | -0.173835237 | 0.00908115 | 0.5448184 |
| Mvd | 0.83339361 | -0.262930064 | 0.00908877 | 0.5448184 |
| Pcolce | 0.86315421 | -0.212309763 | 0.00917296 | 0.5453825 |

| | | | | |
|---|---|---|---|---|
| Col9a2 | 0.74809677 | -0.418703187 | 0.00919054 | 0.5453825 |
| Ednrb | 0.85634293 | -0.223739437 | 0.00928942 | 0.5476468 |
| Map3k7cl | 1.99371734 | 0.995460882 | 0.00932145 | 0.5476468 |
| Srd5a2 | 10.4835148 | 3.390050586 | 0.00945476 | 0.5527289 |
| Elp6 | 1.23521078 | 0.304757252 | 0.00959036 | 0.5571745 |
| Tp53i13 | 0.70112966 | -0.512246819 | 0.00962517 | 0.5571745 |
| Ghdc | 0.76248027 | -0.391228085 | 0.01005354 | 0.5791327 |
| Col4a1 | 0.88793117 | -0.171480254 | 0.01013926 | 0.5812355 |
| Aatk | 0.8894632 | -0.168993179 | 0.01038675 | 0.5908462 |
| Sult1a1 | 1.15852433 | 0.212288341 | 0.01040698 | 0.5908462 |
| Prlr | 2.07860145 | 1.055613162 | 0.0106687 | 0.5953581 |
| Dlgap3 | 0.84591294 | -0.241418902 | 0.01069395 | 0.5953581 |
| Gpx1 | 1.13883651 | 0.187560651 | 0.01070439 | 0.5953581 |
| Rin2 | 1.19051703 | 0.251588258 | 0.01070787 | 0.5953581 |
| Tec | 1.27481062 | 0.35028294 | 0.01082667 | 0.5953581 |
| Fam221a | 0.50835628 | -0.976088148 | 0.01082708 | 0.5953581 |
| Ankrd6 | 0.82059899 | -0.285250714 | 0.01085036 | 0.5953581 |
| Snx14 | 0.84895898 | -0.236233251 | 0.01093419 | 0.5953581 |
| Rit2 | 1.14630388 | 0.196989542 | 0.01094019 | 0.5953581 |
| LOC360231 | 1.57603444 | 0.656299058 | 0.01106913 | 0.5976639 |
| Slc27a4 | 0.85718862 | -0.222315395 | 0.01112299 | 0.5976639 |
| Sgk1 | 0.86072143 | -0.216381698 | 0.01117309 | 0.5976639 |
| Dnase1l3 | 2.02170702 | 1.015573941 | 0.01122969 | 0.5976639 |
| RT1-S2 | 1.69907536 | 0.764749839 | 0.01123562 | 0.5976639 |
| Siglec1 | 1.57031539 | 0.651054348 | 0.01133165 | 0.6000691 |
| Ptms | 0.89045984 | -0.167377554 | 0.01146255 | 0.6039076 |
| Cd53 | 1.2350187 | 0.304532891 | 0.01150641 | 0.6039076 |
| Ackr2 | 1.60198154 | 0.67985752 | 0.01170793 | 0.6117654 |
| Dagla | 0.87295235 | -0.196025182 | 0.0117607 | 0.6118156 |
| Npr3 | 0.66254359 | -0.59391273 | 0.01196399 | 0.6173367 |
| Rtp4 | 1.47507388 | 0.560787214 | 0.01199845 | 0.6173367 |
| Fbxo41 | 0.77306497 | -0.371338431 | 0.01210453 | 0.6173367 |
| Cd4 | 1.24304214 | 0.313875209 | 0.01212594 | 0.6173367 |
| Hmgcr | 0.89042978 | -0.167426246 | 0.0121465 | 0.6173367 |
| Ctnnd2 | 0.85327272 | -0.228921168 | 0.01221696 | 0.6173367 |
| Cnp | 0.89132498 | -0.165976551 | 0.01223277 | 0.6173367 |
| Arsb | 0.84468341 | -0.24351738 | 0.01239548 | 0.6212965 |
| Usp2 | 0.6911598 | -0.532908789 | 0.01241646 | 0.6212965 |
| Lrif1 | 1.20146786 | 0.264798053 | 0.01272368 | 0.6330313 |
| Orai1 | 1.29265386 | 0.37033601 | 0.01280817 | 0.6330313 |

| | | | | |
|---|---|---|---|---|
| Agps | 0.83325256 | -0.263174247 | 0.01288006 | 0.6330313 |
| Mpeg1 | 1.18960057 | 0.250477247 | 0.01290181 | 0.6330313 |
| Slitrk2 | 0.81493609 | -0.295241163 | 0.01291901 | 0.6330313 |
| Nid2 | 0.87076115 | -0.199651054 | 0.01297988 | 0.6333859 |
| Cx3cr1 | 1.69990328 | 0.765452663 | 0.01310929 | 0.6370685 |
| RGD1305938 | 0.87455644 | -0.193376596 | 0.01327099 | 0.6422833 |
| Madd | 0.88838287 | -0.170746518 | 0.0135817 | 0.6528714 |
| Slfn2 | 1.26325813 | 0.33714947 | 0.01367086 | 0.6528714 |
| Olfml3 | 0.84502887 | -0.242927459 | 0.0136984 | 0.6528714 |
| B3galt5 | 1.21898597 | 0.285681517 | 0.01372177 | 0.6528714 |
| Scn10a | 0.89158975 | -0.165548062 | 0.01383771 | 0.6528714 |
| Cd180 | 1.75029383 | 0.80759713 | 0.01386415 | 0.6528714 |
| Slc4a4 | 0.8566972 | -0.223142728 | 0.01387768 | 0.6528714 |
| Mboat2 | 0.87214057 | -0.197367412 | 0.01401061 | 0.6528714 |
| Car1 | 1.81132034 | 0.857041716 | 0.01401508 | 0.6528714 |
| Pclo | 0.89064514 | -0.167077366 | 0.01404262 | 0.6528714 |
| Dnajb1 | 0.87662215 | -0.189972957 | 0.01421491 | 0.6544204 |
| LOC498675 | 0.53972434 | -0.889705356 | 0.01422682 | 0.6544204 |
| Fut7 | 0.58206405 | -0.780750168 | 0.01424998 | 0.6544204 |
| Rdm1 | 1.47868674 | 0.564316454 | 0.01429761 | 0.6544204 |
| Tmem132a | 0.83945986 | -0.252466753 | 0.01448472 | 0.6563146 |
| Tlr3 | 1.28443028 | 0.36112858 | 0.01454855 | 0.6563146 |
| Fadd | 2.07019444 | 1.049766275 | 0.01455577 | 0.6563146 |
| F13a1 | 1.21581822 | 0.281927546 | 0.01460541 | 0.6563146 |
| Myof | 1.1835479 | 0.243118092 | 0.01461688 | 0.6563146 |
| Bst2 | 1.24059643 | 0.311033883 | 0.01529473 | 0.6806416 |
| Tanc1 | 0.88057707 | -0.183478814 | 0.01532469 | 0.6806416 |
| Ahsa2 | 0.82587788 | -0.27599962 | 0.01535031 | 0.6806416 |
| Tbx18 | 1.34553484 | 0.428179745 | 0.01538922 | 0.6806416 |
| Pecam1 | 1.23016861 | 0.298856067 | 0.01547132 | 0.6817194 |
| RT1-N3 | 1.49814928 | 0.583181386 | 0.01555188 | 0.6827219 |
| Ly6e | 1.1502402 | 0.201935162 | 0.01569516 | 0.6864597 |
| Osr1 | 1.34477773 | 0.427367742 | 0.01595791 | 0.6953762 |
| Tbkbp1 | 0.81162332 | -0.301117775 | 0.0160697 | 0.6969059 |
| Phf11b | 2.27258372 | 1.184333443 | 0.01614299 | 0.6969059 |
| Lnc215 | 1.17026145 | 0.226830879 | 0.01617006 | 0.6969059 |
| Kcnc4 | 0.86591 | -0.207711008 | 0.0163209 | 0.7008491 |
| Peak1 | 0.87443881 | -0.193570666 | 0.0165737 | 0.7066856 |
| Ebf2 | 1.1990942 | 0.261945005 | 0.0165765 | 0.7066856 |
| Adam19 | 0.89194842 | -0.164967808 | 0.01682216 | 0.7145785 |

| | | | | |
|---|---|---|---|---|
| Birc3 | 1.75517221 | 0.811612586 | 0.0169494 | 0.7150672 |
| Ptpdc1 | 0.85065889 | -0.233347361 | 0.01695476 | 0.7150672 |
| Cilp | 1.25202464 | 0.324262951 | 0.01713653 | 0.7201609 |
| Pex12 | 1.25191682 | 0.324138708 | 0.01731245 | 0.7249741 |
| C1s | 1.11835778 | 0.161381799 | 0.01739214 | 0.725046 |
| St6galnac5 | 0.64311884 | -0.636842751 | 0.01748474 | 0.725046 |
| Ntrk2 | 0.8970279 | -0.156775238 | 0.0175556 | 0.725046 |
| Ism1 | 1.65193367 | 0.724155757 | 0.01763364 | 0.725046 |
| Dapp1 | 1.52713009 | 0.610822961 | 0.0176865 | 0.725046 |
| Fxyd1 | 1.13754514 | 0.185923792 | 0.017744 | 0.725046 |
| Lrat | 1.78600363 | 0.836735016 | 0.01780145 | 0.725046 |
| Gpd1 | 0.85158077 | -0.231784717 | 0.01781771 | 0.725046 |
| Mrps35 | 1.17691639 | 0.235011837 | 0.01788161 | 0.725046 |
| Crot | 0.88783729 | -0.171632784 | 0.01792814 | 0.725046 |
| Sparcl1 | 1.11675193 | 0.159308752 | 0.01807683 | 0.7285641 |
| Pnpla1 | 1.39846836 | 0.48384761 | 0.01849413 | 0.7412579 |
| Tagap | 0.66291519 | -0.593103787 | 0.01852108 | 0.7412579 |
| Ncs1 | 0.86621969 | -0.207195131 | 0.0185866 | 0.7412579 |
| Vtn | 1.18108014 | 0.240106858 | 0.01868906 | 0.7412579 |
| Smpx | 1.27818151 | 0.354092728 | 0.01871705 | 0.7412579 |
| Alpl | 0.85424343 | -0.227280847 | 0.01877752 | 0.7412579 |
| Gstm7 | 1.1535901 | 0.206130684 | 0.01883118 | 0.7412579 |
| Cep104 | 0.87292509 | -0.196070234 | 0.01895927 | 0.7414275 |
| Lrfn4 | 0.81064667 | -0.302854866 | 0.01898398 | 0.7414275 |
| Gucy1a3 | 1.31187815 | 0.391633729 | 0.01902398 | 0.7414275 |
| Ankrd52 | 0.87856294 | -0.186782448 | 0.01908662 | 0.7414275 |
| Prex1 | 0.87206403 | -0.197494035 | 0.01918804 | 0.7429231 |
| Fbxl16 | 0.82835853 | -0.271672771 | 0.01942244 | 0.7495412 |
| Slc16a10 | 1.96484869 | 0.97441822 | 0.01960815 | 0.7520753 |
| Acad10 | 1.36021621 | 0.443835992 | 0.01961548 | 0.7520753 |
| Uba7 | 1.44109902 | 0.527169472 | 0.01988592 | 0.7599767 |
| Tf | 1.12496652 | 0.169882062 | 0.02005124 | 0.763823 |
| Spib | 0.65349915 | -0.613742736 | 0.02012485 | 0.7641619 |
| Nrp2 | 0.87627053 | -0.190551749 | 0.02039013 | 0.7717532 |
| Rgs8 | 0.69916648 | -0.516292076 | 0.02049349 | 0.7731872 |
| Gsk3a | 0.88272317 | -0.179967032 | 0.02065198 | 0.7766854 |
| Col11a2 | 0.75229033 | -0.410638542 | 0.02075149 | 0.7779503 |
| Cdo1 | 1.16894956 | 0.225212674 | 0.02088049 | 0.7803093 |
| Gsn | 1.11809272 | 0.161039833 | 0.02103274 | 0.7833953 |
| Mmp2 | 0.89535357 | -0.159470592 | 0.02109575 | 0.7833953 |

| | | | | |
|---|---|---|---|---|
| Kcna6 | 0.88660819 | -0.173631406 | 0.02153882 | 0.7973415 |
| Akr1b8 | 1.7114602 | 0.775227747 | 0.02168554 | 0.7989878 |
| Nox4 | 1.69711314 | 0.763082747 | 0.02171861 | 0.7989878 |
| Taf1a | 0.69932991 | -0.51595489 | 0.02187284 | 0.8008687 |
| Itgam | 1.40968223 | 0.495369983 | 0.02190538 | 0.8008687 |
| Phka2 | 0.65388249 | -0.612896711 | 0.02218621 | 0.802256 |
| Zbtb3 | 1.58458004 | 0.664100535 | 0.02231921 | 0.802256 |
| Sf1 | 0.88506633 | -0.176142516 | 0.0223432 | 0.802256 |
| St6galnac3 | 0.72547076 | -0.46301062 | 0.02246629 | 0.802256 |
| Samsn1 | 1.83155106 | 0.873065923 | 0.0225754 | 0.802256 |
| Map3k2 | 0.71638207 | -0.481198872 | 0.02262346 | 0.802256 |
| Fads2 | 0.88878064 | -0.170100704 | 0.0226433 | 0.802256 |
| Ces1d | 1.5017142 | 0.586610267 | 0.02265096 | 0.802256 |
| Arhgef7 | 0.87998866 | -0.184443166 | 0.02265156 | 0.802256 |
| Cacng5 | 0.69474808 | -0.525438163 | 0.02267607 | 0.802256 |
| Kctd7 | 0.76448641 | -0.38743724 | 0.02274137 | 0.802256 |
| Nrap | 0.42288961 | -1.241646995 | 0.022799 | 0.802256 |
| Rab36 | 0.75858907 | -0.398609503 | 0.0228628 | 0.802256 |
| Usp33 | 1.1116205 | 0.152664346 | 0.02289443 | 0.802256 |
| Cntn2 | 0.90097015 | -0.150448788 | 0.02340113 | 0.8083752 |
| Zfp746 | 0.79784957 | -0.325811332 | 0.02342121 | 0.8083752 |
| Prr7 | 0.64197328 | -0.639414852 | 0.02348631 | 0.8083752 |
| Sh3bp1 | 1.50733569 | 0.592000749 | 0.02354222 | 0.8083752 |
| Csrnp2 | 1.20454077 | 0.268483221 | 0.02357813 | 0.8083752 |
| Itga8 | 1.28847189 | 0.365661063 | 0.02361111 | 0.8083752 |
| Mib1 | 0.87026671 | -0.200470492 | 0.02362012 | 0.8083752 |
| Il17rc | 1.40775251 | 0.493393719 | 0.02365172 | 0.8083752 |
| Hmg1l1 | 1.18949961 | 0.250354797 | 0.02372577 | 0.8083752 |
| Map4k3 | 0.86953923 | -0.201676979 | 0.02375359 | 0.8083752 |
| RT1-CE10 | 1.17746426 | 0.235683269 | 0.02386504 | 0.8098341 |
| Dcdc2 | 1.41672165 | 0.502556338 | 0.02410394 | 0.815597 |
| Pank3 | 0.90189944 | -0.148961505 | 0.02436931 | 0.8222207 |
| Phlda3 | 0.88749621 | -0.172187139 | 0.02450441 | 0.8244233 |
| C6 | 1.45948755 | 0.545461901 | 0.02476968 | 0.8300439 |
| Exosc7 | 1.3738929 | 0.458269547 | 0.02481205 | 0.8300439 |
| Cfb | 1.2507994 | 0.322850429 | 0.02497037 | 0.8329806 |
| Cd3e | 0.56962227 | -0.81192255 | 0.02508127 | 0.8343233 |
| Hemgn | 1.41455863 | 0.500351972 | 0.02523834 | 0.8353303 |
| Srsf5 | 1.1115635 | 0.152590371 | 0.02525302 | 0.8353303 |
| Prg3 | 1.8426948 | 0.881817144 | 0.02541393 | 0.8383048 |

| | | | | |
|---|---|---|---|---|
| Mmgt2 | 1.36340416 | 0.447213291 | 0.02556851 | 0.8403299 |
| Ehd4 | 1.13549113 | 0.183316442 | 0.02561765 | 0.8403299 |
| Acss1 | 1.20818992 | 0.27284725 | 0.02585633 | 0.8437987 |
| Atp13a2 | 0.90050024 | -0.15120144 | 0.02592163 | 0.8437987 |
| Cpxm1 | 0.77787767 | -0.362384802 | 0.02593775 | 0.8437987 |
| Tmem184b | 0.89658492 | -0.157487858 | 0.02607917 | 0.8458398 |
| Disp2 | 0.90082609 | -0.150679487 | 0.02614375 | 0.8458398 |
| Wipf3 | 0.88328934 | -0.179041997 | 0.02647034 | 0.8537316 |
| Rarres2 | 0.87611879 | -0.190801599 | 0.02653226 | 0.8537316 |
| Pfkl | 0.87289231 | -0.196124424 | 0.02667877 | 0.856113 |
| Set | 0.88895716 | -0.169814197 | 0.02684568 | 0.8566906 |
| Elmo3 | 1.5245882 | 0.608419616 | 0.02691287 | 0.8566906 |
| Spn | 1.39751725 | 0.482866086 | 0.02691441 | 0.8566906 |
| Taf6l | 1.41477528 | 0.500572916 | 0.02699832 | 0.8567516 |
| Ly6g6e | 0.62246991 | -0.683923992 | 0.02715502 | 0.8567516 |
| Oprm1 | 0.82186455 | -0.28302745 | 0.02731905 | 0.8567516 |
| Ltbp3 | 0.89650781 | -0.157611943 | 0.02733098 | 0.8567516 |
| Arap2 | 1.13131306 | 0.177998216 | 0.02744572 | 0.8567516 |
| Ndst1 | 0.85591092 | -0.224467437 | 0.0274972 | 0.8567516 |
| Nacc1 | 0.86039377 | -0.216931016 | 0.02750402 | 0.8567516 |
| Hspa1a | 0.75484815 | -0.405741645 | 0.02756053 | 0.8567516 |
| Dhrs3 | 1.21366032 | 0.279364698 | 0.02769059 | 0.8567516 |
| Fam150b | 0.52379204 | -0.932933959 | 0.02769425 | 0.8567516 |
| Eng | 1.15166246 | 0.20371794 | 0.02782625 | 0.8567516 |
| Ugt8 | 0.9038681 | -0.14581584 | 0.02783988 | 0.8567516 |
| Cpxm2 | 0.90090436 | -0.15055413 | 0.02804589 | 0.8567516 |
| Cyp2s1 | 0.84284848 | -0.246654803 | 0.0281579 | 0.8567516 |
| Fggy | 0.77222249 | -0.372911527 | 0.02817134 | 0.8567516 |
| Galr2 | 0.7799375 | -0.358569585 | 0.028272 | 0.8567516 |
| Prss35 | 1.36986534 | 0.454034084 | 0.02831223 | 0.8567516 |
| Bin2 | 1.489777 | 0.575096392 | 0.02839896 | 0.8567516 |
| Fam89a | 0.83136322 | -0.266449164 | 0.02840972 | 0.8567516 |
| RGD1565059 | 1.38237576 | 0.467149827 | 0.02841491 | 0.8567516 |
| Rag1 | 0.57810843 | -0.790587989 | 0.02843989 | 0.8567516 |
| Ptprz1 | 0.89980856 | -0.152310002 | 0.02851891 | 0.8569459 |
| Fndc5 | 0.86115801 | -0.215650124 | 0.02875908 | 0.859499 |
| Vipr2 | 0.78224517 | -0.354307247 | 0.02888524 | 0.859499 |
| Mab21l1 | 0.75945295 | -0.396967508 | 0.02890566 | 0.859499 |
| Slc2a12 | 1.36281186 | 0.446586412 | 0.02892749 | 0.859499 |
| Tfpi2 | 1.85105607 | 0.888348597 | 0.02896779 | 0.859499 |

| | | | | |
|---|---|---|---|---|
| Nupr2 | 1.54887508 | 0.631220789 | 0.02931619 | 0.8676565 |
| Map7d3 | 1.23069734 | 0.299476013 | 0.02965907 | 0.87561 |
| Sema5a | 0.90321801 | -0.146853844 | 0.0299188 | 0.881075 |
| Tph1 | 0.74372757 | -0.427153834 | 0.03021924 | 0.8814563 |
| Slc9a3r1 | 0.88781066 | -0.17167606 | 0.0302228 | 0.8814563 |
| Mdfi | 0.53735025 | -0.896065349 | 0.03022688 | 0.8814563 |
| LOC24906 | 1.65135074 | 0.723646579 | 0.03036701 | 0.8814563 |
| Abca2 | 0.90588456 | -0.142600884 | 0.03039343 | 0.8814563 |
| Spta1 | 1.46140334 | 0.547354406 | 0.03042764 | 0.8814563 |
| Zfyve28 | 0.88657492 | -0.173685551 | 0.03049061 | 0.8814563 |
| Cd47 | 0.88337957 | -0.178894619 | 0.03052889 | 0.8814563 |
| Fgfr4 | 0.65269701 | -0.615514667 | 0.0306574 | 0.883008 |
| Adipoq | 1.65655205 | 0.728183532 | 0.03078639 | 0.8830421 |
| Pcdhgc3 | 0.88513309 | -0.176033691 | 0.03088243 | 0.8830421 |
| Nrgn | 1.77889479 | 0.830981183 | 0.03088292 | 0.8830421 |
| RGD1359290 | 1.14770507 | 0.198751956 | 0.03105314 | 0.8840758 |
| Jph4 | 0.86140778 | -0.215231736 | 0.0310688 | 0.8840758 |
| Drgx | 0.88779301 | -0.171704752 | 0.03116982 | 0.884525 |
| Fcgr2a | 1.48163459 | 0.567189689 | 0.03125975 | 0.884525 |
| Pou2af1 | 0.63820809 | -0.647901209 | 0.03130929 | 0.884525 |
| Pla2g7 | 1.11792836 | 0.160827743 | 0.03159524 | 0.8904731 |
| Igfbp2 | 0.86835162 | -0.203648742 | 0.03192236 | 0.8931451 |
| Ckm | 0.72235375 | -0.469222571 | 0.03201687 | 0.8931451 |
| Cd302 | 1.16550936 | 0.220960597 | 0.03203089 | 0.8931451 |
| Ppp2r5b | 0.90239353 | -0.148171371 | 0.03203644 | 0.8931451 |
| Atp13a5 | 1.60722573 | 0.684572561 | 0.032174 | 0.8931451 |
| Rufy1 | 0.84386505 | -0.244915785 | 0.03219795 | 0.8931451 |
| Kbtbd11 | 0.83206912 | -0.265224721 | 0.03221948 | 0.8931451 |
| Ciart | 0.76068903 | -0.394621293 | 0.0323843 | 0.8956116 |
| Mis18bp1 | 1.30281853 | 0.38163615 | 0.03252479 | 0.8966152 |
| Ece1 | 0.88528425 | -0.175787341 | 0.03257244 | 0.8966152 |
| Mecom | 1.36242955 | 0.446181631 | 0.03271913 | 0.8985587 |
| Ifit3 | 1.42012116 | 0.506014023 | 0.03283406 | 0.8988568 |
| Zc3h12d | 1.61980342 | 0.695818734 | 0.03299436 | 0.8988568 |
| Rad54l2 | 0.84281855 | -0.246706022 | 0.03308736 | 0.8988568 |
| Parvg | 1.4938581 | 0.579043117 | 0.03313585 | 0.8988568 |
| Sat1 | 1.11722586 | 0.159920867 | 0.03318283 | 0.8988568 |
| Zfp498 | 0.72372018 | -0.466496101 | 0.03318669 | 0.8988568 |
| Ccnb2 | 0.64308076 | -0.636928177 | 0.0333308 | 0.9004595 |
| Tlr2 | 1.38532213 | 0.470221485 | 0.03339836 | 0.9004595 |

| | | | | |
|---|---|---|---|---|
| Nckap5l | 0.85852914 | -0.220060985 | 0.03356753 | 0.902959 |
| Sell | 0.71537814 | -0.483222067 | 0.03370857 | 0.9046921 |
| Crk | 0.88018566 | -0.184120227 | 0.03396385 | 0.9094763 |
| Ifi47 | 1.30086904 | 0.379475735 | 0.03406528 | 0.9101287 |
| Zfp870 | 1.47045709 | 0.55626468 | 0.03421271 | 0.9120044 |
| Lamp5 | 1.18414307 | 0.243843401 | 0.03429226 | 0.9120659 |
| Cp | 1.12087875 | 0.164630227 | 0.03446221 | 0.9126916 |
| Postn | 0.82905286 | -0.270464001 | 0.03461553 | 0.9126916 |
| Ung | 1.49508732 | 0.58022975 | 0.03462231 | 0.9126916 |
| Kcnj13 | 1.31508157 | 0.395152288 | 0.03464361 | 0.9126916 |
| Gfra2 | 0.89203011 | -0.16483569 | 0.03470222 | 0.9126916 |
| Nup210 | 0.74393734 | -0.426746983 | 0.03493776 | 0.9164648 |
| Pde11a | 0.75133208 | -0.41247739 | 0.03506619 | 0.9164648 |
| Eif3k | 1.16310084 | 0.217976183 | 0.03507851 | 0.9164648 |
| Pnpla7 | 1.19755744 | 0.260094858 | 0.03519494 | 0.9174769 |
| Dmgdh | 0.76027333 | -0.395409913 | 0.03540843 | 0.921009 |
| Igsf3 | 0.90042762 | -0.151317786 | 0.03556089 | 0.9229419 |
| Wsb2 | 0.9046386 | -0.144586537 | 0.03593198 | 0.927265 |
| Prdm9 | 0.62895163 | -0.668979034 | 0.03594626 | 0.927265 |
| Tlr6 | 1.49372741 | 0.578916897 | 0.03596302 | 0.927265 |
| Clec14a | 1.23444206 | 0.303859127 | 0.03620668 | 0.9281395 |
| MGC116121 | 1.27932703 | 0.3553851 | 0.03626363 | 0.9281395 |
| Rps25 | 0.90474438 | -0.14441785 | 0.036289 | 0.9281395 |
| Cacna1b | 0.90317986 | -0.146914777 | 0.03631133 | 0.9281395 |
| Synpo2 | 1.27544102 | 0.350996189 | 0.03645168 | 0.9287141 |
| Coq3 | 1.17115313 | 0.227929719 | 0.0364911 | 0.9287141 |
| Pamr1 | 1.48184504 | 0.567394591 | 0.03682027 | 0.9350765 |
| Aaed1 | 1.50433461 | 0.5891255 | 0.03695941 | 0.9365958 |
| Itgad | 2.63607185 | 1.398389696 | 0.03712052 | 0.9382806 |
| Unc13b | 0.79418255 | -0.332457433 | 0.03725493 | 0.9382806 |
| Hgs | 0.90120706 | -0.150069472 | 0.03730196 | 0.9382806 |
| Rhob | 0.90551043 | -0.14319683 | 0.03736799 | 0.9382806 |
| Ina | 0.89627149 | -0.157992288 | 0.03743449 | 0.9382806 |
| Zfp763 | 1.35476619 | 0.438043888 | 0.03750262 | 0.9382806 |
| Il20rb | 1.64684399 | 0.719703887 | 0.03778385 | 0.9433181 |
| Dyrk1b | 0.82639764 | -0.275091962 | 0.03809583 | 0.9447488 |
| Bloc1s6 | 1.18920341 | 0.24999551 | 0.0381116 | 0.9447488 |
| Cacnb3 | 0.9092166 | -0.137304068 | 0.03812717 | 0.9447488 |
| Rbfox3 | 0.86301571 | -0.212541274 | 0.03816116 | 0.9447488 |
| Gpr146 | 0.80853399 | -0.306619666 | 0.03835365 | 0.9464532 |

| | | | | |
|---|---|---|---|---|
| Cyth4 | 1.264827 | 0.338940071 | 0.0383903 | 0.9464532 |
| Bcl10 | 0.85744317 | -0.221887048 | 0.03849214 | 0.9469868 |
| Lat | 0.60716128 | -0.71984831 | 0.03890935 | 0.953255 |
| Hhex | 1.66668398 | 0.736980581 | 0.03897196 | 0.953255 |
| Cdnf | 1.55178511 | 0.633928792 | 0.03898909 | 0.953255 |
| Alkbh3 | 1.11462003 | 0.156551983 | 0.03915137 | 0.9537669 |
| Scrt2 | 0.77767788 | -0.362755393 | 0.03917156 | 0.9537669 |
| Plod1 | 0.89386389 | -0.161872924 | 0.03927453 | 0.9543063 |
| Adgrg1 | 0.90862521 | -0.138242756 | 0.0394713 | 0.9571183 |
| Med25 | 0.86348138 | -0.211763028 | 0.03957306 | 0.9576193 |
| Bnip3 | 1.09967753 | 0.137080525 | 0.03971935 | 0.9578221 |
| C3ar1 | 1.64019504 | 0.713867377 | 0.03974366 | 0.9578221 |
| Mrps28 | 0.64497617 | -0.632682229 | 0.04004146 | 0.9630338 |
| Ifitm3 | 1.13282388 | 0.179923588 | 0.04027686 | 0.9634059 |
| Rbl2 | 1.13517474 | 0.182914392 | 0.04046228 | 0.9634059 |
| Aif1 | 1.23049904 | 0.299243529 | 0.04060616 | 0.9634059 |
| Igtp | 1.28685432 | 0.363848744 | 0.04065944 | 0.9634059 |
| Ckap4 | 0.87181779 | -0.19790145 | 0.0407195 | 0.9634059 |
| Mpz | 0.90266432 | -0.147738507 | 0.04079265 | 0.9634059 |
| Was | 1.56172807 | 0.643143275 | 0.04100593 | 0.9634059 |
| Mcpt8 | 1.48066288 | 0.566243201 | 0.04101668 | 0.9634059 |
| Car14 | 0.61769514 | -0.695033126 | 0.04106625 | 0.9634059 |
| Pcdhac2 | 0.90813046 | -0.139028527 | 0.04116138 | 0.9634059 |
| Map1s | 0.86470735 | -0.209716135 | 0.04120193 | 0.9634059 |
| Natd1 | 0.83694917 | -0.256788091 | 0.04120694 | 0.9634059 |
| Irgm | 1.35605073 | 0.439411153 | 0.0412292 | 0.9634059 |
| Fam213a | 0.84249078 | -0.247267192 | 0.04124179 | 0.9634059 |
| Hebp2 | 1.10133318 | 0.13925099 | 0.0413158 | 0.9634059 |
| Prkca | 0.90445017 | -0.144887077 | 0.04142139 | 0.9634059 |
| Ctps2 | 1.1258208 | 0.170977213 | 0.04146633 | 0.9634059 |
| Fhl1 | 1.10195124 | 0.140060392 | 0.04159083 | 0.9634059 |
| Pcdhga2 | 1.20413895 | 0.268001883 | 0.041607 | 0.9634059 |
| Acer2 | 1.20178651 | 0.265180628 | 0.04192571 | 0.9637836 |
| Tsen34l1 | 1.30354239 | 0.382437497 | 0.04201851 | 0.9637836 |
| Mgat4d | 1.64057798 | 0.714204174 | 0.04201928 | 0.9637836 |
| Ogdh | 0.9111568 | -0.134228752 | 0.04202528 | 0.9637836 |
| Tead2 | 0.65880942 | -0.602066921 | 0.04203138 | 0.9637836 |
| Rce1 | 1.25469226 | 0.327333553 | 0.04238033 | 0.9690959 |
| Pno1 | 1.1675912 | 0.223535238 | 0.04251274 | 0.9690959 |
| Ccbl2 | 1.23033712 | 0.29905368 | 0.04253239 | 0.9690959 |

| | | | | |
|---|---|---|---|---|
| Plk3 | 1.22687157 | 0.294984239 | 0.04261931 | 0.9690959 |
| Myo1a | 0.85994772 | -0.217679141 | 0.04275246 | 0.9690959 |
| Dab2ip | 0.90186011 | -0.149024418 | 0.04275544 | 0.9690959 |
| Bscl2 | 0.90667864 | -0.141336801 | 0.0431044 | 0.9743351 |
| Mrpl38 | 0.84358395 | -0.245396444 | 0.0431516 | 0.9743351 |
| Rcsd1 | 1.18404635 | 0.243725553 | 0.04327783 | 0.9753203 |
| Efl1 | 1.17761223 | 0.235864562 | 0.04343307 | 0.9754553 |
| Orai2 | 0.87386277 | -0.194521357 | 0.04352374 | 0.9754553 |
| H1f0 | 1.11657542 | 0.159080696 | 0.04358583 | 0.9754553 |
| Por | 0.89423853 | -0.16126838 | 0.0436524 | 0.9754553 |
| Pde7a | 1.24565916 | 0.316909368 | 0.04378668 | 0.9754553 |
| Rab3b | 1.28409703 | 0.36075422 | 0.0439911 | 0.9754553 |
| Rps4y2 | 1.40562636 | 0.491213151 | 0.04406986 | 0.9754553 |
| Slc12a2 | 1.09670264 | 0.133172404 | 0.04413949 | 0.9754553 |
| Itpr3 | 0.90696501 | -0.140881198 | 0.04414471 | 0.9754553 |
| Aamdc | 1.44465762 | 0.530727616 | 0.04417399 | 0.9754553 |
| Kif18a | 1.63863563 | 0.71249509 | 0.04420379 | 0.9754553 |
| Ccne1 | 0.78534665 | -0.348598494 | 0.04429229 | 0.9754553 |
| Scn8a | 0.91088456 | -0.134659873 | 0.04440182 | 0.9754553 |
| Pitrm1 | 1.10584581 | 0.145150243 | 0.04444025 | 0.9754553 |
| Atp6ap1l | 0.74943759 | -0.416119754 | 0.04459651 | 0.9765076 |
| Slc9a5 | 1.34859611 | 0.431458336 | 0.04465358 | 0.9765076 |
| Dkk2 | 1.37311333 | 0.457450699 | 0.04481707 | 0.9778804 |
| Pmp22 | 0.90627439 | -0.141980176 | 0.04492806 | 0.9778804 |
| Ldlr | 0.90877606 | -0.138003271 | 0.04496478 | 0.9778804 |
| Map3k1 | 1.19093123 | 0.252090105 | 0.04561838 | 0.9898361 |
| Syt1 | 0.90354622 | -0.146329698 | 0.04568216 | 0.9898361 |
| Aldh1a1 | 0.88113305 | -0.182568209 | 0.04580244 | 0.9899427 |
| Gpx7 | 0.81308503 | -0.298521859 | 0.04591018 | 0.9899427 |
| Aspm | 0.73625804 | -0.441716613 | 0.04610431 | 0.9899427 |
| Tmem202 | 0.61454015 | -0.702420825 | 0.04614608 | 0.9899427 |
| Pyroxd1 | 1.19680341 | 0.259186192 | 0.04621178 | 0.9899427 |
| Bub1 | 0.7613031 | -0.393457133 | 0.04626451 | 0.9899427 |
| Omg | 1.14159044 | 0.191045153 | 0.04627389 | 0.9899427 |
| RGD1311517 | 1.2859508 | 0.362835445 | 0.04641546 | 0.9911756 |
| Dio2 | 0.6363974 | -0.652000149 | 0.04669336 | 0.9928878 |
| Tmeff1 | 0.85080056 | -0.233107116 | 0.04672961 | 0.9928878 |
| Gas8 | 1.31205606 | 0.391829358 | 0.04687052 | 0.9928878 |
| Jund | 0.90432813 | -0.145081756 | 0.04691036 | 0.9928878 |
| Angptl2 | 0.87668345 | -0.189872078 | 0.04691603 | 0.9928878 |

| | | | | |
|---|---|---|---|---|
| Cars2 | 1.20692892 | 0.271340713 | 0.04718915 | 0.9968813 |
| Ptgdr | 0.50959851 | -0.972567024 | 0.04733331 | 0.9976525 |
| Kcnb1 | 0.88429446 | -0.177401251 | 0.04739462 | 0.9976525 |
| Aacs | 0.90397192 | -0.145650141 | 0.04761932 | 1 |
| Etfb | 1.13955372 | 0.18846894 | 0.04785972 | 1 |
| Sepp1 | 1.09781369 | 0.13463323 | 0.04793427 | 1 |
| Nr2c1 | 1.18203509 | 0.241272862 | 0.04796792 | 1 |
| Id4 | 1.15224146 | 0.204443074 | 0.04817286 | 1 |
| Crtac1 | 1.17679777 | 0.234866418 | 0.04819359 | 1 |
| Slc27a1 | 0.89798932 | -0.155229802 | 0.04821264 | 1 |
| Hp | 1.48074781 | 0.566325956 | 0.04838335 | 1 |
| Abcd2 | 1.12555773 | 0.170640051 | 0.04850573 | 1 |
| Itgal | 1.31152985 | 0.39125064 | 0.04913364 | 1 |
| Pak6 | 0.6029265 | -0.729945954 | 0.04916366 | 1 |
| Prep | 0.89785236 | -0.155449856 | 0.0492923 | 1 |
| Kcng1 | 0.81928192 | -0.287568121 | 0.0493086 | 1 |
| Setd7 | 1.11851527 | 0.161584953 | 0.0495152 | 1 |
| Inpp5j | 0.87977677 | -0.184790583 | 0.04954127 | 1 |
| Id1 | 0.84760406 | -0.238537605 | 0.04991226 | 1 |
| Gpnmb | 1.19296688 | 0.25455399 | 0.04997366 | 1 |
| Papss2 | 1.27129565 | 0.346299582 | 0.05001308 | 1 |
| Scn3b | 0.90046685 | -0.151254933 | 0.05003352 | 1 |
| RT1-A3 | 0.70185261 | -0.510760005 | 0.05013349 | 1 |
| P2ry1 | 0.88219036 | -0.180838095 | 0.05018891 | 1 |
| Ccl24 | 1.64266593 | 0.716039104 | 0.05026328 | 1 |
| RGD1563365 | 0.73745992 | -0.439363448 | 0.05039904 | 1 |
| Dbp | 0.89943105 | -0.152915411 | 0.0505586 | 1 |
| Rfx2 | 1.50407353 | 0.588875096 | 0.05098318 | 1 |
| Med15 | 0.8797946 | -0.184761341 | 0.05101415 | 1 |
| Nnat | 0.87019766 | -0.200584955 | 0.05106207 | 1 |
| Lmod1 | 1.31537888 | 0.395478412 | 0.05119205 | 1 |
| Mblac2 | 0.8530399 | -0.229314869 | 0.05126568 | 1 |
| Cd55 | 0.91433994 | -0.129197462 | 0.05127592 | 1 |
| Agt | 0.75617627 | -0.40320551 | 0.05146105 | 1 |
| RatNP-3b | 1.39195349 | 0.47711101 | 0.05156504 | 1 |
| Aoah | 1.90778201 | 0.931896336 | 0.05170973 | 1 |
| Bmp6 | 1.21723291 | 0.283605242 | 0.0517229 | 1 |
| P2ry14 | 1.36273371 | 0.446503678 | 0.05195926 | 1 |
| Usp34 | 0.91456803 | -0.128837608 | 0.05197921 | 1 |
| Selp | 1.52863328 | 0.612242341 | 0.05222361 | 1 |

| | | | | |
|---|---|---|---|---|
| Csf1r | 1.12286595 | 0.167185701 | 0.05242563 | 1 |
| Cbx6 | 0.91089305 | -0.134646421 | 0.05246215 | 1 |
| Bcl6 | 0.87794188 | -0.187802656 | 0.052656 | 1 |
| Csrp1 | 1.09298892 | 0.128278781 | 0.05270195 | 1 |
| Myot | 0.43457566 | -1.202320738 | 0.05276249 | 1 |
| Dusp10 | 1.3426427 | 0.425075429 | 0.0528663 | 1 |
| Mcpt9 | 2.25979103 | 1.176189371 | 0.05294813 | 1 |
| Emc2 | 1.09715402 | 0.133766064 | 0.05299603 | 1 |
| RT1-CE16 | 0.85791035 | -0.221101194 | 0.05305842 | 1 |
| Dtymk | 1.15125373 | 0.203205834 | 0.05307246 | 1 |
| Syt15 | 1.30010043 | 0.378623074 | 0.05320353 | 1 |
| Acbd4 | 1.1705348 | 0.227167823 | 0.05333941 | 1 |
| RT1-Db1 | 1.13935211 | 0.18821367 | 0.05340453 | 1 |
| Sox10 | 0.90033126 | -0.151472179 | 0.05342186 | 1 |
| Haus8 | 1.30113558 | 0.379771304 | 0.05345223 | 1 |
| Col14a1 | 0.90878301 | -0.137992227 | 0.05355622 | 1 |
| Adra2a | 0.72870945 | -0.456584393 | 0.05365439 | 1 |
| Plppr3 | 0.77815869 | -0.361863696 | 0.05371663 | 1 |
| Ubqln1 | 0.91256769 | -0.13199652 | 0.05378342 | 1 |
| Skap1 | 0.6166591 | -0.69745494 | 0.05384595 | 1 |
| Cdc26 | 0.86417369 | -0.21060678 | 0.05394582 | 1 |
| Sorcs1 | 0.88133133 | -0.182243597 | 0.05423509 | 1 |
| Inhba | 1.61343856 | 0.690138643 | 0.05432041 | 1 |
| Hid1 | 0.90993828 | -0.136159409 | 0.05443653 | 1 |
| Rpap2 | 1.16788167 | 0.223894109 | 0.05458116 | 1 |
| Ap4e1 | 1.18535697 | 0.245321589 | 0.0547245 | 1 |
| Ankzf1 | 1.15154872 | 0.203575455 | 0.05487749 | 1 |
| Tef | 0.913355 | -0.130752387 | 0.05494993 | 1 |
| Mtmr1 | 0.85777307 | -0.221332069 | 0.05503108 | 1 |
| Lrrn3 | 0.89106639 | -0.166395173 | 0.05504733 | 1 |
| Tspan14 | 0.86993977 | -0.201012582 | 0.05517924 | 1 |
| Bmper | 0.82497782 | -0.277572755 | 0.0553047 | 1 |
| Srcin1 | 0.84285336 | -0.24664645 | 0.05544399 | 1 |
| Rasgrp1 | 0.86518782 | -0.208914733 | 0.05546277 | 1 |
| Foxk2 | 0.88283901 | -0.179777712 | 0.05551793 | 1 |
| Hs6st3 | 0.77918059 | -0.359970347 | 0.05579305 | 1 |
| Clstn3 | 0.91348569 | -0.130545969 | 0.05589939 | 1 |
| Cog7 | 1.13798966 | 0.186487447 | 0.05600389 | 1 |
| Cox6b2 | 1.68454849 | 0.752361958 | 0.05601838 | 1 |
| Ubr4 | 0.91563277 | -0.127158997 | 0.0560355 | 1 |

| | | | | |
|---|---|---|---|---|
| Lgals9 | 1.19342111 | 0.255103196 | 0.05616243 | 1 |
| Ephb6 | 0.87509083 | -0.192495317 | 0.05619146 | 1 |
| Mrpl41 | 1.12179571 | 0.165809973 | 0.05619845 | 1 |
| Apc2 | 0.91386352 | -0.129949378 | 0.05629881 | 1 |
| Caskin1 | 0.91358276 | -0.130392676 | 0.05642243 | 1 |
| Nxph1 | 1.26622881 | 0.340538127 | 0.05643116 | 1 |
| Hltf | 1.13593174 | 0.183876139 | 0.05645724 | 1 |
| Slc14a1 | 0.67000716 | -0.577751592 | 0.05670386 | 1 |
| Ccdc125 | 1.49049401 | 0.575790581 | 0.05690162 | 1 |
| Tnn | 0.45369831 | -1.140194806 | 0.05701054 | 1 |
| Mid1 | 1.30521541 | 0.38428793 | 0.05710593 | 1 |
| Cacna1h | 0.89021813 | -0.167769207 | 0.05714013 | 1 |
| Itgb7 | 0.68834819 | -0.538789572 | 0.05726345 | 1 |
| Slit2 | 0.90364653 | -0.146169542 | 0.05747198 | 1 |
| Tmem72 | 0.85798481 | -0.220975982 | 0.05751133 | 1 |
| Casc3 | 0.88573907 | -0.175046341 | 0.0575609 | 1 |
| Dirc2 | 0.89184303 | -0.165138293 | 0.05765335 | 1 |
| Eps15l1 | 1.11896329 | 0.162162708 | 0.05777075 | 1 |
| Adgrl1 | 0.90661886 | -0.141431916 | 0.05779092 | 1 |
| Lpcat1 | 0.87868611 | -0.186580203 | 0.05780807 | 1 |
| Gnao1 | 0.91695881 | -0.125071158 | 0.05783041 | 1 |
| Tceanc2 | 1.19360413 | 0.25532443 | 0.05784853 | 1 |
| Csrp2 | 0.83076855 | -0.2674815 | 0.05788316 | 1 |
| Scd2 | 0.91006185 | -0.135963501 | 0.05798384 | 1 |
| Gins1 | 0.70961548 | -0.494890611 | 0.05803577 | 1 |
| Lyn | 1.23208834 | 0.301105703 | 0.05807753 | 1 |
| Ncan | 0.83483467 | -0.260437572 | 0.05808203 | 1 |
| Myzap | 1.42724962 | 0.513237679 | 0.05816983 | 1 |
| Oas1a | 1.37083992 | 0.455060111 | 0.05853511 | 1 |
| Gfra3 | 0.86801474 | -0.204208551 | 0.05859096 | 1 |
| Polr2g | 1.12857481 | 0.174502054 | 0.05876551 | 1 |
| Csad | 0.84973878 | -0.23490868 | 0.05889476 | 1 |
| Kel | 1.59982834 | 0.677917115 | 0.05929158 | 1 |
| Jup | 0.91484207 | -0.128405385 | 0.05956908 | 1 |
| Cd93 | 1.19927823 | 0.262166404 | 0.05969759 | 1 |
| Dtd1 | 1.12704869 | 0.172549847 | 0.05969905 | 1 |
| Ntng2 | 0.89293802 | -0.16336806 | 0.05978045 | 1 |
| Pnmal2 | 0.88070403 | -0.183270832 | 0.05986577 | 1 |
| Clcn4 | 1.0916889 | 0.126561789 | 0.05986757 | 1 |
| Lgr5 | 0.88123042 | -0.182408795 | 0.0599257 | 1 |

| | | | | |
|---|---|---|---|---|
| Abcb4 | 1.454436 | 0.540459818 | 0.06038201 | 1 |
| Fa2h | 0.88751805 | -0.17215164 | 0.06038943 | 1 |
| Mppe1 | 1.18678186 | 0.247054779 | 0.06039613 | 1 |
| Bre | 1.11991758 | 0.163392563 | 0.06046233 | 1 |
| Taf9b | 1.14625601 | 0.196929298 | 0.06054996 | 1 |
| Mul1 | 1.17090663 | 0.227626037 | 0.06065275 | 1 |
| Lsm10 | 1.35559152 | 0.438922516 | 0.06065778 | 1 |
| Pcdhga11 | 0.82135627 | -0.283919949 | 0.06097725 | 1 |
| Bmp5 | 1.23606664 | 0.305756521 | 0.06101521 | 1 |
| Ripk4 | 0.70995612 | -0.494198229 | 0.06112218 | 1 |
| Cadm1 | 0.91429512 | -0.129268178 | 0.06120788 | 1 |
| Fat1 | 0.90926043 | -0.13723453 | 0.06152503 | 1 |
| Atp8b2 | 0.91384392 | -0.129980309 | 0.06169365 | 1 |
| Sparc | 0.91250588 | -0.132094234 | 0.06179292 | 1 |
| Tex15 | 0.80734471 | -0.308743303 | 0.06191636 | 1 |
| Ino80e | 0.86560264 | -0.208223192 | 0.06226263 | 1 |
| Tp53inp2 | 0.90770936 | -0.139697659 | 0.06288247 | 1 |
| Slc25a14 | 1.16566895 | 0.221158124 | 0.06294786 | 1 |
| Tsc2 | 0.89902979 | -0.153559177 | 0.06294822 | 1 |
| Tsc22d3 | 0.91732039 | -0.124502386 | 0.0629812 | 1 |
| Errfi1 | 0.89077265 | -0.166870833 | 0.06311928 | 1 |
| P2rx6 | 1.14026657 | 0.18937114 | 0.06316738 | 1 |
| Myc | 0.78640722 | -0.346651519 | 0.06336859 | 1 |
| Gemin7 | 0.84292784 | -0.246518963 | 0.06339091 | 1 |
| Tm4sf1 | 1.14174793 | 0.191244177 | 0.06343928 | 1 |
| Tnk2 | 0.91239168 | -0.132274809 | 0.06345317 | 1 |
| Frem2 | 1.50637632 | 0.591082224 | 0.06348716 | 1 |
| Fcrl2 | 0.76105389 | -0.393929483 | 0.06357356 | 1 |
| Tmem86a | 1.31195287 | 0.391715896 | 0.06358327 | 1 |
| Lgals3bp | 1.10377068 | 0.14244047 | 0.06372048 | 1 |
| Inpp1 | 1.16659999 | 0.222309968 | 0.0637481 | 1 |
| Tmem33 | 0.90588511 | -0.142600005 | 0.06385429 | 1 |
| Plat | 1.09142164 | 0.126208556 | 0.06387252 | 1 |
| Dgki | 0.90033517 | -0.151465914 | 0.06403342 | 1 |
| Eif2s3y | 1.09247451 | 0.127599616 | 0.0642788 | 1 |
| Loxl1 | 0.87217133 | -0.197316536 | 0.06429266 | 1 |
| Xk | 0.81323871 | -0.2982492 | 0.06467703 | 1 |
| Cpsf1 | 0.88614365 | -0.174387511 | 0.06472456 | 1 |
| Klf1 | 1.55887509 | 0.64050533 | 0.06472467 | 1 |
| Tmem176a | 0.91853679 | -0.122590589 | 0.0647665 | 1 |

| | | | | |
|---|---|---|---|---|
| Aph1b | 0.86383037 | -0.211180061 | 0.06480933 | 1 |
| Pum2 | 0.91699269 | -0.125017863 | 0.06481596 | 1 |
| Tnnt3 | 0.78990659 | -0.340246041 | 0.06506778 | 1 |
| Isl2 | 0.88616608 | -0.174350997 | 0.06515761 | 1 |
| Rpl34 | 1.1041731 | 0.142966361 | 0.06519692 | 1 |
| Hspa5 | 0.91648064 | -0.125823684 | 0.06523445 | 1 |
| Ncf1 | 1.22962245 | 0.298215413 | 0.06538544 | 1 |
| Pdlim5 | 0.84989186 | -0.234648812 | 0.06546792 | 1 |
| Fcgbp | 1.42914912 | 0.515156453 | 0.06551437 | 1 |
| Dab2 | 1.12699595 | 0.172482329 | 0.06553885 | 1 |
| Dpf1 | 1.3868174 | 0.471777839 | 0.06554513 | 1 |
| Sec31a | 0.90737488 | -0.14022938 | 0.06556428 | 1 |
| Tmem164 | 0.90480368 | -0.144323304 | 0.06568806 | 1 |
| Nlgn3 | 0.86556512 | -0.208285738 | 0.06590922 | 1 |
| Ndufc2 | 1.10461807 | 0.143547638 | 0.06606232 | 1 |
| Tmem209 | 0.83770522 | -0.255485425 | 0.06623207 | 1 |
| Zfp39 | 1.39935013 | 0.484756983 | 0.06629898 | 1 |
| Rnase4 | 1.12628653 | 0.171573897 | 0.06636553 | 1 |
| Tmc8 | 0.65761885 | -0.604676439 | 0.06654094 | 1 |
| Glrb | 1.09019593 | 0.124587433 | 0.06654282 | 1 |
| Commd9 | 1.1403065 | 0.18942166 | 0.0666349 | 1 |
| Commd10 | 1.14325048 | 0.193141519 | 0.06672464 | 1 |
| Hivep1 | 0.90205106 | -0.148718993 | 0.06676373 | 1 |
| Jmjd4 | 1.43873348 | 0.52479936 | 0.06681583 | 1 |
| Hecw2 | 0.82568955 | -0.276328652 | 0.06681804 | 1 |
| Vamp2 | 0.91850726 | -0.122636976 | 0.06721978 | 1 |
| Rbm33 | 0.86627604 | -0.207101275 | 0.06735083 | 1 |
| Isyna1 | 1.12408255 | 0.168747982 | 0.06737323 | 1 |
| Ttc38 | 0.75839646 | -0.398975868 | 0.06737903 | 1 |
| Laptm4b | 1.08914836 | 0.12320048 | 0.06762633 | 1 |
| Ahsp | 1.29502714 | 0.372982329 | 0.06806775 | 1 |
| Psmb8 | 1.21857546 | 0.285195598 | 0.06816422 | 1 |
| Padi2 | 0.88561063 | -0.175255559 | 0.06843465 | 1 |
| Ddx56 | 0.83789936 | -0.25515112 | 0.06849748 | 1 |
| Zfp956 | 1.32796608 | 0.409218296 | 0.06852454 | 1 |
| Mrpl15 | 1.1645245 | 0.219740992 | 0.06874543 | 1 |
| Lmna | 0.91701724 | -0.124979239 | 0.06879523 | 1 |
| Mcm7 | 1.20115986 | 0.264428166 | 0.06894546 | 1 |
| Mphosph8 | 1.0932025 | 0.128560662 | 0.06897797 | 1 |
| Faah | 0.79542863 | -0.330195606 | 0.06899853 | 1 |

| | | | | |
|---|---|---|---|---|
| Donson | 0.81977516 | -0.286699811 | 0.06910767 | 1 |
| Adcyap1 | 0.87127545 | -0.198799204 | 0.06929968 | 1 |
| Ppa1 | 1.09016239 | 0.124543053 | 0.06934983 | 1 |
| Vof16 | 1.230551 | 0.29930445 | 0.0694388 | 1 |
| RGD1564379 | 0.89175291 | -0.165284069 | 0.06956871 | 1 |
| Wdr92 | 1.17282547 | 0.229988344 | 0.06957542 | 1 |
| Tcerg1l | 1.31014417 | 0.389725577 | 0.06968435 | 1 |
| Golga5 | 1.12906841 | 0.175132899 | 0.06970846 | 1 |
| Smoc2 | 0.9023649 | -0.148217148 | 0.0697521 | 1 |
| Hnrnpl | 0.91434945 | -0.129182448 | 0.06978337 | 1 |
| Ace2 | 1.24229013 | 0.313002143 | 0.06982706 | 1 |
| Slc36a4 | 0.83769789 | -0.255498049 | 0.06990629 | 1 |
| Tap2 | 1.17656772 | 0.23458436 | 0.07012743 | 1 |
| Sgpp2 | 0.88227792 | -0.180694913 | 0.07019283 | 1 |
| Exoc8 | 0.88279043 | -0.179857101 | 0.07027017 | 1 |
| Pltp | 1.1383803 | 0.186982595 | 0.0705272 | 1 |
| Cyb561a3 | 1.18006143 | 0.238861965 | 0.07057426 | 1 |
| LOC100365921 | 1.24464787 | 0.315737636 | 0.07059712 | 1 |
| Rn5-8s | 1.44762222 | 0.533685155 | 0.07076389 | 1 |
| Atrn | 0.92054266 | -0.119443514 | 0.07077557 | 1 |
| Ache | 0.92022608 | -0.119939749 | 0.07078212 | 1 |
| Zfp213 | 1.2645947 | 0.338675078 | 0.07080577 | 1 |
| Cd22 | 1.45522126 | 0.541238525 | 0.07094606 | 1 |
| Ubap2 | 0.91513667 | -0.127940876 | 0.07126655 | 1 |
| Armc7 | 0.79082389 | -0.338571634 | 0.07148309 | 1 |
| Pih1d1 | 1.22295491 | 0.290371213 | 0.07159741 | 1 |
| Mag | 0.85669379 | -0.223148464 | 0.07161617 | 1 |
| Plac8 | 1.19182325 | 0.253170293 | 0.07161911 | 1 |
| Abcg1 | 0.9113832 | -0.133870315 | 0.07170844 | 1 |
| Klf15 | 0.87081286 | -0.199565388 | 0.07184941 | 1 |
| Tor1a | 0.86941834 | -0.201877563 | 0.07186478 | 1 |
| Plk2 | 0.883239 | -0.179124224 | 0.07214671 | 1 |
| Bex3 | 1.13305533 | 0.180218311 | 0.07220728 | 1 |
| Ptprn | 0.91941109 | -0.121218022 | 0.07223877 | 1 |
| Gpm6b | 0.9208454 | -0.118969125 | 0.07224409 | 1 |
| Sema3g | 0.89384312 | -0.161906453 | 0.07237408 | 1 |
| Rftn1 | 1.29824315 | 0.376560611 | 0.07238257 | 1 |
| Msr1 | 1.38524735 | 0.470143608 | 0.07282717 | 1 |
| Dgkz | 0.92089714 | -0.118888077 | 0.07306501 | 1 |
| Nxpe3 | 1.09954054 | 0.136900791 | 0.07325015 | 1 |

| | | | | |
|---|---|---|---|---|
| Ank1 | 0.92144678 | -0.118027256 | 0.07334221 | 1 |
| Tcaim | 1.15184577 | 0.203947551 | 0.07356848 | 1 |
| Nfatc3 | 0.89205272 | -0.164799119 | 0.07366537 | 1 |
| Ccsap | 1.50227857 | 0.587152362 | 0.07367243 | 1 |
| Traf3 | 0.90693451 | -0.140929713 | 0.0738544 | 1 |
| Adamts7 | 0.81755499 | -0.290612329 | 0.07443648 | 1 |
| Kctd1 | 0.8600766 | -0.217462942 | 0.07443763 | 1 |
| Snta1 | 0.87653944 | -0.19010909 | 0.07448064 | 1 |
| Ptch1 | 1.11681413 | 0.159389097 | 0.07454212 | 1 |
| Aox4 | 1.21644354 | 0.282669357 | 0.07457588 | 1 |
| Rdh16 | 1.41965485 | 0.50554022 | 0.07469452 | 1 |
| Cbfa2t3 | 0.68904532 | -0.537329217 | 0.0747296 | 1 |
| Napb | 1.09694352 | 0.133489247 | 0.07475693 | 1 |
| Sepsecs | 1.14968436 | 0.201237828 | 0.07482899 | 1 |
| Irgq | 0.91846993 | -0.122695601 | 0.07489649 | 1 |
| Ube2z | 0.91834514 | -0.122891628 | 0.07501732 | 1 |
| Prorsd1 | 1.48207931 | 0.567622651 | 0.07525267 | 1 |
| Chpt1 | 1.20639923 | 0.270707415 | 0.07525809 | 1 |
| Tspan18 | 0.88659912 | -0.173646163 | 0.07527373 | 1 |
| Clic1 | 1.0977284 | 0.134521144 | 0.07527592 | 1 |
| Spg7 | 1.10969114 | 0.150158189 | 0.07530083 | 1 |
| Crtc2 | 0.86503094 | -0.209176358 | 0.07539629 | 1 |
| Clip3 | 0.92154176 | -0.117878554 | 0.07549159 | 1 |
| Senp8 | 1.26162712 | 0.33528558 | 0.07568649 | 1 |
| Plekhg2 | 0.89817074 | -0.154938374 | 0.07595947 | 1 |
| Purb | 0.88013111 | -0.184209643 | 0.07606592 | 1 |
| Mpzl2 | 1.1813247 | 0.240405556 | 0.0762319 | 1 |
| Pxmp2 | 1.29332015 | 0.371079446 | 0.07633958 | 1 |
| Aph1a | 0.8878012 | -0.171691437 | 0.07634758 | 1 |
| C1qa | 1.12725446 | 0.172813223 | 0.07635219 | 1 |
| Mterf3 | 0.86723098 | -0.205511802 | 0.07637978 | 1 |
| Ptprcap | 0.62636679 | -0.674920372 | 0.07647352 | 1 |
| Slc9a2 | 1.20553709 | 0.269676037 | 0.0768286 | 1 |
| Efhc2 | 1.28188422 | 0.358265968 | 0.07690647 | 1 |
| Speg | 0.87110808 | -0.199076367 | 0.07723186 | 1 |
| Kcnd3 | 0.91890605 | -0.122010728 | 0.07723525 | 1 |
| Pla2g2d | 0.63511745 | -0.654904683 | 0.07731746 | 1 |
| Engase | 0.84614827 | -0.241017612 | 0.0773404 | 1 |
| Tsen15 | 0.88304562 | -0.17944013 | 0.07747598 | 1 |
| Gpr35 | 0.76340538 | -0.389478746 | 0.07749828 | 1 |

| | | | | |
|---|---|---|---|---|
| Cdh1 | 0.92249234 | -0.116391157 | 0.07749934 | 1 |
| Tigd5 | 1.51341088 | 0.597803724 | 0.07762914 | 1 |
| Cebpz | 1.10814318 | 0.148144301 | 0.07772045 | 1 |
| Map7d2 | 1.08605558 | 0.11909794 | 0.07781725 | 1 |
| Smim3 | 1.17045825 | 0.227073479 | 0.07790222 | 1 |
| Sec14l1 | 0.91982882 | -0.120562695 | 0.07793553 | 1 |
| Scyl1 | 0.91196345 | -0.132952086 | 0.07795667 | 1 |
| Usp31 | 0.91195323 | -0.132968252 | 0.07830931 | 1 |
| Gpc3 | 1.0959092 | 0.132128265 | 0.07832639 | 1 |
| Crmp1 | 0.90960744 | -0.136684036 | 0.07852343 | 1 |
| Dusp5 | 0.84562228 | -0.241914702 | 0.07862397 | 1 |
| Zfp423 | 1.15125857 | 0.203211897 | 0.07871073 | 1 |
| Trim34 | 1.19511532 | 0.257149837 | 0.0788106 | 1 |
| Slc11a2 | 1.13512611 | 0.182852585 | 0.07889506 | 1 |
| Chka | 0.89051152 | -0.167293822 | 0.07912855 | 1 |
| Mmd | 0.89361331 | -0.162277425 | 0.07941897 | 1 |
| Lap3 | 1.09961216 | 0.136994768 | 0.07950006 | 1 |
| Reg3a | 0.31540385 | -1.664727843 | 0.07963001 | 1 |
| Ckmt1b | 1.08452949 | 0.117069277 | 0.07971634 | 1 |
| Irak3 | 1.17031397 | 0.226895626 | 0.07972469 | 1 |
| Hacd4 | 1.24564112 | 0.316888476 | 0.07983694 | 1 |
| Tmem231 | 1.13602055 | 0.183988929 | 0.07985798 | 1 |
| Rcn1 | 0.92023528 | -0.119925321 | 0.07996622 | 1 |
| Vsir | 1.15517268 | 0.208108532 | 0.08007546 | 1 |
| Lum | 0.92223356 | -0.116795928 | 0.08010735 | 1 |
| Clcn6 | 0.91458859 | -0.128805166 | 0.08019291 | 1 |
| Ly49s7 | 0.59540726 | -0.748051279 | 0.08025544 | 1 |
| Csdc2 | 1.10440249 | 0.143266045 | 0.0807478 | 1 |
| Apbb1ip | 1.25743528 | 0.330484146 | 0.08083353 | 1 |
| Cdc42bpb | 0.91662479 | -0.125596786 | 0.080948 | 1 |
| S100a9 | 1.12765578 | 0.173326745 | 0.08106759 | 1 |
| Casp12 | 1.29351817 | 0.37130032 | 0.08118643 | 1 |
| Fut10 | 0.7159282 | -0.482113182 | 0.08126837 | 1 |
| Ddrgk1 | 1.11217348 | 0.153381845 | 0.08137317 | 1 |
| Foxp2 | 1.21816787 | 0.284712963 | 0.08138287 | 1 |
| Dpp9 | 0.91070359 | -0.134946527 | 0.08150858 | 1 |
| Sv2c | 0.92337563 | -0.115010433 | 0.08179043 | 1 |
| Abat | 1.14524646 | 0.195658098 | 0.08188785 | 1 |
| Coq2 | 1.15476134 | 0.207594719 | 0.08190385 | 1 |
| Slc17a8 | 0.73628388 | -0.441665976 | 0.08190849 | 1 |

| | | | | |
|---|---|---|---|---|
| Nsd1 | 0.92027772 | -0.119858793 | 0.08191367 | 1 |
| Smarca4 | 0.91958299 | -0.120948312 | 0.08195917 | 1 |
| Syngap1 | 0.86026326 | -0.217149873 | 0.08201935 | 1 |
| Socs2 | 0.79833395 | -0.324935723 | 0.08202646 | 1 |
| Fcna | 1.28276962 | 0.359262094 | 0.08202821 | 1 |
| Slit1 | 0.9130263 | -0.131271672 | 0.08245261 | 1 |
| Il17re | 1.37144578 | 0.455697588 | 0.08246015 | 1 |
| Vars | 0.90234214 | -0.148253538 | 0.08251494 | 1 |
| Scn1a | 1.08351641 | 0.115721005 | 0.08253124 | 1 |
| Etfbkmt | 1.28168704 | 0.358044029 | 0.0825645 | 1 |
| Xpr1 | 0.87644627 | -0.190262452 | 0.08284589 | 1 |
| Mtor | 0.91528639 | -0.127704859 | 0.08287034 | 1 |
| Gtse1 | 1.40698447 | 0.492606406 | 0.08290836 | 1 |
| Cyp4f4 | 1.53099141 | 0.614466192 | 0.08306984 | 1 |
| Nelfb | 0.91950967 | -0.121063349 | 0.08327476 | 1 |
| Zc3h7b | 0.92175923 | -0.117538138 | 0.08334463 | 1 |
| RGD1310587 | 1.12602679 | 0.171241146 | 0.08339118 | 1 |
| Zbtb6 | 1.18323934 | 0.242741919 | 0.08344174 | 1 |
| Mrpl14 | 1.16602511 | 0.221598862 | 0.08360223 | 1 |
| Dmap1 | 1.15239438 | 0.204634531 | 0.08374231 | 1 |
| Palm | 0.91741762 | -0.124349472 | 0.08377117 | 1 |
| Gadd45b | 1.19760994 | 0.260158103 | 0.08396763 | 1 |
| Tpx2 | 0.78093231 | -0.356730593 | 0.08420742 | 1 |
| Traf1 | 1.44713536 | 0.53319987 | 0.08446244 | 1 |
| Arg1 | 0.65549768 | -0.60933742 | 0.08448218 | 1 |
| Zswim5 | 0.87486976 | -0.192859834 | 0.08474218 | 1 |
| Gga1 | 0.89617073 | -0.158154489 | 0.08492061 | 1 |
| Actr8 | 1.11760503 | 0.160410414 | 0.08492366 | 1 |
| Epas1 | 0.91516508 | -0.127896094 | 0.08496392 | 1 |
| Adgrf5 | 1.12061941 | 0.164296385 | 0.08499605 | 1 |
| Smug1 | 0.76160685 | -0.39288164 | 0.08503489 | 1 |
| Ddhd1 | 0.89960805 | -0.15263152 | 0.08506397 | 1 |
| Erf | 0.86385236 | -0.211143325 | 0.08509158 | 1 |
| Plcl1 | 1.13929768 | 0.188144749 | 0.08510661 | 1 |
| Ddx51 | 1.19846881 | 0.261192365 | 0.08525135 | 1 |
| Ch25h | 1.33895696 | 0.421109592 | 0.08528075 | 1 |
| Crbn | 1.08262609 | 0.114535067 | 0.0853834 | 1 |
| Lyrm1 | 1.15175074 | 0.203828529 | 0.08556598 | 1 |
| Arid3b | 0.7571228 | -0.401400772 | 0.08577186 | 1 |
| Ubald1 | 0.87905021 | -0.185982531 | 0.08579365 | 1 |

| | | | | |
|---|---|---|---|---|
| Fam46c | 1.29348309 | 0.371261193 | 0.08579487 | 1 |
| Prkx | 0.90331162 | -0.146704323 | 0.08586303 | 1 |
| Spg21 | 0.8940156 | -0.161628096 | 0.08598322 | 1 |
| Nptx1 | 0.92397867 | -0.11406854 | 0.08608438 | 1 |
| Perp | 1.1610893 | 0.215478939 | 0.08615426 | 1 |
| Abcc8 | 0.8460274 | -0.241223711 | 0.08660831 | 1 |
| Myh9 | 0.91675075 | -0.125398547 | 0.08682542 | 1 |
| ST7 | 1.19392235 | 0.255709015 | 0.08699352 | 1 |
| Tmem229a | 0.92100544 | -0.118718418 | 0.08715738 | 1 |
| Mybpc1 | 0.7379665 | -0.438372771 | 0.08716926 | 1 |
| RT1-S3 | 1.13238625 | 0.179366141 | 0.08735632 | 1 |
| Atp1a3 | 0.92378153 | -0.114376397 | 0.08745807 | 1 |
| Haus1 | 1.17641804 | 0.234400814 | 0.08759339 | 1 |
| Kctd8 | 0.90761553 | -0.139846797 | 0.08762473 | 1 |
| Pcdha11 | 0.78163451 | -0.355433923 | 0.08775414 | 1 |
| Ret | 0.92279531 | -0.115917425 | 0.08800938 | 1 |
| Hoxc4 | 1.25097383 | 0.323051608 | 0.08818718 | 1 |
| Pef1 | 1.09900334 | 0.136195777 | 0.0883963 | 1 |
| Pcdhac1 | 0.90352568 | -0.146362492 | 0.08841364 | 1 |
| Il1rl1 | 1.54705451 | 0.629524033 | 0.08844196 | 1 |
| Slc46a2 | 0.73089076 | -0.4522723 | 0.08844276 | 1 |
| Gmip | 1.18067084 | 0.239606806 | 0.08850322 | 1 |
| Sesn3 | 0.90601468 | -0.142393672 | 0.08867573 | 1 |
| Trmt61a | 0.86340256 | -0.211894723 | 0.08871005 | 1 |
| Cfh | 1.08576207 | 0.118707985 | 0.08881245 | 1 |
| Aste1 | 0.73458992 | -0.444988986 | 0.08883515 | 1 |
| Grik3 | 0.764383 | -0.387632398 | 0.0888902 | 1 |
| Rbks | 1.30816518 | 0.38754472 | 0.0889383 | 1 |
| Fcnb | 1.33101937 | 0.412531571 | 0.08908332 | 1 |
| Ccdc89 | 1.38319145 | 0.468000853 | 0.0892026 | 1 |
| Sptbn4 | 0.9079762 | -0.139273621 | 0.08930098 | 1 |
| Pcdh9 | 1.0864303 | 0.119595622 | 0.08938404 | 1 |
| Magi2 | 0.85567582 | -0.224863774 | 0.08954919 | 1 |
| Htr3b | 1.09022048 | 0.124619931 | 0.08958121 | 1 |
| Farp1 | 0.88331828 | -0.178994728 | 0.08965133 | 1 |
| Snx29 | 0.82728921 | -0.27353633 | 0.08971897 | 1 |
| Rngtt | 0.89080574 | -0.166817234 | 0.08976374 | 1 |
| Ddb2 | 1.40271679 | 0.488223756 | 0.08987794 | 1 |
| Cd3eap | 0.77458911 | -0.368496876 | 0.08989805 | 1 |
| Plscr3 | 0.88966228 | -0.168670305 | 0.08991156 | 1 |

| | | | | |
|---|---|---|---|---|
| Atp5o | 1.08107174 | 0.112462269 | 0.09067422 | 1 |
| Gem | 0.74502848 | -0.424632519 | 0.09070066 | 1 |
| Pgls | 0.85167859 | -0.231619007 | 0.09102541 | 1 |
| Akt1 | 0.91895453 | -0.121934624 | 0.09102795 | 1 |
| Shc3 | 0.71692429 | -0.480107325 | 0.09156925 | 1 |
| Plekha3 | 0.8788539 | -0.186304746 | 0.09168398 | 1 |
| Rbm4b | 0.83726565 | -0.256242652 | 0.09178117 | 1 |
| Slfn13 | 1.14708987 | 0.19797843 | 0.09186355 | 1 |
| Htra1 | 1.08161114 | 0.113181916 | 0.09188117 | 1 |
| Lnx2 | 0.85080436 | -0.233100676 | 0.0919375 | 1 |
| Zfp35 | 1.32190518 | 0.402618698 | 0.0919503 | 1 |
| E2f4 | 1.1609685 | 0.215328826 | 0.09206893 | 1 |
| Epb41l4a | 0.84329149 | -0.245896695 | 0.09229256 | 1 |
| Prr5l | 1.57189047 | 0.652500697 | 0.09233004 | 1 |
| Slc6a9 | 1.32688705 | 0.408045565 | 0.09236139 | 1 |
| Fam151a | 0.76557217 | -0.385389712 | 0.09246059 | 1 |
| Rcbtb2 | 1.1668225 | 0.222585116 | 0.09252064 | 1 |
| Zbtb7c | 1.23905463 | 0.309239795 | 0.09257434 | 1 |
| Isg15 | 1.34346699 | 0.425960869 | 0.09258933 | 1 |
| Slc2a6 | 0.89361478 | -0.162275052 | 0.09260495 | 1 |
| RGD1311739 | 0.89541547 | -0.159370851 | 0.09263194 | 1 |
| Chrna4 | 0.65507923 | -0.610258689 | 0.09268429 | 1 |
| Nedd4l | 0.92290436 | -0.115746943 | 0.09304291 | 1 |
| Adck3 | 1.15785943 | 0.211460117 | 0.09314599 | 1 |
| Efnb1 | 1.12086291 | 0.164609839 | 0.09336883 | 1 |
| G0s2 | 0.79579725 | -0.329527188 | 0.09344159 | 1 |
| Galt | 0.85532197 | -0.225460492 | 0.09374817 | 1 |
| Snph | 1.08999381 | 0.124319936 | 0.09387891 | 1 |
| Snapin | 1.0927643 | 0.127982259 | 0.09396736 | 1 |
| Arhgef12 | 0.91208566 | -0.132758763 | 0.0940266 | 1 |
| Olfm4 | 1.93038882 | 0.948891466 | 0.09405311 | 1 |
| Bves | 0.78778288 | -0.344130036 | 0.09409396 | 1 |
| Ddx10 | 1.111144 | 0.152045801 | 0.09413906 | 1 |
| Mfge8 | 1.10230395 | 0.140522092 | 0.09414812 | 1 |
| Iqsec3 | 0.89674091 | -0.157236873 | 0.09426874 | 1 |
| Mks1 | 1.23508906 | 0.30461507 | 0.09452082 | 1 |
| Dot1l | 0.87990663 | -0.184577658 | 0.09457757 | 1 |
| Xrn1 | 1.15261181 | 0.204906705 | 0.09457774 | 1 |
| Pou3f1 | 0.7782645 | -0.361667538 | 0.09460674 | 1 |
| Hdac4 | 0.85948487 | -0.218455859 | 0.0947964 | 1 |

| | | | | |
|---|---|---|---|---|
| Pcdha4 | 0.84976045 | -0.234871898 | 0.09494639 | 1 |
| Erg | 1.33579798 | 0.417701835 | 0.09529864 | 1 |
| Bcam | 0.92615663 | -0.110671896 | 0.09575208 | 1 |
| Hoxb8 | 1.320907 | 0.401528895 | 0.09581722 | 1 |
| Slc5a5 | 0.62318535 | -0.682266781 | 0.09585194 | 1 |
| Tesk1 | 0.88345654 | -0.178768937 | 0.09594441 | 1 |
| Tie1 | 1.13849305 | 0.187125481 | 0.09628102 | 1 |
| Kcna2 | 0.92368922 | -0.114520567 | 0.09630616 | 1 |
| Il7r | 0.86700501 | -0.20588776 | 0.09641562 | 1 |
| Lppos | 0.71375264 | -0.486503912 | 0.09675296 | 1 |
| Anp32e | 1.10859962 | 0.148738416 | 0.09683215 | 1 |
| Rps7 | 1.08023443 | 0.111344439 | 0.0968647 | 1 |
| Syt5 | 0.87914326 | -0.185829824 | 0.09692346 | 1 |
| Ndst4 | 1.10728769 | 0.147030106 | 0.0969604 | 1 |
| Cmtm4 | 0.82765972 | -0.27289034 | 0.09712343 | 1 |
| Bcl2l12 | 1.38280047 | 0.467592998 | 0.09723214 | 1 |
| Cdk2ap2 | 1.14576747 | 0.196314284 | 0.09739617 | 1 |
| Sec11c | 1.1108628 | 0.151680641 | 0.09772245 | 1 |
| Rc3h2 | 1.13458186 | 0.182160699 | 0.09776091 | 1 |
| Slc52a2 | 0.82654164 | -0.274840591 | 0.09801066 | 1 |
| Kcnc2 | 1.15431783 | 0.207040514 | 0.09822061 | 1 |
| Gal | 0.80887888 | -0.306004406 | 0.09823357 | 1 |
| Ntrk1 | 0.92515813 | -0.11222812 | 0.09838496 | 1 |
| Ptpn23 | 0.89657005 | -0.157511784 | 0.09844214 | 1 |
| Scand1 | 0.88407424 | -0.177760572 | 0.09849949 | 1 |
| Epha5 | 0.86744324 | -0.205158734 | 0.09886344 | 1 |
| Msmo1 | 0.92717045 | -0.10909351 | 0.09894539 | 1 |
| Tbc1d24 | 0.85229157 | -0.230581025 | 0.09905472 | 1 |
| Dynlt3 | 1.080485 | 0.111679046 | 0.09927638 | 1 |
| RGD1565355 | 1.54993062 | 0.632203636 | 0.09933439 | 1 |
| Polr3c | 1.12547913 | 0.170539307 | 0.09952809 | 1 |
| Ubxn1 | 1.09016772 | 0.124550107 | 0.09956728 | 1 |
| Ncln | 0.90449438 | -0.144816564 | 0.09973383 | 1 |
| Klrk1 | 0.68046941 | -0.555397794 | 0.09999834 | 1 |
| Lyl1 | 1.36974344 | 0.453905691 | 0.09999911 | 1 |
| Slc16a6 | 0.87271455 | -0.196418252 | 0.10002872 | 1 |
| Lrrc8b | 0.90038566 | -0.151385019 | 0.1001668 | 1 |
| Marcks | 0.91513256 | -0.127947355 | 0.10039802 | 1 |
| Fam45a | 1.09408531 | 0.12972524 | 0.10045546 | 1 |
| Slc44a1 | 0.92764631 | -0.108353244 | 0.10061666 | 1 |

| | | | | |
|---|---|---|---|---|
| Hirip3 | 1.1490712 | 0.200468198 | 0.10061875 | 1 |
| Scrg1 | 0.74262167 | -0.429300676 | 0.10062957 | 1 |
| Slc25a22 | 0.91468587 | -0.128651737 | 0.10063383 | 1 |
| Adm | 1.26555699 | 0.339772478 | 0.10064246 | 1 |
| Dclk1 | 0.92667016 | -0.109872186 | 0.10088211 | 1 |
| Ms4a6bl | 1.17785898 | 0.236166824 | 0.10111539 | 1 |
| Pde1c | 0.91271491 | -0.131763789 | 0.1011901 | 1 |
| Adam11 | 0.92351947 | -0.114785722 | 0.10131799 | 1 |
| Ak3 | 0.90600831 | -0.142403813 | 0.10146261 | 1 |
| Pdilt | 1.49683197 | 0.58191228 | 0.10149897 | 1 |
| Dnajc19 | 0.84010077 | -0.251365697 | 0.1019122 | 1 |
| Elovl6 | 0.91749698 | -0.124224682 | 0.10202924 | 1 |
| Ptprs | 0.92743557 | -0.108681035 | 0.10223082 | 1 |
| Zic1 | 1.24068182 | 0.31113317 | 0.1023251 | 1 |
| Mfhas1 | 0.89191627 | -0.165019821 | 0.10256484 | 1 |
| Gpsm1 | 0.88978161 | -0.168476814 | 0.10272472 | 1 |
| Bri3 | 0.8803609 | -0.18383302 | 0.10289618 | 1 |
| Efcab1 | 1.30807075 | 0.38744058 | 0.10290497 | 1 |
| Insig1 | 0.92733429 | -0.108838594 | 0.10293265 | 1 |
| P2rx3 | 0.92699604 | -0.109364924 | 0.10294873 | 1 |
| Tmem55a | 0.92095741 | -0.118793657 | 0.10298938 | 1 |
| Mup5 | 0.70077704 | -0.512972588 | 0.10301654 | 1 |
| Sez6l2 | 0.9253773 | -0.111886382 | 0.10313974 | 1 |
| Pggt1b | 1.15890622 | 0.212763822 | 0.10316572 | 1 |
| Cntln | 1.17928963 | 0.237918087 | 0.10327323 | 1 |
| Ehd1 | 0.92069511 | -0.119204609 | 0.10330048 | 1 |
| Camk2b | 0.90828582 | -0.138781735 | 0.10350519 | 1 |
| Serinc5 | 0.92076552 | -0.119094291 | 0.10366873 | 1 |
| Arhgef6 | 0.88778245 | -0.17172191 | 0.10378049 | 1 |
| Tbcb | 1.0871482 | 0.120548618 | 0.10393939 | 1 |
| Erich1 | 1.20910783 | 0.273942912 | 0.10395725 | 1 |
| Slc5a7 | 0.80625241 | -0.310696535 | 0.10402278 | 1 |
| Hoxd9 | 1.28886078 | 0.366096434 | 0.10403054 | 1 |
| Alg10 | 0.86960413 | -0.201569305 | 0.10404618 | 1 |
| Dctn1 | 0.92670286 | -0.109821265 | 0.10420456 | 1 |
| Hnrnpul2 | 0.92225398 | -0.116763986 | 0.10423213 | 1 |
| Piwil2 | 1.1574136 | 0.210904497 | 0.10423649 | 1 |
| Angpt2 | 1.17772222 | 0.235999308 | 0.10451513 | 1 |
| Fcmr | 1.23723602 | 0.307120738 | 0.10470763 | 1 |
| Ostc | 1.09445321 | 0.130210279 | 0.10490475 | 1 |

| | | | | |
|---|---|---|---|---|
| Cacna2d1 | 0.91321533 | -0.130973013 | 0.10527188 | 1 |
| Hmmr | 1.20969606 | 0.274644616 | 0.10539902 | 1 |
| Kcnq3 | 0.88620083 | -0.174294409 | 0.10544615 | 1 |
| Inafm1 | 1.20431207 | 0.268209286 | 0.1054531 | 1 |
| Gpr19 | 1.28606 | 0.362957956 | 0.10549182 | 1 |
| Camk2n2 | 0.81736535 | -0.290947016 | 0.10554364 | 1 |
| Cnr2 | 1.49573893 | 0.580858385 | 0.10582617 | 1 |
| Guk1 | 1.08388595 | 0.116212961 | 0.1059025 | 1 |
| Cdc25c | 1.23984601 | 0.310160944 | 0.10590887 | 1 |
| Slc16a13 | 1.2926323 | 0.37031195 | 0.10604955 | 1 |
| Gjb2 | 1.18932208 | 0.250139461 | 0.10615523 | 1 |
| Pvrl1 | 0.88388705 | -0.17806608 | 0.10629775 | 1 |
| Ugt1a6 | 1.19487231 | 0.256856457 | 0.10640875 | 1 |
| Nkd1 | 0.8356138 | -0.259091776 | 0.10643227 | 1 |
| Plcg2 | 1.289131 | 0.366398871 | 0.10648631 | 1 |
| Sema3b | 1.07680004 | 0.106750371 | 0.1065201 | 1 |
| Asrgl1 | 0.90405785 | -0.145513008 | 0.10664702 | 1 |
| Hip1 | 0.92006114 | -0.120198364 | 0.10665171 | 1 |
| Btg3 | 1.16235632 | 0.217052396 | 0.10669059 | 1 |
| Kcnj3 | 1.12482489 | 0.169700423 | 0.10672905 | 1 |
| Gtf3c6 | 1.20033962 | 0.263442649 | 0.10675649 | 1 |
| Mtss1l | 0.90425899 | -0.145192061 | 0.10691736 | 1 |
| Hcfc1r1 | 1.08949813 | 0.123663715 | 0.10726572 | 1 |
| Syn1 | 1.07920953 | 0.109974991 | 0.10734308 | 1 |
| Abtb2 | 0.90016205 | -0.151743358 | 0.10736745 | 1 |
| Sav1 | 0.8831175 | -0.179322689 | 0.1074153 | 1 |
| Rsl24d1 | 0.90468415 | -0.144513897 | 0.10756368 | 1 |
| Ankrd13c | 0.91843744 | -0.122746637 | 0.10768507 | 1 |
| Thoc3 | 1.13016367 | 0.17653172 | 0.10787976 | 1 |
| Tnfrsf11b | 1.25426053 | 0.326837052 | 0.10803135 | 1 |
| Cdpf1 | 0.77259316 | -0.372219195 | 0.10811489 | 1 |
| Ddt | 1.09960707 | 0.136988093 | 0.1083064 | 1 |
| Creld2 | 0.86214962 | -0.213989826 | 0.10849472 | 1 |
| Cnot3 | 1.14218336 | 0.191794275 | 0.10850148 | 1 |
| Gjb6 | 1.18929746 | 0.250109595 | 0.1085348 | 1 |
| Apobr | 1.38150966 | 0.466245648 | 0.10883691 | 1 |
| Icam1 | 1.19931598 | 0.262211815 | 0.10909373 | 1 |
| Mnda | 1.34900994 | 0.431900975 | 0.1091966 | 1 |
| Cope | 1.10658699 | 0.146116864 | 0.10926429 | 1 |
| Higd2a | 1.10383423 | 0.142523535 | 0.10932376 | 1 |

| | | | | |
|---|---|---|---|---|
| Klf16 | 0.84793694 | -0.237971119 | 0.1094404 | 1 |
| Rpl12 | 1.11278579 | 0.154175897 | 0.10998131 | 1 |
| Kcnn2 | 1.19062613 | 0.25172046 | 0.11005238 | 1 |
| Cyb5b | 0.91553812 | -0.127308138 | 0.11014662 | 1 |
| Nipsnap1 | 1.08169223 | 0.113290072 | 0.11024818 | 1 |
| Tatdn3 | 0.85064851 | -0.233364968 | 0.11029269 | 1 |
| Prkacb | 1.09275077 | 0.127964401 | 0.11043104 | 1 |
| Cdc42ep4 | 0.90521766 | -0.143663361 | 0.11048017 | 1 |
| Gcn1l1 | 0.91778972 | -0.123764451 | 0.11049408 | 1 |
| RGD1562136 | 1.08907656 | 0.12310538 | 0.11051709 | 1 |
| Eln | 1.13529872 | 0.183071947 | 0.1105845 | 1 |
| Igsf11 | 0.91117676 | -0.134197145 | 0.11065463 | 1 |
| LOC361016 | 1.24977233 | 0.321665308 | 0.11074254 | 1 |
| Acp5 | 0.78944943 | -0.34108124 | 0.11079725 | 1 |
| Vmp1 | 1.08228305 | 0.114077863 | 0.11089433 | 1 |
| Rbm7 | 1.14158025 | 0.191032285 | 0.11097433 | 1 |
| Wdfy3 | 0.92980619 | -0.104998071 | 0.11117428 | 1 |
| Arhgef1 | 0.91043321 | -0.135374912 | 0.11118218 | 1 |
| Zdhhc18 | 0.89159815 | -0.165534478 | 0.11136912 | 1 |
| Gas2l1 | 0.89320228 | -0.162941153 | 0.11141822 | 1 |
| Maf | 1.13353557 | 0.180829659 | 0.11144152 | 1 |
| Slc45a1 | 0.91415588 | -0.129487903 | 0.11149298 | 1 |
| Ednra | 0.86290943 | -0.212718955 | 0.11150621 | 1 |
| Creb3l1 | 1.16360277 | 0.218598639 | 0.11155712 | 1 |
| LOC361646 | 0.87609638 | -0.190838509 | 0.11161485 | 1 |
| Rasl11b | 0.84673867 | -0.240011323 | 0.11162336 | 1 |
| Luc7l3 | 1.07867038 | 0.109254069 | 0.11163466 | 1 |
| Gdpd1 | 1.12159952 | 0.165557642 | 0.11172662 | 1 |
| Tmem51 | 1.23044703 | 0.299182557 | 0.1118458 | 1 |
| Ppp1r9b | 0.9231474 | -0.115367065 | 0.1118664 | 1 |
| Akap10 | 0.89636202 | -0.157846575 | 0.11193854 | 1 |
| Pde9a | 0.78761974 | -0.344428821 | 0.11203568 | 1 |
| Tmem184c | 1.0767269 | 0.106652366 | 0.11213932 | 1 |
| Krt19 | 0.80338251 | -0.315841037 | 0.11214554 | 1 |
| Kptn | 1.3539156 | 0.437137808 | 0.11221854 | 1 |
| Spr | 0.85852762 | -0.220063553 | 0.11227229 | 1 |
| Tlr7 | 1.21197374 | 0.277358446 | 0.11233168 | 1 |
| Cacnb1 | 0.88764713 | -0.171941817 | 0.11239772 | 1 |
| Pten | 0.91281285 | -0.131608989 | 0.11242815 | 1 |
| RGD1564053 | 1.13534067 | 0.183125253 | 0.11248691 | 1 |

| | | | | |
|---|---|---|---|---|
| Tsta3 | 1.12716521 | 0.172698991 | 0.11256115 | 1 |
| Gbp5 | 1.20185077 | 0.265257767 | 0.11286443 | 1 |
| Rabl3 | 1.14370214 | 0.193711373 | 0.11293121 | 1 |
| Adgre4 | 1.82970604 | 0.871611887 | 0.11299481 | 1 |
| Rnf139 | 1.12107815 | 0.164886851 | 0.11319253 | 1 |
| Zbtb16 | 0.82313303 | -0.280802484 | 0.11325697 | 1 |
| Marcksl1 | 0.91802257 | -0.123398479 | 0.11333608 | 1 |
| Cd5 | 0.63802871 | -0.648306752 | 0.11337123 | 1 |
| Qk | 0.90430089 | -0.145125217 | 0.11377233 | 1 |
| Ppic | 0.87994791 | -0.184509974 | 0.11382491 | 1 |
| Xiap | 0.89702566 | -0.156778839 | 0.11384347 | 1 |
| Rhobtb2 | 0.89822899 | -0.154844817 | 0.11385889 | 1 |
| Kcnj6 | 0.8737239 | -0.194750633 | 0.11396572 | 1 |
| Htr1b | 0.81910351 | -0.287882324 | 0.11422414 | 1 |
| S1pr1 | 0.86869689 | -0.20307522 | 0.11422944 | 1 |
| Irf3 | 1.09872624 | 0.135831964 | 0.11425064 | 1 |
| Des | 1.23449172 | 0.303917159 | 0.11431777 | 1 |
| Stt3b | 0.91428081 | -0.129290757 | 0.11434759 | 1 |
| Tmem200a | 0.85459309 | -0.226690441 | 0.1146424 | 1 |
| Mbtps2 | 1.217746 | 0.284213243 | 0.11473339 | 1 |
| Mgrn1 | 0.91533862 | -0.127622544 | 0.11482531 | 1 |
| Cacna1e | 0.74662478 | -0.421544694 | 0.11490748 | 1 |
| Tpm1 | 1.07521519 | 0.104625422 | 0.11492516 | 1 |
| Prdx2 | 1.07591619 | 0.105565701 | 0.1150502 | 1 |
| Fam219a | 0.91493708 | -0.128255562 | 0.11531804 | 1 |
| Fam129b | 0.91559582 | -0.127217212 | 0.1153744 | 1 |
| Gstm4 | 1.11425052 | 0.156073628 | 0.11538281 | 1 |
| Fras1 | 0.87973285 | -0.184862612 | 0.11547991 | 1 |
| Nectin4 | 1.35009718 | 0.433063252 | 0.11569261 | 1 |
| Cab39l | 1.10128114 | 0.139182819 | 0.11571722 | 1 |
| Gstm6 | 1.28219374 | 0.358614268 | 0.11577559 | 1 |
| Enpp3 | 0.88712048 | -0.17279805 | 0.11588703 | 1 |
| Usp11 | 1.09794576 | 0.13480678 | 0.11598681 | 1 |
| Lpar6 | 1.19462975 | 0.256563553 | 0.11601623 | 1 |
| Notch1 | 0.9175648 | -0.124118051 | 0.11607116 | 1 |
| Galnt2 | 0.91349451 | -0.130532038 | 0.1162229 | 1 |
| B4galt5 | 0.89376891 | -0.162026233 | 0.11656082 | 1 |
| Ing1 | 1.18121494 | 0.240271504 | 0.11681269 | 1 |
| Slc37a1 | 0.88833504 | -0.170824202 | 0.11686005 | 1 |
| Tln1 | 0.92998057 | -0.104727516 | 0.11693695 | 1 |

| | | | | |
|---|---|---|---|---|
| Clic2 | 1.15904626 | 0.212938146 | 0.11699375 | 1 |
| RGD1559896 | 0.91361854 | -0.130336175 | 0.11703514 | 1 |
| Grasp | 1.30821185 | 0.387596189 | 0.11718908 | 1 |
| Ranbp17 | 0.82335498 | -0.280413523 | 0.11740616 | 1 |
| Tox2 | 0.87070657 | -0.199741479 | 0.11751595 | 1 |
| Fam46a | 1.34594189 | 0.428616126 | 0.11797784 | 1 |
| Ap2a1 | 0.92790094 | -0.107957292 | 0.11820888 | 1 |
| Vwf | 1.09536122 | 0.131406705 | 0.1183163 | 1 |
| Hbp1 | 1.09369845 | 0.129215022 | 0.11837812 | 1 |
| Ero1b | 0.83532657 | -0.259587767 | 0.11845414 | 1 |
| Mustn1 | 1.20686852 | 0.271268508 | 0.11860236 | 1 |
| Mak16 | 1.13157749 | 0.17833538 | 0.11868016 | 1 |
| Cd274 | 1.20606779 | 0.270311006 | 0.11868497 | 1 |
| Reln | 0.92967881 | -0.105195728 | 0.11905582 | 1 |
| Igfbp5 | 1.07540166 | 0.104875599 | 0.11912601 | 1 |
| Nap1l5 | 1.07508399 | 0.10444938 | 0.11927783 | 1 |
| Tph2 | 0.87401561 | -0.194269053 | 0.11939157 | 1 |
| Neurl4 | 0.91222281 | -0.132541852 | 0.11946845 | 1 |
| Scg5 | 1.07457217 | 0.103762384 | 0.11959439 | 1 |
| Mkks | 1.09638775 | 0.132758117 | 0.11974098 | 1 |
| Fam180a | 1.15115868 | 0.203086718 | 0.11976916 | 1 |
| Nhlrc2 | 0.87161653 | -0.198234546 | 0.11981843 | 1 |
| Nrep | 0.84162886 | -0.248743918 | 0.12003564 | 1 |
| Gimap9 | 1.35436525 | 0.437616866 | 0.12004456 | 1 |
| Glyr1 | 0.91067708 | -0.134988513 | 0.12004544 | 1 |
| Cd27 | 0.66614861 | -0.586084038 | 0.12016343 | 1 |
| Tagln3 | 1.0785905 | 0.109147237 | 0.12018393 | 1 |
| Extl2 | 1.08665524 | 0.11989429 | 0.12030614 | 1 |
| Cirbp | 1.11570716 | 0.157958415 | 0.12032832 | 1 |
| Dgat2 | 0.87283641 | -0.196216813 | 0.1204072 | 1 |
| Tbc1d19 | 1.09295332 | 0.128231786 | 0.12108005 | 1 |
| Bet1l | 1.13708392 | 0.185338731 | 0.12121028 | 1 |
| Arl11 | 1.33141442 | 0.412959702 | 0.1212351 | 1 |
| LOC365985 | 0.92342043 | -0.114940439 | 0.12128405 | 1 |
| Pappa | 0.78904383 | -0.341822661 | 0.1214355 | 1 |
| Secisbp2l | 0.93062408 | -0.103729575 | 0.12160792 | 1 |
| Anks1b | 0.89200189 | -0.164881322 | 0.12174652 | 1 |
| Slc12a1 | 0.7446777 | -0.425311938 | 0.12180535 | 1 |
| Poc5 | 1.16838762 | 0.224518971 | 0.12185116 | 1 |
| Ctsg | 0.60634978 | -0.721777815 | 0.12194708 | 1 |

| | | | | |
|---|---|---|---|---|
| Gas6 | 1.07363265 | 0.102500456 | 0.12199719 | 1 |
| Ubqln2 | 0.92229335 | -0.116702404 | 0.12219356 | 1 |
| Pabpn1 | 0.92436609 | -0.113463752 | 0.12222819 | 1 |
| LOC100302372 | 0.80116108 | -0.319835763 | 0.12223452 | 1 |
| Apobec1 | 1.36105828 | 0.444728844 | 0.12227759 | 1 |
| Cops3 | 1.08547572 | 0.118327455 | 0.12256093 | 1 |
| Chgb | 1.07351973 | 0.1023487 | 0.12266922 | 1 |
| Npy5r | 0.77897997 | -0.360341855 | 0.12309148 | 1 |
| Unc13a | 0.9141393 | -0.129514077 | 0.12309776 | 1 |
| Rhbdl3 | 1.14422113 | 0.194365892 | 0.12311776 | 1 |
| Naalad2 | 1.5172225 | 0.601432669 | 0.12311835 | 1 |
| Cubn | 1.20169506 | 0.265070843 | 0.12331423 | 1 |
| Rab31 | 0.92035185 | -0.11974258 | 0.12347354 | 1 |
| Add2 | 0.92795206 | -0.10787782 | 0.12348918 | 1 |
| Shox2 | 1.13168515 | 0.178472638 | 0.12350724 | 1 |
| Tmem220 | 1.29917303 | 0.377593592 | 0.12359621 | 1 |
| Abcg3l3 | 1.13598299 | 0.18394123 | 0.12368119 | 1 |
| Cog4 | 1.07935404 | 0.110168162 | 0.12369825 | 1 |
| Mrps7 | 1.10354733 | 0.142148506 | 0.12377812 | 1 |
| Prpf31 | 1.10718747 | 0.146899519 | 0.12385741 | 1 |
| Sema4g | 0.87607336 | -0.190876411 | 0.12404691 | 1 |
| Atp6v1f | 1.07738612 | 0.107535386 | 0.12412463 | 1 |
| Nadsyn1 | 1.16557578 | 0.221042807 | 0.12412677 | 1 |
| Taf13 | 1.08397905 | 0.116336881 | 0.12420382 | 1 |
| Sdf4 | 1.07553491 | 0.10505435 | 0.1242911 | 1 |
| Fbxl12 | 1.28283201 | 0.359332256 | 0.1243077 | 1 |
| Slco2b1 | 1.12659668 | 0.171971119 | 0.12444887 | 1 |
| Polr2c | 1.12169802 | 0.165684333 | 0.12454156 | 1 |
| Adamts2 | 0.88966326 | -0.168668724 | 0.1245469 | 1 |
| Lrrc51 | 0.74762681 | -0.419609788 | 0.12457599 | 1 |
| Sdr39u1 | 1.13109515 | 0.1777203 | 0.12481322 | 1 |
| Thbs1 | 0.90456099 | -0.144710315 | 0.12481451 | 1 |
| Dcaf12 | 0.86636687 | -0.206950027 | 0.12484047 | 1 |
| Tppp3 | 1.07628633 | 0.106061934 | 0.12502066 | 1 |
| Arvcf | 0.91408484 | -0.129600022 | 0.12511504 | 1 |
| Abcb1 | 1.10026021 | 0.137844763 | 0.12518289 | 1 |
| Clic5 | 0.83947595 | -0.252439095 | 0.12521135 | 1 |
| Cdca3 | 0.63529901 | -0.654492327 | 0.12550163 | 1 |
| Wdr13 | 1.1237215 | 0.168284522 | 0.12556445 | 1 |
| Cpt1b | 0.76415466 | -0.388063431 | 0.12580775 | 1 |

| | | | | |
|---|---|---|---|---|
| Lamc1 | 0.93223021 | -0.101241829 | 0.12583104 | 1 |
| Henmt1 | 1.31002749 | 0.389597085 | 0.12583395 | 1 |
| Pde2a | 0.91938566 | -0.121257931 | 0.12588891 | 1 |
| Nuf2 | 0.69398412 | -0.527025454 | 0.12601597 | 1 |
| Fgd4 | 1.12334114 | 0.167796122 | 0.12620858 | 1 |
| Dusp15 | 0.92741111 | -0.108719088 | 0.1262573 | 1 |
| RGD1309779 | 1.27540397 | 0.350954274 | 0.12661823 | 1 |
| Hpca | 0.8415871 | -0.248815504 | 0.12669298 | 1 |
| Nat8f5 | 0.72751884 | -0.458943492 | 0.12674608 | 1 |
| Fez1 | 1.0734997 | 0.102321788 | 0.12689017 | 1 |
| Usp20 | 0.9312624 | -0.102740366 | 0.12742196 | 1 |
| Kctd4 | 0.80820591 | -0.307205187 | 0.12761561 | 1 |
| Sncaip | 0.81495744 | -0.295203381 | 0.12765626 | 1 |
| Rnf168 | 1.11416981 | 0.155969126 | 0.12774557 | 1 |
| Azin1 | 0.93216104 | -0.101348875 | 0.1277945 | 1 |
| Zmat3 | 0.92224129 | -0.116783838 | 0.12779816 | 1 |
| Fes | 1.2985865 | 0.37694212 | 0.12795675 | 1 |
| B3gnt7 | 0.7092079 | -0.495719496 | 0.12796954 | 1 |
| Enkd1 | 1.35762219 | 0.441082053 | 0.12798712 | 1 |
| Cpne2 | 0.88055919 | -0.183508111 | 0.12807164 | 1 |
| Amigo3 | 0.90399998 | -0.145605353 | 0.1280967 | 1 |
| Bloc1s5 | 1.09389656 | 0.129476321 | 0.12811613 | 1 |
| Ppp1r3c | 1.13986499 | 0.188862954 | 0.12814898 | 1 |
| Mtmr4 | 0.93229803 | -0.101136879 | 0.12839042 | 1 |
| Rbm22 | 0.87381288 | -0.194603717 | 0.12852987 | 1 |
| Aldh16a1 | 1.14687814 | 0.197712106 | 0.12853214 | 1 |
| Naa20 | 1.09255687 | 0.127708379 | 0.1286229 | 1 |
| Rab27a | 1.1097141 | 0.150188033 | 0.12866239 | 1 |
| Nup107 | 1.11732581 | 0.160049935 | 0.12866879 | 1 |
| Mylip | 1.12396446 | 0.168596412 | 0.12871311 | 1 |
| Hus1 | 1.11872478 | 0.161855154 | 0.12884605 | 1 |
| Prr3 | 0.89041919 | -0.167443405 | 0.1290895 | 1 |
| Hormad2 | 1.47983433 | 0.565435668 | 0.12909997 | 1 |
| Hspb7 | 1.4665264 | 0.55240304 | 0.12922988 | 1 |
| Prrx2 | 0.77702218 | -0.363972321 | 0.12930436 | 1 |
| Selm | 1.07601563 | 0.105699029 | 0.1293746 | 1 |
| Car3 | 1.33698357 | 0.418981735 | 0.12941823 | 1 |
| Dclre1c | 1.22566347 | 0.293562909 | 0.12960649 | 1 |
| Hn1l | 0.88656031 | -0.173709325 | 0.12966217 | 1 |
| Lrfn1 | 0.87176826 | -0.197983425 | 0.12992451 | 1 |

| | | | | |
|---|---|---|---|---|
| Hbegf | 1.14680904 | 0.197625184 | 0.12995109 | 1 |
| Rwdd3 | 0.80871442 | -0.306297764 | 0.1300671 | 1 |
| Tmem255b | 1.30460104 | 0.383608687 | 0.13041747 | 1 |
| Pcdhga1 | 1.25118525 | 0.323295413 | 0.13049732 | 1 |
| Mgp | 0.93279281 | -0.100371427 | 0.13065025 | 1 |
| LOC500959 | 1.08377067 | 0.116059507 | 0.13068222 | 1 |
| Iscu | 1.07168278 | 0.099877929 | 0.1307397 | 1 |
| Cdh11 | 0.93102983 | -0.1031007 | 0.13088661 | 1 |
| Mab21l3 | 0.79023685 | -0.339642973 | 0.13090632 | 1 |
| S100a8 | 1.13528876 | 0.183059288 | 0.13091946 | 1 |
| Aurkb | 1.54100731 | 0.623873709 | 0.13094671 | 1 |
| MGC108823 | 1.23794984 | 0.307952858 | 0.13102106 | 1 |
| Dnaaf5 | 0.8391971 | -0.252918395 | 0.13103446 | 1 |
| Nosip | 1.11361739 | 0.155253651 | 0.13122678 | 1 |
| Utp4 | 0.89309758 | -0.163110287 | 0.13139815 | 1 |
| Flna | 1.07157709 | 0.099735645 | 0.13144816 | 1 |
| Dhx58 | 1.29224692 | 0.369881764 | 0.13150254 | 1 |
| Ifit2 | 1.14559407 | 0.196095933 | 0.13159121 | 1 |
| Cdk13 | 0.90629166 | -0.141952692 | 0.13179616 | 1 |
| Urb2 | 0.84425873 | -0.24424291 | 0.13198214 | 1 |
| Sema3e | 0.87464332 | -0.193233292 | 0.1320871 | 1 |
| Hdc | 1.48810674 | 0.573478014 | 0.13220431 | 1 |
| Prnp | 1.07473445 | 0.103980232 | 0.13220642 | 1 |
| Ngly1 | 1.0840834 | 0.116475749 | 0.13226461 | 1 |
| Dcakd | 0.89740408 | -0.156170344 | 0.13230794 | 1 |
| Mettl13 | 0.88420124 | -0.177553344 | 0.13241036 | 1 |
| Pitpnm2 | 0.93356744 | -0.099173851 | 0.13241113 | 1 |
| Zfp202 | 1.218537 | 0.285150056 | 0.13257946 | 1 |
| Ell3 | 0.54823558 | -0.867132141 | 0.13260524 | 1 |
| Ptprf | 0.93181262 | -0.101888218 | 0.13273608 | 1 |
| Cyb561 | 0.87807267 | -0.187587751 | 0.13281107 | 1 |
| Rnd1 | 0.81693864 | -0.291700373 | 0.13291241 | 1 |
| Anapc7 | 0.89359523 | -0.162306609 | 0.13299105 | 1 |
| Tgif1 | 1.29265237 | 0.370334351 | 0.13302704 | 1 |
| Tmem223 | 1.10404437 | 0.142798159 | 0.13328054 | 1 |
| Qtrt1 | 1.29074693 | 0.368206163 | 0.13331069 | 1 |
| Ier5l | 0.79608343 | -0.329008452 | 0.13341052 | 1 |
| Erc1 | 0.89093909 | -0.166601286 | 0.13352004 | 1 |
| Add3 | 0.93378504 | -0.098837614 | 0.13373399 | 1 |
| Traf3ip2 | 0.85836996 | -0.220328516 | 0.13378067 | 1 |

| | | | | |
|---|---|---|---|---|
| Casc1 | 1.15400036 | 0.206643678 | 0.13381417 | 1 |
| Psmd13 | 1.07582259 | 0.105440182 | 0.13399344 | 1 |
| Rgcc | 0.87963614 | -0.185021217 | 0.13402337 | 1 |
| Hivep2 | 0.92426874 | -0.113615708 | 0.13426229 | 1 |
| RGD1359508 | 1.31260747 | 0.392435545 | 0.13445867 | 1 |
| Chrnb2 | 0.92160467 | -0.117780061 | 0.1345289 | 1 |
| Slc15a4 | 0.89383679 | -0.161916663 | 0.13453776 | 1 |
| Maob | 0.92343361 | -0.11491985 | 0.13470596 | 1 |
| Med23 | 1.10111278 | 0.138962241 | 0.13490801 | 1 |
| Slc12a7 | 0.91140735 | -0.133832092 | 0.13498143 | 1 |
| Plxnb2 | 1.08704026 | 0.120405371 | 0.13524297 | 1 |
| Phex | 1.33679381 | 0.418776962 | 0.13535254 | 1 |
| Syndig1 | 1.16974744 | 0.226197072 | 0.13538574 | 1 |
| Ccdc32 | 0.91677591 | -0.125358956 | 0.13541641 | 1 |
| Trub2 | 0.88267744 | -0.180041774 | 0.13542308 | 1 |
| Zcchc2 | 0.88234096 | -0.18059183 | 0.13561349 | 1 |
| Slc25a42 | 0.88402844 | -0.177835318 | 0.13587096 | 1 |
| Gpr183 | 1.34069875 | 0.422985104 | 0.13587305 | 1 |
| Tfap4 | 1.2807056 | 0.356938875 | 0.13593244 | 1 |
| Fbxo27 | 0.83197523 | -0.265387519 | 0.13609385 | 1 |
| Lamtor2 | 1.10179836 | 0.139860218 | 0.1361036 | 1 |
| Atg12 | 1.07908978 | 0.109814895 | 0.13614065 | 1 |
| Nts | 1.39154008 | 0.476682463 | 0.13628203 | 1 |
| Dpt | 1.082509 | 0.114379022 | 0.1362896 | 1 |
| LOC500956 | 1.13361994 | 0.18093704 | 0.1365042 | 1 |
| Lrrcc1 | 1.1514498 | 0.203451521 | 0.13652105 | 1 |
| Pdgfrl | 0.86042039 | -0.216886373 | 0.13665292 | 1 |
| Zgpat | 1.1370296 | 0.185269812 | 0.13672186 | 1 |
| Prrt3 | 0.90385671 | -0.145834022 | 0.13682395 | 1 |
| Ppp1r15b | 0.91121452 | -0.134137359 | 0.1369092 | 1 |
| Rrp9 | 0.85312199 | -0.229176038 | 0.13713438 | 1 |
| F2rl1 | 0.73229851 | -0.449496239 | 0.13719445 | 1 |
| Agtr1b | 1.48770279 | 0.573086336 | 0.13765203 | 1 |
| Coro2a | 0.89901779 | -0.153578427 | 0.13770059 | 1 |
| Nsmaf | 1.12003224 | 0.16354026 | 0.13789606 | 1 |
| Slc43a1 | 1.31948011 | 0.399969603 | 0.13792277 | 1 |
| Sla | 1.34118961 | 0.423513211 | 0.1380831 | 1 |
| Trpv1 | 0.92787749 | -0.107993761 | 0.13819838 | 1 |
| Atox1 | 0.90460922 | -0.144633389 | 0.13834775 | 1 |
| Prx | 0.93252575 | -0.100784537 | 0.13844513 | 1 |

| | | | | |
|---|---|---|---|---|
| Mrps18a | 1.09406629 | 0.129700159 | 0.13859248 | 1 |
| Hopx | 1.07472369 | 0.10396579 | 0.13865474 | 1 |
| Ctbp2 | 0.87534525 | -0.19207595 | 0.13880812 | 1 |
| Chek1 | 0.80265286 | -0.317151927 | 0.13883508 | 1 |
| Foxo4 | 1.11991231 | 0.163385778 | 0.13891064 | 1 |
| Bclaf1 | 1.0753616 | 0.104821855 | 0.13935123 | 1 |
| Pdzd8 | 0.93475271 | -0.097343343 | 0.13959243 | 1 |
| Aldoc | 1.07173359 | 0.099946328 | 0.13977238 | 1 |
| Polr2m | 0.93105743 | -0.103057939 | 0.13994895 | 1 |
| Acvr2a | 0.88240897 | -0.180480646 | 0.14004097 | 1 |
| Gpr162 | 0.89377811 | -0.162011389 | 0.14024902 | 1 |
| Lrrc75b | 0.84598642 | -0.241293591 | 0.14025722 | 1 |
| Cep85l | 0.84943186 | -0.235429865 | 0.14051451 | 1 |
| Ndp | 1.13792123 | 0.186400693 | 0.14051959 | 1 |
| Hipk3 | 0.91679525 | -0.125328533 | 0.14074727 | 1 |
| Fam195a | 1.34985678 | 0.432806347 | 0.14074892 | 1 |
| Ankra2 | 0.88387125 | -0.178091854 | 0.14079089 | 1 |
| Reck | 1.08186854 | 0.11352521 | 0.14085164 | 1 |
| Cnksr3 | 1.20876289 | 0.273531276 | 0.1408672 | 1 |
| Abl1 | 0.91740573 | -0.124368178 | 0.14128372 | 1 |
| Tmem151b | 0.86215774 | -0.213976249 | 0.14128881 | 1 |
| Hyal2 | 0.88616514 | -0.174352518 | 0.14157305 | 1 |
| Frs3 | 1.16368816 | 0.218704505 | 0.14159897 | 1 |
| Bin3 | 1.11757886 | 0.160376642 | 0.14169627 | 1 |
| Tsr2 | 1.07780348 | 0.108094144 | 0.1417511 | 1 |
| RGD735029 | 1.09012768 | 0.124497119 | 0.14176693 | 1 |
| Tmem151a | 0.92551983 | -0.111664188 | 0.14182013 | 1 |
| Mon1a | 0.88961202 | -0.168751818 | 0.14183111 | 1 |
| Ttc9c | 1.0964917 | 0.132894894 | 0.14192652 | 1 |
| Pcdha10 | 1.20231794 | 0.265818453 | 0.14219621 | 1 |
| Elk4 | 0.91381106 | -0.130032198 | 0.14236889 | 1 |
| Srd5a1 | 0.8822093 | -0.180807127 | 0.14262949 | 1 |
| Sdhaf3 | 0.76407237 | -0.388218804 | 0.14288458 | 1 |
| Trappc4 | 1.08648524 | 0.119668572 | 0.14292643 | 1 |
| Alg14 | 1.10918312 | 0.149497567 | 0.14293074 | 1 |
| Dnajc17 | 1.23908277 | 0.309272567 | 0.14312257 | 1 |
| Mettl3 | 1.09792248 | 0.134776198 | 0.1432919 | 1 |
| Tlk2 | 1.09288604 | 0.128142975 | 0.1433344 | 1 |
| Pck2 | 0.89824893 | -0.154812783 | 0.1433503 | 1 |
| Ccpg1os | 1.34211567 | 0.424509013 | 0.14350186 | 1 |

| | | | | |
|---|---|---|---|---|
| Olfm2 | 0.85340563 | -0.228696463 | 0.14356639 | 1 |
| Samd4b | 0.90328118 | -0.14675295 | 0.1435827 | 1 |
| Plpp3 | 1.07220511 | 0.100580917 | 0.14367922 | 1 |
| Phospho2 | 1.11543591 | 0.157607625 | 0.14373092 | 1 |
| Gla | 1.08650975 | 0.119701123 | 0.1437438 | 1 |
| Igsf9 | 1.31395472 | 0.393915557 | 0.14400609 | 1 |
| Fam57b | 1.06960323 | 0.097075725 | 0.14413052 | 1 |
| Kif20b | 1.28024544 | 0.356420421 | 0.14463382 | 1 |
| Snrpb2 | 1.1028157 | 0.14119171 | 0.14493838 | 1 |
| Ddx31 | 1.17276821 | 0.229917902 | 0.14499413 | 1 |
| Tcf4 | 1.14046623 | 0.189623723 | 0.14500373 | 1 |
| Psmc5 | 1.06952995 | 0.096976887 | 0.14506161 | 1 |
| Hk2 | 1.21929442 | 0.286046528 | 0.14513151 | 1 |
| Arhgap12 | 1.08186129 | 0.113515539 | 0.14513568 | 1 |
| Fzd6 | 1.24742306 | 0.318950836 | 0.14513984 | 1 |
| Cyp1b1 | 1.18077482 | 0.239733861 | 0.14523175 | 1 |
| Ybx1-ps3 | 0.93298675 | -0.100071503 | 0.14525677 | 1 |
| Trmt6 | 1.11968272 | 0.163089982 | 0.14532473 | 1 |
| Parn | 1.12680767 | 0.172241291 | 0.1453746 | 1 |
| Tek | 1.11130375 | 0.152253196 | 0.14539631 | 1 |
| Fkbp3 | 1.0694184 | 0.096826409 | 0.14556069 | 1 |
| Cbwd1 | 1.19089112 | 0.252041512 | 0.14564803 | 1 |
| Chmp3 | 1.07401037 | 0.103007926 | 0.14567019 | 1 |
| Gfra1 | 0.93437516 | -0.097926176 | 0.1458783 | 1 |
| Timp4 | 1.19030381 | 0.251329847 | 0.14596273 | 1 |
| Mrpl40 | 1.12655043 | 0.171911902 | 0.14600978 | 1 |
| Ybx3 | 1.07568003 | 0.105248999 | 0.1460325 | 1 |
| Zfp703 | 0.8663385 | -0.206997258 | 0.14629433 | 1 |
| Mex3b | 0.81016999 | -0.30370344 | 0.1463777 | 1 |
| Clptm1 | 0.9337626 | -0.098872296 | 0.14639682 | 1 |
| Aldh1a3 | 1.1741666 | 0.23163712 | 0.14642078 | 1 |
| Mtf1 | 0.88327746 | -0.179061396 | 0.14651396 | 1 |
| Nmnat1 | 0.847192 | -0.239239131 | 0.14657207 | 1 |
| Tuba1c | 0.92855477 | -0.106941081 | 0.14659499 | 1 |
| Kcnt1 | 0.91798495 | -0.12345759 | 0.1466194 | 1 |
| Grin3a | 1.26991173 | 0.344728225 | 0.14674759 | 1 |
| Ncor1 | 0.93085719 | -0.103368238 | 0.14693713 | 1 |
| Gm2a | 1.08620432 | 0.119295511 | 0.14694062 | 1 |
| Fkrp | 0.89779895 | -0.155535693 | 0.14702096 | 1 |
| Hyou1 | 0.93318797 | -0.09976038 | 0.14704113 | 1 |

| | | | | |
|---|---|---|---|---|
| Wdr45 | 1.12398987 | 0.168629036 | 0.14713287 | 1 |
| Ctxn3 | 0.90821941 | -0.138887232 | 0.14731593 | 1 |
| Plcd3 | 1.15344128 | 0.205944564 | 0.14751108 | 1 |
| Recql | 1.09892117 | 0.136087896 | 0.14757944 | 1 |
| Mcc | 1.17993894 | 0.238712201 | 0.14768512 | 1 |
| Il1r1 | 1.22568078 | 0.293583283 | 0.14785814 | 1 |
| Actn4 | 0.93603731 | -0.095362052 | 0.14790148 | 1 |
| Ankrd34c | 1.35054215 | 0.433538668 | 0.14790283 | 1 |
| Stom | 1.13053967 | 0.17701162 | 0.14806054 | 1 |
| Anapc10 | 1.17654937 | 0.234561856 | 0.14827068 | 1 |
| Ing3 | 1.13814334 | 0.186682266 | 0.14832235 | 1 |
| Nectin3 | 0.87265825 | -0.196511319 | 0.14833741 | 1 |
| Plk5 | 0.71227363 | -0.489496511 | 0.14839524 | 1 |
| Crh | 0.67931205 | -0.557853659 | 0.14841328 | 1 |
| Surf2 | 1.10690974 | 0.146537591 | 0.14850636 | 1 |
| Nxnl2 | 1.39066018 | 0.475769923 | 0.14855854 | 1 |
| Clu | 1.07221099 | 0.100588828 | 0.14871144 | 1 |
| Bud31 | 1.0972134 | 0.133844151 | 0.1487774 | 1 |
| Fkbp4 | 0.93510991 | -0.096792152 | 0.14894969 | 1 |
| Zbtb7b | 0.83492665 | -0.260278635 | 0.14901858 | 1 |
| Safb | 0.92209201 | -0.117017377 | 0.1490767 | 1 |
| Lactb | 1.14206822 | 0.191648836 | 0.14917612 | 1 |
| Cyp46a1 | 0.78670532 | -0.346104758 | 0.14930684 | 1 |
| Ccdc90b | 1.12294011 | 0.167280985 | 0.14936639 | 1 |
| Car11 | 0.90438398 | -0.144992653 | 0.14939539 | 1 |
| Gnl3 | 1.10274841 | 0.141103678 | 0.14951299 | 1 |
| Mfap3 | 0.84771718 | -0.238345077 | 0.14954684 | 1 |
| Gtf3c1 | 0.93447181 | -0.097776951 | 0.1497419 | 1 |
| Sbf1 | 0.93018398 | -0.104412001 | 0.14979509 | 1 |
| Rpl38 | 1.07149936 | 0.099630992 | 0.14982184 | 1 |
| Dhh | 0.92060323 | -0.119348596 | 0.15003516 | 1 |
| Ifi44 | 1.10352536 | 0.142119785 | 0.15006412 | 1 |
| Myh7b | 1.17816971 | 0.236547364 | 0.15009401 | 1 |
| Cnnm1 | 0.90669127 | -0.141316699 | 0.15010612 | 1 |
| Hmgn5 | 1.24135156 | 0.311911756 | 0.15056643 | 1 |
| Rgs18 | 1.30865773 | 0.388087822 | 0.15062976 | 1 |
| Prpf4 | 0.88809385 | -0.17121596 | 0.1506733 | 1 |
| Skp1 | 1.07133244 | 0.099406226 | 0.15067989 | 1 |
| Pxdc1 | 0.88451644 | -0.177039133 | 0.15081136 | 1 |
| Adora1 | 0.88783241 | -0.171640717 | 0.15100073 | 1 |

| | | | | |
|---|---|---|---|---|
| Atp2b3 | 0.92627065 | -0.110494294 | 0.15109929 | 1 |
| Asb1 | 0.89611069 | -0.158251153 | 0.1512226 | 1 |
| Gjc1 | 0.75618733 | -0.403184415 | 0.15126385 | 1 |
| Lhfpl5 | 1.10688539 | 0.146505846 | 0.15148657 | 1 |
| Trim9 | 0.88409984 | -0.177718789 | 0.15150445 | 1 |
| Polr1b | 0.88554886 | -0.175356193 | 0.15157307 | 1 |
| Plaur | 1.16869112 | 0.224893685 | 0.15181957 | 1 |
| Adamts9 | 0.87373664 | -0.194729599 | 0.15183116 | 1 |
| Mmp15 | 0.93308911 | -0.099913224 | 0.15189778 | 1 |
| Sptlc2 | 0.92851024 | -0.107010275 | 0.15190538 | 1 |
| Ppm1d | 1.15862254 | 0.212410631 | 0.15207273 | 1 |
| Cdk2 | 0.79726661 | -0.32686585 | 0.15213512 | 1 |
| Tpm3 | 0.9327137 | -0.100493788 | 0.15221794 | 1 |
| Tnnt2 | 1.18776893 | 0.248254196 | 0.15239509 | 1 |
| Slc38a1 | 1.09759606 | 0.134347202 | 0.15277081 | 1 |
| Tacc2 | 0.92631602 | -0.110423623 | 0.1528048 | 1 |
| Rnf138 | 0.8697318 | -0.201357504 | 0.15302475 | 1 |
| Pcdhb6 | 1.44755028 | 0.533613461 | 0.15328097 | 1 |
| Gsdmd | 1.22607515 | 0.294047407 | 0.15394367 | 1 |
| Fmo4 | 1.25880734 | 0.332057493 | 0.15455744 | 1 |
| RT1-CE14 | 1.15573046 | 0.208804973 | 0.15456353 | 1 |
| Plxna3 | 0.92192523 | -0.117278339 | 0.15459105 | 1 |
| Apol9a | 1.25540644 | 0.328154513 | 0.15461553 | 1 |
| Slamf9 | 1.31728732 | 0.397570049 | 0.15469508 | 1 |
| Kdelr3 | 0.90117846 | -0.150115269 | 0.15485316 | 1 |
| St14 | 1.3197426 | 0.400256572 | 0.1550818 | 1 |
| Kctd13 | 0.86547387 | -0.208437826 | 0.15513229 | 1 |
| Apba1 | 0.93624084 | -0.095048398 | 0.15517077 | 1 |
| Oxr1 | 1.06884799 | 0.096056695 | 0.15525549 | 1 |
| Cenpk | 1.33299757 | 0.414674154 | 0.15569815 | 1 |
| Aida | 1.07419695 | 0.103258532 | 0.15574538 | 1 |
| Ahsa1 | 1.06777974 | 0.094614079 | 0.15581296 | 1 |
| Naf1 | 1.22304122 | 0.290473023 | 0.15594039 | 1 |
| Npm1 | 1.07176759 | 0.099992096 | 0.15604145 | 1 |
| Ptpn6 | 1.16935526 | 0.225713299 | 0.1560874 | 1 |
| Gchfr | 1.24987375 | 0.321782377 | 0.1561688 | 1 |
| Synj2 | 0.8611444 | -0.215672919 | 0.15617288 | 1 |
| Cdc7 | 0.80502401 | -0.312896283 | 0.15624462 | 1 |
| Wdr72 | 1.38574145 | 0.470658107 | 0.1562703 | 1 |
| Mrpl49 | 1.10043332 | 0.138071733 | 0.15629508 | 1 |

| | | | | |
|---|---|---|---|---|
| Clec4a1 | 1.24993237 | 0.321850038 | 0.1563821 | 1 |
| Diaph3 | 0.82191156 | -0.282944937 | 0.15641931 | 1 |
| Rnf216 | 0.90121451 | -0.15005756 | 0.15642259 | 1 |
| Ier2 | 0.86864429 | -0.203162583 | 0.15644226 | 1 |
| Cluap1 | 1.10675078 | 0.146330386 | 0.15649657 | 1 |
| Myh7 | 0.68307938 | -0.549874863 | 0.15653262 | 1 |
| Cdh23 | 0.68813316 | -0.539240319 | 0.15656454 | 1 |
| Tab1 | 1.11306956 | 0.15454375 | 0.15660249 | 1 |
| Pcdhga9 | 0.81920965 | -0.287695377 | 0.15677451 | 1 |
| Ephx2 | 1.24172817 | 0.312349383 | 0.15703597 | 1 |
| Folr2 | 1.21604294 | 0.282194176 | 0.15707877 | 1 |
| Taf1c | 0.85576933 | -0.224706122 | 0.15715817 | 1 |
| S100b | 1.07104816 | 0.099023357 | 0.15727146 | 1 |
| Ipo13 | 1.09117162 | 0.125878023 | 0.157288 | 1 |
| Taldo1 | 0.93478259 | -0.097297228 | 0.15751919 | 1 |
| Arhgap25 | 1.15822929 | 0.21192089 | 0.15771011 | 1 |
| Myh14 | 0.93383302 | -0.098763497 | 0.1577321 | 1 |
| Sirt4 | 1.2934105 | 0.371180222 | 0.1578079 | 1 |
| Asb7 | 0.80048935 | -0.321045891 | 0.15787599 | 1 |
| Tcf12 | 1.11522003 | 0.157328379 | 0.15788437 | 1 |
| Wnt5a | 1.15228367 | 0.20449592 | 0.15794521 | 1 |
| Ska2 | 1.22126398 | 0.288375072 | 0.15799796 | 1 |
| Rgs7 | 0.92181106 | -0.11745701 | 0.15806069 | 1 |
| Cited2 | 0.88341951 | -0.178829407 | 0.15818567 | 1 |
| Manea | 0.8933898 | -0.162638311 | 0.1582116 | 1 |
| Klhl42 | 0.89291169 | -0.163410599 | 0.15837594 | 1 |
| Cmss1 | 1.17071985 | 0.227395887 | 0.15842117 | 1 |
| Prickle2 | 1.09446486 | 0.13022564 | 0.15845181 | 1 |
| Mapt | 0.93707075 | -0.093770115 | 0.15859402 | 1 |
| Tubb2b | 0.93655665 | -0.094561838 | 0.15869815 | 1 |
| Eci1 | 1.13865729 | 0.187333599 | 0.15876188 | 1 |
| Bnip1 | 0.8699123 | -0.201058138 | 0.1588234 | 1 |
| Trpc1 | 0.91521808 | -0.127812541 | 0.15887708 | 1 |
| Cse1l | 1.0725563 | 0.101053378 | 0.15898227 | 1 |
| Atp2a2 | 0.93628632 | -0.094978316 | 0.1590117 | 1 |
| Rtn4rl1 | 0.93163105 | -0.102169364 | 0.15920635 | 1 |
| Thbd | 1.08130483 | 0.112773285 | 0.15927591 | 1 |
| Prrc1 | 1.18497481 | 0.244856387 | 0.15931828 | 1 |
| Snrk | 0.91538036 | -0.127556759 | 0.15942436 | 1 |
| Tulp4 | 0.91468866 | -0.128647326 | 0.15951323 | 1 |

| | | | | | |
|---|---|---|---|---|---|
| L1cam | | 0.93656062 | -0.094555715 | 0.15957598 | 1 |
| Mark3 | | 0.92168124 | -0.117660212 | 0.15959476 | 1 |
| Mybl1 | | 1.21124411 | 0.276489649 | 0.15960635 | 1 |
| Copz2 | | 0.85837873 | -0.220313773 | 0.15965089 | 1 |
| Reep2 | | 0.93660831 | -0.094482259 | 0.15966965 | 1 |
| Arhgap31 | | 0.90599488 | -0.142425196 | 0.1596763 | 1 |
| Stambp | | 0.91045197 | -0.135345181 | 0.15968635 | 1 |
| Htra2 | | 0.9055328 | -0.143161191 | 0.15972914 | 1 |
| Trim10 | | 1.66216095 | 0.733060087 | 0.15984676 | 1 |
| Kti12 | | 0.86056144 | -0.216649894 | 0.15995981 | 1 |
| Cln6 | | 0.90336646 | -0.14661675 | 0.16023022 | 1 |
| Traf2 | | 1.1393689 | 0.188234934 | 0.16026975 | 1 |
| Naa50 | | 0.90501703 | -0.143983154 | 0.16032456 | 1 |
| Tmem179b | | 1.22373282 | 0.291288604 | 0.16039641 | 1 |
| Nrm | | 1.26682697 | 0.341219482 | 0.16060083 | 1 |
| Msn | | 1.06669231 | 0.093144092 | 0.16071004 | 1 |
| Slc35f6 | | 0.9068565 | -0.141053816 | 0.16071751 | 1 |
| Limch1 | | 0.92455979 | -0.113161482 | 0.16088226 | 1 |
| Rasl10b | | 0.90091981 | -0.150529401 | 0.16095968 | 1 |
| Slc35b4 | | 1.07562886 | 0.105180368 | 0.16108379 | 1 |
| Kctd16 | | 0.85691802 | -0.222770904 | 0.16113659 | 1 |
| Atpif1 | | 1.06744942 | 0.094167711 | 0.16114795 | 1 |
| Ptges3l1 | | 0.89666199 | -0.157363859 | 0.16115818 | 1 |
| Nop9 | | 0.87268379 | -0.196469092 | 0.16115896 | 1 |
| Tmem88 | | 0.80940761 | -0.305061683 | 0.16117193 | 1 |
| Hsd11b1 | | 1.21236721 | 0.277826737 | 0.16139752 | 1 |
| Aspn | | 1.08070767 | 0.111976326 | 0.16140021 | 1 |
| Dnajc4 | | 1.1927073 | 0.254240042 | 0.16155808 | 1 |
| Lipt2 | | 0.74927768 | -0.416427615 | 0.16174365 | 1 |
| Zbtb37 | | 1.37696158 | 0.461488302 | 0.16186706 | 1 |
| Trim23 | | 1.07492198 | 0.104231951 | 0.16198854 | 1 |
| Cul3 | | 0.93668327 | -0.0943668 | 0.16201487 | 1 |
| Klhl4 | | 1.26825055 | 0.342839785 | 0.16246935 | 1 |
| St3gal1 | | 1.21135215 | 0.276618333 | 0.16273992 | 1 |
| Htr1d | | 1.1350624 | 0.182771606 | 0.1627921 | 1 |
| Ube2r2 | | 0.92035277 | -0.11974115 | 0.16279653 | 1 |
| Slamf6 | | 0.65989025 | -0.599701987 | 0.16285161 | 1 |
| Hrc | | 0.65231922 | -0.616349962 | 0.16285608 | 1 |
| Ccnd2 | | 0.8484432 | -0.237110011 | 0.16289375 | 1 |
| | 8-Sep | 0.93653624 | -0.094593274 | 0.16289552 | 1 |

| | | | | |
|---|---|---|---|---|
| Exd1 | 1.33463087 | 0.416440784 | 0.16298732 | 1 |
| Zfp438 | 1.2297762 | 0.298395793 | 0.16308162 | 1 |
| Psmb5 | 1.06993382 | 0.097521558 | 0.16319463 | 1 |
| Rfwd2 | 0.91118938 | -0.134177157 | 0.16325545 | 1 |
| Gpr34 | 1.28610597 | 0.363009522 | 0.1635916 | 1 |
| Itsn1 | 0.92018331 | -0.120006811 | 0.16382606 | 1 |
| Abcg3l1 | 1.14048198 | 0.189643646 | 0.16387532 | 1 |
| Sgce | 1.09731553 | 0.133978424 | 0.16394447 | 1 |
| Gpbp1l1 | 1.09978966 | 0.137227631 | 0.16410827 | 1 |
| Pou3f2 | 1.14644829 | 0.197171289 | 0.16420961 | 1 |
| Crlf1 | 1.19532725 | 0.257405648 | 0.16422609 | 1 |
| Cntn1 | 1.06742687 | 0.094137235 | 0.16424896 | 1 |
| Napsa | 1.27695933 | 0.352712572 | 0.16429449 | 1 |
| Agmo | 1.13867138 | 0.187351441 | 0.1643093 | 1 |
| Tyrp1 | 1.06702837 | 0.09359853 | 0.16439623 | 1 |
| Ptprv | 0.71750381 | -0.478941593 | 0.16440496 | 1 |
| Smad1 | 0.90296252 | -0.147261988 | 0.16445636 | 1 |
| Rwdd2b | 1.15527389 | 0.208234925 | 0.16458447 | 1 |
| Dis3l | 1.09914371 | 0.136380023 | 0.1645977 | 1 |
| Elf4 | 1.28296441 | 0.359481153 | 0.16467791 | 1 |
| Wdr73 | 0.85208799 | -0.230925678 | 0.16470954 | 1 |
| Atad2 | 0.88370131 | -0.178369274 | 0.16477081 | 1 |
| Glrx | 1.06891567 | 0.096148034 | 0.16477524 | 1 |
| Tesk2 | 1.17799801 | 0.2363371 | 0.1651562 | 1 |
| LOC100910945 | 0.74995739 | -0.415119471 | 0.16516998 | 1 |
| Slc25a26 | 0.81681425 | -0.291920065 | 0.16517523 | 1 |
| Mlycd | 0.89542224 | -0.159359939 | 0.16530516 | 1 |
| Fermt3 | 1.19853342 | 0.261270135 | 0.16533197 | 1 |
| Gja1 | 0.91112703 | -0.134275887 | 0.16559792 | 1 |
| Rbx1 | 1.07275732 | 0.101323745 | 0.16560097 | 1 |
| Pdia5 | 0.80751689 | -0.308435662 | 0.16571436 | 1 |
| Fam136a | 1.15902076 | 0.212906413 | 0.16578253 | 1 |
| Unc119b | 0.91513805 | -0.12793871 | 0.16588242 | 1 |
| Gpr68 | 0.8948056 | -0.160353811 | 0.1659042 | 1 |
| Fastkd2 | 1.11918067 | 0.162442957 | 0.16591889 | 1 |
| Mtmr6 | 1.06634405 | 0.092672993 | 0.16608732 | 1 |
| Sox5 | 1.38271871 | 0.467507694 | 0.16621079 | 1 |
| Rpa1 | 1.08943812 | 0.123584248 | 0.16630897 | 1 |
| Vipas39 | 1.08365346 | 0.115903477 | 0.16640794 | 1 |
| Ltb | 1.30867204 | 0.388103594 | 0.16643628 | 1 |

| | | | | |
|---|---|---|---|---|
| Mmp23 | 1.24898441 | 0.320755467 | 0.16655643 | 1 |
| Grcc10 | 1.07921136 | 0.109977446 | 0.16659489 | 1 |
| Slc47a1 | 1.13291395 | 0.180038282 | 0.16664002 | 1 |
| Kdr | 0.90421563 | -0.14526124 | 0.16690082 | 1 |
| Maoa | 1.07403696 | 0.103043643 | 0.16693318 | 1 |
| Adam9 | 1.06810932 | 0.09505931 | 0.16698966 | 1 |
| Fam219b | 0.84716714 | -0.239281459 | 0.1671426 | 1 |
| Commd1 | 1.08294041 | 0.114953854 | 0.16726237 | 1 |
| Slc35e1 | 0.90903931 | -0.137585419 | 0.16727531 | 1 |
| Mmp8 | 1.28134164 | 0.357655192 | 0.16733825 | 1 |
| Smtn | 0.9113133 | -0.133980972 | 0.16745454 | 1 |
| Phf10 | 1.13930382 | 0.18815253 | 0.16754357 | 1 |
| Has2 | 1.31435257 | 0.394352327 | 0.16787946 | 1 |
| Kif19 | 0.92455721 | -0.113165504 | 0.16788588 | 1 |
| Dnajc12 | 0.87629891 | -0.190505029 | 0.16794013 | 1 |
| Vps13a | 0.93490192 | -0.097113076 | 0.16807263 | 1 |
| Sarm1 | 0.92782346 | -0.108077775 | 0.16828627 | 1 |
| Pdk2 | 1.08942169 | 0.123562492 | 0.16832336 | 1 |
| Dhx16 | 0.8953728 | -0.1594396 | 0.16836 | 1 |
| Scaf1 | 0.93192949 | -0.101707285 | 0.16845384 | 1 |
| Ddx50 | 1.08367379 | 0.115930541 | 0.16855193 | 1 |
| Rasgrf1 | 0.8862353 | -0.174238297 | 0.16863486 | 1 |
| Usp22 | 0.92573634 | -0.111326735 | 0.16871397 | 1 |
| Slc7a8 | 0.89809684 | -0.155057081 | 0.16876252 | 1 |
| Dynll1 | 0.93892621 | -0.090916308 | 0.16883924 | 1 |
| Syngr1 | 0.90557291 | -0.143097293 | 0.16900142 | 1 |
| Cpne6 | 1.06549452 | 0.091523169 | 0.16902215 | 1 |
| Pip5k1c | 0.93822396 | -0.091995753 | 0.16917298 | 1 |
| Cnot6 | 0.89851639 | -0.154383278 | 0.16928619 | 1 |
| B4galt2 | 0.83661955 | -0.257356386 | 0.16939067 | 1 |
| Atg13 | 1.0793372 | 0.110145649 | 0.16953765 | 1 |
| Ano6 | 0.93082715 | -0.103414805 | 0.1695603 | 1 |
| RT1-T24-4 | 1.1160015 | 0.158338969 | 0.16985741 | 1 |
| LOC100912041 | 1.180998 | 0.240006521 | 0.16991021 | 1 |
| Pawr | 1.20181251 | 0.265211842 | 0.17002541 | 1 |
| Hpdl | 0.83419584 | -0.261541981 | 0.17008266 | 1 |
| Cuedc2 | 0.87060442 | -0.199910752 | 0.17008929 | 1 |
| Npr1 | 0.84298196 | -0.24642633 | 0.17016181 | 1 |
| Ubtd1 | 0.88903358 | -0.169690179 | 0.17056386 | 1 |
| Abhd2 | 0.9346722 | -0.097467614 | 0.17068946 | 1 |

| | | | | |
|---|---|---|---|---|
| B3gnt2 | 0.92789787 | -0.107962078 | 0.17084286 | 1 |
| Actr6 | 1.09523658 | 0.13124254 | 0.17093031 | 1 |
| Ptprn2 | 0.9196308 | -0.12087331 | 0.17101352 | 1 |
| Gpr158 | 1.06564297 | 0.091724167 | 0.17111501 | 1 |
| Zw10 | 1.12301878 | 0.167382055 | 0.17119795 | 1 |
| Ctsw | 0.66536549 | -0.587781062 | 0.171245 | 1 |
| Slc29a1 | 0.87495384 | -0.192721196 | 0.171322 | 1 |
| Bmp1 | 0.91887188 | -0.122064373 | 0.17144545 | 1 |
| Nmnat2 | 0.91222347 | -0.132540798 | 0.17150314 | 1 |
| Car5b | 1.17193309 | 0.228890201 | 0.17152319 | 1 |
| Pcbp3 | 0.93828468 | -0.09190239 | 0.17153297 | 1 |
| Mme | 0.88916062 | -0.169484043 | 0.17164774 | 1 |
| Fbxo5 | 1.30212632 | 0.380869416 | 0.17165073 | 1 |
| Cul9 | 0.91653054 | -0.12574514 | 0.17183841 | 1 |
| Atp9b | 1.06745118 | 0.094170096 | 0.17189763 | 1 |
| Ibsp | 0.72642796 | -0.461108361 | 0.1719536 | 1 |
| Slc48a1 | 0.9230424 | -0.11553118 | 0.17225948 | 1 |
| Usp53 | 1.08275609 | 0.114708288 | 0.17227269 | 1 |
| Tlr1 | 1.44982135 | 0.535875136 | 0.17230616 | 1 |
| Syt6 | 0.8871011 | -0.172829559 | 0.17234649 | 1 |
| Tmem163 | 1.10418184 | 0.142977775 | 0.17235383 | 1 |
| Fancl | 0.82858837 | -0.27127252 | 0.1727295 | 1 |
| Lrp10 | 0.92856626 | -0.106923231 | 0.17274774 | 1 |
| Trim13 | 1.13996128 | 0.188984819 | 0.17284038 | 1 |
| Ptk7 | 0.91167521 | -0.133408142 | 0.17314273 | 1 |
| Ttc7b | 0.92438226 | -0.113438519 | 0.17315721 | 1 |
| Id2 | 1.14660677 | 0.197370698 | 0.17329159 | 1 |
| Aifm1 | 1.09002351 | 0.124359255 | 0.17334812 | 1 |
| Tsku | 0.85881494 | -0.219580809 | 0.17343123 | 1 |
| Calm2 | 1.06803146 | 0.094954149 | 0.17357683 | 1 |
| Kpnb1 | 0.93756867 | -0.093003738 | 0.17357873 | 1 |
| Rnmtl1 | 0.86232661 | -0.213693698 | 0.17378696 | 1 |
| Dag1 | 0.93875557 | -0.091178526 | 0.17390828 | 1 |
| Rab39a | 0.88894318 | -0.169836893 | 0.17396028 | 1 |
| Rmnd1 | 0.89163443 | -0.165475765 | 0.17405465 | 1 |
| Srd5a3 | 1.11612049 | 0.158492781 | 0.1741147 | 1 |
| Icmt | 0.91268214 | -0.131815592 | 0.17414752 | 1 |
| Map4k4 | 0.93898457 | -0.090826648 | 0.17423607 | 1 |
| Samd11 | 0.84095648 | -0.249896951 | 0.17424703 | 1 |
| Pbld1 | 1.19444402 | 0.256339238 | 0.17439353 | 1 |

| | | | | |
|---|---|---|---|---|
| Tdg | 0.91267891 | -0.131820699 | 0.17462815 | 1 |
| Ptbp3 | 1.09579008 | 0.131971445 | 0.17476066 | 1 |
| Lin54 | 0.90157084 | -0.149487234 | 0.1749255 | 1 |
| Stbd1 | 1.08498425 | 0.117674094 | 0.1749659 | 1 |
| Acot1 | 0.85270781 | -0.229876631 | 0.17500199 | 1 |
| Kdm4b | 0.91935971 | -0.121298656 | 0.17500267 | 1 |
| Pigt | 0.93290769 | -0.100193761 | 0.1751727 | 1 |
| Ghr | 1.09455614 | 0.130345949 | 0.17521884 | 1 |
| Cdkn2b | 0.85946351 | -0.218491705 | 0.17523108 | 1 |
| Rnf14 | 0.93854278 | -0.091505594 | 0.17525643 | 1 |
| Ap1s3 | 0.82279279 | -0.281398936 | 0.17528503 | 1 |
| Slc51a | 0.85366137 | -0.228264196 | 0.17529852 | 1 |
| Pak1 | 1.06879718 | 0.0959881 | 0.17543581 | 1 |
| Tecr | 1.06543983 | 0.091449115 | 0.17544032 | 1 |
| Ncoa6 | 0.92522951 | -0.11211682 | 0.17544166 | 1 |
| Mrpl18 | 0.9153211 | -0.127650162 | 0.17544588 | 1 |
| Vps39 | 1.06741494 | 0.094121112 | 0.17552298 | 1 |
| Poll | 1.10967346 | 0.150135208 | 0.17559137 | 1 |
| Cd24 | 0.93877099 | -0.091154836 | 0.17559469 | 1 |
| Tmed9 | 1.07367161 | 0.102552809 | 0.17582328 | 1 |
| Flii | 0.93933963 | -0.090281213 | 0.17583602 | 1 |
| LOC365238 | 1.18475344 | 0.244586849 | 0.17589792 | 1 |
| Ankrd27 | 1.0753742 | 0.104838771 | 0.17590235 | 1 |
| Dapk1 | 0.92546123 | -0.111755548 | 0.17591247 | 1 |
| Cep19 | 1.07239577 | 0.100837438 | 0.17612609 | 1 |
| Zdhhc1 | 0.88925386 | -0.169332762 | 0.17631409 | 1 |
| Hs6st1 | 0.92692483 | -0.109475749 | 0.17632105 | 1 |
| Prkce | 0.93003087 | -0.104649496 | 0.17689832 | 1 |
| Plekhh1 | 0.91886429 | -0.122076298 | 0.17690938 | 1 |
| Fbxo36 | 0.7696982 | -0.377635226 | 0.17695075 | 1 |
| Tlr4 | 1.17263846 | 0.229758276 | 0.17710661 | 1 |
| Cytl1 | 1.15944311 | 0.213432031 | 0.17712946 | 1 |
| Dolk | 0.90684245 | -0.141076164 | 0.17764207 | 1 |
| Gab1 | 1.07176732 | 0.099991734 | 0.17776725 | 1 |
| Ctbp1 | 0.93917073 | -0.090540655 | 0.17780771 | 1 |
| Slc25a10 | 0.89973749 | -0.152423962 | 0.17785845 | 1 |
| Olfml2b | 0.90462618 | -0.144606343 | 0.17797009 | 1 |
| R3hdml | 1.35828121 | 0.441782192 | 0.17798933 | 1 |
| Cd14 | 1.24470961 | 0.315809202 | 0.1780739 | 1 |
| Mtmr2 | 0.93916788 | -0.09054502 | 0.17829845 | 1 |

| | | | | |
|---|---|---|---|---|
| Akap2 | 1.0661336 | 0.092388234 | 0.17836789 | 1 |
| Klf12 | 0.81059879 | -0.302940078 | 0.17844494 | 1 |
| Alcam | 0.93996392 | -0.089322714 | 0.17845843 | 1 |
| Celf2 | 0.88424569 | -0.177480816 | 0.1785032 | 1 |
| Fam69a | 1.09188535 | 0.126821374 | 0.17850962 | 1 |
| Pde7b | 0.82761121 | -0.272974916 | 0.17862838 | 1 |
| Cdk5rap2 | 1.0973274 | 0.133994039 | 0.17865533 | 1 |
| LOC100158225 | 0.79750489 | -0.326434728 | 0.17887045 | 1 |
| Marveld1 | 0.89896366 | -0.153665296 | 0.17929672 | 1 |
| Adgre1 | 1.13026845 | 0.176665467 | 0.17951662 | 1 |
| Nap1l3 | 0.92913673 | -0.106037186 | 0.17962904 | 1 |
| Gsk3b | 0.92573668 | -0.111326204 | 0.17963074 | 1 |
| Defa5 | 1.2701187 | 0.344963327 | 0.17981816 | 1 |
| LOC100361083 | 0.88062674 | -0.183397439 | 0.17983623 | 1 |
| Plec | 0.93920053 | -0.09049488 | 0.18000406 | 1 |
| Rpe | 1.08882532 | 0.122772515 | 0.18015696 | 1 |
| Fgf12 | 1.0721965 | 0.100569325 | 0.1801967 | 1 |
| Kdm4c | 1.10288705 | 0.141285054 | 0.18021959 | 1 |
| Hs6st2 | 0.90848314 | -0.138468348 | 0.18053107 | 1 |
| Cacna1c | 0.91077601 | -0.134831803 | 0.18080983 | 1 |
| Uqcr10 | 1.07269365 | 0.101238112 | 0.18088046 | 1 |
| Cbarp | 0.92176474 | -0.117529512 | 0.18093695 | 1 |
| Dhcr24 | 0.93846899 | -0.091619022 | 0.18094864 | 1 |
| Neto2 | 0.87395745 | -0.194365048 | 0.18101171 | 1 |
| Necab3 | 1.06708907 | 0.093680604 | 0.18106484 | 1 |
| Apool | 1.11873588 | 0.16186947 | 0.18111061 | 1 |
| Sin3b | 0.90809466 | -0.139085404 | 0.18120413 | 1 |
| Lnc134 | 1.272889 | 0.348106622 | 0.1812519 | 1 |
| Dusp8 | 0.87672494 | -0.189803805 | 0.18129975 | 1 |
| Mapk1ip1l | 0.89411485 | -0.161467937 | 0.18195773 | 1 |
| Alkbh2 | 1.2695836 | 0.344355393 | 0.18233821 | 1 |
| Top1 | 0.90720966 | -0.140492092 | 0.18239447 | 1 |
| Map1lc3b | 1.06505771 | 0.090931601 | 0.18281512 | 1 |
| Hcfc1 | 0.93559161 | -0.09604917 | 0.18289547 | 1 |
| Fut11 | 0.84929024 | -0.235670428 | 0.18294858 | 1 |
| Dpysl5 | 0.93926725 | -0.090392394 | 0.18318904 | 1 |
| Fech | 1.08621093 | 0.119304285 | 0.18323152 | 1 |
| Sdhd | 1.06754921 | 0.094302574 | 0.18334893 | 1 |
| Chchd4 | 1.14563133 | 0.196142846 | 0.18346294 | 1 |
| Nle1 | 0.84053473 | -0.250620663 | 0.18353672 | 1 |

| | | | | |
|---|---|---|---|---|
| Pik3r2 | 0.92159761 | -0.117791113 | 0.18363431 | 1 |
| Clvs1 | 1.10991759 | 0.150452565 | 0.18364294 | 1 |
| Vps52 | 0.92771112 | -0.108252463 | 0.18373601 | 1 |
| Ublcp1 | 1.06405913 | 0.089578318 | 0.18373717 | 1 |
| Gss | 1.10111881 | 0.138970148 | 0.18380403 | 1 |
| Stk11ip | 0.88910384 | -0.169576168 | 0.18386038 | 1 |
| Cyfip2 | 0.94054357 | -0.088433317 | 0.18390272 | 1 |
| Ino80d | 0.82293966 | -0.281141437 | 0.18400161 | 1 |
| Atp6v1e1 | 1.06332121 | 0.088577472 | 0.18412974 | 1 |
| Ptpn2 | 1.16945998 | 0.225842498 | 0.18420292 | 1 |
| Tnip2 | 0.85007853 | -0.234331964 | 0.1842289 | 1 |
| Foxo3 | 0.89927034 | -0.153173201 | 0.18425272 | 1 |
| Cep44 | 1.17781453 | 0.236112375 | 0.18427642 | 1 |
| Dvl3 | 0.89227812 | -0.164434637 | 0.18449678 | 1 |
| Mnat1 | 1.09198884 | 0.126958115 | 0.18456676 | 1 |
| Phf8 | 0.92356435 | -0.114715611 | 0.18465224 | 1 |
| Kcnq2 | 0.93343489 | -0.0993787 | 0.18482816 | 1 |
| Sipa1l2 | 0.9229634 | -0.115654649 | 0.18521762 | 1 |
| Thsd1 | 0.85498633 | -0.226026748 | 0.18524809 | 1 |
| Soga3 | 0.94108623 | -0.08760117 | 0.18525146 | 1 |
| Gtf2ird2 | 1.27250933 | 0.347676234 | 0.18525392 | 1 |
| Armc10 | 1.09509314 | 0.131053576 | 0.18530103 | 1 |
| LOC684871 | 0.85656141 | -0.223371407 | 0.18538967 | 1 |
| Rbm11 | 1.07136915 | 0.099455662 | 0.18553812 | 1 |
| Wdr46 | 0.88829235 | -0.170893534 | 0.18558751 | 1 |
| Map2k6 | 1.13734871 | 0.185674654 | 0.18564148 | 1 |
| Celf5 | 0.8658836 | -0.207754989 | 0.1857176 | 1 |
| Dnajc28 | 1.18132758 | 0.240409079 | 0.18610542 | 1 |
| Msh3 | 1.11598073 | 0.158312113 | 0.18611111 | 1 |
| Mmp19 | 1.24178861 | 0.31241961 | 0.1864094 | 1 |
| Phyh | 1.07512948 | 0.104510411 | 0.18645479 | 1 |
| Creb3l2 | 0.90177184 | -0.149165633 | 0.18645627 | 1 |
| Tmem140 | 1.23238997 | 0.301458844 | 0.18653129 | 1 |
| Sec24b | 0.92558132 | -0.111568352 | 0.18657639 | 1 |
| Sdsl | 0.75535231 | -0.40477839 | 0.18666908 | 1 |
| Scmh1 | 0.8788228 | -0.186355793 | 0.18668565 | 1 |
| Gramd1a | 0.93028782 | -0.104250962 | 0.18673453 | 1 |
| Ccdc61 | 1.18511822 | 0.245030986 | 0.18673791 | 1 |
| Parva | 1.08855073 | 0.122408643 | 0.18675553 | 1 |
| Rhbdf1 | 0.91403529 | -0.129678223 | 0.1867719 | 1 |

| | | | | |
|---|---|---|---|---|
| Mpi | 1.08097808 | 0.112337264 | 0.18682149 | 1 |
| Col27a1 | 0.88537297 | -0.175642761 | 0.18703154 | 1 |
| Apba3 | 0.87622618 | -0.190624767 | 0.18729436 | 1 |
| Pcdhga7 | 1.1169586 | 0.159575708 | 0.18731896 | 1 |
| Spef1 | 0.87247615 | -0.196812404 | 0.18733821 | 1 |
| Dpp10 | 1.06345435 | 0.08875811 | 0.18748857 | 1 |
| Mapk14 | 0.92519682 | -0.11216778 | 0.18752969 | 1 |
| Phf19 | 0.76719675 | -0.382331485 | 0.18753039 | 1 |
| Pdxk | 0.81656406 | -0.292362017 | 0.18757341 | 1 |
| Map7d1 | 0.9395304 | -0.089988248 | 0.18757866 | 1 |
| Ngfr | 0.93992586 | -0.089381126 | 0.18764962 | 1 |
| Dpy19l1 | 0.93000352 | -0.104691925 | 0.18765773 | 1 |
| Htr7 | 0.7927103 | -0.335134379 | 0.18766792 | 1 |
| Ccndbp1 | 1.08522822 | 0.117998473 | 0.18769379 | 1 |
| Mlf1 | 1.1970272 | 0.259455932 | 0.18777542 | 1 |
| Nfasc | 0.94109627 | -0.087585778 | 0.18779351 | 1 |
| Atp6v1d | 1.0630417 | 0.088198197 | 0.18780874 | 1 |
| Acyp1 | 1.12045423 | 0.164083715 | 0.18783992 | 1 |
| Nppa | 1.27628479 | 0.351950291 | 0.18793693 | 1 |
| Fam220a | 0.90506494 | -0.143906778 | 0.18804313 | 1 |
| Cdh2 | 0.9393864 | -0.09020938 | 0.18812142 | 1 |
| Zfp14 | 0.8611712 | -0.215628019 | 0.18813468 | 1 |
| LOC106631776 | 0.84649866 | -0.240420319 | 0.18829554 | 1 |
| Ubr2 | 0.93657538 | -0.094532984 | 0.18834221 | 1 |
| Kank4 | 0.92854896 | -0.106950112 | 0.18837967 | 1 |
| Ccdc80 | 0.92546901 | -0.11174342 | 0.18838884 | 1 |
| Mapk4 | 0.9028364 | -0.147463513 | 0.1885312 | 1 |
| Ston2 | 1.20682773 | 0.271219754 | 0.1885435 | 1 |
| Diras1 | 0.9285998 | -0.106871117 | 0.18857771 | 1 |
| Cklf | 1.36799194 | 0.452059727 | 0.18860058 | 1 |
| Hoxc9 | 2.01499362 | 1.010775268 | 0.18871364 | 1 |
| Epc1 | 1.09456057 | 0.130351793 | 0.18876727 | 1 |
| Thoc1 | 1.11907707 | 0.162309392 | 0.18877605 | 1 |
| Megf6 | 1.18890695 | 0.249635801 | 0.18889046 | 1 |
| Prrx1 | 1.19824533 | 0.26092332 | 0.18889653 | 1 |
| Myo5b | 0.85347385 | -0.228581143 | 0.18892075 | 1 |
| Ndufa11 | 1.08198692 | 0.113683061 | 0.18892409 | 1 |
| Dixdc1 | 0.88782407 | -0.171654277 | 0.18893395 | 1 |
| Mrps16 | 0.89957855 | -0.152678835 | 0.18896087 | 1 |
| Idua | 1.09505024 | 0.130997055 | 0.18897716 | 1 |

| | | | | |
|---|---|---|---|---|
| Megf9 | 0.93933011 | -0.090295844 | 0.18911251 | 1 |
| Rer1 | 0.92816178 | -0.107551804 | 0.18919967 | 1 |
| Hprt1 | 1.06824096 | 0.095237104 | 0.1892432 | 1 |
| Panx1 | 0.88792097 | -0.171496819 | 0.18926359 | 1 |
| Ggt7 | 0.93908199 | -0.090676966 | 0.18944273 | 1 |
| LOC499770 | 0.88420315 | -0.177550221 | 0.18947273 | 1 |
| RGD1563941 | 1.31063236 | 0.390263058 | 0.18953031 | 1 |
| Appbp2 | 0.9332122 | -0.099722922 | 0.18954133 | 1 |
| Stk40 | 0.9178535 | -0.1236642 | 0.1895704 | 1 |
| Slf1 | 1.13755912 | 0.185941526 | 0.18966139 | 1 |
| Nmral1 | 0.83712685 | -0.256481843 | 0.18987851 | 1 |
| Rad52 | 1.17968805 | 0.238405407 | 0.18989446 | 1 |
| Spryd4 | 1.15009646 | 0.201754871 | 0.18997238 | 1 |
| RT1-Db2 | 1.29395442 | 0.371786796 | 0.1900724 | 1 |
| Snrpa1 | 1.13189668 | 0.178742272 | 0.19014119 | 1 |
| Arhgap29 | 1.09622373 | 0.132542269 | 0.19014946 | 1 |
| Lyrm7 | 1.11263109 | 0.153975325 | 0.19018818 | 1 |
| Gria1 | 0.77215622 | -0.37303534 | 0.19026926 | 1 |
| Cc2d1a | 0.90682586 | -0.141102566 | 0.19043561 | 1 |
| Gng2 | 0.94162547 | -0.086774746 | 0.19052391 | 1 |
| Ppp2r3c | 1.09258249 | 0.127742211 | 0.19074341 | 1 |
| Brpf3 | 0.82122436 | -0.284151668 | 0.19086672 | 1 |
| Serpine1 | 0.8660944 | -0.207403814 | 0.19092602 | 1 |
| Galnt16 | 1.17049839 | 0.227122944 | 0.19106652 | 1 |
| Pla1a | 1.24879554 | 0.320537284 | 0.19108382 | 1 |
| Rasd1 | 0.81373663 | -0.297366158 | 0.19117267 | 1 |
| Cacng4 | 0.80074351 | -0.320587896 | 0.19128922 | 1 |
| Megf8 | 0.93997163 | -0.089310883 | 0.19143819 | 1 |
| Arl10 | 1.28933825 | 0.366630796 | 0.19146705 | 1 |
| Slc2a4 | 1.29245678 | 0.37011604 | 0.19148172 | 1 |
| Tubgcp5 | 1.08375583 | 0.116039748 | 0.19152754 | 1 |
| Minpp1 | 1.08768086 | 0.121255316 | 0.19158016 | 1 |
| Pcdh17 | 0.906884 | -0.141010068 | 0.19164725 | 1 |
| Dram2 | 0.90168753 | -0.149300529 | 0.19173182 | 1 |
| Dhx38 | 1.09818251 | 0.135117846 | 0.19173932 | 1 |
| Borcs6 | 0.90274766 | -0.147605326 | 0.19181461 | 1 |
| Shroom4 | 1.1915875 | 0.252884893 | 0.19204109 | 1 |
| Wbscr16 | 1.16076282 | 0.215073218 | 0.19211658 | 1 |
| RGD1562987 | 1.06487887 | 0.090689328 | 0.19224424 | 1 |
| RGD1560108 | 0.90810346 | -0.139071416 | 0.19234212 | 1 |

| | | | | |
|---|---|---|---|---|
| Kcnk4 | 0.85995056 | -0.217674368 | 0.19253948 | 1 |
| Crls1 | 0.89488613 | -0.160223984 | 0.19261281 | 1 |
| Pced1b | 1.19546347 | 0.25757004 | 0.19266094 | 1 |
| Decr1 | 1.10796267 | 0.14790927 | 0.19270979 | 1 |
| Cpq | 1.07396042 | 0.102940828 | 0.19274024 | 1 |
| B3gnt5 | 0.87197606 | -0.197639562 | 0.19276355 | 1 |
| Ankrd34b | 1.19850301 | 0.261233538 | 0.19289048 | 1 |
| Ankrd54 | 0.84274649 | -0.246829383 | 0.19291504 | 1 |
| Gtf2ird1 | 0.88939407 | -0.169105304 | 0.19300708 | 1 |
| Zfp68 | 1.07968684 | 0.110612924 | 0.19304223 | 1 |
| Rbms1 | 0.9186724 | -0.12237761 | 0.19309102 | 1 |
| Slc7a6os | 1.12547176 | 0.170529855 | 0.19312414 | 1 |
| Med11 | 1.19481291 | 0.256784734 | 0.19316935 | 1 |
| Nfyb | 1.1066169 | 0.146155864 | 0.1932165 | 1 |
| P2ry10 | 1.31530741 | 0.395400026 | 0.19323999 | 1 |
| Fgfr2 | 1.14677757 | 0.197585597 | 0.19325143 | 1 |
| Fkbp7 | 0.87548117 | -0.19185195 | 0.19337952 | 1 |
| Tmcc2 | 1.08456134 | 0.117111645 | 0.19346611 | 1 |
| Cdh8 | 1.20704183 | 0.27147567 | 0.19361714 | 1 |
| Atg9a | 0.93599072 | -0.095433868 | 0.19365952 | 1 |
| Igbp1 | 1.08865509 | 0.122546947 | 0.19367711 | 1 |
| Cep83 | 1.0779687 | 0.108315293 | 0.19368033 | 1 |
| Ppdpf | 0.92035701 | -0.119734493 | 0.19372208 | 1 |
| Crym | 0.88364585 | -0.178459816 | 0.19373853 | 1 |
| Tk2 | 1.11025398 | 0.150889742 | 0.19398903 | 1 |
| Prss12 | 1.07839311 | 0.108883186 | 0.19405 | 1 |
| Astn2 | 0.89846986 | -0.154457979 | 0.19407582 | 1 |
| Akr1b10 | 0.87213746 | -0.19737256 | 0.19411324 | 1 |
| Myt1 | 0.91430755 | -0.129248567 | 0.19428326 | 1 |
| Numbl | 0.90820075 | -0.138916873 | 0.19432789 | 1 |
| Tcf7l1 | 0.86333399 | -0.212009312 | 0.19436725 | 1 |
| Slitrk4 | 0.91674992 | -0.125399855 | 0.19452266 | 1 |
| Ddx18 | 1.11407293 | 0.155843683 | 0.19472124 | 1 |
| Gpr4 | 0.80606499 | -0.311031928 | 0.19474017 | 1 |
| Repin1 | 1.14575641 | 0.19630036 | 0.1951552 | 1 |
| Eif3e | 1.07089179 | 0.098812705 | 0.19524055 | 1 |
| Papola | 0.93905273 | -0.090721928 | 0.19539712 | 1 |
| Rtn4rl2 | 0.88075596 | -0.183185767 | 0.19559674 | 1 |
| Gabbr1 | 0.94142823 | -0.087076983 | 0.19561961 | 1 |
| Umps | 1.09830058 | 0.135272936 | 0.19567533 | 1 |

| | | | | |
|---|---|---|---|---|
| Pcdha2 | 0.87049659 | -0.200089452 | 0.19570065 | 1 |
| Ccnd3 | 0.92104995 | -0.11864869 | 0.1958005 | 1 |
| Ice2 | 1.14807165 | 0.199212679 | 0.19604025 | 1 |
| Wdr89 | 0.82893603 | -0.270667328 | 0.19609811 | 1 |
| Map1a | 0.9388134 | -0.091089655 | 0.19616739 | 1 |
| Prelid3b | 1.08513227 | 0.117870906 | 0.19620765 | 1 |
| Syt7 | 0.90971173 | -0.136518646 | 0.19633908 | 1 |
| Tlr8 | 1.20149719 | 0.264833271 | 0.19635156 | 1 |
| Sec16b | 1.16086838 | 0.215204407 | 0.19640001 | 1 |
| Map4 | 0.94182225 | -0.086473286 | 0.19643872 | 1 |
| Tm2d1 | 1.08750091 | 0.121016609 | 0.19664644 | 1 |
| Dcc | 0.6437629 | -0.635398657 | 0.19669244 | 1 |
| Pim3 | 0.91902295 | -0.121827212 | 0.19705506 | 1 |
| Isoc2b | 1.21509519 | 0.281069339 | 0.19707601 | 1 |
| Psmb4 | 1.06217374 | 0.087019773 | 0.19709387 | 1 |
| Trak1 | 0.93930661 | -0.090331935 | 0.19712004 | 1 |
| Foxp4 | 0.86019502 | -0.217264323 | 0.1972481 | 1 |
| Cluh | 0.94085086 | -0.087962041 | 0.19749881 | 1 |
| Tcirg1 | 1.10524315 | 0.144363793 | 0.19749969 | 1 |
| Stx4 | 1.10272537 | 0.141073536 | 0.19751598 | 1 |
| Islr | 1.06561749 | 0.091689669 | 0.19753585 | 1 |
| Mat2a | 0.92952053 | -0.105441363 | 0.19759642 | 1 |
| Atp2a1 | 0.80449176 | -0.313850446 | 0.19772583 | 1 |
| Slc22a3 | 1.27431233 | 0.349718921 | 0.19782733 | 1 |
| Ttbk1 | 0.91167093 | -0.133414917 | 0.19783653 | 1 |
| Fance | 0.87541361 | -0.19196328 | 0.19783926 | 1 |
| B4galt6 | 1.08114633 | 0.112561804 | 0.19789816 | 1 |
| Katnal1 | 0.88862157 | -0.17035894 | 0.19793056 | 1 |
| Klf13 | 0.79286021 | -0.334861566 | 0.19810648 | 1 |
| Eif3i | 1.08915969 | 0.12321549 | 0.19826464 | 1 |
| Pdlim2 | 1.1778159 | 0.236114054 | 0.19850914 | 1 |
| Pde6b | 1.32514029 | 0.4061451 | 0.19863823 | 1 |
| Mboat4 | 0.74120579 | -0.432053946 | 0.19923658 | 1 |
| Pfdn2 | 1.07849343 | 0.109017388 | 0.19923721 | 1 |
| Trpm7 | 0.94031419 | -0.08878521 | 0.199273 | 1 |
| Rtel1 | 0.92137667 | -0.118137024 | 0.1994256 | 1 |
| Hmgcs1 | 0.94017792 | -0.088994293 | 0.19947926 | 1 |
| Ptger3 | 0.79137327 | -0.337569749 | 0.19954357 | 1 |
| Rprm | 1.08218826 | 0.113951497 | 0.19970147 | 1 |
| Adcy1 | 0.88353214 | -0.178645481 | 0.1998542 | 1 |

| | | | | |
|---|---|---|---|---|
| Isca1 | 1.06230497 | 0.087197994 | 0.19989685 | 1 |
| Epn1 | 0.93433213 | -0.097992613 | 0.19989914 | 1 |
| Flrt1 | 1.20882238 | 0.273602279 | 0.19991607 | 1 |
| Pnp | 1.08032033 | 0.111459156 | 0.20006809 | 1 |
| Fgl2 | 0.94249592 | -0.085441728 | 0.20050127 | 1 |
| Aox1 | 1.0832193 | 0.115325346 | 0.20057097 | 1 |
| Mknk2 | 0.92850669 | -0.107015791 | 0.20062937 | 1 |
| Cnppd1 | 0.92379317 | -0.114358211 | 0.20091286 | 1 |
| Nkain3 | 1.37678229 | 0.461300441 | 0.20098988 | 1 |
| Kremen1 | 1.18714064 | 0.247490859 | 0.20106253 | 1 |
| Mrpl48 | 1.07704549 | 0.107079191 | 0.20112803 | 1 |
| Ccdc137 | 1.12255889 | 0.166791128 | 0.20117896 | 1 |
| Kitlg | 1.0618675 | 0.086603762 | 0.20118272 | 1 |
| Th | 1.36412093 | 0.447971551 | 0.20134007 | 1 |
| Hoxd1 | 0.90811969 | -0.139045643 | 0.20137063 | 1 |
| Pbdc1 | 1.10858901 | 0.148724615 | 0.2015016 | 1 |
| Spry2 | 0.87593818 | -0.191099041 | 0.20160601 | 1 |
| Acadm | 1.06673101 | 0.093196429 | 0.20172396 | 1 |
| Skp2 | 0.81825027 | -0.289385927 | 0.20186769 | 1 |
| Irf8 | 1.28591657 | 0.362797047 | 0.20190419 | 1 |
| Grpel1 | 1.07930939 | 0.110108475 | 0.20223662 | 1 |
| Cr2 | 1.59069112 | 0.669653724 | 0.20227796 | 1 |
| Ddx19a | 1.09035938 | 0.124803724 | 0.20229841 | 1 |
| Slc29a4 | 0.85461475 | -0.226653875 | 0.20233208 | 1 |
| Rnf112 | 0.93278559 | -0.100382586 | 0.20245126 | 1 |
| Oplah | 0.89064856 | -0.167071829 | 0.20246098 | 1 |
| Zcchc12 | 1.1218785 | 0.165916439 | 0.20276724 | 1 |
| Strn | 0.91927761 | -0.121427496 | 0.20278898 | 1 |
| Tbc1d20 | 0.89499891 | -0.160042173 | 0.20286423 | 1 |
| Usp39 | 1.10813317 | 0.148131263 | 0.20306282 | 1 |
| Gabbr2 | 0.94289164 | -0.084836119 | 0.20331688 | 1 |
| Fry | 0.94241575 | -0.085564444 | 0.20335742 | 1 |
| Uqcrh | 1.06174099 | 0.086431868 | 0.20340296 | 1 |
| Srgap3 | 0.93455892 | -0.097642472 | 0.20341389 | 1 |
| Srgap2 | 1.09307521 | 0.128392671 | 0.20356877 | 1 |
| Tm7sf2 | 0.90944259 | -0.136945525 | 0.20363203 | 1 |
| Sgsm1 | 0.93036484 | -0.104131518 | 0.20363991 | 1 |
| Psmd4 | 1.06326018 | 0.088494667 | 0.20367328 | 1 |
| Grm7 | 0.90977433 | -0.136419365 | 0.20367411 | 1 |
| Eif1 | 0.94339809 | -0.08406142 | 0.20370871 | 1 |

| | | | | |
|---|---|---|---|---|
| Rcn3 | 0.92145328 | -0.118017074 | 0.20371441 | 1 |
| Serp1 | 0.9407328 | -0.088143095 | 0.20379303 | 1 |
| S100a13 | 1.10293675 | 0.14135006 | 0.20384033 | 1 |
| RGD1311805 | 1.06939148 | 0.096790088 | 0.20390382 | 1 |
| Ndufa10 | 1.0624129 | 0.087344563 | 0.20392863 | 1 |
| Lace1 | 0.83567079 | -0.258993394 | 0.2039705 | 1 |
| Ahcyl2 | 1.06176937 | 0.086470425 | 0.20398212 | 1 |
| Jakmip2 | 1.07891635 | 0.109583014 | 0.20401264 | 1 |
| Mapk8ip3 | 0.94319428 | -0.084373123 | 0.20406175 | 1 |
| Slc25a27 | 1.09009225 | 0.124450235 | 0.20415817 | 1 |
| Lman2 | 1.08699436 | 0.120344461 | 0.20424845 | 1 |
| Twf1 | 0.93534419 | -0.096430751 | 0.2042583 | 1 |
| B4galt1 | 0.89575646 | -0.158821559 | 0.20458831 | 1 |
| Lpl | 1.13353677 | 0.18083119 | 0.20473388 | 1 |
| Tpst1 | 1.11030943 | 0.150961788 | 0.20479717 | 1 |
| Tmem259 | 0.92005675 | -0.120205251 | 0.20492771 | 1 |
| Afap1l2 | 1.0708291 | 0.098728256 | 0.20500935 | 1 |
| Zfp608 | 1.1154587 | 0.157637098 | 0.20504335 | 1 |
| Pex19 | 1.06912374 | 0.096428841 | 0.20511364 | 1 |
| Ncoa4 | 1.05994516 | 0.083989627 | 0.20524472 | 1 |
| Slc16a4 | 1.24215031 | 0.312839758 | 0.20534224 | 1 |
| Nr4a1 | 0.92265583 | -0.116135508 | 0.20538327 | 1 |
| Acat1 | 1.0772542 | 0.107358727 | 0.20548737 | 1 |
| Gk | 1.0811066 | 0.112508781 | 0.20573909 | 1 |
| Ccdc117 | 0.88202408 | -0.181110056 | 0.20589972 | 1 |
| Nkap | 1.13148917 | 0.17822278 | 0.20597385 | 1 |
| Lsamp | 0.9051736 | -0.143733581 | 0.2060195 | 1 |
| Peli2 | 0.9023911 | -0.148175254 | 0.20603695 | 1 |
| RGD1566265 | 0.89207591 | -0.164761618 | 0.20612267 | 1 |
| Ncdn | 0.94271878 | -0.085100627 | 0.20626528 | 1 |
| Grn | 0.93332326 | -0.099551242 | 0.20682101 | 1 |
| Ralgds | 0.9285251 | -0.106987192 | 0.20683337 | 1 |
| Gtf2b | 1.09087438 | 0.125484982 | 0.20685462 | 1 |
| Aff1 | 0.88937537 | -0.169135639 | 0.20685742 | 1 |
| Sema5b | 0.81150532 | -0.301327536 | 0.20709252 | 1 |
| Pth1r | 1.13052321 | 0.176990613 | 0.2074286 | 1 |
| Avpr1a | 0.76996717 | -0.37713117 | 0.2074592 | 1 |
| Susd3 | 1.32321463 | 0.404047096 | 0.20745934 | 1 |
| Acap2 | 0.92222971 | -0.116801955 | 0.20751577 | 1 |
| Zcchc4 | 1.20207985 | 0.26553273 | 0.20752789 | 1 |

| | | | | |
|---|---|---|---|---|
| Slc22a15 | 1.15529106 | 0.208256364 | 0.20765965 | 1 |
| Dlat | 0.93885085 | -0.091032113 | 0.20766202 | 1 |
| Tuft1 | 0.85986426 | -0.217819167 | 0.20791764 | 1 |
| Car6 | 1.37862168 | 0.463226605 | 0.20815397 | 1 |
| Aen | 0.86481593 | -0.209534998 | 0.20822638 | 1 |
| Colq | 0.804349 | -0.314106479 | 0.2084019 | 1 |
| Kcna1 | 0.94193712 | -0.086297338 | 0.20843151 | 1 |
| MGC95210 | 0.8462127 | -0.240907764 | 0.20848873 | 1 |
| Nxt1 | 0.83416892 | -0.261588538 | 0.20850558 | 1 |
| Ikbke | 1.1492015 | 0.200631779 | 0.20852449 | 1 |
| Pspc1 | 0.91398019 | -0.129765201 | 0.20853773 | 1 |
| Cndp2 | 1.06211775 | 0.086943713 | 0.20860777 | 1 |
| Cox20 | 0.89572904 | -0.158865718 | 0.2086095 | 1 |
| Anp32a | 1.0785715 | 0.109121823 | 0.20873163 | 1 |
| Pax9 | 1.22877332 | 0.297218802 | 0.20882754 | 1 |
| Ppp1r3d | 0.91241805 | -0.132233103 | 0.20883607 | 1 |
| Adcy7 | 0.90322308 | -0.146845741 | 0.20910297 | 1 |
| RGD1304884 | 0.94405283 | -0.0830605 | 0.2091571 | 1 |
| Sphkap | 1.07618911 | 0.105931614 | 0.20918823 | 1 |
| Epn2 | 0.94207578 | -0.086084988 | 0.20923384 | 1 |
| RGD1308117 | 0.85678834 | -0.222989244 | 0.2092437 | 1 |
| Fdxacb1 | 0.82511388 | -0.277334849 | 0.20925646 | 1 |
| Arhgef15 | 0.81209801 | -0.300274249 | 0.20970665 | 1 |
| Cabp7 | 1.18549458 | 0.24548907 | 0.20973183 | 1 |
| Fam189b | 0.94385485 | -0.083363086 | 0.20994192 | 1 |
| Tgoln2 | 1.05936519 | 0.083200003 | 0.21018073 | 1 |
| Arhgef37 | 0.89118833 | -0.166197759 | 0.21048034 | 1 |
| Emp2 | 0.92530105 | -0.112005273 | 0.21050567 | 1 |
| Mbd3 | 0.9195759 | -0.120959441 | 0.21067955 | 1 |
| Slc36a2 | 1.1069893 | 0.146641274 | 0.21097007 | 1 |
| Zdhhc3 | 1.07774621 | 0.108017491 | 0.21097401 | 1 |
| Hddc2 | 1.13092133 | 0.177498575 | 0.21122757 | 1 |
| A3galt2 | 0.94264479 | -0.085213864 | 0.21134243 | 1 |
| Me3 | 0.88698121 | -0.173024548 | 0.21139296 | 1 |
| Tmc7 | 0.90472455 | -0.144449468 | 0.21149889 | 1 |
| Dennd5a | 0.94447816 | -0.082410656 | 0.21167599 | 1 |
| Cacng7 | 0.91221159 | -0.132559598 | 0.21180239 | 1 |
| Tprn | 0.82654356 | -0.27483724 | 0.21190194 | 1 |
| Man2a1 | 0.92007148 | -0.120182144 | 0.21190421 | 1 |
| Glod4 | 1.06690887 | 0.093436958 | 0.21192642 | 1 |

| | | | | |
|---|---|---|---|---|
| Mrc1 | 1.08250865 | 0.114378547 | 0.21193117 | 1 |
| Slc2a10 | 0.79670415 | -0.327884006 | 0.21194985 | 1 |
| Nr2f1 | 0.91473883 | -0.128568195 | 0.21197306 | 1 |
| Abhd8 | 0.93491811 | -0.097088087 | 0.21197929 | 1 |
| Mrps17 | 1.10770551 | 0.147574384 | 0.21202705 | 1 |
| Slco1c1 | 0.86744491 | -0.205155956 | 0.21208101 | 1 |
| Plekha5 | 0.93487843 | -0.09714933 | 0.21229093 | 1 |
| Rnf113a2 | 0.86512103 | -0.209026112 | 0.21255217 | 1 |
| Ccng2 | 1.08728814 | 0.120734315 | 0.2127813 | 1 |
| Mxd4 | 0.90380499 | -0.145916578 | 0.21303133 | 1 |
| Meis2 | 1.11342846 | 0.155008867 | 0.21309116 | 1 |
| Bckdk | 1.08684475 | 0.120145875 | 0.21353408 | 1 |
| Tmem150c | 0.93930126 | -0.090340156 | 0.21356204 | 1 |
| Supt4h1 | 1.09058296 | 0.125099517 | 0.21385376 | 1 |
| Stc1 | 0.81167777 | -0.301020997 | 0.21392019 | 1 |
| Kcnip1 | 0.93353288 | -0.099227255 | 0.21392211 | 1 |
| Lrrc9 | 1.23317312 | 0.30237535 | 0.21396559 | 1 |
| Ubxn11 | 0.77655803 | -0.364834353 | 0.21428122 | 1 |
| Cradd | 1.2969011 | 0.375068463 | 0.21431392 | 1 |
| Uqcc1 | 1.08231808 | 0.11412455 | 0.21455289 | 1 |
| Ebpl | 1.14327099 | 0.193167404 | 0.21469714 | 1 |
| Sufu | 0.8242666 | -0.278817054 | 0.21481126 | 1 |
| Ciapin1 | 1.07411551 | 0.103149148 | 0.21482544 | 1 |
| Stat4 | 0.73479333 | -0.444589568 | 0.21484541 | 1 |
| Mccc1 | 1.11581027 | 0.15809173 | 0.21497935 | 1 |
| Mal | 0.94493732 | -0.081709465 | 0.21506353 | 1 |
| Clec4a3 | 1.19897266 | 0.261798759 | 0.21510725 | 1 |
| Tbck | 0.90726898 | -0.140397761 | 0.21516292 | 1 |
| RGD1305733 | 1.05991567 | 0.083949478 | 0.21522894 | 1 |
| Rnf208 | 0.9014793 | -0.149633733 | 0.21529841 | 1 |
| Ace | 1.14442793 | 0.194626615 | 0.21531331 | 1 |
| Glg1 | 0.94453702 | -0.082320757 | 0.21541594 | 1 |
| Arhgap26 | 0.83408652 | -0.261731045 | 0.21548792 | 1 |
| Ddx54 | 0.91760402 | -0.124056377 | 0.21550428 | 1 |
| Shc4 | 0.91476863 | -0.128521197 | 0.21570863 | 1 |
| Arhgef28 | 0.92464229 | -0.113032749 | 0.21577836 | 1 |
| LOC500077 | 1.21680356 | 0.283096278 | 0.21579109 | 1 |
| Tnfaip2 | 1.16734067 | 0.223225653 | 0.21595667 | 1 |
| Stx18 | 1.08672626 | 0.119988576 | 0.2160426 | 1 |
| Hs3st1 | 1.18948195 | 0.250333378 | 0.2164706 | 1 |

| | | | | |
|---|---|---|---|---|
| Oprl1 | 1.08091182 | 0.112248831 | 0.21674384 | 1 |
| Map7 | 0.93828964 | -0.091894761 | 0.2170744 | 1 |
| Nkr-p1c | 1.53115982 | 0.614624881 | 0.21727953 | 1 |
| Lamtor3 | 1.08649893 | 0.119686753 | 0.21746725 | 1 |
| Ptpn12 | 0.93134203 | -0.102617003 | 0.2175688 | 1 |
| Ndufs1 | 1.05832257 | 0.081779417 | 0.21760038 | 1 |
| Fam227b | 1.32481727 | 0.405793383 | 0.21769734 | 1 |
| Neu3 | 1.26720469 | 0.341649585 | 0.21772742 | 1 |
| Dnmt3b | 0.76423261 | -0.387916269 | 0.21773259 | 1 |
| Tmem205 | 1.09987644 | 0.13734146 | 0.21776562 | 1 |
| Rab38 | 1.21851933 | 0.285129133 | 0.21800395 | 1 |
| Dolpp1 | 0.91515351 | -0.127914331 | 0.21817049 | 1 |
| Arl3 | 1.07111177 | 0.099109034 | 0.21817269 | 1 |
| Cep95 | 1.11547203 | 0.157654341 | 0.21833468 | 1 |
| Pdzd7 | 1.1534701 | 0.205980613 | 0.21843716 | 1 |
| Trappc1 | 1.06502402 | 0.090885964 | 0.21851575 | 1 |
| Trpm2 | 0.89217727 | -0.164597699 | 0.2188761 | 1 |
| Akap7 | 0.80894542 | -0.305885727 | 0.21900833 | 1 |
| Ccnb1 | 0.84636025 | -0.240656221 | 0.21982073 | 1 |
| Slc22a17 | 0.94454137 | -0.082314113 | 0.21990856 | 1 |
| Plekhd1 | 0.93961634 | -0.089856298 | 0.21991667 | 1 |
| Eif4a2 | 1.06055358 | 0.084817505 | 0.22005055 | 1 |
| Cyb5r2 | 1.12403968 | 0.168692961 | 0.22008178 | 1 |
| Mefv | 1.25823714 | 0.331403847 | 0.22022324 | 1 |
| Tgm2 | 1.07752756 | 0.107724765 | 0.2203348 | 1 |
| Slc27a6 | 1.20401893 | 0.267858072 | 0.22035835 | 1 |
| Cdadc1 | 1.08559854 | 0.118490682 | 0.22044324 | 1 |
| Adck5 | 0.8813226 | -0.182257888 | 0.22055764 | 1 |
| Snrpn | 1.0785832 | 0.109137469 | 0.22097118 | 1 |
| E2f3 | 1.18009954 | 0.238908556 | 0.22119103 | 1 |
| Mal2 | 0.86348485 | -0.211757227 | 0.22134412 | 1 |
| Gimap7 | 0.79433119 | -0.332187442 | 0.22145341 | 1 |
| Cdk5r1 | 0.87037304 | -0.200294233 | 0.2215575 | 1 |
| Kif16b | 1.06817872 | 0.095153045 | 0.22161701 | 1 |
| Ndufaf6 | 1.1656966 | 0.221192346 | 0.22163569 | 1 |
| Taf2 | 0.92639849 | -0.110295198 | 0.22167031 | 1 |
| Lig3 | 1.0936214 | 0.129113379 | 0.22174727 | 1 |
| Rnf181 | 1.06764014 | 0.094425447 | 0.22180985 | 1 |
| Ppp4r1 | 0.92371716 | -0.114476929 | 0.22186564 | 1 |
| Arnt2 | 0.92356095 | -0.114720917 | 0.22199279 | 1 |

| | | | | |
|---|---|---|---|---|
| Zwint | 1.06071852 | 0.085041859 | 0.22210071 | 1 |
| Hars | 1.05958109 | 0.083493997 | 0.2222614 | 1 |
| Zbed3 | 1.20781329 | 0.272397455 | 0.22227974 | 1 |
| Hhip | 1.33561831 | 0.417507779 | 0.22247279 | 1 |
| Ccl21 | 0.89254844 | -0.163997619 | 0.22264498 | 1 |
| LOC308990 | 1.30923554 | 0.388724668 | 0.22266429 | 1 |
| Vdac1 | 1.05796089 | 0.081286292 | 0.22282452 | 1 |
| Pqlc3 | 1.06677972 | 0.093262302 | 0.22286078 | 1 |
| Hmx1 | 1.36560242 | 0.449537519 | 0.22324071 | 1 |
| Cd59 | 1.05840156 | 0.08188709 | 0.22340622 | 1 |
| Gucd1 | 0.83384268 | -0.262152883 | 0.22373316 | 1 |
| Rgs1 | 1.20321814 | 0.26689822 | 0.22389876 | 1 |
| Chml | 0.83446485 | -0.261076815 | 0.22395793 | 1 |
| Dqx1 | 0.83922776 | -0.252865692 | 0.22436525 | 1 |
| Vkorc1 | 0.90083467 | -0.15066575 | 0.22462995 | 1 |
| Mfap5 | 0.9318552 | -0.101822296 | 0.22495028 | 1 |
| Slc35a3 | 0.90214582 | -0.148567443 | 0.22496336 | 1 |
| Vwa5a | 1.05899756 | 0.082699268 | 0.22501666 | 1 |
| Baz1b | 0.93971479 | -0.089705146 | 0.22505841 | 1 |
| Pwwp2a | 1.11859672 | 0.161690009 | 0.22508138 | 1 |
| Cables1 | 0.89549276 | -0.159246333 | 0.22519935 | 1 |
| Fndc1 | 0.89620455 | -0.158100046 | 0.22520445 | 1 |
| RGD1560436 | 0.75690216 | -0.401821264 | 0.22529928 | 1 |
| Numb | 0.91125033 | -0.134080657 | 0.22551868 | 1 |
| Oprk1 | 0.87811075 | -0.187525181 | 0.22552015 | 1 |
| Slc45a3 | 0.81256823 | -0.299439144 | 0.22566431 | 1 |
| Prdx1 | 1.05975345 | 0.083728667 | 0.22567708 | 1 |
| Mthfs | 0.86169164 | -0.214756412 | 0.22571312 | 1 |
| Ufsp2 | 1.07512586 | 0.10450556 | 0.22616605 | 1 |
| Uggt1 | 0.93007159 | -0.104586322 | 0.22628245 | 1 |
| Cerk | 0.94207485 | -0.086086412 | 0.2263063 | 1 |
| Snx13 | 1.06904148 | 0.096317838 | 0.22640098 | 1 |
| Zbtb26 | 1.18898672 | 0.249732604 | 0.22647445 | 1 |
| Esm1 | 1.16373887 | 0.218767371 | 0.22653999 | 1 |
| Tvp23a | 0.89287645 | -0.163467536 | 0.22657054 | 1 |
| Casp4 | 1.1309485 | 0.177533236 | 0.22662335 | 1 |
| F2rl2 | 0.89578482 | -0.158775874 | 0.22665645 | 1 |
| Shtn1 | 0.91517202 | -0.127885151 | 0.22668277 | 1 |
| Slc35c1 | 0.90830936 | -0.138744349 | 0.22668516 | 1 |
| Blvrb | 1.11245488 | 0.153746822 | 0.22676983 | 1 |

| | | | | |
|---|---|---|---|---|
| Exosc8 | 1.19572213 | 0.25788217 | 0.22683635 | 1 |
| Spef2 | 1.42080014 | 0.506703632 | 0.22693779 | 1 |
| Rpl17 | 0.94344685 | -0.083986847 | 0.22708193 | 1 |
| Traf4 | 0.82379742 | -0.279638491 | 0.22716282 | 1 |
| Mfsd12 | 0.90972245 | -0.136501646 | 0.22717274 | 1 |
| Pan2 | 0.89598352 | -0.158455896 | 0.22717364 | 1 |
| St6galnac6 | 0.92723443 | -0.108993953 | 0.22718472 | 1 |
| Dus4l | 0.77229017 | -0.372785081 | 0.22719994 | 1 |
| Xpnpep1 | 0.93763392 | -0.092903331 | 0.2272311 | 1 |
| Rhoj | 1.0873701 | 0.120843067 | 0.22727414 | 1 |
| Mfng | 1.28819123 | 0.365346777 | 0.22735057 | 1 |
| Pla2g5 | 1.14807743 | 0.199219944 | 0.22736747 | 1 |
| Agl | 1.05758035 | 0.08076728 | 0.22756376 | 1 |
| Pold2 | 0.86552519 | -0.208352282 | 0.22773448 | 1 |
| Tlcd1 | 0.87103411 | -0.199198886 | 0.22800927 | 1 |
| PCOLCE2 | 1.09176553 | 0.126663058 | 0.22821064 | 1 |
| Tecpr1 | 0.93163214 | -0.102167691 | 0.22822932 | 1 |
| Tmem130 | 0.94635372 | -0.079548575 | 0.22835341 | 1 |
| Snx33 | 1.11029302 | 0.150940466 | 0.22840933 | 1 |
| Pon3 | 0.79095892 | -0.338325335 | 0.22844206 | 1 |
| Cdk5r2 | 0.93683067 | -0.094139788 | 0.22846228 | 1 |
| Csnk1g2 | 0.94094434 | -0.087818707 | 0.22850083 | 1 |
| RT1-A2 | 1.06757077 | 0.094331709 | 0.22858407 | 1 |
| Setd2 | 0.93696699 | -0.093929877 | 0.22859275 | 1 |
| Shank3 | 0.90627598 | -0.141977645 | 0.22859643 | 1 |
| Bcap31 | 1.05695827 | 0.079918423 | 0.22880995 | 1 |
| Ptgfrn | 0.93226984 | -0.101180506 | 0.22882524 | 1 |
| Wdr5 | 0.91250962 | -0.13208833 | 0.22891164 | 1 |
| Gna12 | 0.94321596 | -0.084339964 | 0.22905975 | 1 |
| Cog3 | 0.91682936 | -0.125274845 | 0.22908772 | 1 |
| Fyn | 0.9338093 | -0.098800143 | 0.22911156 | 1 |
| Gaa | 0.94465953 | -0.082133643 | 0.22916651 | 1 |
| Thg1l | 1.20189699 | 0.265313252 | 0.22934693 | 1 |
| Alox5ap | 1.1768757 | 0.234961956 | 0.22936219 | 1 |
| Rnf38 | 0.88007627 | -0.18429953 | 0.22936451 | 1 |
| Tmem9 | 0.92189789 | -0.117321136 | 0.22948523 | 1 |
| Cyp51 | 0.94587845 | -0.080273289 | 0.22955211 | 1 |
| Cep120 | 1.07251082 | 0.100992204 | 0.22960623 | 1 |
| Cox7b | 1.05869995 | 0.08229377 | 0.2297285 | 1 |
| Shc2 | 0.93519176 | -0.096665884 | 0.22972892 | 1 |

| | | | | |
|---|---|---|---|---|
| Cpa2 | 0.73282391 | -0.448461517 | 0.22994607 | 1 |
| Alad | 1.11304744 | 0.154515089 | 0.23000544 | 1 |
| Xrcc2 | 1.20941275 | 0.27430669 | 0.23004883 | 1 |
| Fcgr2b | 1.23638514 | 0.306128224 | 0.23005801 | 1 |
| Psen1 | 1.08968728 | 0.123914172 | 0.23008823 | 1 |
| Tnc | 0.92843549 | -0.107126423 | 0.23009419 | 1 |
| Noc4l | 1.12785798 | 0.173585412 | 0.23030426 | 1 |
| Upp1 | 0.77721079 | -0.363622161 | 0.23040882 | 1 |
| Rbbp9 | 1.16633291 | 0.221979638 | 0.23049902 | 1 |
| Lztr1 | 0.93788735 | -0.092513437 | 0.23054828 | 1 |
| Tjp1 | 1.05939922 | 0.083246356 | 0.23090475 | 1 |
| Ptpn14 | 0.85876456 | -0.219665442 | 0.23098451 | 1 |
| G3bp1 | 0.94363044 | -0.083706141 | 0.23116136 | 1 |
| Cnnm4 | 0.92420967 | -0.113707913 | 0.23132524 | 1 |
| Cops5 | 1.0582653 | 0.081701353 | 0.23145518 | 1 |
| RGD1305645 | 1.14334916 | 0.193266041 | 0.23154187 | 1 |
| Ankib1 | 0.93792545 | -0.092454843 | 0.23156733 | 1 |
| Rec114 | 1.36702905 | 0.451043897 | 0.23161121 | 1 |
| Pdgfc | 1.30677153 | 0.386006932 | 0.23178825 | 1 |
| Pcdhga10 | 0.90066972 | -0.150929936 | 0.23183635 | 1 |
| Frmd4a | 0.91240394 | -0.132255423 | 0.23193378 | 1 |
| Tank | 1.0977323 | 0.134526273 | 0.23202504 | 1 |
| Mroh1 | 0.88317977 | -0.179220977 | 0.2320832 | 1 |
| Smim8 | 1.1164931 | 0.158974338 | 0.23223752 | 1 |
| Rgl3 | 1.23225243 | 0.30129782 | 0.23230083 | 1 |
| Nmb | 1.08236428 | 0.114186136 | 0.23243064 | 1 |
| Serpini1 | 1.06628906 | 0.092598593 | 0.23245788 | 1 |
| Slc25a18 | 0.76761721 | -0.381541044 | 0.23249908 | 1 |
| Spata7 | 1.11334839 | 0.154905112 | 0.23262292 | 1 |
| Dpy19l4 | 0.91790532 | -0.12358274 | 0.23269045 | 1 |
| Dcp1a | 1.20438176 | 0.268292758 | 0.23273506 | 1 |
| Dact2 | 1.12619781 | 0.171460248 | 0.23274623 | 1 |
| Ylpm1 | 0.93222038 | -0.101257037 | 0.23293496 | 1 |
| Ntm | 1.11796379 | 0.160873456 | 0.2330403 | 1 |
| Agap1 | 0.92902722 | -0.106207226 | 0.23330773 | 1 |
| Doc2a | 1.12920686 | 0.175309794 | 0.23332212 | 1 |
| Rptor | 0.93596327 | -0.095476182 | 0.23337197 | 1 |
| C1galt1 | 0.8782628 | -0.187275405 | 0.23342562 | 1 |
| Sez6 | 0.90803727 | -0.139176581 | 0.23352781 | 1 |
| Ahr | 0.91667026 | -0.125525234 | 0.23357125 | 1 |

| | | | | |
|---|---|---|---|---|
| Fah | 1.15739042 | 0.210875604 | 0.23365539 | 1 |
| Enah | 0.93426494 | -0.098096366 | 0.23367844 | 1 |
| Bcor | 1.11106023 | 0.151937024 | 0.23368193 | 1 |
| Lims1 | 1.09098265 | 0.125628152 | 0.23369447 | 1 |
| Gab2 | 0.9072853 | -0.140371813 | 0.23369845 | 1 |
| Dcun1d3 | 0.88100596 | -0.182776313 | 0.23373201 | 1 |
| Nell1 | 0.94066924 | -0.088240559 | 0.23378156 | 1 |
| Gba2 | 1.07942423 | 0.110261977 | 0.23402222 | 1 |
| Abhd14a | 1.13787437 | 0.186341276 | 0.23407577 | 1 |
| Fbxo22 | 1.08281454 | 0.114786166 | 0.23410617 | 1 |
| Rnf10 | 0.94647449 | -0.079364479 | 0.23433171 | 1 |
| Mrps22 | 1.09137354 | 0.126144965 | 0.23438709 | 1 |
| Actn3 | 0.77538478 | -0.36701568 | 0.23438926 | 1 |
| Ccs | 0.904088 | -0.145464889 | 0.23443348 | 1 |
| Ifitm1 | 1.12263061 | 0.166883307 | 0.2344462 | 1 |
| Sgtb | 1.06856947 | 0.095680698 | 0.23455457 | 1 |
| Mt2A | 1.19019157 | 0.251193804 | 0.23464756 | 1 |
| Csf1 | 1.09431648 | 0.130030031 | 0.2346868 | 1 |
| Klf5 | 1.07914515 | 0.109888923 | 0.23471195 | 1 |
| Tgds | 1.1238159 | 0.168405723 | 0.23472125 | 1 |
| B4galt4 | 0.91157428 | -0.133567875 | 0.23488549 | 1 |
| Txlng | 1.18892144 | 0.24965339 | 0.23492427 | 1 |
| Dhcr7 | 0.92636873 | -0.110341537 | 0.23492905 | 1 |
| Top1mt | 0.88140587 | -0.182121593 | 0.23506678 | 1 |
| Rars2 | 0.85871356 | -0.219751127 | 0.23566854 | 1 |
| Toe1 | 1.20842658 | 0.273129821 | 0.23566934 | 1 |
| Man1c1 | 1.10784952 | 0.147761931 | 0.23585547 | 1 |
| Lingo1 | 0.93209756 | -0.101447126 | 0.23588738 | 1 |
| Uhrf1 | 0.7521127 | -0.410979237 | 0.23591155 | 1 |
| Ndufaf7 | 1.08091729 | 0.112256134 | 0.23603189 | 1 |
| Cenpj | 1.09711357 | 0.133712882 | 0.2360413 | 1 |
| Cdh6 | 0.91630732 | -0.126096555 | 0.23606934 | 1 |
| Scrn2 | 1.19104979 | 0.252233726 | 0.23608391 | 1 |
| Tbxas1 | 1.2301069 | 0.298783701 | 0.23632188 | 1 |
| Sfxn1 | 0.94085516 | -0.087955458 | 0.23632966 | 1 |
| Tnfrsf11a | 1.19990458 | 0.262919688 | 0.23638159 | 1 |
| Slco1a2 | 1.07265935 | 0.10119198 | 0.23650646 | 1 |
| Nmd3 | 1.09867652 | 0.135766687 | 0.2369726 | 1 |
| Tfg | 0.94576991 | -0.080438845 | 0.23707031 | 1 |
| Nek6 | 0.92750557 | -0.108572147 | 0.23711171 | 1 |

| | | | | |
|---|---|---|---|---|
| Phtf1 | 1.06072273 | 0.085047587 | 0.2371969 | 1 |
| Limd1 | 1.09616866 | 0.132469793 | 0.23751838 | 1 |
| Itgb3 | 0.8407581 | -0.250237328 | 0.23763321 | 1 |
| Nod1 | 1.09741545 | 0.134109794 | 0.23809398 | 1 |
| Atp5h | 1.05751246 | 0.080674662 | 0.23821386 | 1 |
| Als2cr12 | 1.33675804 | 0.418738359 | 0.23821793 | 1 |
| Hexim2 | 1.15874793 | 0.212566762 | 0.23842472 | 1 |
| Smc1a | 1.05686159 | 0.079786442 | 0.23851591 | 1 |
| Pde4c | 1.17072703 | 0.227404734 | 0.23855082 | 1 |
| Smox | 1.0966534 | 0.133107628 | 0.23866686 | 1 |
| Sbno2 | 0.90824813 | -0.13884161 | 0.23867727 | 1 |
| Gda | 1.06304368 | 0.08820088 | 0.23871124 | 1 |
| Stxbp6 | 1.19207877 | 0.253479567 | 0.23885822 | 1 |
| Rom1 | 1.14384104 | 0.193886579 | 0.23891467 | 1 |
| Pparg | 1.2192594 | 0.286005089 | 0.23902822 | 1 |
| Tmem170a | 1.17801197 | 0.236354205 | 0.23904061 | 1 |
| Cox4i1 | 1.05631827 | 0.079044588 | 0.23919396 | 1 |
| Zbtb11 | 0.93467355 | -0.097465526 | 0.23920178 | 1 |
| Flrt3 | 1.09772812 | 0.134520776 | 0.23928704 | 1 |
| Cyp4b1 | 1.33288161 | 0.414548638 | 0.23931083 | 1 |
| Spc25 | 1.32900526 | 0.410346811 | 0.2393848 | 1 |
| Pld1 | 1.08533235 | 0.118136888 | 0.23956122 | 1 |
| Shd | 1.10863212 | 0.148780708 | 0.23957108 | 1 |
| Atp5j | 1.0575461 | 0.080720559 | 0.239606 | 1 |
| Peli1 | 0.90056263 | -0.151101486 | 0.23968733 | 1 |
| Rab22a | 0.9282468 | -0.107419659 | 0.2398072 | 1 |
| Med13 | 0.92349461 | -0.114824553 | 0.23982054 | 1 |
| Pcsk6 | 0.920926 | -0.118842866 | 0.23989588 | 1 |
| Scn4a | 0.87745835 | -0.188597448 | 0.2399245 | 1 |
| RGD1308428 | 0.93366566 | -0.099022072 | 0.2400883 | 1 |
| Tmem208 | 1.10361586 | 0.142238099 | 0.24008894 | 1 |
| Sumf2 | 0.84943491 | -0.235424701 | 0.24012107 | 1 |
| Txlna | 0.92807552 | -0.107685891 | 0.24015271 | 1 |
| Syk | 1.16102968 | 0.21540485 | 0.2402279 | 1 |
| Icam2 | 1.17588147 | 0.233742649 | 0.24027295 | 1 |
| Itpa | 1.0721805 | 0.1005478 | 0.24028002 | 1 |
| Lsg1 | 1.09418263 | 0.129853554 | 0.24036104 | 1 |
| Gldn | 0.92267645 | -0.116103262 | 0.24038427 | 1 |
| Akr1a1 | 1.05637445 | 0.079121309 | 0.24040286 | 1 |
| Tmem183a | 1.06735248 | 0.094036693 | 0.24042123 | 1 |

| | | | | |
|---|---|---|---|---|
| Scrib | 0.92201545 | -0.117137162 | 0.24051146 | 1 |
| Snhg11 | 1.05899838 | 0.082700384 | 0.24059817 | 1 |
| Rabep1 | 1.05749852 | 0.080655646 | 0.24082083 | 1 |
| Tmem237 | 1.08814646 | 0.121872751 | 0.24100632 | 1 |
| Eno3 | 0.85776208 | -0.221350559 | 0.24103593 | 1 |
| Klc4 | 1.08925447 | 0.123341033 | 0.24122641 | 1 |
| Abhd10 | 1.09703799 | 0.133613483 | 0.24125921 | 1 |
| Itih3 | 1.31785247 | 0.398188873 | 0.24139854 | 1 |
| Pald1 | 1.1003269 | 0.137932199 | 0.24146321 | 1 |
| Myo9b | 0.94165656 | -0.086727119 | 0.24170636 | 1 |
| Bcas3 | 0.9344044 | -0.097881025 | 0.24197164 | 1 |
| Ikbkap | 0.93584991 | -0.095650928 | 0.24201071 | 1 |
| Scyl3 | 1.10876702 | 0.148956245 | 0.24218723 | 1 |
| Elmod3 | 0.80741424 | -0.308619058 | 0.24265976 | 1 |
| Psma5 | 1.06181316 | 0.086529929 | 0.24269516 | 1 |
| Fmo5 | 0.80667846 | -0.30993436 | 0.24275889 | 1 |
| Rpl6 | 1.05524936 | 0.077583956 | 0.24284824 | 1 |
| Slc7a4 | 1.14337961 | 0.193304463 | 0.24285987 | 1 |
| Fnbp4 | 1.07770128 | 0.107957348 | 0.24302383 | 1 |
| Arpp21 | 1.08853108 | 0.122382595 | 0.24305415 | 1 |
| Rmdn2 | 1.14088536 | 0.190153835 | 0.24306213 | 1 |
| Dars2 | 1.11746669 | 0.160231824 | 0.24311412 | 1 |
| Cmah | 1.37835671 | 0.462949293 | 0.24336019 | 1 |
| Flt1 | 0.92981775 | -0.104980125 | 0.24368332 | 1 |
| Abcg3l4 | 0.90036678 | -0.151415263 | 0.24371373 | 1 |
| Taf5 | 0.85313001 | -0.229162477 | 0.2439268 | 1 |
| Lgr4 | 0.93712079 | -0.093693082 | 0.2440084 | 1 |
| Ctla2a | 1.23427434 | 0.3036631 | 0.24415736 | 1 |
| Pafah2 | 0.79119572 | -0.33789347 | 0.24422344 | 1 |
| Wif1 | 0.83779298 | -0.255334304 | 0.24424372 | 1 |
| Sh3bgrl2 | 1.13816226 | 0.186706249 | 0.2443943 | 1 |
| Kcnk6 | 1.22203363 | 0.289283984 | 0.24439941 | 1 |
| Ccl2 | 0.88708597 | -0.172854172 | 0.24443114 | 1 |
| Myh2 | 0.71615158 | -0.481663106 | 0.24445688 | 1 |
| Pnoc | 1.34323835 | 0.425715323 | 0.24461265 | 1 |
| Prelp | 1.07120117 | 0.099229436 | 0.24475896 | 1 |
| Yif1a | 1.09513009 | 0.131102255 | 0.2451917 | 1 |
| Ndufs4 | 1.06121866 | 0.085721942 | 0.24520825 | 1 |
| Abhd6 | 1.07678148 | 0.106725499 | 0.24532638 | 1 |
| Pld3 | 1.05643832 | 0.079208542 | 0.24535361 | 1 |

| | | | | |
|---|---|---|---|---|
| Dhx9 | 0.94543764 | -0.080945787 | 0.24536693 | 1 |
| Mgat5b | 0.8535429 | -0.228464424 | 0.24540599 | 1 |
| Brf2 | 1.12465835 | 0.16948681 | 0.24542981 | 1 |
| Fam222a | 0.80133692 | -0.31951914 | 0.24545406 | 1 |
| Ppif | 1.13432703 | 0.181836638 | 0.24547122 | 1 |
| Tdrd1 | 0.74679244 | -0.421220765 | 0.24556431 | 1 |
| Alox5 | 1.22800722 | 0.296319042 | 0.24569548 | 1 |
| Cmtm5 | 0.92513005 | -0.112271905 | 0.24579698 | 1 |
| Pfn2 | 1.05579744 | 0.078333079 | 0.24583885 | 1 |
| Slc35f5 | 0.92547551 | -0.111733282 | 0.24608538 | 1 |
| Adhfe1 | 1.26662343 | 0.340987674 | 0.24610532 | 1 |
| Dnajc22 | 1.21716077 | 0.283519741 | 0.24617505 | 1 |
| Cript | 1.06303684 | 0.088191595 | 0.24617877 | 1 |
| RGD1559747 | 0.88092796 | -0.182904046 | 0.24628951 | 1 |
| Setdb1 | 1.0765904 | 0.10646947 | 0.24656797 | 1 |
| Popdc3 | 0.90099479 | -0.150409338 | 0.24669502 | 1 |
| Chrdl1 | 1.10896347 | 0.149211844 | 0.24673743 | 1 |
| Adra2c | 0.88057668 | -0.18347945 | 0.24674509 | 1 |
| Prdx4 | 0.92114272 | -0.1185034 | 0.24681368 | 1 |
| Grap2 | 1.30387749 | 0.382808325 | 0.2468827 | 1 |
| Amn1 | 1.10090987 | 0.138696357 | 0.24697861 | 1 |
| Slc1a3 | 1.08927481 | 0.123367969 | 0.24701987 | 1 |
| Ctdspl | 0.91494954 | -0.128235914 | 0.24704474 | 1 |
| Ddx25 | 1.07610551 | 0.105819544 | 0.24706636 | 1 |
| Gnb4 | 0.92095706 | -0.1187942 | 0.24711576 | 1 |
| Abl2 | 1.13763897 | 0.186042795 | 0.24729465 | 1 |
| Bub3 | 1.08567827 | 0.118596643 | 0.2474648 | 1 |
| Cep162 | 0.88888199 | -0.169936195 | 0.2476235 | 1 |
| Ltbp1 | 0.92039723 | -0.119671451 | 0.24774034 | 1 |
| 11-Mar | 0.87667063 | -0.189893183 | 0.24800656 | 1 |
| Ruvbl2 | 1.08890745 | 0.122881346 | 0.24830972 | 1 |
| Tnfsf10 | 1.26160785 | 0.335263539 | 0.24834367 | 1 |
| Ngdn | 0.90788636 | -0.139416371 | 0.24839087 | 1 |
| Resp18 | 1.06026423 | 0.084423845 | 0.24846811 | 1 |
| Cst7 | 1.49357947 | 0.578773999 | 0.24852044 | 1 |
| Uaca | 1.06463414 | 0.090357733 | 0.24860186 | 1 |
| Pidd1 | 0.82378704 | -0.279656662 | 0.24867648 | 1 |
| Tnip1 | 1.07272922 | 0.101285951 | 0.24886921 | 1 |
| B3galt2 | 1.12575329 | 0.17089069 | 0.2490069 | 1 |
| RT1-DMa | 1.1357121 | 0.183597163 | 0.24915519 | 1 |

| | | | | |
|---|---|---|---|---|
| Hist3h2ba | 1.11004173 | 0.150613918 | 0.24916045 | 1 |
| Rnf150 | 0.87264732 | -0.196529384 | 0.24922667 | 1 |
| Mb21d2 | 1.14406708 | 0.194171646 | 0.24924239 | 1 |
| Abca7 | 0.94693533 | -0.078662189 | 0.24937469 | 1 |
| Acadvl | 1.06205789 | 0.086862405 | 0.2494081 | 1 |
| Phc2 | 0.93892455 | -0.090918861 | 0.24943791 | 1 |
| Ndufs5 | 1.06107892 | 0.08553197 | 0.24968506 | 1 |
| Sh3bp5 | 1.08498584 | 0.117676221 | 0.24985994 | 1 |
| Plekhf2 | 0.90190376 | -0.1489546 | 0.24994582 | 1 |
| Ccna2 | 0.77184451 | -0.373617844 | 0.24999214 | 1 |
| Wdr3 | 1.09998492 | 0.13748375 | 0.25000093 | 1 |
| Kif26b | 0.89629664 | -0.157951804 | 0.25012261 | 1 |
| Galnt9 | 0.91493903 | -0.12825249 | 0.25013128 | 1 |
| Smarcd1 | 0.92856377 | -0.106927108 | 0.25016742 | 1 |
| Zfp292 | 1.07534972 | 0.104805926 | 0.25016905 | 1 |
| Tfcp2 | 0.90858259 | -0.138310438 | 0.25017854 | 1 |
| Clta | 1.05443275 | 0.076467084 | 0.2503734 | 1 |
| Wdr47 | 1.05528977 | 0.077639197 | 0.25044458 | 1 |
| Pkm | 1.05632644 | 0.079055745 | 0.25045299 | 1 |
| Cox7c | 1.05425227 | 0.076220133 | 0.25050671 | 1 |
| Lrrc16b | 0.93176792 | -0.101957428 | 0.25059936 | 1 |
| Msc | 0.76985497 | -0.377341409 | 0.2506912 | 1 |
| Bola1 | 0.86877052 | -0.20295295 | 0.25096556 | 1 |
| Rims1 | 0.92742286 | -0.108700803 | 0.25107533 | 1 |
| Mfn1 | 1.07246905 | 0.100936011 | 0.25110602 | 1 |
| Mettl7a | 1.09008739 | 0.124443801 | 0.25119541 | 1 |
| Gpr81 | 1.2704397 | 0.345327901 | 0.25123372 | 1 |
| Zfp133 | 1.11594293 | 0.158263251 | 0.25132405 | 1 |
| Gpd1l | 1.10624755 | 0.145674266 | 0.25152555 | 1 |
| Sos1 | 0.94635876 | -0.079540896 | 0.25170962 | 1 |
| Wnk3 | 1.24419809 | 0.315216196 | 0.25176446 | 1 |
| Cpd | 0.94421525 | -0.082812304 | 0.25227894 | 1 |
| Rmdn1 | 1.11596291 | 0.158289085 | 0.25231453 | 1 |
| Fcrla | 0.77450528 | -0.368653024 | 0.25262367 | 1 |
| Rhbdd1 | 0.81797607 | -0.289869462 | 0.25274315 | 1 |
| Ganc | 1.09676283 | 0.13325158 | 0.2527622 | 1 |
| Pragmin | 0.85690196 | -0.222797946 | 0.25288825 | 1 |
| Srm | 0.90843921 | -0.138538124 | 0.25291817 | 1 |
| Stxbp1 | 0.94719514 | -0.078266422 | 0.25294343 | 1 |
| RGD1309808 | 1.34084009 | 0.423137194 | 0.25299522 | 1 |

| | | | | |
|---|---|---|---|---|
| Timm23 | 1.06641338 | 0.092766784 | 0.25330806 | 1 |
| Parvb | 0.9422726 | -0.085783596 | 0.25330913 | 1 |
| Adrbk1 | 0.94061273 | -0.088327237 | 0.2533435 | 1 |
| Sap30bp | 1.08042478 | 0.111598637 | 0.25337978 | 1 |
| Acvr1b | 0.91839606 | -0.122811639 | 0.2533875 | 1 |
| Colec12 | 1.08349103 | 0.115687215 | 0.25343664 | 1 |
| Pja1 | 0.94906497 | -0.075421236 | 0.25372395 | 1 |
| Myh8 | 1.20588515 | 0.270092515 | 0.25378396 | 1 |
| Atp1a1 | 0.94545231 | -0.080923401 | 0.25378605 | 1 |
| Atp6v1c2 | 1.30906945 | 0.388541644 | 0.25399085 | 1 |
| Rnf13 | 1.05382661 | 0.075637513 | 0.25407954 | 1 |
| Eif3j | 1.05651288 | 0.079310359 | 0.25418711 | 1 |
| Scamp5 | 0.92566691 | -0.111434938 | 0.25459075 | 1 |
| Col6a2 | 1.05632972 | 0.079060227 | 0.25470439 | 1 |
| Nit2 | 1.12247979 | 0.166689475 | 0.25476298 | 1 |
| Cd34 | 1.08715305 | 0.120555059 | 0.25477817 | 1 |
| B9d2 | 1.1552755 | 0.208236931 | 0.25480392 | 1 |
| Art3 | 0.94698789 | -0.078582113 | 0.25497417 | 1 |
| B4galt3 | 1.10029079 | 0.137884852 | 0.2550232 | 1 |
| Abcb1b | 1.29314496 | 0.370884011 | 0.25522611 | 1 |
| Pdk3 | 1.07357875 | 0.102428016 | 0.25528727 | 1 |
| Zfp61 | 1.10651771 | 0.146026544 | 0.25532183 | 1 |
| Lca5 | 0.84933548 | -0.235593577 | 0.25539414 | 1 |
| Zcchc24 | 0.94337874 | -0.084091012 | 0.25540112 | 1 |
| Chfr | 1.07233868 | 0.100760635 | 0.25567356 | 1 |
| Lrfn3 | 0.84197697 | -0.248147316 | 0.25588968 | 1 |
| Tfip11 | 1.0754863 | 0.104989146 | 0.25605324 | 1 |
| RGD1306746 | 1.09797629 | 0.134846898 | 0.25615947 | 1 |
| Cald1 | 1.05527772 | 0.077622723 | 0.25631902 | 1 |
| Upf3b | 1.0820086 | 0.113711964 | 0.25647448 | 1 |
| Kxd1 | 0.92918355 | -0.105964488 | 0.25651238 | 1 |
| Sorcs3 | 1.08287298 | 0.114864031 | 0.2566018 | 1 |
| Sclt1 | 1.12923485 | 0.175345556 | 0.25664574 | 1 |
| Ss18 | 0.93185539 | -0.101822012 | 0.25664733 | 1 |
| Wbscr27 | 0.80483299 | -0.313238649 | 0.25673538 | 1 |
| Mark4 | 0.93278955 | -0.100376461 | 0.25685576 | 1 |
| RGD1559482 | 1.18459095 | 0.244388967 | 0.25691206 | 1 |
| LOC100125367 | 1.15140104 | 0.203390418 | 0.25692287 | 1 |
| Hoxc8 | 1.2918457 | 0.369433761 | 0.25698216 | 1 |
| Cntf | 1.06130117 | 0.085834109 | 0.25701719 | 1 |

| | | | | |
|---|---|---|---|---|
| Dhrsx | 0.86462598 | -0.209851913 | 0.25724919 | 1 |
| Adgre5 | 0.90446278 | -0.144866962 | 0.25724938 | 1 |
| Tnks2 | 0.94569719 | -0.080549791 | 0.25729143 | 1 |
| Uxt | 0.88314001 | -0.179285926 | 0.25735832 | 1 |
| D2hgdh | 0.88944126 | -0.169028769 | 0.25747008 | 1 |
| Gnal | 0.94903252 | -0.07547057 | 0.2575348 | 1 |
| Ttc39c | 1.07815178 | 0.108560297 | 0.25757201 | 1 |
| Kdelc2 | 1.06683242 | 0.093333576 | 0.25782614 | 1 |
| Pigv | 0.86771268 | -0.20471069 | 0.25785383 | 1 |
| Creb5 | 1.16351181 | 0.218485851 | 0.25790972 | 1 |
| Dnajc11 | 1.066831 | 0.093331658 | 0.25799612 | 1 |
| Cercam | 0.87096616 | -0.199311435 | 0.25803817 | 1 |
| Rasip1 | 0.88880731 | -0.170057419 | 0.25822445 | 1 |
| Tmem134 | 1.10319278 | 0.141684919 | 0.25865383 | 1 |
| Rps6ka1 | 0.909595 | -0.136703768 | 0.25868986 | 1 |
| Fbxo42 | 0.89519202 | -0.159730922 | 0.25870228 | 1 |
| Psmb3 | 1.05817959 | 0.081584503 | 0.25889577 | 1 |
| Nkiras1 | 1.05916542 | 0.08292793 | 0.25921481 | 1 |
| Ctss | 1.07633311 | 0.106124639 | 0.25951029 | 1 |
| Pja2 | 1.05558211 | 0.078038802 | 0.25954669 | 1 |
| Mmp24 | 0.93121127 | -0.102819578 | 0.25961441 | 1 |
| Slc23a2 | 0.94602462 | -0.080050363 | 0.25996011 | 1 |
| Gpr155 | 0.93358331 | -0.099149329 | 0.25997378 | 1 |
| Hibadh | 1.06897386 | 0.096226579 | 0.2600826 | 1 |
| Api5 | 1.0544387 | 0.076475231 | 0.26009385 | 1 |
| Zfp90 | 1.22071044 | 0.287721029 | 0.2602511 | 1 |
| Fosb | 0.81858935 | -0.2887882 | 0.26029592 | 1 |
| Sod1 | 1.05292368 | 0.074400871 | 0.26029841 | 1 |
| Pld4 | 1.12853421 | 0.174450149 | 0.26032135 | 1 |
| Emid1 | 0.86102204 | -0.215877929 | 0.26047864 | 1 |
| Abcb7 | 1.11212745 | 0.15332213 | 0.26048128 | 1 |
| S100a16 | 0.91360859 | -0.130351885 | 0.26051884 | 1 |
| Trex1 | 0.87902396 | -0.186025605 | 0.26053553 | 1 |
| Ptpn5 | 0.8843769 | -0.177266745 | 0.26059446 | 1 |
| Cpne7 | 0.80871383 | -0.306298814 | 0.26065866 | 1 |
| Fcer1g | 1.12104156 | 0.164839765 | 0.26071567 | 1 |
| Flt3 | 1.07856359 | 0.10911124 | 0.2607909 | 1 |
| Fam60a | 1.15639554 | 0.209634953 | 0.26089488 | 1 |
| Ptprg | 0.91576799 | -0.126945955 | 0.26103447 | 1 |
| Tob1 | 1.09125736 | 0.125991386 | 0.26112893 | 1 |

| | | | | |
|---|---|---|---|---|
| Cox18 | 0.88964665 | -0.168695653 | 0.26119436 | 1 |
| Camta1 | 0.94935505 | -0.074980356 | 0.26138592 | 1 |
| Gzmm | 1.21565441 | 0.281733158 | 0.26139917 | 1 |
| Ago3 | 0.88936878 | -0.169146341 | 0.2614863 | 1 |
| Wbp1 | 0.92211421 | -0.116982642 | 0.26149601 | 1 |
| Slc29a2 | 1.15077604 | 0.202607089 | 0.26167829 | 1 |
| Dbi | 0.94967186 | -0.074498996 | 0.26174109 | 1 |
| Ttc27 | 1.0834875 | 0.115682516 | 0.26185595 | 1 |
| Mstn | 0.8161161 | -0.293153684 | 0.2619374 | 1 |
| Pcdh8 | 1.26439072 | 0.338442348 | 0.26206918 | 1 |
| Pgm2 | 1.1015843 | 0.139579901 | 0.26220007 | 1 |
| Gspt2 | 1.110239 | 0.150870272 | 0.26221572 | 1 |
| Rps26 | 1.05554876 | 0.077993219 | 0.26233822 | 1 |
| Nupr1 | 1.06633784 | 0.092664594 | 0.26250147 | 1 |
| Tmem230 | 1.05982858 | 0.083830936 | 0.26265721 | 1 |
| Exosc2 | 0.87543908 | -0.191921306 | 0.26271459 | 1 |
| Nlgn1 | 1.10224298 | 0.14044229 | 0.26332795 | 1 |
| Fastkd3 | 1.10951593 | 0.149930376 | 0.26334051 | 1 |
| Galnt7 | 0.84952394 | -0.235273486 | 0.26334491 | 1 |
| Lurap1 | 1.17372205 | 0.231090802 | 0.26340173 | 1 |
| Nalcn | 0.9480448 | -0.076972861 | 0.26350152 | 1 |
| RGD1305713 | 1.12034234 | 0.163939642 | 0.26358164 | 1 |
| Ubn1 | 0.92887248 | -0.106447549 | 0.26369639 | 1 |
| Snx12 | 1.10776337 | 0.147649737 | 0.26382427 | 1 |
| Fbxl17 | 0.91061197 | -0.135091668 | 0.26387422 | 1 |
| Dnajb14 | 1.09008122 | 0.12443563 | 0.26392341 | 1 |
| RT1-CE6 | 1.32519399 | 0.40620357 | 0.26416627 | 1 |
| Abcc1 | 0.93637134 | -0.09484731 | 0.26423045 | 1 |
| Eif1a | 1.06052698 | 0.084781323 | 0.26428829 | 1 |
| Ap4s1 | 1.11641451 | 0.158872782 | 0.26440463 | 1 |
| Adat3 | 0.8087335 | -0.30626373 | 0.26474564 | 1 |
| Rnf111 | 0.930456 | -0.103990163 | 0.26490196 | 1 |
| Eef1b2 | 1.05491817 | 0.077131093 | 0.26505348 | 1 |
| Nlgn2 | 0.9359525 | -0.095492786 | 0.26511917 | 1 |
| Tnik | 0.92417004 | -0.113769781 | 0.2651262 | 1 |
| Ptprj | 1.06910717 | 0.096406481 | 0.26514492 | 1 |
| Ercc8 | 0.91490245 | -0.128310173 | 0.26515428 | 1 |
| Rttn | 0.8542429 | -0.227281744 | 0.26515815 | 1 |
| Ntrk3 | 0.91572301 | -0.127016821 | 0.2652788 | 1 |
| Fam129c | 0.73528806 | -0.443618532 | 0.26539426 | 1 |

| | | | | |
|---|---|---|---|---|
| Ociad1 | 1.05235988 | 0.07362815 | 0.26554986 | 1 |
| Ahcyl1 | 0.94979099 | -0.074318023 | 0.26564479 | 1 |
| Wdr18 | 1.09325782 | 0.128633662 | 0.2657373 | 1 |
| Gpr156 | 0.84962267 | -0.235105841 | 0.26580083 | 1 |
| Nin | 1.09846221 | 0.135485238 | 0.26580558 | 1 |
| Ptgr1 | 1.18578538 | 0.24584292 | 0.26605859 | 1 |
| Ndufv2 | 1.05365617 | 0.075404167 | 0.26608963 | 1 |
| Tomm20 | 1.05467669 | 0.076800815 | 0.26627546 | 1 |
| Sh3gl1 | 0.92817768 | -0.107527085 | 0.26651447 | 1 |
| Htr4 | 1.18540772 | 0.245383361 | 0.26667428 | 1 |
| Afmid | 0.78255365 | -0.353738433 | 0.26710103 | 1 |
| Slc25a44 | 0.92629195 | -0.110461122 | 0.26712607 | 1 |
| Ulk2 | 0.94824478 | -0.076668576 | 0.2677144 | 1 |
| Nfe2 | 1.27679777 | 0.35253004 | 0.2678399 | 1 |
| Mfsd5 | 1.08207441 | 0.113799705 | 0.2678646 | 1 |
| Pid1 | 1.15775321 | 0.211327757 | 0.26795962 | 1 |
| Ikzf3 | 1.39571146 | 0.481000718 | 0.26812223 | 1 |
| Ccny | 0.93209308 | -0.101454057 | 0.26826284 | 1 |
| Igf2 | 0.93823733 | -0.091975186 | 0.26838242 | 1 |
| Fto | 1.07544902 | 0.104939132 | 0.26848796 | 1 |
| Ocln | 1.15314631 | 0.205575576 | 0.26852391 | 1 |
| Sntb2 | 0.83017274 | -0.268516539 | 0.26854371 | 1 |
| Aff4 | 0.95012251 | -0.073814552 | 0.26854883 | 1 |
| Gnl1 | 0.93567448 | -0.095921391 | 0.26865573 | 1 |
| Pdk4 | 0.87215212 | -0.197348306 | 0.26875217 | 1 |
| Wwox | 1.15683953 | 0.210188754 | 0.26879695 | 1 |
| Pex14 | 0.93098123 | -0.103176021 | 0.26882782 | 1 |
| Prg4 | 1.08839248 | 0.122198888 | 0.26884388 | 1 |
| Pkib | 1.09423023 | 0.129916313 | 0.26900399 | 1 |
| Egr3 | 0.7894103 | -0.341152761 | 0.26919414 | 1 |
| Pcsk1n | 0.94985898 | -0.074214756 | 0.26922307 | 1 |
| Hsd17b10 | 1.07332183 | 0.102082732 | 0.26929569 | 1 |
| Psma3l | 1.07693907 | 0.106936632 | 0.26931716 | 1 |
| Agpat4 | 1.06908136 | 0.096371644 | 0.26951063 | 1 |
| Scfd2 | 1.09698413 | 0.133542656 | 0.2698737 | 1 |
| Tmem159 | 0.90406555 | -0.145500712 | 0.27002787 | 1 |
| Slc22a18 | 1.21323768 | 0.278862207 | 0.27003252 | 1 |
| Gpn3 | 1.09677562 | 0.133268401 | 0.2701604 | 1 |
| Slc4a11 | 1.104332 | 0.14317396 | 0.27016884 | 1 |
| Scaf4 | 1.08762876 | 0.121186206 | 0.2702862 | 1 |

| | | | | |
|---|---|---|---|---|
| Slc24a1 | 1.16835874 | 0.224483315 | 0.27040688 | 1 |
| Chp2 | 0.91215345 | -0.132651543 | 0.27060821 | 1 |
| Strn4 | 0.93374746 | -0.098895684 | 0.2707324 | 1 |
| Crcp | 1.09035129 | 0.124793017 | 0.27087473 | 1 |
| Wdr25 | 1.16997599 | 0.226478925 | 0.27095146 | 1 |
| Slc6a1 | 1.08608658 | 0.119139114 | 0.27104965 | 1 |
| Zfp111 | 0.83604452 | -0.258348319 | 0.27121015 | 1 |
| Pxdn | 0.94781 | -0.07733021 | 0.27124432 | 1 |
| Cdc42se1 | 0.93961476 | -0.089858717 | 0.27137654 | 1 |
| RGD1560455 | 1.41919675 | 0.505074613 | 0.27154088 | 1 |
| Pla2g3 | 1.09336064 | 0.128769345 | 0.27162236 | 1 |
| Rap1gap2 | 0.9427212 | -0.085096918 | 0.27181513 | 1 |
| Prrc2a | 0.94985522 | -0.074220468 | 0.27200813 | 1 |
| LOC100912312 | 0.81381958 | -0.297219099 | 0.2720425 | 1 |
| Zkscan1 | 1.09699616 | 0.133558476 | 0.27206352 | 1 |
| Plscr2 | 0.87072858 | -0.199705014 | 0.27208041 | 1 |
| Ninj1 | 0.91949087 | -0.121092839 | 0.27211339 | 1 |
| Hdgfrp2 | 0.93558321 | -0.09606212 | 0.2721252 | 1 |
| Ubox5 | 1.08658536 | 0.119801508 | 0.27220945 | 1 |
| Lamp1 | 0.95021567 | -0.073673093 | 0.27225963 | 1 |
| Krr1 | 0.90309822 | -0.147045197 | 0.27231487 | 1 |
| Piezo1 | 0.90920461 | -0.137323097 | 0.27242675 | 1 |
| Nif3l1 | 0.88963167 | -0.168719955 | 0.2726365 | 1 |
| Kcnc1 | 0.92456809 | -0.113148519 | 0.27266007 | 1 |
| Dus3l | 0.90890508 | -0.137798459 | 0.27284906 | 1 |
| Frmd6 | 1.06260581 | 0.087606505 | 0.27287693 | 1 |
| Cnot2 | 1.08116876 | 0.112591728 | 0.27293096 | 1 |
| Dbnl | 1.05518274 | 0.077492867 | 0.27304186 | 1 |
| Rfc1 | 1.05997297 | 0.084027477 | 0.27337375 | 1 |
| Rad17 | 0.91711478 | -0.124825795 | 0.27337636 | 1 |
| Tdo2 | 1.23307647 | 0.302262273 | 0.27337725 | 1 |
| Eif1ad | 1.08617047 | 0.119250543 | 0.27338737 | 1 |
| Dync1h1 | 0.94819951 | -0.07673745 | 0.27339047 | 1 |
| Inpp5e | 0.91381824 | -0.130020858 | 0.27341809 | 1 |
| RGD1305110 | 1.05695438 | 0.079913106 | 0.27351266 | 1 |
| Lsm1 | 1.10943123 | 0.149820245 | 0.27354739 | 1 |
| Zfp36l1 | 0.93402282 | -0.098470294 | 0.27368372 | 1 |
| Mtx1 | 0.90375955 | -0.145989101 | 0.27379843 | 1 |
| Lonp2 | 1.06152611 | 0.086139863 | 0.27396277 | 1 |
| Tamm41 | 1.11895341 | 0.162149963 | 0.2741464 | 1 |

| | | | | |
|---|---|---|---|---|
| Reep4 | 0.8562447 | -0.223904939 | 0.27416797 | 1 |
| Tmod4 | 1.24229784 | 0.313011103 | 0.27434076 | 1 |
| Dnajc24 | 1.14104869 | 0.190360358 | 0.27442888 | 1 |
| Cdc42ep2 | 1.0748913 | 0.104190778 | 0.27450159 | 1 |
| Ftsj2 | 0.83564479 | -0.259038272 | 0.27459691 | 1 |
| Slc6a8 | 0.94360516 | -0.083744787 | 0.27464244 | 1 |
| Fam49a | 1.14112486 | 0.19045666 | 0.27488488 | 1 |
| Nt5e | 1.10230301 | 0.140520857 | 0.27499867 | 1 |
| Cep295 | 1.11308915 | 0.154569152 | 0.27531797 | 1 |
| Sec61a1 | 0.94841222 | -0.076413843 | 0.27538893 | 1 |
| Csmd1 | 0.94022571 | -0.088920969 | 0.27543684 | 1 |
| Ccnc | 1.07141694 | 0.099520015 | 0.27544905 | 1 |
| Chrna3 | 0.94536686 | -0.081053798 | 0.2755516 | 1 |
| Cd72 | 0.85903493 | -0.219211297 | 0.27559662 | 1 |
| Pard6a | 1.15393898 | 0.206566934 | 0.27575507 | 1 |
| Nrip3 | 1.0514711 | 0.072409202 | 0.27579365 | 1 |
| Nav2 | 0.93353667 | -0.099221407 | 0.27589169 | 1 |
| Arhgef25 | 1.06725112 | 0.093899671 | 0.27593965 | 1 |
| Pccb | 0.9244941 | -0.113263984 | 0.27602506 | 1 |
| Commd4 | 1.08516412 | 0.117913246 | 0.27603997 | 1 |
| Syn2 | 0.94838658 | -0.07645285 | 0.27607635 | 1 |
| Mad1l1 | 0.91690604 | -0.125154198 | 0.27618697 | 1 |
| Mcts1 | 0.93239496 | -0.100986893 | 0.27627298 | 1 |
| Nat8b | 0.81361561 | -0.297580728 | 0.27657019 | 1 |
| Lrrc47 | 0.92503686 | -0.112417236 | 0.27661824 | 1 |
| Akr1c14 | 1.1225653 | 0.166799372 | 0.27668918 | 1 |
| Acaa1 | 0.91934929 | -0.121315004 | 0.27672142 | 1 |
| Slc11a1 | 1.28567395 | 0.362524822 | 0.27683694 | 1 |
| Ccdc77 | 0.83760191 | -0.255663356 | 0.27686525 | 1 |
| Mreg | 1.28347979 | 0.360060576 | 0.27699466 | 1 |
| Pcp4 | 1.05180838 | 0.072871893 | 0.27724644 | 1 |
| LOC500300 | 0.87584749 | -0.191248411 | 0.27732354 | 1 |
| Gpr75 | 1.17247972 | 0.229562975 | 0.27733293 | 1 |
| Rhog | 1.07803737 | 0.108407187 | 0.27734991 | 1 |
| Rabl6 | 0.94814085 | -0.076826702 | 0.27747296 | 1 |
| Ermp1 | 0.94912983 | -0.075322653 | 0.27762812 | 1 |
| Slc25a1 | 0.93637797 | -0.094837103 | 0.27767527 | 1 |
| C1qtnf2 | 1.0990089 | 0.136203066 | 0.27791566 | 1 |
| Ubxn8 | 1.09899596 | 0.136186077 | 0.27797329 | 1 |
| Gfi1b | 1.62644116 | 0.701718633 | 0.27801686 | 1 |

| | | | | |
|---|---|---|---|---|
| Pex5l | 0.93783348 | -0.092596319 | 0.27804935 | 1 |
| Mblac1 | 1.14502669 | 0.195381225 | 0.27805898 | 1 |
| Polr2b | 1.05699363 | 0.079966687 | 0.27806757 | 1 |
| Rassf4 | 0.94972553 | -0.074417453 | 0.27812757 | 1 |
| Slco4a1 | 1.31753128 | 0.397837209 | 0.27818901 | 1 |
| Kcnk18 | 0.87188568 | -0.197789118 | 0.27821199 | 1 |
| Tmub2 | 0.94095721 | -0.087798977 | 0.27827995 | 1 |
| Stard4 | 0.92683524 | -0.109615199 | 0.27843436 | 1 |
| Zkscan3 | 0.90551876 | -0.143183567 | 0.27867445 | 1 |
| Clec4e | 0.72269903 | -0.468533139 | 0.27878583 | 1 |
| Eri2 | 1.13221608 | 0.179149315 | 0.27885244 | 1 |
| Spag8 | 1.27357048 | 0.348878807 | 0.27902423 | 1 |
| Dph2 | 1.16572384 | 0.221226056 | 0.27916983 | 1 |
| Fbxw4 | 0.91850568 | -0.122639456 | 0.27932304 | 1 |
| Rab14 | 0.94021447 | -0.088938206 | 0.27937457 | 1 |
| Dynlt1 | 0.91461058 | -0.128770485 | 0.27941953 | 1 |
| Dock1 | 0.9352335 | -0.096601489 | 0.27957797 | 1 |
| Smarcb1 | 0.93189307 | -0.101763674 | 0.2796191 | 1 |
| Trmt10c | 1.08431609 | 0.116785383 | 0.27968791 | 1 |
| Tbk1 | 1.06140001 | 0.085968468 | 0.27981326 | 1 |
| Zfp161 | 0.90985104 | -0.136297732 | 0.27989917 | 1 |
| Ppie | 0.87636185 | -0.190401418 | 0.28000677 | 1 |
| Pcyt1a | 1.07165005 | 0.099833863 | 0.28002381 | 1 |
| Ptprk | 0.91973641 | -0.120707645 | 0.28020403 | 1 |
| Glyctk | 1.15248119 | 0.2047432 | 0.28043766 | 1 |
| Mb | 0.78373518 | -0.351561833 | 0.28047248 | 1 |
| Arhgap27 | 1.1227586 | 0.167047767 | 0.28059047 | 1 |
| LOC100911483 | 1.06570407 | 0.091806873 | 0.28059262 | 1 |
| Zzz3 | 0.93101511 | -0.103123516 | 0.28070442 | 1 |
| Ntan1 | 1.05630373 | 0.079024724 | 0.28123271 | 1 |
| Atp5i | 1.05989071 | 0.083915513 | 0.28125465 | 1 |
| Ndufb2 | 1.07120539 | 0.099235124 | 0.28147537 | 1 |
| Qpct | 0.87438646 | -0.193657036 | 0.28150347 | 1 |
| Actr1b | 0.95194545 | -0.071049197 | 0.28160566 | 1 |
| Kcnj12 | 0.91830749 | -0.122950778 | 0.28164407 | 1 |
| Sema4d | 0.93061706 | -0.103740461 | 0.28174023 | 1 |
| Prex2 | 1.07166003 | 0.099847308 | 0.28174694 | 1 |
| Gins2 | 1.30778513 | 0.387125526 | 0.2818464 | 1 |
| Fmnl1 | 0.92598068 | -0.110946004 | 0.2819269 | 1 |
| Minos1 | 0.9402485 | -0.088886002 | 0.28203592 | 1 |

| | | | | |
|---|---|---|---|---|
| Dgkg | 0.92173154 | -0.117581471 | 0.28206954 | 1 |
| Timm22 | 1.07350514 | 0.102329101 | 0.28212236 | 1 |
| Tmem189 | 0.95088298 | -0.072660283 | 0.2821991 | 1 |
| Igf1 | 0.91141645 | -0.133817692 | 0.28228724 | 1 |
| Dapk3 | 0.93075008 | -0.103534259 | 0.28235302 | 1 |
| Mgmt | 1.14137455 | 0.190772305 | 0.28243121 | 1 |
| Fam184b | 0.89130868 | -0.166002946 | 0.28248796 | 1 |
| Mical1 | 0.94541025 | -0.080987584 | 0.28252139 | 1 |
| Psma6 | 1.0518881 | 0.072981244 | 0.28253898 | 1 |
| RGD1306941 | 1.05625675 | 0.078960567 | 0.28263425 | 1 |
| Fpgs | 0.89864315 | -0.154179765 | 0.28289846 | 1 |
| Sec22a | 1.12302113 | 0.167385074 | 0.28291375 | 1 |
| Evc | 1.11782383 | 0.160692832 | 0.28296206 | 1 |
| Dvl2 | 0.87994563 | -0.184513708 | 0.28299156 | 1 |
| Gimap8 | 0.88057635 | -0.183479991 | 0.2830417 | 1 |
| Hsd3b7 | 1.16057335 | 0.21483771 | 0.28316095 | 1 |
| Mapk12 | 1.06427887 | 0.089876221 | 0.28335006 | 1 |
| Nomo1 | 0.95016234 | -0.073754073 | 0.28338995 | 1 |
| Mitd1 | 1.11790595 | 0.160798822 | 0.28344436 | 1 |
| Ttc4 | 1.11334958 | 0.154906654 | 0.28364389 | 1 |
| Apoh | 1.27715372 | 0.352932183 | 0.28369199 | 1 |
| Igsf10 | 0.86121584 | -0.215553238 | 0.28386974 | 1 |
| Map9 | 1.07822111 | 0.108653063 | 0.28391107 | 1 |
| Prkar1b | 0.95143178 | -0.071827887 | 0.28393248 | 1 |
| Aff3 | 0.91747373 | -0.124261238 | 0.28402881 | 1 |
| Mcam | 0.95195352 | -0.071036956 | 0.28406161 | 1 |
| Prr15 | 1.2103521 | 0.275426796 | 0.28415165 | 1 |
| Snx19 | 0.95098591 | -0.072504133 | 0.28439011 | 1 |
| Cybrd1 | 1.1341892 | 0.18166132 | 0.28451501 | 1 |
| Misp | 0.81293542 | -0.298787354 | 0.28455035 | 1 |
| Top2b | 0.94069925 | -0.088194546 | 0.28456881 | 1 |
| Gria2 | 0.93856362 | -0.091473558 | 0.28460019 | 1 |
| Tspan8 | 1.05074633 | 0.071414424 | 0.28461006 | 1 |
| Sox2 | 0.9026004 | -0.147840679 | 0.28469436 | 1 |
| Rgma | 1.1214548 | 0.16537147 | 0.28470674 | 1 |
| Rbm48 | 0.87510755 | -0.192467763 | 0.28473788 | 1 |
| Lnpep | 0.93333032 | -0.099540338 | 0.28481816 | 1 |
| Derl3 | 1.22260937 | 0.28996353 | 0.28493295 | 1 |
| Csf3r | 1.21345494 | 0.279120535 | 0.2850206 | 1 |
| Bok | 0.92366573 | -0.114557253 | 0.2851979 | 1 |

| | | | | |
|---|---|---|---|---|
| Cyhr1 | 1.09950977 | 0.136860429 | 0.28524618 | 1 |
| Ada | 0.88041049 | -0.183751764 | 0.28526668 | 1 |
| Stx1a | 0.9147162 | -0.1286039 | 0.28530019 | 1 |
| Coasy | 1.07410193 | 0.103130906 | 0.28537402 | 1 |
| Gng10 | 0.94068852 | -0.088210996 | 0.28545067 | 1 |
| Cbx5 | 1.09816511 | 0.135094975 | 0.28547368 | 1 |
| Mapkapk3 | 1.0967539 | 0.133239843 | 0.28558517 | 1 |
| Pex7 | 0.87880248 | -0.186389154 | 0.28583217 | 1 |
| Pgd | 1.05424139 | 0.076205244 | 0.28603965 | 1 |
| Igsf8 | 1.08774103 | 0.12133512 | 0.28608042 | 1 |
| E2f1 | 1.11204343 | 0.153213126 | 0.28633775 | 1 |
| R3hdm1 | 0.9340729 | -0.098392951 | 0.28650631 | 1 |
| Rrn3 | 0.94427489 | -0.082721182 | 0.2865278 | 1 |
| Aimp2 | 0.92430135 | -0.113564803 | 0.28667082 | 1 |
| Nat1 | 1.18085281 | 0.239829153 | 0.28676072 | 1 |
| Pomp | 1.05361865 | 0.075352784 | 0.28691598 | 1 |
| Rsad2 | 1.10978847 | 0.150284717 | 0.28692272 | 1 |
| Larp1b | 0.93075014 | -0.103534169 | 0.28696284 | 1 |
| Mvb12b | 0.91566147 | -0.127113782 | 0.28700484 | 1 |
| Fkbp14 | 1.09854483 | 0.135593739 | 0.28706059 | 1 |
| Ugt1a7c | 0.82589646 | -0.275967167 | 0.28707036 | 1 |
| Klhl7 | 1.05796851 | 0.081296687 | 0.28713158 | 1 |
| Morc3 | 1.06648082 | 0.092858022 | 0.2872089 | 1 |
| Hadhb | 1.05012787 | 0.070565003 | 0.28725146 | 1 |
| Ccdc25 | 1.07897833 | 0.109665886 | 0.28727515 | 1 |
| Myo1b | 0.94042821 | -0.088610278 | 0.28732598 | 1 |
| Ankrd42 | 0.91875314 | -0.122250823 | 0.28749627 | 1 |
| Tmem5 | 1.08798491 | 0.121658541 | 0.28772185 | 1 |
| Pgm5 | 1.14936611 | 0.200838413 | 0.28776288 | 1 |
| Ttpal | 0.94066499 | -0.088247079 | 0.28807772 | 1 |
| Borcs7 | 1.06963007 | 0.097111923 | 0.28813056 | 1 |
| Scrn1 | 0.94994339 | -0.074086546 | 0.28832892 | 1 |
| B2m | 1.05203551 | 0.073183399 | 0.28844008 | 1 |
| Lrrc63 | 0.81940142 | -0.287357709 | 0.28844568 | 1 |
| Nsun5 | 1.10548148 | 0.144674855 | 0.28847156 | 1 |
| Aga | 0.93113437 | -0.102938725 | 0.2887187 | 1 |
| Pitpna | 0.95268796 | -0.069924337 | 0.28887219 | 1 |
| Tada2a | 1.08935516 | 0.123474384 | 0.28896277 | 1 |
| Evi5 | 0.95191232 | -0.071099401 | 0.28897864 | 1 |
| Zfp219 | 0.93135216 | -0.102601314 | 0.28899348 | 1 |

| | | | | |
|---|---|---|---|---|
| Oaz1 | 0.95250783 | -0.07019714 | 0.28908737 | 1 |
| Tufm | 1.05717531 | 0.08021463 | 0.28917717 | 1 |
| Dcaf13 | 0.92562029 | -0.11150761 | 0.28919042 | 1 |
| Stk16 | 1.07754871 | 0.107753087 | 0.28925571 | 1 |
| Sfxn5 | 0.88715172 | -0.172747245 | 0.28930814 | 1 |
| Micb | 0.76700945 | -0.382683738 | 0.28949933 | 1 |
| Ccdc126 | 1.06474203 | 0.090503925 | 0.28963285 | 1 |
| Nus1 | 0.95061542 | -0.073066292 | 0.28967597 | 1 |
| Vamp1 | 1.05267843 | 0.074064797 | 0.28973701 | 1 |
| Ldhb | 1.05074921 | 0.071418376 | 0.28974276 | 1 |
| Edn3 | 1.34541374 | 0.428049901 | 0.28981466 | 1 |
| Nsmf | 0.94611518 | -0.079912274 | 0.29010145 | 1 |
| Ethe1 | 1.10546762 | 0.144656767 | 0.2901424 | 1 |
| Lsm14a | 0.94072108 | -0.088161065 | 0.29029018 | 1 |
| Serpine2 | 0.94979308 | -0.074314854 | 0.29032362 | 1 |
| Fads3 | 0.92436359 | -0.113467663 | 0.29038884 | 1 |
| Stk32c | 0.92267619 | -0.116103661 | 0.2904608 | 1 |
| Dlgap5 | 0.86861067 | -0.203218423 | 0.29056132 | 1 |
| Sbds | 1.05575323 | 0.078272665 | 0.29058195 | 1 |
| Camk1g | 0.86378566 | -0.211254732 | 0.2908739 | 1 |
| Hdhd2 | 0.91891492 | -0.121996801 | 0.29091084 | 1 |
| Chpf | 0.94720522 | -0.078251065 | 0.2910879 | 1 |
| Scpep1 | 1.06490362 | 0.09072286 | 0.29112489 | 1 |
| Mbip | 1.11786319 | 0.160743639 | 0.29118323 | 1 |
| Idi1 | 0.94939242 | -0.074923566 | 0.29138443 | 1 |
| Zic4 | 0.75499708 | -0.405457027 | 0.29152378 | 1 |
| Kcnh5 | 0.87889577 | -0.186236017 | 0.29155017 | 1 |
| Efcab7 | 1.11744872 | 0.160208622 | 0.29170224 | 1 |
| Cisd3 | 1.07964143 | 0.110552247 | 0.29174008 | 1 |
| Grik1 | 0.95105321 | -0.072402032 | 0.29181248 | 1 |
| Ncor2 | 0.95095811 | -0.072546299 | 0.29193864 | 1 |
| Ccnk | 1.08571724 | 0.11864842 | 0.29205906 | 1 |
| Lox | 1.17775179 | 0.236035529 | 0.29210505 | 1 |
| Fbxo33 | 0.91910128 | -0.121704244 | 0.2925748 | 1 |
| Pgk1 | 1.04977448 | 0.070079431 | 0.29268639 | 1 |
| Vsnl1 | 1.04967803 | 0.06994688 | 0.29270746 | 1 |
| Inpp5a | 0.92086092 | -0.118944808 | 0.29272051 | 1 |
| Yod1 | 1.17023412 | 0.226797182 | 0.2927469 | 1 |
| Dimt1 | 0.86026742 | -0.217142902 | 0.29275175 | 1 |
| Pank2 | 0.88886189 | -0.169968821 | 0.29276014 | 1 |

| | | | | |
|---|---|---|---|---|
| Sertm1 | 1.07369972 | 0.102590579 | 0.29283964 | 1 |
| Slc3a2 | 1.05169355 | 0.07271439 | 0.29288732 | 1 |
| Ttll12 | 1.10793388 | 0.147871792 | 0.29312591 | 1 |
| Nubp1 | 0.90667048 | -0.141349784 | 0.29317497 | 1 |
| Ndufb6 | 1.05867467 | 0.082259323 | 0.29320797 | 1 |
| Rbm12 | 0.91675299 | -0.125395024 | 0.29335286 | 1 |
| Rasal3 | 0.89100593 | -0.166493059 | 0.29352516 | 1 |
| Cilp2 | 1.39696888 | 0.482299881 | 0.29356855 | 1 |
| Timp2 | 0.95227847 | -0.070544587 | 0.29371343 | 1 |
| Gltscr1 | 0.90876318 | -0.138023707 | 0.29382297 | 1 |
| Clrn1 | 1.09080133 | 0.125388369 | 0.29393568 | 1 |
| Rybp | 0.93642814 | -0.094759806 | 0.29394353 | 1 |
| Gja5 | 0.79627663 | -0.328658374 | 0.29397321 | 1 |
| Arhgap20 | 1.11822394 | 0.161209137 | 0.29421659 | 1 |
| Zfp641 | 0.87837945 | -0.187083793 | 0.29437306 | 1 |
| Mrpl17 | 1.05256119 | 0.073904111 | 0.29441659 | 1 |
| Rufy2 | 0.93986376 | -0.089476457 | 0.29446808 | 1 |
| Rock2 | 0.9516602 | -0.071481564 | 0.29458936 | 1 |
| Ppp1r1a | 0.94345134 | -0.083979991 | 0.29459192 | 1 |
| Nabp2 | 0.93322992 | -0.099695533 | 0.29475859 | 1 |
| Gclc | 0.93847858 | -0.091604282 | 0.2947854 | 1 |
| Fbxl7 | 1.17390154 | 0.231311411 | 0.29479641 | 1 |
| Klhdc10 | 0.9210596 | -0.118633582 | 0.29480525 | 1 |
| Rtfdc1 | 1.05594389 | 0.078533178 | 0.29498188 | 1 |
| Tgfbr3 | 0.9430212 | -0.084637887 | 0.29502206 | 1 |
| Rprd1b | 1.08386371 | 0.116183353 | 0.29505866 | 1 |
| Map2 | 1.04973673 | 0.070027546 | 0.29510306 | 1 |
| Ndufa4 | 1.04923487 | 0.069337658 | 0.29511932 | 1 |
| Col5a2 | 0.95268817 | -0.069924022 | 0.29513309 | 1 |
| Etnk1 | 0.95317478 | -0.069187318 | 0.29514145 | 1 |
| Dgkb | 1.07409222 | 0.10311786 | 0.29515921 | 1 |
| RGD1310262 | 0.78145376 | -0.355767585 | 0.29518861 | 1 |
| Sema3c | 1.05041056 | 0.070953328 | 0.29526511 | 1 |
| Cpne4 | 1.05174783 | 0.072788837 | 0.29526877 | 1 |
| Arhgdib | 1.08404333 | 0.116422428 | 0.29527508 | 1 |
| Nampt | 0.93902434 | -0.090765546 | 0.29528827 | 1 |
| Ap3d1 | 0.95314439 | -0.069233313 | 0.29558062 | 1 |
| Pced1a | 1.10752908 | 0.147344583 | 0.29561989 | 1 |
| Camkv | 0.8661687 | -0.207280053 | 0.29571703 | 1 |
| Kcnc3 | 0.94935913 | -0.074974146 | 0.29574508 | 1 |

| | | | | |
|---|---|---|---|---|
| Snip1 | 1.09349949 | 0.128952553 | 0.29603624 | 1 |
| Atf6b | 0.93849297 | -0.091582159 | 0.29611235 | 1 |
| Dhtkd1 | 1.0941099 | 0.129757664 | 0.2961539 | 1 |
| Pgap2 | 1.10688266 | 0.146502287 | 0.29623247 | 1 |
| Hsp90aa1 | 1.05317667 | 0.074747473 | 0.29630598 | 1 |
| Pde5a | 1.11302234 | 0.154482551 | 0.29632179 | 1 |
| Sowahc | 0.93548268 | -0.09621715 | 0.29680564 | 1 |
| Celsr3 | 0.93870769 | -0.091252119 | 0.29698068 | 1 |
| Hrasls | 1.21501727 | 0.280976822 | 0.29702169 | 1 |
| Sema4c | 0.93478396 | -0.09729511 | 0.29713923 | 1 |
| Rps17 | 1.05340168 | 0.075055659 | 0.29718185 | 1 |
| Pigw | 1.12701717 | 0.172509497 | 0.29727787 | 1 |
| Nab1 | 0.94643494 | -0.079424758 | 0.29762936 | 1 |
| Lifr | 0.95257114 | -0.070101251 | 0.29776082 | 1 |
| Zfp316 | 0.88533101 | -0.17571114 | 0.29790155 | 1 |
| Rgs7bp | 0.94155447 | -0.086883536 | 0.29794241 | 1 |
| Impdh2 | 1.06076851 | 0.085109859 | 0.29796945 | 1 |
| Bche | 1.1696077 | 0.226024711 | 0.29797635 | 1 |
| Qrsl1 | 1.12659837 | 0.171973293 | 0.29807672 | 1 |
| Wars | 1.04959261 | 0.069829471 | 0.29814488 | 1 |
| Rbp1 | 1.11004373 | 0.150616508 | 0.29820481 | 1 |
| Rnase2 | 1.46884456 | 0.554681733 | 0.29821128 | 1 |
| Prph | 0.95077133 | -0.072829695 | 0.29832914 | 1 |
| RGD1309534 | 1.08750044 | 0.121015981 | 0.29837013 | 1 |
| Prelid1 | 0.94802611 | -0.077001297 | 0.29838745 | 1 |
| Rnf144a | 0.90204099 | -0.148735103 | 0.29841113 | 1 |
| Rell2 | 0.94849001 | -0.076295512 | 0.29846499 | 1 |
| Als2 | 0.93226707 | -0.101184787 | 0.29861452 | 1 |
| Maz | 0.92394747 | -0.114117257 | 0.29867107 | 1 |
| Nyap1 | 0.93554782 | -0.096116698 | 0.29871057 | 1 |
| Brpf1 | 1.09629106 | 0.132630881 | 0.29893195 | 1 |
| Vps41 | 1.0496797 | 0.069949176 | 0.29910217 | 1 |
| Ankrd24 | 1.1207475 | 0.164461286 | 0.29911763 | 1 |
| Slc39a13 | 1.07108272 | 0.099069902 | 0.29932391 | 1 |
| Mrpl3 | 0.9358012 | -0.095726012 | 0.29933375 | 1 |
| Plpp1 | 1.05381722 | 0.075624653 | 0.29933928 | 1 |
| Hadha | 1.04876885 | 0.068696745 | 0.29937642 | 1 |
| Fads1 | 0.95367035 | -0.068437429 | 0.29941119 | 1 |
| Epyc | 0.75201026 | -0.411175754 | 0.29942758 | 1 |
| Kcna3 | 0.81076285 | -0.302648111 | 0.29944577 | 1 |

| | | | | |
|---|---|---|---|---|
| Slc2a3 | 1.0745257 | 0.10369999 | 0.29948181 | 1 |
| Ttc23 | 1.11293715 | 0.154372125 | 0.29949927 | 1 |
| Nup43 | 1.12376524 | 0.168340676 | 0.29958799 | 1 |
| Lats2 | 0.94708448 | -0.078434971 | 0.2996252 | 1 |
| Heyl | 0.91204031 | -0.132830513 | 0.29974449 | 1 |
| Nat14 | 0.86560427 | -0.208220477 | 0.30037353 | 1 |
| Nlrp3 | 1.20814958 | 0.27279909 | 0.30066156 | 1 |
| Htatip2 | 1.06602173 | 0.092236849 | 0.30066204 | 1 |
| Sf3a2 | 0.89805239 | -0.155128491 | 0.30087318 | 1 |
| Ccdc134 | 1.19583232 | 0.258015113 | 0.30088458 | 1 |
| Btaf1 | 0.93727066 | -0.093462368 | 0.30090936 | 1 |
| Emc10 | 0.94920657 | -0.075206014 | 0.3009716 | 1 |
| rnf141 | 0.92180033 | -0.117473806 | 0.30098008 | 1 |
| Emc1 | 0.94718359 | -0.078284001 | 0.30106879 | 1 |
| Aadat | 0.80458122 | -0.31369004 | 0.30116994 | 1 |
| Med27 | 1.0791919 | 0.109951423 | 0.30121886 | 1 |
| Foxc2 | 1.12117407 | 0.165010283 | 0.30122174 | 1 |
| Fam65b | 1.05755877 | 0.080737845 | 0.30132297 | 1 |
| Cutc | 0.88867201 | -0.170277048 | 0.30136431 | 1 |
| Zc3h6 | 1.1334633 | 0.18073768 | 0.30145363 | 1 |
| Egflam | 0.9308752 | -0.10334033 | 0.30152931 | 1 |
| Fam127b | 0.95001397 | -0.073979362 | 0.30153095 | 1 |
| Pls1 | 1.06068579 | 0.084997346 | 0.30153182 | 1 |
| Eif5a2 | 1.04950255 | 0.069705673 | 0.3016856 | 1 |
| Avpi1 | 0.9208802 | -0.118914603 | 0.30186905 | 1 |
| Zfp157 | 1.10059679 | 0.138286021 | 0.30192037 | 1 |
| Dis3l2 | 0.90932345 | -0.137134536 | 0.30192396 | 1 |
| Hspb1 | 1.04848521 | 0.068306509 | 0.30196824 | 1 |
| Apold1 | 0.88087704 | -0.182987441 | 0.30208544 | 1 |
| Srrm2 | 0.95256321 | -0.07011327 | 0.3021148 | 1 |
| Egln2 | 0.94776729 | -0.077395227 | 0.30239064 | 1 |
| Cish | 0.90933697 | -0.137113083 | 0.30240759 | 1 |
| Rpl35a | 0.94396615 | -0.083192962 | 0.30249606 | 1 |
| Ptgdrl | 0.9067503 | -0.141222783 | 0.30287056 | 1 |
| Thrb | 0.9085288 | -0.13839585 | 0.30291178 | 1 |
| Gusb | 1.11444566 | 0.156326268 | 0.30312327 | 1 |
| Cacybp | 1.04832387 | 0.068084493 | 0.30319589 | 1 |
| Sfpq | 0.94657227 | -0.079215433 | 0.30328828 | 1 |
| Exoc3l1 | 1.18880234 | 0.249508862 | 0.30331092 | 1 |
| Abhd5 | 0.91380314 | -0.130044698 | 0.30332609 | 1 |

| | | | | |
|---|---|---|---|---|
| Smad9 | 1.08376168 | 0.116047543 | 0.30337345 | 1 |
| Cc2d1b | 0.94396151 | -0.083200054 | 0.30347353 | 1 |
| Tpst2 | 0.92479207 | -0.112799074 | 0.30360832 | 1 |
| Sorl1 | 0.95247021 | -0.07025413 | 0.30398727 | 1 |
| Irf5 | 1.1634192 | 0.218371017 | 0.3041233 | 1 |
| Gmeb1 | 1.12851577 | 0.174426579 | 0.30418603 | 1 |
| Timeless | 1.21652706 | 0.282768412 | 0.30421716 | 1 |
| Myrf | 1.12343681 | 0.167918975 | 0.30422601 | 1 |
| Dgcr8 | 0.92880614 | -0.106550587 | 0.30423813 | 1 |
| Enpp5 | 1.05096332 | 0.07171232 | 0.30426511 | 1 |
| Ift172 | 1.06344971 | 0.088751816 | 0.30430767 | 1 |
| Il1rap | 0.89049816 | -0.167315469 | 0.30439887 | 1 |
| Anapc5 | 1.04825678 | 0.067992163 | 0.30443845 | 1 |
| Cachd1 | 0.91223813 | -0.132517627 | 0.30450792 | 1 |
| Orc5 | 1.07148879 | 0.099616758 | 0.30451072 | 1 |
| Chmp5 | 1.05006754 | 0.070482131 | 0.3045705 | 1 |
| Tmem106c | 1.09605591 | 0.132321396 | 0.30462532 | 1 |
| Svil | 1.05974779 | 0.083720958 | 0.30472857 | 1 |
| B3galnt1 | 0.93291016 | -0.100189938 | 0.30475554 | 1 |
| Ttbk2 | 0.95016371 | -0.073751991 | 0.30490341 | 1 |
| Hlx | 1.24616911 | 0.317499866 | 0.30499885 | 1 |
| Pcdhb5 | 1.21162153 | 0.276939118 | 0.3050449 | 1 |
| Ctsa | 1.04813955 | 0.067830813 | 0.30513904 | 1 |
| Pcdha9 | 1.16645611 | 0.222132029 | 0.30514802 | 1 |
| E2f7 | 1.36334343 | 0.44714903 | 0.30549006 | 1 |
| Cct2 | 1.04807692 | 0.067744607 | 0.30555736 | 1 |
| Slc9a1 | 0.94479303 | -0.081929776 | 0.30563004 | 1 |
| Acp6 | 1.12524389 | 0.170237733 | 0.30581464 | 1 |
| Ccna1 | 0.81122717 | -0.301822115 | 0.30583372 | 1 |
| Tmem165 | 1.07341005 | 0.102201304 | 0.30588385 | 1 |
| Gpr108 | 1.06779387 | 0.094633178 | 0.30601069 | 1 |
| B3gat3 | 1.05041207 | 0.070955396 | 0.30611095 | 1 |
| Rabggta | 0.91780319 | -0.123743268 | 0.30641551 | 1 |
| Sh2d3c | 0.92017233 | -0.120024024 | 0.30641985 | 1 |
| Ctns | 1.07894534 | 0.109621785 | 0.30654484 | 1 |
| Map2k3 | 0.91040167 | -0.135424894 | 0.30670948 | 1 |
| Svop | 0.92794703 | -0.107885644 | 0.30679919 | 1 |
| Brd1 | 0.93220986 | -0.101273315 | 0.30690549 | 1 |
| Zbtb8b | 1.24500621 | 0.316152937 | 0.30691972 | 1 |
| Fbxw17 | 1.11802403 | 0.160951193 | 0.30694904 | 1 |

| | | | | |
|---|---|---|---|---|
| Obfc1 | 0.91932102 | -0.121359359 | 0.30702172 | 1 |
| Cdkn2aip | 1.0952537 | 0.131265091 | 0.30706758 | 1 |
| Tinagl1 | 1.10862643 | 0.148773312 | 0.30712908 | 1 |
| Scn7a | 1.05074784 | 0.071416492 | 0.30713409 | 1 |
| Heatr1 | 0.91733877 | -0.124473477 | 0.30716341 | 1 |
| Got1 | 0.95426682 | -0.067535385 | 0.30718102 | 1 |
| Whsc1l1 | 0.93123373 | -0.102784786 | 0.30734767 | 1 |
| Cryab | 0.95329959 | -0.068998424 | 0.30738407 | 1 |
| Homer1 | 1.09082895 | 0.125424899 | 0.30739193 | 1 |
| Ebf1 | 0.93787551 | -0.092531658 | 0.30745994 | 1 |
| Stx1b | 1.05337886 | 0.075024407 | 0.30748922 | 1 |
| Lhfp | 1.0656556 | 0.091741267 | 0.30752129 | 1 |
| Polg | 0.93760876 | -0.092942047 | 0.30756484 | 1 |
| LOC500227 | 1.25647016 | 0.329376413 | 0.30764568 | 1 |
| Il6r | 0.9137688 | -0.130098914 | 0.30772845 | 1 |
| Synpr | 0.9543141 | -0.067463905 | 0.3077514 | 1 |
| Mrap2 | 0.94105712 | -0.087645798 | 0.30842137 | 1 |
| Eif4e3 | 1.0901792 | 0.124565298 | 0.30842558 | 1 |
| Nup214 | 0.93526783 | -0.096548533 | 0.30857904 | 1 |
| Selplg | 0.82827073 | -0.271825696 | 0.30885273 | 1 |
| Lypd3 | 0.75389494 | -0.407564597 | 0.30887655 | 1 |
| RGD1305350 | 0.93850694 | -0.091560676 | 0.30888976 | 1 |
| Pcif1 | 1.11980649 | 0.163249446 | 0.30891482 | 1 |
| Ns5atp9 | 1.40441376 | 0.489968032 | 0.30895343 | 1 |
| Podxl2 | 0.93238815 | -0.100997421 | 0.30897277 | 1 |
| Ndufb9 | 1.05095085 | 0.071695201 | 0.30898204 | 1 |
| Ppfibp1 | 1.06224684 | 0.087119051 | 0.30908672 | 1 |
| Fhod3 | 1.05692148 | 0.079868207 | 0.30911349 | 1 |
| Angpt1 | 1.14660138 | 0.197363926 | 0.30916879 | 1 |
| Siva1 | 1.15203016 | 0.204178481 | 0.30918682 | 1 |
| Plek | 1.15759332 | 0.211128506 | 0.30930605 | 1 |
| Rad51c | 0.8455678 | -0.242007657 | 0.30933151 | 1 |
| LOC100125362 | 1.04765175 | 0.067159232 | 0.30948637 | 1 |
| Ltbp2 | 1.0924859 | 0.127614656 | 0.30953438 | 1 |
| Rasgrf2 | 1.19527019 | 0.257336771 | 0.30967088 | 1 |
| Plcd4 | 0.95048444 | -0.073265092 | 0.3098364 | 1 |
| Nr2f2 | 1.07064474 | 0.098479847 | 0.30997222 | 1 |
| Usp16 | 1.07595664 | 0.105619943 | 0.30998872 | 1 |
| Kbtbd4 | 0.85338689 | -0.228728154 | 0.31007089 | 1 |
| Prkg2 | 1.11720183 | 0.159889842 | 0.31009029 | 1 |

| | | | | |
|---|---|---|---|---|
| Snap91 | 0.95458153 | -0.067059672 | 0.31016178 | 1 |
| Stxbp4 | 1.16282688 | 0.217636327 | 0.31020962 | 1 |
| Pmfbp1 | 1.17112247 | 0.227891955 | 0.31025159 | 1 |
| Pkd1 | 0.95355789 | -0.068607566 | 0.31028273 | 1 |
| Samhd1 | 1.13359248 | 0.180902097 | 0.31028425 | 1 |
| H19 | 1.10211559 | 0.140275536 | 0.3103388 | 1 |
| Cd276 | 1.08953983 | 0.123718937 | 0.31042957 | 1 |
| Mtus2 | 1.17223882 | 0.229266524 | 0.31054476 | 1 |
| LOC100125364 | 0.94183667 | -0.086451193 | 0.31064106 | 1 |
| Rwdd1 | 1.08348903 | 0.115684552 | 0.31102375 | 1 |
| Bdkrb2 | 1.09111677 | 0.125805509 | 0.31114557 | 1 |
| Bend5 | 1.17978206 | 0.238520376 | 0.31118263 | 1 |
| Eci3 | 0.9137181 | -0.130178959 | 0.31136594 | 1 |
| Cntnap4 | 0.91863098 | -0.122442657 | 0.31154393 | 1 |
| Rbbp5 | 1.06274773 | 0.087799171 | 0.31178252 | 1 |
| Wbp11 | 0.9356402 | -0.09597425 | 0.31179622 | 1 |
| Hexdc | 0.9056244 | -0.143015261 | 0.31180363 | 1 |
| Ccp110 | 1.05538178 | 0.07776498 | 0.31196744 | 1 |
| Zfp287 | 1.09250701 | 0.127642539 | 0.31214585 | 1 |
| Osbpl2 | 0.94163437 | -0.086761116 | 0.3122334 | 1 |
| Fkbpl | 0.91195281 | -0.132968928 | 0.31236766 | 1 |
| Tns1 | 0.95446294 | -0.06723892 | 0.3124541 | 1 |
| Desi1 | 0.93314584 | -0.099825512 | 0.31248042 | 1 |
| Cxcl12 | 1.0677416 | 0.094562548 | 0.31252292 | 1 |
| Enpep | 1.08119231 | 0.11262315 | 0.31252856 | 1 |
| Gna15 | 1.2535594 | 0.326030355 | 0.31254353 | 1 |
| Tbx15 | 1.21201893 | 0.277412231 | 0.31267676 | 1 |
| Vcan | 0.92763787 | -0.108366372 | 0.31272992 | 1 |
| Nkg7 | 1.38153192 | 0.466268894 | 0.31274243 | 1 |
| Cxcr2 | 1.40050662 | 0.485948808 | 0.31289472 | 1 |
| Naa25 | 0.9313771 | -0.102562681 | 0.31291584 | 1 |
| Lbh | 0.94717401 | -0.078298608 | 0.31296204 | 1 |
| Cxcl14 | 1.30851549 | 0.387931003 | 0.31299126 | 1 |
| Sema4f | 0.94018655 | -0.088981054 | 0.31313547 | 1 |
| Fbxl8 | 0.82168785 | -0.283337662 | 0.3131489 | 1 |
| Cox4i2 | 1.25539984 | 0.328146927 | 0.31317377 | 1 |
| Rpl27 | 1.05297845 | 0.074475913 | 0.31322068 | 1 |
| Rapgef2 | 0.93794934 | -0.092418094 | 0.31336727 | 1 |
| Capn15 | 0.92060608 | -0.119344123 | 0.31343996 | 1 |
| Rab40b | 0.85320626 | -0.229033549 | 0.31346814 | 1 |

| | | | | |
|---|---|---|---|---|
| Brk1 | 1.049907 | 0.070261535 | 0.31361255 | 1 |
| Dgcr6 | 1.06409755 | 0.089630409 | 0.31369724 | 1 |
| Eno4 | 0.81136975 | -0.301568583 | 0.31373785 | 1 |
| Cmip | 0.94999051 | -0.074014999 | 0.3137735 | 1 |
| Aspa | 1.06912131 | 0.096425556 | 0.31379484 | 1 |
| Cd19 | 1.2491952 | 0.320998929 | 0.31388852 | 1 |
| Mrpl53 | 1.08271149 | 0.114648862 | 0.3139301 | 1 |
| Kcnk16 | 1.30425493 | 0.383225886 | 0.31401841 | 1 |
| Sirpa | 1.07144838 | 0.099562341 | 0.31412208 | 1 |
| Zbtb10 | 0.88369513 | -0.178379365 | 0.31417793 | 1 |
| Pcsk4 | 0.84719942 | -0.239226491 | 0.31432683 | 1 |
| Hnmt | 1.28967718 | 0.367009994 | 0.31442534 | 1 |
| Slco3a1 | 0.92440063 | -0.113409852 | 0.31455088 | 1 |
| Hspe1 | 1.06760865 | 0.094382893 | 0.31479451 | 1 |
| Astn1 | 0.94790123 | -0.077191355 | 0.31492378 | 1 |
| Hint2 | 1.0870137 | 0.120370122 | 0.31505038 | 1 |
| Pkp2 | 0.81494 | -0.295234246 | 0.31506821 | 1 |
| Bank1 | 1.33115575 | 0.412679384 | 0.31507207 | 1 |
| Nme1 | 1.0527596 | 0.074176025 | 0.3151796 | 1 |
| Tbc1d25 | 0.91251167 | -0.13208509 | 0.3154117 | 1 |
| Ndufa1 | 1.06609163 | 0.092331445 | 0.31555005 | 1 |
| Slc25a38 | 1.07857389 | 0.109125021 | 0.31557169 | 1 |
| Rpa3 | 0.8917964 | -0.165213724 | 0.31571174 | 1 |
| Atraid | 0.91358607 | -0.130387441 | 0.31572715 | 1 |
| Ube2h | 0.95372343 | -0.068357138 | 0.3157495 | 1 |
| Ppargc1b | 0.83803544 | -0.254916843 | 0.31575214 | 1 |
| Tm2d3 | 1.14768263 | 0.198723751 | 0.31575725 | 1 |
| Smarcd2 | 0.93734345 | -0.093350336 | 0.31604361 | 1 |
| Rnf182 | 0.85006303 | -0.234358277 | 0.31655801 | 1 |
| Mib2 | 0.93364192 | -0.09905875 | 0.31658544 | 1 |
| Sema6b | 0.8916883 | -0.165388604 | 0.31663044 | 1 |
| P3h2 | 1.14953013 | 0.201044275 | 0.31691709 | 1 |
| Shisa5 | 0.94377638 | -0.083483032 | 0.31696462 | 1 |
| Asb8 | 1.05360864 | 0.075339079 | 0.31711397 | 1 |
| Urb1 | 0.93285009 | -0.10028284 | 0.31714794 | 1 |
| Ppp3cc | 1.07255804 | 0.101055714 | 0.3171664 | 1 |
| Smpd5 | 0.90331049 | -0.146706138 | 0.31724854 | 1 |
| Hnrnpa1 | 1.04811674 | 0.067799417 | 0.31729848 | 1 |
| Lzts3 | 0.91103556 | -0.134420725 | 0.31730119 | 1 |
| Myadm | 0.94929421 | -0.075072814 | 0.31741847 | 1 |

| | | | | |
|---|---|---|---|---|
| Ampd2 | 0.93758502 | -0.092978579 | 0.3176588 | 1 |
| Imp4 | 0.92269389 | -0.116075988 | 0.31778431 | 1 |
| Gmeb2 | 1.17015087 | 0.226694558 | 0.31790844 | 1 |
| Zc3h8 | 0.8956386 | -0.159011385 | 0.31792126 | 1 |
| Snn | 0.95276749 | -0.069803911 | 0.31799892 | 1 |
| Elp2 | 1.04878191 | 0.068714711 | 0.31814269 | 1 |
| RGD1305464 | 0.85485339 | -0.226251074 | 0.31837972 | 1 |
| Cttnbp2nl | 0.9458547 | -0.080309524 | 0.31839498 | 1 |
| Zfp395 | 1.11566468 | 0.157903483 | 0.31841872 | 1 |
| Stag1 | 0.93243403 | -0.100926433 | 0.31842453 | 1 |
| Mif | 0.94935213 | -0.074984784 | 0.31853765 | 1 |
| Slc8a3 | 0.91018503 | -0.135768232 | 0.31872257 | 1 |
| Slc35f1 | 0.94760082 | -0.077648648 | 0.31874747 | 1 |
| Nkain4 | 0.88249679 | -0.180337058 | 0.31887286 | 1 |
| Ddit4l2 | 1.30899719 | 0.388462 | 0.31887481 | 1 |
| Sdhc | 1.05124841 | 0.072103613 | 0.3188955 | 1 |
| Pqbp1 | 0.92566008 | -0.111445597 | 0.31935814 | 1 |
| Srebf1 | 0.9434176 | -0.084031577 | 0.31952546 | 1 |
| Fam192a | 1.07176877 | 0.099993681 | 0.31963999 | 1 |
| Plod3 | 0.93499175 | -0.096974459 | 0.32000305 | 1 |
| Trim8 | 0.9422638 | -0.085797079 | 0.32000808 | 1 |
| Fign | 0.8814976 | -0.181971451 | 0.3200448 | 1 |
| Thrap3 | 1.05014799 | 0.070592652 | 0.32009186 | 1 |
| Itfg2 | 1.13285148 | 0.179958729 | 0.32012458 | 1 |
| RGD1561931 | 1.16142929 | 0.21590132 | 0.32023589 | 1 |
| Zc3h10 | 0.90208652 | -0.148662291 | 0.32043786 | 1 |
| Mprip | 0.95535391 | -0.065892819 | 0.32046953 | 1 |
| Ubr7 | 1.07022915 | 0.097919735 | 0.32064119 | 1 |
| Ccdc62 | 0.8954336 | -0.159341634 | 0.3206924 | 1 |
| Adam15 | 0.94500008 | -0.081613646 | 0.32074619 | 1 |
| Coprs | 0.91424544 | -0.129346566 | 0.32074658 | 1 |
| Map2k7 | 1.09375954 | 0.129295597 | 0.32076973 | 1 |
| Eefsec | 0.9096825 | -0.136565 | 0.32078166 | 1 |
| Idh3B | 1.04688576 | 0.066104019 | 0.32085239 | 1 |
| Espl1 | 1.25899163 | 0.332268692 | 0.32085913 | 1 |
| Nrsn1 | 0.94584883 | -0.080318472 | 0.32097883 | 1 |
| Lin7c | 0.8979769 | -0.155249758 | 0.32098119 | 1 |
| Man2b1 | 0.94279334 | -0.084986521 | 0.32105493 | 1 |
| Tspan11 | 0.92237069 | -0.116581431 | 0.32106721 | 1 |
| Xpo1 | 0.95110465 | -0.072324004 | 0.32131939 | 1 |

| | | | | | |
|---|---|---|---|---|---|
| Il16 | | 1.0691677 | 0.096488156 | 0.32140042 | 1 |
| Vcam1 | | 1.0982888 | 0.135257464 | 0.32169924 | 1 |
| Kcnd2 | | 0.78247709 | -0.353879591 | 0.32177298 | 1 |
| Ppp2r1b | | 1.10238085 | 0.140622732 | 0.32220496 | 1 |
| Zfp398 | | 1.12087898 | 0.164630517 | 0.32233487 | 1 |
| Atmin | | 0.9452086 | -0.081295335 | 0.32234006 | 1 |
| Tug1 | | 0.94888664 | -0.075692358 | 0.32235115 | 1 |
| Ttc30a1 | | 0.83237881 | -0.264687853 | 0.3226046 | 1 |
| Ndor1 | | 0.90731215 | -0.140329114 | 0.32260903 | 1 |
| Hmgb3 | | 0.93526802 | -0.096548234 | 0.32285035 | 1 |
| Pign | | 1.09117351 | 0.12588052 | 0.32297862 | 1 |
| Ect2 | | 0.80282992 | -0.316833703 | 0.32300174 | 1 |
| Hal | | 0.80561112 | -0.311844492 | 0.32305511 | 1 |
| Pou4f2 | | 0.91598279 | -0.126607599 | 0.32321364 | 1 |
| Banp | | 0.92995886 | -0.104761199 | 0.32330655 | 1 |
| Pcbp4 | | 0.9551376 | -0.066219503 | 0.32333383 | 1 |
| Glud1 | | 0.9558242 | -0.065182796 | 0.32334696 | 1 |
| Pdzrn3 | | 0.92769318 | -0.108280358 | 0.32338713 | 1 |
| Elavl1 | | 0.9246673 | -0.112993729 | 0.32348284 | 1 |
| Runx1 | | 0.90091676 | -0.150534278 | 0.32358712 | 1 |
| Tmem252 | | 0.84675733 | -0.239979532 | 0.32359641 | 1 |
| Cadm2 | | 0.89160755 | -0.165519262 | 0.3237621 | 1 |
| Pou4f3 | | 1.07115113 | 0.099162048 | 0.32391232 | 1 |
| Myh9l1 | | 0.95579398 | -0.065228412 | 0.32393093 | 1 |
| Lypla2 | | 0.94395263 | -0.083213633 | 0.32397697 | 1 |
| Ppp2r3b | | 1.13946018 | 0.188350512 | 0.3240466 | 1 |
| Map4k1 | | 1.19409459 | 0.255917123 | 0.32423104 | 1 |
| Rbm14 | | 0.91710151 | -0.124846659 | 0.32431282 | 1 |
| Maea | | 1.04752959 | 0.066990999 | 0.32457257 | 1 |
| Plekha1 | | 0.94989458 | -0.074160689 | 0.32468791 | 1 |
| Rad23a | | 1.07779001 | 0.108076116 | 0.32478791 | 1 |
| Fbxo11 | | 0.94954595 | -0.074690274 | 0.32486775 | 1 |
| Tmod3 | | 0.9360759 | -0.095302581 | 0.32501301 | 1 |
| | 7-Sep | 1.04646486 | 0.06552387 | 0.32504351 | 1 |
| Add1 | | 0.95570223 | -0.065366913 | 0.3251107 | 1 |
| Slc35g2 | | 1.08624161 | 0.119345036 | 0.3251415 | 1 |
| Sympk | | 0.94677274 | -0.078909933 | 0.32523259 | 1 |
| Fundc1 | | 1.05154145 | 0.072505715 | 0.32528339 | 1 |
| Atp6v1h | | 1.04701779 | 0.066285962 | 0.32533979 | 1 |
| Syt9 | | 0.93742335 | -0.093227366 | 0.32550366 | 1 |

| | | | | |
|---|---|---|---|---|
| Pacsin1 | 0.94661316 | -0.079153116 | 0.32575199 | 1 |
| Dclk2 | 1.05374681 | 0.075528259 | 0.32584335 | 1 |
| Nfkbie | 0.89622719 | -0.158063598 | 0.3258982 | 1 |
| Edn1 | 1.25594485 | 0.328773112 | 0.32598842 | 1 |
| Sephs2 | 0.93315526 | -0.099810957 | 0.32613572 | 1 |
| Nedd8 | 1.04923 | 0.069330965 | 0.32615928 | 1 |
| Ly96 | 1.28178078 | 0.358149539 | 0.32616473 | 1 |
| Mdga1 | 0.91062924 | -0.135064303 | 0.32618664 | 1 |
| Chst5 | 0.87409526 | -0.194137583 | 0.32622495 | 1 |
| Med29 | 1.08698707 | 0.120334778 | 0.32626 | 1 |
| Rpl9 | 1.04635559 | 0.065373214 | 0.32631495 | 1 |
| Gtpbp10 | 1.14741296 | 0.198384724 | 0.32640543 | 1 |
| Rab2a | 1.05323358 | 0.074825419 | 0.32646098 | 1 |
| Ctsc | 0.93132288 | -0.102646667 | 0.3264904 | 1 |
| Smad3 | 1.10022477 | 0.137798292 | 0.32658947 | 1 |
| Polr3d | 1.06407193 | 0.089595676 | 0.32661577 | 1 |
| Galnt6 | 0.86988028 | -0.201111239 | 0.32700641 | 1 |
| Tal1 | 1.19755616 | 0.260093319 | 0.32704627 | 1 |
| Mn1 | 0.79557834 | -0.329924101 | 0.32715952 | 1 |
| Zfp518a | 1.07751845 | 0.107712573 | 0.32722472 | 1 |
| Rnf144b | 1.15872727 | 0.212541037 | 0.32727325 | 1 |
| Ppargc1a | 1.09824738 | 0.135203054 | 0.32742098 | 1 |
| Bag3 | 0.9537257 | -0.068353707 | 0.32743628 | 1 |
| St3gal2 | 0.94697392 | -0.078603404 | 0.32747233 | 1 |
| Chst11 | 0.90318457 | -0.146907254 | 0.32748573 | 1 |
| Npap60 | 0.93852445 | -0.091533758 | 0.3275167 | 1 |
| Nek11 | 0.84253048 | -0.24719921 | 0.3277113 | 1 |
| Pes1 | 0.94414486 | -0.082919861 | 0.32776271 | 1 |
| Phactr2 | 1.06698047 | 0.093533764 | 0.32778112 | 1 |
| Ddx21 | 1.05823129 | 0.081654983 | 0.32778739 | 1 |
| Mrps30 | 1.06296624 | 0.088095784 | 0.32784239 | 1 |
| Mctp2 | 1.06032695 | 0.08450919 | 0.32794835 | 1 |
| Gtpbp4 | 1.0859723 | 0.118987311 | 0.32821688 | 1 |
| Kdm2a | 0.95250358 | -0.070203587 | 0.32822638 | 1 |
| Tbp | 1.10973261 | 0.150212096 | 0.32838773 | 1 |
| Papd7 | 0.92815808 | -0.107557559 | 0.32838987 | 1 |
| Mrap | 1.09513158 | 0.131104222 | 0.32854134 | 1 |
| Ddx6 | 0.94421422 | -0.082813891 | 0.32860985 | 1 |
| Hipk1 | 0.95482494 | -0.066691844 | 0.32874056 | 1 |
| Meox1 | 1.26344266 | 0.337360188 | 0.32877952 | 1 |

| | | | | |
|---|---|---|---|---|
| Prkci | 0.9519195 | -0.071088525 | 0.32879787 | 1 |
| Csrnp1 | 0.83374589 | -0.262320355 | 0.32888955 | 1 |
| Mbd2 | 0.95042926 | -0.073348845 | 0.32908204 | 1 |
| Cdc6 | 0.75816648 | -0.399413422 | 0.32909806 | 1 |
| Mdh1 | 1.04712159 | 0.06642897 | 0.32920313 | 1 |
| Psmd10 | 0.89305303 | -0.163182247 | 0.32923475 | 1 |
| Tead1 | 0.87029696 | -0.200420337 | 0.32939286 | 1 |
| RGD1309821 | 0.9314716 | -0.102416308 | 0.32939446 | 1 |
| Rnf167 | 1.05754692 | 0.080721679 | 0.32967096 | 1 |
| Nsmce2 | 0.91011573 | -0.135878078 | 0.32975963 | 1 |
| Kcnk12 | 1.13615042 | 0.184153851 | 0.32993919 | 1 |
| Spred3 | 0.84071563 | -0.250310203 | 0.32994528 | 1 |
| Hemk1 | 1.15061596 | 0.20240639 | 0.33005136 | 1 |
| Taf11 | 1.08409214 | 0.116487385 | 0.33006237 | 1 |
| C5ar2 | 1.25344949 | 0.325903862 | 0.33016869 | 1 |
| Plcxd3 | 0.93661631 | -0.09446994 | 0.33023734 | 1 |
| Msl3 | 1.06485937 | 0.09066291 | 0.3302438 | 1 |
| Zfp18 | 1.12139782 | 0.165298172 | 0.33025165 | 1 |
| Aldh3b1 | 0.86988411 | -0.201104885 | 0.33067193 | 1 |
| Pygl | 1.07055543 | 0.098359502 | 0.33073059 | 1 |
| Tom1l2 | 0.95286979 | -0.069649013 | 0.33090558 | 1 |
| Hspb11 | 1.13933004 | 0.188185727 | 0.33100052 | 1 |
| Cox8a | 1.04679514 | 0.065979129 | 0.33106983 | 1 |
| Cntn3 | 0.91048049 | -0.135299985 | 0.33117787 | 1 |
| Tm9sf2 | 0.95611398 | -0.064745473 | 0.33127813 | 1 |
| Evi2b | 1.20409022 | 0.267943494 | 0.33136223 | 1 |
| Ppil1 | 1.09140486 | 0.126186377 | 0.33139975 | 1 |
| Dnajc27 | 0.95380318 | -0.068236509 | 0.33145452 | 1 |
| Slc35e3 | 0.94783354 | -0.077294386 | 0.33152751 | 1 |
| Atp5sl | 0.93693337 | -0.093981646 | 0.33155682 | 1 |
| Scx | 1.12523453 | 0.170225733 | 0.3315963 | 1 |
| Nupl2 | 1.12272196 | 0.167000692 | 0.33176821 | 1 |
| Sipa1l3 | 0.89420545 | -0.161321757 | 0.33177944 | 1 |
| Ldhd | 1.11449772 | 0.156393671 | 0.33184727 | 1 |
| Mrpl27 | 0.93526709 | -0.096549668 | 0.33201659 | 1 |
| Manf | 0.91630633 | -0.126098116 | 0.3320603 | 1 |
| Cyb5d2 | 1.13641307 | 0.184487326 | 0.33212511 | 1 |
| Zfp167 | 0.84764641 | -0.238465521 | 0.33214056 | 1 |
| Anxa2 | 1.0476708 | 0.067185467 | 0.33231893 | 1 |
| LOC683897 | 1.08354579 | 0.115760116 | 0.33255135 | 1 |

| | | | | |
|---|---|---|---|---|
| Hmox2 | 1.04897995 | 0.068987109 | 0.33259894 | 1 |
| Ppp4r4 | 1.0649854 | 0.090833649 | 0.33277368 | 1 |
| Edc4 | 1.06088824 | 0.085272678 | 0.33279916 | 1 |
| Rhd | 1.21943187 | 0.286209151 | 0.33286718 | 1 |
| Rfxap | 0.91819356 | -0.123129776 | 0.33290118 | 1 |
| Mterf2 | 1.12705153 | 0.172553477 | 0.33302162 | 1 |
| Ncmap | 0.93504108 | -0.09689835 | 0.33306304 | 1 |
| Tap1 | 1.10375874 | 0.142424861 | 0.33340997 | 1 |
| Scaf11 | 1.05223214 | 0.073453021 | 0.3334321 | 1 |
| Tp53i11 | 0.92530789 | -0.111994605 | 0.33370791 | 1 |
| Mrpl55 | 0.92188071 | -0.117348019 | 0.33373046 | 1 |
| Adam23 | 1.04526893 | 0.063874169 | 0.33377986 | 1 |
| Tiam1 | 0.92514372 | -0.112250598 | 0.33381959 | 1 |
| Slc44a2 | 0.9558297 | -0.0651745 | 0.33395028 | 1 |
| Ube2m | 0.95272459 | -0.069868874 | 0.33414828 | 1 |
| Cox5b | 1.04792667 | 0.067537763 | 0.33427967 | 1 |
| RGD1565033 | 1.11392718 | 0.155654924 | 0.33432621 | 1 |
| Sh3bgrl3 | 0.95628148 | -0.064492766 | 0.33437715 | 1 |
| Rnf166 | 1.09596796 | 0.132205617 | 0.33438438 | 1 |
| Setmar | 1.12238486 | 0.166567456 | 0.33453133 | 1 |
| Tiprl | 1.06146923 | 0.086062549 | 0.33458826 | 1 |
| Clpx | 1.07139989 | 0.099497057 | 0.33465581 | 1 |
| Npepl1 | 0.92197651 | -0.117198105 | 0.33474176 | 1 |
| Tmem248 | 1.0652905 | 0.091246902 | 0.33477944 | 1 |
| Stat1 | 1.04851635 | 0.068349353 | 0.33479749 | 1 |
| Kansl3 | 0.95050619 | -0.073232068 | 0.33480945 | 1 |
| Ap5b1 | 0.90714775 | -0.140590551 | 0.33486668 | 1 |
| Slc1a4 | 0.93286048 | -0.100266771 | 0.33498434 | 1 |
| Fgf7 | 1.16397882 | 0.219064804 | 0.33501069 | 1 |
| Lars | 1.04614442 | 0.065082031 | 0.33521961 | 1 |
| Ldlrap1 | 1.13480613 | 0.182445854 | 0.33525364 | 1 |
| Ppm1h | 0.94532464 | -0.081118229 | 0.33525776 | 1 |
| Rnf25 | 1.07748888 | 0.107672975 | 0.33526205 | 1 |
| Ofd1 | 0.91613764 | -0.126363724 | 0.33530675 | 1 |
| Kcnq5 | 1.05898594 | 0.082683437 | 0.3354076 | 1 |
| Clstn1 | 0.95616394 | -0.064670098 | 0.33542526 | 1 |
| Cdca7 | 1.24635099 | 0.317710404 | 0.33549153 | 1 |
| Eri1 | 1.08683468 | 0.12013251 | 0.33553513 | 1 |
| Ccdc94 | 1.08069058 | 0.111953512 | 0.33555351 | 1 |
| Heca | 0.94049522 | -0.088507485 | 0.33568874 | 1 |

| | | | | |
|---|---|---|---|---|
| Nek2 | 0.78013234 | -0.358209217 | 0.3357782 | 1 |
| Itgb1bp1 | 1.05673521 | 0.079613918 | 0.33580731 | 1 |
| Tctn2 | 0.88189031 | -0.181328877 | 0.33584671 | 1 |
| Tpt1 | 1.04654288 | 0.065631418 | 0.33601436 | 1 |
| Gnpda1 | 1.08352811 | 0.115736584 | 0.33615195 | 1 |
| Disp1 | 1.07834108 | 0.108813579 | 0.33616372 | 1 |
| Nr1d2 | 0.9553692 | -0.065869726 | 0.33622971 | 1 |
| Dnaaf3 | 1.21789485 | 0.284389584 | 0.33623358 | 1 |
| Plxnb3 | 0.94756467 | -0.077703687 | 0.33625634 | 1 |
| Fas | 0.86762561 | -0.204855456 | 0.33632653 | 1 |
| Rnf125 | 0.91446822 | -0.128995067 | 0.33646226 | 1 |
| Psmd5 | 1.05919464 | 0.082967732 | 0.33658027 | 1 |
| Ndufs7 | 1.05972486 | 0.083689744 | 0.33658615 | 1 |
| Cnbd2 | 1.1963459 | 0.258634581 | 0.33676477 | 1 |
| Sos2 | 0.9411221 | -0.087546186 | 0.33677739 | 1 |
| Bag6 | 0.95337121 | -0.068890033 | 0.33678873 | 1 |
| Slc43a3 | 1.08745277 | 0.120952748 | 0.33679505 | 1 |
| Cryl1 | 1.14235788 | 0.192014695 | 0.33692623 | 1 |
| Cyld | 0.93810871 | -0.092172986 | 0.33697878 | 1 |
| Mtrf1l | 0.89138795 | -0.165874629 | 0.33699786 | 1 |
| LOC100911576 | 1.0477761 | 0.067330459 | 0.33713684 | 1 |
| Ergic3 | 0.94692237 | -0.07868194 | 0.33714063 | 1 |
| Rpl23a | 1.04498296 | 0.063479424 | 0.33717246 | 1 |
| Snx17 | 0.9376755 | -0.09283936 | 0.33719179 | 1 |
| Eif3c | 1.0450816 | 0.063615589 | 0.33725402 | 1 |
| Polr3h | 1.0891755 | 0.12323643 | 0.33728041 | 1 |
| Pgap1 | 0.90326707 | -0.146775486 | 0.33736208 | 1 |
| Kpna4 | 0.94325402 | -0.084281756 | 0.33743501 | 1 |
| Dctn4 | 1.05737611 | 0.080488636 | 0.33752043 | 1 |
| Rac1 | 0.95487448 | -0.066616996 | 0.33768327 | 1 |
| Gbp4 | 1.19021451 | 0.251221606 | 0.33770995 | 1 |
| Fam81a | 0.90796582 | -0.139290102 | 0.33776738 | 1 |
| Dpm2 | 1.0824345 | 0.114279729 | 0.3378797 | 1 |
| Bzw2 | 0.92071479 | -0.11917378 | 0.33794796 | 1 |
| Psma2 | 1.04684952 | 0.066054077 | 0.33812314 | 1 |
| Nln | 1.07620703 | 0.105955639 | 0.33812454 | 1 |
| Dnaaf2 | 1.10260671 | 0.140918289 | 0.33813046 | 1 |
| Pm20d1 | 0.81618296 | -0.293035504 | 0.33815455 | 1 |
| Gnai2 | 0.9570962 | -0.063264147 | 0.33839294 | 1 |
| Rab2b | 1.06206108 | 0.086866741 | 0.33840715 | 1 |

| | | | | |
|---|---|---|---|---|
| Sdhb | 0.95431317 | -0.067465316 | 0.33840795 | 1 |
| Tmem196 | 1.11923898 | 0.162518117 | 0.33854769 | 1 |
| Bmpr1b | 1.1891768 | 0.249963226 | 0.33864154 | 1 |
| Ndufa2 | 1.06497516 | 0.090819783 | 0.33870274 | 1 |
| RGD1564036 | 1.19970139 | 0.262675358 | 0.33871409 | 1 |
| Lrg1 | 1.12834667 | 0.174210379 | 0.3387348 | 1 |
| Ltbr | 0.92936447 | -0.105683602 | 0.33883264 | 1 |
| Adh5 | 1.05254287 | 0.073878994 | 0.33889452 | 1 |
| Tbc1d9 | 0.95577378 | -0.065258909 | 0.33898159 | 1 |
| Pdgfd | 1.11311383 | 0.154601139 | 0.3390681 | 1 |
| Igf2r | 0.95492181 | -0.066545479 | 0.33907385 | 1 |
| Clec11a | 1.137014 | 0.185250021 | 0.33926745 | 1 |
| Palm2 | 0.88732793 | -0.17246072 | 0.3392949 | 1 |
| Paqr7 | 0.9212103 | -0.118397554 | 0.33930176 | 1 |
| Trpc5 | 1.20840996 | 0.273109982 | 0.33935781 | 1 |
| Sestd1 | 0.94006992 | -0.089160023 | 0.33945421 | 1 |
| Sox13 | 0.91952102 | -0.121045537 | 0.33949654 | 1 |
| Atxn2l | 0.95397802 | -0.067972067 | 0.33953494 | 1 |
| Rps6 | 1.04501195 | 0.063519439 | 0.33960189 | 1 |
| Zfp397 | 0.86351908 | -0.211700044 | 0.33974322 | 1 |
| Kdm1b | 0.91901527 | -0.121839258 | 0.33975613 | 1 |
| Rab13 | 0.87680927 | -0.18966505 | 0.33981554 | 1 |
| Pskh1 | 1.17431428 | 0.231818572 | 0.33995071 | 1 |
| Ulk3 | 0.86387813 | -0.211100294 | 0.34011901 | 1 |
| Cep131 | 0.92487003 | -0.112677451 | 0.34023427 | 1 |
| Slamf8 | 1.29854978 | 0.376901326 | 0.34034685 | 1 |
| Ints4 | 0.94462917 | -0.082180015 | 0.34038672 | 1 |
| Dbndd2 | 0.94519074 | -0.081322598 | 0.34040852 | 1 |
| Nfx1 | 1.05976438 | 0.083743548 | 0.34047372 | 1 |
| Ccdc64 | 0.91993001 | -0.120403994 | 0.34054036 | 1 |
| Trpm3 | 0.95128836 | -0.072045366 | 0.34058838 | 1 |
| Rcor1 | 0.93816783 | -0.092082067 | 0.34059868 | 1 |
| RGD1308147 | 1.06112315 | 0.085592106 | 0.34063287 | 1 |
| Zcchc17 | 1.0561489 | 0.078813241 | 0.34070674 | 1 |
| Gypc | 1.11028915 | 0.150935448 | 0.34073926 | 1 |
| Cmtm7 | 0.88544297 | -0.175528703 | 0.34076852 | 1 |
| Zbtb33 | 0.91614042 | -0.12635935 | 0.34082922 | 1 |
| Ppp1r35 | 1.17576239 | 0.233596536 | 0.34094325 | 1 |
| Grik2 | 1.08669307 | 0.11994452 | 0.34095987 | 1 |
| Cir1 | 1.07621804 | 0.105970398 | 0.3410585 | 1 |

| | | | | |
|---|---|---|---|---|
| Wtap | 1.04755719 | 0.067029011 | 0.34117048 | 1 |
| Zfp295 | 0.91304379 | -0.131244045 | 0.34119849 | 1 |
| Mgat3 | 0.94627499 | -0.0796686 | 0.34126018 | 1 |
| Hmgn2 | 0.94729752 | -0.078110488 | 0.34128767 | 1 |
| Pabpc4 | 0.93631071 | -0.094940735 | 0.34161376 | 1 |
| Xpo6 | 0.9522112 | -0.070646492 | 0.34181298 | 1 |
| Fzd4 | 0.9287541 | -0.106631422 | 0.34200253 | 1 |
| Dtx3l | 1.08179584 | 0.113428254 | 0.34209669 | 1 |
| Scube1 | 1.08317243 | 0.11526293 | 0.34237567 | 1 |
| Rragd | 0.89483185 | -0.16031148 | 0.34243078 | 1 |
| Atp2b2 | 0.95672082 | -0.0638301 | 0.34248105 | 1 |
| Klhl31 | 0.66372726 | -0.591337562 | 0.34257736 | 1 |
| Sertad4 | 0.91825872 | -0.123027405 | 0.34258097 | 1 |
| Slc7a11 | 0.90562603 | -0.143012663 | 0.34259693 | 1 |
| Rnf135 | 1.19664231 | 0.258991974 | 0.34272542 | 1 |
| Crlf2 | 0.87614802 | -0.190753471 | 0.34272726 | 1 |
| Cox16 | 1.09898263 | 0.13616858 | 0.34290896 | 1 |
| Nelfe | 0.92055251 | -0.119428082 | 0.34306173 | 1 |
| Gan | 1.1102893 | 0.150935634 | 0.34306621 | 1 |
| Nsa2 | 1.04946537 | 0.069654566 | 0.34314337 | 1 |
| Hsd17b11 | 0.95368686 | -0.068412458 | 0.3433375 | 1 |
| Slc38a3 | 1.18579888 | 0.245859336 | 0.34364551 | 1 |
| LOC691921 | 1.19507354 | 0.257099393 | 0.34380258 | 1 |
| Sik3 | 0.93922169 | -0.090462375 | 0.34408033 | 1 |
| Lrwd1 | 0.89680119 | -0.157139898 | 0.34426125 | 1 |
| Ldb1 | 0.94201643 | -0.086175872 | 0.34426389 | 1 |
| Dpysl3 | 0.95637466 | -0.064352188 | 0.34443853 | 1 |
| Zpbp2 | 1.27352766 | 0.348830298 | 0.34459796 | 1 |
| Jade2 | 0.92759352 | -0.108435346 | 0.34459981 | 1 |
| Hr | 0.93096802 | -0.103196479 | 0.34494767 | 1 |
| Rere | 0.95125921 | -0.072089581 | 0.34501167 | 1 |
| Slc22a6 | 1.18792718 | 0.248446402 | 0.34506814 | 1 |
| Kcnn4 | 1.11748151 | 0.160250962 | 0.34508082 | 1 |
| Cntnap1 | 0.95777221 | -0.062245514 | 0.34511733 | 1 |
| Zfp2 | 1.19680217 | 0.25918469 | 0.34522926 | 1 |
| Ankrd10 | 0.92910157 | -0.10609178 | 0.34527856 | 1 |
| Anxa7 | 1.04615027 | 0.065090093 | 0.34563131 | 1 |
| Spi1 | 1.17493901 | 0.232585873 | 0.34572654 | 1 |
| Apoc1 | 1.24272125 | 0.313502729 | 0.3458979 | 1 |
| Yeats4 | 1.07398506 | 0.102973929 | 0.3459363 | 1 |

| | | | | |
|---|---|---|---|---|
| Cdh5 | 1.06687711 | 0.093394004 | 0.3459488 | 1 |
| Pdcd4 | 1.05531583 | 0.077674822 | 0.34610198 | 1 |
| Zfp653 | 0.88563628 | -0.175213777 | 0.34614709 | 1 |
| Irak1 | 0.9465892 | -0.079189639 | 0.34623919 | 1 |
| Ptpn1 | 1.10867893 | 0.148841629 | 0.34628831 | 1 |
| MAST1 | 0.95742845 | -0.062763423 | 0.34631299 | 1 |
| Cpa3 | 1.23411731 | 0.303479534 | 0.34637896 | 1 |
| Rpl29 | 1.04423374 | 0.062444677 | 0.34641275 | 1 |
| H2afz | 0.94466434 | -0.082126296 | 0.34644929 | 1 |
| Nt5dc3 | 0.91550381 | -0.127362211 | 0.34645348 | 1 |
| Samd10 | 1.13749693 | 0.185862646 | 0.34646437 | 1 |
| Prkab1 | 1.09625461 | 0.132582904 | 0.34656919 | 1 |
| Slc38a9 | 1.10669134 | 0.146252904 | 0.34658003 | 1 |
| Tmem129 | 0.92720793 | -0.10903519 | 0.346619 | 1 |
| Cdh15 | 0.94271664 | -0.085103902 | 0.3468679 | 1 |
| Mlc1 | 1.20002711 | 0.263067 | 0.34691193 | 1 |
| Atl2 | 1.05819592 | 0.081606765 | 0.34695443 | 1 |
| Tspan4 | 0.93203468 | -0.101544459 | 0.34698049 | 1 |
| Ranbp10 | 0.94558177 | -0.080725869 | 0.34718715 | 1 |
| Hsd17b8 | 1.12651073 | 0.171861056 | 0.34721671 | 1 |
| Ptpn9 | 1.05895109 | 0.082635963 | 0.34723479 | 1 |
| Pcdhb4 | 0.83429299 | -0.261373979 | 0.34724233 | 1 |
| Mvp | 0.94538869 | -0.081020486 | 0.34725822 | 1 |
| Plac9 | 0.89924388 | -0.153215657 | 0.34740608 | 1 |
| Mgst1 | 0.94107682 | -0.087615593 | 0.34771896 | 1 |
| Sgpl1 | 1.06314828 | 0.088342823 | 0.34771908 | 1 |
| Arhgap9 | 1.16780308 | 0.223797026 | 0.34786112 | 1 |
| Rarg | 0.93316053 | -0.099802809 | 0.34789925 | 1 |
| Prmt5 | 0.94519474 | -0.081316488 | 0.34791183 | 1 |
| Mpv17l2 | 1.13223928 | 0.179178881 | 0.34791799 | 1 |
| Ppp1r12a | 0.94556214 | -0.080755826 | 0.34800571 | 1 |
| Clpb | 0.93428439 | -0.098066328 | 0.34808102 | 1 |
| Elovl1 | 0.95375601 | -0.06830785 | 0.34808248 | 1 |
| Orc3 | 1.07847702 | 0.108995432 | 0.34816425 | 1 |
| Trip11 | 0.94936435 | -0.074966222 | 0.34817529 | 1 |
| G3bp2 | 1.04514619 | 0.063704751 | 0.34840813 | 1 |
| Prtfdc1 | 1.0739248 | 0.102892973 | 0.34848363 | 1 |
| Tmem101 | 1.08435029 | 0.116830877 | 0.34849197 | 1 |
| Hrh3 | 0.85890924 | -0.219422398 | 0.34856027 | 1 |
| Gata1 | 1.45193604 | 0.537977906 | 0.34858163 | 1 |

| | | | | |
|---|---|---|---|---|
| Srf | 1.08038919 | 0.111551106 | 0.34863068 | 1 |
| Mchr1 | 1.24335993 | 0.314243992 | 0.34890155 | 1 |
| Tmem14c | 1.05293776 | 0.074420154 | 0.34893739 | 1 |
| Epha3 | 0.83076622 | -0.267485533 | 0.3489518 | 1 |
| LOC687399 | 0.81439108 | -0.29620634 | 0.34929397 | 1 |
| Bax | 1.08646555 | 0.119642429 | 0.3493071 | 1 |
| Gpr176 | 1.08206323 | 0.113784801 | 0.3494145 | 1 |
| Ppp1r11 | 0.95364673 | -0.068473159 | 0.34941563 | 1 |
| S1pr4 | 0.80280776 | -0.316873534 | 0.34957677 | 1 |
| Rsl1d1 | 1.06772228 | 0.094536448 | 0.3496378 | 1 |
| Il13ra2 | 1.20886333 | 0.273651151 | 0.34974984 | 1 |
| Cacnb4 | 1.05575587 | 0.078276263 | 0.34993165 | 1 |
| Gpsm3 | 1.17173372 | 0.228644746 | 0.34994162 | 1 |
| Lsr | 1.1371134 | 0.18537614 | 0.34997084 | 1 |
| Lrrc16a | 0.93782993 | -0.092601767 | 0.34998954 | 1 |
| Atp6v1g1 | 1.04413711 | 0.062311172 | 0.3500125 | 1 |
| Dnm1l | 1.0438427 | 0.061904325 | 0.35012348 | 1 |
| Zfyve19 | 1.10137502 | 0.139305789 | 0.35020938 | 1 |
| C2cd3 | 0.92524317 | -0.112095516 | 0.35022059 | 1 |
| Lims2 | 0.94636639 | -0.079529262 | 0.35036882 | 1 |
| Dusp22 | 1.08920984 | 0.123281925 | 0.35039843 | 1 |
| Slc16a1 | 1.04741386 | 0.066831606 | 0.35043795 | 1 |
| Adipor1 | 1.04408658 | 0.062241349 | 0.3505019 | 1 |
| Ccdc163 | 1.14068185 | 0.189896463 | 0.35074747 | 1 |
| Jrk | 1.15944685 | 0.213436685 | 0.35079208 | 1 |
| Stim1 | 0.95187998 | -0.071148414 | 0.35095008 | 1 |
| Pdgfa | 0.92467787 | -0.112977238 | 0.35114135 | 1 |
| Slc35e2b | 0.91290525 | -0.131462969 | 0.35122303 | 1 |
| Vps26b | 0.9505326 | -0.07319199 | 0.35130984 | 1 |
| Med10 | 1.07127759 | 0.099332356 | 0.35131871 | 1 |
| Xpo4 | 0.94258691 | -0.08530245 | 0.35151219 | 1 |
| Vrk1 | 1.09013156 | 0.124502252 | 0.35155327 | 1 |
| Nthl1 | 1.14428606 | 0.194447753 | 0.35162505 | 1 |
| Lrrc46 | 0.85377244 | -0.22807651 | 0.3516574 | 1 |
| Shoc2 | 0.95035733 | -0.073458035 | 0.35165877 | 1 |
| Fam168a | 0.95596998 | -0.064962774 | 0.3516892 | 1 |
| Wdr60 | 0.917478 | -0.124254527 | 0.35172252 | 1 |
| Gpr182 | 1.23564967 | 0.305269772 | 0.35186527 | 1 |
| Zfp867 | 1.22611489 | 0.29409417 | 0.35196467 | 1 |
| Naa35 | 1.05902053 | 0.082730552 | 0.3519838 | 1 |

| | | | | |
|---|---|---|---|---|
| Arhgap39 | 0.94812896 | -0.076844794 | 0.35200521 | 1 |
| Gosr2 | 1.05325319 | 0.07485228 | 0.35202525 | 1 |
| RGD1307704 | 0.87918161 | -0.185766891 | 0.35210538 | 1 |
| Ephb1 | 0.92513005 | -0.112271916 | 0.35214934 | 1 |
| Psma3 | 0.94182568 | -0.086468039 | 0.35215116 | 1 |
| Hmgcs2 | 0.94425816 | -0.082746744 | 0.3524781 | 1 |
| Rrp7a | 1.05170749 | 0.072733507 | 0.3524856 | 1 |
| Arrb2 | 1.06600145 | 0.092209398 | 0.35248722 | 1 |
| Zfp597 | 1.12311679 | 0.167507953 | 0.35250588 | 1 |
| Gorab | 1.09773731 | 0.134532863 | 0.35267682 | 1 |
| Rph3a | 0.95828041 | -0.061480218 | 0.35284485 | 1 |
| Cx3cl1 | 0.87351359 | -0.195097948 | 0.35297746 | 1 |
| Tmco1 | 1.05363643 | 0.075377135 | 0.35301129 | 1 |
| Ebf3 | 0.94405482 | -0.083057456 | 0.35316566 | 1 |
| Steap2 | 0.90219087 | -0.148495408 | 0.35317229 | 1 |
| Angptl1 | 1.15163294 | 0.203680956 | 0.35318246 | 1 |
| Atp23 | 1.16816494 | 0.224243989 | 0.35329155 | 1 |
| Samd8 | 1.07812538 | 0.108524968 | 0.35331786 | 1 |
| Ror2 | 0.84967303 | -0.235020324 | 0.35344372 | 1 |
| Phlpp2 | 0.93144104 | -0.10246365 | 0.35361664 | 1 |
| Serpinb10 | 1.2345615 | 0.303998702 | 0.3537971 | 1 |
| Hba-a1 | 0.92862858 | -0.106826405 | 0.35380854 | 1 |
| Zfp57 | 0.85746507 | -0.221850188 | 0.3538162 | 1 |
| Ptger2 | 0.80599267 | -0.311161375 | 0.35401438 | 1 |
| Ogfod2 | 1.13086905 | 0.17743188 | 0.3542824 | 1 |
| Rcor2 | 0.90560809 | -0.143041249 | 0.35428469 | 1 |
| Elp4 | 0.88461539 | -0.176877748 | 0.3543091 | 1 |
| Cabin1 | 0.94349342 | -0.083915634 | 0.35434476 | 1 |
| Slc39a10 | 0.95459309 | -0.067042208 | 0.35439745 | 1 |
| Adap1 | 0.95539697 | -0.065827793 | 0.35441563 | 1 |
| Bcs1l | 1.09907844 | 0.136294348 | 0.35446874 | 1 |
| Ythdf1 | 1.04972309 | 0.070008806 | 0.35458479 | 1 |
| Klra1 | 1.18201679 | 0.241250529 | 0.35459341 | 1 |
| Nepro | 0.8718995 | -0.197766237 | 0.35482125 | 1 |
| Smarcal1 | 1.08489701 | 0.11755809 | 0.35489669 | 1 |
| Ccdc127 | 0.93410755 | -0.098339426 | 0.35493825 | 1 |
| Taf9 | 0.94945597 | -0.074826999 | 0.35497886 | 1 |
| Chrna5 | 1.13104202 | 0.177652532 | 0.35498337 | 1 |
| Plekhm1 | 0.92518128 | -0.112192014 | 0.3550414 | 1 |
| Otub1 | 0.95392009 | -0.068059673 | 0.35507324 | 1 |

| | | | | |
|---|---|---|---|---|
| Thumpd2 | 1.13501797 | 0.182715143 | 0.35511429 | 1 |
| Arl2 | 1.0669781 | 0.093530571 | 0.35517393 | 1 |
| Kcnn3 | 0.89661654 | -0.157436989 | 0.35521892 | 1 |
| Ccdc71 | 1.08437224 | 0.116860092 | 0.35532777 | 1 |
| Acp2 | 1.05377784 | 0.075570752 | 0.3555734 | 1 |
| Rp2 | 1.11198412 | 0.153136182 | 0.35565392 | 1 |
| Gpr173 | 0.81078008 | -0.302617452 | 0.35573795 | 1 |
| Sf3b4 | 0.9285437 | -0.106958288 | 0.35588054 | 1 |
| Snx9 | 0.93385793 | -0.098725005 | 0.3560187 | 1 |
| Eif4b | 1.04367476 | 0.0616722 | 0.3560216 | 1 |
| Eif3d | 0.94640499 | -0.079470418 | 0.35605925 | 1 |
| Lactb2 | 1.06051521 | 0.084765308 | 0.35613509 | 1 |
| Gopc | 0.95266082 | -0.069965433 | 0.35624317 | 1 |
| Zfp592 | 0.94229536 | -0.085748755 | 0.35625957 | 1 |
| Gpx4 | 1.04328027 | 0.061126774 | 0.35628135 | 1 |
| Mpg | 1.15224402 | 0.204446274 | 0.35628296 | 1 |
| Retnlg | 0.75703571 | -0.401566735 | 0.35629305 | 1 |
| Alms1 | 1.06288777 | 0.087989269 | 0.35631396 | 1 |
| Cyyr1 | 1.07422401 | 0.103294868 | 0.35641432 | 1 |
| Atp5s | 0.91770252 | -0.123901529 | 0.35654563 | 1 |
| Wdr20 | 0.92888559 | -0.106427176 | 0.35660283 | 1 |
| Arel1 | 1.04452011 | 0.062840264 | 0.3566467 | 1 |
| Arrdc1 | 0.93967079 | -0.089772693 | 0.35673261 | 1 |
| Sgcg | 1.25451705 | 0.32713208 | 0.35680514 | 1 |
| Cnot7 | 1.05019695 | 0.070659916 | 0.35700042 | 1 |
| Prr5 | 1.09099236 | 0.125641005 | 0.35711592 | 1 |
| Rplp2 | 1.06549238 | 0.091520272 | 0.35722506 | 1 |
| Xbp1 | 0.9539727 | -0.067980118 | 0.35729521 | 1 |
| Dffb | 0.84378474 | -0.245053104 | 0.35730827 | 1 |
| Sh3bgr | 1.11955002 | 0.162918989 | 0.35748701 | 1 |
| Bhlhb9 | 1.05641652 | 0.07917877 | 0.35760185 | 1 |
| Usp10 | 0.93938495 | -0.09021161 | 0.35768664 | 1 |
| Ubxn7 | 1.21387343 | 0.279618004 | 0.35784385 | 1 |
| Myo1e | 0.95696693 | -0.063459023 | 0.35786592 | 1 |
| Lrrc45 | 1.06328118 | 0.088523167 | 0.3578981 | 1 |
| Smc2 | 0.93416375 | -0.098252637 | 0.35790414 | 1 |
| Rap2b | 0.80739974 | -0.30864497 | 0.35791862 | 1 |
| Wdr45b | 1.06941067 | 0.096815982 | 0.35795 | 1 |
| Ttc9 | 0.9576673 | -0.062403554 | 0.35820768 | 1 |
| Neurl1 | 0.87718632 | -0.189044776 | 0.35836806 | 1 |

| | | | | |
|---|---|---|---|---|
| Kcnn1 | 0.90135821 | -0.149827527 | 0.35841674 | 1 |
| Nae1 | 1.06038856 | 0.084593007 | 0.35851576 | 1 |
| Fxyd7 | 1.0437023 | 0.061710262 | 0.35868604 | 1 |
| Polk | 1.09558149 | 0.131696793 | 0.35871808 | 1 |
| Hpgds | 1.33483028 | 0.416656319 | 0.35889629 | 1 |
| Blcap | 1.0568872 | 0.079821405 | 0.35891896 | 1 |
| Eif2s2 | 1.04657969 | 0.065682164 | 0.35897824 | 1 |
| Tmtc2 | 0.93423612 | -0.098140865 | 0.35913684 | 1 |
| Gga3 | 0.94516258 | -0.081365575 | 0.35914909 | 1 |
| Spata5 | 0.90660976 | -0.141446396 | 0.35927498 | 1 |
| Ier5 | 1.12714526 | 0.172673457 | 0.35949474 | 1 |
| Araf | 1.04751327 | 0.066968525 | 0.35954773 | 1 |
| Ipmk | 0.93196368 | -0.101654364 | 0.35977745 | 1 |
| Entpd4 | 1.04317887 | 0.060986546 | 0.3597801 | 1 |
| Tmem229b | 0.95894426 | -0.06048113 | 0.36006619 | 1 |
| Smc6 | 0.9515288 | -0.071680778 | 0.36020785 | 1 |
| Clgn | 0.95621568 | -0.064592028 | 0.36025576 | 1 |
| Col9a1 | 0.79511332 | -0.330767608 | 0.36028098 | 1 |
| Aars | 1.04315671 | 0.060955899 | 0.36029973 | 1 |
| Wash1 | 0.93992466 | -0.089382971 | 0.36034999 | 1 |
| Atg7 | 1.10600428 | 0.145356975 | 0.36036463 | 1 |
| Calm3 | 1.04318175 | 0.060990538 | 0.36038016 | 1 |
| Cpsf6 | 0.95465965 | -0.066941607 | 0.36039252 | 1 |
| Aldh4a1 | 1.07939735 | 0.110226052 | 0.36049215 | 1 |
| Reep1 | 0.95771125 | -0.062337351 | 0.36049399 | 1 |
| Dazap1 | 1.07539716 | 0.104869571 | 0.36050134 | 1 |
| Sh3kbp1 | 1.048849 | 0.068806996 | 0.36072787 | 1 |
| Pus1 | 0.91818381 | -0.123145098 | 0.36075053 | 1 |
| Mypop | 0.92280244 | -0.115906281 | 0.36082999 | 1 |
| Med26 | 0.86505649 | -0.20913375 | 0.36102069 | 1 |
| Rap2a | 1.22143382 | 0.2885757 | 0.3610261 | 1 |
| Tmem127 | 0.95838051 | -0.061329523 | 0.36115923 | 1 |
| Cnpy3 | 0.94295554 | -0.08473834 | 0.36120961 | 1 |
| Fam196a | 0.87237754 | -0.196975465 | 0.36121449 | 1 |
| RGD1310553 | 0.94016009 | -0.089021657 | 0.36124532 | 1 |
| Stoml2 | 1.06063454 | 0.084927641 | 0.36125526 | 1 |
| Osbpl7 | 1.07910519 | 0.109835498 | 0.36125621 | 1 |
| Tm6sf1 | 1.04489198 | 0.063353806 | 0.36128807 | 1 |
| Zswim8 | 0.95040453 | -0.073386376 | 0.36136319 | 1 |
| Galns | 1.1691511 | 0.225461393 | 0.36138967 | 1 |

| | | | | |
|---|---|---|---|---|
| Mars | 1.04866777 | 0.068557685 | 0.36142688 | 1 |
| RGD1311756 | 1.10619634 | 0.145607471 | 0.3614777 | 1 |
| Lage3 | 1.05325415 | 0.074853606 | 0.361517 | 1 |
| Echdc2 | 1.20320884 | 0.266887072 | 0.36164299 | 1 |
| Cyp27a1 | 1.08007602 | 0.11113286 | 0.36169727 | 1 |
| Lsm8 | 1.08581909 | 0.118783759 | 0.36199998 | 1 |
| Pdxp | 0.92526576 | -0.112060294 | 0.36206183 | 1 |
| Tmem161a | 1.09012026 | 0.124487301 | 0.36209009 | 1 |
| Pbxip1 | 1.04841083 | 0.068204164 | 0.36219343 | 1 |
| Zbtb38 | 0.94199551 | -0.086207907 | 0.36243449 | 1 |
| Syn3 | 1.11175829 | 0.152843158 | 0.36247512 | 1 |
| Sema4b | 0.93213614 | -0.101387411 | 0.36259043 | 1 |
| Jmjd1c | 1.04756566 | 0.067040673 | 0.36259963 | 1 |
| Trmt112 | 1.06297206 | 0.088103678 | 0.36280731 | 1 |
| Pomgnt1 | 0.94760077 | -0.077648725 | 0.3629435 | 1 |
| Ezh1 | 1.0442177 | 0.062422524 | 0.3630443 | 1 |
| Itga7 | 0.95918113 | -0.060124825 | 0.36372357 | 1 |
| M6pr | 1.04339283 | 0.061282427 | 0.36373958 | 1 |
| Hoxa7 | 1.14643904 | 0.197159639 | 0.36374439 | 1 |
| Arl8a | 0.95916665 | -0.060146597 | 0.36389475 | 1 |
| Jmjd6 | 0.91163765 | -0.133467583 | 0.36394333 | 1 |
| Dhrs4 | 1.1007961 | 0.13854727 | 0.36394386 | 1 |
| Ckap2 | 0.80733027 | -0.308769107 | 0.36395137 | 1 |
| Slc39a14 | 0.93585986 | -0.095635578 | 0.36397619 | 1 |
| Tnks | 0.944074 | -0.083028142 | 0.36398607 | 1 |
| Zbed4 | 0.91501416 | -0.128134021 | 0.36404266 | 1 |
| Tbrg4 | 0.93654938 | -0.094573026 | 0.36404762 | 1 |
| Mfsd8 | 0.89078774 | -0.166846392 | 0.36405325 | 1 |
| Eif3b | 0.95585486 | -0.06513652 | 0.3642587 | 1 |
| Cenpl | 1.12136197 | 0.165252047 | 0.36427708 | 1 |
| Arhgap30 | 0.87654614 | -0.190098061 | 0.36432036 | 1 |
| Gdap1 | 1.04246523 | 0.059999272 | 0.36444391 | 1 |
| Rnf152 | 1.13051963 | 0.176986043 | 0.36468398 | 1 |
| Mrps33 | 1.06872363 | 0.095888821 | 0.36487826 | 1 |
| Tspan1 | 1.19466907 | 0.256611041 | 0.36493934 | 1 |
| Fn3krp | 1.0756818 | 0.105251373 | 0.36506707 | 1 |
| LOC688765 | 0.86210426 | -0.214065737 | 0.36508745 | 1 |
| Paf1 | 1.06331357 | 0.088567112 | 0.36510995 | 1 |
| Trpc3 | 0.92668282 | -0.109852469 | 0.36524931 | 1 |
| Ccdc174 | 1.0925993 | 0.127764407 | 0.36528426 | 1 |

| | | | | |
|---|---|---|---|---|
| Tmem120b | 0.91476747 | -0.128523025 | 0.36533511 | 1 |
| Jakmip3 | 0.93019745 | -0.104391106 | 0.36534298 | 1 |
| Pagr1 | 1.06119728 | 0.085692877 | 0.36536348 | 1 |
| Mgll | 0.94895017 | -0.075595756 | 0.36540325 | 1 |
| Klhl28 | 0.91568031 | -0.127084093 | 0.36555106 | 1 |
| Fars2 | 1.09128704 | 0.12603062 | 0.36558721 | 1 |
| Pianp | 0.92930115 | -0.105781903 | 0.36568389 | 1 |
| Ik | 1.04436279 | 0.062622961 | 0.36575053 | 1 |
| Znhit1 | 0.93391694 | -0.098633852 | 0.36585736 | 1 |
| Tmem135 | 0.94207505 | -0.086086105 | 0.36601054 | 1 |
| Atp7a | 0.92871486 | -0.106692376 | 0.36622001 | 1 |
| Mettl4 | 0.8237644 | -0.279696308 | 0.36632404 | 1 |
| Stmn2 | 1.04548249 | 0.064168902 | 0.36635621 | 1 |
| Pde12 | 0.89381017 | -0.161959638 | 0.36637187 | 1 |
| Nme6 | 1.17948852 | 0.238161377 | 0.36655668 | 1 |
| Chrdl2 | 0.85045978 | -0.233685082 | 0.36658212 | 1 |
| Kcnf1 | 0.9145989 | -0.128788918 | 0.36659607 | 1 |
| Sf3a1 | 0.94655611 | -0.079240062 | 0.36667307 | 1 |
| Atoh8 | 1.22163997 | 0.288819173 | 0.36667462 | 1 |
| Actr10 | 1.04377327 | 0.061808366 | 0.36675459 | 1 |
| Pstpip1 | 0.87533387 | -0.192094693 | 0.36678456 | 1 |
| Aqp11 | 0.90455053 | -0.144727004 | 0.3668035 | 1 |
| Mx1 | 1.09112249 | 0.125813069 | 0.36684617 | 1 |
| Rhobtb1 | 1.1339036 | 0.181297994 | 0.36685156 | 1 |
| Timm13 | 1.05651575 | 0.079314268 | 0.36691021 | 1 |
| Fap | 1.10223071 | 0.140426226 | 0.36697358 | 1 |
| Cyp2j10 | 0.91448697 | -0.128965474 | 0.3670922 | 1 |
| Chmp6 | 0.93970475 | -0.089720549 | 0.36720302 | 1 |
| Mef2d | 0.92611987 | -0.110729159 | 0.36721726 | 1 |
| St8sia1 | 1.08187176 | 0.113529497 | 0.36725873 | 1 |
| Vps53 | 1.04706091 | 0.066345367 | 0.36739786 | 1 |
| Sulf1 | 1.06780676 | 0.094650582 | 0.36754844 | 1 |
| Ddn | 0.90239571 | -0.148167893 | 0.36761123 | 1 |
| Wwp2 | 0.93340996 | -0.099417231 | 0.36773105 | 1 |
| Ilf3 | 1.06272862 | 0.087773231 | 0.36774938 | 1 |
| Fgfr1 | 0.95795767 | -0.061966187 | 0.36786881 | 1 |
| Mrm1 | 0.88636273 | -0.174030868 | 0.36789883 | 1 |
| Pik3cd | 0.9445361 | -0.08232216 | 0.36792123 | 1 |
| Mepce | 0.93111824 | -0.102963716 | 0.3679942 | 1 |
| RGD1560212 | 1.04501246 | 0.063520145 | 0.36799715 | 1 |

| | | | | |
|---|---|---|---|---|
| Tle1 | 1.0612903 | 0.085819335 | 0.36807695 | 1 |
| Cpz | 1.12551038 | 0.170579367 | 0.36813657 | 1 |
| Max | 0.90846555 | -0.138496288 | 0.36824241 | 1 |
| Scrt1 | 0.94081642 | -0.088014859 | 0.36827571 | 1 |
| Sox12 | 0.88894421 | -0.169835213 | 0.36829719 | 1 |
| Dse | 1.06425293 | 0.089841061 | 0.36837787 | 1 |
| Ubxn6 | 0.94472927 | -0.082027134 | 0.3684737 | 1 |
| Slc17a5 | 1.06860064 | 0.095722782 | 0.36849137 | 1 |
| Ccl6 | 1.12928684 | 0.175411984 | 0.36852025 | 1 |
| Ash1l | 0.95822875 | -0.061557989 | 0.36869089 | 1 |
| Fam46b | 1.23888791 | 0.309045661 | 0.36880821 | 1 |
| Praf2 | 1.04362386 | 0.061601836 | 0.36884954 | 1 |
| Ech1 | 1.04833308 | 0.068097169 | 0.36899422 | 1 |
| Flot1 | 1.04319958 | 0.061015197 | 0.36905294 | 1 |
| LOC306079 | 1.14062012 | 0.189818382 | 0.36914015 | 1 |
| Mafg | 0.93827841 | -0.091912025 | 0.36917018 | 1 |
| Peli3 | 1.11752539 | 0.160307612 | 0.36932108 | 1 |
| Stap2 | 0.85042931 | -0.233736769 | 0.36947222 | 1 |
| Mical3 | 0.94159562 | -0.086820485 | 0.36955213 | 1 |
| Dnase1l1 | 1.08891439 | 0.12289053 | 0.36958625 | 1 |
| Lfng | 0.8965884 | -0.157482263 | 0.36993424 | 1 |
| Pygm | 1.09365353 | 0.129155763 | 0.37006688 | 1 |
| Snw1 | 1.05607637 | 0.078714163 | 0.37009088 | 1 |
| Hdac11 | 0.94494773 | -0.081693567 | 0.37013539 | 1 |
| Tceb3 | 1.06231454 | 0.087210997 | 0.37016667 | 1 |
| C2cd2l | 0.9497472 | -0.074384544 | 0.37038436 | 1 |
| Smn1 | 0.92051008 | -0.119494568 | 0.37041973 | 1 |
| Actr3 | 0.95965824 | -0.059407381 | 0.37063023 | 1 |
| Dcn | 1.04435825 | 0.062616683 | 0.3708452 | 1 |
| Cep70 | 1.10554446 | 0.14475705 | 0.37089085 | 1 |
| Pdzd2 | 0.94364056 | -0.083690663 | 0.37094629 | 1 |
| Fgfbp3 | 1.12894595 | 0.174976421 | 0.37107898 | 1 |
| Fbxl19 | 0.92874527 | -0.106645144 | 0.37113199 | 1 |
| H2afv | 1.09125218 | 0.125984534 | 0.37125028 | 1 |
| Cux2 | 0.89663553 | -0.157406428 | 0.3712898 | 1 |
| Fam120b | 0.95412105 | -0.067755774 | 0.37129247 | 1 |
| Mroh7 | 1.15884695 | 0.212690046 | 0.37144128 | 1 |
| Sf3b2 | 0.95845194 | -0.061221996 | 0.37152404 | 1 |
| Prima1 | 0.9255101 | -0.111679366 | 0.37154898 | 1 |
| Ilvbl | 1.05411667 | 0.076034549 | 0.3715518 | 1 |

| | | | | |
|---|---|---|---|---|
| Ndufa6 | 1.04983683 | 0.070165117 | 0.37158089 | 1 |
| Alkbh5 | 0.92239585 | -0.116542075 | 0.37177226 | 1 |
| Nfat5 | 0.95855076 | -0.061073268 | 0.37194662 | 1 |
| Dnaja3 | 1.05307655 | 0.074610318 | 0.37203041 | 1 |
| Unc5a | 0.85758686 | -0.221645291 | 0.37231945 | 1 |
| Tspan33 | 1.16839764 | 0.224531351 | 0.37235425 | 1 |
| Wdr12 | 1.06640622 | 0.092757099 | 0.3724309 | 1 |
| Plcd1 | 1.07239792 | 0.100840332 | 0.37259152 | 1 |
| Slc22a8 | 1.11319573 | 0.154707278 | 0.37286238 | 1 |
| LOC100909675 | 0.78192055 | -0.354906076 | 0.37291652 | 1 |
| Gng12 | 1.10798978 | 0.147944571 | 0.37302301 | 1 |
| Slc5a6 | 0.95402277 | -0.06790439 | 0.37302569 | 1 |
| Apol3 | 1.13296723 | 0.180106136 | 0.3730403 | 1 |
| Nlk | 0.94756019 | -0.077710511 | 0.37313074 | 1 |
| Suclg2 | 1.05793752 | 0.081254421 | 0.37333772 | 1 |
| Atf7 | 0.92624686 | -0.110531356 | 0.3734463 | 1 |
| Pip4k2c | 1.06896334 | 0.096212373 | 0.37358189 | 1 |
| Tor1b | 0.92927602 | -0.10582091 | 0.373672 | 1 |
| Xab2 | 0.94062305 | -0.088311415 | 0.37373922 | 1 |
| Fam135a | 1.05620274 | 0.078886784 | 0.37380546 | 1 |
| Dnajc25 | 0.87564144 | -0.191587866 | 0.3738311 | 1 |
| Dap3 | 0.93940181 | -0.09018572 | 0.37388831 | 1 |
| Zbtb41 | 0.91800775 | -0.123421766 | 0.37404153 | 1 |
| Rarres1 | 1.0468755 | 0.066089877 | 0.37415371 | 1 |
| Zcchc7 | 0.93501235 | -0.096942667 | 0.37418259 | 1 |
| Card6 | 1.08341663 | 0.115588143 | 0.3742284 | 1 |
| Ssu72 | 1.04304356 | 0.060799405 | 0.37428414 | 1 |
| Gale | 1.0730928 | 0.101774842 | 0.37428575 | 1 |
| Glb1l2 | 1.10780831 | 0.147708267 | 0.37452146 | 1 |
| Rpl24 | 1.04201152 | 0.05937123 | 0.37456997 | 1 |
| Abcc10 | 0.89831163 | -0.154712077 | 0.37467799 | 1 |
| Hsd17b4 | 1.04420442 | 0.06240417 | 0.37470554 | 1 |
| Mtpn | 0.95736689 | -0.062856184 | 0.37487108 | 1 |
| Mrpl2 | 0.92748488 | -0.108604336 | 0.3750177 | 1 |
| Dab1 | 0.8764257 | -0.190296305 | 0.37502353 | 1 |
| Faf2 | 1.05899598 | 0.082697117 | 0.37506838 | 1 |
| Zfp598 | 0.93927645 | -0.090378257 | 0.37509258 | 1 |
| Cyb5a | 1.04765196 | 0.067159524 | 0.37521107 | 1 |
| Prmt7 | 1.07423526 | 0.103309983 | 0.37532887 | 1 |
| Pemt | 1.1671695 | 0.223014091 | 0.37541716 | 1 |

| | | | | |
|---|---|---|---|---|
| Commd2 | 1.07791698 | 0.108246062 | 0.375419 | 1 |
| Nucb2 | 1.05277326 | 0.074194756 | 0.37558353 | 1 |
| RT1-CE2 | 1.10972438 | 0.150201397 | 0.37627919 | 1 |
| Abra | 1.25484598 | 0.327510295 | 0.37629163 | 1 |
| Mipep | 0.92889438 | -0.106413531 | 0.37632154 | 1 |
| Rarb | 1.09843033 | 0.135443366 | 0.37636999 | 1 |
| Man2c1 | 1.06044345 | 0.084667694 | 0.37644202 | 1 |
| Slc2a13 | 0.94100122 | -0.087731501 | 0.37651907 | 1 |
| Tnnc1 | 1.21976756 | 0.286606254 | 0.37653399 | 1 |
| Prdx5 | 1.04262207 | 0.060216308 | 0.37663619 | 1 |
| LOC297756 | 1.04275332 | 0.060397912 | 0.37671275 | 1 |
| Ppm1a | 1.04306867 | 0.060834142 | 0.37680417 | 1 |
| Zfp668 | 0.88407051 | -0.177766653 | 0.37694233 | 1 |
| Scn3a | 1.09289506 | 0.128154879 | 0.37695712 | 1 |
| Lemd2 | 0.94163308 | -0.086763086 | 0.37696236 | 1 |
| Bmx | 1.28349288 | 0.360075288 | 0.37703074 | 1 |
| Prpsap1 | 0.94413892 | -0.08292895 | 0.37707732 | 1 |
| Chd1 | 1.06299621 | 0.088136447 | 0.37717532 | 1 |
| Homer2 | 0.90480125 | -0.144327173 | 0.37721121 | 1 |
| Ushbp1 | 1.15890148 | 0.21275793 | 0.3773263 | 1 |
| Scp2 | 0.96022739 | -0.058552013 | 0.37740767 | 1 |
| Spsb1 | 0.91783195 | -0.123698071 | 0.37749625 | 1 |
| Stmn3 | 1.04256532 | 0.060137776 | 0.37773418 | 1 |
| Phactr1 | 0.92910086 | -0.10609287 | 0.3779125 | 1 |
| Ercc3 | 1.04864474 | 0.068525998 | 0.3781993 | 1 |
| Pdlim7 | 0.94451384 | -0.082356159 | 0.37835997 | 1 |
| RGD1312005 | 1.06327555 | 0.088515529 | 0.37837527 | 1 |
| Dhrs1 | 1.07832028 | 0.108785743 | 0.37840832 | 1 |
| Thoc6 | 1.0950056 | 0.130938251 | 0.37852808 | 1 |
| Cers2 | 0.95624278 | -0.064551149 | 0.3786434 | 1 |
| Asun | 1.07663839 | 0.106533769 | 0.37882728 | 1 |
| Terf2ip | 1.04979805 | 0.070111826 | 0.37890484 | 1 |
| 5-Sep | 0.96035808 | -0.058355669 | 0.37922485 | 1 |
| Mnt | 1.06509139 | 0.09097723 | 0.37928702 | 1 |
| Pi4k2a | 0.94864834 | -0.076054715 | 0.37950926 | 1 |
| Gabarapl2 | 1.04122287 | 0.058278901 | 0.37964941 | 1 |
| Slc6a15 | 1.04115475 | 0.058184518 | 0.37984395 | 1 |
| Sord | 1.04872897 | 0.068641881 | 0.37988166 | 1 |
| Camkk2 | 0.89706033 | -0.156723079 | 0.37991505 | 1 |
| Zcchc11 | 1.08761224 | 0.121164291 | 0.37998203 | 1 |

| | | | | |
|---|---|---|---|---|
| Dpm1 | 1.05577427 | 0.078301406 | 0.38022822 | 1 |
| Zbtb40 | 1.10852999 | 0.1486478 | 0.3802466 | 1 |
| Ccnl1 | 1.05901839 | 0.082727638 | 0.38040515 | 1 |
| Mknk1 | 1.08031445 | 0.111451297 | 0.38040837 | 1 |
| Rad9b | 1.20348014 | 0.267212332 | 0.38049387 | 1 |
| Scly | 0.90294509 | -0.147289832 | 0.38071982 | 1 |
| Calr | 1.04184467 | 0.0591402 | 0.38072791 | 1 |
| Ascc1 | 0.93258225 | -0.100697118 | 0.38108805 | 1 |
| Sp4 | 1.10061444 | 0.138309166 | 0.3810922 | 1 |
| Baiap3 | 1.2897967 | 0.367143679 | 0.38109585 | 1 |
| Atp13a4 | 1.15237459 | 0.204609754 | 0.38113241 | 1 |
| RGD1559786 | 1.06019142 | 0.084324766 | 0.38116245 | 1 |
| Fmo2 | 0.94455979 | -0.082285975 | 0.38118181 | 1 |
| Cry1 | 0.90806493 | -0.139132631 | 0.38150589 | 1 |
| LOC310926 | 1.1670561 | 0.222873916 | 0.38158357 | 1 |
| Papolg | 1.0740614 | 0.103076465 | 0.38158968 | 1 |
| Cxxc4 | 1.07830042 | 0.108759182 | 0.38167994 | 1 |
| Cwc15 | 0.94835842 | -0.07649568 | 0.38171879 | 1 |
| Sft2d1 | 0.93376125 | -0.098874368 | 0.38174194 | 1 |
| Pgp | 0.9504691 | -0.073288364 | 0.38182441 | 1 |
| Syp | 0.96019895 | -0.058594742 | 0.38186027 | 1 |
| Clec1a | 1.11445744 | 0.156341527 | 0.38186389 | 1 |
| Trmt2a | 1.07833819 | 0.108809706 | 0.38190997 | 1 |
| Atp6v0e2 | 1.04304341 | 0.060799198 | 0.38213168 | 1 |
| Sfmbt1 | 1.08065285 | 0.111903151 | 0.38213648 | 1 |
| Spryd7 | 1.05482486 | 0.077003471 | 0.38216381 | 1 |
| Wasl | 0.95762655 | -0.06246495 | 0.38230598 | 1 |
| Zdhhc2 | 1.04114563 | 0.058171874 | 0.38245887 | 1 |
| Nmt1 | 1.04540059 | 0.064055878 | 0.38251767 | 1 |
| Ankrd40 | 0.96046356 | -0.058197223 | 0.38264559 | 1 |
| RGD1563986 | 1.12994718 | 0.176255337 | 0.38272137 | 1 |
| Ammecr1l | 0.94290007 | -0.084823214 | 0.38283293 | 1 |
| Irs1 | 1.13648678 | 0.184580904 | 0.3828441 | 1 |
| Pgam2 | 0.75623704 | -0.403089585 | 0.38285283 | 1 |
| Ap3m1 | 1.04611656 | 0.065043614 | 0.38297328 | 1 |
| Uba3 | 1.04243307 | 0.059954755 | 0.38297682 | 1 |
| Ttll3 | 0.83859316 | -0.253957035 | 0.38307517 | 1 |
| Cd69 | 0.78806906 | -0.343606032 | 0.38312493 | 1 |
| Kcnh8 | 1.15293564 | 0.205311978 | 0.38321703 | 1 |
| Rqcd1 | 1.06698553 | 0.093540615 | 0.38334181 | 1 |

| | | | | |
|---|---|---|---|---|
| Hira | 0.93696401 | -0.093934461 | 0.3833605 | 1 |
| Tgfbr1 | 1.05927767 | 0.083080819 | 0.38348907 | 1 |
| Atp1b2 | 0.95706239 | -0.063315126 | 0.3835886 | 1 |
| Edem1 | 0.95434558 | -0.067416319 | 0.38369043 | 1 |
| mrpl24 | 1.06155357 | 0.086177183 | 0.38389824 | 1 |
| Gltscr2 | 1.05041175 | 0.070954955 | 0.38391382 | 1 |
| Zdhhc15 | 0.942618 | -0.085254858 | 0.38400783 | 1 |
| Rnpepl1 | 0.93095188 | -0.103221492 | 0.38423727 | 1 |
| Dlg3 | 0.88825325 | -0.170957029 | 0.38441938 | 1 |
| Srgn | 1.07842982 | 0.10893229 | 0.38444077 | 1 |
| Wasf2 | 1.09642396 | 0.132805761 | 0.3847504 | 1 |
| Dek | 1.05667162 | 0.079527103 | 0.38480956 | 1 |
| Wdr81 | 0.93387292 | -0.098701854 | 0.38495016 | 1 |
| Bckdhb | 1.06887234 | 0.09608955 | 0.38498444 | 1 |
| Cbx2 | 0.87949393 | -0.185254477 | 0.38505645 | 1 |
| Shank2 | 0.93830891 | -0.091865134 | 0.38516155 | 1 |
| Nedd9 | 1.06918398 | 0.096510131 | 0.38517867 | 1 |
| Nde1 | 1.09965722 | 0.137053888 | 0.38543762 | 1 |
| Foxs1 | 1.12208301 | 0.166179411 | 0.38560357 | 1 |
| Tmsbl1 | 1.06599335 | 0.092198443 | 0.3856362 | 1 |
| Sntb1 | 0.92882215 | -0.106525723 | 0.38570148 | 1 |
| Plekhm2 | 0.94972097 | -0.07442438 | 0.38575761 | 1 |
| Dpp4 | 0.93504674 | -0.096889605 | 0.38576944 | 1 |
| Fam173a | 0.91880981 | -0.122161832 | 0.38586001 | 1 |
| Fgf1 | 1.04136795 | 0.058479911 | 0.38586244 | 1 |
| Ccdc28a | 1.05494662 | 0.077169994 | 0.38601518 | 1 |
| Kif20a | 1.20524237 | 0.269323295 | 0.38604126 | 1 |
| Nup210l | 1.29652741 | 0.374652704 | 0.3860427 | 1 |
| Eif3f | 0.95518996 | -0.066140426 | 0.38609392 | 1 |
| Mapk8ip1 | 0.96046709 | -0.058191921 | 0.38630885 | 1 |
| Elp5 | 1.0776953 | 0.107949343 | 0.38636764 | 1 |
| Carm1 | 0.94850434 | -0.076273721 | 0.38655468 | 1 |
| Flrt2 | 1.11074103 | 0.15152249 | 0.38674353 | 1 |
| Cenpq | 1.168504 | 0.224662676 | 0.38692535 | 1 |
| Bcas1 | 1.05600849 | 0.078621428 | 0.38717186 | 1 |
| Mov10 | 1.13582316 | 0.183738239 | 0.38731103 | 1 |
| Cul5 | 0.94523211 | -0.081259459 | 0.38741065 | 1 |
| Ociad2 | 1.12000346 | 0.163503188 | 0.38748886 | 1 |
| Tubgcp3 | 1.05795409 | 0.081277026 | 0.38751617 | 1 |
| Rbfa | 0.92493587 | -0.112574755 | 0.38752799 | 1 |

| | | | | |
|---|---|---|---|---|
| Nsmce1 | 0.92494643 | -0.11255829 | 0.38771591 | 1 |
| Pla2g12a | 0.91993488 | -0.120396349 | 0.38780601 | 1 |
| LOC294154 | 0.96048556 | -0.058164171 | 0.38787608 | 1 |
| Ccnf | 0.85164474 | -0.231676349 | 0.38787933 | 1 |
| Rhoc | 0.95000772 | -0.073988856 | 0.38794591 | 1 |
| Card19 | 0.93741479 | -0.093240539 | 0.38795734 | 1 |
| Id3 | 1.04948577 | 0.069682604 | 0.38804537 | 1 |
| Prkcdbp | 0.95192088 | -0.071086421 | 0.38816089 | 1 |
| Ptprm | 1.0518008 | 0.072861501 | 0.38819392 | 1 |
| Uqcr11 | 1.04498352 | 0.06348019 | 0.38822633 | 1 |
| Xylb | 0.93832329 | -0.091843021 | 0.38842018 | 1 |
| Cyp4f18 | 1.39526372 | 0.480537834 | 0.38849396 | 1 |
| Mrgprf | 0.9156189 | -0.127180844 | 0.38854408 | 1 |
| Foxm1 | 1.17775344 | 0.236037541 | 0.38887363 | 1 |
| Hgh1 | 0.92759345 | -0.108435457 | 0.38899494 | 1 |
| Nat9 | 0.92802245 | -0.107768387 | 0.38910395 | 1 |
| RGD1304694 | 0.95286394 | -0.069657869 | 0.38936132 | 1 |
| Rpp21 | 0.89983871 | -0.152261658 | 0.38937922 | 1 |
| Prkd3 | 1.05884436 | 0.082490543 | 0.38942214 | 1 |
| Map4k2 | 1.06013669 | 0.084250296 | 0.3894263 | 1 |
| Hspa12b | 1.11121571 | 0.152138905 | 0.38952072 | 1 |
| Triqk | 1.06118899 | 0.085681608 | 0.38966093 | 1 |
| Gba | 1.04199252 | 0.059344925 | 0.3896766 | 1 |
| Mitf | 1.09008976 | 0.124446928 | 0.39001863 | 1 |
| Hsf2 | 1.05236793 | 0.073639194 | 0.39004535 | 1 |
| Camk4 | 0.91572522 | -0.127013337 | 0.39008915 | 1 |
| Hey1 | 0.88286231 | -0.179739646 | 0.39016737 | 1 |
| Zfp777 | 0.92802475 | -0.107764811 | 0.39022916 | 1 |
| Npl | 0.92078443 | -0.119064663 | 0.39028301 | 1 |
| Myo9a | 0.95978989 | -0.059209483 | 0.39029497 | 1 |
| Alg3 | 1.08095489 | 0.112306323 | 0.39052932 | 1 |
| Ccdc115 | 1.08178359 | 0.113411921 | 0.39053302 | 1 |
| Usp12 | 0.94097732 | -0.087768142 | 0.39053925 | 1 |
| Btbd6 | 0.95494029 | -0.066517568 | 0.39057512 | 1 |
| Sdk2 | 0.89310944 | -0.163091122 | 0.39071379 | 1 |
| Sv2b | 1.04075814 | 0.057634836 | 0.39071939 | 1 |
| Usp36 | 0.92762561 | -0.108385443 | 0.39076734 | 1 |
| Zfp580 | 1.11198708 | 0.153140028 | 0.39078055 | 1 |
| Plp2 | 0.93778343 | -0.092673307 | 0.39083663 | 1 |
| Cmtm3 | 1.1067838 | 0.146373432 | 0.39092114 | 1 |

| | | | | |
|---|---|---|---|---|
| Ado | 0.88571903 | -0.175078984 | 0.39111264 | 1 |
| Trem2 | 1.14788448 | 0.198977466 | 0.39114625 | 1 |
| Usf2 | 0.94488679 | -0.081786611 | 0.39118742 | 1 |
| Sf3b6 | 1.07619172 | 0.105935115 | 0.39136251 | 1 |
| Ppfibp2 | 1.06666382 | 0.093105555 | 0.39153803 | 1 |
| RGD1307554 | 0.9119166 | -0.133026211 | 0.39155686 | 1 |
| Pcdhb9 | 1.1146792 | 0.156628568 | 0.39166345 | 1 |
| Nab2 | 0.91176142 | -0.133271734 | 0.39176295 | 1 |
| Plaa | 0.95514173 | -0.066213269 | 0.39189322 | 1 |
| Sstr2 | 0.81270999 | -0.29918747 | 0.39190619 | 1 |
| Csf2ra | 1.14424497 | 0.194395953 | 0.39196003 | 1 |
| Fmo1 | 0.94647151 | -0.079369018 | 0.39196422 | 1 |
| Ifngr2 | 1.06441091 | 0.090055209 | 0.39207615 | 1 |
| Slc39a9 | 0.91211547 | -0.132711613 | 0.39210644 | 1 |
| Pex10 | 1.13031571 | 0.176725784 | 0.39218195 | 1 |
| Snapc2 | 0.94423166 | -0.082787236 | 0.39226204 | 1 |
| Lrrtm3 | 1.12720611 | 0.172751341 | 0.39229791 | 1 |
| Atp6v1g2 | 1.04025611 | 0.056938758 | 0.39245832 | 1 |
| Myo1c | 0.95220453 | -0.070656604 | 0.39252652 | 1 |
| Iba57 | 1.12234547 | 0.166516821 | 0.39260806 | 1 |
| Ghitm | 1.04011724 | 0.056746149 | 0.39260834 | 1 |
| Mmp16 | 0.90505631 | -0.143920538 | 0.39277961 | 1 |
| Gcdh | 1.05499654 | 0.077238265 | 0.39281353 | 1 |
| Atp5l | 1.04035244 | 0.057072356 | 0.39282661 | 1 |
| Lmo4 | 0.95109356 | -0.072340822 | 0.39283479 | 1 |
| Zfp521 | 1.05882038 | 0.082457871 | 0.3928865 | 1 |
| RGD1307315 | 0.88921944 | -0.169388604 | 0.39301731 | 1 |
| Sesn1 | 1.05426731 | 0.076240715 | 0.39317711 | 1 |
| Naaa | 1.07472313 | 0.103965042 | 0.39327725 | 1 |
| Pop4 | 1.06331676 | 0.088571433 | 0.39329784 | 1 |
| Tmem88b | 0.84818227 | -0.237553769 | 0.39335287 | 1 |
| LOC100910973 | 1.06951526 | 0.09695707 | 0.39336513 | 1 |
| Ddr1 | 0.95141443 | -0.071854184 | 0.39358563 | 1 |
| Prr16 | 0.83158768 | -0.266059706 | 0.39368474 | 1 |
| Rida | 1.06751354 | 0.094254365 | 0.39384419 | 1 |
| Lbhd1 | 1.08635606 | 0.119497035 | 0.3938967 | 1 |
| Nrg1 | 1.04201219 | 0.059372158 | 0.39391984 | 1 |
| Tpr | 1.04058487 | 0.057394633 | 0.3940253 | 1 |
| Clec7a | 1.13814964 | 0.18669025 | 0.39415082 | 1 |
| A2m | 0.95547447 | -0.065710765 | 0.39415672 | 1 |

| | | | | |
|---|---|---|---|---|
| Gls2 | 0.94616223 | -0.079840524 | 0.39453109 | 1 |
| N4bp2l2 | 1.06244183 | 0.087383856 | 0.39468029 | 1 |
| Zfp428 | 1.09715572 | 0.133768309 | 0.39476144 | 1 |
| Rab10 | 0.96178025 | -0.056220786 | 0.39491052 | 1 |
| Diexf | 1.06608495 | 0.092322405 | 0.39493711 | 1 |
| Usp21 | 0.92199897 | -0.117162962 | 0.39495289 | 1 |
| Wsb1 | 1.04648359 | 0.065549687 | 0.39495615 | 1 |
| Smcr8 | 0.93510488 | -0.096799904 | 0.39503843 | 1 |
| Tex30 | 1.11105282 | 0.151927399 | 0.39506615 | 1 |
| Nacad | 0.96181459 | -0.056169278 | 0.39539581 | 1 |
| Rangrf | 0.89112048 | -0.1663076 | 0.39543738 | 1 |
| Skiv2l | 0.94458642 | -0.082245302 | 0.3954876 | 1 |
| Clec4a | 1.2677886 | 0.342314195 | 0.39549705 | 1 |
| Cst6 | 1.12461014 | 0.169424965 | 0.39556462 | 1 |
| Sema3d | 1.05606205 | 0.078694603 | 0.39557165 | 1 |
| C1qtnf6 | 0.87360076 | -0.194953977 | 0.39573331 | 1 |
| Slc8a1 | 1.06516417 | 0.09107581 | 0.3957868 | 1 |
| Zbtb39 | 0.91814138 | -0.123211772 | 0.39582441 | 1 |
| Polr3k | 1.07511834 | 0.104495473 | 0.39584623 | 1 |
| Fbxo31 | 0.95367978 | -0.06842317 | 0.39586252 | 1 |
| Srp19 | 1.0548917 | 0.077094896 | 0.39590152 | 1 |
| Cldn19 | 0.95257484 | -0.070095646 | 0.39607639 | 1 |
| Fbxo21 | 1.04575706 | 0.064547745 | 0.39614781 | 1 |
| Fez2 | 1.05803329 | 0.081385016 | 0.39624019 | 1 |
| Tomm40l | 1.08145581 | 0.112974721 | 0.39634464 | 1 |
| Trub1 | 0.8960174 | -0.158401349 | 0.39641739 | 1 |
| Gstk1 | 1.06729518 | 0.093959236 | 0.39646489 | 1 |
| Rpusd4 | 1.08582258 | 0.118788391 | 0.39648947 | 1 |
| Sncg | 1.04217167 | 0.059592944 | 0.39649292 | 1 |
| Gcc2 | 1.04389051 | 0.061970406 | 0.39656628 | 1 |
| Mcat | 0.91351092 | -0.130506124 | 0.39673757 | 1 |
| Zic2 | 1.07910995 | 0.109841872 | 0.396793 | 1 |
| Taf1 | 0.9605396 | -0.058083006 | 0.39687236 | 1 |
| Etaa1 | 1.07427015 | 0.103356842 | 0.39691469 | 1 |
| Parp2 | 1.08538612 | 0.118208368 | 0.39701573 | 1 |
| Sec13 | 1.05365846 | 0.075407296 | 0.39702164 | 1 |
| Kif5a | 0.96055496 | -0.058059934 | 0.39706996 | 1 |
| Sp110 | 1.0801665 | 0.111253705 | 0.3972094 | 1 |
| Mettl2b | 0.90060374 | -0.151035625 | 0.39721733 | 1 |
| Clvs2 | 1.11978288 | 0.163219033 | 0.3973814 | 1 |

| | | | | |
|---|---|---|---|---|
| Shq1 | 0.84164497 | -0.248716296 | 0.39760681 | 1 |
| Birc6 | 0.96194006 | -0.05598109 | 0.39761537 | 1 |
| Zfp622 | 1.06181777 | 0.086536197 | 0.39787767 | 1 |
| Zfand4 | 1.12744431 | 0.173056177 | 0.39789769 | 1 |
| Kank3 | 0.90973493 | -0.136481844 | 0.39790042 | 1 |
| Adk | 1.04115362 | 0.058182953 | 0.39799756 | 1 |
| Zfyve21 | 0.89731393 | -0.15631529 | 0.39800929 | 1 |
| A4galt | 0.93955963 | -0.089943367 | 0.39808749 | 1 |
| Depdc7 | 1.13261607 | 0.179658899 | 0.39814607 | 1 |
| Acvrl1 | 1.08202812 | 0.113737992 | 0.3982217 | 1 |
| Elmsan1 | 1.1068972 | 0.146521246 | 0.39824503 | 1 |
| Polr1d | 0.93771293 | -0.092781773 | 0.39832207 | 1 |
| Efnb2 | 1.09506392 | 0.131015083 | 0.39834659 | 1 |
| Szrd1 | 0.94712309 | -0.078376156 | 0.39837527 | 1 |
| Vps16 | 1.05399584 | 0.075869175 | 0.39858126 | 1 |
| Enpp1 | 0.92517964 | -0.112194583 | 0.39867467 | 1 |
| Fjx1 | 0.84918601 | -0.23584749 | 0.39870531 | 1 |
| Srpx | 1.07057338 | 0.098383683 | 0.39875719 | 1 |
| Tysnd1 | 0.9154553 | -0.127438649 | 0.39882942 | 1 |
| Poldip3 | 1.04626879 | 0.065253533 | 0.39884441 | 1 |
| Brca1 | 1.12539327 | 0.170429241 | 0.39887627 | 1 |
| Grhpr | 1.11626665 | 0.158681697 | 0.39898883 | 1 |
| Tbrg1 | 1.06010586 | 0.084208336 | 0.39910847 | 1 |
| Aatf | 1.0802277 | 0.111335447 | 0.39913848 | 1 |
| Kif13a | 0.94686835 | -0.078764246 | 0.3991965 | 1 |
| Fgd2 | 0.87212445 | -0.197394069 | 0.3992966 | 1 |
| Taf3 | 0.92362229 | -0.114625104 | 0.39932528 | 1 |
| Mad2l2 | 0.89193183 | -0.164994639 | 0.39945788 | 1 |
| RGD1566320 | 1.07479215 | 0.104057691 | 0.39953802 | 1 |
| Arsk | 1.09315222 | 0.128494304 | 0.39967401 | 1 |
| Serhl2 | 0.8419419 | -0.248207417 | 0.39976075 | 1 |
| Uso1 | 0.96132965 | -0.056896857 | 0.39981189 | 1 |
| Ppp2r5a | 0.93850241 | -0.09156765 | 0.39991576 | 1 |
| Limk1 | 0.96110373 | -0.05723595 | 0.40002237 | 1 |
| Ecel1 | 0.79543633 | -0.33018163 | 0.40003812 | 1 |
| Asl | 0.93639287 | -0.094814145 | 0.40009002 | 1 |
| Nell2 | 1.06087632 | 0.085256467 | 0.40009744 | 1 |
| Spg11 | 1.06058593 | 0.084861514 | 0.40012088 | 1 |
| Ankrd29 | 1.0489577 | 0.068956507 | 0.40044286 | 1 |
| Pex1 | 1.06201211 | 0.086800214 | 0.4004901 | 1 |

| | | | | |
|---|---|---|---|---|
| Lzts1 | 0.94835119 | -0.076506686 | 0.40051212 | 1 |
| Bcdin3d | 0.87623573 | -0.190609047 | 0.40064967 | 1 |
| Fyco1 | 0.94193626 | -0.086298654 | 0.40070905 | 1 |
| Fli1 | 1.10467162 | 0.14361757 | 0.40079887 | 1 |
| Agfg2 | 0.9347211 | -0.097392128 | 0.40081012 | 1 |
| Arid4b | 1.05218799 | 0.073392492 | 0.4008251 | 1 |
| Kif1b | 0.96059732 | -0.057996313 | 0.40084773 | 1 |
| Grpel2 | 1.06997089 | 0.097571541 | 0.40085245 | 1 |
| Tgs1 | 1.07228581 | 0.100689497 | 0.40100798 | 1 |
| Lysmd1 | 1.07984772 | 0.110827884 | 0.40103266 | 1 |
| Alyref | 0.93487294 | -0.097157801 | 0.40106969 | 1 |
| Atg14 | 1.07639812 | 0.10621178 | 0.40108045 | 1 |
| Grid1 | 0.90993131 | -0.136170451 | 0.40111006 | 1 |
| Grem1 | 1.13359047 | 0.180899536 | 0.40115744 | 1 |
| Rev1 | 1.06368051 | 0.089064888 | 0.4013769 | 1 |
| Msln | 0.86291151 | -0.212715476 | 0.40152703 | 1 |
| Naprt | 1.10726057 | 0.146994774 | 0.40153376 | 1 |
| Dhx35 | 1.07690304 | 0.106888357 | 0.40158176 | 1 |
| Pvalb | 1.04087709 | 0.057799715 | 0.40166762 | 1 |
| Tnni3k | 1.22180354 | 0.28901233 | 0.40174178 | 1 |
| Slc35a4 | 1.04743448 | 0.066860005 | 0.40178182 | 1 |
| Kdm3a | 1.05345488 | 0.075128519 | 0.40219386 | 1 |
| Prkag2 | 0.95887491 | -0.060585468 | 0.40235519 | 1 |
| Slc2a9 | 1.13003648 | 0.176369345 | 0.40256796 | 1 |
| Mpp3 | 0.88769204 | -0.171868842 | 0.40258038 | 1 |
| Brsk1 | 0.96176733 | -0.056240168 | 0.40265122 | 1 |
| Foxc1 | 1.07326559 | 0.102007124 | 0.4026541 | 1 |
| Mkrn2 | 1.05391556 | 0.075759288 | 0.4026995 | 1 |
| Ifi30 | 0.9286654 | -0.106769214 | 0.40275278 | 1 |
| Ypel2 | 1.06997186 | 0.097572849 | 0.40289079 | 1 |
| Kcne4 | 0.84572725 | -0.241735633 | 0.40309822 | 1 |
| Csnk2a1 | 1.04278937 | 0.060447781 | 0.40310672 | 1 |
| Mgat2 | 1.05804923 | 0.081406751 | 0.40325733 | 1 |
| Bdh2 | 0.86784302 | -0.204493994 | 0.40326906 | 1 |
| Rfc2 | 1.10207848 | 0.140226965 | 0.40328891 | 1 |
| Lrrc17 | 1.12688939 | 0.172345909 | 0.40332873 | 1 |
| Hhatl | 1.05518122 | 0.077490797 | 0.40339706 | 1 |
| Gdf11 | 0.94384812 | -0.083373374 | 0.40339865 | 1 |
| Smyd2 | 1.04372661 | 0.061743864 | 0.4034632 | 1 |
| Slc40a1 | 1.07894377 | 0.10961968 | 0.40354256 | 1 |

| | | | | |
|---|---|---|---|---|
| Ddx23 | 0.95703459 | -0.063357031 | 0.40359352 | 1 |
| Wdr34 | 0.89514731 | -0.159802976 | 0.40379933 | 1 |
| Prkcq | 0.93175348 | -0.101979798 | 0.40380219 | 1 |
| Rangap1 | 0.96113702 | -0.057185975 | 0.40381259 | 1 |
| Mrpl1 | 1.05981742 | 0.083815743 | 0.40386211 | 1 |
| Adpgk | 1.07110887 | 0.09910512 | 0.40429858 | 1 |
| Clic4 | 0.94039954 | -0.088654263 | 0.40442372 | 1 |
| Bccip | 1.06777495 | 0.094607611 | 0.4046133 | 1 |
| Sema4a | 1.10437753 | 0.143233441 | 0.40464194 | 1 |
| Rara | 0.90102054 | -0.150368098 | 0.40472021 | 1 |
| Amt | 0.92490736 | -0.112619221 | 0.40472746 | 1 |
| Vav2 | 0.95137058 | -0.071920676 | 0.40473179 | 1 |
| Cmpk2 | 1.06396497 | 0.089450651 | 0.40475128 | 1 |
| Zfp260 | 1.0537803 | 0.075574109 | 0.4047823 | 1 |
| Gnl3l | 0.95878956 | -0.060713895 | 0.40484156 | 1 |
| Supv3l1 | 1.06197835 | 0.086754358 | 0.40493495 | 1 |
| Dnajb11 | 0.94862421 | -0.076091406 | 0.40518337 | 1 |
| Ydjc | 1.12791676 | 0.173660603 | 0.40527525 | 1 |
| Epha4 | 0.91983069 | -0.120559758 | 0.40530781 | 1 |
| Tmem54 | 1.07825423 | 0.108697374 | 0.40539069 | 1 |
| Stk38 | 0.94078617 | -0.088061243 | 0.40550362 | 1 |
| Cd99l2 | 1.03885889 | 0.054999702 | 0.40556755 | 1 |
| Bid | 1.06682964 | 0.093329814 | 0.40562493 | 1 |
| Slc6a2 | 1.25375513 | 0.326255609 | 0.40576605 | 1 |
| Gabarapl1 | 1.04045423 | 0.057213508 | 0.40590802 | 1 |
| Gpatch2l | 1.08670039 | 0.119954233 | 0.40605728 | 1 |
| Sfr1 | 1.04335007 | 0.061223301 | 0.40625817 | 1 |
| Smad7 | 0.89370228 | -0.162133797 | 0.40628958 | 1 |
| Atxn1 | 1.10910194 | 0.149391978 | 0.40630217 | 1 |
| Cpsf2 | 0.95550627 | -0.065662756 | 0.40634109 | 1 |
| Pir | 1.06610769 | 0.09235318 | 0.40637452 | 1 |
| Mrpl19 | 1.06866707 | 0.095812464 | 0.40640468 | 1 |
| Camk2a | 0.9344755 | -0.097771258 | 0.40642362 | 1 |
| Hsf4 | 0.94003 | -0.089221296 | 0.40689629 | 1 |
| Eif2a | 1.05152761 | 0.072486729 | 0.40703027 | 1 |
| Pbx3 | 0.93828076 | -0.091908408 | 0.40713064 | 1 |
| Hexa | 0.956149 | -0.064692639 | 0.40717317 | 1 |
| Rcc1 | 0.87385843 | -0.194528528 | 0.40718532 | 1 |
| Nans | 1.04554128 | 0.064250029 | 0.40728058 | 1 |
| Abcb6 | 1.05776978 | 0.081025669 | 0.40737384 | 1 |

| | | | | |
|---|---|---|---|---|
| Mta1 | 0.94499145 | -0.081626823 | 0.40742106 | 1 |
| Gipc1 | 0.95557865 | -0.065553471 | 0.40772277 | 1 |
| Ppfia3 | 0.95345537 | -0.068762681 | 0.40778815 | 1 |
| Tnr | 1.08283605 | 0.114814818 | 0.40786847 | 1 |
| Sh3rf2 | 1.33028661 | 0.411737102 | 0.4078993 | 1 |
| Rfng | 1.06885768 | 0.096069762 | 0.40798291 | 1 |
| Sp2 | 1.11059636 | 0.151334576 | 0.40807177 | 1 |
| Nudt22 | 0.91152208 | -0.133650485 | 0.40815725 | 1 |
| Polr1e | 1.09709327 | 0.133686176 | 0.40818293 | 1 |
| Dhfr | 1.10915757 | 0.149464331 | 0.40819571 | 1 |
| Rev3l | 0.94744102 | -0.077891959 | 0.40824194 | 1 |
| Cyb5r4 | 1.06090013 | 0.085288858 | 0.40825786 | 1 |
| Spint2 | 1.07001683 | 0.097633494 | 0.40828738 | 1 |
| Abi2 | 0.95950034 | -0.059644774 | 0.40860994 | 1 |
| Fhl2 | 1.10778282 | 0.147675074 | 0.40868259 | 1 |
| Med24 | 1.04950074 | 0.069703185 | 0.40869682 | 1 |
| Calcb | 0.96055376 | -0.058061738 | 0.40886564 | 1 |
| Sardh | 0.91244298 | -0.132193692 | 0.40898284 | 1 |
| Pdcl | 0.94522413 | -0.08127164 | 0.4089868 | 1 |
| Smim11 | 0.88862218 | -0.170357938 | 0.40900774 | 1 |
| Yars2 | 1.11770954 | 0.160545321 | 0.40912454 | 1 |
| Swap70 | 0.95309338 | -0.069310529 | 0.40916212 | 1 |
| Rrm2 | 1.12292134 | 0.167256871 | 0.40939369 | 1 |
| Lrrc61 | 1.0648535 | 0.09065496 | 0.40949773 | 1 |
| Fbxl21 | 1.20900036 | 0.27381467 | 0.40952435 | 1 |
| Metrn | 0.94585486 | -0.080309273 | 0.40953553 | 1 |
| Mfap2 | 0.89481111 | -0.16034493 | 0.40954847 | 1 |
| Plppr4 | 0.92863189 | -0.106821274 | 0.40965332 | 1 |
| Rgs2 | 1.0838269 | 0.116134354 | 0.40969041 | 1 |
| Srfbp1 | 0.92350481 | -0.114808616 | 0.40972626 | 1 |
| Serinc3 | 1.03926948 | 0.055569783 | 0.40976909 | 1 |
| Lpgat1 | 0.95360027 | -0.068543456 | 0.40988306 | 1 |
| Galnt11 | 1.04820246 | 0.067917403 | 0.40994741 | 1 |
| Tmeff2 | 1.05178977 | 0.072846373 | 0.41016437 | 1 |
| Vash2 | 1.18765478 | 0.248115543 | 0.41023039 | 1 |
| Tbxa2r | 1.18391484 | 0.243565313 | 0.4104185 | 1 |
| Fam134a | 0.95791729 | -0.062027006 | 0.41046678 | 1 |
| Dnajc7 | 1.03965283 | 0.056101845 | 0.41052954 | 1 |
| Limd2 | 0.93159693 | -0.102222211 | 0.41060727 | 1 |
| Ap5m1 | 0.93175514 | -0.101977229 | 0.41068044 | 1 |

| | | | | |
|---|---|---|---|---|
| N4bp2l1 | 1.15918422 | 0.213109867 | 0.41088775 | 1 |
| Lsp1 | 1.07816273 | 0.108574941 | 0.41089807 | 1 |
| Mbtps1 | 1.04130503 | 0.058392741 | 0.41090435 | 1 |
| Coa7 | 0.89826894 | -0.154780647 | 0.41098309 | 1 |
| Kifc2 | 1.05578588 | 0.078317275 | 0.41119099 | 1 |
| Efs | 1.0942611 | 0.129957017 | 0.41125291 | 1 |
| Dyrk2 | 0.92782701 | -0.108072244 | 0.41138142 | 1 |
| Igfbp4 | 1.04204493 | 0.05941749 | 0.41145965 | 1 |
| Cep78 | 0.89996719 | -0.152055682 | 0.41146973 | 1 |
| Lrp3 | 0.9540897 | -0.067803185 | 0.41172752 | 1 |
| Fig4 | 1.05211204 | 0.073288347 | 0.41186451 | 1 |
| Chmp2a | 0.95290503 | -0.069595664 | 0.41190435 | 1 |
| Osbp | 0.95776941 | -0.062249739 | 0.41192856 | 1 |
| Maf1 | 1.04838102 | 0.06816314 | 0.41198336 | 1 |
| Alg13 | 1.10193832 | 0.140043478 | 0.41201604 | 1 |
| Nos3 | 0.89269411 | -0.16376219 | 0.41206371 | 1 |
| Pcsk7 | 0.94358024 | -0.083782883 | 0.41208159 | 1 |
| Osbp2 | 1.05993797 | 0.083979836 | 0.41209669 | 1 |
| Mtif2 | 1.07096759 | 0.098914817 | 0.41214599 | 1 |
| Eif4a3 | 0.95343648 | -0.068791265 | 0.4121783 | 1 |
| Sars2 | 0.90496699 | -0.14406293 | 0.41218048 | 1 |
| Usb1 | 1.09197107 | 0.126934632 | 0.41234623 | 1 |
| Hey2 | 0.94532436 | -0.081118668 | 0.41243683 | 1 |
| Cipc | 0.9568679 | -0.06360833 | 0.41245148 | 1 |
| Asb6 | 1.06742359 | 0.094132798 | 0.41256718 | 1 |
| Dpf3 | 1.22953316 | 0.298110644 | 0.41258921 | 1 |
| Gfap | 0.93186118 | -0.101813047 | 0.41260346 | 1 |
| Nrxn1 | 0.96139157 | -0.056803936 | 0.41265787 | 1 |
| Atg16l1 | 0.95360948 | -0.068529516 | 0.41273637 | 1 |
| Dhx57 | 0.96084327 | -0.057626967 | 0.41332062 | 1 |
| Ncl | 1.03851338 | 0.054519798 | 0.41342481 | 1 |
| Slc35a5 | 1.05698318 | 0.079952417 | 0.41355207 | 1 |
| Bnip2 | 1.0690956 | 0.096390861 | 0.41358318 | 1 |
| Rsbn1l | 1.07183713 | 0.100085695 | 0.41363756 | 1 |
| Ttc39a | 0.92123374 | -0.118360841 | 0.41367482 | 1 |
| Snx18 | 1.04858285 | 0.068440856 | 0.41367797 | 1 |
| Pard6g | 0.89385869 | -0.16188132 | 0.41373914 | 1 |
| Paqr5 | 1.07171522 | 0.099921593 | 0.41383036 | 1 |
| Krt75 | 1.18503838 | 0.244933781 | 0.4138874 | 1 |
| Nck1 | 1.05602417 | 0.078642849 | 0.41392728 | 1 |

| | | | | |
|---|---|---|---|---|
| Brms1 | 0.95116476 | -0.07223283 | 0.41402608 | 1 |
| Insrr | 1.08124155 | 0.112688862 | 0.41406946 | 1 |
| Zfyve9 | 0.95891579 | -0.060523975 | 0.41411233 | 1 |
| Begain | 1.14047203 | 0.189631064 | 0.41419946 | 1 |
| Plxdc1 | 0.88734172 | -0.172438293 | 0.41430143 | 1 |
| Fam98c | 0.91211827 | -0.132707188 | 0.41430344 | 1 |
| Rps19 | 1.0468613 | 0.066070308 | 0.41463777 | 1 |
| Gnaq | 0.95202249 | -0.070932439 | 0.41468488 | 1 |
| LOC102550394 | 0.848582 | -0.236874018 | 0.41472536 | 1 |
| Sp100 | 1.11802026 | 0.160946337 | 0.41473652 | 1 |
| Ttc5 | 1.05498235 | 0.077218864 | 0.41496092 | 1 |
| Pdap1 | 0.9591117 | -0.060229257 | 0.41497087 | 1 |
| Cdc73 | 0.91378024 | -0.13008085 | 0.41506835 | 1 |
| Osbpl11 | 0.94053919 | -0.088440037 | 0.41507151 | 1 |
| Pcdhga12 | 0.87240481 | -0.196930363 | 0.41512128 | 1 |
| Fbxl20 | 0.91057926 | -0.135143502 | 0.41515721 | 1 |
| Zfp512b | 1.05816563 | 0.081565462 | 0.41528949 | 1 |
| Vangl1 | 0.90317054 | -0.146929671 | 0.41541251 | 1 |
| Rad18 | 1.09797511 | 0.134845347 | 0.41550659 | 1 |
| Plekhh2 | 0.90269985 | -0.147681733 | 0.41557285 | 1 |
| Aebp2 | 0.90227191 | -0.148365817 | 0.41558736 | 1 |
| Kifc3 | 0.92622212 | -0.110569889 | 0.41560083 | 1 |
| Drd2 | 0.93479961 | -0.097270958 | 0.41568841 | 1 |
| Dcaf15 | 0.92214963 | -0.116927234 | 0.41570972 | 1 |
| Plppr2 | 1.0619204 | 0.086675623 | 0.41571566 | 1 |
| Rap1gap | 0.96076552 | -0.057743713 | 0.41583378 | 1 |
| Lrrc56 | 1.20397281 | 0.267802816 | 0.41593075 | 1 |
| Tmem100 | 0.9426909 | -0.085143295 | 0.41596556 | 1 |
| Tsen2 | 1.1079558 | 0.147900335 | 0.41597551 | 1 |
| Fam3c | 1.04750419 | 0.066956019 | 0.41618842 | 1 |
| Iqgap1 | 1.04000435 | 0.056589563 | 0.41619635 | 1 |
| Slc6a13 | 1.08031313 | 0.111449546 | 0.41647176 | 1 |
| Prpf40a | 1.04840627 | 0.068197885 | 0.41647185 | 1 |
| Ppp2r2d | 1.05480242 | 0.076972789 | 0.41652049 | 1 |
| Hnrnph3 | 1.05773909 | 0.080983807 | 0.41655876 | 1 |
| Tbc1d2b | 0.9492089 | -0.075202464 | 0.41656163 | 1 |
| Wdpcp | 1.10499566 | 0.144040703 | 0.41656825 | 1 |
| Cnnm2 | 0.9272537 | -0.108963981 | 0.41658867 | 1 |
| Rhag | 0.82625162 | -0.275346894 | 0.41658994 | 1 |
| Dact3 | 0.93466639 | -0.097476573 | 0.4166328 | 1 |

| | | | | |
|---|---|---|---|---|
| Arhgap22 | 0.93516838 | -0.096701943 | 0.41663737 | 1 |
| Zfp418 | 0.89547767 | -0.159270645 | 0.41672549 | 1 |
| Gsta1 | 0.9595287 | -0.059602129 | 0.41681159 | 1 |
| Chst12 | 1.06881018 | 0.09600566 | 0.41682052 | 1 |
| Gen1 | 0.83901706 | -0.253227949 | 0.41689699 | 1 |
| Ttyh1 | 0.96178134 | -0.056219158 | 0.41720426 | 1 |
| Fscn1 | 0.93633733 | -0.094899718 | 0.41736286 | 1 |
| Nav3 | 0.91766042 | -0.123967708 | 0.41746145 | 1 |
| Trpm8 | 0.95726998 | -0.063002224 | 0.41747806 | 1 |
| Zbtb48 | 0.89155871 | -0.165598297 | 0.41760563 | 1 |
| Adrm1 | 0.95276363 | -0.069809748 | 0.41772034 | 1 |
| Grin2d | 0.78166698 | -0.355373993 | 0.41774988 | 1 |
| Fntb | 1.04860596 | 0.068472646 | 0.41776548 | 1 |
| Tusc3 | 0.95859466 | -0.061007198 | 0.41784915 | 1 |
| Aak1 | 0.96365129 | -0.053416908 | 0.41792046 | 1 |
| Tmem68 | 1.05565456 | 0.078137827 | 0.41799631 | 1 |
| Slc35b2 | 0.93894908 | -0.090881172 | 0.41800379 | 1 |
| Pik3c2g | 1.14345656 | 0.193401564 | 0.41802324 | 1 |
| Ccdc97 | 1.06014334 | 0.084259339 | 0.41821531 | 1 |
| Cd300a | 0.82578796 | -0.276156706 | 0.4182896 | 1 |
| Agpat2 | 1.19576038 | 0.257928314 | 0.41845705 | 1 |
| Bcl6b | 1.176137 | 0.234056124 | 0.4184764 | 1 |
| Hdac8 | 1.09736986 | 0.134049854 | 0.41848316 | 1 |
| Fibp | 1.05654054 | 0.079348128 | 0.41851275 | 1 |
| Pxmp4 | 1.09838066 | 0.135378123 | 0.41854567 | 1 |
| Zfp691 | 1.10086301 | 0.138634959 | 0.4186387 | 1 |
| Lamc2 | 1.13109811 | 0.177724077 | 0.41871558 | 1 |
| Ptpro | 0.93240224 | -0.100975632 | 0.4188255 | 1 |
| Nbr1 | 1.0378735 | 0.053630611 | 0.4189369 | 1 |
| Gpt | 1.12267996 | 0.166946719 | 0.41899503 | 1 |
| Plcg1 | 0.96128263 | -0.056967423 | 0.41901477 | 1 |
| Hmg20a | 1.04623342 | 0.065204765 | 0.41906476 | 1 |
| Serinc1 | 1.03996014 | 0.056528237 | 0.41913624 | 1 |
| Ehbp1l1 | 1.0532239 | 0.074812168 | 0.41926105 | 1 |
| Qsox1 | 0.95100446 | -0.072475984 | 0.41946118 | 1 |
| Arid5a | 1.13989445 | 0.188900242 | 0.41952815 | 1 |
| Ank3 | 0.96355911 | -0.05355492 | 0.41958351 | 1 |
| Tle3 | 0.94459651 | -0.082229896 | 0.41961437 | 1 |
| Dnajb9 | 0.95787802 | -0.062086151 | 0.41982586 | 1 |
| Smagp | 0.83159244 | -0.266051444 | 0.41991367 | 1 |

| | | | | |
|---|---|---|---|---|
| Irf2bp1 | 0.93843139 | -0.091676821 | 0.41993843 | 1 |
| Stx12 | 1.03771849 | 0.053415123 | 0.41998575 | 1 |
| Slc6a17 | 0.9634831 | -0.053668736 | 0.42009325 | 1 |
| Rps11 | 1.03805833 | 0.053887509 | 0.42009402 | 1 |
| Npepo | 1.11718631 | 0.159869796 | 0.42035058 | 1 |
| Trim21 | 0.89632443 | -0.15790708 | 0.42035807 | 1 |
| Shc1 | 0.96203847 | -0.055833514 | 0.42040652 | 1 |
| Capn1 | 0.96287139 | -0.054584979 | 0.42041992 | 1 |
| Abcd4 | 1.12147621 | 0.165399021 | 0.42060552 | 1 |
| Acot11 | 1.13084649 | 0.177403094 | 0.42064192 | 1 |
| Snx11 | 1.05317505 | 0.074745256 | 0.42065482 | 1 |
| Lrrc24 | 0.93805158 | -0.092260841 | 0.42077736 | 1 |
| Dok2 | 0.84494441 | -0.243071674 | 0.42095273 | 1 |
| Mfsd2a | 0.94032458 | -0.088769258 | 0.42120936 | 1 |
| Appl2 | 0.94429797 | -0.082685922 | 0.42143313 | 1 |
| Slc25a45 | 1.07822005 | 0.108651643 | 0.42156985 | 1 |
| Tac1 | 0.96385215 | -0.053116233 | 0.42166613 | 1 |
| Rab12 | 0.89266424 | -0.163810456 | 0.42179355 | 1 |
| Rgs19 | 0.90894121 | -0.137741115 | 0.42184877 | 1 |
| Psmb6 | 1.04899541 | 0.069008363 | 0.42186982 | 1 |
| Notch4 | 1.09634399 | 0.132700532 | 0.42194316 | 1 |
| Sox6 | 0.86963511 | -0.201517905 | 0.42195653 | 1 |
| Rnh1 | 1.04240783 | 0.059919821 | 0.42215892 | 1 |
| Emc4 | 1.04163879 | 0.05885508 | 0.42217519 | 1 |
| Hs2st1 | 0.95822927 | -0.061557208 | 0.42217572 | 1 |
| Celf4 | 0.9488439 | -0.075757337 | 0.42229184 | 1 |
| Rb1 | 0.95448724 | -0.067202191 | 0.42236994 | 1 |
| Mvk | 0.94428361 | -0.082707859 | 0.42238619 | 1 |
| Hmgb2 | 1.09279747 | 0.12802605 | 0.42243554 | 1 |
| Magee1 | 0.96265384 | -0.054910987 | 0.4224858 | 1 |
| Zfx | 1.08420541 | 0.116638109 | 0.42286727 | 1 |
| Eif4a1 | 1.03740515 | 0.052979436 | 0.42287825 | 1 |
| Omd | 1.05337942 | 0.075025174 | 0.42292217 | 1 |
| Nr1h3 | 1.10048239 | 0.138136064 | 0.42300787 | 1 |
| Taok2 | 0.95687647 | -0.063595411 | 0.42303969 | 1 |
| Klhl23 | 1.08688555 | 0.120200033 | 0.42304067 | 1 |
| Cdc25a | 0.92480966 | -0.112771626 | 0.42318484 | 1 |
| Psmg2 | 1.06566528 | 0.09175437 | 0.42326657 | 1 |
| Zfp280b | 0.90344575 | -0.146490118 | 0.42334851 | 1 |
| Gabrb2 | 0.87672222 | -0.189808276 | 0.42336272 | 1 |

| | | | | |
|---|---|---|---|---|
| Pdzrn4 | 1.20918408 | 0.274033892 | 0.42336955 | 1 |
| RGD1566239 | 1.07366766 | 0.102547495 | 0.42372896 | 1 |
| Psmc3 | 1.0377273 | 0.053427368 | 0.42384698 | 1 |
| Nploc4 | 0.96229219 | -0.055453079 | 0.42393877 | 1 |
| Phc3 | 0.93959457 | -0.089889722 | 0.42395794 | 1 |
| Dcaf17 | 0.89358376 | -0.162325123 | 0.42396449 | 1 |
| Alox15 | 1.08921386 | 0.123287246 | 0.42396534 | 1 |
| Slc39a7 | 1.04435381 | 0.06261055 | 0.42398139 | 1 |
| Per3 | 0.94640375 | -0.079472311 | 0.42435015 | 1 |
| Golgb1 | 0.96406219 | -0.052801877 | 0.42438279 | 1 |
| Vps4a | 0.96065027 | -0.057916791 | 0.42439497 | 1 |
| Yipf6 | 1.04414502 | 0.062322097 | 0.42448045 | 1 |
| Ltbp4 | 0.95258773 | -0.070076132 | 0.42451846 | 1 |
| Ostf1 | 1.03798581 | 0.053786725 | 0.4246101 | 1 |
| Apln | 0.83927687 | -0.252781276 | 0.42464825 | 1 |
| Raver2 | 0.88974628 | -0.168534106 | 0.42494724 | 1 |
| Birc2 | 1.05565353 | 0.078136415 | 0.42498029 | 1 |
| Mcur1 | 0.90549059 | -0.143228445 | 0.42520536 | 1 |
| Prss23 | 1.06452371 | 0.090208082 | 0.42521316 | 1 |
| Ggnbp1 | 1.27127192 | 0.346272653 | 0.42532247 | 1 |
| Scamp2 | 1.06569326 | 0.09179224 | 0.42532902 | 1 |
| Zfyve1 | 0.95248684 | -0.070228931 | 0.42536011 | 1 |
| Tars2 | 0.94852111 | -0.076248215 | 0.4255252 | 1 |
| St6gal1 | 1.04586768 | 0.064700337 | 0.4259335 | 1 |
| Slc12a3 | 0.89445998 | -0.160911154 | 0.42597462 | 1 |
| Dmrtc1a | 1.11145559 | 0.152450301 | 0.42604828 | 1 |
| Cdh13 | 1.03892784 | 0.055095458 | 0.42622834 | 1 |
| Pik3ap1 | 0.85935567 | -0.218672738 | 0.42627697 | 1 |
| Ndufa8 | 1.04484712 | 0.06329186 | 0.42636979 | 1 |
| Zfp426 | 1.10312509 | 0.141596399 | 0.4263751 | 1 |
| Ypel3 | 1.0379255 | 0.053702889 | 0.42641239 | 1 |
| RGD1564541 | 1.06611029 | 0.092356699 | 0.42643844 | 1 |
| Mrps25 | 0.94105035 | -0.087656178 | 0.42646347 | 1 |
| Pou6f1 | 0.89943925 | -0.152902257 | 0.42653837 | 1 |
| Pip4k2b | 0.95088355 | -0.072659424 | 0.42655539 | 1 |
| Gtf2i | 0.96393967 | -0.05298524 | 0.42662772 | 1 |
| Flt4 | 0.84452577 | -0.243786641 | 0.42701353 | 1 |
| Sec16a | 0.95929584 | -0.05995229 | 0.42705011 | 1 |
| Tyms | 1.1003102 | 0.13791031 | 0.4271064 | 1 |
| Slc25a37 | 1.10896524 | 0.149214142 | 0.42716055 | 1 |

| | | | | | |
|---|---|---|---|---|---|
| Socs4 | | 1.10211821 | 0.140278978 | 0.42720051 | 1 |
| Uck1 | | 0.9440513 | -0.083062843 | 0.42724575 | 1 |
| Poglut1 | | 1.04776727 | 0.067318302 | 0.42731616 | 1 |
| Dcun1d5 | | 0.9504356 | -0.073339215 | 0.42747698 | 1 |
| Atp1b3 | | 0.96310519 | -0.054234717 | 0.4274897 | 1 |
| Fbxo9 | | 0.95660447 | -0.064005557 | 0.42755241 | 1 |
| Abcb10 | | 0.93333521 | -0.099532769 | 0.42758579 | 1 |
| Mettl23 | | 0.88576391 | -0.175005873 | 0.42770426 | 1 |
| Gmpr | | 0.9521185 | -0.070786952 | 0.42797816 | 1 |
| Ythdf3 | | 1.04213442 | 0.059541379 | 0.42814208 | 1 |
| Nhej1 | | 0.85659965 | -0.223307003 | 0.42819313 | 1 |
| Ndufb4 | | 1.05090216 | 0.071628365 | 0.42822146 | 1 |
| Btbd3 | | 0.95797074 | -0.06194651 | 0.4284204 | 1 |
| Cwf19l1 | | 1.07189936 | 0.100169457 | 0.4285356 | 1 |
| Soat1 | | 1.06470212 | 0.090449852 | 0.42858478 | 1 |
| Vat1 | | 0.96368672 | -0.053363877 | 0.42859651 | 1 |
| Krt26 | | 0.8271679 | -0.273747901 | 0.42866999 | 1 |
| Tcea1 | | 1.04324417 | 0.061076864 | 0.42876062 | 1 |
| Adnp | | 0.96153179 | -0.056593538 | 0.42887449 | 1 |
| Pde4a | | 1.14253325 | 0.192236156 | 0.42896558 | 1 |
| Dusp3 | | 0.96438656 | -0.052316544 | 0.42900604 | 1 |
| Hoxd10 | | 0.51004017 | -0.971317208 | 0.42921299 | 1 |
| Mrpl10 | | 1.05370144 | 0.075466142 | 0.42931624 | 1 |
| | 2-Mar | 1.03883028 | 0.054959968 | 0.42951101 | 1 |
| Opcml | | 0.93259085 | -0.100683817 | 0.42951214 | 1 |
| Akap8 | | 1.04417219 | 0.062359645 | 0.42951322 | 1 |
| Exosc3 | | 1.16959714 | 0.226011691 | 0.42957613 | 1 |
| Ahdc1 | | 1.0639146 | 0.089382347 | 0.42974329 | 1 |
| Fibin | | 1.08527325 | 0.118058326 | 0.42980926 | 1 |
| Dip2c | | 0.95832974 | -0.061405957 | 0.42984633 | 1 |
| Alkbh6 | | 1.09344817 | 0.128884833 | 0.43003836 | 1 |
| Zfp414 | | 0.9261499 | -0.110682372 | 0.4302227 | 1 |
| Tbc1d10b | | 0.95790325 | -0.062048153 | 0.4303313 | 1 |
| Cadm4 | | 0.95613761 | -0.064709818 | 0.43041246 | 1 |
| Pigm | | 1.05194674 | 0.073061657 | 0.43061648 | 1 |
| Nrd1 | | 0.96268929 | -0.054857853 | 0.43096085 | 1 |
| Rrm1 | | 1.05502081 | 0.077271461 | 0.43100135 | 1 |
| Ftsj3 | | 0.94574598 | -0.080475356 | 0.43109634 | 1 |
| Stxbp5l | | 1.06723775 | 0.093881604 | 0.43118037 | 1 |
| Cenpa | | 0.73109361 | -0.451871945 | 0.43137772 | 1 |

| | | | | |
|---|---|---|---|---|
| Cdc25b | 0.95821616 | -0.061576944 | 0.43151445 | 1 |
| Gdap1l1 | 0.95070447 | -0.072931147 | 0.43163313 | 1 |
| Chst9 | 1.13753807 | 0.185914827 | 0.43207968 | 1 |
| Ston1 | 1.06488016 | 0.090691074 | 0.43216423 | 1 |
| Snf8 | 0.95005406 | -0.073918491 | 0.43221051 | 1 |
| Fam110b | 1.06340867 | 0.088696132 | 0.4322426 | 1 |
| Gramd1c | 0.80599636 | -0.311154769 | 0.43227108 | 1 |
| Slitrk5 | 1.05816231 | 0.081560942 | 0.43230536 | 1 |
| Fuk | 1.08507102 | 0.117789471 | 0.43240924 | 1 |
| Slc12a4 | 0.95439662 | -0.067339164 | 0.43271959 | 1 |
| Rab34 | 0.9434557 | -0.08397331 | 0.43273731 | 1 |
| RGD1561149 | 0.9561501 | -0.064690974 | 0.43275291 | 1 |
| Psd | 0.8895452 | -0.168860178 | 0.43276649 | 1 |
| Acot7 | 0.96462427 | -0.051960981 | 0.43287164 | 1 |
| Idnk | 0.89972421 | -0.152445259 | 0.43294273 | 1 |
| Klhl11 | 0.95459546 | -0.067038622 | 0.43302014 | 1 |
| LOC317456 | 0.91434953 | -0.129182315 | 0.4331111 | 1 |
| Trmt11 | 1.13026125 | 0.176656282 | 0.43323843 | 1 |
| Bspry | 1.20270897 | 0.266287586 | 0.43346598 | 1 |
| Aebp1 | 1.03992154 | 0.05647468 | 0.43381401 | 1 |
| LOC361990 | 1.03864377 | 0.054700935 | 0.4338535 | 1 |
| Rgs16 | 0.81994719 | -0.286397093 | 0.43386438 | 1 |
| Nefl | 1.0404957 | 0.057270999 | 0.43402324 | 1 |
| Rundc3b | 0.94652025 | -0.07929472 | 0.43404918 | 1 |
| Aldh1l2 | 0.85926951 | -0.218817391 | 0.43452856 | 1 |
| Ccng1 | 0.96472298 | -0.051813365 | 0.43454632 | 1 |
| Syde1 | 1.07181463 | 0.100055409 | 0.43466501 | 1 |
| Lipa | 1.03656546 | 0.051811233 | 0.43467502 | 1 |
| Gbf1 | 0.96412766 | -0.052703906 | 0.43474872 | 1 |
| Mfap1a | 1.05175267 | 0.072795485 | 0.43475685 | 1 |
| Pam | 0.96475165 | -0.051770489 | 0.43476484 | 1 |
| Tgif2 | 0.9025772 | -0.147877764 | 0.43478741 | 1 |
| Vegfc | 0.86232101 | -0.213703063 | 0.43499431 | 1 |
| l7Rn6 | 1.05947185 | 0.083345251 | 0.43506489 | 1 |
| Nsdhl | 0.95995647 | -0.05895911 | 0.43509845 | 1 |
| Anapc11 | 1.0558045 | 0.078342718 | 0.43517242 | 1 |
| Ints7 | 1.06580398 | 0.091942128 | 0.43525239 | 1 |
| Slc26a10 | 1.18602309 | 0.246132097 | 0.43531424 | 1 |
| Bri3bp | 1.17631841 | 0.23427863 | 0.43541383 | 1 |
| Supt7l | 1.10548926 | 0.144685006 | 0.43543721 | 1 |

| | | | | | |
|---|---|---|---|---|---|
| Myl6l | | 1.03670131 | 0.052000297 | 0.43544563 | 1 |
| Nipsnap3b | | 1.05582336 | 0.078368496 | 0.43548536 | 1 |
| Uqcrc1 | | 1.03691749 | 0.052301095 | 0.43555254 | 1 |
| Smim17 | | 0.90779452 | -0.13956231 | 0.43557821 | 1 |
| Lcmt1 | | 0.94843956 | -0.07637225 | 0.43558302 | 1 |
| Arhgap21 | | 0.96360211 | -0.05349054 | 0.43558965 | 1 |
| Mob3b | | 0.93482043 | -0.097238834 | 0.43560021 | 1 |
| Psme4 | | 1.0365458 | 0.051783868 | 0.43568717 | 1 |
| Cdt1 | | 0.82846045 | -0.271495264 | 0.43570476 | 1 |
| Chpf2 | | 0.95606943 | -0.064812703 | 0.4357169 | 1 |
| Coq10a | | 1.06892871 | 0.096165634 | 0.43582615 | 1 |
| Skap2 | | 1.06896934 | 0.096220473 | 0.43583585 | 1 |
| Trps1 | | 0.89078291 | -0.166854212 | 0.43610841 | 1 |
| Clcn1 | | 1.16167383 | 0.216205051 | 0.43616409 | 1 |
| Mrgprx3 | | 0.94226296 | -0.085798365 | 0.43630119 | 1 |
| Coq10b | | 0.94681498 | -0.078845563 | 0.43633417 | 1 |
| Thnsl2 | | 0.92170098 | -0.117629318 | 0.43642577 | 1 |
| Kdm8 | | 1.07604139 | 0.10573357 | 0.43647982 | 1 |
| Pde4dip | | 1.03719138 | 0.052682117 | 0.43656526 | 1 |
| Pigx | | 1.06859887 | 0.095720402 | 0.43662536 | 1 |
| Pik3r6 | | 1.18793062 | 0.248450583 | 0.43663991 | 1 |
| LOC500124 | | 0.83527272 | -0.259680775 | 0.43664481 | 1 |
| Tmem178b | | 0.9628316 | -0.054644602 | 0.43664968 | 1 |
| Dnajb6 | | 1.04521396 | 0.063798298 | 0.43667871 | 1 |
| LOC691141 | | 1.22471677 | 0.292448151 | 0.43669591 | 1 |
| Atl1 | | 1.03655917 | 0.051802471 | 0.43672791 | 1 |
| Coq5 | | 0.94549876 | -0.080852533 | 0.43679518 | 1 |
| Wdr4 | | 0.90975591 | -0.136448582 | 0.43729417 | 1 |
| Tmem168 | | 1.05117644 | 0.07200484 | 0.43736834 | 1 |
| Myo18a | | 0.96473052 | -0.051802081 | 0.43758615 | 1 |
| Abca13 | | 0.78728337 | -0.345045086 | 0.43770307 | 1 |
| Chd2 | | 1.04895087 | 0.068947113 | 0.43776876 | 1 |
| Mapre2 | | 0.96477553 | -0.051734778 | 0.43783055 | 1 |
| | 7-Mar | 1.04919986 | 0.069289516 | 0.4378331 | 1 |
| Ppp1r1c | | 0.96489942 | -0.051549526 | 0.43784004 | 1 |
| Man1a2 | | 0.95912934 | -0.06020272 | 0.43786933 | 1 |
| RGD1304624 | | 0.86471251 | -0.209707541 | 0.43795481 | 1 |
| Prr13 | | 1.0427389 | 0.060377959 | 0.43819647 | 1 |
| Entpd3 | | 1.03980402 | 0.056311631 | 0.43827767 | 1 |
| Ppm1e | | 1.05911386 | 0.082857695 | 0.4383453 | 1 |

| | | | | |
|---|---|---|---|---|
| Nr2c2 | 0.9457069 | -0.080534977 | 0.43835073 | 1 |
| Plk4 | 0.91666014 | -0.125541152 | 0.43852727 | 1 |
| Trak2 | 0.964892 | -0.051560621 | 0.43864162 | 1 |
| Eogt | 1.09295703 | 0.128236686 | 0.43869406 | 1 |
| Rfk | 1.04126602 | 0.058338685 | 0.43872109 | 1 |
| Pla2g16 | 0.95791382 | -0.06203223 | 0.43882543 | 1 |
| Iqgap3 | 0.8883926 | -0.170730726 | 0.43897473 | 1 |
| Rapgef6 | 0.95594219 | -0.065004727 | 0.43900401 | 1 |
| RGD1561778 | 1.23511808 | 0.304648968 | 0.43947685 | 1 |
| Zfp955a | 1.07602067 | 0.10570579 | 0.4394934 | 1 |
| Dnmt1 | 0.94429382 | -0.082692262 | 0.43949506 | 1 |
| Ccnh | 1.06771463 | 0.094526102 | 0.43969287 | 1 |
| Mapkapk2 | 1.05486976 | 0.077064884 | 0.43986653 | 1 |
| Comtd1 | 0.94149788 | -0.086970256 | 0.44000165 | 1 |
| Clns1a | 1.05063258 | 0.071258221 | 0.44008179 | 1 |
| Tmed7 | 1.03796571 | 0.053758782 | 0.4401201 | 1 |
| Nckap1l | 1.07570251 | 0.105279148 | 0.44029869 | 1 |
| Ipp | 1.07616947 | 0.105905281 | 0.44035727 | 1 |
| Dhx40 | 1.0446581 | 0.063030845 | 0.4404932 | 1 |
| Man1b1 | 0.96399949 | -0.052895718 | 0.44059426 | 1 |
| Polr2e | 1.04852752 | 0.068364726 | 0.44063414 | 1 |
| Aftph | 1.03938327 | 0.05572774 | 0.44069047 | 1 |
| Ifi27 | 1.03600065 | 0.051024912 | 0.44072898 | 1 |
| Adap2 | 1.16052112 | 0.214772776 | 0.440793 | 1 |
| Tfpi | 1.05109053 | 0.071886932 | 0.44085676 | 1 |
| Cpne5 | 0.86760798 | -0.20488477 | 0.44095423 | 1 |
| Trim3 | 0.95605099 | -0.064840529 | 0.44096002 | 1 |
| Arfgap1 | 1.05955071 | 0.083452636 | 0.44106734 | 1 |
| Ddx3x | 1.03675311 | 0.052072374 | 0.44112674 | 1 |
| Frmd3 | 0.94843628 | -0.076377247 | 0.44117638 | 1 |
| Mcm3 | 0.88125389 | -0.18237037 | 0.44126831 | 1 |
| Pcm1 | 0.96468516 | -0.051869925 | 0.44153396 | 1 |
| RT1-CE4 | 1.0948072 | 0.130676829 | 0.44157489 | 1 |
| Inpp4a | 0.96310676 | -0.054232365 | 0.44159267 | 1 |
| Fubp1 | 1.04606101 | 0.064966993 | 0.44179328 | 1 |
| Pcdhb3 | 0.90268938 | -0.147698461 | 0.44215687 | 1 |
| Cyb5r3 | 1.03600177 | 0.051026474 | 0.44225484 | 1 |
| Retsat | 0.94996459 | -0.074054358 | 0.44229225 | 1 |
| Sod2 | 1.03590558 | 0.050892511 | 0.44237852 | 1 |
| RT1-CE7 | 0.90648341 | -0.141647476 | 0.44237992 | 1 |

| | | | | |
|---|---|---|---|---|
| Rab3il1 | 1.09881463 | 0.135948029 | 0.44238141 | 1 |
| Prelid3a | 0.90244989 | -0.148081276 | 0.44242099 | 1 |
| Slc20a1 | 1.03704267 | 0.052475253 | 0.44244732 | 1 |
| Ctsb | 1.03835372 | 0.054297987 | 0.44269122 | 1 |
| Tspan2 | 0.96249683 | -0.055146304 | 0.44284958 | 1 |
| Zeb2 | 0.95978926 | -0.05921043 | 0.44293588 | 1 |
| Msi1 | 1.15128005 | 0.203238817 | 0.44294218 | 1 |
| Elovl4 | 1.04605683 | 0.064961234 | 0.44302805 | 1 |
| Abr | 0.96554147 | -0.050589869 | 0.44308073 | 1 |
| Mccc2 | 1.06590821 | 0.0920832 | 0.44308335 | 1 |
| Ppp1r2 | 0.96469016 | -0.051862438 | 0.44327231 | 1 |
| Def8 | 0.94875146 | -0.075897887 | 0.44328165 | 1 |
| Rpl21 | 1.03740254 | 0.052975802 | 0.44329868 | 1 |
| Dysf | 0.96392156 | -0.053012339 | 0.44334476 | 1 |
| Egfr | 1.08393043 | 0.116272163 | 0.44337792 | 1 |
| Cstf3 | 1.07289868 | 0.101513845 | 0.44395776 | 1 |
| H2afj | 0.91149798 | -0.133688631 | 0.44395899 | 1 |
| Ap2a2 | 0.96550964 | -0.050637431 | 0.44403192 | 1 |
| Ccnt2 | 1.04032417 | 0.057033149 | 0.44414896 | 1 |
| Clip2 | 0.9649755 | -0.051435783 | 0.44417845 | 1 |
| Heatr6 | 1.07889705 | 0.109557212 | 0.44434598 | 1 |
| Pde4b | 1.06442516 | 0.09007452 | 0.44434906 | 1 |
| Nos1 | 1.13663432 | 0.184768179 | 0.44463271 | 1 |
| Bcl2l2 | 0.96507186 | -0.051291723 | 0.44464203 | 1 |
| Mtus1 | 1.03686932 | 0.052234083 | 0.44464452 | 1 |
| Zfp524 | 0.89270068 | -0.163751572 | 0.44475559 | 1 |
| Pcdha6 | 1.07474661 | 0.103996562 | 0.44492094 | 1 |
| Stra6 | 1.08362943 | 0.115871486 | 0.44493663 | 1 |
| Pcgf3 | 0.93110268 | -0.102987815 | 0.44518548 | 1 |
| Ccdc53 | 1.05751441 | 0.080677322 | 0.44523046 | 1 |
| Med6 | 1.06872454 | 0.095890047 | 0.44544776 | 1 |
| Jam2 | 1.03821805 | 0.054109473 | 0.44578031 | 1 |
| Krcc1 | 0.95284178 | -0.069691428 | 0.44579108 | 1 |
| Zfp787 | 0.9095353 | -0.136798458 | 0.44584405 | 1 |
| Evpl | 1.1259655 | 0.171162623 | 0.44608747 | 1 |
| Sppl2b | 0.95116052 | -0.072239257 | 0.44614805 | 1 |
| Rcan2 | 0.96543567 | -0.050747968 | 0.44618318 | 1 |
| Rbsn | 1.04606761 | 0.064976102 | 0.44618932 | 1 |
| Nasp | 1.07347418 | 0.102287489 | 0.44629017 | 1 |
| Wdtc1 | 0.96125856 | -0.057003554 | 0.44635229 | 1 |

| | | | | | |
|---|---|---|---|---|---|
| Cdkn2c | | 1.10871158 | 0.148884115 | 0.44635731 | 1 |
| Cad | | 0.9487116 | -0.075958511 | 0.44637775 | 1 |
| Pla2g4a | | 0.91425689 | -0.1293285 | 0.44655737 | 1 |
| Crabp2 | | 1.05778296 | 0.081043645 | 0.44656406 | 1 |
| F8 | | 0.94207679 | -0.086083429 | 0.44672743 | 1 |
| Plp1 | | 0.96449917 | -0.052148103 | 0.44673669 | 1 |
| Copz1 | | 0.96269604 | -0.054847742 | 0.44690238 | 1 |
| Angel1 | | 0.95329279 | -0.069008709 | 0.44703582 | 1 |
| Ptn | | 0.96493042 | -0.051503185 | 0.44708803 | 1 |
| Lix1 | | 1.10125906 | 0.13915389 | 0.44714035 | 1 |
| Adcy3 | | 0.95115245 | -0.072251505 | 0.44722753 | 1 |
| Dclre1b | | 0.92856038 | -0.106932365 | 0.4472802 | 1 |
| Ddx52 | | 1.07483805 | 0.104119304 | 0.44734005 | 1 |
| Arpc5 | | 0.96531776 | -0.050924169 | 0.44737285 | 1 |
| Tfb1m | | 0.94967276 | -0.074497618 | 0.4473899 | 1 |
| Mrpl30 | | 1.0576684 | 0.080887389 | 0.44742221 | 1 |
| Dnal1 | | 1.11517871 | 0.157274921 | 0.44745853 | 1 |
| Cpeb1 | | 0.96445179 | -0.052218965 | 0.44750703 | 1 |
| Abcc4 | | 0.94018195 | -0.088988106 | 0.44781582 | 1 |
| S100a10 | | 1.03657534 | 0.051824971 | 0.44789908 | 1 |
| Tmcc3 | | 0.94935684 | -0.074977637 | 0.44803738 | 1 |
| Rnaseh2b | | 1.07003058 | 0.097652023 | 0.44811852 | 1 |
| Lamtor4 | | 1.0476217 | 0.067117842 | 0.44813615 | 1 |
| Rph3al | | 0.85432018 | -0.227151236 | 0.44815619 | 1 |
| Amd1 | | 1.03669129 | 0.051986343 | 0.44817478 | 1 |
| Gnl2 | | 1.04508758 | 0.063623847 | 0.44830709 | 1 |
| Rpp14 | | 1.05805268 | 0.081411464 | 0.44848367 | 1 |
| Fam58b | | 0.91537364 | -0.12756734 | 0.44852515 | 1 |
| Cadps | | 0.96591992 | -0.050024503 | 0.44884507 | 1 |
| Amigo2 | | 0.93930175 | -0.0903394 | 0.44885858 | 1 |
| Fam103a1 | | 1.05316374 | 0.074729749 | 0.44893839 | 1 |
| Mogat2 | | 0.90615819 | -0.142165175 | 0.44902233 | 1 |
| Eno1 | | 0.96560547 | -0.050494242 | 0.44915438 | 1 |
| Dnajc18 | | 1.05173962 | 0.072777581 | 0.44930592 | 1 |
| | 9-Mar | 1.0922288 | 0.1272751 | 0.44932909 | 1 |
| Zbtb24 | | 1.0891186 | 0.123161062 | 0.44933225 | 1 |
| Kctd9 | | 1.03856707 | 0.054594389 | 0.44947935 | 1 |
| Mcf2l | | 0.9615008 | -0.05664003 | 0.44968322 | 1 |
| Tubgcp6 | | 0.94267401 | -0.085169147 | 0.44973306 | 1 |
| Sash3 | | 0.88389342 | -0.178055681 | 0.44979074 | 1 |

| | | | | |
|---|---|---|---|---|
| Kcnip2 | 0.96223517 | -0.055538564 | 0.44986458 | 1 |
| Asf1a | 1.06543833 | 0.091447094 | 0.44995576 | 1 |
| Rcan3 | 0.89122222 | -0.166142885 | 0.45018757 | 1 |
| RGD1560289 | 1.15489619 | 0.207763181 | 0.45020319 | 1 |
| Rps6ka2 | 0.96447805 | -0.052179693 | 0.45021452 | 1 |
| Zfp282 | 0.93494329 | -0.097049232 | 0.45025486 | 1 |
| Paox | 1.05654566 | 0.079355117 | 0.45028677 | 1 |
| Asah2 | 1.04801154 | 0.067654604 | 0.45038508 | 1 |
| Mrpl16 | 1.05953315 | 0.083428721 | 0.45055876 | 1 |
| Fdxr | 1.11462461 | 0.156557919 | 0.45056112 | 1 |
| Slc9a3 | 0.91328248 | -0.13086693 | 0.45067199 | 1 |
| Mmadhc | 1.05007678 | 0.070494813 | 0.4509915 | 1 |
| Drc1 | 1.12946432 | 0.175638697 | 0.45115236 | 1 |
| Mertk | 0.94107872 | -0.087612686 | 0.45117186 | 1 |
| Zfp317 | 1.06816118 | 0.095129363 | 0.4512196 | 1 |
| Nfyc | 1.05578022 | 0.078309548 | 0.45139882 | 1 |
| Pmm2 | 1.04486168 | 0.063311968 | 0.45146657 | 1 |
| Apobec2 | 0.72671372 | -0.460540949 | 0.45160082 | 1 |
| Scn2b | 0.91490474 | -0.128306553 | 0.451601 | 1 |
| Gal3st3 | 0.86087767 | -0.216119852 | 0.4516089 | 1 |
| Itfg1 | 1.03552869 | 0.050367524 | 0.45184881 | 1 |
| Slc25a23 | 0.93613163 | -0.095216693 | 0.45196825 | 1 |
| Tmem86b | 0.83061869 | -0.267741759 | 0.45212471 | 1 |
| Hspb6 | 1.07565111 | 0.105210218 | 0.45218272 | 1 |
| Anxa4 | 1.03648368 | 0.051697396 | 0.45222701 | 1 |
| Il33 | 0.96470785 | -0.051835995 | 0.45230152 | 1 |
| Daglb | 0.94087144 | -0.087930482 | 0.45243572 | 1 |
| Hoga1 | 0.85336534 | -0.228764583 | 0.45252492 | 1 |
| MGC94199 | 1.05911371 | 0.082857487 | 0.45260054 | 1 |
| Pgm3 | 1.06468172 | 0.090422213 | 0.45260679 | 1 |
| Nfkbil1 | 0.92593807 | -0.111012391 | 0.45274663 | 1 |
| Cmtm6 | 0.9044611 | -0.144869644 | 0.45276028 | 1 |
| Arf6 | 0.95149298 | -0.071735082 | 0.45276381 | 1 |
| Ctsz | 1.04633747 | 0.065348237 | 0.45277855 | 1 |
| Sp140 | 0.92266426 | -0.116122314 | 0.45293726 | 1 |
| Twist1 | 1.10465496 | 0.143595809 | 0.45309937 | 1 |
| LRRTM1 | 0.918148 | -0.123201373 | 0.45330186 | 1 |
| Copg1 | 1.03515406 | 0.049845492 | 0.45332274 | 1 |
| Zfp532 | 1.04928338 | 0.069404357 | 0.4533332 | 1 |
| Kalrn | 0.95001724 | -0.073974394 | 0.45333344 | 1 |

| | | | | | |
|---|---|---|---|---|---|
| Sidt2 | | 0.96190577 | -0.056032529 | 0.45342356 | 1 |
| Rexo1 | | 0.94734225 | -0.078042366 | 0.45345112 | 1 |
| Entpd2 | | 0.95673465 | -0.06380925 | 0.45345887 | 1 |
| Snapc1 | | 0.93159866 | -0.102219525 | 0.45353421 | 1 |
| | 6-Mar | 0.96625609 | -0.049522498 | 0.4535604 | 1 |
| Nsl1 | | 1.18577204 | 0.245826682 | 0.45361195 | 1 |
| Gnai3 | | 0.96487712 | -0.05158287 | 0.45365073 | 1 |
| Gpat4 | | 0.96176752 | -0.056239891 | 0.45373467 | 1 |
| Mzf1 | | 0.91168446 | -0.133393516 | 0.45399678 | 1 |
| Cbr3 | | 0.89654782 | -0.157547559 | 0.45401442 | 1 |
| Cplx2 | | 0.94909828 | -0.075370614 | 0.45402357 | 1 |
| Pcdhb12 | | 0.8515355 | -0.231861412 | 0.45407144 | 1 |
| Cd101 | | 1.22426615 | 0.291917226 | 0.45414382 | 1 |
| Runx1t1 | | 0.92763256 | -0.108374642 | 0.45421473 | 1 |
| Tmx1 | | 0.96351991 | -0.05361361 | 0.45432836 | 1 |
| Zmat2 | | 1.04061341 | 0.057434207 | 0.45445078 | 1 |
| Fbn2 | | 0.87318796 | -0.195635858 | 0.45447783 | 1 |
| Cacna1g | | 0.87717386 | -0.189065273 | 0.45453153 | 1 |
| Tnfaip8 | | 0.91908573 | -0.121728651 | 0.45456556 | 1 |
| Actr5 | | 1.11736937 | 0.160106173 | 0.45470788 | 1 |
| Serpinb6 | | 1.03539617 | 0.05018288 | 0.4548033 | 1 |
| Synj2bp | | 0.96560621 | -0.050493144 | 0.45525399 | 1 |
| Pelp1 | | 0.94897635 | -0.075555955 | 0.45529409 | 1 |
| Rufy3 | | 1.03508711 | 0.04975219 | 0.45529533 | 1 |
| Cox5a | | 0.96295868 | -0.054454205 | 0.45533135 | 1 |
| Pole | | 0.90063889 | -0.150979327 | 0.45542692 | 1 |
| Txndc17 | | 1.05645064 | 0.079225365 | 0.45543828 | 1 |
| Higd1a | | 0.95592432 | -0.065031688 | 0.45547201 | 1 |
| Tmem120a | | 0.94768803 | -0.077515879 | 0.4555124 | 1 |
| Ptprt | | 0.92905034 | -0.106171328 | 0.45555911 | 1 |
| Kif5c | | 1.03531458 | 0.0500692 | 0.45567019 | 1 |
| Abca4 | | 1.17249509 | 0.229581884 | 0.45571783 | 1 |
| Atp11a | | 1.03612832 | 0.051202692 | 0.4557364 | 1 |
| Snx6 | | 1.0359841 | 0.051001862 | 0.45575252 | 1 |
| Pnrc1 | | 1.0523477 | 0.073611459 | 0.45585509 | 1 |
| Fam122b | | 0.9204601 | -0.119572913 | 0.45590685 | 1 |
| Fam133b | | 1.05681115 | 0.079717588 | 0.45600775 | 1 |
| Map1b | | 0.96285214 | -0.054613832 | 0.4561776 | 1 |
| Esyt1 | | 0.96645061 | -0.049232094 | 0.45638198 | 1 |
| Adprm | | 0.94300539 | -0.084662085 | 0.4563966 | 1 |

| | | | | |
|---|---|---|---|---|
| Cops4 | 1.03690579 | 0.052284817 | 0.456409 | 1 |
| Smpdl3a | 1.05740729 | 0.080531182 | 0.45644495 | 1 |
| Sar1b | 1.04818545 | 0.067893992 | 0.45654104 | 1 |
| Sdf2l1 | 1.09587759 | 0.13208666 | 0.45658107 | 1 |
| Gadd45g | 0.93285804 | -0.100270541 | 0.45707323 | 1 |
| Wdr7 | 0.96489741 | -0.05155254 | 0.45707856 | 1 |
| Dda1 | 0.95953839 | -0.059587565 | 0.45708129 | 1 |
| Rasgrp2 | 0.89902435 | -0.153567911 | 0.45722257 | 1 |
| LOC689064 | 0.96639692 | -0.049312234 | 0.45727753 | 1 |
| Pogz | 0.9511611 | -0.072238383 | 0.45728264 | 1 |
| Rbm34 | 0.93373467 | -0.098915436 | 0.45749532 | 1 |
| Denr | 1.0500419 | 0.070446899 | 0.45750754 | 1 |
| Myo16 | 0.86343447 | -0.211841408 | 0.45755937 | 1 |
| Reep6 | 1.08850656 | 0.122350109 | 0.45758131 | 1 |
| Aldh9a1 | 1.04918516 | 0.06926931 | 0.45760451 | 1 |
| Pbk | 0.78340993 | -0.352160683 | 0.45762632 | 1 |
| Degs1 | 1.03516732 | 0.049863982 | 0.45765611 | 1 |
| Krt25 | 1.17670489 | 0.234752545 | 0.45768341 | 1 |
| Blmh | 1.04557861 | 0.064301536 | 0.45775732 | 1 |
| Tor4a | 1.12183172 | 0.165856283 | 0.45776554 | 1 |
| Mcm4 | 1.06388314 | 0.089339693 | 0.45794726 | 1 |
| Spry3 | 1.13074893 | 0.177278629 | 0.45801081 | 1 |
| Fam175b | 1.04378677 | 0.061827027 | 0.45804982 | 1 |
| Plcb1 | 0.94548503 | -0.080873476 | 0.45815095 | 1 |
| Mrgprd | 0.93787297 | -0.092535571 | 0.45825555 | 1 |
| Krt28 | 1.10303807 | 0.141482587 | 0.45830034 | 1 |
| Pdgfra | 1.03980858 | 0.056317961 | 0.45842927 | 1 |
| Nt5c3b | 1.0692568 | 0.096608377 | 0.4584953 | 1 |
| Amph | 1.03615211 | 0.051235817 | 0.45850102 | 1 |
| Znhit2 | 0.93702525 | -0.093840166 | 0.45885715 | 1 |
| Rbpms | 1.15240539 | 0.204648319 | 0.4590317 | 1 |
| Zbtb17 | 1.06915349 | 0.096468986 | 0.45919677 | 1 |
| Zswim3 | 1.08048833 | 0.11168349 | 0.45924181 | 1 |
| Kdm2b | 0.92684709 | -0.109596752 | 0.45929972 | 1 |
| Dync2li1 | 1.07212317 | 0.100470662 | 0.45930516 | 1 |
| Dram1 | 1.11934055 | 0.162649032 | 0.45934426 | 1 |
| Tmem11 | 1.05795235 | 0.081274645 | 0.45945159 | 1 |
| Fbxw2 | 0.96091821 | -0.057514449 | 0.4595066 | 1 |
| Selenbp1 | 1.07653617 | 0.106396788 | 0.45954362 | 1 |
| Sqstm1 | 0.96606504 | -0.049807774 | 0.45959814 | 1 |

| | | | | |
|---|---|---|---|---|
| Dgcr14 | 0.9368055 | -0.094178547 | 0.45967004 | 1 |
| Htt | 0.96512653 | -0.051210002 | 0.45971285 | 1 |
| Slk | 1.04322619 | 0.061052 | 0.45977479 | 1 |
| Gstp1 | 1.04034376 | 0.057060317 | 0.45979748 | 1 |
| Dlgap4 | 0.94910308 | -0.075363311 | 0.45980241 | 1 |
| Tipin | 0.90690457 | -0.140977351 | 0.45984915 | 1 |
| Pick1 | 0.93763486 | -0.092901885 | 0.45995147 | 1 |
| Pla2r1 | 1.17254468 | 0.229642894 | 0.45998992 | 1 |
| Jakmip1 | 0.95934318 | -0.059881096 | 0.46000966 | 1 |
| Elavl3 | 0.95064676 | -0.073018728 | 0.46002098 | 1 |
| Tpk1 | 0.92088387 | -0.118908854 | 0.46012791 | 1 |
| Gapdh | 1.03608779 | 0.051146252 | 0.4602083 | 1 |
| Cldn5 | 1.04966592 | 0.069930226 | 0.46024326 | 1 |
| Capzb | 0.96679056 | -0.048724709 | 0.46061949 | 1 |
| Agtrap | 0.92723227 | -0.108997311 | 0.46077705 | 1 |
| Atat1 | 0.92407765 | -0.113914009 | 0.46104771 | 1 |
| Ankmy2 | 1.04134663 | 0.058450377 | 0.46146342 | 1 |
| 3-Mar | 1.1319428 | 0.178801063 | 0.46169553 | 1 |
| Vps45 | 0.95604707 | -0.064846441 | 0.46170448 | 1 |
| Cryzl1 | 1.05613942 | 0.078800298 | 0.46179736 | 1 |
| Itpkb | 0.95534798 | -0.065901775 | 0.46195944 | 1 |
| Vps72 | 1.06626414 | 0.092564877 | 0.4620728 | 1 |
| Tnfsf12 | 1.06073491 | 0.085064162 | 0.46209528 | 1 |
| Tshz3 | 1.05652939 | 0.079332899 | 0.46230316 | 1 |
| Ddb1 | 0.96687933 | -0.048592242 | 0.4623695 | 1 |
| Fbln1 | 1.04799501 | 0.067631843 | 0.46237704 | 1 |
| Sidt1 | 0.84815228 | -0.237604781 | 0.46241598 | 1 |
| Foxred2 | 0.85429156 | -0.22719956 | 0.46247256 | 1 |
| RGD1559909 | 1.06338591 | 0.088665262 | 0.46251229 | 1 |
| Hmgn5b | 0.85949851 | -0.218432964 | 0.46269088 | 1 |
| Prune | 1.051091 | 0.071887573 | 0.46270432 | 1 |
| Uvrag | 0.93958762 | -0.089900394 | 0.46271207 | 1 |
| Acly | 0.96633249 | -0.049408421 | 0.46274678 | 1 |
| Zc3h11a | 0.96344674 | -0.053723175 | 0.46290709 | 1 |
| Epb41l1 | 0.96651876 | -0.049130358 | 0.46315303 | 1 |
| Pum3 | 1.05850584 | 0.08202923 | 0.46326704 | 1 |
| Tollip | 0.96535696 | -0.050865586 | 0.46330005 | 1 |
| Cuta | 1.0508525 | 0.071560181 | 0.46361609 | 1 |
| Psmd12 | 1.03449908 | 0.048932363 | 0.46362845 | 1 |
| Tnfrsf21 | 0.9668125 | -0.048691974 | 0.46369369 | 1 |

| | | | | |
|---|---|---|---|---|
| Acot2 | 1.08706688 | 0.120440704 | 0.46372133 | 1 |
| Tmem192 | 1.06001798 | 0.084088733 | 0.46382874 | 1 |
| Efhd2 | 0.94682142 | -0.078835757 | 0.46387089 | 1 |
| Pwp1 | 1.07033258 | 0.098059151 | 0.46388388 | 1 |
| Prkrip1 | 0.93786071 | -0.092554431 | 0.46390734 | 1 |
| Cmtr2 | 0.92155838 | -0.117852539 | 0.46394464 | 1 |
| Dync1i1 | 1.0343194 | 0.048681766 | 0.46399938 | 1 |
| Cntnap5c | 0.92128443 | -0.11828146 | 0.46414692 | 1 |
| Zfp354a | 1.13575352 | 0.18364978 | 0.46417171 | 1 |
| Ift88 | 1.05695729 | 0.079917082 | 0.4642129 | 1 |
| Trmt44 | 0.91317099 | -0.131043067 | 0.46423355 | 1 |
| Zfpm1 | 0.93505465 | -0.096877408 | 0.46433923 | 1 |
| Clec5a | 1.20149117 | 0.264826047 | 0.46436336 | 1 |
| Adcy4 | 0.90154382 | -0.149530479 | 0.46439656 | 1 |
| Dctn3 | 1.04098262 | 0.057945988 | 0.46441348 | 1 |
| Mtmr10 | 1.05317064 | 0.07473921 | 0.4644159 | 1 |
| Zfand2b | 0.94438068 | -0.082559565 | 0.46452806 | 1 |
| Tcea3 | 0.85340183 | -0.228702885 | 0.46457698 | 1 |
| Dctn5 | 1.03797977 | 0.053778328 | 0.46460969 | 1 |
| Parp3 | 1.04955442 | 0.069776971 | 0.46473449 | 1 |
| Lonp1 | 0.96264665 | -0.05492175 | 0.46474183 | 1 |
| Ogdhl | 0.92955198 | -0.10539255 | 0.46482615 | 1 |
| Them6 | 1.03482528 | 0.049387208 | 0.46483742 | 1 |
| Arl4d | 0.92819957 | -0.10749306 | 0.46493891 | 1 |
| P3h1 | 0.93546735 | -0.096240793 | 0.4650775 | 1 |
| Zfp516 | 0.93607863 | -0.095298368 | 0.46517379 | 1 |
| Ptprr | 1.04238566 | 0.059889147 | 0.46525891 | 1 |
| Snrnp40 | 0.94127196 | -0.08731648 | 0.4653054 | 1 |
| Cggbp1 | 0.9594534 | -0.059715364 | 0.46551939 | 1 |
| Tmem126b | 1.0619305 | 0.086689355 | 0.46563722 | 1 |
| Wdr44 | 0.95906099 | -0.060305524 | 0.46588646 | 1 |
| Glb1l | 1.07263295 | 0.101156475 | 0.46598619 | 1 |
| Mr1 | 1.14844742 | 0.199684807 | 0.46606545 | 1 |
| Kctd21 | 1.13227985 | 0.179230573 | 0.46610121 | 1 |
| Nr1d1 | 1.04604899 | 0.064950415 | 0.46664488 | 1 |
| Hpcal1 | 0.96065501 | -0.057909675 | 0.46665367 | 1 |
| Hacl1 | 1.06567613 | 0.091769053 | 0.46698427 | 1 |
| Tmco3 | 1.04274146 | 0.060381501 | 0.46706421 | 1 |
| Hcn2 | 0.96345011 | -0.053718128 | 0.46719167 | 1 |
| Zfp709 | 0.87397597 | -0.194334481 | 0.46726186 | 1 |

| | | | | |
|---|---|---|---|---|
| Myadml2 | 0.86504739 | -0.20914893 | 0.46744071 | 1 |
| Mrpl35 | 1.05229814 | 0.073543509 | 0.46744095 | 1 |
| Smyd4 | 1.08304813 | 0.115097362 | 0.46744412 | 1 |
| Arpc3 | 1.03534649 | 0.050113666 | 0.4675823 | 1 |
| Prkaca | 0.96609862 | -0.049757626 | 0.46768517 | 1 |
| Psmc1 | 1.03382063 | 0.047985903 | 0.46772523 | 1 |
| Gcfc2 | 0.90565798 | -0.142961771 | 0.46785518 | 1 |
| Vps33b | 1.05804631 | 0.081402779 | 0.46793537 | 1 |
| Lmcd1 | 0.88730158 | -0.172503563 | 0.46803731 | 1 |
| Sncb | 0.96543411 | -0.050750295 | 0.46804503 | 1 |
| Adamts12 | 1.06577232 | 0.091899266 | 0.46844169 | 1 |
| Mgst3 | 1.03901332 | 0.055214154 | 0.46847224 | 1 |
| Rtn1 | 0.9663488 | -0.049384072 | 0.46878269 | 1 |
| Chi3l1 | 1.10501343 | 0.14406391 | 0.46886075 | 1 |
| Plpp7 | 1.09523299 | 0.131237809 | 0.46898116 | 1 |
| Brinp3 | 1.11708573 | 0.159739913 | 0.46899558 | 1 |
| Myl12b | 1.03522745 | 0.049947783 | 0.4690659 | 1 |
| Sart1 | 0.95334493 | -0.068929802 | 0.4691453 | 1 |
| Smc4 | 1.04367597 | 0.061673872 | 0.46919079 | 1 |
| Ccl7 | 0.85413542 | -0.227463277 | 0.46920473 | 1 |
| Capg | 1.04928414 | 0.069405409 | 0.46936464 | 1 |
| Zfp192 | 1.0617872 | 0.086494661 | 0.46948956 | 1 |
| Hsh2d | 1.30341748 | 0.382299249 | 0.46951738 | 1 |
| Atp5g2 | 1.04013176 | 0.056766302 | 0.46957347 | 1 |
| Ring1 | 1.07024052 | 0.097935063 | 0.4696374 | 1 |
| Fat3 | 0.95200657 | -0.070956563 | 0.46978045 | 1 |
| Gak | 0.96519347 | -0.051109935 | 0.46978214 | 1 |
| Tmem132b | 0.90170912 | -0.14926598 | 0.46981896 | 1 |
| Sap18 | 1.05447067 | 0.076518969 | 0.46982248 | 1 |
| Ddhd2 | 0.96053313 | -0.058092715 | 0.46991919 | 1 |
| Eef1g | 1.03387335 | 0.048059472 | 0.47003975 | 1 |
| Cyp2d4 | 0.9124661 | -0.132157129 | 0.47021487 | 1 |
| Lmnb1 | 1.14250797 | 0.192204228 | 0.47028533 | 1 |
| Brd4 | 0.96063744 | -0.057936061 | 0.47034997 | 1 |
| Gtf2a1 | 0.91054699 | -0.135194619 | 0.47044849 | 1 |
| Cct5 | 1.03387256 | 0.048058362 | 0.47057217 | 1 |
| Gatad2b | 0.92741528 | -0.108712596 | 0.47066375 | 1 |
| Syt4 | 0.96723229 | -0.048065686 | 0.47078467 | 1 |
| Bloc1s4 | 1.0829355 | 0.114947322 | 0.47094254 | 1 |
| MGC105567 | 1.10460654 | 0.143532573 | 0.4709929 | 1 |

| | | | | |
|---|---|---|---|---|
| Bub1b | 0.8210933 | -0.284381933 | 0.47102575 | 1 |
| LOC300308 | 1.09726105 | 0.1339068 | 0.47115505 | 1 |
| Nup153 | 0.96124388 | -0.057025585 | 0.47132041 | 1 |
| Tmem71 | 1.28425824 | 0.360935335 | 0.4713526 | 1 |
| Col18a1 | 0.95301816 | -0.069424393 | 0.47148855 | 1 |
| Cyc1 | 0.9666047 | -0.049002086 | 0.47151106 | 1 |
| Atg4d | 1.05278256 | 0.074207498 | 0.47155616 | 1 |
| Whamm | 0.90589114 | -0.142590408 | 0.47166333 | 1 |
| Aqr | 0.96329792 | -0.053946039 | 0.47179171 | 1 |
| Paip2 | 1.03350664 | 0.047547655 | 0.47180573 | 1 |
| Ube2s | 0.95900616 | -0.060388014 | 0.47181686 | 1 |
| Isl1 | 1.05188841 | 0.072981668 | 0.47191746 | 1 |
| LOC689574 | 0.95037903 | -0.07342509 | 0.47194183 | 1 |
| Tmub1 | 0.9341519 | -0.098270938 | 0.4719808 | 1 |
| Kctd17 | 0.95452305 | -0.067148051 | 0.47208864 | 1 |
| Stam | 1.03660438 | 0.051865392 | 0.47229792 | 1 |
| Bmi1 | 1.07062297 | 0.098450512 | 0.47252313 | 1 |
| Axl | 1.04488399 | 0.063342775 | 0.47254017 | 1 |
| Nes | 0.96672084 | -0.048828748 | 0.47255165 | 1 |
| Dennd1c | 0.88817883 | -0.171077914 | 0.47259031 | 1 |
| Tst | 1.06655207 | 0.092954403 | 0.47312369 | 1 |
| Zswim7 | 1.16623051 | 0.221852976 | 0.47315019 | 1 |
| Lgi4 | 1.03476307 | 0.049300465 | 0.47333817 | 1 |
| Rnf185 | 0.95109979 | -0.072331374 | 0.47336544 | 1 |
| Aldh7a1 | 0.95318214 | -0.069176176 | 0.47340291 | 1 |
| Cabp1 | 1.03510734 | 0.049780377 | 0.47361559 | 1 |
| Acad9 | 1.04414261 | 0.062318775 | 0.47362287 | 1 |
| Zdhhc12 | 0.90236908 | -0.14821046 | 0.47387315 | 1 |
| Hpse | 0.95301583 | -0.069427918 | 0.47387831 | 1 |
| Ago1 | 0.9484779 | -0.076313935 | 0.4739063 | 1 |
| Slc39a11 | 0.86996191 | -0.200975866 | 0.4739512 | 1 |
| Csnk1g3 | 1.03600824 | 0.051035476 | 0.47398164 | 1 |
| Cherp | 0.94666435 | -0.079075097 | 0.47409027 | 1 |
| Elf1 | 1.07126334 | 0.099313166 | 0.47417294 | 1 |
| Naa10 | 0.93188972 | -0.101768861 | 0.47423691 | 1 |
| Def6 | 0.91587236 | -0.126781542 | 0.47428158 | 1 |
| Cep290 | 1.04773758 | 0.067277421 | 0.47430569 | 1 |
| Gtf3c2 | 1.04452202 | 0.062842909 | 0.47449393 | 1 |
| Usp24 | 0.96201563 | -0.055867758 | 0.47452824 | 1 |
| Tmem263 | 1.09309196 | 0.128414775 | 0.47455933 | 1 |

| | | | | |
|---|---|---|---|---|
| Cops6 | 0.96380588 | -0.053185489 | 0.47456936 | 1 |
| H2afy | 0.95401637 | -0.067914074 | 0.47481112 | 1 |
| Tmem147 | 0.94131497 | -0.087250558 | 0.47489549 | 1 |
| Zbtb49 | 0.8842134 | -0.177533499 | 0.47498604 | 1 |
| Dennd6a | 1.04199077 | 0.059342494 | 0.47506856 | 1 |
| Pknox1 | 0.90055036 | -0.151121132 | 0.47510212 | 1 |
| Tll1 | 1.05498274 | 0.077219399 | 0.4751482 | 1 |
| Rcl1 | 0.92445828 | -0.113319883 | 0.47515973 | 1 |
| Chmp7 | 0.96096972 | -0.057437119 | 0.47536807 | 1 |
| Erbb4 | 0.85323182 | -0.228990331 | 0.47544199 | 1 |
| Gng3 | 0.96762177 | -0.047484867 | 0.47559863 | 1 |
| Cndp1 | 1.06747531 | 0.0942027 | 0.47561904 | 1 |
| Calcrl | 1.06856208 | 0.095670721 | 0.47581839 | 1 |
| Rassf8 | 0.92297134 | -0.11564225 | 0.47604359 | 1 |
| Phka1 | 1.05201695 | 0.073157946 | 0.4761438 | 1 |
| Crebrf | 1.03782239 | 0.053559573 | 0.47632071 | 1 |
| Upf2 | 1.04610184 | 0.065023302 | 0.4764745 | 1 |
| Capza1 | 0.96717431 | -0.048152178 | 0.47653987 | 1 |
| Dnajc6 | 1.03317364 | 0.04708274 | 0.47672388 | 1 |
| Serpina11 | 1.08138068 | 0.112874493 | 0.47681161 | 1 |
| Slc2a1 | 1.03740233 | 0.052975513 | 0.47707455 | 1 |
| Vkorc1l1 | 0.9241862 | -0.11374455 | 0.47722193 | 1 |
| Atpaf1 | 1.04570206 | 0.064471853 | 0.47723274 | 1 |
| Tspan12 | 1.03519988 | 0.049909357 | 0.47733159 | 1 |
| Fcho1 | 0.94244889 | -0.085513717 | 0.47737749 | 1 |
| Ube2j2 | 0.95560115 | -0.065519499 | 0.47746832 | 1 |
| Snd1 | 0.96487009 | -0.051593391 | 0.47757208 | 1 |
| Lmbrd1 | 1.03361049 | 0.047692614 | 0.4775746 | 1 |
| Dusp11 | 1.0698868 | 0.097458156 | 0.47770106 | 1 |
| Hsph1 | 0.96709301 | -0.048273449 | 0.4777312 | 1 |
| Bbs4 | 1.0494108 | 0.069579548 | 0.47786116 | 1 |
| Emb | 1.04184196 | 0.059136446 | 0.47809275 | 1 |
| Tgfbi | 1.03427912 | 0.048625582 | 0.4781132 | 1 |
| Psap | 1.03532557 | 0.050084514 | 0.47821917 | 1 |
| Ckmt2 | 0.88144297 | -0.182060871 | 0.47830903 | 1 |
| Dgkq | 1.04176247 | 0.059026366 | 0.47845607 | 1 |
| Sh3gl2 | 1.03545825 | 0.050269388 | 0.47851006 | 1 |
| RT1-Ba | 1.04742271 | 0.066843787 | 0.4785192 | 1 |
| Ttc1 | 1.03410647 | 0.048384733 | 0.47854828 | 1 |
| Tnpo2 | 0.96754596 | -0.047597907 | 0.47861782 | 1 |

| | | | | |
|---|---|---|---|---|
| Rhbdd2 | 0.94249813 | -0.085438346 | 0.47864411 | 1 |
| Otud7b | 1.04115442 | 0.058184066 | 0.47871751 | 1 |
| Fan1 | 0.92242318 | -0.11649933 | 0.47877021 | 1 |
| Traf3ip3 | 0.85571673 | -0.224794795 | 0.47891088 | 1 |
| Cbx3 | 1.03682975 | 0.052179021 | 0.47893254 | 1 |
| RGD1563888 | 1.14475567 | 0.195039707 | 0.47909238 | 1 |
| Magt1 | 1.06375226 | 0.0891622 | 0.47922799 | 1 |
| F11r | 1.06334165 | 0.088605212 | 0.47923467 | 1 |
| Cdk17 | 1.05386375 | 0.075688365 | 0.479332 | 1 |
| Inpp4b | 1.10104204 | 0.138869555 | 0.4794046 | 1 |
| Slc35d1 | 0.93431199 | -0.098023707 | 0.47955657 | 1 |
| Atf6 | 1.04257449 | 0.060150468 | 0.47956902 | 1 |
| Iffo1 | 0.96169706 | -0.056345593 | 0.47982915 | 1 |
| Htra3 | 1.04960691 | 0.069849121 | 0.4800109 | 1 |
| C1ql3 | 1.1314104 | 0.178122343 | 0.48010992 | 1 |
| Ankrd12 | 0.96585383 | -0.050123219 | 0.48019085 | 1 |
| Smo | 0.93238299 | -0.101005408 | 0.48022829 | 1 |
| Tp53inp1 | 1.11508869 | 0.157158467 | 0.48033096 | 1 |
| Eml2 | 1.03653729 | 0.051772013 | 0.48075855 | 1 |
| Fam118a | 1.05974026 | 0.083710702 | 0.48102534 | 1 |
| Prkcd | 0.96602171 | -0.049872483 | 0.48112869 | 1 |
| Coq4 | 1.10760684 | 0.14744587 | 0.48116099 | 1 |
| Zfp9 | 1.08338941 | 0.115551899 | 0.48124773 | 1 |
| Vav3 | 1.05811017 | 0.081489847 | 0.4812669 | 1 |
| Anapc2 | 1.0340653 | 0.048327286 | 0.4813072 | 1 |
| Atxn7l3b | 0.96833209 | -0.046426182 | 0.48146326 | 1 |
| Ilk | 0.96746187 | -0.047723286 | 0.48149758 | 1 |
| Mrps21 | 1.05333529 | 0.074964741 | 0.48174175 | 1 |
| Tgfbr2 | 0.96238147 | -0.055319232 | 0.48184964 | 1 |
| Prkcg | 0.86139665 | -0.215250389 | 0.48200426 | 1 |
| Rnf5 | 0.95601131 | -0.064900407 | 0.48211973 | 1 |
| Vps13b | 0.96593229 | -0.050006028 | 0.48229868 | 1 |
| Tnnc2 | 0.85200883 | -0.231059715 | 0.48232855 | 1 |
| Elk3 | 1.06093008 | 0.085329586 | 0.48238003 | 1 |
| Fcgr1a | 1.10198618 | 0.140106135 | 0.48250782 | 1 |
| Vps28 | 0.9615969 | -0.056495852 | 0.48259522 | 1 |
| Kcns3 | 1.03450472 | 0.048940229 | 0.48267425 | 1 |
| Trmt5 | 1.0584075 | 0.081895186 | 0.48315313 | 1 |
| RGD1307752 | 0.95754267 | -0.062591322 | 0.48316066 | 1 |
| Grin3b | 0.83635119 | -0.257819227 | 0.48320197 | 1 |

| | | | | |
|---|---|---|---|---|
| Rassf9 | 1.14618036 | 0.196834085 | 0.48322268 | 1 |
| Kcns1 | 1.04198901 | 0.059340062 | 0.48329343 | 1 |
| Farsa | 0.95285485 | -0.069671639 | 0.4833548 | 1 |
| Gpatch4 | 0.89127872 | -0.166051435 | 0.48336286 | 1 |
| Herc2 | 0.9683263 | -0.046434821 | 0.48339532 | 1 |
| Rell1 | 1.09075199 | 0.125323106 | 0.48353875 | 1 |
| Cep85 | 1.05152977 | 0.072489693 | 0.48363977 | 1 |
| Fbxo7 | 1.04134923 | 0.058453972 | 0.48373541 | 1 |
| Ppfia1 | 0.96058132 | -0.058020337 | 0.48378674 | 1 |
| Lrguk | 1.11348834 | 0.155086457 | 0.48378942 | 1 |
| LOC501110 | 1.07346469 | 0.102274734 | 0.48384138 | 1 |
| Rab43 | 1.16043868 | 0.214670285 | 0.48390214 | 1 |
| Cdk16 | 0.9679784 | -0.046953242 | 0.48391587 | 1 |
| Dtnbp1 | 1.0786692 | 0.10925249 | 0.48392361 | 1 |
| Cdc45 | 0.8242111 | -0.278914205 | 0.48413745 | 1 |
| Dhx32 | 0.95343991 | -0.06878608 | 0.48423125 | 1 |
| Hsbp1l1 | 0.84705545 | -0.239471674 | 0.48426728 | 1 |
| Nova1 | 0.92145427 | -0.118015523 | 0.48427459 | 1 |
| Gpc4 | 1.07075751 | 0.098631792 | 0.48437714 | 1 |
| Dnah1 | 0.88618984 | -0.174312314 | 0.48446685 | 1 |
| Gpat3 | 1.09944129 | 0.13677057 | 0.48447584 | 1 |
| Tut1 | 0.93983032 | -0.089527791 | 0.48455789 | 1 |
| Ccnd1 | 1.04058375 | 0.05739308 | 0.48477956 | 1 |
| Lancl2 | 0.95594135 | -0.065005984 | 0.48514704 | 1 |
| Inha | 0.90841046 | -0.138583775 | 0.48522711 | 1 |
| Ppp4r3b | 1.03850801 | 0.054512342 | 0.48538811 | 1 |
| Tfe3 | 0.95124994 | -0.072103633 | 0.48564417 | 1 |
| Fam134b | 0.96819345 | -0.046632767 | 0.48573217 | 1 |
| Thap11 | 0.93537004 | -0.096390872 | 0.48577467 | 1 |
| Slc16a12 | 1.07345744 | 0.102264993 | 0.4858596 | 1 |
| Tor1aip2 | 0.96400483 | -0.052887713 | 0.48587052 | 1 |
| Tatdn2 | 0.95727538 | -0.062994091 | 0.48593682 | 1 |
| Rtbdn | 0.88353676 | -0.17863793 | 0.48603467 | 1 |
| Clcn2 | 1.05038953 | 0.07092444 | 0.48607517 | 1 |
| Eaf1 | 0.96220759 | -0.055579921 | 0.48624957 | 1 |
| Rmdn3 | 1.04659719 | 0.065706288 | 0.48635504 | 1 |
| Trappc2l | 1.05281286 | 0.074249019 | 0.48641008 | 1 |
| Gorasp2 | 0.96627531 | -0.049493795 | 0.4864967 | 1 |
| Serpinb6b | 0.86374022 | -0.211330625 | 0.48654541 | 1 |
| Smg6 | 1.05533431 | 0.077700095 | 0.48674249 | 1 |

| | | | | |
|---|---|---|---|---|
| Lima1 | 1.05730102 | 0.080386184 | 0.48688757 | 1 |
| Arrb1 | 0.9517496 | -0.071346032 | 0.48691775 | 1 |
| Khdrbs1 | 0.96601746 | -0.049878834 | 0.48705293 | 1 |
| Ppip5k1 | 1.04610095 | 0.06502208 | 0.48709207 | 1 |
| Ddi2 | 0.95837543 | -0.061337175 | 0.4871869 | 1 |
| Dtd2 | 0.90401198 | -0.145586204 | 0.4872496 | 1 |
| LOC498154 | 0.92132281 | -0.118221359 | 0.48732356 | 1 |
| Ppia | 1.03414079 | 0.048432617 | 0.4875208 | 1 |
| Slc27a3 | 1.07509063 | 0.104458278 | 0.48760639 | 1 |
| Rps18 | 1.03620973 | 0.051316036 | 0.4882122 | 1 |
| Ppp2r3a | 1.04784232 | 0.06742164 | 0.48827976 | 1 |
| Sepw1 | 1.03222498 | 0.045757455 | 0.48845627 | 1 |
| Arl8b | 0.9684333 | -0.046275402 | 0.48850999 | 1 |
| Tmem175 | 1.04726618 | 0.06662818 | 0.48876586 | 1 |
| Mospd2 | 0.94996924 | -0.07404729 | 0.48876972 | 1 |
| Ralgps2 | 1.06186283 | 0.086597411 | 0.48883485 | 1 |
| Dhx34 | 0.93101242 | -0.103127682 | 0.48884681 | 1 |
| Csnk1g1 | 0.88700334 | -0.172988562 | 0.48890956 | 1 |
| Dnajc2 | 1.04287463 | 0.060565729 | 0.48896308 | 1 |
| Vps54 | 1.04612777 | 0.065059062 | 0.48898293 | 1 |
| Acap3 | 0.9556106 | -0.065505243 | 0.4890543 | 1 |
| Dnpep | 1.03641316 | 0.051599244 | 0.48930479 | 1 |
| Invs | 1.06638274 | 0.092725335 | 0.48934135 | 1 |
| Nop2 | 1.05676387 | 0.079653051 | 0.48936538 | 1 |
| Hapln2 | 1.09919459 | 0.136446805 | 0.48939706 | 1 |
| Sigmar1 | 0.9661991 | -0.049607588 | 0.48942852 | 1 |
| Gimap6 | 1.13904476 | 0.187824443 | 0.48948542 | 1 |
| Sptlc3 | 0.94506565 | -0.081513546 | 0.48957699 | 1 |
| Caly | 1.07715333 | 0.107223634 | 0.48962515 | 1 |
| Lpar3 | 1.06770393 | 0.094511653 | 0.48962832 | 1 |
| Tardbp | 1.03320141 | 0.047121523 | 0.48968201 | 1 |
| Arntl2 | 1.15188196 | 0.203992879 | 0.48972172 | 1 |
| Sytl1 | 0.83459164 | -0.260857634 | 0.4897516 | 1 |
| Cpsf7 | 0.95863874 | -0.060940855 | 0.48980037 | 1 |
| Serpinf1 | 0.96750984 | -0.047651753 | 0.48982913 | 1 |
| Fam167a | 1.08966598 | 0.123885962 | 0.48988229 | 1 |
| RGD1309104 | 0.95811659 | -0.061726866 | 0.48989462 | 1 |
| H3f3b | 1.03223502 | 0.045771488 | 0.48995345 | 1 |
| Prepl | 1.03213191 | 0.04562737 | 0.49002906 | 1 |
| Sptb | 0.9670329 | -0.048363116 | 0.49012315 | 1 |

| | | | | |
|---|---|---|---|---|
| Slc30a4 | 0.95454364 | -0.067116933 | 0.49013814 | 1 |
| Rbbp7 | 1.03252871 | 0.046181893 | 0.49018573 | 1 |
| Ndufaf1 | 0.94642713 | -0.079436662 | 0.49031193 | 1 |
| Lynx1 | 0.96833118 | -0.046427538 | 0.49042691 | 1 |
| Slc19a1 | 0.92308307 | -0.115467607 | 0.49056802 | 1 |
| Entpd6 | 0.95559308 | -0.065531696 | 0.49064812 | 1 |
| Rnf44 | 0.94261035 | -0.085266575 | 0.49084124 | 1 |
| RGD1311747 | 0.95405208 | -0.067860068 | 0.49096079 | 1 |
| Setd6 | 1.12030595 | 0.163892777 | 0.49100786 | 1 |
| Snrnp35 | 0.94249992 | -0.085435603 | 0.49106334 | 1 |
| Pik3c2b | 1.07137144 | 0.099458739 | 0.49110712 | 1 |
| Sdccag8 | 0.92359095 | -0.114674052 | 0.49119089 | 1 |
| Suox | 1.05712257 | 0.080142665 | 0.49119099 | 1 |
| Zfp644 | 0.96331058 | -0.05392709 | 0.49120137 | 1 |
| Serpinb9 | 1.03473254 | 0.049257912 | 0.49120307 | 1 |
| Rragc | 0.9673298 | -0.04792025 | 0.491349 | 1 |
| Med12 | 0.95922783 | -0.060054579 | 0.491385 | 1 |
| Cdk20 | 0.91685009 | -0.125242229 | 0.49178317 | 1 |
| Nsfl1c | 1.03346968 | 0.04749607 | 0.49180449 | 1 |
| Hk1 | 0.96900644 | -0.045421843 | 0.49182917 | 1 |
| Rxra | 0.91782983 | -0.123701393 | 0.49204749 | 1 |
| Cacna2d3 | 0.96564109 | -0.050441027 | 0.49205469 | 1 |
| Fam117a | 0.93839036 | -0.091739907 | 0.49228733 | 1 |
| Chrnb3 | 0.96078395 | -0.057716047 | 0.49234127 | 1 |
| Wdr37 | 0.96378946 | -0.053210069 | 0.49242879 | 1 |
| Arid1a | 0.96340039 | -0.053792594 | 0.49256732 | 1 |
| Renbp | 1.09130756 | 0.126057749 | 0.49262305 | 1 |
| Srsf2 | 0.9683659 | -0.046375811 | 0.49264351 | 1 |
| Rab11fip1 | 1.10161791 | 0.139623926 | 0.49269204 | 1 |
| Nmt2 | 1.04555579 | 0.064270048 | 0.49279874 | 1 |
| Mrvi1 | 1.06487113 | 0.090678847 | 0.49281952 | 1 |
| Tspan6 | 0.95428884 | -0.067502099 | 0.49284255 | 1 |
| Cmbl | 0.94097253 | -0.087775492 | 0.49287062 | 1 |
| Gpr52 | 1.17176783 | 0.228686743 | 0.49287934 | 1 |
| Chst14 | 0.89377121 | -0.162022513 | 0.49290546 | 1 |
| Tpi1 | 1.03230829 | 0.045873886 | 0.49306476 | 1 |
| Olfm1 | 0.96893473 | -0.045528607 | 0.49331493 | 1 |
| Fbll1 | 0.93924275 | -0.090430025 | 0.49342698 | 1 |
| Usp14 | 1.03229897 | 0.045860858 | 0.49346086 | 1 |
| Rab29 | 1.06621134 | 0.092493432 | 0.49347368 | 1 |

| | | | | |
|---|---|---|---|---|
| Cplx1 | 1.03322063 | 0.04714836 | 0.49352375 | 1 |
| Asic3 | 1.03465919 | 0.049155634 | 0.49361221 | 1 |
| Eif6 | 1.04127338 | 0.058348894 | 0.49361382 | 1 |
| Tceb1 | 1.03321508 | 0.047140609 | 0.49373079 | 1 |
| Magi1 | 0.96391602 | -0.053020637 | 0.49378187 | 1 |
| Wdr82 | 1.04285798 | 0.060542694 | 0.49382647 | 1 |
| Cacna1a | 0.9651814 | -0.051127976 | 0.49387329 | 1 |
| RGD1566099 | 0.86940414 | -0.201901121 | 0.49390523 | 1 |
| Ndufs8 | 0.95977344 | -0.059234204 | 0.49397934 | 1 |
| Wipf2 | 1.07746289 | 0.107638187 | 0.49402231 | 1 |
| Cd320 | 1.05780151 | 0.081068941 | 0.49402689 | 1 |
| Nmi | 1.08136663 | 0.112855738 | 0.49408479 | 1 |
| Myo7a | 1.16601656 | 0.221588284 | 0.49426868 | 1 |
| Camkk1 | 1.03828166 | 0.05419786 | 0.49429422 | 1 |
| Plcl2 | 1.07052828 | 0.098322913 | 0.49431099 | 1 |
| Sgms1 | 0.95193687 | -0.07106219 | 0.49438583 | 1 |
| Ano4 | 0.96638704 | -0.049326981 | 0.49442166 | 1 |
| Micu1 | 1.03265674 | 0.046360772 | 0.49456623 | 1 |
| Lgals1 | 1.03262992 | 0.046323311 | 0.4946064 | 1 |
| Fam104a | 1.04092862 | 0.057871137 | 0.49469424 | 1 |
| Odf2l | 1.08007942 | 0.111137402 | 0.49487306 | 1 |
| Fchsd1 | 0.94848841 | -0.076297951 | 0.49488511 | 1 |
| Ubac1 | 0.96370068 | -0.053342965 | 0.49489743 | 1 |
| Glcci1 | 0.94486961 | -0.081812836 | 0.4948975 | 1 |
| Osgepl1 | 1.05465541 | 0.076771705 | 0.49490144 | 1 |
| Rad51ap1 | 0.88557373 | -0.175315665 | 0.49491665 | 1 |
| Ncstn | 0.96594277 | -0.049990378 | 0.49501523 | 1 |
| LOC100912071 | 0.84862934 | -0.236793543 | 0.49505682 | 1 |
| Fam101b | 1.07474104 | 0.103989084 | 0.49507544 | 1 |
| Zfp286a | 1.11457618 | 0.156495232 | 0.49508967 | 1 |
| Fam234b | 0.96568125 | -0.050381022 | 0.49512763 | 1 |
| Scrn3 | 0.94181244 | -0.086488316 | 0.49518198 | 1 |
| Scarb1 | 0.951627 | -0.071531887 | 0.49519848 | 1 |
| Zdhhc16 | 1.06894085 | 0.096182027 | 0.49533948 | 1 |
| Dnajc14 | 0.96263861 | -0.054933806 | 0.49534174 | 1 |
| Adrbk2 | 0.92008721 | -0.120157484 | 0.49537316 | 1 |
| Rpl5 | 1.03168782 | 0.045006486 | 0.49554637 | 1 |
| Fam120a | 0.96863304 | -0.045977883 | 0.49564631 | 1 |
| Rbbp8 | 1.04993075 | 0.070294178 | 0.49586824 | 1 |
| Sema6c | 1.06817952 | 0.095154123 | 0.49593006 | 1 |

| | | | | |
|---|---|---|---|---|
| Ncbp1 | 0.96168146 | -0.056368984 | 0.495932 | 1 |
| Mthfd1l | 0.95863724 | -0.06094311 | 0.49594349 | 1 |
| Apbb3 | 1.03826386 | 0.054173136 | 0.49606973 | 1 |
| Asnsd1 | 1.03292456 | 0.046734896 | 0.4960915 | 1 |
| Syngr3 | 0.9664411 | -0.049246286 | 0.49617337 | 1 |
| Bag2 | 1.08309677 | 0.115162148 | 0.49621895 | 1 |
| Zfp496 | 1.07038234 | 0.098126223 | 0.49623991 | 1 |
| Siah2 | 0.91250722 | -0.132092127 | 0.49628175 | 1 |
| Abi3 | 0.86187162 | -0.214455103 | 0.49630301 | 1 |
| Aldh3a1 | 1.1214551 | 0.165371864 | 0.49633722 | 1 |
| Arl4c | 0.95769743 | -0.06235816 | 0.49657103 | 1 |
| Ube2e2 | 1.05886735 | 0.08252186 | 0.49671252 | 1 |
| Brf1 | 0.94343668 | -0.084002406 | 0.49672591 | 1 |
| Sgol2 | 0.86330873 | -0.212051515 | 0.49679629 | 1 |
| Golt1b | 0.94758443 | -0.077673603 | 0.49687724 | 1 |
| Acadsb | 0.95788255 | -0.06207932 | 0.49702429 | 1 |
| Tasp1 | 0.9311376 | -0.102933716 | 0.49712662 | 1 |
| Phb2 | 0.96309186 | -0.054254683 | 0.4971405 | 1 |
| Lonrf3 | 0.87650881 | -0.190159504 | 0.49725602 | 1 |
| Anapc15 | 1.06616213 | 0.092426837 | 0.49742009 | 1 |
| Rpl41 | 0.96891246 | -0.045561773 | 0.49742972 | 1 |
| Fstl1 | 1.0334416 | 0.047456858 | 0.49748604 | 1 |
| Lcp1 | 1.03211653 | 0.045605871 | 0.49749613 | 1 |
| Slc35e4 | 1.05326973 | 0.074874937 | 0.49759602 | 1 |
| Tmf1 | 0.96694171 | -0.048499169 | 0.49767306 | 1 |
| Map3k10 | 0.95838351 | -0.061325013 | 0.49776057 | 1 |
| Tyro3 | 0.96106497 | -0.057294133 | 0.49788822 | 1 |
| Prokr2 | 0.89323516 | -0.162888059 | 0.49801645 | 1 |
| Itgb4 | 1.03239823 | 0.045999572 | 0.49802876 | 1 |
| Klhl6 | 0.87616273 | -0.190729252 | 0.49814664 | 1 |
| Celf3 | 1.12945561 | 0.175627572 | 0.49817711 | 1 |
| Abcg2 | 1.0471446 | 0.066460677 | 0.49823975 | 1 |
| Naa40 | 0.90587413 | -0.142617499 | 0.49830494 | 1 |
| Pnisr | 1.0325171 | 0.046165672 | 0.49833356 | 1 |
| Pdhb | 1.03174713 | 0.045089432 | 0.49843793 | 1 |
| Wbp4 | 0.95372906 | -0.068348624 | 0.49855012 | 1 |
| Tada1 | 1.06278943 | 0.087855787 | 0.49855148 | 1 |
| Pphln1 | 1.04450467 | 0.062818946 | 0.49857693 | 1 |
| Mphosph6 | 1.06346187 | 0.08876831 | 0.49860704 | 1 |
| Zc3h3 | 0.93393284 | -0.098609282 | 0.49873608 | 1 |

| | | | | |
|---|---|---|---|---|
| Arfgef2 | 0.96759344 | -0.047527102 | 0.49880488 | 1 |
| Gzf1 | 1.05614022 | 0.078801385 | 0.49881542 | 1 |
| Adat1 | 0.89203357 | -0.164830087 | 0.49882457 | 1 |
| Sh3gl3 | 0.92532569 | -0.111966846 | 0.49906552 | 1 |
| Hist1h4b | 1.05537418 | 0.077754588 | 0.49915463 | 1 |
| Atp5c1 | 1.03172559 | 0.0450593 | 0.49916936 | 1 |
| Zc3h12a | 0.87872747 | -0.186512296 | 0.49917147 | 1 |
| Cnep1r1 | 1.04837431 | 0.068153901 | 0.49924968 | 1 |
| Gtpbp3 | 0.91662602 | -0.125594851 | 0.49927845 | 1 |
| Srr | 0.93671893 | -0.094311881 | 0.49930837 | 1 |
| Ncoa1 | 0.96279964 | -0.054692486 | 0.4993372 | 1 |
| Btrc | 0.95998804 | -0.058911659 | 0.49935895 | 1 |
| Tbcc | 0.94665488 | -0.079089532 | 0.49937525 | 1 |
| Rrp1 | 0.95672997 | -0.063816301 | 0.49948723 | 1 |
| Rtn2 | 0.96623888 | -0.049548185 | 0.49950004 | 1 |
| Vwa9 | 1.04398813 | 0.062105311 | 0.49981542 | 1 |
| Cr1l | 1.04001007 | 0.056597498 | 0.49994974 | 1 |
| Lpar1 | 0.96745183 | -0.047738259 | 0.49999775 | 1 |
| Yy1 | 0.94003918 | -0.089207211 | 0.50004196 | 1 |
| Aldh1l1 | 1.12002615 | 0.163532421 | 0.50011921 | 1 |
| Zfp251 | 1.09214375 | 0.127162762 | 0.50014893 | 1 |
| Ppil4 | 1.05770088 | 0.08093169 | 0.50020245 | 1 |
| Smarce1 | 0.96306543 | -0.054294274 | 0.5002036 | 1 |
| Trim47 | 0.91881945 | -0.122146703 | 0.50024938 | 1 |
| Lilra5 | 1.16163644 | 0.216158618 | 0.50025804 | 1 |
| Hfe | 1.12877653 | 0.174759902 | 0.50028362 | 1 |
| Eml5 | 0.93031857 | -0.104203268 | 0.50034136 | 1 |
| Ralbp1 | 0.96756701 | -0.047566521 | 0.50050819 | 1 |
| Mllt3 | 0.93516431 | -0.096708229 | 0.50095561 | 1 |
| Snapc5 | 0.94605135 | -0.080009606 | 0.50102761 | 1 |
| Polr2d | 1.07411021 | 0.103142027 | 0.50124264 | 1 |
| Met | 1.1172446 | 0.159945072 | 0.50130571 | 1 |
| Pld2 | 1.05015107 | 0.070596876 | 0.50133293 | 1 |
| Slc4a7 | 0.91699617 | -0.12501238 | 0.5016252 | 1 |
| Slc25a39 | 0.96588275 | -0.050080028 | 0.50163823 | 1 |
| Traf3ip1 | 0.92005269 | -0.120211617 | 0.50166682 | 1 |
| Ctdsp1 | 0.96343761 | -0.053736845 | 0.50169399 | 1 |
| B4galnt4 | 0.94565823 | -0.080609217 | 0.50173862 | 1 |
| Prodh | 1.05392722 | 0.075775239 | 0.50175375 | 1 |
| Zbtb11os1 | 0.89116375 | -0.166237539 | 0.50177411 | 1 |

| | | | | |
|---|---|---|---|---|
| Cyp2t1 | 0.9005598 | -0.151106011 | 0.50186157 | 1 |
| Psmb7 | 1.03131373 | 0.044483279 | 0.50195028 | 1 |
| Dyx1c1 | 1.16946377 | 0.225847164 | 0.50200354 | 1 |
| Wwc2 | 0.95596704 | -0.064967223 | 0.50207422 | 1 |
| Jkamp | 1.03658452 | 0.051837751 | 0.50230468 | 1 |
| Pcdh7 | 1.0429721 | 0.060700565 | 0.50235699 | 1 |
| Fam131b | 0.96855608 | -0.046092507 | 0.50251416 | 1 |
| Faxc | 1.05763719 | 0.080844808 | 0.50260878 | 1 |
| Gsap | 1.09983549 | 0.137287741 | 0.50281885 | 1 |
| Oasl2 | 1.10113436 | 0.138990516 | 0.5028489 | 1 |
| Fam181b | 0.90432623 | -0.145084782 | 0.50292593 | 1 |
| Iqcb1 | 1.0542115 | 0.076164336 | 0.50296664 | 1 |
| Plvap | 1.07368234 | 0.102567215 | 0.50315261 | 1 |
| Sh3bp4 | 1.04643014 | 0.065476002 | 0.50322389 | 1 |
| Calca | 0.96853621 | -0.046122108 | 0.50326666 | 1 |
| Prkar1a | 1.03128258 | 0.044439694 | 0.50326896 | 1 |
| Puf60 | 1.03303891 | 0.046894589 | 0.50334723 | 1 |
| Reep5 | 0.96810271 | -0.046767983 | 0.50341245 | 1 |
| Camta2 | 0.96770368 | -0.047362749 | 0.5034234 | 1 |
| Itga4 | 0.90960067 | -0.13669477 | 0.50347006 | 1 |
| Celf1 | 0.95034343 | -0.073479128 | 0.50347813 | 1 |
| Zscan21 | 0.93900923 | -0.090788763 | 0.50352772 | 1 |
| Ctxn1 | 0.95033151 | -0.073497232 | 0.50353129 | 1 |
| Mcm3ap | 0.96475204 | -0.051769903 | 0.50353983 | 1 |
| Snca | 1.03496886 | 0.049587356 | 0.50358377 | 1 |
| Trim45 | 1.07161005 | 0.099780022 | 0.50373616 | 1 |
| Clk4 | 0.94958636 | -0.074628879 | 0.50382229 | 1 |
| Pex11g | 1.20695736 | 0.271374703 | 0.50395522 | 1 |
| RGD1563348 | 1.06596768 | 0.092163699 | 0.50398037 | 1 |
| Zfp91 | 0.96874019 | -0.045818295 | 0.50411013 | 1 |
| Lbr | 1.08659792 | 0.119818185 | 0.50430195 | 1 |
| Slco5a1 | 0.95200509 | -0.070958806 | 0.5044286 | 1 |
| Galk2 | 1.05786398 | 0.081154141 | 0.50443335 | 1 |
| Dus2 | 0.92949347 | -0.105483362 | 0.50450669 | 1 |
| Afap1l1 | 0.94392787 | -0.083251478 | 0.50452987 | 1 |
| Wdr75 | 1.04813858 | 0.067829477 | 0.5045574 | 1 |
| Afg3l2 | 1.03585461 | 0.050821524 | 0.50456244 | 1 |
| Napepld | 0.896765 | -0.157198116 | 0.50472192 | 1 |
| Rpf2 | 1.05731394 | 0.080403808 | 0.50475007 | 1 |
| Med18 | 1.08300115 | 0.115034773 | 0.5049884 | 1 |

| | | | | |
|---|---|---|---|---|
| Ubiad1 | 0.9170321 | -0.124955857 | 0.50506569 | 1 |
| Apc | 0.96987083 | -0.044135471 | 0.50506793 | 1 |
| Gmppb | 0.93933408 | -0.090289744 | 0.50506886 | 1 |
| Ercc2 | 1.05880808 | 0.082441108 | 0.50513048 | 1 |
| Hk3 | 1.31096001 | 0.390623678 | 0.50515531 | 1 |
| Habp4 | 0.96470634 | -0.051838252 | 0.50518069 | 1 |
| Pycrl | 1.06155024 | 0.086172652 | 0.50528626 | 1 |
| Zmat4 | 0.9130712 | -0.13120073 | 0.50530908 | 1 |
| Dok4 | 0.96935385 | -0.0449047 | 0.50531892 | 1 |
| Gas2 | 1.04918221 | 0.069265244 | 0.50539701 | 1 |
| Fnip2 | 1.07333477 | 0.102100123 | 0.50549372 | 1 |
| Mrps36 | 1.04140801 | 0.058535402 | 0.50550335 | 1 |
| Ssfa2 | 1.03128101 | 0.044437496 | 0.50567627 | 1 |
| Etf1 | 0.96677169 | -0.048752866 | 0.50568298 | 1 |
| Slc16a14 | 0.90741397 | -0.140167225 | 0.50570742 | 1 |
| Patz1 | 0.94601958 | -0.080058057 | 0.50575105 | 1 |
| Ubxn2b | 0.91719382 | -0.124701464 | 0.50575441 | 1 |
| Slc35a1 | 0.94700276 | -0.07855946 | 0.50576196 | 1 |
| Rmnd5b | 0.94307483 | -0.084555852 | 0.50578664 | 1 |
| Ezh2 | 1.09597903 | 0.132220189 | 0.5058581 | 1 |
| Atp13a1 | 0.9583536 | -0.061370041 | 0.50589055 | 1 |
| Pllp | 0.95929014 | -0.059960865 | 0.50599278 | 1 |
| Mtch1 | 0.9699528 | -0.044013548 | 0.50606076 | 1 |
| Bbof1 | 1.10597082 | 0.145313326 | 0.50607169 | 1 |
| Pacs1 | 0.95420562 | -0.067627915 | 0.50607604 | 1 |
| Polr3gl | 0.90320046 | -0.146881875 | 0.50616347 | 1 |
| Calcoco1 | 1.03901501 | 0.055216493 | 0.50636453 | 1 |
| Atn1 | 0.96082997 | -0.057646948 | 0.50657663 | 1 |
| Ddx58 | 1.06490531 | 0.09072515 | 0.50666417 | 1 |
| P2ry12 | 1.11689233 | 0.159490111 | 0.50671804 | 1 |
| Mif4gd | 1.08926798 | 0.12335893 | 0.50690489 | 1 |
| Cd248 | 1.06627767 | 0.092583187 | 0.50696889 | 1 |
| Mavs | 1.05309889 | 0.074640915 | 0.50718914 | 1 |
| Asgr1 | 1.16836979 | 0.22449696 | 0.50728867 | 1 |
| Msl3l2 | 0.89838708 | -0.154590921 | 0.50735763 | 1 |
| Apoo | 0.95693642 | -0.063505023 | 0.50755964 | 1 |
| Rab8a | 0.95900777 | -0.060385596 | 0.50765732 | 1 |
| Dhrs11 | 1.08912095 | 0.123164175 | 0.50778656 | 1 |
| Cln8 | 1.06011524 | 0.084221102 | 0.50781378 | 1 |
| C1qc | 1.04294407 | 0.060661793 | 0.50784692 | 1 |

| | | | | |
|---|---|---|---|---|
| Prkcb | 0.96775277 | -0.047289557 | 0.50786702 | 1 |
| Hjurp | 1.17760571 | 0.235856578 | 0.50787498 | 1 |
| Sft2d3 | 1.09614074 | 0.132433047 | 0.50800847 | 1 |
| Sf3a3 | 1.04350486 | 0.061437318 | 0.50806999 | 1 |
| Cd164l2 | 1.12628 | 0.171565535 | 0.50809663 | 1 |
| Mrpl32 | 1.0522912 | 0.073534 | 0.5081424 | 1 |
| Rasgrp3 | 1.08072911 | 0.11200495 | 0.50829287 | 1 |
| Mlec | 1.04217625 | 0.059599285 | 0.50835175 | 1 |
| Susd2 | 0.96937783 | -0.044869012 | 0.5083941 | 1 |
| Cstf2t | 0.95134344 | -0.071961833 | 0.50868662 | 1 |
| Nap1l2 | 1.03132352 | 0.044496967 | 0.50877509 | 1 |
| RGD1306954 | 1.04834177 | 0.068109129 | 0.50877944 | 1 |
| Myd88 | 1.07399533 | 0.102987725 | 0.5088 | 1 |
| Snx8 | 0.93333913 | -0.099526711 | 0.50883219 | 1 |
| Gpaa1 | 0.95501854 | -0.066399356 | 0.50888739 | 1 |
| Stx11 | 0.89697397 | -0.15686197 | 0.50891675 | 1 |
| Sprtn | 0.90650703 | -0.141609887 | 0.50901791 | 1 |
| Trpt1 | 1.08182706 | 0.113469888 | 0.50921278 | 1 |
| Slc24a3 | 0.96035871 | -0.058354711 | 0.50934224 | 1 |
| Aurka | 0.92107231 | -0.118613667 | 0.5093507 | 1 |
| Btbd10 | 0.96025914 | -0.058504299 | 0.50945797 | 1 |
| Pfas | 1.04775219 | 0.067297532 | 0.50949337 | 1 |
| Tpm4 | 0.9658556 | -0.050120586 | 0.50959083 | 1 |
| Prpf8 | 0.97013761 | -0.043738696 | 0.50959515 | 1 |
| Fat4 | 0.93841214 | -0.091706415 | 0.50959642 | 1 |
| Commd7 | 1.05814428 | 0.081536361 | 0.5096938 | 1 |
| Nebl | 1.0557884 | 0.07832072 | 0.5098689 | 1 |
| Tbc1d10a | 0.94374141 | -0.083536489 | 0.51011008 | 1 |
| Nol10 | 1.04954892 | 0.069769417 | 0.51020212 | 1 |
| Hspa12a | 0.96902133 | -0.045399673 | 0.51025444 | 1 |
| Cib1 | 0.93666813 | -0.094390117 | 0.51033246 | 1 |
| Sccpdh | 1.03092166 | 0.043934711 | 0.51034623 | 1 |
| Plxna2 | 0.9698415 | -0.044179101 | 0.51034726 | 1 |
| Lrrn1 | 1.03090307 | 0.043908693 | 0.51034848 | 1 |
| Brms1l | 0.96312791 | -0.05420068 | 0.51040754 | 1 |
| Fam69b | 1.03715714 | 0.0526345 | 0.51042164 | 1 |
| Aass | 1.08604718 | 0.119086773 | 0.51060833 | 1 |
| Trpm4 | 1.06141609 | 0.085990328 | 0.51063218 | 1 |
| Rgs5 | 0.93878007 | -0.091140882 | 0.51063612 | 1 |
| Sub1 | 0.9678365 | -0.047164751 | 0.51079984 | 1 |

| | | | | | |
|---|---|---|---|---|---|
| Mcmbp | | 0.94921693 | -0.075190268 | 0.51085366 | 1 |
| Ggta1 | | 1.05414743 | 0.076076657 | 0.51099087 | 1 |
| Cep57 | | 0.94434618 | -0.082612269 | 0.51099272 | 1 |
| Foxj2 | | 1.04923231 | 0.069334142 | 0.5111024 | 1 |
| Snai1 | | 0.87915482 | -0.185810848 | 0.51111638 | 1 |
| Ap3s2 | | 1.03499357 | 0.0496218 | 0.51113336 | 1 |
| Pak2 | | 1.04238978 | 0.059894844 | 0.51126744 | 1 |
| Rap2c | | 0.96686085 | -0.048619818 | 0.51132987 | 1 |
| Arfgap3 | | 1.03775613 | 0.053467448 | 0.51139005 | 1 |
| Drosha | | 0.96531686 | -0.050925519 | 0.51142357 | 1 |
| | 5-Mar | 0.961612 | -0.056473197 | 0.51144585 | 1 |
| Tp53i3 | | 0.93058358 | -0.103792359 | 0.51144932 | 1 |
| Nipa1 | | 1.03755535 | 0.0531883 | 0.5117824 | 1 |
| Topors | | 1.04730177 | 0.066677205 | 0.51187211 | 1 |
| Ppard | | 0.94368066 | -0.08362936 | 0.51190477 | 1 |
| Psmd1 | | 1.03075686 | 0.043704059 | 0.51193706 | 1 |
| Kcnk1 | | 1.03492465 | 0.049525732 | 0.51212324 | 1 |
| Golph3 | | 1.03228188 | 0.045836969 | 0.51214537 | 1 |
| Mmachc | | 1.07094963 | 0.098890626 | 0.51214927 | 1 |
| Ints5 | | 1.0603088 | 0.084484488 | 0.51218638 | 1 |
| Ppef1 | | 1.03138101 | 0.044577383 | 0.51221108 | 1 |
| Gabrb3 | | 1.04475898 | 0.063170157 | 0.51226853 | 1 |
| Rasa2 | | 1.05876008 | 0.082375703 | 0.51236848 | 1 |
| Katnbl1 | | 1.05211266 | 0.073289202 | 0.51244566 | 1 |
| Rbl1 | | 0.87285917 | -0.196179185 | 0.51246826 | 1 |
| Sgsm3 | | 1.04528418 | 0.063895224 | 0.51258892 | 1 |
| Thap1 | | 0.90686312 | -0.141043284 | 0.51260172 | 1 |
| Ap1m1 | | 1.03428093 | 0.048628098 | 0.51265395 | 1 |
| Cbfb | | 1.03052349 | 0.043377387 | 0.51265407 | 1 |
| Igf1r | | 0.93406588 | -0.098403793 | 0.51265801 | 1 |
| Ndufb10 | | 1.03231123 | 0.045877999 | 0.51267466 | 1 |
| Kctd5 | | 0.96192659 | -0.056001291 | 0.51267689 | 1 |
| Khk | | 0.94397948 | -0.083172595 | 0.51270753 | 1 |
| Cfi | | 1.07234089 | 0.100763608 | 0.51271898 | 1 |
| Clptm1l | | 0.96523008 | -0.051055221 | 0.51274881 | 1 |
| RGD735065 | | 0.95932591 | -0.059907068 | 0.51275588 | 1 |
| Tarsl2 | | 1.05516695 | 0.077471277 | 0.51294466 | 1 |
| RGD1307621 | | 0.90175616 | -0.149190724 | 0.51300417 | 1 |
| Ddx47 | | 1.04180331 | 0.05908293 | 0.51317768 | 1 |
| Fbxw8 | | 0.95934615 | -0.059876641 | 0.51332217 | 1 |

| | | | | |
|---|---|---|---|---|
| Col4a4 | 0.85668546 | -0.223162486 | 0.51332702 | 1 |
| RGD1305089 | 1.05850283 | 0.082025132 | 0.51353884 | 1 |
| Ifitm2 | 0.9602582 | -0.058505715 | 0.5136108 | 1 |
| Chchd6 | 1.04556636 | 0.064284636 | 0.513615 | 1 |
| Zmiz2 | 0.96499507 | -0.051406519 | 0.51363063 | 1 |
| Ift43 | 0.92055402 | -0.119425709 | 0.51369105 | 1 |
| Kcnt2 | 0.93579203 | -0.095740151 | 0.51371283 | 1 |
| Kifc1 | 1.15843735 | 0.212180029 | 0.51372808 | 1 |
| Capns1 | 1.03076091 | 0.043709727 | 0.51378367 | 1 |
| Usp25 | 0.96722291 | -0.048079682 | 0.51383679 | 1 |
| Ift27 | 1.06497968 | 0.0908259 | 0.51383778 | 1 |
| Nars | 1.03075558 | 0.043702266 | 0.51405428 | 1 |
| Prpf19 | 1.03034817 | 0.043131922 | 0.51420439 | 1 |
| Pcnp | 1.04355327 | 0.061504252 | 0.51439038 | 1 |
| Fam8a1 | 0.96993565 | -0.044039053 | 0.51442197 | 1 |
| Cygb | 0.94867692 | -0.076011251 | 0.51448154 | 1 |
| Ephx4 | 0.87849856 | -0.186888181 | 0.51449652 | 1 |
| St5 | 1.04493536 | 0.063413694 | 0.51480228 | 1 |
| Gdi1 | 1.03119743 | 0.044320571 | 0.5148067 | 1 |
| Tusc5 | 0.96989442 | -0.04410038 | 0.51500409 | 1 |
| Esco1 | 1.0467806 | 0.065959094 | 0.51506583 | 1 |
| Alg11 | 1.04258411 | 0.060163771 | 0.51508241 | 1 |
| Eif2ak1 | 1.03534055 | 0.050105386 | 0.51530302 | 1 |
| Rita1 | 0.95625731 | -0.064529218 | 0.51545082 | 1 |
| Spast | 0.9622952 | -0.055448565 | 0.51568961 | 1 |
| Slpi | 1.19832235 | 0.26101605 | 0.51572154 | 1 |
| Kcne3 | 1.13822532 | 0.186786183 | 0.51599136 | 1 |
| Kit | 0.95895803 | -0.060460423 | 0.51601743 | 1 |
| Agtr1a | 0.91666912 | -0.125527021 | 0.51623417 | 1 |
| Ccar1 | 0.96665532 | -0.048926533 | 0.51623826 | 1 |
| Camk2n1 | 1.07687528 | 0.106851166 | 0.5164498 | 1 |
| Impdh1 | 0.97056736 | -0.043099748 | 0.51647198 | 1 |
| Nat8l | 0.97067812 | -0.042935125 | 0.51661376 | 1 |
| Arhgap15 | 1.11372622 | 0.155394629 | 0.51663706 | 1 |
| Otc | 1.19784544 | 0.260441762 | 0.51663963 | 1 |
| Surf4 | 1.03175537 | 0.045100948 | 0.51664345 | 1 |
| Snx3 | 0.97073906 | -0.042844545 | 0.51664996 | 1 |
| Mre11a | 0.93636433 | -0.094858112 | 0.51666873 | 1 |
| Mybl2 | 0.81499279 | -0.295140801 | 0.5166927 | 1 |
| RT1-DMb | 1.07393008 | 0.102900067 | 0.51671321 | 1 |

| | | | | |
|---|---|---|---|---|
| Rb1cc1 | 1.03049647 | 0.043339565 | 0.51693147 | 1 |
| Arpp19 | 1.04081691 | 0.057716303 | 0.51710827 | 1 |
| Prrt1 | 1.05578918 | 0.07832179 | 0.51711923 | 1 |
| Acat2 | 0.96411043 | -0.052729691 | 0.51722363 | 1 |
| Nubpl | 0.91866178 | -0.122394279 | 0.51734191 | 1 |
| Large | 0.97055488 | -0.043118309 | 0.51746953 | 1 |
| Rabac1 | 0.9695587 | -0.044599847 | 0.51748848 | 1 |
| Copb2 | 1.03225871 | 0.045804593 | 0.51752251 | 1 |
| Slc5a3 | 0.90314847 | -0.146964928 | 0.5175557 | 1 |
| Pex13 | 0.95120776 | -0.072167616 | 0.517682 | 1 |
| Uqcrfs1 | 1.03227718 | 0.045830403 | 0.51777792 | 1 |
| Kcnmb1 | 1.21581332 | 0.281921729 | 0.51796517 | 1 |
| Wdr61 | 1.0417085 | 0.05895163 | 0.51798084 | 1 |
| Rexo2 | 0.96462425 | -0.05196102 | 0.51799111 | 1 |
| Edf1 | 1.03164095 | 0.044940951 | 0.5180214 | 1 |
| Kif2a | 0.96785262 | -0.047140723 | 0.51813021 | 1 |
| Rp9 | 1.12329991 | 0.167743167 | 0.51814069 | 1 |
| Trim46 | 1.05042787 | 0.070977102 | 0.51821353 | 1 |
| Zdhhc9 | 0.96134836 | -0.056868789 | 0.51829329 | 1 |
| Zmym3 | 0.96634817 | -0.04938501 | 0.51831318 | 1 |
| Gtpbp6 | 0.92574695 | -0.1113102 | 0.51835503 | 1 |
| RGD1565784 | 0.95061303 | -0.073069913 | 0.51837028 | 1 |
| Cys1 | 0.93771769 | -0.092774444 | 0.51860631 | 1 |
| Pdgfb | 1.102076 | 0.140223712 | 0.51864246 | 1 |
| Gde1 | 1.03232354 | 0.045895193 | 0.51867053 | 1 |
| Fam101a | 1.13430742 | 0.18181169 | 0.51884049 | 1 |
| Cct6a | 0.97042251 | -0.043315072 | 0.51889571 | 1 |
| Csnk1e | 0.95120058 | -0.072178508 | 0.51895413 | 1 |
| Lhfpl2 | 0.96538814 | -0.050818997 | 0.51898176 | 1 |
| Cd300le | 1.09835831 | 0.13534877 | 0.51908751 | 1 |
| Acsl3 | 0.96877303 | -0.045769388 | 0.51926472 | 1 |
| Entpd7 | 1.11037114 | 0.15104197 | 0.51931937 | 1 |
| Xpo7 | 1.03251815 | 0.046167146 | 0.51933878 | 1 |
| Lcat | 1.05728802 | 0.080368434 | 0.51935966 | 1 |
| Banf1 | 1.04143059 | 0.05856669 | 0.51941396 | 1 |
| Apbb1 | 0.97095718 | -0.042520427 | 0.5195312 | 1 |
| Eif4e | 1.03319112 | 0.047107156 | 0.51954977 | 1 |
| Bst1 | 0.89955726 | -0.152712974 | 0.51958982 | 1 |
| Jak3 | 1.08709904 | 0.120483385 | 0.51959261 | 1 |
| Dynlrb1 | 1.03000215 | 0.042647351 | 0.51971167 | 1 |

| | | | | | |
|---|---|---|---|---|---|
| Tmem53 | | 1.09149934 | 0.126311261 | 0.5197172 | 1 |
| | 3-Sep | 1.05306532 | 0.074594933 | 0.51995294 | 1 |
| Med1 | | 1.044221 | 0.062427084 | 0.51995895 | 1 |
| Cops7a | | 1.03272595 | 0.04645746 | 0.52003581 | 1 |
| Gls | | 0.97065022 | -0.042976596 | 0.52003981 | 1 |
| Stxbp3 | | 1.04269768 | 0.060320917 | 0.52008736 | 1 |
| Caskin2 | | 0.96240562 | -0.055283022 | 0.52015735 | 1 |
| Aars2 | | 1.08127408 | 0.112732255 | 0.52020926 | 1 |
| Mapk3 | | 0.97048703 | -0.043219159 | 0.52034624 | 1 |
| Efemp1 | | 1.03148243 | 0.044719244 | 0.52036991 | 1 |
| Stmn1 | | 0.97103205 | -0.042409183 | 0.52045563 | 1 |
| Srrm3 | | 0.94999068 | -0.074014739 | 0.52046867 | 1 |
| Lrrc4 | | 1.06387032 | 0.089322308 | 0.52056773 | 1 |
| Laptm5 | | 1.04864487 | 0.068526178 | 0.52067958 | 1 |
| Rnf2 | | 0.94734855 | -0.078032773 | 0.52068883 | 1 |
| Zfp672 | | 1.05876946 | 0.082388486 | 0.52100209 | 1 |
| Adra1b | | 1.14649476 | 0.197229761 | 0.52129614 | 1 |
| Ash2l | | 1.04004911 | 0.056651654 | 0.52138124 | 1 |
| Ptgr2 | | 0.95283336 | -0.069704172 | 0.52140132 | 1 |
| Mier1 | | 1.03749677 | 0.053106845 | 0.52153405 | 1 |
| Cand1 | | 0.97110137 | -0.04230619 | 0.52162352 | 1 |
| Ehhadh | | 1.09814232 | 0.135065045 | 0.52168926 | 1 |
| Dcaf5 | | 0.96563344 | -0.050452457 | 0.52179635 | 1 |
| Pak7 | | 1.11493981 | 0.156965831 | 0.52179856 | 1 |
| Ece2 | | 1.03732192 | 0.052863687 | 0.52185382 | 1 |
| Il31ra | | 1.04549758 | 0.064189728 | 0.52203155 | 1 |
| Myl6b | | 1.03555125 | 0.050398955 | 0.52212555 | 1 |
| Slirp | | 1.05043108 | 0.070981504 | 0.52215229 | 1 |
| Hbs1l | | 1.03019578 | 0.042918535 | 0.52229311 | 1 |
| Tspan15 | | 0.95762168 | -0.062472287 | 0.52266048 | 1 |
| Cacna1i | | 0.90001453 | -0.151979806 | 0.52266702 | 1 |
| Ubac2 | | 0.95566411 | -0.065424456 | 0.52280098 | 1 |
| Ttc30a | | 1.13754165 | 0.18591937 | 0.52287396 | 1 |
| Pcdhga5 | | 0.9056365 | -0.14299599 | 0.52287638 | 1 |
| Mien1 | | 1.03844497 | 0.054424766 | 0.52290481 | 1 |
| Arl6ip1 | | 1.02998886 | 0.042628731 | 0.52293582 | 1 |
| Kcnj14 | | 0.90905449 | -0.13756132 | 0.5230007 | 1 |
| Nup98 | | 0.96516533 | -0.051152007 | 0.52320101 | 1 |
| Pop5 | | 1.08055113 | 0.111767341 | 0.52320993 | 1 |
| Ptpn7 | | 1.19443562 | 0.256329091 | 0.52321371 | 1 |

| | | | | |
|---|---|---|---|---|
| Arhgap5 | 0.96531426 | -0.050929406 | 0.52324398 | 1 |
| Plce1 | 0.96237463 | -0.055329486 | 0.52333703 | 1 |
| Rnf6 | 0.96725792 | -0.048027458 | 0.52335691 | 1 |
| Fer | 1.06804568 | 0.094973347 | 0.52339569 | 1 |
| Cdyl2 | 0.91884921 | -0.122099964 | 0.52339926 | 1 |
| Mfsd13a | 0.936994 | -0.093888283 | 0.52340099 | 1 |
| Dnaja2 | 1.03012624 | 0.04282115 | 0.52340596 | 1 |
| Zfp280d | 1.0411158 | 0.058130543 | 0.52343392 | 1 |
| Alkbh4 | 0.88785912 | -0.171597325 | 0.52344962 | 1 |
| Slc7a5 | 0.96051085 | -0.05812619 | 0.52374362 | 1 |
| Kirrel | 0.93815957 | -0.092094764 | 0.52380381 | 1 |
| Sema7a | 0.97038553 | -0.043370059 | 0.52397127 | 1 |
| Acot3 | 0.92578423 | -0.111252104 | 0.52408964 | 1 |
| Adamts10 | 0.95934009 | -0.059885751 | 0.52411034 | 1 |
| Lgi3 | 1.02989388 | 0.042495688 | 0.52415172 | 1 |
| Gpr135 | 0.86686498 | -0.206120794 | 0.52415331 | 1 |
| Kctd15 | 1.0522568 | 0.07348684 | 0.52427355 | 1 |
| Acacb | 1.07328979 | 0.102039659 | 0.52443632 | 1 |
| Apeh | 1.03702223 | 0.052446827 | 0.52450048 | 1 |
| Mmp17 | 0.9329587 | -0.100114877 | 0.52461598 | 1 |
| Cmpk1 | 1.03027465 | 0.043028982 | 0.52471347 | 1 |
| RGD1563349 | 0.97003605 | -0.043889734 | 0.52472364 | 1 |
| Klhl26 | 0.94614255 | -0.079870538 | 0.52476226 | 1 |
| S1pr3 | 1.03105444 | 0.044120511 | 0.52480915 | 1 |
| Ap3b1 | 0.964676 | -0.051883614 | 0.52482073 | 1 |
| Itpr1 | 1.04868362 | 0.06857949 | 0.5248557 | 1 |
| Hps5 | 1.04472537 | 0.063123751 | 0.52497176 | 1 |
| Slc39a3 | 0.93357529 | -0.099161714 | 0.52506035 | 1 |
| Epsti1 | 1.12669664 | 0.172099127 | 0.52515907 | 1 |
| Cept1 | 0.96603556 | -0.049851801 | 0.52522394 | 1 |
| Stard13 | 0.96168885 | -0.056357909 | 0.52557961 | 1 |
| Pigc | 1.04715999 | 0.06648188 | 0.52561752 | 1 |
| Vstm2b | 1.0620167 | 0.086806447 | 0.52575469 | 1 |
| Phf2 | 0.95199356 | -0.070976279 | 0.52576605 | 1 |
| Pdik1l | 0.9378748 | -0.092532746 | 0.52585568 | 1 |
| Tmed4 | 0.96400429 | -0.05288853 | 0.52587132 | 1 |
| LOC100910827 | 0.97084998 | -0.042679718 | 0.52602928 | 1 |
| Lias | 1.05196195 | 0.073082516 | 0.52613679 | 1 |
| Dhx37 | 0.94011158 | -0.089096101 | 0.52633834 | 1 |
| Cd46 | 1.12964899 | 0.175874558 | 0.52646463 | 1 |

| | | | | |
|---|---|---|---|---|
| Ppp1r8 | 1.04764753 | 0.067153419 | 0.52650165 | 1 |
| Gli1 | 1.05258836 | 0.073941349 | 0.52668455 | 1 |
| Slc35a2 | 0.95428934 | -0.06750134 | 0.52674456 | 1 |
| Bfsp1 | 0.89210146 | -0.164720293 | 0.52677649 | 1 |
| Pttg1ip | 1.03091704 | 0.04392824 | 0.52679854 | 1 |
| Lgalsl | 0.96889546 | -0.045587083 | 0.52687899 | 1 |
| Lysmd2 | 1.03885811 | 0.054998623 | 0.52688672 | 1 |
| Tbc1d14 | 1.03269152 | 0.046409366 | 0.52696115 | 1 |
| Psma4 | 1.03113224 | 0.044229373 | 0.52705918 | 1 |
| Kif13b | 1.03914161 | 0.055392266 | 0.52710409 | 1 |
| Wdr54 | 1.04765738 | 0.06716698 | 0.52717692 | 1 |
| Nuak2 | 1.1348997 | 0.1825648 | 0.52747084 | 1 |
| Ctsk | 0.9582714 | -0.061493781 | 0.52770935 | 1 |
| Zfp868 | 1.07474127 | 0.103989392 | 0.52773357 | 1 |
| Tmem63c | 0.97001265 | -0.043924526 | 0.52774138 | 1 |
| RGD1310429 | 1.04278619 | 0.06044338 | 0.52794216 | 1 |
| Arfip1 | 1.05011716 | 0.070550294 | 0.52799365 | 1 |
| Lmtk2 | 0.96131964 | -0.056911886 | 0.52845597 | 1 |
| Ptprc | 0.95507312 | -0.066316898 | 0.52852551 | 1 |
| Gng5 | 0.95419807 | -0.067639325 | 0.52853841 | 1 |
| Yeats2 | 1.05311946 | 0.074669092 | 0.52857157 | 1 |
| Pcdha13 | 1.06622198 | 0.092507834 | 0.5285854 | 1 |
| Egfem1 | 1.11338883 | 0.154957509 | 0.52865661 | 1 |
| Palmd | 1.0412493 | 0.058315531 | 0.52878509 | 1 |
| Htr2c | 1.11322559 | 0.154745982 | 0.52887078 | 1 |
| Sbsn | 1.11660379 | 0.159117356 | 0.52898137 | 1 |
| Cdk6 | 1.11314951 | 0.154647381 | 0.5290617 | 1 |
| Armc1 | 1.03569324 | 0.050596753 | 0.52907161 | 1 |
| Txnrd2 | 1.0714298 | 0.099537323 | 0.52911152 | 1 |
| Slc33a1 | 0.95613739 | -0.064710162 | 0.52920411 | 1 |
| Sugp2 | 0.96683267 | -0.048661875 | 0.52922415 | 1 |
| Slc37a2 | 1.11195389 | 0.153096959 | 0.52924821 | 1 |
| Wisp2 | 1.05219109 | 0.073396734 | 0.52931185 | 1 |
| Apmap | 0.96123315 | -0.057041691 | 0.52931839 | 1 |
| Carhsp1 | 0.96475372 | -0.051767389 | 0.52947871 | 1 |
| Aldh1b1 | 0.89691301 | -0.156960023 | 0.52954778 | 1 |
| Nmrk1 | 0.92901899 | -0.10622001 | 0.52957026 | 1 |
| Adnp2 | 0.95081145 | -0.072768821 | 0.52957878 | 1 |
| Col4a3 | 0.89966425 | -0.152541405 | 0.52961076 | 1 |
| Ttc3 | 1.03050964 | 0.043357998 | 0.52963295 | 1 |

| | | | | |
|---|---|---|---|---|
| Tex10 | 1.04576237 | 0.06455506 | 0.5296628 | 1 |
| Lrfn5 | 0.96403896 | -0.052836649 | 0.52972039 | 1 |
| Pfdn1 | 0.96898537 | -0.045453215 | 0.52985637 | 1 |
| Tcta | 1.03722171 | 0.052724302 | 0.52990541 | 1 |
| Dgcr2 | 0.96057879 | -0.058024136 | 0.53017403 | 1 |
| Trib2 | 0.9678134 | -0.047199184 | 0.53025302 | 1 |
| Sec11a | 0.95923757 | -0.060039931 | 0.53026537 | 1 |
| Cdkn2aipnl | 1.04227792 | 0.059740017 | 0.53031333 | 1 |
| Irf1 | 1.06434346 | 0.089963785 | 0.5303145 | 1 |
| RGD1306063 | 0.95811202 | -0.061733758 | 0.53035436 | 1 |
| Suclg1 | 1.03001362 | 0.042663416 | 0.53042484 | 1 |
| Oser1 | 1.04664433 | 0.065771269 | 0.53043195 | 1 |
| Vezf1 | 1.03893366 | 0.05510354 | 0.53050703 | 1 |
| Vgll4 | 0.95274404 | -0.069839413 | 0.53067268 | 1 |
| Ntsr2 | 1.11092381 | 0.151759882 | 0.53067943 | 1 |
| Ehd3 | 0.97174525 | -0.041349947 | 0.531166 | 1 |
| Fgfrl1 | 0.8973886 | -0.156195246 | 0.53126185 | 1 |
| Pcdha3 | 0.93833561 | -0.091824083 | 0.53132125 | 1 |
| Rab3ip | 0.95770183 | -0.062351535 | 0.53134 | 1 |
| Ide | 0.96589136 | -0.050067163 | 0.5316389 | 1 |
| Trip13 | 0.84642492 | -0.240545995 | 0.531688 | 1 |
| Tbce | 1.03310217 | 0.046982941 | 0.53176336 | 1 |
| Kiaa0895l | 0.96136755 | -0.056839982 | 0.53182758 | 1 |
| Gtf2h1 | 0.96142823 | -0.056748925 | 0.53203545 | 1 |
| Ppp4r2 | 0.96710572 | -0.048254492 | 0.5320518 | 1 |
| Ucp2 | 0.97077565 | -0.042790175 | 0.53213508 | 1 |
| Kcnab1 | 1.0290396 | 0.041298509 | 0.53224379 | 1 |
| Trmt12 | 1.07516989 | 0.104564646 | 0.53232417 | 1 |
| B3galt4 | 0.8865167 | -0.173780292 | 0.53240346 | 1 |
| Ccdc102a | 0.94409677 | -0.082993344 | 0.53250192 | 1 |
| Sec14l5 | 0.88287789 | -0.179714189 | 0.53260718 | 1 |
| Tmem67 | 1.05194886 | 0.073064572 | 0.53277027 | 1 |
| Bach1 | 0.9488609 | -0.075731483 | 0.53282429 | 1 |
| B3gat2 | 0.93892323 | -0.090920899 | 0.53289796 | 1 |
| Lrrk2 | 1.03268652 | 0.046402383 | 0.53295864 | 1 |
| Mpc2 | 1.04895026 | 0.068946273 | 0.5331193 | 1 |
| Tsga10 | 0.92468393 | -0.112967776 | 0.5332205 | 1 |
| Klhl36 | 1.0995123 | 0.136863746 | 0.53341003 | 1 |
| Ogfod3 | 1.0870261 | 0.120386576 | 0.53343425 | 1 |
| LOC654482 | 1.09952207 | 0.136876568 | 0.53372074 | 1 |

| | | | | |
|---|---|---|---|---|
| Cma1 | 1.201132 | 0.264394711 | 0.53374993 | 1 |
| Snurf | 0.97181843 | -0.041241309 | 0.53397077 | 1 |
| Serinc2 | 0.94191847 | -0.086325912 | 0.53397341 | 1 |
| Cxxc5 | 0.96056475 | -0.058045221 | 0.53397475 | 1 |
| Vps4b | 1.03000341 | 0.042649113 | 0.53398881 | 1 |
| Sspn | 1.04445246 | 0.062746823 | 0.53400721 | 1 |
| Kbtbd3 | 1.03778654 | 0.053509724 | 0.53415541 | 1 |
| Gstt3 | 0.9548846 | -0.066601711 | 0.534318 | 1 |
| G4 | 0.93025408 | -0.104303275 | 0.53436642 | 1 |
| Cacul1 | 0.96852062 | -0.046145331 | 0.53440315 | 1 |
| Rbm42 | 0.96009874 | -0.058745312 | 0.53457881 | 1 |
| Dpagt1 | 1.03546658 | 0.050280997 | 0.53501259 | 1 |
| Pias4 | 0.94940077 | -0.074910868 | 0.53507903 | 1 |
| Smg9 | 0.93967217 | -0.08977057 | 0.53511579 | 1 |
| Prkar2b | 1.02995914 | 0.042587107 | 0.53512426 | 1 |
| Fbxo25 | 0.9589801 | -0.060427212 | 0.53544877 | 1 |
| Cask | 0.96239376 | -0.055300809 | 0.53568002 | 1 |
| Fxr2 | 0.97116485 | -0.042211892 | 0.53583353 | 1 |
| Trove2 | 1.04397322 | 0.062084705 | 0.53583649 | 1 |
| Rpl22 | 1.04222213 | 0.059662788 | 0.53587506 | 1 |
| Ankrd50 | 1.03836014 | 0.054306902 | 0.53591347 | 1 |
| Slc9a8 | 0.95110276 | -0.072326874 | 0.53593094 | 1 |
| Cln3 | 1.0610131 | 0.085442467 | 0.53595637 | 1 |
| Mcm8 | 1.08317974 | 0.115272662 | 0.53602061 | 1 |
| Grk4 | 0.89438069 | -0.16103906 | 0.53619432 | 1 |
| Nudt19 | 0.94292747 | -0.08478129 | 0.5362095 | 1 |
| Ankrd13a | 0.96704096 | -0.048351103 | 0.53632954 | 1 |
| Pcyt2 | 0.96264824 | -0.054919369 | 0.53632978 | 1 |
| Elfn1 | 0.94679329 | -0.078878611 | 0.53640911 | 1 |
| Pofut1 | 0.91511395 | -0.127976697 | 0.53647365 | 1 |
| Slc1a5 | 1.09867277 | 0.135761754 | 0.53648375 | 1 |
| Tmem18 | 0.93936188 | -0.090247048 | 0.5366439 | 1 |
| Rraga | 1.02966499 | 0.042175022 | 0.53677778 | 1 |
| Llgl2 | 1.13337984 | 0.180631448 | 0.53689705 | 1 |
| Trhde | 1.11050433 | 0.151215014 | 0.53704843 | 1 |
| Hcfc2 | 0.94470605 | -0.082062604 | 0.53705709 | 1 |
| Prkch | 1.07261148 | 0.101127602 | 0.53719175 | 1 |
| Msra | 0.93452769 | -0.097690684 | 0.53735671 | 1 |
| Creld1 | 0.96911367 | -0.045262198 | 0.53737124 | 1 |
| Spns1 | 0.95923556 | -0.060042949 | 0.5374973 | 1 |

| | | | | |
|---|---|---|---|---|
| Dhrs7b | 1.04858201 | 0.068439695 | 0.53770421 | 1 |
| Cxcl16 | 1.05364809 | 0.075393101 | 0.53778199 | 1 |
| Ets1 | 1.04054136 | 0.057334307 | 0.53782348 | 1 |
| Chid1 | 0.95444923 | -0.067259643 | 0.53786024 | 1 |
| Ccdc12 | 0.94139316 | -0.087130732 | 0.53790722 | 1 |
| Tp53rk | 0.91258605 | -0.131967489 | 0.53793489 | 1 |
| Src | 0.95089586 | -0.072640744 | 0.53795122 | 1 |
| Dmd | 0.96586709 | -0.050103421 | 0.5379848 | 1 |
| Tmem63a | 1.08314004 | 0.115219781 | 0.53805635 | 1 |
| Snap29 | 1.03395505 | 0.048173473 | 0.53814281 | 1 |
| Kdm5b | 0.96239648 | -0.05529673 | 0.53820504 | 1 |
| Zic5 | 1.2280799 | 0.296404422 | 0.53822479 | 1 |
| Pafah1b3 | 1.05686203 | 0.079787056 | 0.53825348 | 1 |
| Cep41 | 1.09116425 | 0.125868279 | 0.53829686 | 1 |
| Cd2ap | 1.0335646 | 0.047628557 | 0.5385387 | 1 |
| Srebf2 | 0.9710285 | -0.042414456 | 0.53888841 | 1 |
| Slc20a2 | 1.04271489 | 0.060344738 | 0.5389638 | 1 |
| Acot9 | 1.03416715 | 0.048469382 | 0.53901887 | 1 |
| Arhgdia | 0.97168356 | -0.04144153 | 0.53924025 | 1 |
| Sh3tc1 | 1.12279315 | 0.167092173 | 0.53927068 | 1 |
| Rlim | 1.04282614 | 0.060498657 | 0.53938929 | 1 |
| Ebna1bp2 | 1.04134526 | 0.058448474 | 0.53945897 | 1 |
| Ttc13 | 0.95724347 | -0.063042187 | 0.53957391 | 1 |
| Lmo2 | 1.06905261 | 0.096332853 | 0.54002701 | 1 |
| Tmod2 | 0.96997553 | -0.043979739 | 0.54005777 | 1 |
| Ddx3y | 1.08372978 | 0.116005079 | 0.54007935 | 1 |
| RGD1560010 | 1.07030388 | 0.098020467 | 0.54014896 | 1 |
| Zfhx4 | 0.94207855 | -0.086080736 | 0.54015496 | 1 |
| Extl1 | 1.14496266 | 0.195300554 | 0.5402699 | 1 |
| Cecr2 | 1.09615366 | 0.132450051 | 0.54028374 | 1 |
| Gfm1 | 0.96269375 | -0.054851171 | 0.54042227 | 1 |
| Osr2 | 1.13310312 | 0.180279161 | 0.54050835 | 1 |
| Spag7 | 0.95955313 | -0.059565412 | 0.54056365 | 1 |
| Zfp386 | 1.05991322 | 0.083946154 | 0.54085922 | 1 |
| Lats1 | 1.03941238 | 0.055768148 | 0.54092662 | 1 |
| Tle4 | 0.96428378 | -0.052470314 | 0.54093453 | 1 |
| Hbb | 1.02999875 | 0.042642593 | 0.54101326 | 1 |
| Ctif | 0.96188666 | -0.056061186 | 0.54111354 | 1 |
| Hacd1 | 0.88353256 | -0.178644796 | 0.54113878 | 1 |
| Magee2 | 0.90152177 | -0.149565766 | 0.54117536 | 1 |

| | | | | |
|---|---|---|---|---|
| Sesn2 | 0.92860662 | -0.10686053 | 0.54118348 | 1 |
| Epdr1 | 1.02847355 | 0.040504687 | 0.54119608 | 1 |
| Cyth1 | 1.05598785 | 0.078593232 | 0.54124371 | 1 |
| Mycn | 1.1241974 | 0.168895388 | 0.54127695 | 1 |
| Zfp319 | 0.92455479 | -0.113169274 | 0.54130211 | 1 |
| Pdcd5 | 1.03683738 | 0.05218964 | 0.54146779 | 1 |
| Tnrc18 | 1.03959711 | 0.056024529 | 0.54149616 | 1 |
| Hdgfrp3 | 1.03702358 | 0.052448697 | 0.54150684 | 1 |
| Cdc123 | 1.03515902 | 0.049852404 | 0.54158443 | 1 |
| Prcc | 0.95950179 | -0.059642604 | 0.54160339 | 1 |
| Zadh2 | 1.05060459 | 0.071219794 | 0.54167839 | 1 |
| Tm4sf4 | 1.17526942 | 0.232991522 | 0.54185573 | 1 |
| Erap1 | 0.9659115 | -0.050037091 | 0.54189355 | 1 |
| Kif3a | 1.02841073 | 0.040416563 | 0.54212035 | 1 |
| Bbs2 | 0.96047453 | -0.058180744 | 0.5421305 | 1 |
| Mms22l | 1.15979443 | 0.21386911 | 0.54215011 | 1 |
| Ggct | 1.08202044 | 0.113727755 | 0.54249196 | 1 |
| Vamp7 | 1.04710974 | 0.066412645 | 0.5425024 | 1 |
| Nrarp | 1.04865173 | 0.068535622 | 0.54253674 | 1 |
| Lrba | 0.95737283 | -0.062847224 | 0.5425518 | 1 |
| Znrd1 | 1.07696289 | 0.106968543 | 0.5425596 | 1 |
| Ppp2r2c | 0.96771927 | -0.04733951 | 0.54257266 | 1 |
| LOC100233176 | 0.95543965 | -0.06576334 | 0.54259001 | 1 |
| Tmed2 | 0.97162707 | -0.041525409 | 0.54269979 | 1 |
| Pnpla2 | 0.96172997 | -0.056296219 | 0.54280418 | 1 |
| App | 0.97162026 | -0.041535524 | 0.5428386 | 1 |
| Hip1r | 0.96421571 | -0.052572161 | 0.54289699 | 1 |
| Prpf40b | 0.96071815 | -0.05781485 | 0.54299531 | 1 |
| Morc2 | 0.95855716 | -0.061063636 | 0.54301793 | 1 |
| Stub1 | 1.02962116 | 0.042113608 | 0.54305131 | 1 |
| Hmgb1 | 0.96799457 | -0.046929142 | 0.54306877 | 1 |
| Sumf1 | 0.96383053 | -0.053148593 | 0.54311304 | 1 |
| Gpld1 | 0.92777915 | -0.108146664 | 0.54325236 | 1 |
| Plxnd1 | 1.03249786 | 0.04613879 | 0.54338005 | 1 |
| Ssh3 | 1.04796963 | 0.067596902 | 0.54347489 | 1 |
| Idh3g | 1.02952101 | 0.041973271 | 0.54353169 | 1 |
| Hnrnpa2b1 | 1.02872721 | 0.040860468 | 0.54364909 | 1 |
| Coq7 | 0.93130942 | -0.102667531 | 0.54365548 | 1 |
| Ppp1r16b | 0.93108985 | -0.103007706 | 0.54381374 | 1 |
| Ksr1 | 0.94691943 | -0.078686417 | 0.54400534 | 1 |

| | | | | |
|---|---|---|---|---|
| Stx3 | 0.9039386 | -0.145703322 | 0.54411002 | 1 |
| Ophn1 | 1.0395545 | 0.055965394 | 0.5441196 | 1 |
| Gxylt1 | 0.96153867 | -0.056583214 | 0.5442544 | 1 |
| Fgf10 | 1.13107642 | 0.177696406 | 0.54440451 | 1 |
| Dock8 | 1.0644444 | 0.0901006 | 0.54441535 | 1 |
| Kcnh7 | 0.96540659 | -0.050791416 | 0.54452612 | 1 |
| Pnpo | 1.04324274 | 0.061074882 | 0.54456489 | 1 |
| Atp2b1 | 0.97264534 | -0.040014253 | 0.54457052 | 1 |
| Clcnkb | 1.13340096 | 0.180658333 | 0.54465454 | 1 |
| Rbm28 | 0.9526868 | -0.069926095 | 0.54470631 | 1 |
| Adprh | 0.95789391 | -0.062062212 | 0.54479916 | 1 |
| Bin1 | 0.97134043 | -0.04195108 | 0.54483427 | 1 |
| Cast | 1.02820105 | 0.04012239 | 0.54484164 | 1 |
| Ewsr1 | 0.96966283 | -0.044444906 | 0.54497462 | 1 |
| Wnk4 | 1.05298653 | 0.074486983 | 0.54511829 | 1 |
| Rhpn1 | 0.96216726 | -0.055640385 | 0.54519482 | 1 |
| Tomm40 | 0.95823015 | -0.06155589 | 0.54533727 | 1 |
| Hint1 | 1.02893261 | 0.041148491 | 0.5454341 | 1 |
| Rheb | 1.02911035 | 0.041397682 | 0.54552689 | 1 |
| Irs3 | 1.11764074 | 0.160456516 | 0.54559708 | 1 |
| Usp46 | 0.96966694 | -0.044438792 | 0.54561579 | 1 |
| Kpna1 | 0.96275306 | -0.054762299 | 0.54566214 | 1 |
| Slc25a16 | 0.9621329 | -0.055691909 | 0.54566614 | 1 |
| Spcs2 | 0.96916403 | -0.045187234 | 0.54572098 | 1 |
| Trim26 | 0.95943266 | -0.059746546 | 0.54578819 | 1 |
| Tbc1d10c | 0.87265606 | -0.19651494 | 0.54588483 | 1 |
| Trpc6 | 0.92194581 | -0.117246142 | 0.54592174 | 1 |
| Ybx1 | 0.96196372 | -0.055945604 | 0.54603135 | 1 |
| Mob2 | 0.95956207 | -0.059551967 | 0.54616714 | 1 |
| Casr | 0.94630441 | -0.079623746 | 0.54617138 | 1 |
| Crim1 | 0.94959235 | -0.074619783 | 0.54642081 | 1 |
| Taz | 0.95293673 | -0.069547668 | 0.54643157 | 1 |
| Mical2 | 0.95801925 | -0.061873446 | 0.54652922 | 1 |
| Akirin2 | 1.03676435 | 0.052088018 | 0.54658182 | 1 |
| Fancb | 1.11276298 | 0.154146326 | 0.54664035 | 1 |
| Pdrg1 | 1.0355695 | 0.050424376 | 0.54670077 | 1 |
| Nos1ap | 0.923757 | -0.114414696 | 0.54672498 | 1 |
| Uhrf2 | 0.96711306 | -0.048243541 | 0.54678732 | 1 |
| Cep112 | 0.90670578 | -0.141293621 | 0.54681732 | 1 |
| Nt5c | 1.14868134 | 0.199978634 | 0.54710572 | 1 |

| | | | | |
|---|---|---|---|---|
| Slc34a2 | 0.88446394 | -0.177124766 | 0.54720781 | 1 |
| Scgn | 1.24900507 | 0.320779332 | 0.54744834 | 1 |
| Sqle | 0.97249785 | -0.040233038 | 0.54759966 | 1 |
| Dpp3 | 0.96864732 | -0.045956614 | 0.54766698 | 1 |
| Zmynd11 | 1.03096114 | 0.04398996 | 0.54770154 | 1 |
| Cacna1d | 1.05911584 | 0.08286039 | 0.54779586 | 1 |
| RGD1304567 | 1.05968236 | 0.083631881 | 0.54785082 | 1 |
| Lrrfip2 | 0.96087133 | -0.057584843 | 0.5479115 | 1 |
| Tti1 | 1.04584499 | 0.064669042 | 0.5479359 | 1 |
| Ptp4a1 | 0.97010612 | -0.043785526 | 0.5479472 | 1 |
| Zfp275 | 0.95069481 | -0.072945814 | 0.54822405 | 1 |
| Tmem17 | 1.06844915 | 0.095518252 | 0.54827005 | 1 |
| Ppp1r3e | 0.88972548 | -0.168567833 | 0.54831383 | 1 |
| Spire2 | 0.96462984 | -0.051952661 | 0.54838612 | 1 |
| Fundc2 | 1.04597345 | 0.064846233 | 0.54843932 | 1 |
| S100a3 | 1.11592767 | 0.15824352 | 0.54844703 | 1 |
| Ptbp2 | 1.03352934 | 0.047579343 | 0.5485348 | 1 |
| Psmd6 | 1.02783804 | 0.039612953 | 0.54869015 | 1 |
| Kansl2 | 0.96852785 | -0.046134568 | 0.54909709 | 1 |
| Rrm2b | 0.94814233 | -0.076824444 | 0.54914796 | 1 |
| Fdft1 | 0.97298101 | -0.039516443 | 0.54941083 | 1 |
| Mettl6 | 1.06454898 | 0.090242328 | 0.549445 | 1 |
| Pln | 1.11192718 | 0.153062314 | 0.54947571 | 1 |
| Spdya | 1.13552195 | 0.183355596 | 0.54950637 | 1 |
| Fam160b2 | 0.96468665 | -0.0518677 | 0.54968948 | 1 |
| Polr3e | 0.9554139 | -0.065802226 | 0.54978234 | 1 |
| Rabif | 1.04827667 | 0.068019531 | 0.55007151 | 1 |
| Tmem200c | 0.93846304 | -0.091628166 | 0.55026068 | 1 |
| Fiz1 | 1.05543855 | 0.077842584 | 0.55030157 | 1 |
| Clspn | 0.82025649 | -0.285852985 | 0.55030497 | 1 |
| Mtch2 | 1.02849352 | 0.040532697 | 0.55034703 | 1 |
| Mast2 | 1.03195848 | 0.045384928 | 0.55035149 | 1 |
| Knop1 | 0.92750594 | -0.108571574 | 0.55053271 | 1 |
| Tnni2 | 0.86481591 | -0.209535038 | 0.55057871 | 1 |
| Hspbap1 | 1.08760819 | 0.121158923 | 0.55059719 | 1 |
| Atp6v0a1 | 0.97292518 | -0.039599226 | 0.55063727 | 1 |
| Diras2 | 1.06934346 | 0.096725299 | 0.55065171 | 1 |
| Stk25 | 1.02903599 | 0.04129344 | 0.55077896 | 1 |
| Tmbim4 | 1.0427444 | 0.060385559 | 0.55080586 | 1 |
| Chrnb4 | 0.94338649 | -0.084079156 | 0.55084632 | 1 |

| | | | | |
|---|---|---|---|---|
| Esam | 0.94186346 | -0.086410159 | 0.55094328 | 1 |
| Hspa4l | 0.97266836 | -0.039980103 | 0.5509821 | 1 |
| Itgb8 | 1.03946349 | 0.055839083 | 0.55100593 | 1 |
| B3galt6 | 0.93317002 | -0.09978813 | 0.55102106 | 1 |
| Nr2c2ap | 0.9500402 | -0.073939541 | 0.55115186 | 1 |
| Spidr | 1.09024017 | 0.124645986 | 0.55122336 | 1 |
| Ap2s1 | 1.02981822 | 0.042389696 | 0.5513996 | 1 |
| Hagh | 1.03140873 | 0.044616167 | 0.55145629 | 1 |
| Slc35f2 | 0.94107289 | -0.087621627 | 0.55169575 | 1 |
| Chrm2 | 1.05451171 | 0.076575118 | 0.55174453 | 1 |
| P4ha2 | 0.95439579 | -0.067340413 | 0.55175312 | 1 |
| Il13ra1 | 1.03638577 | 0.051561111 | 0.55176533 | 1 |
| Pgrmc2 | 1.03389087 | 0.048083915 | 0.55177564 | 1 |
| Gtf2e2 | 1.04589855 | 0.064742917 | 0.55179477 | 1 |
| Cpne8 | 0.96264133 | -0.054929729 | 0.55193886 | 1 |
| Tubg1 | 0.96601894 | -0.049876619 | 0.55203006 | 1 |
| Xylt2 | 0.9640792 | -0.05277643 | 0.55203637 | 1 |
| Taf6 | 0.9556631 | -0.065425987 | 0.55218468 | 1 |
| Irf4 | 0.87456479 | -0.193362828 | 0.55225599 | 1 |
| Tmem222 | 1.0452324 | 0.063823747 | 0.55230855 | 1 |
| Phrf1 | 0.96400242 | -0.052891331 | 0.5524825 | 1 |
| Ndufaf5 | 1.04616092 | 0.065104787 | 0.55254 | 1 |
| Ppp1r16a | 0.9518316 | -0.071221742 | 0.55254763 | 1 |
| Ltv1 | 1.04599635 | 0.064877811 | 0.55271017 | 1 |
| Pigu | 0.95582648 | -0.065179364 | 0.55294367 | 1 |
| RGD1311188 | 1.0915598 | 0.126391164 | 0.55304097 | 1 |
| Stk36 | 0.92357787 | -0.114694486 | 0.55307217 | 1 |
| Gtpbp2 | 0.96050185 | -0.058139695 | 0.55320205 | 1 |
| Plscr4 | 1.04729833 | 0.066672457 | 0.55320943 | 1 |
| Dscam | 0.93611292 | -0.095245523 | 0.55325155 | 1 |
| Fam172a | 1.03566961 | 0.050563842 | 0.55346345 | 1 |
| Card9 | 0.88073586 | -0.183218693 | 0.55360273 | 1 |
| Oxsr1 | 0.97104545 | -0.042389271 | 0.55361983 | 1 |
| Kbtbd7 | 0.95577197 | -0.06526163 | 0.55373598 | 1 |
| Rbmxrtl | 0.96084809 | -0.057619737 | 0.55381213 | 1 |
| Arnt | 0.95028046 | -0.07357473 | 0.55382942 | 1 |
| Tmem37 | 1.172217 | 0.229239661 | 0.55384033 | 1 |
| Erlin2 | 1.04421903 | 0.06242435 | 0.55406577 | 1 |
| Spon1 | 0.96116242 | -0.057147853 | 0.55408072 | 1 |
| LOC688869 | 1.02819177 | 0.040109375 | 0.55415771 | 1 |

| | | | | |
|---|---|---|---|---|
| Adck2 | 1.05353008 | 0.075231511 | 0.55425712 | 1 |
| Casc5 | 1.0770898 | 0.10713854 | 0.5546215 | 1 |
| Etv6 | 0.89753658 | -0.15595735 | 0.55474441 | 1 |
| Csnk1a1 | 0.97088955 | -0.042620915 | 0.55502678 | 1 |
| Ftl1 | 1.02811588 | 0.040002888 | 0.55504873 | 1 |
| Six4 | 1.03099229 | 0.044033542 | 0.5553315 | 1 |
| Myh1 | 0.91974743 | -0.120690352 | 0.5553493 | 1 |
| Romo1 | 1.04507928 | 0.063612396 | 0.55539971 | 1 |
| Zfp444 | 0.93552489 | -0.096152063 | 0.55542515 | 1 |
| Depdc1 | 1.19630221 | 0.258581889 | 0.55545678 | 1 |
| Sobp | 0.94500719 | -0.081602795 | 0.55573628 | 1 |
| Trmt1 | 0.95892755 | -0.060506271 | 0.55584322 | 1 |
| Aar2 | 0.9512687 | -0.072075182 | 0.55596985 | 1 |
| Kdelc1 | 1.05780504 | 0.081073758 | 0.55598243 | 1 |
| Prmt1 | 0.96114164 | -0.057179036 | 0.55621901 | 1 |
| Cdk12 | 0.95961427 | -0.059473485 | 0.55629665 | 1 |
| Rai1 | 0.9622479 | -0.055519475 | 0.55634776 | 1 |
| Dcbld2 | 1.04504287 | 0.063562122 | 0.55639389 | 1 |
| Smpdl3b | 1.12254556 | 0.166773999 | 0.55654541 | 1 |
| Tmem199 | 0.96593882 | -0.049996281 | 0.55665403 | 1 |
| Zfp281 | 0.94217263 | -0.085936677 | 0.55666312 | 1 |
| Supt6h | 0.97316248 | -0.03924739 | 0.55694837 | 1 |
| Shisa4 | 0.95709331 | -0.063268514 | 0.55716695 | 1 |
| Duox2 | 1.11449588 | 0.156391284 | 0.55717396 | 1 |
| Bbx | 0.9393724 | -0.090230896 | 0.55723702 | 1 |
| Zbtb9 | 0.942353 | -0.085660506 | 0.55723706 | 1 |
| Txn1 | 1.02850554 | 0.040549563 | 0.55726983 | 1 |
| Arhgef2 | 0.96247749 | -0.055175302 | 0.55728242 | 1 |
| Rft1 | 1.07305333 | 0.101721772 | 0.55747695 | 1 |
| Slc12a9 | 0.93164311 | -0.102150692 | 0.55749655 | 1 |
| Loxl3 | 1.06969572 | 0.097200477 | 0.55752526 | 1 |
| Abhd12 | 0.97353238 | -0.038699135 | 0.55756433 | 1 |
| LOC102549726 | 1.04900162 | 0.069016903 | 0.55770945 | 1 |
| Crebbp | 1.04238606 | 0.059889695 | 0.55779772 | 1 |
| Pxk | 0.96205456 | -0.055809385 | 0.55780935 | 1 |
| Foxp1 | 0.92504975 | -0.112397131 | 0.55785798 | 1 |
| Phykpl | 0.88624404 | -0.174224078 | 0.55792537 | 1 |
| Med21 | 0.93885662 | -0.091023242 | 0.55813477 | 1 |
| Fam195b | 1.03966347 | 0.056116618 | 0.55828281 | 1 |
| Tmem14a | 1.03181772 | 0.045188134 | 0.55848995 | 1 |

| | | | | |
|---|---|---|---|---|
| Kpna3 | 0.96224129 | -0.055529392 | 0.55876129 | 1 |
| Cdkn1b | 1.03030122 | 0.043066184 | 0.55876609 | 1 |
| Cdkn3 | 0.83909241 | -0.253098387 | 0.55883099 | 1 |
| Tra2a | 0.96822812 | -0.046581096 | 0.55892965 | 1 |
| Psma1 | 1.02852692 | 0.040579548 | 0.55904061 | 1 |
| mrpl11 | 0.95198382 | -0.070991048 | 0.55906699 | 1 |
| Armcx3 | 0.97338304 | -0.038920459 | 0.55912319 | 1 |
| Gnas | 0.97238937 | -0.040393971 | 0.55917899 | 1 |
| Iffo2 | 1.1070909 | 0.146773679 | 0.55923281 | 1 |
| Pcdha12 | 1.07960819 | 0.11050782 | 0.55930426 | 1 |
| Cpped1 | 1.05310093 | 0.074643712 | 0.55948174 | 1 |
| Zcchc10 | 1.09786497 | 0.134700628 | 0.55957123 | 1 |
| Fsd1 | 0.94769362 | -0.077507373 | 0.55967232 | 1 |
| Snrpd1 | 1.04632919 | 0.065336815 | 0.55968435 | 1 |
| Tagln2 | 1.02710077 | 0.038577739 | 0.55971351 | 1 |
| Eps8l2 | 0.9070152 | -0.140801371 | 0.55978514 | 1 |
| Cbx8 | 0.91242386 | -0.132223928 | 0.56005731 | 1 |
| Stat3 | 0.97364254 | -0.038535895 | 0.56007224 | 1 |
| Slc4a10 | 0.87960266 | -0.18507613 | 0.56011773 | 1 |
| Pqlc2 | 1.10845215 | 0.148546494 | 0.56013175 | 1 |
| Prkrir | 1.03768485 | 0.05336835 | 0.56021117 | 1 |
| Zfp112 | 0.91283055 | -0.131581013 | 0.56028077 | 1 |
| Vps37c | 0.95968409 | -0.059368525 | 0.56028144 | 1 |
| Hps1 | 1.07127235 | 0.09932531 | 0.56049953 | 1 |
| Sdad1 | 0.94212157 | -0.086014861 | 0.56051469 | 1 |
| Tmem25 | 1.03215691 | 0.045662312 | 0.56051653 | 1 |
| Dclre1a | 1.05564593 | 0.078126024 | 0.56052167 | 1 |
| Desi2 | 0.93758238 | -0.092982641 | 0.56078239 | 1 |
| Mcoln1 | 0.96359738 | -0.053497617 | 0.56106161 | 1 |
| RGD1560394 | 0.91917132 | -0.121594315 | 0.56107904 | 1 |
| Zfp629 | 0.94517526 | -0.081346224 | 0.56109023 | 1 |
| Pter | 1.10014838 | 0.137698118 | 0.56111327 | 1 |
| Scyl2 | 1.02946658 | 0.041896997 | 0.5612228 | 1 |
| Spen | 0.96277825 | -0.05472455 | 0.56128294 | 1 |
| Stk19 | 1.06413804 | 0.089685315 | 0.56131326 | 1 |
| Slc12a6 | 0.97073854 | -0.042845328 | 0.56135031 | 1 |
| Ccr1 | 1.11072002 | 0.1514952 | 0.5614178 | 1 |
| Gabarap | 0.97350594 | -0.03873832 | 0.56142808 | 1 |
| Pik3ip1 | 0.96746704 | -0.047715582 | 0.56159442 | 1 |
| Terf1 | 1.06048747 | 0.084727577 | 0.56161267 | 1 |

| | | | | |
|---|---|---|---|---|
| Rgs4 | 0.9724341 | -0.040327607 | 0.56170074 | 1 |
| Nprl3 | 0.95237278 | -0.070401706 | 0.56176293 | 1 |
| Ppm1f | 0.96038829 | -0.058310275 | 0.56185343 | 1 |
| Uchl5 | 1.04303773 | 0.060791346 | 0.56187319 | 1 |
| Dedd | 1.04981585 | 0.070136291 | 0.56199962 | 1 |
| Fgf18 | 1.08585585 | 0.118832589 | 0.56208744 | 1 |
| Ubc | 0.97261234 | -0.040063204 | 0.56218382 | 1 |
| Tyrobp | 0.92710988 | -0.109187763 | 0.56218797 | 1 |
| L3mbtl2 | 0.96669809 | -0.048862705 | 0.56232823 | 1 |
| Mrpl43 | 1.04205606 | 0.059432893 | 0.5624038 | 1 |
| Vdac2 | 1.02703941 | 0.038491541 | 0.56245938 | 1 |
| Laptm4a | 1.02719094 | 0.038704387 | 0.5626444 | 1 |
| Nrp1 | 0.9668155 | -0.048687495 | 0.56272153 | 1 |
| Gsr | 0.96765469 | -0.047435785 | 0.56273158 | 1 |
| Arse | 1.07192801 | 0.100208021 | 0.56294146 | 1 |
| Fbxo30 | 0.96209009 | -0.055756106 | 0.56297864 | 1 |
| Pbx1 | 1.05105499 | 0.071838149 | 0.5631638 | 1 |
| Meis1 | 0.91388708 | -0.129912182 | 0.56318861 | 1 |
| Rab6b | 0.97260905 | -0.040068084 | 0.56364921 | 1 |
| Mospd1 | 1.03624851 | 0.051370022 | 0.5636899 | 1 |
| Ppp1r3a | 1.30959532 | 0.389121075 | 0.56373619 | 1 |
| Cdc16 | 1.03662275 | 0.051890956 | 0.5637897 | 1 |
| Ptk2b | 0.95785078 | -0.062127177 | 0.56388204 | 1 |
| Srsf6 | 0.97287685 | -0.039670899 | 0.56388858 | 1 |
| Esrra | 0.96208921 | -0.055757421 | 0.56395281 | 1 |
| Atcay | 1.02764969 | 0.039348551 | 0.56403495 | 1 |
| Mns1 | 1.10571932 | 0.144985208 | 0.56403994 | 1 |
| Tor2a | 1.05857245 | 0.082120016 | 0.56423518 | 1 |
| Fbxl3 | 1.03026573 | 0.043016492 | 0.5644462 | 1 |
| Pde1b | 0.90220459 | -0.148473469 | 0.56466608 | 1 |
| Mon2 | 0.97071312 | -0.042883106 | 0.56473352 | 1 |
| Lrrn2 | 0.96582373 | -0.050168179 | 0.56488347 | 1 |
| Mfsd14b | 0.96676733 | -0.048759367 | 0.56493266 | 1 |
| Ns5atp4 | 1.12207698 | 0.166171658 | 0.56497485 | 1 |
| Gpr165 | 1.13784803 | 0.186307887 | 0.56502826 | 1 |
| Acot13 | 1.0638602 | 0.089308584 | 0.56507976 | 1 |
| Snx30 | 1.06900371 | 0.096266856 | 0.56531838 | 1 |
| Slc10a3 | 0.9487551 | -0.075892361 | 0.56541517 | 1 |
| Ywhae | 0.97313521 | -0.039287825 | 0.56545179 | 1 |
| Erbb3 | 0.96949568 | -0.044693621 | 0.56548795 | 1 |

| | | | | |
|---|---|---|---|---|
| Dstn | 1.02696188 | 0.038382627 | 0.56549149 | 1 |
| Fgd3 | 0.88460314 | -0.176897728 | 0.56550586 | 1 |
| Fnbp1 | 0.97176143 | -0.041325921 | 0.56555692 | 1 |
| Fip1l1 | 0.96078959 | -0.057707581 | 0.56560418 | 1 |
| Map2k1 | 0.97307838 | -0.039372078 | 0.56573305 | 1 |
| Tkt | 0.97403459 | -0.037955086 | 0.565765 | 1 |
| Letm2 | 1.05546557 | 0.07787952 | 0.56580357 | 1 |
| Morf4l2 | 1.02690063 | 0.038296582 | 0.56586215 | 1 |
| Nbl1 | 0.9740929 | -0.037868719 | 0.56596976 | 1 |
| Pkmyt1 | 0.88014752 | -0.184182746 | 0.56618053 | 1 |
| Kdm5a | 1.03125983 | 0.044407876 | 0.56620271 | 1 |
| Sdccag3 | 0.96897859 | -0.045463305 | 0.56634204 | 1 |
| LOC690871 | 0.9318692 | -0.101800629 | 0.56635744 | 1 |
| RGD1304587 | 1.06165721 | 0.086318024 | 0.56639069 | 1 |
| Exoc2 | 0.97314192 | -0.039277881 | 0.56639786 | 1 |
| Fbxl6 | 1.07911451 | 0.109847961 | 0.56643586 | 1 |
| Rps24 | 1.02737037 | 0.038956369 | 0.56646396 | 1 |
| Ppp1r9a | 1.04767979 | 0.067197848 | 0.56655291 | 1 |
| Acox3 | 0.96145305 | -0.056711679 | 0.56656213 | 1 |
| Ccnj | 0.93663167 | -0.09444628 | 0.56659585 | 1 |
| Zfp949 | 0.92065067 | -0.119274247 | 0.56663232 | 1 |
| Gria3 | 0.95277024 | -0.069799741 | 0.56663238 | 1 |
| Fanca | 1.08804765 | 0.121741744 | 0.56669447 | 1 |
| Klf2 | 0.9310897 | -0.10300794 | 0.56670708 | 1 |
| Dopey2 | 0.97082213 | -0.042721103 | 0.56677139 | 1 |
| Plpp6 | 0.95318256 | -0.069175538 | 0.56679456 | 1 |
| Hist3h2a | 1.04217316 | 0.059595003 | 0.566838 | 1 |
| Senp3 | 1.03562862 | 0.050506737 | 0.56701856 | 1 |
| Chkb | 0.9597647 | -0.05924734 | 0.56708623 | 1 |
| Usp19 | 0.97215405 | -0.04074315 | 0.5671609 | 1 |
| Ppip5k2 | 1.04268478 | 0.060303074 | 0.5671745 | 1 |
| Jph3 | 0.97270696 | -0.039922849 | 0.56718145 | 1 |
| Ccdc17 | 1.10256057 | 0.14085791 | 0.56727546 | 1 |
| Gcs1 | 1.04823818 | 0.067966556 | 0.56729742 | 1 |
| Zbtb46 | 0.92345548 | -0.11488568 | 0.56730869 | 1 |
| Ticam2 | 1.15210876 | 0.204276917 | 0.56733399 | 1 |
| Zc3h12c | 0.92302049 | -0.115565414 | 0.56742992 | 1 |
| Il18bp | 1.10109642 | 0.138940812 | 0.56745574 | 1 |
| Sbk1 | 0.96643692 | -0.04925253 | 0.56748151 | 1 |
| Spon2 | 1.07345826 | 0.1022661 | 0.56755467 | 1 |

| | | | | |
|---|---|---|---|---|
| Snx24 | 0.95732204 | -0.062923773 | 0.56765165 | 1 |
| St8sia4 | 0.92113275 | -0.118519013 | 0.56781413 | 1 |
| Mrps27 | 1.04892823 | 0.068915974 | 0.56785921 | 1 |
| Acp1 | 1.02713215 | 0.038621812 | 0.56791172 | 1 |
| Bcl3 | 1.10580436 | 0.14509616 | 0.56804045 | 1 |
| Sec61a2 | 1.03118218 | 0.044299236 | 0.56806853 | 1 |
| Rpp38 | 1.06943807 | 0.09685294 | 0.56809639 | 1 |
| Usp1 | 1.04758285 | 0.067064352 | 0.56812325 | 1 |
| RGD1309594 | 0.96440676 | -0.05228633 | 0.56812885 | 1 |
| Tbc1d22b | 0.95523887 | -0.066066555 | 0.56814731 | 1 |
| LOC103689992 | 1.02864553 | 0.040745918 | 0.56829061 | 1 |
| Rps9 | 1.02864553 | 0.040745918 | 0.56829061 | 1 |
| Lime1 | 0.94019768 | -0.088963973 | 0.56839917 | 1 |
| Sh3glb1 | 0.96823532 | -0.046570374 | 0.56850691 | 1 |
| Tmem45b | 1.03971111 | 0.056182716 | 0.56852834 | 1 |
| Nckipsd | 0.96532426 | -0.050914465 | 0.56854512 | 1 |
| Il1b | 1.15160898 | 0.203650945 | 0.56855471 | 1 |
| Gch1 | 1.14145188 | 0.190870036 | 0.56885696 | 1 |
| Dock7 | 0.97188036 | -0.041149368 | 0.56905009 | 1 |
| Sec24c | 0.97166944 | -0.041462504 | 0.56907204 | 1 |
| Apba2 | 0.9687458 | -0.045809952 | 0.56908981 | 1 |
| Rnf111 | 1.03285278 | 0.046634626 | 0.56915663 | 1 |
| Dhodh | 0.92014648 | -0.120064543 | 0.56916246 | 1 |
| Eral1 | 1.05064883 | 0.071280546 | 0.56925073 | 1 |
| Acsl1 | 1.0281374 | 0.040033084 | 0.56944443 | 1 |
| Mtfmt | 1.06090994 | 0.08530219 | 0.56963326 | 1 |
| Acadl | 1.0278445 | 0.039622022 | 0.5696678 | 1 |
| Ppp6r2 | 0.9699424 | -0.044029016 | 0.56977881 | 1 |
| Rundc3a | 0.97362875 | -0.038556321 | 0.56991113 | 1 |
| Thbs2 | 0.95783282 | -0.062154221 | 0.5699125 | 1 |
| Dmtn | 0.97398796 | -0.038024161 | 0.56994962 | 1 |
| Cct8 | 1.02655182 | 0.037806458 | 0.57000082 | 1 |
| Mok | 1.09706963 | 0.133655091 | 0.57001477 | 1 |
| Ankrd46 | 1.02639353 | 0.037583982 | 0.57001992 | 1 |
| Dck | 1.06915244 | 0.096467572 | 0.57004017 | 1 |
| Gar1 | 0.91826204 | -0.123022183 | 0.57020206 | 1 |
| Mturn | 0.97336101 | -0.038953108 | 0.57024367 | 1 |
| Golga2 | 0.973078 | -0.039372637 | 0.57027336 | 1 |
| Mast3 | 0.96419034 | -0.052610121 | 0.57052529 | 1 |
| Fgfr1op | 0.95231295 | -0.070492348 | 0.57064761 | 1 |

| | | | | |
|---|---|---|---|---|
| Spats2 | 1.03582073 | 0.05077434 | 0.57083121 | 1 |
| Usp32 | 0.97427342 | -0.037601381 | 0.57086673 | 1 |
| Psmd11 | 0.97209957 | -0.040824 | 0.57091212 | 1 |
| Aoc3 | 1.03492522 | 0.049526529 | 0.57095501 | 1 |
| Myo10 | 1.03879727 | 0.054914122 | 0.57098888 | 1 |
| Serpinb1a | 0.9742585 | -0.037623477 | 0.57105811 | 1 |
| Sgta | 0.96584845 | -0.050131262 | 0.57106231 | 1 |
| Ccm2 | 0.96266124 | -0.054899897 | 0.57117676 | 1 |
| Arfgef1 | 0.97436863 | -0.037460405 | 0.57124211 | 1 |
| Alg2 | 0.9737207 | -0.038420079 | 0.57124913 | 1 |
| Dusp7 | 0.95229042 | -0.070526477 | 0.57126319 | 1 |
| Mtbp | 1.11853356 | 0.161608541 | 0.57126551 | 1 |
| Nek4 | 1.05662967 | 0.079469823 | 0.57136999 | 1 |
| Hoxd3 | 0.90726178 | -0.140409213 | 0.57149077 | 1 |
| Cd9 | 0.9740847 | -0.037880867 | 0.57157328 | 1 |
| Mau2 | 0.96637852 | -0.049339711 | 0.57178828 | 1 |
| Phf3 | 1.03254395 | 0.046203187 | 0.57183267 | 1 |
| Stim2 | 0.97081031 | -0.04273866 | 0.57186333 | 1 |
| Arhgef40 | 0.97161959 | -0.041536524 | 0.57189657 | 1 |
| Rxrg | 0.95110635 | -0.072321423 | 0.5720005 | 1 |
| Tmem138 | 0.91370595 | -0.130198138 | 0.57203322 | 1 |
| Eepd1 | 0.97265784 | -0.039995715 | 0.57211826 | 1 |
| Pdlim1 | 0.96704339 | -0.048347472 | 0.57222004 | 1 |
| B3galnt2 | 0.95799776 | -0.061905812 | 0.57222203 | 1 |
| Cdk2ap1 | 0.97213253 | -0.04077508 | 0.57224362 | 1 |
| Smu1 | 1.03612888 | 0.051203463 | 0.57232076 | 1 |
| Cebpd | 1.06217465 | 0.087021003 | 0.5723365 | 1 |
| Tmem39a | 1.05806304 | 0.08142559 | 0.57235775 | 1 |
| Nr3c2 | 1.03691726 | 0.052300781 | 0.57235885 | 1 |
| Apoe | 1.02849168 | 0.04053012 | 0.57249422 | 1 |
| Gal3st4 | 1.09397199 | 0.129575802 | 0.57253094 | 1 |
| Tmem123 | 0.96043423 | -0.058241277 | 0.57273658 | 1 |
| Sprn | 0.90808439 | -0.139101721 | 0.57288885 | 1 |
| Itch | 1.03228778 | 0.045845227 | 0.57300011 | 1 |
| Cdc42bpa | 0.974013 | -0.037987074 | 0.57302582 | 1 |
| Mapk10 | 1.02613091 | 0.037214799 | 0.57306421 | 1 |
| MGC95208 | 0.94155175 | -0.086887701 | 0.57319175 | 1 |
| Msl1 | 0.96950915 | -0.044673581 | 0.57340775 | 1 |
| Pkd2 | 0.96593365 | -0.050003999 | 0.57343965 | 1 |
| Slc8a2 | 1.06734973 | 0.094032971 | 0.57345655 | 1 |

| | | | | |
|---|---|---|---|---|
| Rpl31 | 1.02913931 | 0.041438287 | 0.57352084 | 1 |
| Stox2 | 0.95139533 | -0.071883151 | 0.5736721 | 1 |
| Leng1 | 1.06080063 | 0.085153539 | 0.57374797 | 1 |
| Ero1a | 0.96413698 | -0.052689958 | 0.5738475 | 1 |
| Trip6 | 0.9393807 | -0.090218146 | 0.57402967 | 1 |
| Rps12 | 1.03562896 | 0.05050722 | 0.57403179 | 1 |
| Mt3 | 1.02743618 | 0.039048782 | 0.57412001 | 1 |
| Mcm5 | 0.88601291 | -0.174600376 | 0.57420669 | 1 |
| F8a1 | 1.04963297 | 0.069884947 | 0.57437055 | 1 |
| Stk11 | 1.03831253 | 0.054240751 | 0.57447557 | 1 |
| Ddx17 | 0.97368967 | -0.038466062 | 0.57448489 | 1 |
| Eef1d | 0.97129833 | -0.042013608 | 0.57454113 | 1 |
| Mmp28 | 1.08154811 | 0.113097836 | 0.57461264 | 1 |
| Swt1 | 0.95583488 | -0.065166686 | 0.57462543 | 1 |
| Btd | 0.95859981 | -0.060999446 | 0.5746564 | 1 |
| Plrg1 | 1.03657628 | 0.051826281 | 0.57466097 | 1 |
| Sfi1 | 1.04228036 | 0.059743398 | 0.57467986 | 1 |
| Alg1 | 0.94653803 | -0.079267616 | 0.57480295 | 1 |
| Tomm70 | 1.02906464 | 0.041333607 | 0.57481144 | 1 |
| Zfp180 | 1.02620093 | 0.03731324 | 0.57482822 | 1 |
| Hps6 | 0.95115953 | -0.072240765 | 0.57485809 | 1 |
| Cpox | 1.05090369 | 0.071630464 | 0.57500412 | 1 |
| Sertad2 | 0.93327132 | -0.099631539 | 0.57501339 | 1 |
| Mff | 1.02605005 | 0.037101104 | 0.57506825 | 1 |
| Cstf1 | 0.95464519 | -0.066963472 | 0.57510363 | 1 |
| Gstm5 | 1.05081389 | 0.071507173 | 0.57510608 | 1 |
| Abcf1 | 0.97076121 | -0.042811638 | 0.57514611 | 1 |
| Itgae | 1.08371078 | 0.115979789 | 0.57517535 | 1 |
| Usp42 | 0.95230195 | -0.070509001 | 0.57526345 | 1 |
| Rhbdf2 | 0.93503407 | -0.096909168 | 0.57538482 | 1 |
| Rem1 | 1.09621349 | 0.132528792 | 0.57543262 | 1 |
| Tcerg1 | 1.02718774 | 0.038699889 | 0.57544063 | 1 |
| Col5a1 | 1.02954242 | 0.042003275 | 0.57559147 | 1 |
| Lrrc58 | 0.96369433 | -0.053352475 | 0.57561102 | 1 |
| Gna14 | 1.02714128 | 0.038634628 | 0.57565733 | 1 |
| Rpl18 | 1.02785462 | 0.039636222 | 0.57570193 | 1 |
| Hspa13 | 0.9610994 | -0.057242442 | 0.57581417 | 1 |
| Tgfb1 | 1.07247853 | 0.100948761 | 0.57603032 | 1 |
| Ogfrl1 | 0.96791244 | -0.047051555 | 0.57606524 | 1 |
| Scamp4 | 0.96096896 | -0.057438263 | 0.57628167 | 1 |

| | | | | |
|---|---|---|---|---|
| St3gal6 | 0.95983238 | -0.059145611 | 0.57635325 | 1 |
| Trappc6b | 1.03081943 | 0.043791639 | 0.57642334 | 1 |
| Csnk1d | 0.97473957 | -0.036911281 | 0.57646051 | 1 |
| Rpn1 | 1.02591509 | 0.036911329 | 0.57664388 | 1 |
| Endog | 0.91444423 | -0.129032906 | 0.57666613 | 1 |
| Socs6 | 1.05459289 | 0.07668618 | 0.57670698 | 1 |
| Mllt11 | 0.97465875 | -0.037030903 | 0.57671103 | 1 |
| Itga5 | 0.94183328 | -0.086456387 | 0.5768276 | 1 |
| Abca17 | 0.87508794 | -0.192500098 | 0.57688556 | 1 |
| Stx2 | 0.95766818 | -0.062402226 | 0.5768949 | 1 |
| Tmem97 | 0.95540727 | -0.065812245 | 0.57694681 | 1 |
| Rfesd | 0.94915652 | -0.075282085 | 0.57695608 | 1 |
| Mmp11 | 0.9096015 | -0.136693455 | 0.57699787 | 1 |
| Thtpa | 0.96886323 | -0.045635069 | 0.57706628 | 1 |
| Spry1 | 0.91142389 | -0.133805902 | 0.57708249 | 1 |
| Med9 | 1.04240826 | 0.059920415 | 0.57709534 | 1 |
| Ddah2 | 1.0333787 | 0.047369053 | 0.57747823 | 1 |
| RGD1311899 | 0.9727628 | -0.039840036 | 0.57748286 | 1 |
| Slc25a20 | 1.0340226 | 0.048267715 | 0.57754782 | 1 |
| Prosc | 1.04678794 | 0.06596921 | 0.57764483 | 1 |
| Pacsin3 | 1.04849309 | 0.068317355 | 0.57769455 | 1 |
| Gart | 0.96698844 | -0.048429456 | 0.57773157 | 1 |
| Foxj1 | 0.9001266 | -0.151800162 | 0.57789445 | 1 |
| Ctps1 | 0.97168463 | -0.041439944 | 0.57796595 | 1 |
| Pcmt1 | 1.02614401 | 0.03723321 | 0.57800028 | 1 |
| Nme2 | 1.02690555 | 0.038303495 | 0.57805907 | 1 |
| Lilrb3a | 1.12961091 | 0.17582593 | 0.57815544 | 1 |
| Lilrb3b | 1.12961091 | 0.17582593 | 0.57815544 | 1 |
| Pde10a | 0.94161675 | -0.086788111 | 0.57818814 | 1 |
| Uqcrb | 1.03070235 | 0.043627764 | 0.57819515 | 1 |
| Rcbtb1 | 1.02992294 | 0.042536395 | 0.57829925 | 1 |
| Tnrc6b | 1.02877247 | 0.040923947 | 0.57830332 | 1 |
| Fbxl14 | 0.95714276 | -0.063193973 | 0.57832699 | 1 |
| Mrps2 | 0.94953893 | -0.074700944 | 0.57858029 | 1 |
| RGD1304810 | 0.92600161 | -0.110913395 | 0.57858157 | 1 |
| Card10 | 0.93185239 | -0.101826652 | 0.57863887 | 1 |
| Cysltr2 | 0.9415111 | -0.08694999 | 0.57865751 | 1 |
| Elf2 | 0.95745962 | -0.062716456 | 0.57873086 | 1 |
| Daxx | 0.9539495 | -0.068015196 | 0.57890985 | 1 |
| Dr1 | 1.03562118 | 0.050496382 | 0.57930802 | 1 |

| | | | | |
|---|---|---|---|---|
| Samd14 | 0.96589178 | -0.05006654 | 0.57934834 | 1 |
| Xpot | 0.9693037 | -0.04497934 | 0.57964752 | 1 |
| Cwc25 | 0.95281243 | -0.069735865 | 0.57973996 | 1 |
| Rilpl1 | 1.04736786 | 0.066768242 | 0.57987945 | 1 |
| RGD1359108 | 1.02824316 | 0.04018147 | 0.57995579 | 1 |
| Scg3 | 0.97450535 | -0.037257995 | 0.58013153 | 1 |
| Gnb5 | 0.96134995 | -0.056866393 | 0.58026163 | 1 |
| Emc8 | 0.94652199 | -0.079292072 | 0.58041822 | 1 |
| Pnrc2 | 1.03003889 | 0.042698814 | 0.58055441 | 1 |
| Ctf1 | 0.90796865 | -0.139285616 | 0.58062895 | 1 |
| Ppme1 | 1.02637536 | 0.037558438 | 0.58078444 | 1 |
| Asns | 0.97173714 | -0.041361979 | 0.58079044 | 1 |
| Ostm1 | 0.96244719 | -0.055220708 | 0.58082736 | 1 |
| Frem1 | 1.12703636 | 0.172534065 | 0.58085956 | 1 |
| Ptpn18 | 1.12212256 | 0.16623026 | 0.58111182 | 1 |
| Kmt5a | 0.95877765 | -0.060731817 | 0.58141853 | 1 |
| Prkab2 | 0.96043105 | -0.058246052 | 0.58144561 | 1 |
| Prmt2 | 0.96720363 | -0.048108441 | 0.58152119 | 1 |
| Ywhah | 0.97345655 | -0.038811511 | 0.58157014 | 1 |
| Jrkl | 1.05611082 | 0.078761224 | 0.5816204 | 1 |
| Tsr3 | 1.05773234 | 0.080974595 | 0.58163722 | 1 |
| Nphp4 | 0.94145927 | -0.087029419 | 0.58174206 | 1 |
| Fam173b | 0.94884571 | -0.075754587 | 0.58177576 | 1 |
| Gtf2h3 | 0.94529463 | -0.081164039 | 0.5819637 | 1 |
| Dync2h1 | 0.96477303 | -0.051738517 | 0.58201738 | 1 |
| Akr1b1 | 1.02636359 | 0.037541898 | 0.58213207 | 1 |
| Ptcd3 | 1.03337806 | 0.047368154 | 0.58235097 | 1 |
| Bcl11a | 1.12070502 | 0.164406595 | 0.58236599 | 1 |
| Plekha7 | 1.0983052 | 0.135279006 | 0.58279439 | 1 |
| Esf1 | 1.03780666 | 0.053537701 | 0.58290595 | 1 |
| LOC500684 | 0.93294038 | -0.100143206 | 0.58291623 | 1 |
| Afg3l1 | 0.95833104 | -0.061403999 | 0.5829942 | 1 |
| Vezt | 1.02689564 | 0.038289566 | 0.58301465 | 1 |
| Ints12 | 0.93779273 | -0.092659004 | 0.58304466 | 1 |
| Rasal2 | 0.96237356 | -0.055331095 | 0.58350645 | 1 |
| Gtf2h5 | 0.96458072 | -0.052026114 | 0.5836131 | 1 |
| Sfrp1 | 1.03474486 | 0.049275078 | 0.58363799 | 1 |
| Tmem117 | 1.030447 | 0.043270306 | 0.58376373 | 1 |
| Gli3 | 1.11729124 | 0.160005303 | 0.5838962 | 1 |
| Dtx3 | 1.04239616 | 0.05990367 | 0.58399271 | 1 |

| | | | | |
|---|---|---|---|---|
| Cited1 | 1.13070087 | 0.177217311 | 0.5840098 | 1 |
| Zfp579 | 0.93043157 | -0.104028041 | 0.58406007 | 1 |
| Ppm1b | 0.97407117 | -0.037900911 | 0.5842504 | 1 |
| Oprd1 | 0.88210372 | -0.180979787 | 0.58455747 | 1 |
| Fam160a1 | 1.19611938 | 0.258361386 | 0.58456304 | 1 |
| Ccdc93 | 1.044531 | 0.062855309 | 0.58466407 | 1 |
| Lck | 0.90844264 | -0.138532678 | 0.58467424 | 1 |
| Fnip1 | 1.03398945 | 0.048221468 | 0.58479659 | 1 |
| Jtb | 1.04176647 | 0.059031913 | 0.5848986 | 1 |
| Tmem69 | 1.04949459 | 0.069694725 | 0.58492301 | 1 |
| Elmod2 | 1.03356891 | 0.047634578 | 0.58495741 | 1 |
| Crkl | 1.03484921 | 0.049420565 | 0.58500157 | 1 |
| Cab39 | 1.02562556 | 0.036504118 | 0.58506398 | 1 |
| Letm1 | 0.97093945 | -0.042546769 | 0.58510276 | 1 |
| Prdx3 | 1.02678977 | 0.038140834 | 0.58517944 | 1 |
| Uba6 | 1.0271067 | 0.038586065 | 0.5853664 | 1 |
| Cchcr1 | 0.95262972 | -0.070012536 | 0.58541144 | 1 |
| Insr | 0.95424633 | -0.067566367 | 0.58547879 | 1 |
| Vopp1 | 0.97400812 | -0.037994294 | 0.58549237 | 1 |
| Ptpru | 1.06241109 | 0.087342112 | 0.58550136 | 1 |
| Tnpo3 | 0.970992 | -0.042468693 | 0.58560489 | 1 |
| Pls3 | 1.02566147 | 0.036554636 | 0.58561422 | 1 |
| Atp1a4 | 0.94837165 | -0.07647556 | 0.58577445 | 1 |
| Eno2 | 1.02618111 | 0.03728538 | 0.58583677 | 1 |
| Znrd1as1 | 1.09242427 | 0.127533268 | 0.5858823 | 1 |
| Stard10 | 0.96863156 | -0.045980082 | 0.58599749 | 1 |
| Fcgr3a | 0.87625719 | -0.190573718 | 0.58607278 | 1 |
| Wdr26 | 0.97395085 | -0.038079133 | 0.58607761 | 1 |
| Rassf3 | 0.95248319 | -0.07023446 | 0.58614963 | 1 |
| Wasf1 | 0.96782424 | -0.047183015 | 0.58620017 | 1 |
| Tsc1 | 1.04233153 | 0.059814228 | 0.58622451 | 1 |
| Neto1 | 0.90383556 | -0.145867783 | 0.58627421 | 1 |
| Casp6 | 1.1054734 | 0.144664313 | 0.58639411 | 1 |
| LOC498276 | 0.92352547 | -0.114776339 | 0.58642586 | 1 |
| Gpr174 | 0.86676746 | -0.206283094 | 0.58651529 | 1 |
| Ppt1 | 1.02950082 | 0.041944984 | 0.58657042 | 1 |
| Mest | 0.96559314 | -0.050512665 | 0.58659542 | 1 |
| Ddit3 | 1.04253715 | 0.060098798 | 0.58659716 | 1 |
| Ccdc8 | 0.93871656 | -0.091238491 | 0.58666469 | 1 |
| Cldn22 | 1.09836405 | 0.135356305 | 0.58668465 | 1 |

| | | | | | |
|---|---|---|---|---|---|
| Stk24 | | 0.96528948 | -0.050966433 | 0.58673353 | 1 |
| Lancl3 | | 1.08423593 | 0.116678719 | 0.58674085 | 1 |
| Pi4ka | | 1.02524192 | 0.035964369 | 0.58681534 | 1 |
| Timm21 | | 0.95851464 | -0.061127627 | 0.58702756 | 1 |
| Stt3a | | 1.02730915 | 0.038870397 | 0.58720131 | 1 |
| Aagab | | 1.05199103 | 0.073122408 | 0.58739106 | 1 |
| Ccpg1 | | 1.02587494 | 0.03685487 | 0.58767371 | 1 |
| Stard3 | | 0.96278663 | -0.054711986 | 0.58773443 | 1 |
| Mmaa | | 0.93348826 | -0.099296215 | 0.58774197 | 1 |
| Lmx1a | | 0.87765064 | -0.188281317 | 0.58783198 | 1 |
| Acsf2 | | 1.04686557 | 0.066076196 | 0.58789687 | 1 |
| Plin2 | | 1.02950182 | 0.041946374 | 0.5879514 | 1 |
| Ubfd1 | | 1.04387688 | 0.061951568 | 0.58797523 | 1 |
| Rfc5 | | 1.07183049 | 0.100076763 | 0.58799323 | 1 |
| Ppp2r2a | | 0.97202531 | -0.040934216 | 0.58806155 | 1 |
| Hps4 | | 0.95573775 | -0.065313298 | 0.58815287 | 1 |
| Cfap20 | | 1.03920537 | 0.05548079 | 0.58818837 | 1 |
| Tnxa-ps1 | | 0.95842496 | -0.061262615 | 0.58820694 | 1 |
| Psma7 | | 1.02639666 | 0.037588376 | 0.58821119 | 1 |
| | 10-Mar | 1.13103013 | 0.17763736 | 0.58825819 | 1 |
| Insig2 | | 1.06816786 | 0.095138387 | 0.58826126 | 1 |
| Aco1 | | 0.96696523 | -0.048464083 | 0.5882891 | 1 |
| Rnf40 | | 1.03866403 | 0.054729067 | 0.58838967 | 1 |
| Dpm3 | | 0.94045164 | -0.088574341 | 0.58839595 | 1 |
| Nup205 | | 1.03825793 | 0.054164896 | 0.5883965 | 1 |
| Bcl2a1 | | 1.12834029 | 0.17420223 | 0.58840991 | 1 |
| H6pd | | 1.06154494 | 0.08616545 | 0.58855604 | 1 |
| Brap | | 0.97430337 | -0.037557043 | 0.58861934 | 1 |
| Frk | | 1.09143503 | 0.126226253 | 0.58882491 | 1 |
| Cisd1 | | 1.02832009 | 0.040289404 | 0.58909297 | 1 |
| Kif2c | | 1.1757297 | 0.233556425 | 0.589112 | 1 |
| Klhl22 | | 1.03764559 | 0.053313778 | 0.58911278 | 1 |
| Mdh2 | | 1.02511688 | 0.035788412 | 0.58928269 | 1 |
| Ergic2 | | 1.03435909 | 0.048737127 | 0.58938419 | 1 |
| Ankrd16 | | 1.08412088 | 0.116525625 | 0.58943896 | 1 |
| Katna1 | | 1.06256949 | 0.087557199 | 0.58953688 | 1 |
| Nfib | | 0.9448833 | -0.081791944 | 0.58970861 | 1 |
| Hmgxb4 | | 1.06722926 | 0.093870131 | 0.58973408 | 1 |
| Mpp7 | | 1.0601125 | 0.084217368 | 0.58985195 | 1 |
| Eif5b | | 1.03081624 | 0.043787165 | 0.58997566 | 1 |

| | | | | |
|---|---|---|---|---|
| Akt1s1 | 0.96703883 | -0.048354279 | 0.59019062 | 1 |
| Snrpe | 1.04717879 | 0.066507782 | 0.59022955 | 1 |
| Trim16 | 1.04191366 | 0.059235735 | 0.59029847 | 1 |
| Chrna6 | 1.03915352 | 0.055408801 | 0.59036817 | 1 |
| Cops8 | 1.02731964 | 0.038885132 | 0.59039172 | 1 |
| Dbndd1 | 1.05311636 | 0.074664851 | 0.59051824 | 1 |
| RT1-CE5 | 1.03424401 | 0.048576597 | 0.5905535 | 1 |
| Arl14ep | 1.04397596 | 0.062088497 | 0.5906172 | 1 |
| Celf6 | 0.95373167 | -0.068344678 | 0.59066763 | 1 |
| Acy3 | 0.93781282 | -0.092628093 | 0.59069433 | 1 |
| Bcat2 | 1.03219079 | 0.045709663 | 0.59082406 | 1 |
| Pms1 | 1.05528555 | 0.077633436 | 0.59084435 | 1 |
| Smco4 | 0.87777207 | -0.188081726 | 0.59094974 | 1 |
| Clasrp | 0.96175646 | -0.056256473 | 0.59105153 | 1 |
| Smg7 | 0.97075653 | -0.042818591 | 0.59107178 | 1 |
| Snap25 | 1.02641959 | 0.037620612 | 0.59110377 | 1 |
| Enoph1 | 0.96840627 | -0.04631568 | 0.59113036 | 1 |
| Gca | 0.96220834 | -0.055578786 | 0.59124364 | 1 |
| Metrnl | 1.05116359 | 0.071987212 | 0.59127512 | 1 |
| Fen1 | 1.04918736 | 0.069272329 | 0.59133187 | 1 |
| Gsta3 | 1.14375507 | 0.193778137 | 0.59148768 | 1 |
| LOC361985 | 1.02881505 | 0.040983646 | 0.59150742 | 1 |
| Taf7 | 0.94443203 | -0.082481131 | 0.59170292 | 1 |
| Pus3 | 1.05178599 | 0.072841188 | 0.59172442 | 1 |
| Impact | 0.97485502 | -0.036740411 | 0.59198431 | 1 |
| Ift20 | 1.03653922 | 0.0517747 | 0.59204095 | 1 |
| Gipc2 | 0.92067943 | -0.119229182 | 0.59219872 | 1 |
| Dll4 | 0.91204964 | -0.132815754 | 0.59223883 | 1 |
| Cox15 | 1.03245517 | 0.046079147 | 0.5923095 | 1 |
| Gpr139 | 1.10510083 | 0.144178012 | 0.59231432 | 1 |
| Rbmx | 0.9589233 | -0.060512676 | 0.59249335 | 1 |
| Hoxb4 | 1.09606468 | 0.132332935 | 0.59257516 | 1 |
| Ptcd1 | 1.04857064 | 0.068424061 | 0.59261055 | 1 |
| Taco1 | 0.92974849 | -0.105087599 | 0.59266585 | 1 |
| Magoh | 0.95506055 | -0.066335891 | 0.59266944 | 1 |
| Spata5l1 | 0.91213485 | -0.13268097 | 0.59267881 | 1 |
| Nfkb2 | 0.94011345 | -0.089093225 | 0.59268005 | 1 |
| Casq1 | 0.89070081 | -0.166987184 | 0.59272895 | 1 |
| Apopt1 | 0.95756667 | -0.062555158 | 0.59276338 | 1 |
| Itpr2 | 1.04301055 | 0.060753754 | 0.59293425 | 1 |

| | | | | |
|---|---|---|---|---|
| P3h3 | 1.03370143 | 0.047819546 | 0.59312773 | 1 |
| Nudt7 | 1.04237726 | 0.059877516 | 0.59314782 | 1 |
| Slc25a19 | 1.05658698 | 0.079411537 | 0.59315749 | 1 |
| Rab11fip4 | 0.96761441 | -0.047495836 | 0.59316109 | 1 |
| Tspyl1 | 0.97465521 | -0.037036147 | 0.59323205 | 1 |
| Fth1 | 1.02683687 | 0.038206999 | 0.59323568 | 1 |
| Pmpca | 1.03029441 | 0.043056654 | 0.59324428 | 1 |
| Anxa11 | 0.97472356 | -0.036934983 | 0.5932966 | 1 |
| Zfp846 | 1.04457193 | 0.062911837 | 0.59331219 | 1 |
| Mcfd2 | 1.03756457 | 0.053201127 | 0.59331923 | 1 |
| Ccni | 0.97577501 | -0.03537956 | 0.59353723 | 1 |
| Josd2 | 0.96288868 | -0.054559072 | 0.59355935 | 1 |
| Cbll1 | 1.03912061 | 0.055363116 | 0.59357238 | 1 |
| Glrx3 | 1.02526519 | 0.035997125 | 0.59370808 | 1 |
| Lhfpl1 | 0.95447185 | -0.067225447 | 0.59375862 | 1 |
| Bivm | 1.0388461 | 0.054981945 | 0.59379356 | 1 |
| Foxn3 | 0.95634712 | -0.064393731 | 0.59382817 | 1 |
| Ralgapa1 | 1.02524677 | 0.035971195 | 0.59395367 | 1 |
| Pak3 | 1.04651213 | 0.065589038 | 0.5939555 | 1 |
| Zfp523 | 1.04612008 | 0.065048463 | 0.5940012 | 1 |
| Ppp1r14b | 1.04452635 | 0.062848892 | 0.59401834 | 1 |
| Pros1 | 0.96539242 | -0.050812603 | 0.59402001 | 1 |
| Jag1 | 0.96891179 | -0.045562771 | 0.59410919 | 1 |
| Padi4 | 1.22640712 | 0.294437976 | 0.59411112 | 1 |
| Rcn2 | 0.97313739 | -0.039284591 | 0.59415436 | 1 |
| Psmd9 | 1.03817349 | 0.054047559 | 0.59423583 | 1 |
| Srsf10 | 0.95713314 | -0.063208477 | 0.5943058 | 1 |
| Krtcap2 | 0.96638948 | -0.049323344 | 0.59430924 | 1 |
| Rela | 1.03909585 | 0.055328746 | 0.59431385 | 1 |
| Cdh7 | 1.0645388 | 0.09022853 | 0.59452686 | 1 |
| Cltb | 1.02516872 | 0.035861372 | 0.59455434 | 1 |
| Gabra3 | 1.05038655 | 0.070920345 | 0.59461906 | 1 |
| Hdgf | 0.96937557 | -0.044872376 | 0.59465019 | 1 |
| Fkbp8 | 0.97572434 | -0.035454477 | 0.59469542 | 1 |
| Akna | 1.0566344 | 0.079476284 | 0.5947048 | 1 |
| Hectd2 | 1.04784789 | 0.06742931 | 0.59475149 | 1 |
| Ccnt1 | 1.07549028 | 0.104994481 | 0.59479088 | 1 |
| Map10 | 1.11306925 | 0.154543359 | 0.59502703 | 1 |
| Slc25a11 | 0.96858548 | -0.046048724 | 0.59507122 | 1 |
| Cdc42 | 1.02482803 | 0.035381843 | 0.59512466 | 1 |

| | | | | |
|---|---|---|---|---|
| Fbxw11 | 0.97536298 | -0.035988876 | 0.59515878 | 1 |
| Faim | 1.06144473 | 0.086029253 | 0.59523727 | 1 |
| Myo1g | 1.09167169 | 0.12653905 | 0.59528177 | 1 |
| Bcar3 | 0.96216645 | -0.055641599 | 0.59539934 | 1 |
| Fzd9 | 1.13032087 | 0.17673238 | 0.59547486 | 1 |
| Prkcsh | 0.97445324 | -0.037335131 | 0.59552312 | 1 |
| Nudt9 | 0.96573562 | -0.050299803 | 0.59575284 | 1 |
| Abcc12 | 1.08248665 | 0.114349234 | 0.59588282 | 1 |
| Ei24 | 0.97325528 | -0.039109834 | 0.59588403 | 1 |
| Polr2k | 1.03398499 | 0.048215249 | 0.595928 | 1 |
| Dph3 | 1.04313769 | 0.060929604 | 0.59595936 | 1 |
| Tmtc3 | 0.96597926 | -0.049935883 | 0.59605656 | 1 |
| Yipf4 | 1.03005792 | 0.04272546 | 0.59605742 | 1 |
| Pex16 | 1.04888875 | 0.068861664 | 0.59611912 | 1 |
| LOC100294508 | 0.97281175 | -0.039767447 | 0.59612855 | 1 |
| Rhoq | 1.02569582 | 0.03660295 | 0.59617793 | 1 |
| Itgb5 | 1.02775353 | 0.039494329 | 0.59637257 | 1 |
| Riok2 | 0.95747847 | -0.062688042 | 0.59648609 | 1 |
| Xrcc4 | 1.06513697 | 0.091038969 | 0.59650787 | 1 |
| Slc35g1 | 0.89024139 | -0.167731518 | 0.59664983 | 1 |
| Mecr | 0.93834254 | -0.091813427 | 0.59672348 | 1 |
| Snrpc | 1.0503818 | 0.07091382 | 0.59673492 | 1 |
| Zdhhc4 | 0.94412254 | -0.082953968 | 0.59681519 | 1 |
| Vnn1 | 0.86655232 | -0.206641234 | 0.59691147 | 1 |
| Scube3 | 0.90295751 | -0.147270002 | 0.59718133 | 1 |
| Nsun6 | 1.07737814 | 0.107524702 | 0.59720874 | 1 |
| Timp1 | 0.96712769 | -0.048221719 | 0.59721561 | 1 |
| Rspo2 | 0.95663994 | -0.063952065 | 0.59735552 | 1 |
| Stard6 | 1.09651676 | 0.132927867 | 0.59737358 | 1 |
| Calu | 1.02449506 | 0.034913025 | 0.59763799 | 1 |
| Oxct1 | 0.97615789 | -0.034813573 | 0.59779363 | 1 |
| Lgi1 | 1.02805131 | 0.039912273 | 0.59783039 | 1 |
| Col2a1 | 0.94941539 | -0.07488866 | 0.59808482 | 1 |
| Zfp207 | 1.02743021 | 0.039040395 | 0.59825464 | 1 |
| Ube2j1 | 0.97286053 | -0.039695099 | 0.59832556 | 1 |
| Specc1l | 1.02785427 | 0.039635735 | 0.59845994 | 1 |
| Cpeb4 | 0.96545404 | -0.050720516 | 0.59859201 | 1 |
| Usp8 | 1.02464947 | 0.035130458 | 0.59866812 | 1 |
| Rtn4r | 0.94103348 | -0.087682041 | 0.59868707 | 1 |
| Cntn6 | 1.07118275 | 0.099204627 | 0.59882548 | 1 |

| | | | | |
|---|---|---|---|---|
| Tmem143 | 0.94569517 | -0.08055286 | 0.59885957 | 1 |
| Plekha4 | 0.97580347 | -0.035337487 | 0.5988925 | 1 |
| Abcb9 | 0.9687504 | -0.045803093 | 0.59896771 | 1 |
| Paqr6 | 0.96042917 | -0.05824887 | 0.59901112 | 1 |
| Tmtc4 | 0.97081382 | -0.042733449 | 0.59914276 | 1 |
| Mtif3 | 0.94657693 | -0.079208339 | 0.59924749 | 1 |
| Scaf8 | 0.9683506 | -0.046398606 | 0.59939635 | 1 |
| Moap1 | 0.94970793 | -0.074444192 | 0.59941182 | 1 |
| Atp6v1b2 | 1.02506227 | 0.035711558 | 0.59941294 | 1 |
| E4f1 | 1.05083737 | 0.071539409 | 0.59949214 | 1 |
| Phactr3 | 0.95052481 | -0.073203804 | 0.59953706 | 1 |
| Tmem30b | 0.94356494 | -0.083806279 | 0.59957249 | 1 |
| Lrrtm2 | 0.95174523 | -0.071352663 | 0.59970501 | 1 |
| Pdk1 | 1.05650486 | 0.07929941 | 0.59979691 | 1 |
| Ccbl1 | 0.94780457 | -0.077338479 | 0.59980101 | 1 |
| Sfxn3 | 1.02542227 | 0.036218133 | 0.60015114 | 1 |
| RGD1307461 | 0.96079978 | -0.057692282 | 0.60019328 | 1 |
| Kdm4d | 1.10770088 | 0.147568349 | 0.60026047 | 1 |
| Slc13a3 | 0.9618639 | -0.056095323 | 0.60034802 | 1 |
| Dynll2 | 0.97526625 | -0.036131967 | 0.60047445 | 1 |
| Cactin | 1.04796953 | 0.067596773 | 0.60052796 | 1 |
| Rasef | 0.90818734 | -0.138938169 | 0.60074659 | 1 |
| Akr1e2 | 1.04412864 | 0.062299466 | 0.60092213 | 1 |
| Grik4 | 1.04352392 | 0.061463674 | 0.60101672 | 1 |
| Tceal1 | 1.03817939 | 0.054055753 | 0.60103539 | 1 |
| Slc16a2 | 0.92498818 | -0.112493167 | 0.6010419 | 1 |
| Tnfaip6 | 1.12076719 | 0.164486623 | 0.60110175 | 1 |
| Abhd13 | 1.04155413 | 0.05873782 | 0.60114943 | 1 |
| Ubap1 | 1.03112454 | 0.04421859 | 0.6012357 | 1 |
| Cnbp | 1.02429979 | 0.034638021 | 0.60146026 | 1 |
| Mxd1 | 0.94634044 | -0.07956882 | 0.60146407 | 1 |
| Kcnh2 | 0.97425053 | -0.037635282 | 0.60149036 | 1 |
| Rad23b | 1.02426963 | 0.034595541 | 0.60163341 | 1 |
| Taok3 | 0.95808288 | -0.061777631 | 0.60165558 | 1 |
| Cd84 | 1.09081881 | 0.125411485 | 0.60180013 | 1 |
| Cdc14a | 0.95866428 | -0.06090241 | 0.60185559 | 1 |
| Vegfa | 0.97343751 | -0.038839724 | 0.60197254 | 1 |
| Rnmt | 1.03467444 | 0.049176893 | 0.60201638 | 1 |
| Papd4 | 1.04934351 | 0.069487038 | 0.60202141 | 1 |
| Lmo7 | 0.96391572 | -0.053021086 | 0.6020625 | 1 |

| | | | | |
|---|---|---|---|---|
| Cdk1 | 1.12265958 | 0.166920531 | 0.60216422 | 1 |
| Faf1 | 1.03243247 | 0.046047424 | 0.60227571 | 1 |
| Chordc1 | 0.97485274 | -0.036743785 | 0.60228924 | 1 |
| Ggnbp2 | 1.02574716 | 0.036675158 | 0.60231547 | 1 |
| Baz2b | 1.03443104 | 0.048837474 | 0.60241934 | 1 |
| Vsig10 | 0.94874703 | -0.075904626 | 0.60243485 | 1 |
| Sox11 | 0.8853646 | -0.175656407 | 0.60243704 | 1 |
| Rsad1 | 0.90056707 | -0.151094366 | 0.6024561 | 1 |
| Tram1 | 1.02823343 | 0.040167824 | 0.60247194 | 1 |
| Lin7a | 0.92320924 | -0.115270435 | 0.60261322 | 1 |
| Hspd1 | 1.02421109 | 0.03451308 | 0.60271342 | 1 |
| Spata24 | 1.10863272 | 0.148781489 | 0.60279148 | 1 |
| Fbxo16 | 1.07227239 | 0.100671441 | 0.60281091 | 1 |
| Nol3 | 0.96584121 | -0.050142073 | 0.60284352 | 1 |
| Tox | 0.93942727 | -0.090146617 | 0.60286078 | 1 |
| Efemp2 | 0.95760664 | -0.062494936 | 0.60287149 | 1 |
| Trappc5 | 0.95397953 | -0.067969781 | 0.60292982 | 1 |
| Rpl28 | 0.97529285 | -0.03609262 | 0.60293412 | 1 |
| Elac2 | 0.96436419 | -0.052350022 | 0.60297327 | 1 |
| RGD1564450 | 1.04106727 | 0.058063298 | 0.60314708 | 1 |
| Napg | 1.02427549 | 0.03460379 | 0.60314979 | 1 |
| Farp2 | 1.03458342 | 0.049049974 | 0.60319137 | 1 |
| Mapk8 | 1.02604159 | 0.037089218 | 0.60332931 | 1 |
| Cct4 | 1.024075 | 0.034321376 | 0.60336362 | 1 |
| Morn1 | 1.08020402 | 0.111303819 | 0.60348527 | 1 |
| Vapa | 1.02407671 | 0.034323793 | 0.60353484 | 1 |
| Tex264 | 0.96963156 | -0.04449144 | 0.60372895 | 1 |
| Tcea2 | 1.03193989 | 0.045358932 | 0.60380969 | 1 |
| Aip | 0.96865522 | -0.04594484 | 0.60381146 | 1 |
| Pgm2l1 | 0.97600668 | -0.035037067 | 0.60418727 | 1 |
| Tspyl4 | 1.02424065 | 0.03455472 | 0.60422879 | 1 |
| Shfm1 | 0.96753228 | -0.047618297 | 0.60426863 | 1 |
| Tmem35 | 0.97570597 | -0.03548164 | 0.60430973 | 1 |
| St13 | 1.02402372 | 0.034249138 | 0.60432045 | 1 |
| Rnaseh1 | 1.05644167 | 0.079213118 | 0.60453018 | 1 |
| Zfp110 | 1.03229092 | 0.045849613 | 0.60455211 | 1 |
| Smc3 | 1.02789452 | 0.039692222 | 0.6046224 | 1 |
| Ubqln4 | 0.97064311 | -0.042987165 | 0.60463716 | 1 |
| Tspan17 | 0.97652605 | -0.034269559 | 0.60464887 | 1 |
| Slc39a1 | 1.03184064 | 0.045220171 | 0.60465171 | 1 |

| | | | | |
|---|---|---|---|---|
| Cnnm3 | 1.04082025 | 0.057720937 | 0.60469742 | 1 |
| Skiv2l2 | 0.97221892 | -0.040646882 | 0.60483014 | 1 |
| Ncapd3 | 1.05055825 | 0.071156151 | 0.6050181 | 1 |
| Acbd6 | 1.04544534 | 0.064117628 | 0.60504117 | 1 |
| Socs3 | 0.90027885 | -0.151556162 | 0.60505693 | 1 |
| Arhgef11 | 0.97416097 | -0.037767914 | 0.60507675 | 1 |
| Lrrc71 | 1.08715299 | 0.120554982 | 0.60513759 | 1 |
| Chn1 | 0.97454336 | -0.037201711 | 0.60527559 | 1 |
| Slitrk3 | 1.02819579 | 0.040115009 | 0.60527853 | 1 |
| Lrrc42 | 1.04297085 | 0.060698834 | 0.60531834 | 1 |
| Ift22 | 1.05006837 | 0.070483267 | 0.60532879 | 1 |
| Elac1 | 1.06298566 | 0.088122136 | 0.60535803 | 1 |
| Grxcr2 | 0.92082358 | -0.119003317 | 0.6053709 | 1 |
| Lmnb2 | 0.93248548 | -0.100846837 | 0.60544317 | 1 |
| Hdac1 | 1.03843584 | 0.054412077 | 0.60559394 | 1 |
| Ufd1l | 1.02777075 | 0.039518496 | 0.60563542 | 1 |
| Thap7 | 0.95538458 | -0.065846509 | 0.60569696 | 1 |
| Sars | 1.02391146 | 0.034090968 | 0.60570514 | 1 |
| Fkbp10 | 0.95822037 | -0.061570606 | 0.60584808 | 1 |
| Armc9 | 1.03434744 | 0.048720879 | 0.60591036 | 1 |
| Otud5 | 0.97307183 | -0.039381795 | 0.60598917 | 1 |
| Ica1 | 1.03102324 | 0.044076849 | 0.60604023 | 1 |
| Zfp36l2 | 1.03010123 | 0.042786115 | 0.6060626 | 1 |
| Cckar | 1.08000145 | 0.111033245 | 0.60620417 | 1 |
| Trrap | 0.97550126 | -0.035784356 | 0.60626732 | 1 |
| Man2a2 | 0.96834582 | -0.046405728 | 0.60628036 | 1 |
| Pin1 | 1.02477185 | 0.035302756 | 0.60647412 | 1 |
| Akap6 | 0.97603884 | -0.03498953 | 0.60648262 | 1 |
| Arf3 | 0.97652321 | -0.034273764 | 0.60659099 | 1 |
| Dlg1 | 0.96273483 | -0.054789611 | 0.60670433 | 1 |
| Tcf19 | 0.92398749 | -0.11405477 | 0.60684784 | 1 |
| Grem2 | 0.97038481 | -0.043371125 | 0.60685327 | 1 |
| Atg3 | 0.97147529 | -0.041750791 | 0.60697567 | 1 |
| Cant1 | 1.03436685 | 0.048747939 | 0.60714096 | 1 |
| Mrps26 | 0.96248662 | -0.055161604 | 0.60731769 | 1 |
| Egfl8 | 0.97468739 | -0.036988511 | 0.60734964 | 1 |
| Pde4d | 0.95435504 | -0.067402011 | 0.60735621 | 1 |
| Prkar2a | 0.96512035 | -0.051219235 | 0.60735648 | 1 |
| Llph | 1.04491154 | 0.063380818 | 0.60744526 | 1 |
| Car8 | 0.94515363 | -0.08137925 | 0.60747531 | 1 |

| | | | | |
|---|---|---|---|---|
| Tm7sf3 | 0.96504982 | -0.051324669 | 0.60756371 | 1 |
| Utp3 | 1.03369145 | 0.047805613 | 0.60759445 | 1 |
| Tmem214 | 0.96759931 | -0.047518348 | 0.60774589 | 1 |
| Smarca5 | 0.97465877 | -0.037030882 | 0.60775381 | 1 |
| Alk | 0.95969346 | -0.059354438 | 0.6078475 | 1 |
| Tmem198b | 1.0676914 | 0.094494717 | 0.60790332 | 1 |
| Timm10b | 1.0566787 | 0.07953677 | 0.60793222 | 1 |
| LOC361635 | 1.02806994 | 0.039938421 | 0.60793761 | 1 |
| Jdp2 | 1.07599591 | 0.105672599 | 0.60795598 | 1 |
| Dnaja1 | 0.97628423 | -0.034626863 | 0.60812462 | 1 |
| Etv3 | 1.06227505 | 0.087157361 | 0.60831872 | 1 |
| Gpr27 | 0.89355819 | -0.16236641 | 0.60844607 | 1 |
| Fra10ac1 | 1.04371426 | 0.061726789 | 0.60863712 | 1 |
| Lrrc36 | 1.10552149 | 0.144727066 | 0.60882374 | 1 |
| Xkr7 | 0.92977617 | -0.105044644 | 0.60887363 | 1 |
| Pet100 | 1.04451544 | 0.062833814 | 0.60888949 | 1 |
| Bcl2l1 | 0.97061438 | -0.043029859 | 0.60893845 | 1 |
| Lrrc49 | 1.02374588 | 0.033857645 | 0.60901458 | 1 |
| Adgrb3 | 0.96725675 | -0.048029207 | 0.60901625 | 1 |
| Itgbl1 | 1.02920507 | 0.04153047 | 0.60917034 | 1 |
| Tomm34 | 1.02490206 | 0.035486046 | 0.60919674 | 1 |
| Wiz | 0.96510133 | -0.051247664 | 0.60925244 | 1 |
| Dok6 | 0.93836149 | -0.091784294 | 0.60931666 | 1 |
| Rps3 | 1.02396385 | 0.034164782 | 0.60935811 | 1 |
| Fgd1 | 0.94130221 | -0.087270118 | 0.6094888 | 1 |
| Slc36a1 | 1.03154612 | 0.044808331 | 0.60956069 | 1 |
| Zfp566 | 1.11034047 | 0.151002126 | 0.60959565 | 1 |
| Rps16 | 1.02368344 | 0.033769658 | 0.60966037 | 1 |
| Cpne3 | 0.97091767 | -0.042579127 | 0.6097217 | 1 |
| Glmn | 1.04322133 | 0.061045276 | 0.60973044 | 1 |
| Wbp1l | 0.97154005 | -0.041654628 | 0.60975205 | 1 |
| Itga1 | 1.05132194 | 0.072204521 | 0.60982336 | 1 |
| Ust | 0.9587986 | -0.060700291 | 0.60983899 | 1 |
| N6amt1 | 1.07521319 | 0.104622743 | 0.60986569 | 1 |
| Otud4 | 1.03756871 | 0.053206875 | 0.60991836 | 1 |
| Ptch2 | 1.06092726 | 0.085325743 | 0.60996976 | 1 |
| Parm1 | 1.02471856 | 0.03522773 | 0.61018534 | 1 |
| Cops2 | 1.02369697 | 0.033788716 | 0.61031164 | 1 |
| Lrig2 | 0.96691442 | -0.048539885 | 0.6103918 | 1 |
| Cdc42ep5 | 1.10013554 | 0.137681275 | 0.61042906 | 1 |

| | | | | |
|---|---|---|---|---|
| Rab21 | 0.97015024 | -0.043719908 | 0.61052466 | 1 |
| Fam214b | 0.96301297 | -0.054372862 | 0.61057406 | 1 |
| Med8 | 0.91575067 | -0.126973241 | 0.61062707 | 1 |
| Ror1 | 0.92857591 | -0.106908246 | 0.61110655 | 1 |
| Nrsn2 | 1.08883907 | 0.122790735 | 0.61113297 | 1 |
| Hmces | 1.04119259 | 0.058236954 | 0.61119018 | 1 |
| Car2 | 1.02512711 | 0.035802802 | 0.6112248 | 1 |
| Flnc | 0.95425415 | -0.06755454 | 0.61124606 | 1 |
| Ahctf1 | 0.97135763 | -0.041925541 | 0.61132268 | 1 |
| LOC100910996 | 1.0434022 | 0.061295384 | 0.61134499 | 1 |
| Iqub | 0.91564866 | -0.127133956 | 0.61140514 | 1 |
| Ndst3 | 1.06123835 | 0.085748714 | 0.61141026 | 1 |
| Mrps34 | 1.04348867 | 0.061414936 | 0.61163289 | 1 |
| Ddx3 | 1.02621434 | 0.037332091 | 0.61165324 | 1 |
| Pigh | 0.94586679 | -0.080291073 | 0.61193064 | 1 |
| Dohh | 1.03711275 | 0.052572752 | 0.61194638 | 1 |
| Trnau1ap | 1.05618752 | 0.078866001 | 0.61194659 | 1 |
| Leprotl1 | 1.03588968 | 0.050870374 | 0.61205583 | 1 |
| Igfbp7 | 1.02353742 | 0.033563849 | 0.61213888 | 1 |
| Cdk8 | 0.96497267 | -0.051440005 | 0.6121496 | 1 |
| Spata6 | 0.96016718 | -0.058642468 | 0.61221417 | 1 |
| Rpl36 | 0.96872631 | -0.045838965 | 0.61226554 | 1 |
| Med7 | 1.04793668 | 0.067551553 | 0.61227535 | 1 |
| Cd81 | 1.02437258 | 0.034740542 | 0.61228229 | 1 |
| Ift52 | 1.03155716 | 0.044823764 | 0.61232821 | 1 |
| Arg2 | 1.04303938 | 0.060793632 | 0.61234609 | 1 |
| Nrn1 | 1.02371326 | 0.033811677 | 0.61256244 | 1 |
| Slc9a9 | 1.0234894 | 0.033496161 | 0.61258048 | 1 |
| Pttg1 | 1.14608344 | 0.196712085 | 0.61275866 | 1 |
| Gstm1 | 0.97185088 | -0.04119313 | 0.61284528 | 1 |
| Rala | 1.03453293 | 0.048979569 | 0.61284931 | 1 |
| Brca2 | 0.94414645 | -0.082917438 | 0.61290834 | 1 |
| Fzd3 | 1.05547402 | 0.077891072 | 0.61300897 | 1 |
| Mpdu1 | 1.02922169 | 0.041553769 | 0.61316777 | 1 |
| Nlrp1a | 1.07415833 | 0.10320666 | 0.61341525 | 1 |
| Tspan5 | 0.96831751 | -0.046447916 | 0.61342546 | 1 |
| Htr3a | 0.97602897 | -0.035004121 | 0.61349922 | 1 |
| Hnrnpf | 0.97667485 | -0.034049742 | 0.61351448 | 1 |
| Cars | 0.96562435 | -0.050466042 | 0.61363744 | 1 |
| Man2b2 | 0.96456402 | -0.052051105 | 0.61373581 | 1 |

| | | | | |
|---|---|---|---|---|
| Tmem204 | 1.05761037 | 0.080808221 | 0.61374103 | 1 |
| Eif2b4 | 1.03644139 | 0.051638538 | 0.61379518 | 1 |
| Phldb3 | 1.1254353 | 0.170483115 | 0.61392007 | 1 |
| Klhl5 | 0.96760074 | -0.047516229 | 0.61395728 | 1 |
| Nek9 | 1.02395231 | 0.034148531 | 0.61423161 | 1 |
| Cinp | 0.96218406 | -0.055615193 | 0.61454393 | 1 |
| Rpl39 | 0.9761294 | -0.034855687 | 0.61463633 | 1 |
| Pex6 | 0.95894291 | -0.060483171 | 0.61464009 | 1 |
| Eif2s1 | 0.97218958 | -0.040690416 | 0.61469954 | 1 |
| Ankrd44 | 0.95906275 | -0.060302881 | 0.61471603 | 1 |
| Pnn | 1.02689026 | 0.038282018 | 0.61477551 | 1 |
| RGD1311164 | 1.04484509 | 0.063289059 | 0.61478183 | 1 |
| Pdzd4 | 1.0379798 | 0.053778368 | 0.614812 | 1 |
| Arhgef3 | 1.03206579 | 0.045534946 | 0.6149096 | 1 |
| Nudt6 | 0.90709121 | -0.140680464 | 0.61493582 | 1 |
| Armc8 | 0.97163016 | -0.041520828 | 0.61494309 | 1 |
| Wdr6 | 0.97600852 | -0.035034357 | 0.61502342 | 1 |
| Grb10 | 0.9709724 | -0.042497811 | 0.61510568 | 1 |
| Slc10a7 | 1.06141806 | 0.085993007 | 0.61522888 | 1 |
| Cdk9 | 0.9708875 | -0.042623958 | 0.61569674 | 1 |
| Atl3 | 1.03178432 | 0.045141421 | 0.61570908 | 1 |
| Rxrb | 0.96590313 | -0.050049584 | 0.61570937 | 1 |
| Gstz1 | 1.06185259 | 0.086583506 | 0.61584797 | 1 |
| Bcl2 | 0.92217489 | -0.116887709 | 0.61591447 | 1 |
| Rad54l | 1.14320689 | 0.193086518 | 0.61593503 | 1 |
| Aoc2-ps1 | 0.90968129 | -0.136566916 | 0.61596059 | 1 |
| Rps6kb2 | 1.0474171 | 0.06683607 | 0.61601719 | 1 |
| Abcg3 | 1.07674273 | 0.106673582 | 0.61606438 | 1 |
| Syndig1l | 0.94794791 | -0.077120313 | 0.6161252 | 1 |
| Stpg1 | 0.91120575 | -0.134151249 | 0.61614862 | 1 |
| Ubl7 | 0.97044659 | -0.043279285 | 0.61626893 | 1 |
| Phkg2 | 1.02965487 | 0.042160848 | 0.61629073 | 1 |
| Srsf3 | 1.02414305 | 0.034417237 | 0.61640214 | 1 |
| Slc26a6 | 1.06027418 | 0.084437382 | 0.61644207 | 1 |
| Sec24d | 0.96375937 | -0.053255112 | 0.61646825 | 1 |
| Camk1 | 1.03777373 | 0.053491925 | 0.61646988 | 1 |
| Pop7 | 1.04461519 | 0.062971589 | 0.61647524 | 1 |
| Fam109b | 1.10900096 | 0.149260618 | 0.61651067 | 1 |
| Ccdc92 | 0.97720408 | -0.033268209 | 0.61670621 | 1 |
| Lars2 | 1.04380253 | 0.061848805 | 0.61671076 | 1 |

| | | | | |
|---|---|---|---|---|
| Ndufa5 | 0.97327854 | -0.039075351 | 0.61674926 | 1 |
| Bmp7 | 1.06185168 | 0.086582259 | 0.61691777 | 1 |
| Ppib | 0.97600443 | -0.035040402 | 0.61698705 | 1 |
| Atp2b4 | 0.96352109 | -0.053611848 | 0.6172613 | 1 |
| Rbp2 | 0.88427701 | -0.17742972 | 0.61728375 | 1 |
| Slc19a2 | 1.03524998 | 0.049979171 | 0.61738876 | 1 |
| MGC94335 | 0.95553728 | -0.065615929 | 0.61742587 | 1 |
| Sphk2 | 0.97337984 | -0.038925194 | 0.61793739 | 1 |
| Fancd2 | 1.08079528 | 0.11209328 | 0.61802396 | 1 |
| Qdpr | 1.02391418 | 0.034094806 | 0.61802437 | 1 |
| Sumo2 | 1.02409699 | 0.034352353 | 0.6180472 | 1 |
| Grwd1 | 0.95028858 | -0.07356241 | 0.6180814 | 1 |
| Rab3d | 1.02886365 | 0.041051808 | 0.61819205 | 1 |
| Spopl | 0.95314856 | -0.069226996 | 0.61820966 | 1 |
| G6pc3 | 0.97312341 | -0.039305319 | 0.6183665 | 1 |
| Yipf2 | 1.03829014 | 0.054209645 | 0.61841593 | 1 |
| Opn4 | 1.1114066 | 0.152386707 | 0.61848455 | 1 |
| Gcnt2 | 1.02652979 | 0.037775489 | 0.61854842 | 1 |
| Surf1 | 1.04118466 | 0.058225957 | 0.61876203 | 1 |
| Thbs3 | 1.05577296 | 0.078299627 | 0.61883549 | 1 |
| Plekhh3 | 0.94854616 | -0.076210108 | 0.61894165 | 1 |
| Ap1b1 | 1.02371978 | 0.03382087 | 0.61907604 | 1 |
| Tfap2a | 1.11661131 | 0.15912707 | 0.61915255 | 1 |
| Zfp710 | 1.06449496 | 0.090169124 | 0.61915419 | 1 |
| Hccs | 1.03556785 | 0.050422081 | 0.6192113 | 1 |
| Hcn4 | 0.94545559 | -0.080918402 | 0.61929181 | 1 |
| Stap1 | 1.14552532 | 0.196009352 | 0.61931361 | 1 |
| Sass6 | 1.10130787 | 0.13921783 | 0.61935643 | 1 |
| Rlbp1 | 0.89254719 | -0.163999641 | 0.61942445 | 1 |
| Efr3a | 1.02312161 | 0.032977632 | 0.61948909 | 1 |
| Catsper2 | 1.07422511 | 0.103296355 | 0.61959065 | 1 |
| Ciao1 | 1.03037159 | 0.043164721 | 0.61968535 | 1 |
| Dmtf1 | 0.97465291 | -0.037039557 | 0.6197187 | 1 |
| Srek1 | 1.04011957 | 0.056749385 | 0.61972366 | 1 |
| Cnksr2 | 1.05127162 | 0.072135471 | 0.61972579 | 1 |
| Zc3h14 | 0.97388 | -0.038184085 | 0.61977649 | 1 |
| Ly86 | 1.08660893 | 0.119832813 | 0.61979152 | 1 |
| Akirin1 | 0.97652167 | -0.034276038 | 0.61980801 | 1 |
| Ralgapb | 0.97753204 | -0.032784099 | 0.61992148 | 1 |
| Ntsr1 | 0.93313405 | -0.09984375 | 0.62021224 | 1 |

| | | | | |
|---|---|---|---|---|
| Adcy2 | 1.0399662 | 0.056536642 | 0.62022699 | 1 |
| Naa60 | 0.97176681 | -0.04131793 | 0.62023641 | 1 |
| Ehd2 | 0.96710592 | -0.048254191 | 0.62033251 | 1 |
| Rhoh | 0.88935835 | -0.169163247 | 0.6204978 | 1 |
| Gas7 | 0.97755955 | -0.032743505 | 0.6205109 | 1 |
| Ptcd2 | 1.04764945 | 0.067156068 | 0.62051154 | 1 |
| Cd8a | 0.89697111 | -0.156866584 | 0.62053685 | 1 |
| Ppp1r14a | 1.0884172 | 0.122231666 | 0.62059313 | 1 |
| Rassf1 | 0.95561003 | -0.065506101 | 0.62064907 | 1 |
| Lat2 | 0.85206816 | -0.230959245 | 0.62067034 | 1 |
| Slc7a7 | 0.97218254 | -0.040700864 | 0.62068524 | 1 |
| Tnrc6a | 1.02490617 | 0.035491839 | 0.62068716 | 1 |
| Ikbip | 1.04973847 | 0.070029938 | 0.62072577 | 1 |
| Chodl | 0.93768395 | -0.092826358 | 0.62076179 | 1 |
| B4galt7 | 0.96154367 | -0.056575713 | 0.62077226 | 1 |
| Fcho2 | 0.96256926 | -0.055037737 | 0.62091481 | 1 |
| LOC100363521 | 1.03484504 | 0.049414751 | 0.62095944 | 1 |
| LOC500475 | 1.08630472 | 0.119428851 | 0.62099265 | 1 |
| Tmx2 | 1.02294861 | 0.032733677 | 0.62102525 | 1 |
| Insc | 0.95329554 | -0.069004546 | 0.62119397 | 1 |
| Prkaa1 | 1.02366956 | 0.033750094 | 0.62128799 | 1 |
| Arhgap4 | 0.94279919 | -0.084977577 | 0.6213329 | 1 |
| Hoxb5 | 0.93264238 | -0.100604108 | 0.62134045 | 1 |
| Cul4a | 1.02744493 | 0.039061066 | 0.6213544 | 1 |
| Itga10 | 1.12262991 | 0.1668824 | 0.62136727 | 1 |
| Fermt2 | 0.97763838 | -0.032627174 | 0.62144792 | 1 |
| Lhfpl4 | 0.96769185 | -0.047380386 | 0.62146348 | 1 |
| Eif4ebp3 | 1.07776765 | 0.108046183 | 0.62146378 | 1 |
| Rnf103 | 0.97331444 | -0.039022134 | 0.62151306 | 1 |
| Srxn1 | 0.97046683 | -0.043249197 | 0.62152552 | 1 |
| Kazald1 | 0.91777551 | -0.123786789 | 0.6216722 | 1 |
| Abcd1 | 1.05764151 | 0.08085071 | 0.62169511 | 1 |
| LOC100365289 | 1.04615945 | 0.065102762 | 0.62169887 | 1 |
| Sart3 | 1.03846172 | 0.054448041 | 0.62178695 | 1 |
| Scarf1 | 0.91615167 | -0.126341636 | 0.62184723 | 1 |
| Gatm | 0.97266121 | -0.039990718 | 0.62186576 | 1 |
| Tmbim6 | 0.97514051 | -0.036317976 | 0.62189848 | 1 |
| Faap24 | 1.0971427 | 0.133751188 | 0.62193269 | 1 |
| Psmg4 | 0.94033426 | -0.088754408 | 0.62195817 | 1 |
| Kcnj10 | 0.96590806 | -0.050042221 | 0.62198873 | 1 |

| | | | | |
|---|---|---|---|---|
| Fkbp11 | 1.09095163 | 0.125587144 | 0.6220094 | 1 |
| LOC497899 | 0.91832153 | -0.122928722 | 0.62202747 | 1 |
| Phpt1 | 1.03578992 | 0.050731417 | 0.62215559 | 1 |
| Grk6 | 0.96840707 | -0.046314488 | 0.6221924 | 1 |
| Eed | 1.04072154 | 0.057584108 | 0.62223999 | 1 |
| Tmem266 | 1.11139661 | 0.15237374 | 0.62235599 | 1 |
| Hspa14 | 0.95930322 | -0.059941192 | 0.62254622 | 1 |
| Anks3 | 1.04418508 | 0.062377452 | 0.62256674 | 1 |
| Calhm2 | 0.94524036 | -0.081246869 | 0.62262109 | 1 |
| Ptgis | 1.06498856 | 0.09083793 | 0.62271796 | 1 |
| Necab2 | 0.92249174 | -0.116392097 | 0.62296314 | 1 |
| Hexb | 1.03135993 | 0.044547901 | 0.62312816 | 1 |
| Asmtl | 0.95781085 | -0.062187316 | 0.6232443 | 1 |
| Parp6 | 1.02344414 | 0.033432359 | 0.6233598 | 1 |
| Slc25a29 | 1.06011861 | 0.084225686 | 0.62349972 | 1 |
| Aasdhppt | 0.96817766 | -0.046656286 | 0.62379668 | 1 |
| Snupn | 0.95599082 | -0.064931333 | 0.6238354 | 1 |
| Ubxn4 | 1.02399205 | 0.03420451 | 0.62400114 | 1 |
| Ankrd23 | 1.15394635 | 0.206576154 | 0.62404893 | 1 |
| Cog6 | 1.03795654 | 0.053746042 | 0.62412602 | 1 |
| Uri1 | 0.96490243 | -0.051545029 | 0.62425762 | 1 |
| Zfp143 | 1.04656833 | 0.065666503 | 0.62435942 | 1 |
| Zfp36 | 1.03329937 | 0.047258294 | 0.62446517 | 1 |
| Dbt | 1.03145269 | 0.044677656 | 0.62447233 | 1 |
| Nr3c1 | 0.97605135 | -0.034971042 | 0.62448601 | 1 |
| Dnmt3a | 1.05288993 | 0.074354625 | 0.62458482 | 1 |
| Irf6 | 1.03375094 | 0.047888646 | 0.62460765 | 1 |
| Stx6 | 0.97040355 | -0.04334326 | 0.62464881 | 1 |
| Mmp9 | 0.90160978 | -0.149424933 | 0.62466579 | 1 |
| Fcf1 | 1.03612417 | 0.051196902 | 0.6247406 | 1 |
| Guf1 | 1.03310887 | 0.046992295 | 0.62477423 | 1 |
| Mtap | 1.0620673 | 0.086875182 | 0.62517324 | 1 |
| Ino80 | 0.96715592 | -0.048179604 | 0.62521223 | 1 |
| Tmem243 | 1.06116485 | 0.085648797 | 0.6253248 | 1 |
| Mef2a | 0.9730086 | -0.039475541 | 0.62538066 | 1 |
| Mfn2 | 0.97791144 | -0.03222427 | 0.62545905 | 1 |
| Wfs1 | 0.97479089 | -0.036835326 | 0.62551094 | 1 |
| Bmp15 | 0.87242602 | -0.196895291 | 0.62563603 | 1 |
| Utp14a | 0.96092078 | -0.057510601 | 0.62580293 | 1 |
| Bop1 | 1.03680238 | 0.052140938 | 0.62625673 | 1 |

| | | | | |
|---|---|---|---|---|
| Trim5 | 1.0394439 | 0.055811897 | 0.6264176 | 1 |
| Epm2a | 0.92819043 | -0.107507265 | 0.62642547 | 1 |
| Rps6ka6 | 0.93366951 | -0.099016122 | 0.62651099 | 1 |
| Lrp2 | 0.92987111 | -0.104897333 | 0.62651597 | 1 |
| Polr3g | 0.97737379 | -0.033017681 | 0.62679357 | 1 |
| Matk | 0.95450112 | -0.067181197 | 0.62683615 | 1 |
| Cfap52 | 0.88155397 | -0.181879198 | 0.62709109 | 1 |
| Hist2h2aa3 | 0.95154267 | -0.071659739 | 0.62711107 | 1 |
| Bag1 | 1.02486608 | 0.035435401 | 0.62716681 | 1 |
| Fam50a | 1.03973246 | 0.056212348 | 0.62718429 | 1 |
| Gprc5c | 1.04016837 | 0.056817069 | 0.62719048 | 1 |
| Camp | 1.09039646 | 0.124852785 | 0.62719969 | 1 |
| RGD1565616 | 1.02783491 | 0.03960856 | 0.62721243 | 1 |
| Sel1l | 0.97685495 | -0.033783734 | 0.62725327 | 1 |
| Taf5l | 0.95246797 | -0.070257512 | 0.62727158 | 1 |
| Fpr1 | 1.32438647 | 0.405324179 | 0.62728069 | 1 |
| Ceacam4 | 1.14350608 | 0.19346404 | 0.62737873 | 1 |
| Cops7b | 1.04230417 | 0.059776356 | 0.62741842 | 1 |
| Grb14 | 0.96550745 | -0.050640706 | 0.62746632 | 1 |
| Mrrf | 1.02969108 | 0.042211582 | 0.62757498 | 1 |
| Lsm6 | 1.04633623 | 0.065346526 | 0.62763217 | 1 |
| Galc | 0.94935556 | -0.074979574 | 0.627726 | 1 |
| Rfxank | 1.06666052 | 0.09310109 | 0.62784277 | 1 |
| Dxo | 0.93088509 | -0.103325002 | 0.62791671 | 1 |
| Rabgap1 | 0.97682734 | -0.033824509 | 0.62792767 | 1 |
| Cd37 | 1.06537641 | 0.091363245 | 0.62796982 | 1 |
| Trappc10 | 0.97645607 | -0.034372958 | 0.62798865 | 1 |
| Zfp503 | 0.95400741 | -0.067927623 | 0.6280408 | 1 |
| Bsg | 1.02293051 | 0.032708149 | 0.62832761 | 1 |
| Cers5 | 1.02908045 | 0.041355769 | 0.62847501 | 1 |
| Parg | 1.03214529 | 0.045646069 | 0.62852652 | 1 |
| Ddx49 | 0.96001349 | -0.058873415 | 0.62858828 | 1 |
| Med14 | 1.02849785 | 0.040538783 | 0.62864246 | 1 |
| RGD1309730 | 1.03880457 | 0.054924269 | 0.62867762 | 1 |
| Twist2 | 0.8843007 | -0.177391065 | 0.6288096 | 1 |
| S100a4 | 0.97375341 | -0.038371626 | 0.62897387 | 1 |
| Fbxw9 | 1.06340142 | 0.088686292 | 0.62903869 | 1 |
| Vps37d | 0.91302604 | -0.131272081 | 0.6291364 | 1 |
| Smurf2 | 1.03004519 | 0.042707634 | 0.62928746 | 1 |
| Amz2 | 0.97529667 | -0.036086971 | 0.629344 | 1 |

| | | | | |
|---|---|---|---|---|
| Pusl1 | 0.95484737 | -0.066657956 | 0.62943809 | 1 |
| Gstt1 | 1.07967233 | 0.110593531 | 0.62962716 | 1 |
| Trpv3 | 1.08680828 | 0.120097462 | 0.62963847 | 1 |
| Dsel | 0.96549676 | -0.05065668 | 0.62969418 | 1 |
| Tinf2 | 1.04284801 | 0.06052891 | 0.62974724 | 1 |
| Agpat5 | 0.96899519 | -0.04543859 | 0.62975797 | 1 |
| Wdfy1 | 0.96421383 | -0.052574977 | 0.62981027 | 1 |
| Blvra | 1.03485988 | 0.049435434 | 0.62988017 | 1 |
| Limk2 | 1.0363747 | 0.051545706 | 0.62993076 | 1 |
| Nrxn3 | 1.03439742 | 0.048790582 | 0.6300834 | 1 |
| Lamtor5 | 1.03415828 | 0.048457006 | 0.6301171 | 1 |
| Pdia3 | 0.97786964 | -0.032285944 | 0.6301903 | 1 |
| Idh3a | 0.97808283 | -0.031971455 | 0.63023974 | 1 |
| Nat10 | 1.04192722 | 0.059254513 | 0.63037505 | 1 |
| Wbscr17 | 0.96608858 | -0.049772618 | 0.63041171 | 1 |
| Wdr91 | 1.04011539 | 0.056743588 | 0.63043137 | 1 |
| Pgpep1 | 1.04654056 | 0.065628221 | 0.63043363 | 1 |
| Rbpj | 1.0294174 | 0.041828072 | 0.63045348 | 1 |
| Lsm3 | 0.95859197 | -0.061011248 | 0.63046429 | 1 |
| Il2rb | 0.87547289 | -0.191865598 | 0.63056391 | 1 |
| Inpp5k | 1.03185869 | 0.045245413 | 0.6310364 | 1 |
| Wipi1 | 0.96332159 | -0.053910588 | 0.63107072 | 1 |
| Sox9 | 1.10688116 | 0.146500337 | 0.63130583 | 1 |
| Ngb | 0.92260288 | -0.1162183 | 0.63135171 | 1 |
| Tmem115 | 1.03393792 | 0.048149569 | 0.63137865 | 1 |
| Agbl5 | 1.07822349 | 0.108656242 | 0.63142823 | 1 |
| Nfkb1 | 0.97048372 | -0.043224077 | 0.63165832 | 1 |
| Dusp4 | 0.91007189 | -0.135947584 | 0.6316625 | 1 |
| Inhbb | 0.956826 | -0.063671505 | 0.6317806 | 1 |
| Sat2 | 0.95522614 | -0.066085774 | 0.63186767 | 1 |
| RGD1562114 | 1.06876633 | 0.095946457 | 0.63193366 | 1 |
| Mmp14 | 1.03015005 | 0.04285449 | 0.63199889 | 1 |
| Scn4b | 0.97773593 | -0.032483227 | 0.63208085 | 1 |
| Cdkl3 | 1.07860318 | 0.109164192 | 0.63210772 | 1 |
| Agtpbp1 | 1.02271194 | 0.032399852 | 0.63211229 | 1 |
| Nadk | 1.02943401 | 0.041851354 | 0.63218403 | 1 |
| Rae1 | 0.96172277 | -0.056307017 | 0.63228642 | 1 |
| Cacfd1 | 1.06103872 | 0.085477307 | 0.63251064 | 1 |
| Gkap1 | 1.03811446 | 0.053965515 | 0.63254826 | 1 |
| Nup62 | 1.0364589 | 0.051662904 | 0.63272776 | 1 |

| | | | | | |
|---|---|---|---|---|---|
| Ptges2 | | 1.03735927 | 0.052915628 | 0.63285509 | 1 |
| Mfsd11 | | 0.9693344 | -0.044933637 | 0.63299858 | 1 |
| Clmp | | 0.97695712 | -0.033632856 | 0.63301203 | 1 |
| Ran | | 1.02208804 | 0.031519466 | 0.63301697 | 1 |
| G6pd | | 1.02354152 | 0.033569621 | 0.63304398 | 1 |
| Plgrkt | | 1.04235053 | 0.059840523 | 0.63316215 | 1 |
| Ripk1 | | 1.04143305 | 0.058570099 | 0.63351699 | 1 |
| Uchl3 | | 1.04237763 | 0.059878022 | 0.63358479 | 1 |
| Macf1 | | 0.97828473 | -0.031673668 | 0.63370336 | 1 |
| Eef1a2 | | 0.97791208 | -0.032223331 | 0.63374877 | 1 |
| Cdc14b | | 1.05798604 | 0.081320589 | 0.63377979 | 1 |
| Eef2 | | 0.97727853 | -0.033158291 | 0.63392054 | 1 |
| Metap1d | | 0.93030584 | -0.104223006 | 0.63395152 | 1 |
| Ablim3 | | 0.96623293 | -0.049557068 | 0.63411631 | 1 |
| Psme2 | | 0.95382622 | -0.068201659 | 0.6341711 | 1 |
| Dad1 | | 1.02494466 | 0.035546009 | 0.63442337 | 1 |
| Rps15a | | 0.9723298 | -0.040482359 | 0.63453167 | 1 |
| Sptbn2 | | 0.97824173 | -0.031737081 | 0.63460573 | 1 |
| Nabp1 | | 0.9361129 | -0.095245562 | 0.63463675 | 1 |
| Inip | | 0.95930047 | -0.059945324 | 0.63464072 | 1 |
| Shh | | 1.03678218 | 0.052112824 | 0.6347992 | 1 |
| Sgcd | | 0.9373956 | -0.093270071 | 0.6349035 | 1 |
| Tfeb | | 1.0570156 | 0.079996662 | 0.63495199 | 1 |
| Gprc5a | | 0.86798705 | -0.204254581 | 0.63503743 | 1 |
| | 10-Sep | 1.03504287 | 0.049690522 | 0.63525176 | 1 |
| Tmem56 | | 1.03752253 | 0.053142663 | 0.63528287 | 1 |
| Sec23b | | 1.0364644 | 0.051670571 | 0.63534824 | 1 |
| Chga | | 1.02236337 | 0.031908046 | 0.63539334 | 1 |
| Dnm3 | | 0.97829079 | -0.031664737 | 0.63540263 | 1 |
| Tdrd3 | | 0.96239977 | -0.055291797 | 0.63547799 | 1 |
| LOC688090 | | 1.04910538 | 0.069159603 | 0.63553464 | 1 |
| Ptpn13 | | 0.97427495 | -0.03759912 | 0.63566103 | 1 |
| Clk3 | | 0.96988074 | -0.044120738 | 0.63569138 | 1 |
| Msto1 | | 1.03853572 | 0.054550841 | 0.63574558 | 1 |
| Glul | | 0.97824769 | -0.031728299 | 0.63586281 | 1 |
| Abhd17c | | 0.95660088 | -0.064010978 | 0.6359819 | 1 |
| Oscp1 | | 1.04518129 | 0.063753205 | 0.63629349 | 1 |
| Mcemp1 | | 1.15311796 | 0.205540105 | 0.63649548 | 1 |
| Zfp24 | | 0.96355282 | -0.053564343 | 0.63655314 | 1 |
| Hspb8 | | 1.02191214 | 0.031271167 | 0.63682865 | 1 |

| | | | | |
|---|---|---|---|---|
| Cd151 | 1.02201465 | 0.03141587 | 0.63698576 | 1 |
| Igf2bp2 | 0.90313852 | -0.146980817 | 0.63701996 | 1 |
| Zmym6nb | 0.90771486 | -0.139688915 | 0.63703709 | 1 |
| Matr3 | 1.02197903 | 0.03136559 | 0.63711562 | 1 |
| Tenm4 | 0.95568955 | -0.065386051 | 0.63714217 | 1 |
| Gid8 | 1.03212003 | 0.045610758 | 0.63716104 | 1 |
| Arhgap8 | 0.92958643 | -0.105339091 | 0.63728585 | 1 |
| Jagn1 | 0.9698506 | -0.044165565 | 0.63731622 | 1 |
| Ndfip2 | 0.97863818 | -0.031152526 | 0.63733387 | 1 |
| Veph1 | 1.115195 | 0.157295999 | 0.63735983 | 1 |
| Cd48 | 0.97868761 | -0.031079658 | 0.63748267 | 1 |
| Kat8 | 1.0345457 | 0.048997381 | 0.63750363 | 1 |
| Arhgap11a | 0.93808007 | -0.09221703 | 0.63753374 | 1 |
| Fdx1l | 1.03776165 | 0.053475125 | 0.63753868 | 1 |
| Ssx2ip | 0.97309532 | -0.039346967 | 0.63756379 | 1 |
| Fh | 1.02478413 | 0.035320041 | 0.63772205 | 1 |
| Klhl25 | 0.94413411 | -0.082936293 | 0.6378093 | 1 |
| Tspan13 | 0.97863744 | -0.031153613 | 0.63827364 | 1 |
| Ubl5 | 1.03073073 | 0.04366749 | 0.63831604 | 1 |
| Adh1 | 0.89100547 | -0.166493806 | 0.6383603 | 1 |
| Mllt1 | 1.0393681 | 0.055706685 | 0.63872486 | 1 |
| Manba | 1.04647979 | 0.065544449 | 0.63881276 | 1 |
| Psmb1 | 1.02232312 | 0.031851255 | 0.63883587 | 1 |
| Ints10 | 1.0323914 | 0.045990029 | 0.63904387 | 1 |
| Ube2l6 | 1.03983702 | 0.056357425 | 0.63905656 | 1 |
| Pag1 | 0.91867898 | -0.122367272 | 0.63909063 | 1 |
| Cdk5 | 0.97617365 | -0.034790289 | 0.63919073 | 1 |
| Lcp2 | 1.05508203 | 0.077355176 | 0.6392581 | 1 |
| Wdr24 | 1.03578392 | 0.050723067 | 0.63932492 | 1 |
| Gfm2 | 0.96342694 | -0.053752828 | 0.63933517 | 1 |
| Mylpf | 0.90628054 | -0.141970388 | 0.63954987 | 1 |
| Gbas | 1.03159058 | 0.044870503 | 0.63970887 | 1 |
| Lyar | 1.04654912 | 0.065640028 | 0.63976292 | 1 |
| RGD1308134 | 1.05198899 | 0.073119607 | 0.63982965 | 1 |
| Zfp706 | 0.97089498 | -0.042612841 | 0.63988808 | 1 |
| Cox7a2 | 0.97584502 | -0.035276047 | 0.63996789 | 1 |
| Fam212a | 1.08621728 | 0.119312722 | 0.64000171 | 1 |
| Spata33 | 1.07244339 | 0.100901494 | 0.64005821 | 1 |
| Pfdn5 | 0.97423975 | -0.037651244 | 0.6400864 | 1 |
| Eif2ak3 | 0.97107104 | -0.042351252 | 0.64010247 | 1 |

| | | | | |
|---|---|---|---|---|
| Vhl | 0.97050407 | -0.043193831 | 0.64012446 | 1 |
| Eya3 | 0.97290033 | -0.039636085 | 0.6401565 | 1 |
| Slc2a8 | 1.03309343 | 0.046970738 | 0.64017622 | 1 |
| Zfp507 | 1.04054305 | 0.057336649 | 0.64031912 | 1 |
| Foxk1 | 0.92973252 | -0.10511237 | 0.64043313 | 1 |
| Clec10a | 1.0416584 | 0.058882234 | 0.64045664 | 1 |
| Prrg1 | 1.04357072 | 0.061528367 | 0.64053544 | 1 |
| Mboat7 | 0.96137846 | -0.056823616 | 0.6406149 | 1 |
| Mboat7l1 | 0.96137846 | -0.056823616 | 0.6406149 | 1 |
| Wdr5b | 1.07998397 | 0.111009894 | 0.64076302 | 1 |
| Clstn2 | 0.94965069 | -0.074531155 | 0.64096136 | 1 |
| Trim41 | 0.96947105 | -0.044730282 | 0.64108075 | 1 |
| Trappc2 | 1.02193167 | 0.031298731 | 0.64120536 | 1 |
| Tbx2 | 0.95291605 | -0.069578976 | 0.64130251 | 1 |
| Rab26 | 1.09284098 | 0.128083484 | 0.64137207 | 1 |
| Cyp4f5 | 1.0636098 | 0.088968973 | 0.64160041 | 1 |
| Rock1 | 0.97312443 | -0.039303807 | 0.64164349 | 1 |
| Lpin3 | 0.93906474 | -0.090703473 | 0.64170269 | 1 |
| Nenf | 1.03600337 | 0.0510287 | 0.64194273 | 1 |
| Ap5s1 | 0.92264061 | -0.116159295 | 0.64200225 | 1 |
| Rap1gds1 | 0.97832446 | -0.031615081 | 0.64213458 | 1 |
| Bsn | 0.97385606 | -0.038219549 | 0.64215533 | 1 |
| Ppfia4 | 0.96956773 | -0.044586412 | 0.64215582 | 1 |
| Tnfsf13 | 0.9440409 | -0.083078724 | 0.64221761 | 1 |
| Ciita | 0.91939405 | -0.121244758 | 0.64232536 | 1 |
| Wipi2 | 0.96687126 | -0.048604285 | 0.64234273 | 1 |
| Cth | 1.08328538 | 0.115413356 | 0.64239149 | 1 |
| Slc15a2 | 1.04034072 | 0.057056099 | 0.64239484 | 1 |
| Tstd2 | 1.04736328 | 0.066761931 | 0.64242084 | 1 |
| Fbxo32 | 0.90375845 | -0.145990863 | 0.64265228 | 1 |
| Tmem179 | 1.02238624 | 0.031940322 | 0.64266279 | 1 |
| Gal3st1 | 1.05611588 | 0.078768141 | 0.64268963 | 1 |
| Twistnb | 1.04316239 | 0.060963757 | 0.6426916 | 1 |
| Fam83h | 0.95386226 | -0.068147149 | 0.64279642 | 1 |
| Tsc22d4 | 0.97079192 | -0.042765991 | 0.64287473 | 1 |
| Crlf3 | 0.9689228 | -0.045546371 | 0.64292058 | 1 |
| Pepd | 0.97365119 | -0.038523071 | 0.64333145 | 1 |
| Pafah1b2 | 1.02184021 | 0.031169613 | 0.64345467 | 1 |
| Bad | 1.03820567 | 0.054092275 | 0.64358287 | 1 |
| Wnt2b | 1.07372028 | 0.102618201 | 0.64366343 | 1 |

| | | | | | |
|---|---|---|---|---|---|
| | 11-Sep | 1.02882106 | 0.040992081 | 0.64383357 | 1 |
| Jmjd8 | | 0.96763697 | -0.047462209 | 0.64400051 | 1 |
| Notch2 | | 0.97387327 | -0.038194055 | 0.64400994 | 1 |
| Kcnj2 | | 1.11036971 | 0.151040124 | 0.64412246 | 1 |
| Map3k14 | | 1.06483096 | 0.090624419 | 0.64415675 | 1 |
| Tnfaip1 | | 1.02332992 | 0.033271339 | 0.64418361 | 1 |
| Tars | | 0.97453669 | -0.037211588 | 0.64419183 | 1 |
| Ercc6 | | 0.96342938 | -0.053749179 | 0.64425957 | 1 |
| Rbck1 | | 1.03003538 | 0.042693894 | 0.64428836 | 1 |
| Wdr1 | | 0.97909781 | -0.030475107 | 0.64430448 | 1 |
| Slbp | | 1.02898976 | 0.041228625 | 0.64431171 | 1 |
| Apaf1 | | 1.04512461 | 0.063674959 | 0.64435935 | 1 |
| Sipa1l1 | | 0.97356615 | -0.038649086 | 0.6444044 | 1 |
| Dennd6b | | 0.95279763 | -0.069758273 | 0.64441132 | 1 |
| Slc25a4 | | 1.02221899 | 0.031704304 | 0.64451301 | 1 |
| Tep1 | | 0.97300832 | -0.039475946 | 0.644536 | 1 |
| Ahi1 | | 0.96750435 | -0.047659949 | 0.64476878 | 1 |
| Pf4 | | 1.09610783 | 0.132389732 | 0.64479605 | 1 |
| Bmp3 | | 1.12166995 | 0.165648228 | 0.64486391 | 1 |
| Lpcat2 | | 1.04496351 | 0.063452563 | 0.64488091 | 1 |
| Nradd | | 0.90121511 | -0.150056592 | 0.64493752 | 1 |
| Kars | | 1.02300519 | 0.032813468 | 0.64506757 | 1 |
| Tmem98 | | 1.05854336 | 0.082080366 | 0.6450894 | 1 |
| Slc39a8 | | 1.03854112 | 0.054558344 | 0.64546228 | 1 |
| Ercc6l | | 0.88327872 | -0.179059334 | 0.64547055 | 1 |
| Nrbp1 | | 1.02286865 | 0.032620895 | 0.64551007 | 1 |
| Sigirr | | 1.10049977 | 0.138158847 | 0.6455353 | 1 |
| Spred1 | | 1.04202165 | 0.059385253 | 0.64579532 | 1 |
| Gimap5 | | 0.924129 | -0.113833841 | 0.64579836 | 1 |
| Thsd7b | | 1.04069541 | 0.057547878 | 0.64599014 | 1 |
| Sppl2a | | 0.95877771 | -0.060731723 | 0.64612889 | 1 |
| Actr2 | | 0.97912715 | -0.030431872 | 0.64615088 | 1 |
| Rnf187 | | 0.9790017 | -0.030616726 | 0.64618773 | 1 |
| Nemp1 | | 1.09831989 | 0.135298313 | 0.64623328 | 1 |
| Pcmtd1 | | 0.97833478 | -0.031599863 | 0.64627199 | 1 |
| Epb41l3 | | 1.02234209 | 0.031878018 | 0.64630391 | 1 |
| Atp6v0d1 | | 1.02146445 | 0.030638991 | 0.64637118 | 1 |
| Ikbkb | | 1.03421829 | 0.048540718 | 0.64637653 | 1 |
| Pea15 | | 0.97840063 | -0.031502758 | 0.64639556 | 1 |
| Sfxn4 | | 1.04526439 | 0.063867905 | 0.64669861 | 1 |

| | | | | |
|---|---|---|---|---|
| Tprkb | 0.95872783 | -0.060806786 | 0.64676004 | 1 |
| Abca1 | 0.97609168 | -0.034911434 | 0.64679185 | 1 |
| Atg10 | 1.0696604 | 0.09715284 | 0.64682109 | 1 |
| Ocrl | 1.02912998 | 0.041425207 | 0.64692027 | 1 |
| Lancl1 | 0.97784375 | -0.032324138 | 0.64699502 | 1 |
| Trit1 | 0.96138626 | -0.056811913 | 0.64707863 | 1 |
| Ldah | 0.96919201 | -0.045145585 | 0.64708694 | 1 |
| Negr1 | 1.05795671 | 0.081280591 | 0.64723511 | 1 |
| LOC100909539 | 0.93492763 | -0.097073393 | 0.64740938 | 1 |
| Cyp2u1 | 0.9485324 | -0.076231045 | 0.64754239 | 1 |
| Hcn3 | 0.95003759 | -0.0739435 | 0.64759411 | 1 |
| Popdc2 | 0.90797364 | -0.139277679 | 0.64759899 | 1 |
| Srp9 | 1.02671235 | 0.038032049 | 0.64766876 | 1 |
| Imp3 | 1.03860396 | 0.054645635 | 0.64772185 | 1 |
| Tom1l1 | 0.96517429 | -0.051138604 | 0.64773629 | 1 |
| Gpalpp1 | 0.95849875 | -0.061151548 | 0.6477818 | 1 |
| Nup37 | 1.05394854 | 0.075804431 | 0.64798308 | 1 |
| Bach2 | 1.06467007 | 0.090406427 | 0.64799494 | 1 |
| Dtx2 | 1.05016205 | 0.070611964 | 0.64804009 | 1 |
| Dnajb3 | 0.92993506 | -0.10479812 | 0.64811675 | 1 |
| RGD1562608 | 0.94156282 | -0.086870743 | 0.64812354 | 1 |
| Ppp2r2b | 1.02158697 | 0.030812031 | 0.64816149 | 1 |
| Mdm4 | 0.96402423 | -0.05285868 | 0.64816735 | 1 |
| Abcc9 | 1.05003437 | 0.070436546 | 0.64821334 | 1 |
| Cd44 | 0.97870524 | -0.031053672 | 0.64823126 | 1 |
| Grid2ip | 0.93226001 | -0.10119572 | 0.64829873 | 1 |
| Tmem177 | 0.93629495 | -0.094965019 | 0.64834653 | 1 |
| Tubb3 | 1.02273446 | 0.032431609 | 0.64838898 | 1 |
| Kcnb2 | 1.03611878 | 0.051189404 | 0.64846939 | 1 |
| Bard1 | 0.89061059 | -0.167133323 | 0.64857118 | 1 |
| Glrx2 | 1.03395401 | 0.048172018 | 0.64857328 | 1 |
| Sec23ip | 1.0298054 | 0.042371734 | 0.64858451 | 1 |
| Scml4 | 0.9649973 | -0.051403184 | 0.64871001 | 1 |
| Fam229b | 1.06714598 | 0.093757539 | 0.64885891 | 1 |
| Tdrp | 0.97384196 | -0.038240424 | 0.64895925 | 1 |
| Ntng1 | 1.0267973 | 0.038151409 | 0.64895936 | 1 |
| Zmpste24 | 1.02813376 | 0.04002797 | 0.64904462 | 1 |
| Bcat1 | 0.97825587 | -0.031716229 | 0.64916206 | 1 |
| Parp9 | 1.03480592 | 0.049360219 | 0.64940341 | 1 |
| Dcaf8 | 1.02421378 | 0.034516868 | 0.64945453 | 1 |

| | | | | |
|---|---|---|---|---|
| Oma1 | 1.07625913 | 0.106025473 | 0.64965169 | 1 |
| Ric3 | 1.02660675 | 0.037883648 | 0.64981454 | 1 |
| Osbpl5 | 0.97331157 | -0.039026397 | 0.64997346 | 1 |
| Srrm1 | 0.9764585 | -0.034369368 | 0.64997561 | 1 |
| Hdac2 | 0.97711469 | -0.033400189 | 0.65002117 | 1 |
| Grin1 | 0.97793757 | -0.03218573 | 0.65002223 | 1 |
| Foxj3 | 0.9735514 | -0.038670938 | 0.65002745 | 1 |
| Ing5 | 1.05683349 | 0.079748086 | 0.65011274 | 1 |
| Fxyd2 | 1.02785215 | 0.039632762 | 0.65013511 | 1 |
| Hecw1 | 1.0228779 | 0.032633945 | 0.65014822 | 1 |
| Tob2 | 0.95682194 | -0.063677628 | 0.65015575 | 1 |
| Tefm | 1.06442255 | 0.090070975 | 0.65017878 | 1 |
| Zbtb5 | 0.95701266 | -0.063390082 | 0.65023043 | 1 |
| Ttc12 | 0.93230172 | -0.101131167 | 0.65030445 | 1 |
| Hnrnph1 | 1.02104798 | 0.030050668 | 0.65032284 | 1 |
| Rps8 | 0.97868721 | -0.031080254 | 0.65036975 | 1 |
| Phldb1 | 0.97659547 | -0.03416701 | 0.65037204 | 1 |
| Keap1 | 0.97301471 | -0.039466479 | 0.6503848 | 1 |
| Wnt11 | 0.90449404 | -0.144817099 | 0.65047678 | 1 |
| Aldh5a1 | 1.0323285 | 0.045902121 | 0.65047911 | 1 |
| Vom2r44 | 1.10082957 | 0.138591127 | 0.65049254 | 1 |
| Fam53a | 0.92433539 | -0.113511677 | 0.65068693 | 1 |
| Nrxn2 | 0.97796356 | -0.032147383 | 0.65072712 | 1 |
| Pias1 | 0.96701272 | -0.048393228 | 0.65100357 | 1 |
| Ago2 | 0.9550526 | -0.066347901 | 0.65109397 | 1 |
| Alg9 | 1.03435397 | 0.048729978 | 0.65112331 | 1 |
| Thumpd1 | 0.97275158 | -0.039856683 | 0.65117964 | 1 |
| Lrrc20 | 1.04570533 | 0.064476377 | 0.65118388 | 1 |
| Uchl1 | 1.02239304 | 0.031949925 | 0.65120485 | 1 |
| Gys1 | 0.97530476 | -0.036075001 | 0.65127591 | 1 |
| Mier3 | 1.03588887 | 0.050869243 | 0.65143508 | 1 |
| Lingo4 | 0.94566801 | -0.080594306 | 0.65148047 | 1 |
| Vps50 | 0.97904472 | -0.03055334 | 0.65162787 | 1 |
| Siae | 0.96682309 | -0.048676169 | 0.65168305 | 1 |
| Reep3 | 0.97102611 | -0.042418005 | 0.65172006 | 1 |
| Clybl | 1.04239226 | 0.059898282 | 0.652108 | 1 |
| Pecr | 1.04513663 | 0.063691552 | 0.6521422 | 1 |
| Cep83os | 1.05150263 | 0.072452463 | 0.65218456 | 1 |
| Pcyt1b | 0.97316197 | -0.039248156 | 0.65241598 | 1 |
| Oas1b | 1.09425039 | 0.129942902 | 0.6524567 | 1 |

| | | | | |
|---|---|---|---|---|
| Spice1 | 1.06024979 | 0.084404196 | 0.65253567 | 1 |
| Extl3 | 0.97393301 | -0.03810555 | 0.65257001 | 1 |
| Rilp | 1.09197097 | 0.126934498 | 0.6525748 | 1 |
| Ola1 | 1.02112518 | 0.030159739 | 0.65258587 | 1 |
| Arv1 | 1.04531795 | 0.06394183 | 0.65259105 | 1 |
| Ypel4 | 1.03713609 | 0.052605214 | 0.65267506 | 1 |
| Ankrd49 | 1.03913475 | 0.055382752 | 0.65285469 | 1 |
| Hspa4 | 0.97939025 | -0.030044261 | 0.65298857 | 1 |
| Ncam1 | 0.97773451 | -0.032485322 | 0.65299864 | 1 |
| Bap1 | 0.97894077 | -0.030706516 | 0.65320124 | 1 |
| Actl6b | 0.96022851 | -0.058550324 | 0.65323912 | 1 |
| Uvssa | 0.92526882 | -0.112055516 | 0.6534199 | 1 |
| Dnajc5 | 0.97297573 | -0.039524283 | 0.65343132 | 1 |
| Pyroxd2 | 0.95146648 | -0.071775265 | 0.65353906 | 1 |
| Zfp217 | 1.06762243 | 0.094401519 | 0.65361126 | 1 |
| Acss2 | 1.02240867 | 0.031971972 | 0.65376738 | 1 |
| Slc7a6 | 1.03002574 | 0.042680389 | 0.65380295 | 1 |
| Il17rd | 1.08452544 | 0.117063893 | 0.65381288 | 1 |
| Il10rb | 1.04047107 | 0.057236852 | 0.65389343 | 1 |
| Tmem206 | 0.95572332 | -0.065335068 | 0.65390753 | 1 |
| Patl1 | 0.96816662 | -0.046672734 | 0.65407541 | 1 |
| Tsfm | 1.04065624 | 0.057493585 | 0.65411573 | 1 |
| Maff | 0.93855842 | -0.091481541 | 0.65412999 | 1 |
| Sh2b1 | 0.96956847 | -0.044585318 | 0.65413962 | 1 |
| Clmn | 0.95993993 | -0.058983971 | 0.65420862 | 1 |
| Dapk2 | 0.93395288 | -0.098578324 | 0.65421625 | 1 |
| Jmjd7 | 0.90801479 | -0.139212303 | 0.654345 | 1 |
| Eml4 | 0.97536235 | -0.035989818 | 0.65446245 | 1 |
| Cbln2 | 0.97656421 | -0.034213186 | 0.65446736 | 1 |
| RGD1310110 | 0.97515239 | -0.036300407 | 0.65464843 | 1 |
| Mlh1 | 0.95634441 | -0.064397819 | 0.65465684 | 1 |
| Smim7 | 1.02560447 | 0.036474449 | 0.65475646 | 1 |
| Trnt1 | 0.97168801 | -0.041434923 | 0.65506395 | 1 |
| Alg8 | 0.94970263 | -0.074452247 | 0.65506727 | 1 |
| Nfatc2 | 1.08133303 | 0.112810909 | 0.65522312 | 1 |
| Aplnr | 1.07175753 | 0.099978558 | 0.65530139 | 1 |
| Polr2j | 1.03571426 | 0.050626042 | 0.65534445 | 1 |
| Slc29a3 | 1.05478784 | 0.076952848 | 0.6554038 | 1 |
| RGD1308106 | 0.94605316 | -0.080006848 | 0.65551921 | 1 |
| Mid1ip1 | 0.97950622 | -0.029873437 | 0.65552086 | 1 |

| | | | | |
|---|---|---|---|---|
| Capn6 | 0.94141921 | -0.087090801 | 0.65579764 | 1 |
| Isy1 | 0.95724969 | -0.063032814 | 0.65589431 | 1 |
| Enpp2 | 1.03285015 | 0.04663096 | 0.65591577 | 1 |
| Slx1b | 1.0371438 | 0.052615943 | 0.65610186 | 1 |
| Musk | 1.10948703 | 0.149892807 | 0.65610562 | 1 |
| Dock11 | 0.97970842 | -0.029575661 | 0.65622704 | 1 |
| Vwc2l | 1.04514498 | 0.063703079 | 0.65624122 | 1 |
| Thap6 | 1.06768989 | 0.094492684 | 0.6562675 | 1 |
| Fxyd3 | 1.08813174 | 0.121853228 | 0.65644296 | 1 |
| Hdhd3 | 0.93712987 | -0.093679104 | 0.65655623 | 1 |
| Cyp3a9 | 0.92814532 | -0.107577393 | 0.65661985 | 1 |
| Mrps9 | 0.96965786 | -0.044452306 | 0.65675473 | 1 |
| Fxyd6 | 1.02090946 | 0.029854921 | 0.65687594 | 1 |
| Ctsd | 1.02134259 | 0.030466867 | 0.65689035 | 1 |
| Rpl36a | 0.96462591 | -0.051958536 | 0.65689865 | 1 |
| Prrg4 | 0.94813667 | -0.076833064 | 0.656921 | 1 |
| Tgfb2 | 0.9620982 | -0.055743942 | 0.65696616 | 1 |
| Clip4 | 0.96911574 | -0.045259127 | 0.65715432 | 1 |
| Prdm11 | 1.09486364 | 0.130751206 | 0.65733458 | 1 |
| Pddc1 | 0.96292822 | -0.054499831 | 0.65735604 | 1 |
| Ggact | 1.04468507 | 0.063068098 | 0.65748361 | 1 |
| RGD1565002 | 1.02739578 | 0.03899205 | 0.65775747 | 1 |
| Gfpt2 | 1.03301599 | 0.046862581 | 0.65786195 | 1 |
| Epn3 | 0.97703618 | -0.033516115 | 0.65787322 | 1 |
| Fam227a | 1.0711915 | 0.099216423 | 0.65789988 | 1 |
| Ywhab | 0.97902305 | -0.030585273 | 0.65801293 | 1 |
| Pkig | 1.03151781 | 0.04476873 | 0.65806599 | 1 |
| Fam91a1 | 0.97891385 | -0.03074619 | 0.65808238 | 1 |
| Ssc5d | 1.03792746 | 0.053705617 | 0.65812546 | 1 |
| Brd2 | 0.97912103 | -0.030440885 | 0.65817259 | 1 |
| Lmbr1 | 0.9653998 | -0.050801563 | 0.65818328 | 1 |
| Ubn2 | 1.03812689 | 0.053982801 | 0.65819298 | 1 |
| Fam96b | 1.03253518 | 0.046190937 | 0.65820146 | 1 |
| Tmco4 | 0.94221196 | -0.085876456 | 0.65832067 | 1 |
| Exoc6 | 0.97219286 | -0.040685557 | 0.65841647 | 1 |
| Gatc | 0.96903146 | -0.045384593 | 0.65843547 | 1 |
| Gpr150 | 1.1112221 | 0.152147197 | 0.65849128 | 1 |
| Acap1 | 0.936127 | -0.095223831 | 0.6587299 | 1 |
| Nfs1 | 1.02537631 | 0.036153477 | 0.65873613 | 1 |
| Paip1 | 0.97671254 | -0.033994072 | 0.65879835 | 1 |

| | | | | |
|---|---|---|---|---|
| B4galnt3 | 0.90729909 | -0.140349881 | 0.65882894 | 1 |
| Cdh20 | 1.07043932 | 0.098203013 | 0.65897039 | 1 |
| Kidins220 | 0.97959472 | -0.029743104 | 0.6590471 | 1 |
| Pcdhb20 | 1.07265362 | 0.101184281 | 0.65907999 | 1 |
| Dcp2 | 1.04998111 | 0.07036337 | 0.65937148 | 1 |
| Tgfb3 | 0.97083081 | -0.042708196 | 0.6594194 | 1 |
| Snrpf | 1.05158594 | 0.072566762 | 0.65963609 | 1 |
| Cdc34 | 0.97237293 | -0.040418359 | 0.65976128 | 1 |
| Eef1akmt1 | 0.96038467 | -0.058315725 | 0.65981971 | 1 |
| Aco2 | 1.02081056 | 0.02971516 | 0.65981972 | 1 |
| Smurf1 | 0.97444603 | -0.037345806 | 0.65996338 | 1 |
| Tubb2a | 1.02129927 | 0.030405682 | 0.66002456 | 1 |
| Taf1d | 1.05535561 | 0.077729214 | 0.6601221 | 1 |
| Rpl7 | 0.97998467 | -0.029168909 | 0.66017809 | 1 |
| Manbal | 1.02774118 | 0.039476988 | 0.66024391 | 1 |
| Emd | 0.96090188 | -0.057538971 | 0.66034371 | 1 |
| Arih2 | 0.97464812 | -0.037046646 | 0.66045504 | 1 |
| Col13a1 | 0.93076273 | -0.103514646 | 0.66052507 | 1 |
| Luzp1 | 0.97976111 | -0.029498072 | 0.66067217 | 1 |
| U2af1l4 | 1.03626794 | 0.051397074 | 0.66067635 | 1 |
| Armt1 | 1.0317583 | 0.045105046 | 0.6607212 | 1 |
| Sipa1 | 0.96638707 | -0.049326939 | 0.66075376 | 1 |
| Ephb3 | 0.93761092 | -0.092938728 | 0.66078261 | 1 |
| Ppcdc | 1.04526463 | 0.063868232 | 0.66084172 | 1 |
| Mrps12 | 1.03857995 | 0.05461228 | 0.66089373 | 1 |
| Lrp1b | 1.04192718 | 0.059254446 | 0.66091808 | 1 |
| Faap20 | 1.0553409 | 0.077709095 | 0.66099824 | 1 |
| Slc25a34 | 1.11228 | 0.153520007 | 0.66109278 | 1 |
| Exoc3 | 0.97730667 | -0.033116758 | 0.66112675 | 1 |
| Zfand2a | 0.97284524 | -0.039717777 | 0.66115909 | 1 |
| Mgme1 | 1.07999827 | 0.111029 | 0.66123391 | 1 |
| Ftx | 0.93327837 | -0.099620637 | 0.66123901 | 1 |
| Mea1 | 1.02653707 | 0.037785724 | 0.66128376 | 1 |
| Zfp638 | 0.97928415 | -0.030200558 | 0.66130496 | 1 |
| Sulf2 | 1.02263809 | 0.03229567 | 0.66136807 | 1 |
| Pcca | 0.96490804 | -0.051536645 | 0.66141527 | 1 |
| Txn2 | 0.97416532 | -0.037761471 | 0.66147393 | 1 |
| Zhx1 | 1.02123768 | 0.030318674 | 0.66149205 | 1 |
| Wdr35 | 0.9689662 | -0.045481752 | 0.66155713 | 1 |
| Cntrob | 1.07490198 | 0.104205112 | 0.66161244 | 1 |

| | | | | |
|---|---|---|---|---|
| Trim24 | 1.0428501 | 0.060531801 | 0.66161616 | 1 |
| Dnttip1 | 1.03933558 | 0.055661541 | 0.66164272 | 1 |
| Acot8 | 1.05009769 | 0.070523548 | 0.6616468 | 1 |
| Atp5f1 | 1.0204227 | 0.0291669 | 0.66176002 | 1 |
| Pde3b | 1.03702074 | 0.052444748 | 0.6618283 | 1 |
| Cd200 | 0.97951033 | -0.029867384 | 0.6619444 | 1 |
| Gpr63 | 0.91821912 | -0.123089615 | 0.66197415 | 1 |
| Zranb2 | 1.02074224 | 0.029618604 | 0.6620049 | 1 |
| Zbtb8a | 1.07337259 | 0.102150948 | 0.66218236 | 1 |
| Acaca | 0.97913331 | -0.030422799 | 0.66218459 | 1 |
| Ptpn21 | 1.0418947 | 0.059209474 | 0.66219853 | 1 |
| Lrrc73 | 0.94464781 | -0.082151533 | 0.66241959 | 1 |
| Amdhd2 | 1.05012886 | 0.070566367 | 0.66246011 | 1 |
| Gcsh | 1.02813559 | 0.040030536 | 0.66263082 | 1 |
| Mapk1ip1 | 0.96247309 | -0.055181888 | 0.66265531 | 1 |
| Rab9a | 0.96493352 | -0.051498542 | 0.66277412 | 1 |
| Rps6kb1 | 0.97360525 | -0.03859115 | 0.66278649 | 1 |
| Impad1 | 0.97959194 | -0.029747184 | 0.66278922 | 1 |
| Exoc1 | 1.02431864 | 0.034664574 | 0.66279435 | 1 |
| Nop56 | 1.02355568 | 0.03358959 | 0.66304363 | 1 |
| Cep76 | 1.04845374 | 0.068263214 | 0.66306121 | 1 |
| Fam193b | 1.030382 | 0.043179291 | 0.66310496 | 1 |
| Churc1 | 1.03712013 | 0.052583015 | 0.6631084 | 1 |
| Rrp8 | 0.95055803 | -0.073153396 | 0.66311053 | 1 |
| Mettl14 | 1.02926559 | 0.041615299 | 0.66334615 | 1 |
| Dkc1 | 0.96724773 | -0.048042663 | 0.66335508 | 1 |
| Ube3b | 1.02052689 | 0.029314195 | 0.66341858 | 1 |
| Otub2 | 0.94471414 | -0.08205025 | 0.66355526 | 1 |
| Ttc8 | 0.94436655 | -0.082581148 | 0.66369394 | 1 |
| Zfp513 | 1.03869453 | 0.054771431 | 0.66372856 | 1 |
| Ifi35 | 1.0560981 | 0.078743849 | 0.66376262 | 1 |
| Daam1 | 0.97588838 | -0.03521195 | 0.66382916 | 1 |
| Fmod | 1.03348611 | 0.047519001 | 0.6638634 | 1 |
| Rabgef1 | 1.02996343 | 0.042593112 | 0.66389257 | 1 |
| Rpusd1 | 1.04448155 | 0.062787004 | 0.664013 | 1 |
| Spns2 | 1.03625727 | 0.051382217 | 0.66441822 | 1 |
| Fnbp1l | 1.05772 | 0.080957765 | 0.6645605 | 1 |
| Zfp148 | 0.9727658 | -0.039835591 | 0.66499954 | 1 |
| Ccdc51 | 0.95416498 | -0.06768935 | 0.6650431 | 1 |
| Alas2 | 0.97168899 | -0.041433478 | 0.66505741 | 1 |

| | | | | |
|---|---|---|---|---|
| Fam216a | 1.02932223 | 0.041694687 | 0.66506108 | 1 |
| RGD1307155 | 0.92182452 | -0.117435953 | 0.66513993 | 1 |
| Ctdp1 | 0.96466641 | -0.051897967 | 0.66515533 | 1 |
| Fam134c | 0.97065278 | -0.042972787 | 0.66521165 | 1 |
| Mapre3 | 0.97967184 | -0.029629524 | 0.66527359 | 1 |
| Erh | 1.02660618 | 0.037882857 | 0.66572286 | 1 |
| Phyhd1 | 0.93071796 | -0.103584053 | 0.66572468 | 1 |
| St3gal3 | 1.06028392 | 0.084450636 | 0.66601767 | 1 |
| Catsperg1 | 1.06055658 | 0.084821588 | 0.66616928 | 1 |
| Uqcrq | 1.02404912 | 0.034284918 | 0.66626467 | 1 |
| Atp6v1a | 1.02038513 | 0.029113774 | 0.66628699 | 1 |
| Golga3 | 0.97671598 | -0.033989001 | 0.66635925 | 1 |
| Ttc7a | 1.04085423 | 0.057768038 | 0.66636511 | 1 |
| Osgep | 1.05661186 | 0.07944551 | 0.66648937 | 1 |
| Nhlrc1 | 1.0747058 | 0.103941772 | 0.66653747 | 1 |
| MGC105649 | 1.09769559 | 0.13447802 | 0.66656994 | 1 |
| Calb1 | 1.0556592 | 0.078144164 | 0.66659558 | 1 |
| Tdp1 | 0.94412113 | -0.082956127 | 0.66668645 | 1 |
| Nek1 | 0.9763908 | -0.034469392 | 0.66669657 | 1 |
| Zfp131 | 1.02949462 | 0.041936294 | 0.66672003 | 1 |
| Ube2i | 0.97802156 | -0.032061824 | 0.66682425 | 1 |
| Slc25a2 | 0.92147721 | -0.117979616 | 0.66685853 | 1 |
| Pard3b | 0.93911828 | -0.090621225 | 0.66700307 | 1 |
| Gli2 | 0.93820107 | -0.092030942 | 0.66701943 | 1 |
| Srgap1 | 0.96574471 | -0.050286225 | 0.66713911 | 1 |
| Med31 | 0.95980056 | -0.059193444 | 0.66714044 | 1 |
| Kcnmb4 | 1.0613142 | 0.085851823 | 0.66725508 | 1 |
| Pogk | 0.97684445 | -0.033799246 | 0.66728865 | 1 |
| Slc6a12 | 0.90772115 | -0.139678924 | 0.66735103 | 1 |
| Scn5a | 0.93965824 | -0.089791962 | 0.66751849 | 1 |
| Tbc1d5 | 1.02950522 | 0.041951146 | 0.66757645 | 1 |
| Enho | 1.03760478 | 0.053257037 | 0.66770748 | 1 |
| Blk | 0.87490973 | -0.192793914 | 0.66773174 | 1 |
| Camsap1 | 1.02232055 | 0.031847627 | 0.66775656 | 1 |
| Zdhhc20 | 1.03281948 | 0.046588116 | 0.66788201 | 1 |
| Flad1 | 1.02869512 | 0.040815469 | 0.66788957 | 1 |
| Bex4 | 0.95459124 | -0.067044999 | 0.66790089 | 1 |
| Dis3 | 1.03744557 | 0.053035646 | 0.66801545 | 1 |
| Phb | 1.02177703 | 0.031080416 | 0.66804211 | 1 |
| Ermn | 1.08738255 | 0.120859582 | 0.66813103 | 1 |

| | | | | |
|---|---|---|---|---|
| S100a6 | 1.01997125 | 0.028528488 | 0.66816839 | 1 |
| Dnajc8 | 1.02165643 | 0.030910118 | 0.66828117 | 1 |
| Abcg3l2 | 0.92008954 | -0.120153822 | 0.6682997 | 1 |
| Slc25a40 | 1.04305392 | 0.060813745 | 0.66833729 | 1 |
| Zfat | 0.9614547 | -0.056709203 | 0.66837864 | 1 |
| Gabrg1 | 1.02430249 | 0.034641818 | 0.66842835 | 1 |
| Rpl13 | 1.0197946 | 0.0282786 | 0.66846466 | 1 |
| Fhit | 1.05911408 | 0.082857999 | 0.66858797 | 1 |
| Dpy19l3 | 1.0351877 | 0.049892384 | 0.66864367 | 1 |
| Ier3 | 1.04417974 | 0.062370078 | 0.66867811 | 1 |
| Dlc1 | 0.9737169 | -0.038425718 | 0.66877592 | 1 |
| Smim20 | 1.03342143 | 0.047428708 | 0.66906919 | 1 |
| Cntnap5b | 0.91620793 | -0.126253038 | 0.669092 | 1 |
| Gdap2 | 1.0277531 | 0.039493719 | 0.66909897 | 1 |
| Lrrc28 | 0.97246207 | -0.040286118 | 0.6693117 | 1 |
| Adgrl2 | 1.02649031 | 0.037720005 | 0.66931888 | 1 |
| Slc38a6 | 1.08623632 | 0.11933801 | 0.66949483 | 1 |
| Fau | 1.02024114 | 0.028910185 | 0.66952288 | 1 |
| Zfp26 | 0.9482061 | -0.076727421 | 0.66963272 | 1 |
| Pank4 | 1.03083916 | 0.043819251 | 0.66968429 | 1 |
| Trib1 | 1.05389361 | 0.075729235 | 0.66969032 | 1 |
| RGD1564804 | 1.05094905 | 0.071692723 | 0.66973023 | 1 |
| Pfkfb3 | 1.03065194 | 0.043557204 | 0.66975912 | 1 |
| Tincr | 0.95500456 | -0.066420479 | 0.66989182 | 1 |
| Fam185a | 1.04176479 | 0.059029585 | 0.66990385 | 1 |
| Fn1 | 1.01993218 | 0.02847322 | 0.67002349 | 1 |
| Casp8ap2 | 1.03171685 | 0.04504709 | 0.67004633 | 1 |
| Dld | 1.01978267 | 0.028261727 | 0.6700524 | 1 |
| Tmem119 | 0.92860349 | -0.106865392 | 0.67024754 | 1 |
| Brcc3 | 1.03483271 | 0.049397569 | 0.67033748 | 1 |
| Hic2 | 0.9123598 | -0.132325214 | 0.67039746 | 1 |
| Egr2 | 0.97047917 | -0.043230849 | 0.6703978 | 1 |
| Pi4k2b | 1.04527224 | 0.063878736 | 0.67042696 | 1 |
| Aifm2 | 0.96134094 | -0.056879928 | 0.67042797 | 1 |
| LOC498368 | 1.05465024 | 0.076764627 | 0.67044115 | 1 |
| Cobl | 0.97542387 | -0.035898824 | 0.67053827 | 1 |
| Osbpl6 | 0.97364923 | -0.038525981 | 0.67064665 | 1 |
| Shkbp1 | 1.05280293 | 0.074235408 | 0.67114711 | 1 |
| Ctdnep1 | 1.0292725 | 0.041624985 | 0.67117159 | 1 |
| Ttc33 | 1.03074707 | 0.043690354 | 0.67133333 | 1 |

| | | | | |
|---|---|---|---|---|
| Rbm41 | 1.06465508 | 0.090386117 | 0.67134784 | 1 |
| Gng11 | 1.03516603 | 0.049862179 | 0.67141061 | 1 |
| Bambi | 1.06709028 | 0.093682236 | 0.67141809 | 1 |
| Kcna5 | 0.94881215 | -0.075805606 | 0.67143157 | 1 |
| Mrc2 | 1.03204766 | 0.045509598 | 0.67144325 | 1 |
| Tmem173 | 0.92738043 | -0.108766806 | 0.67165956 | 1 |
| Lrfn2 | 0.93599301 | -0.095430345 | 0.67170298 | 1 |
| Ptrf | 0.97994818 | -0.029222639 | 0.67176409 | 1 |
| Rab1b | 0.98038276 | -0.028582977 | 0.6718548 | 1 |
| Epm2aip1 | 1.01973201 | 0.028190049 | 0.6719596 | 1 |
| Fut4 | 0.94134007 | -0.08721209 | 0.6720738 | 1 |
| Ist1 | 1.02116426 | 0.030214957 | 0.67234154 | 1 |
| Rpl18a | 1.01962952 | 0.028045053 | 0.67238109 | 1 |
| Clec4d | 0.84505523 | -0.242882465 | 0.6726699 | 1 |
| Rnpep | 1.03520073 | 0.049910542 | 0.67270898 | 1 |
| Peg3 | 0.98022177 | -0.028819905 | 0.67275124 | 1 |
| Prkd2 | 1.06111356 | 0.085579066 | 0.6727638 | 1 |
| Mapk7 | 1.03969592 | 0.056161645 | 0.67280962 | 1 |
| Samd5 | 0.91677952 | -0.125353274 | 0.67281174 | 1 |
| Rims3 | 0.96551121 | -0.050635092 | 0.67281935 | 1 |
| Stk39 | 1.01955264 | 0.027936259 | 0.67282302 | 1 |
| Orc6 | 0.9377659 | -0.092700278 | 0.67305604 | 1 |
| Adck4 | 1.0377197 | 0.053416805 | 0.6731155 | 1 |
| Yaf2 | 0.97833521 | -0.03159923 | 0.67315538 | 1 |
| Ppcs | 0.96609154 | -0.049768205 | 0.67319971 | 1 |
| Igsf21 | 1.06076197 | 0.085100962 | 0.67322687 | 1 |
| Slc25a32 | 0.96632354 | -0.049421795 | 0.67330214 | 1 |
| Gon4l | 0.97174483 | -0.041350571 | 0.67330895 | 1 |
| Nudt16 | 0.96570463 | -0.050346102 | 0.67341722 | 1 |
| Psme3 | 0.97649125 | -0.03432098 | 0.67343218 | 1 |
| Abhd14b | 0.9667511 | -0.048783597 | 0.67345686 | 1 |
| Rdh13 | 0.95770823 | -0.062341891 | 0.6735333 | 1 |
| Vimp | 0.9747581 | -0.036883863 | 0.67356948 | 1 |
| Hsf1 | 0.97069582 | -0.042908814 | 0.67374965 | 1 |
| Map1lc3a | 0.9805803 | -0.028292312 | 0.67376775 | 1 |
| Chuk | 0.97021215 | -0.04362785 | 0.6738281 | 1 |
| Fam43a | 1.04488337 | 0.06334192 | 0.67386426 | 1 |
| Trappc8 | 1.02074052 | 0.029616162 | 0.67392399 | 1 |
| Taok1 | 0.97575503 | -0.035409095 | 0.67397956 | 1 |
| Gtpbp8 | 0.94308786 | -0.084535917 | 0.67429206 | 1 |

| | | | | |
|---|---|---|---|---|
| Adamts4 | 0.92131159 | -0.118238927 | 0.67435252 | 1 |
| Opa3 | 0.96677329 | -0.048750473 | 0.67437432 | 1 |
| Hnrnpa3 | 1.01950856 | 0.027873884 | 0.67438079 | 1 |
| Dync1i2 | 0.98040015 | -0.028557386 | 0.674463 | 1 |
| Ost4 | 1.02338382 | 0.033347335 | 0.67454891 | 1 |
| Mab21l2 | 0.97228233 | -0.040552793 | 0.67464521 | 1 |
| Nudt1 | 1.0800303 | 0.111071789 | 0.67471167 | 1 |
| Pcdh18 | 1.03436261 | 0.048742031 | 0.67471986 | 1 |
| Clint1 | 1.02940132 | 0.041805533 | 0.67492464 | 1 |
| Ddx46 | 0.97862192 | -0.031176496 | 0.67511062 | 1 |
| C1qtnf7 | 0.93919925 | -0.090496839 | 0.67540646 | 1 |
| Map2k4 | 1.02237609 | 0.031925997 | 0.67541357 | 1 |
| Tax1bp1 | 1.01965798 | 0.028085322 | 0.67541799 | 1 |
| Rhobtb3 | 1.02704181 | 0.038494909 | 0.67552904 | 1 |
| Snrpb | 1.02918425 | 0.041501279 | 0.67553817 | 1 |
| Cers4 | 1.02447215 | 0.034880766 | 0.67559054 | 1 |
| Flnb | 1.02023729 | 0.028904738 | 0.6756222 | 1 |
| Tmem242 | 1.03609074 | 0.051150364 | 0.67569079 | 1 |
| Mrps15 | 1.02493459 | 0.035531838 | 0.67576473 | 1 |
| Klhl15 | 1.07872296 | 0.109324396 | 0.67576688 | 1 |
| Amigo1 | 0.97256668 | -0.040130935 | 0.67576803 | 1 |
| Arhgap24 | 1.03672637 | 0.052035171 | 0.67581878 | 1 |
| Slc25a30 | 1.04028456 | 0.056978216 | 0.67588904 | 1 |
| Chchd5 | 1.06277305 | 0.087833553 | 0.67595858 | 1 |
| Slc9a7 | 1.0433379 | 0.061206474 | 0.67611583 | 1 |
| Kif1c | 0.97595521 | -0.035113158 | 0.67619705 | 1 |
| Trim59 | 1.05003866 | 0.07044245 | 0.67621322 | 1 |
| Cds2 | 1.02133388 | 0.030454566 | 0.67622789 | 1 |
| Pax5 | 0.7954435 | -0.33016863 | 0.67623224 | 1 |
| Ppm1l | 0.97989927 | -0.029294649 | 0.67627071 | 1 |
| Pkdcc | 1.04042538 | 0.057173497 | 0.67627403 | 1 |
| Ak1 | 1.02185605 | 0.031191972 | 0.67628607 | 1 |
| Arntl | 1.05055114 | 0.071146394 | 0.67636212 | 1 |
| Ddx20 | 1.03641944 | 0.05160798 | 0.67642796 | 1 |
| Pigs | 0.97570672 | -0.035480523 | 0.67652735 | 1 |
| Bles03 | 1.02747568 | 0.039104251 | 0.6765869 | 1 |
| Zfp362 | 0.95051679 | -0.073215976 | 0.67666238 | 1 |
| Lgals8 | 1.01930222 | 0.027581869 | 0.67671014 | 1 |
| L2hgdh | 0.92749356 | -0.10859083 | 0.67683213 | 1 |
| Rbm18 | 1.02245803 | 0.032041627 | 0.67691235 | 1 |

| | | | | |
|---|---|---|---|---|
| Clcn7 | 0.97049919 | -0.043201092 | 0.67691535 | 1 |
| Calr3 | 0.92861803 | -0.106842805 | 0.67709535 | 1 |
| Hivep3 | 0.96321978 | -0.054063084 | 0.67722859 | 1 |
| Fktn | 0.97455983 | -0.037177333 | 0.67723705 | 1 |
| Wdr53 | 1.05059395 | 0.071205187 | 0.6773186 | 1 |
| Wnt4 | 1.10121851 | 0.139100767 | 0.67753595 | 1 |
| C1qtnf5 | 1.03970752 | 0.056177735 | 0.67764978 | 1 |
| Rnf19a | 0.9810699 | -0.027572161 | 0.67770181 | 1 |
| Brd7 | 0.97261419 | -0.040060456 | 0.67770783 | 1 |
| Rdh10 | 0.94685121 | -0.078790364 | 0.67781638 | 1 |
| Bmpr1a | 1.02413026 | 0.034399221 | 0.6779334 | 1 |
| Nop14 | 1.02966804 | 0.042179296 | 0.67821237 | 1 |
| Ccdc85a | 0.96605851 | -0.049817524 | 0.67852248 | 1 |
| Zmiz1 | 0.97865745 | -0.031124125 | 0.67853034 | 1 |
| Gemin2 | 1.04859724 | 0.068460649 | 0.67855171 | 1 |
| Rplp1 | 1.02134341 | 0.030468024 | 0.6785691 | 1 |
| Fis1 | 1.02112245 | 0.030155887 | 0.6785951 | 1 |
| Spata13 | 0.96825593 | -0.046539659 | 0.67879987 | 1 |
| Capn7 | 0.97601469 | -0.035025234 | 0.67882624 | 1 |
| Tmed3 | 0.97315326 | -0.039261057 | 0.67902682 | 1 |
| Pkhd1l1 | 1.04499932 | 0.063502009 | 0.67907721 | 1 |
| Zfp637 | 0.9667709 | -0.048754043 | 0.67914438 | 1 |
| Dnajc15 | 1.0338968 | 0.048092194 | 0.67917133 | 1 |
| Npr2 | 1.02022133 | 0.028882169 | 0.67929262 | 1 |
| Rffl | 0.95424818 | -0.06756356 | 0.67956201 | 1 |
| Nsf | 1.01957037 | 0.027961355 | 0.6795683 | 1 |
| Tcf20 | 0.97541269 | -0.035915356 | 0.67970556 | 1 |
| Hibch | 1.03365828 | 0.047759313 | 0.67973436 | 1 |
| Lin7b | 0.95391142 | -0.068072785 | 0.67987985 | 1 |
| Scap | 1.02276059 | 0.032468479 | 0.6799121 | 1 |
| Edem3 | 1.02698242 | 0.03841149 | 0.68005463 | 1 |
| Cd82 | 0.97993404 | -0.029243453 | 0.68009152 | 1 |
| Maged1 | 1.01917371 | 0.027399965 | 0.68050171 | 1 |
| Vps29 | 0.97992514 | -0.029256555 | 0.68050907 | 1 |
| Gars | 1.01929796 | 0.027575837 | 0.68056498 | 1 |
| Nxpe1 | 1.07428234 | 0.103373212 | 0.68057616 | 1 |
| Fkbp2 | 1.02720594 | 0.038725446 | 0.68058542 | 1 |
| Dync1li2 | 0.98103619 | -0.027621744 | 0.68071875 | 1 |
| Sf3b5 | 1.02564136 | 0.036526351 | 0.68077358 | 1 |
| Chp1 | 0.98120396 | -0.027375037 | 0.68083996 | 1 |

| | | | | |
|---|---|---|---|---|
| Rnf128 | 0.94025613 | -0.088874283 | 0.68084575 | 1 |
| Ccz1b | 1.02650174 | 0.037736071 | 0.68092253 | 1 |
| Ptp4a3 | 0.96436779 | -0.052344626 | 0.68093357 | 1 |
| Tcf7l2 | 1.05033905 | 0.070855109 | 0.68107388 | 1 |
| Zfp772 | 0.96198661 | -0.055911278 | 0.68110027 | 1 |
| Spata2L | 0.93118916 | -0.102853835 | 0.68118206 | 1 |
| Mboat1 | 0.96263749 | -0.05493549 | 0.68123109 | 1 |
| Fam83f | 1.04483725 | 0.063278234 | 0.68124176 | 1 |
| Slc30a9 | 0.98084893 | -0.027897149 | 0.68126027 | 1 |
| Enthd2 | 0.96672049 | -0.048829281 | 0.6814355 | 1 |
| RGD1565222 | 1.05985099 | 0.083861441 | 0.68144008 | 1 |
| Kctd11 | 0.95509438 | -0.066284795 | 0.68144938 | 1 |
| Rpl22l1 | 1.02544254 | 0.036246659 | 0.6814865 | 1 |
| Eif2b5 | 1.02256961 | 0.032199054 | 0.68149383 | 1 |
| Impa2 | 0.95880551 | -0.060689888 | 0.68150604 | 1 |
| Ccdc146 | 1.08116044 | 0.112580634 | 0.68153896 | 1 |
| Riok1 | 1.03161079 | 0.044898773 | 0.68163013 | 1 |
| Fam214a | 0.97151936 | -0.041685355 | 0.68164247 | 1 |
| Sgcz | 0.92042209 | -0.119632487 | 0.68169942 | 1 |
| Psat1 | 0.96722107 | -0.048082424 | 0.68193054 | 1 |
| Itga11 | 1.04017217 | 0.056822344 | 0.68194887 | 1 |
| RGD1562079 | 0.97021232 | -0.04362759 | 0.68200785 | 1 |
| Gpcpd1 | 1.02045699 | 0.02921537 | 0.68204177 | 1 |
| Nf1 | 0.98035683 | -0.028621138 | 0.68212737 | 1 |
| RT1-N1 | 1.1073565 | 0.147119759 | 0.68216123 | 1 |
| Arf2 | 1.03071692 | 0.043648156 | 0.68226461 | 1 |
| Asb2 | 0.91751778 | -0.124191973 | 0.68228858 | 1 |
| Cdca2 | 1.10181101 | 0.139876783 | 0.68242433 | 1 |
| Fgf9 | 1.03348348 | 0.047515326 | 0.68246707 | 1 |
| Arf4 | 1.01935263 | 0.027653218 | 0.68252486 | 1 |
| Hdac10 | 1.04228239 | 0.05974621 | 0.68266742 | 1 |
| Acvr1 | 0.95964518 | -0.059427014 | 0.68273963 | 1 |
| Tdrd7 | 1.0212202 | 0.030293978 | 0.68278676 | 1 |
| Vars2 | 0.95915771 | -0.060160042 | 0.68284803 | 1 |
| Satb1 | 0.97102807 | -0.042415092 | 0.68286045 | 1 |
| Tfb2m | 1.0266769 | 0.037982229 | 0.6829625 | 1 |
| Ctnnbl1 | 1.02810803 | 0.03999187 | 0.68297661 | 1 |
| Vprbp | 0.9737045 | -0.038444079 | 0.68300645 | 1 |
| Qrich1 | 1.0243143 | 0.034658456 | 0.6833391 | 1 |
| Purg | 1.05666011 | 0.079511393 | 0.68337606 | 1 |

| | | | | |
|---|---|---|---|---|
| Ccdc181 | 1.03069525 | 0.043617824 | 0.68340764 | 1 |
| Gemin8 | 1.04198626 | 0.059336253 | 0.68362245 | 1 |
| Cpn1 | 0.91556413 | -0.127267151 | 0.68366075 | 1 |
| B9d1 | 0.94853958 | -0.076220117 | 0.68378413 | 1 |
| Timm8b | 1.02214927 | 0.031605902 | 0.68383539 | 1 |
| Tle2 | 0.94167958 | -0.086691849 | 0.68388007 | 1 |
| Phf7 | 0.95273999 | -0.069845542 | 0.68393246 | 1 |
| LOC681325 | 0.84579148 | -0.241626065 | 0.68395696 | 1 |
| Llgl1 | 0.97239186 | -0.040390277 | 0.68400769 | 1 |
| Eya4 | 0.90012873 | -0.151796757 | 0.68400936 | 1 |
| Etnk2 | 1.09684788 | 0.133363456 | 0.68401968 | 1 |
| Slf2 | 0.97392951 | -0.038110738 | 0.6840234 | 1 |
| Gne | 1.03336843 | 0.047354708 | 0.68417346 | 1 |
| Abcc2 | 1.09149347 | 0.126303504 | 0.68421659 | 1 |
| Ptpmt1 | 0.96877943 | -0.045759859 | 0.6842752 | 1 |
| Tub | 0.91700942 | -0.124991544 | 0.68429154 | 1 |
| Ttc14 | 1.0295124 | 0.041961205 | 0.68445865 | 1 |
| Sec14l2 | 0.97580952 | -0.035328539 | 0.68454789 | 1 |
| Jam3 | 1.02382726 | 0.033972325 | 0.6845823 | 1 |
| Osbpl9 | 1.01885442 | 0.02694792 | 0.68460908 | 1 |
| Nup58 | 1.02947514 | 0.041908992 | 0.68466933 | 1 |
| Eef1e1 | 0.97086638 | -0.042655339 | 0.68479798 | 1 |
| Crispld2 | 0.9797294 | -0.02954476 | 0.68482134 | 1 |
| Zfand6 | 1.02232467 | 0.031853447 | 0.68483892 | 1 |
| Get4 | 1.02600518 | 0.03703801 | 0.68506934 | 1 |
| Abtb1 | 1.02950171 | 0.041946231 | 0.68507145 | 1 |
| Papss1 | 0.98093094 | -0.027776519 | 0.6851068 | 1 |
| Cfl2 | 0.98162743 | -0.026752525 | 0.68522669 | 1 |
| Tpcn1 | 1.02017284 | 0.028813592 | 0.68525489 | 1 |
| Rhno1 | 1.05125748 | 0.072116069 | 0.68529557 | 1 |
| Vwa1 | 1.0194405 | 0.027777578 | 0.68544495 | 1 |
| Ilf2 | 0.9727446 | -0.039867031 | 0.68545729 | 1 |
| Sod3 | 1.02048243 | 0.029251343 | 0.68551747 | 1 |
| Kmt5c | 0.9650202 | -0.051368957 | 0.68561124 | 1 |
| Tkfc | 0.94654888 | -0.079251089 | 0.68563208 | 1 |
| Sytl3 | 0.96567202 | -0.050394815 | 0.6857304 | 1 |
| Rpl30 | 0.97944176 | -0.02996839 | 0.68591605 | 1 |
| Usp6nl | 0.97714055 | -0.033362003 | 0.68599102 | 1 |
| Lrrc1 | 0.94960043 | -0.0746075 | 0.68627251 | 1 |
| Pink1 | 1.02315846 | 0.033029592 | 0.68637787 | 1 |

| | | | | |
|---|---|---|---|---|
| Gtf2h4 | 1.04553473 | 0.064240982 | 0.68639365 | 1 |
| Hmbs | 1.03108132 | 0.044158127 | 0.68641246 | 1 |
| Zdhhc6 | 1.0283544 | 0.04033755 | 0.68646174 | 1 |
| Pdlim4 | 0.96928855 | -0.045001887 | 0.68648885 | 1 |
| Gtdc1 | 0.96731253 | -0.047946013 | 0.68663703 | 1 |
| Ythdc1 | 1.0237077 | 0.033803834 | 0.68664621 | 1 |
| Efcc1 | 1.08217052 | 0.113927851 | 0.68683353 | 1 |
| Rpp30 | 0.9710595 | -0.042368391 | 0.68692904 | 1 |
| Ntn1 | 1.03672311 | 0.05203063 | 0.6869404 | 1 |
| Jun | 0.98088724 | -0.027840803 | 0.68712892 | 1 |
| Mettl25 | 0.9479771 | -0.077075881 | 0.6871953 | 1 |
| Ngrn | 1.01910196 | 0.027298403 | 0.68725758 | 1 |
| Parp16 | 1.06286082 | 0.087952684 | 0.68735471 | 1 |
| Pak1ip1 | 1.05315092 | 0.074712199 | 0.68738969 | 1 |
| Adamts15 | 0.95951278 | -0.059626072 | 0.68744305 | 1 |
| Tbc1d17 | 0.97199588 | -0.04097789 | 0.68763224 | 1 |
| Dusp16 | 0.9413808 | -0.08714966 | 0.68764409 | 1 |
| Rtn3 | 1.01998298 | 0.028545079 | 0.68770611 | 1 |
| Filip1 | 0.97237042 | -0.040422081 | 0.68771544 | 1 |
| R3hdm2 | 0.97869259 | -0.031072313 | 0.68778417 | 1 |
| Usp13 | 0.97840129 | -0.031501793 | 0.68778966 | 1 |
| Pcgf1 | 0.96329713 | -0.053947223 | 0.6878096 | 1 |
| Stat5a | 1.03802714 | 0.053844158 | 0.68783312 | 1 |
| Recql4 | 0.91757397 | -0.124103627 | 0.68785442 | 1 |
| Gstcd | 0.93372532 | -0.098929887 | 0.68797984 | 1 |
| Rai2 | 1.05675052 | 0.079634816 | 0.68808174 | 1 |
| Sarnp | 1.02642126 | 0.037622952 | 0.68809961 | 1 |
| Trappc2b | 0.97345811 | -0.038809197 | 0.68813293 | 1 |
| Lamc3 | 0.94961588 | -0.074584036 | 0.68822025 | 1 |
| Myoz1 | 1.13121202 | 0.177869351 | 0.6882913 | 1 |
| Gtf2f2 | 1.04527433 | 0.063881619 | 0.68830818 | 1 |
| Golga7b | 0.96485451 | -0.051616685 | 0.68835484 | 1 |
| Mafk | 1.03416114 | 0.048460997 | 0.68835886 | 1 |
| Pdpk1 | 0.97547735 | -0.035819718 | 0.68842842 | 1 |
| Klhl9 | 0.98178404 | -0.026522385 | 0.68851983 | 1 |
| Ccnl2 | 0.97174241 | -0.04135416 | 0.68861995 | 1 |
| Robo2 | 0.9817909 | -0.026512301 | 0.68868083 | 1 |
| Tcn2 | 1.02337361 | 0.03333293 | 0.68869545 | 1 |
| Epb42 | 0.91422296 | -0.129382051 | 0.68872971 | 1 |
| Ccdc59 | 1.04207742 | 0.059462463 | 0.68873202 | 1 |

| | | | | |
|---|---|---|---|---|
| 15-Sep | 1.01851693 | 0.026469963 | 0.68877107 | 1 |
| Magi3 | 1.01933635 | 0.027630179 | 0.68880692 | 1 |
| Scfd1 | 0.97731323 | -0.033107077 | 0.68883624 | 1 |
| Oas1k | 1.20944238 | 0.274342034 | 0.68897799 | 1 |
| Mthfd2 | 1.03819578 | 0.05407853 | 0.68915288 | 1 |
| RGD621098 | 1.03266819 | 0.046376773 | 0.68926101 | 1 |
| Fam207a | 1.03212098 | 0.04561209 | 0.68957798 | 1 |
| Bcas2 | 1.02883068 | 0.04100557 | 0.68959387 | 1 |
| Rhbdl1 | 1.06267573 | 0.087701437 | 0.68961058 | 1 |
| Pmm1 | 1.01977745 | 0.028254346 | 0.68961373 | 1 |
| Smarcc1 | 1.02442327 | 0.034811924 | 0.68969337 | 1 |
| Rasgrp4 | 1.10905313 | 0.149328481 | 0.68975541 | 1 |
| Myg1 | 0.96892203 | -0.045547524 | 0.6898394 | 1 |
| Flvcr2 | 1.04969486 | 0.069970003 | 0.68995074 | 1 |
| Tmem38b | 0.95317744 | -0.069183289 | 0.6900423 | 1 |
| Nr4a2 | 1.05138557 | 0.072291846 | 0.69009777 | 1 |
| Dpy30 | 1.02765403 | 0.039354651 | 0.69014122 | 1 |
| Gna11 | 1.04626148 | 0.065243458 | 0.69015609 | 1 |
| Cnot6l | 1.04170809 | 0.058951063 | 0.69037305 | 1 |
| Xrcc1 | 0.96959122 | -0.044551466 | 0.69040857 | 1 |
| Sertad3 | 1.06490245 | 0.090721281 | 0.69055307 | 1 |
| Thy1 | 1.01945144 | 0.027793059 | 0.69055578 | 1 |
| Hyal1 | 0.9455797 | -0.080729027 | 0.6905692 | 1 |
| Sephs1 | 1.02756557 | 0.039230455 | 0.69058873 | 1 |
| Dhps | 1.02770839 | 0.039430957 | 0.69059542 | 1 |
| Gapt | 1.20800515 | 0.272626611 | 0.69073472 | 1 |
| Pex11a | 0.9565526 | -0.06408379 | 0.6907372 | 1 |
| Rpp40 | 0.9654444 | -0.050734919 | 0.69080901 | 1 |
| Atp1b1 | 1.01949675 | 0.027857183 | 0.69085027 | 1 |
| Psmd2 | 1.01852063 | 0.026475206 | 0.69092172 | 1 |
| Mynn | 0.96292244 | -0.054508491 | 0.69094131 | 1 |
| Slc39a6 | 0.97808544 | -0.031967591 | 0.69098968 | 1 |
| Arhgap18 | 1.03739075 | 0.052959415 | 0.69109978 | 1 |
| Hddc3 | 1.04255352 | 0.060121452 | 0.69123674 | 1 |
| Ift80 | 1.02696242 | 0.038383385 | 0.69124537 | 1 |
| Pdcd2l | 0.94913451 | -0.075315537 | 0.69132061 | 1 |
| Taf8 | 1.03506847 | 0.049726209 | 0.69168263 | 1 |
| Aaas | 0.96223803 | -0.05553428 | 0.69171978 | 1 |
| Rpl10 | 1.0184327 | 0.026350651 | 0.69175462 | 1 |
| Bbs10 | 0.96251213 | -0.055123377 | 0.69193773 | 1 |

| | | | | |
|---|---|---|---|---|
| Cuedc1 | 0.96374864 | -0.053271173 | 0.69202079 | 1 |
| Bex2 | 1.01929569 | 0.027572627 | 0.69209085 | 1 |
| Znrf2 | 1.0244071 | 0.034789164 | 0.69213281 | 1 |
| Diaph1 | 1.0290173 | 0.04126724 | 0.69223111 | 1 |
| Haus3 | 0.96484158 | -0.051636014 | 0.69228156 | 1 |
| Rbbp4 | 1.02494499 | 0.035546481 | 0.69233808 | 1 |
| Sgcb | 1.01922367 | 0.027470689 | 0.69244478 | 1 |
| Rapsn | 0.93348129 | -0.099306991 | 0.69252897 | 1 |
| Ppt2 | 0.96506026 | -0.051309066 | 0.69256805 | 1 |
| Ripk3 | 0.92908785 | -0.106113071 | 0.69271756 | 1 |
| Tgfbrap1 | 0.97031335 | -0.043477369 | 0.69282494 | 1 |
| Ncbp2 | 0.97894915 | -0.030694166 | 0.6928808 | 1 |
| Atp5g1 | 1.02246306 | 0.032048726 | 0.69290673 | 1 |
| Deaf1 | 0.97234869 | -0.040454324 | 0.69290803 | 1 |
| Cd177 | 0.91630345 | -0.126102641 | 0.69294597 | 1 |
| Mplkip | 1.04105681 | 0.058048798 | 0.69313837 | 1 |
| Bcl2l11 | 0.94511471 | -0.081438659 | 0.69314639 | 1 |
| Zfp865 | 0.94374153 | -0.0835363 | 0.69324272 | 1 |
| Nphp3 | 0.9458448 | -0.080324616 | 0.69325983 | 1 |
| Abcc3 | 1.05040254 | 0.070942313 | 0.69335991 | 1 |
| Rps3a | 0.98207944 | -0.026088364 | 0.69351042 | 1 |
| Ctnnd1 | 0.98140109 | -0.027085218 | 0.69394587 | 1 |
| Schip1 | 0.97947157 | -0.029924473 | 0.69400365 | 1 |
| Med4 | 1.0342882 | 0.048638237 | 0.69420795 | 1 |
| Akap9 | 0.98166874 | -0.026691815 | 0.69425717 | 1 |
| Kat5 | 1.02431859 | 0.034664504 | 0.69441775 | 1 |
| Rps6ka3 | 1.0200326 | 0.028615262 | 0.69449857 | 1 |
| Rpgrip1l | 1.0295353 | 0.041993298 | 0.69449878 | 1 |
| Vps35 | 0.98199457 | -0.026213049 | 0.69467239 | 1 |
| Eif5a | 1.01816363 | 0.025969444 | 0.69473534 | 1 |
| Sh2d4a | 1.06624196 | 0.092534856 | 0.69488261 | 1 |
| Fgf13 | 1.01851742 | 0.026470649 | 0.69493782 | 1 |
| Wrb | 1.01975758 | 0.02822623 | 0.69494565 | 1 |
| Depdc5 | 1.01984322 | 0.028347387 | 0.69496144 | 1 |
| Ttc9b | 0.96599249 | -0.04991612 | 0.69498955 | 1 |
| Rimbp2 | 0.97050532 | -0.043191972 | 0.69510588 | 1 |
| Pih1d2 | 1.07372167 | 0.102620064 | 0.6951305 | 1 |
| Arpc4 | 1.01843363 | 0.026351962 | 0.69516322 | 1 |
| Rps21 | 0.94332634 | -0.084171151 | 0.69527513 | 1 |
| Ppwd1 | 1.03919005 | 0.055459517 | 0.69542441 | 1 |

| | | | | |
|---|---|---|---|---|
| Rab20 | 0.93761815 | -0.092927595 | 0.69547305 | 1 |
| Nrbf2 | 1.02914663 | 0.041448543 | 0.69576921 | 1 |
| Zfp511 | 1.04661876 | 0.065736023 | 0.69580688 | 1 |
| Gtf2h2 | 1.04233387 | 0.059817462 | 0.69590297 | 1 |
| Creb1 | 0.949546 | -0.074690201 | 0.69591865 | 1 |
| Smpd1 | 0.98152297 | -0.026906069 | 0.69594538 | 1 |
| Mxi1 | 0.97991338 | -0.029273868 | 0.69601922 | 1 |
| Adrb2 | 0.92765572 | -0.108338612 | 0.69613109 | 1 |
| Frmpd4 | 1.02311798 | 0.032972517 | 0.6962533 | 1 |
| Adcy8 | 0.9618889 | -0.056057833 | 0.69633073 | 1 |
| Rgl1 | 0.97965244 | -0.029658096 | 0.69635075 | 1 |
| Pla2g6 | 1.02345778 | 0.033451589 | 0.69637092 | 1 |
| Tgfa | 1.03251441 | 0.046161917 | 0.69658993 | 1 |
| Lcmt2 | 1.0537454 | 0.075526336 | 0.69673782 | 1 |
| Pnkp | 1.04366016 | 0.061652014 | 0.69676722 | 1 |
| Usp40 | 1.0283001 | 0.040261369 | 0.69680971 | 1 |
| Phf11 | 1.08024356 | 0.111356625 | 0.69684311 | 1 |
| Zufsp | 0.96539754 | -0.050804947 | 0.69687595 | 1 |
| Btf3 | 1.01874191 | 0.026788599 | 0.69690883 | 1 |
| Cdk10 | 1.02596137 | 0.036976408 | 0.6969176 | 1 |
| Mrps23 | 1.02551817 | 0.03635305 | 0.69697223 | 1 |
| Mtfp1 | 0.9261545 | -0.110675212 | 0.69699687 | 1 |
| Irs2 | 0.97858253 | -0.031234566 | 0.69701283 | 1 |
| Vwa5b2 | 0.97720858 | -0.033261564 | 0.6970598 | 1 |
| Alkbh1 | 1.04099512 | 0.057963311 | 0.69712858 | 1 |
| Maml3 | 0.93034921 | -0.104155756 | 0.69722554 | 1 |
| Rad51 | 0.94065597 | -0.088260921 | 0.69722765 | 1 |
| Dpp7 | 1.0229587 | 0.032747895 | 0.69726033 | 1 |
| Rchy1 | 1.02667675 | 0.037982024 | 0.69727504 | 1 |
| Nup85 | 1.03510895 | 0.049782631 | 0.6972842 | 1 |
| Nhp2 | 1.03263156 | 0.046325598 | 0.69731968 | 1 |
| Rnf20 | 1.02010176 | 0.028713069 | 0.69733294 | 1 |
| Gemin4 | 1.03309991 | 0.046979784 | 0.69735129 | 1 |
| Snx16 | 1.02329304 | 0.033219342 | 0.69751432 | 1 |
| Snrnp70 | 1.02027932 | 0.028964176 | 0.69753507 | 1 |
| Rtkn | 0.9815839 | -0.026816515 | 0.69768166 | 1 |
| RGD1308706 | 1.0312501 | 0.044394259 | 0.69771213 | 1 |
| Tab2 | 0.97736061 | -0.033037126 | 0.69789153 | 1 |
| Ptdss1 | 0.9805843 | -0.028286436 | 0.69803662 | 1 |
| Prpf3 | 1.02992064 | 0.042533169 | 0.69813349 | 1 |

| | | | | |
|---|---|---|---|---|
| Kdelr2 | 1.02255156 | 0.032173588 | 0.69848558 | 1 |
| Acads | 1.0267603 | 0.038099418 | 0.69878842 | 1 |
| Atrip | 0.96039691 | -0.058297326 | 0.69885005 | 1 |
| Phkb | 0.97251677 | -0.040204973 | 0.69899923 | 1 |
| Zfand5 | 0.98247931 | -0.025501073 | 0.69922177 | 1 |
| Fgr | 0.9370022 | -0.09387566 | 0.69930946 | 1 |
| Uba1 | 0.98211122 | -0.026041688 | 0.69936791 | 1 |
| Sdc4 | 1.02555222 | 0.03640095 | 0.69946175 | 1 |
| Nfic | 0.97743121 | -0.032932922 | 0.69952618 | 1 |
| Ten1 | 0.9512207 | -0.072147991 | 0.69972375 | 1 |
| Arl2bp | 1.01902468 | 0.027188987 | 0.69979527 | 1 |
| Gria4 | 1.03195795 | 0.045384187 | 0.69995281 | 1 |
| Mark1 | 1.02262523 | 0.03227752 | 0.70002299 | 1 |
| Helq | 0.96303354 | -0.05434205 | 0.70002802 | 1 |
| Pi4kb | 0.97891537 | -0.030743949 | 0.7000693 | 1 |
| Vegfb | 1.03857578 | 0.054606485 | 0.70016792 | 1 |
| Col16a1 | 1.02073853 | 0.029613362 | 0.70020777 | 1 |
| Ispd | 0.96966905 | -0.044435658 | 0.70035999 | 1 |
| Atp6v0c | 0.98222199 | -0.025878967 | 0.70047066 | 1 |
| Ssna1 | 0.97303925 | -0.03943009 | 0.70053168 | 1 |
| Patj | 1.01797953 | 0.025708556 | 0.70060067 | 1 |
| Pm20d2 | 0.95513464 | -0.066223971 | 0.70070185 | 1 |
| Rtcd1 | 1.02069009 | 0.029544891 | 0.70074807 | 1 |
| Bcan | 1.02306889 | 0.032903294 | 0.70079291 | 1 |
| Gmfg | 0.9517204 | -0.071390295 | 0.70104237 | 1 |
| Myef2 | 1.01865853 | 0.026670522 | 0.70115436 | 1 |
| Bcorl1 | 0.96114188 | -0.057178677 | 0.70122119 | 1 |
| Cox6a1 | 1.01807205 | 0.025839664 | 0.70125238 | 1 |
| Cdk11b | 1.02143806 | 0.030601715 | 0.70125602 | 1 |
| Fam107b | 0.97409849 | -0.037860439 | 0.70128096 | 1 |
| Etv1 | 0.98250845 | -0.025458277 | 0.70141248 | 1 |
| Defa11 | 0.79850983 | -0.324617927 | 0.70148128 | 1 |
| Fam49b | 0.97763062 | -0.032638619 | 0.70158824 | 1 |
| Nfia | 0.96591095 | -0.050037899 | 0.70171589 | 1 |
| Rbm20 | 0.9385659 | -0.091470051 | 0.70172403 | 1 |
| Metap1 | 1.023465 | 0.033461767 | 0.70190221 | 1 |
| Smap2 | 1.0195014 | 0.027863751 | 0.70196536 | 1 |
| Dpysl2 | 1.01832542 | 0.026198666 | 0.70198588 | 1 |
| C2cd5 | 0.97919546 | -0.030331225 | 0.70205473 | 1 |
| Cahm | 1.09421528 | 0.129896605 | 0.70206265 | 1 |

| | | | | |
|---|---|---|---|---|
| Ube2k | 1.01865111 | 0.026660005 | 0.70223688 | 1 |
| Slmap | 0.97979344 | -0.029450466 | 0.70227065 | 1 |
| Timm44 | 0.97459167 | -0.037130199 | 0.70237866 | 1 |
| Wdr33 | 0.97362459 | -0.038562488 | 0.70238348 | 1 |
| Cdk5rap1 | 1.03072779 | 0.043663381 | 0.70259411 | 1 |
| Ppp3cb | 0.98216755 | -0.025958934 | 0.70266766 | 1 |
| Lrrc59 | 1.01835829 | 0.02624523 | 0.70270328 | 1 |
| Pde8b | 1.04788105 | 0.067474954 | 0.70278225 | 1 |
| Dctd | 0.91995124 | -0.120370695 | 0.70278965 | 1 |
| Rapgef3 | 1.03674704 | 0.052063932 | 0.70280717 | 1 |
| Gyg1 | 1.01770801 | 0.025323699 | 0.70280911 | 1 |
| Fam89b | 1.02000072 | 0.028570174 | 0.70280998 | 1 |
| Csgalnact2 | 1.0294395 | 0.041859049 | 0.70287298 | 1 |
| Tmem107 | 1.06915547 | 0.096471652 | 0.7029014 | 1 |
| Nudt5 | 0.96503525 | -0.051346457 | 0.70303782 | 1 |
| Gtf3c5 | 1.0309884 | 0.044028104 | 0.70321759 | 1 |
| Nt5c1a | 1.11824059 | 0.161230623 | 0.70329895 | 1 |
| Pcdhgb7 | 0.96580132 | -0.050201661 | 0.70335205 | 1 |
| Tfcp2l1 | 1.05202581 | 0.073170104 | 0.70350362 | 1 |
| Arid4a | 1.02823733 | 0.040173292 | 0.70369176 | 1 |
| Ctsl | 0.9826101 | -0.02530902 | 0.70374267 | 1 |
| Dnajc10 | 1.01772942 | 0.02535405 | 0.70382058 | 1 |
| Ipo4 | 0.97992898 | -0.029250899 | 0.70385719 | 1 |
| Rplp0 | 0.98248803 | -0.025488258 | 0.70393111 | 1 |
| Strbp | 1.0187865 | 0.026851746 | 0.70396353 | 1 |
| Pkp1 | 1.02953025 | 0.041986218 | 0.70401939 | 1 |
| P4ha1 | 1.01960251 | 0.028006831 | 0.7040523 | 1 |
| Dgat1 | 0.96740129 | -0.047813636 | 0.70437368 | 1 |
| Lyrm2 | 1.03961129 | 0.056044203 | 0.70446247 | 1 |
| Gpc5 | 0.97083564 | -0.042701023 | 0.7044761 | 1 |
| Itpkc | 1.04477542 | 0.063192863 | 0.70459611 | 1 |
| Zbtb22 | 1.03011379 | 0.042803711 | 0.70461354 | 1 |
| Styxl1 | 0.93721399 | -0.093549611 | 0.70463176 | 1 |
| Rogdi | 0.98067724 | -0.028149699 | 0.70472934 | 1 |
| Alas1 | 0.97733726 | -0.033071601 | 0.70475432 | 1 |
| Nub1 | 1.01783925 | 0.025509728 | 0.70480657 | 1 |
| PVR | 0.95370583 | -0.068383761 | 0.70482707 | 1 |
| Timm9 | 0.96756135 | -0.047574948 | 0.70489703 | 1 |
| Chst15 | 1.02559805 | 0.036465429 | 0.70491119 | 1 |
| Ube2c | 1.11263643 | 0.153982255 | 0.70498305 | 1 |

| | | | | |
|---|---|---|---|---|
| Fgfr3 | 0.92824817 | -0.107417533 | 0.70506093 | 1 |
| Arpin | 0.96288381 | -0.054566377 | 0.70510172 | 1 |
| Hspa2 | 1.02157538 | 0.030795658 | 0.70515769 | 1 |
| Eif4enif1 | 1.02606394 | 0.037120638 | 0.70535143 | 1 |
| Ube2d3 | 1.01750545 | 0.025036524 | 0.70547751 | 1 |
| Pold4 | 1.04506623 | 0.063594374 | 0.7055233 | 1 |
| Ing4 | 1.02601196 | 0.037047542 | 0.70566098 | 1 |
| Spag1 | 1.04768697 | 0.067207724 | 0.70571792 | 1 |
| Atad1 | 1.01828811 | 0.026145808 | 0.7057557 | 1 |
| Erich2 | 0.91543914 | -0.127464121 | 0.70575929 | 1 |
| Rad50 | 0.97915442 | -0.030391694 | 0.7058454 | 1 |
| Echs1 | 1.02052201 | 0.029307299 | 0.70587747 | 1 |
| Bmpr2 | 0.98167369 | -0.026684549 | 0.70596757 | 1 |
| Sin3a | 0.97333544 | -0.03899101 | 0.70598065 | 1 |
| S100a5 | 0.90545193 | -0.14329005 | 0.70601495 | 1 |
| Lama5 | 1.03436189 | 0.04874103 | 0.70602085 | 1 |
| Ppef2 | 1.07941856 | 0.1102544 | 0.70620635 | 1 |
| Masp1 | 0.95871739 | -0.060822496 | 0.70642559 | 1 |
| Naa15 | 0.98204945 | -0.026132419 | 0.70643324 | 1 |
| Galnt1 | 1.0251871 | 0.035887232 | 0.70643891 | 1 |
| Grsf1 | 1.01772073 | 0.025341737 | 0.70647531 | 1 |
| Nuak1 | 1.03371808 | 0.047842777 | 0.70669855 | 1 |
| Nicn1 | 1.02079906 | 0.029698905 | 0.70697299 | 1 |
| Rras2 | 0.9679802 | -0.046950561 | 0.70716994 | 1 |
| Kank2 | 0.97867747 | -0.031094608 | 0.70730572 | 1 |
| RGD1561157 | 0.92620405 | -0.110598027 | 0.70739381 | 1 |
| Nbn | 1.02011093 | 0.028726045 | 0.70744018 | 1 |
| Klf10 | 0.97019445 | -0.043654169 | 0.70753296 | 1 |
| Zc3h18 | 0.97117371 | -0.042198724 | 0.70772412 | 1 |
| Ssr2 | 1.0208452 | 0.029764115 | 0.70775011 | 1 |
| Kcmf1 | 0.98011616 | -0.028975359 | 0.70781676 | 1 |
| Trpv2 | 0.97850112 | -0.031354591 | 0.7078884 | 1 |
| RGD1305537 | 1.05315359 | 0.074715848 | 0.70801935 | 1 |
| Dyrk3 | 1.06943028 | 0.096842427 | 0.70801958 | 1 |
| Ufm1 | 1.02147591 | 0.030655177 | 0.70815642 | 1 |
| Evi2a | 1.05416928 | 0.07610655 | 0.70821256 | 1 |
| St3gal4 | 0.97618506 | -0.03477342 | 0.7082757 | 1 |
| Uap1l1 | 1.0364717 | 0.051680727 | 0.70840011 | 1 |
| Ablim2 | 0.97427619 | -0.037597284 | 0.70844144 | 1 |
| Fam118b | 1.0388038 | 0.054923191 | 0.70845994 | 1 |

| | | | | |
|---|---|---|---|---|
| Zbtb25 | 0.96539663 | -0.050806309 | 0.7085233 | 1 |
| Pdcd6 | 1.02135181 | 0.030479901 | 0.70853728 | 1 |
| P4ha3 | 0.92485569 | -0.112699819 | 0.70859393 | 1 |
| Cbr1 | 0.97815081 | -0.031871177 | 0.7085963 | 1 |
| Pbx4 | 1.07202923 | 0.100344244 | 0.7086045 | 1 |
| Trerf1 | 0.94473423 | -0.082019556 | 0.70877036 | 1 |
| Mpc1 | 0.98254675 | -0.025402039 | 0.70890784 | 1 |
| Trappc9 | 0.97581267 | -0.03532388 | 0.70897672 | 1 |
| Irf2 | 0.9634462 | -0.053723984 | 0.70907113 | 1 |
| Eif4g3 | 0.98276079 | -0.025087801 | 0.70929449 | 1 |
| Podxl | 0.97607717 | -0.034932887 | 0.70934351 | 1 |
| Lig4 | 1.02464645 | 0.035126202 | 0.70946152 | 1 |
| Man1a1 | 1.02877239 | 0.040923836 | 0.70952881 | 1 |
| Cyp4f6 | 1.04387694 | 0.061951653 | 0.70959178 | 1 |
| Sar1a | 0.98254824 | -0.025399852 | 0.70970426 | 1 |
| Dnm2 | 0.97810197 | -0.031943215 | 0.70977831 | 1 |
| Ncaph2 | 1.02254868 | 0.032169521 | 0.71000922 | 1 |
| Nr2f6 | 0.96973616 | -0.044335807 | 0.71003789 | 1 |
| Zfp358 | 0.96174872 | -0.056268085 | 0.71045849 | 1 |
| Galnt14 | 1.02282136 | 0.032554193 | 0.7104863 | 1 |
| Psip1 | 0.97983848 | -0.029384141 | 0.71075964 | 1 |
| Fam122a | 1.04149054 | 0.058649729 | 0.71103067 | 1 |
| Rnpc3 | 0.96886106 | -0.045638308 | 0.71105091 | 1 |
| Lrrc4c | 1.02350533 | 0.033518621 | 0.71117628 | 1 |
| Scarb2 | 0.98138954 | -0.027102195 | 0.71124322 | 1 |
| Hax1 | 0.97066544 | -0.042953963 | 0.71124672 | 1 |
| Adcy6 | 0.97521785 | -0.036203559 | 0.71139439 | 1 |
| Slc41a1 | 0.97509992 | -0.036378038 | 0.71142145 | 1 |
| Ap1s1 | 0.98213419 | -0.026007935 | 0.71146034 | 1 |
| Tmem150a | 1.03706757 | 0.052509902 | 0.71146998 | 1 |
| Sssca1 | 1.02864905 | 0.040750858 | 0.71147215 | 1 |
| Ddost | 0.9808823 | -0.027848058 | 0.71156067 | 1 |
| Adgrl4 | 0.96787549 | -0.047106626 | 0.71156321 | 1 |
| Gsto1 | 0.97838467 | -0.031526297 | 0.7116602 | 1 |
| Rap1b | 0.97746067 | -0.032889435 | 0.71166705 | 1 |
| Tpra1 | 1.03215414 | 0.045658443 | 0.71168024 | 1 |
| Smpd2 | 1.06011943 | 0.084226798 | 0.71173894 | 1 |
| Uxs1 | 0.98138008 | -0.0271161 | 0.7117552 | 1 |
| Cspg4 | 1.02829739 | 0.040257561 | 0.7118016 | 1 |
| Klf3 | 0.97523407 | -0.03617957 | 0.71188672 | 1 |

| | | | | |
|---|---|---|---|---|
| Atg4a | 0.97431585 | -0.03753856 | 0.71189192 | 1 |
| Tmed10 | 1.01762726 | 0.02520922 | 0.71195007 | 1 |
