## Supplementary figures and images for "Neural Inflammation in Thoracic Dorsal Root Ganglia Mediates Cardiopulmonary Spinal Afferent Sensitization in Chronic Heart Failure"

### Supplmental Video

#### Slide 1
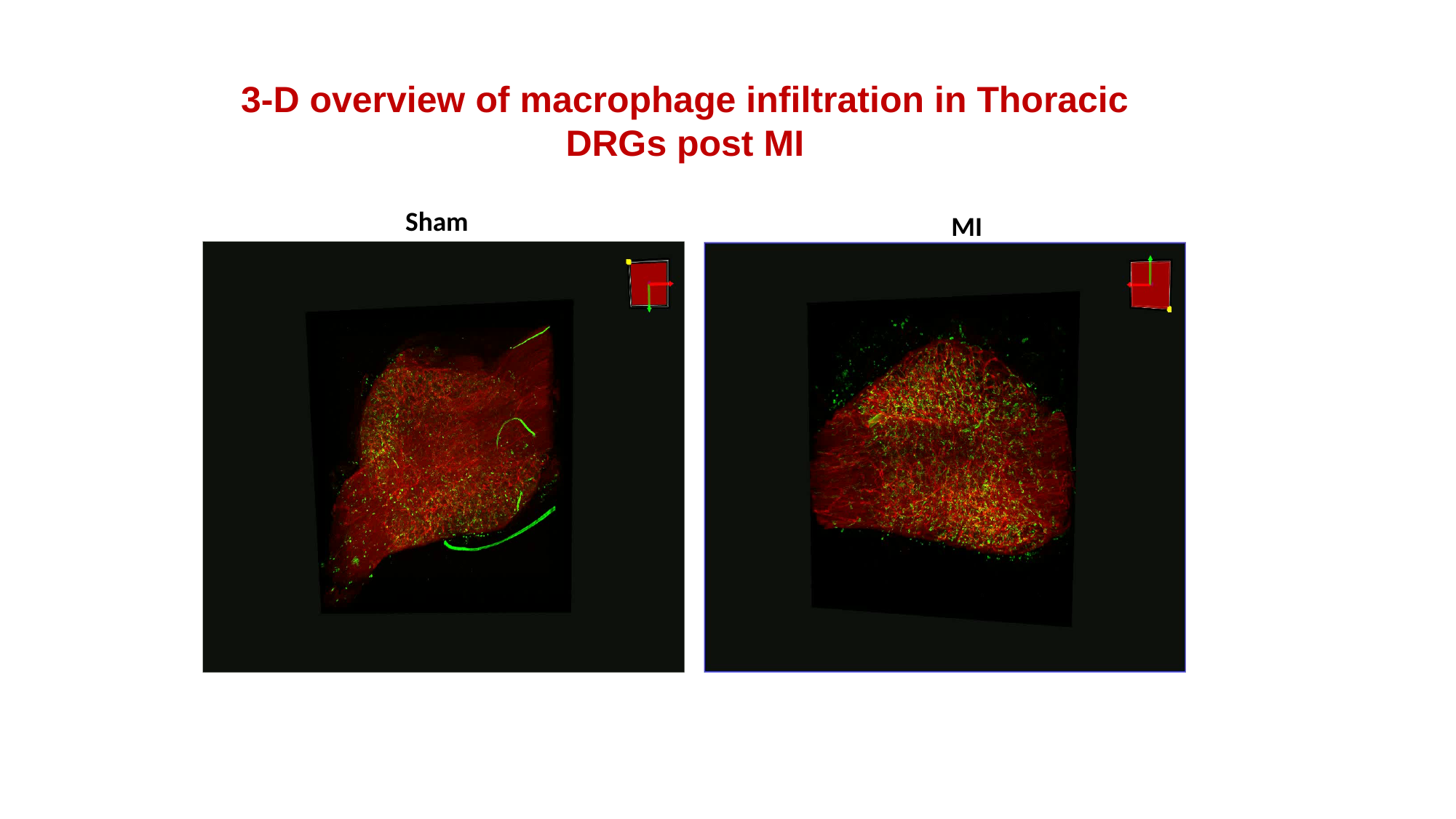

3-D overview of macrophage infiltration in Thoracic DRGs post MI
Sham
MI
